# DNA damage signaling in *Drosophila* macrophages modulates systemic cytokine levels in response to oxidative stress

**DOI:** 10.1101/2023.02.22.529599

**Authors:** Fabian Hersperger, Tim Meyring, Pia Weber, Chintan Chhatbar, Gianni Monaco, Marc S. Dionne, Katrin Paeschke, Marco Prinz, Olaf Groß, Anne-Kathrin Classen, Katrin Kierdorf

## Abstract

Environmental factors, infection, or injury can cause oxidative stress in diverse tissues and loss of tissue homeostasis. Effective stress response cascades, conserved from invertebrates to mammals, ensure reestablishment of homeostasis and tissue repair. Hemocytes, the *Drosophila* blood-like cells, rapidly respond to oxidative stress by immune activation. However, the precise signals how they sense oxidative stress and integrate these signals to modulate and balance the response to oxidative stress in the adult fly are ill-defined. Furthermore, hemocyte diversification was not explored yet on oxidative stress. Here, we employed high throughput single nuclei RNA-sequencing to explore hemocytes and other cell types, such as fat body, during oxidative stress in the adult fly. We identified distinct cellular responder states in plasmatocytes, the *Drosophila* macrophages, associated with immune response and metabolic activation upon oxidative stress. We further define oxidative stress-induced DNA damage signaling as a key sensor and a rate-limiting step in immune-activated plasmatocytes controlling JNK-mediated release of the pro-inflammatory cytokine *unpaired-3*. We subsequently tested the role of this specific immune activated cell stage during oxidative stress and found that inhibition of DNA damage signaling in plasmatocytes, as well as JNK or upd3 overactivation, result in a higher susceptibility to oxidative stress. Our findings uncover that a balanced composition and response of hemocyte subclusters is essential for the survival of adult *Drosophila* on oxidative stress by regulating systemic cytokine levels and cross-talk to other organs, such as the fat body, to control energy mobilization.

## Introduction

Metazoan organisms must constantly adapt to the surrounding environment. Environmental factors, including radiation, pathogens or toxins, permanently lead to the generation of reactive oxygen species (ROS)^1^. In vertebrates and invertebrates, low levels of ROS support cellular pathways like immune cell differentiation and even immune function including antimicrobial defense^2,3^. At high concentrations, ROS lead to oxidative stress, a deleterious process that negatively impacts protein synthesis and stability, lipid metabolism as well as overall genome stability. ROS mediated cellular damage can further result in degenerative diseases or cancer^4^. Therefore, metazoan organisms employ efficient repair and defense mechanisms to circumvent high levels of oxidative stress and subsequent organ damage. Hence, keeping the balance between the production and the antioxidant-dependent clearance of ROS is essential for the survival and well-being of the affected organism^5^. Oxidative stress induced tissue damage is most often followed by a strong local immune response. These molecular processes are conserved across vertebrates and invertebrates such as the fruit fly *Drosophila melanogaster*^6^.

Hemocytes, the *Drosophila* blood cells, are essential cellular components of the immune and stress response machinery. They consist of three different subtypes: crystal cells, lamellocytes and plasmatocytes^7^. Plasmatocytes are considered to be macrophage equivalents in *Drosophila* and comprise up to 95 % of all hemocytes in the adult fly^8^. They share many functions as well as the multilayered ontogeny with vertebrate tissue macrophages^9,10^. Adult plasmatocytes are actively involved in many immune functions such as pathogen recognition, defense and phagocytosis but also chronic immune activation upon metabolic stress by dietary lipids^11,12^. Recent studies highlighted the essential role of adult plasmatocytes as key players in regulation of tissue homeostasis, including the gut by regulating the intestinal stem cell niche or in muscles by controlling glucose homeostasis^13,14^.

Similar to higher eukaryotes, levels of ROS are used as a central antimicrobial defense mechanism of hemocytes against invading pathogens^15^. Moreover, larval hemocytes respond differentially to ROS, with two cell subsets mounting a biphasic immune reaction to invading bacteria by controlling hemocyte spreading and adhesion as well as crystal cell rupture^15^. Larval hemocytes have been also attributed to respond to increased oxidative stress of other tissues, as seen in response to oxidative stress in the eye-antenna imaginal disc^16^. Similarly, by regulating the intestinal stem cell (ISC) proliferation in the damaged midgut during infection-induced or chemical-induced oxidative stress^14^, adult hemocytes were described to release the bone morphogenetic protein (BMP) homologue *decapentaplegic* (*dpp*) to facilitate tissue repair^14^. Other studies pointed to an essential role of hemocyte-derived *unpaired-3* (*upd3*), which is the *Drosophila* orthologue of Interleukin-6, in the regulation of ISC proliferation during septic injury^17^, as well as wound healing during sterile injury^18^. In line with these findings, hemocytes have been implicated in organ-to-organ immune signaling for example upon infection-induced injury in the gut and antimicrobial peptide (AMP) production in the fat body^19^. Hemocyte activation is thus a hallmark of oxidative stress in defined tissues, though their role in modulating and controlling oxidative stress response to facilitate tissue repair and survival of the organism remains undefined.

Here, we demonstrate an essential role of hemocytes to control susceptibility to oxidative stress, where hemocytes orchestrate systemic oxidative stress response by defined cellular states identified by unbiased transcriptomic profiling. Plasmatocytes segregate in oxidative stress responder subclusters, characterized by immune activation and cellular stress response, and potential bystander subsets associated with metabolic gene regulations. Mechanistically, we unravel that the direct immune response to oxidative stress by plasmatocytes is driven by c-Jun N-terminal kinase (JNK) and Jak/STAT pathway activation with subsequent induction of the pro-inflammatory cytokine *upd3*. This inflammatory cascade is to a certain amount regulated by DNA damage signaling in plasmatocytes and essentially defines the susceptibility of the fly to oxidative stress. Deficiencies in the DNA damage signaling machinery in plasmatocytes lead to elevated JNK activation, cytokine release and subsequently a higher susceptibility of the adult fly to oxidative stress. Hence, our data point to a key role of plasmatocytes in balancing systemic cytokine levels and stress response upon oxidative stress. Collectively we provide new insights on the role of macrophages in orchestrating systemic stress responses in other tissues and to balance tissue wasting and energy mobilization.

## Results

### Hemocytes control susceptibility to oxidative stress

In vertebrates and invertebrates, prolonged oxidative stress can cause severe tissue damage, hence controlled activation of stress signaling cascades are essential for the survival of the organism and the resolution of oxidative stress with subsequent tissue repair. Yet the precise role of macrophages in the initiation and control of oxidative stress signaling needs to be determined.

To analyze the role of macrophages in signaling cascades to oxidative stress, we took advantage of the well-established Paraquat (PQ)-feeding model in adult *Drosophila melanogaster* to induce systemic oxidative stress^20^. Feeding of 7-days old *w;Hml-Gal4,UAS-2xeGFP/+* (*Hml/+*) flies with PQ-supplemented sucrose solution led to an increased lethality of adult *Drosophila* after 18 hours in a dose dependent manner (0mM: 1.25 ± 1.25 %, 2mM: 2.5 ± 1.6 4%, 15mM: 27.78 ± 5.47 %, and 30mM: 38.75 ± 6.67 %) (Figure S1A). In agreement with previous studies^20^, 15mM PQ was chosen for further experiments (Figure 1A). First, we determined which systemic changes are associated with susceptibility to oxidative stress. Infection induced damage in enterocytes can result in gut leakage and subsequent death^17^. We tested if the death of flies after 18 hours on 15mM PQ food was associated with PQ-induced damage of gut enterocytes and resulting gut leakage, but we found no indication of gut leakage on 15mM PQ food (Figure S1B). Only very few flies on control food and PQ indicated gut leakage of the food color to thorax and abdomen which was unrelated to the treatment and not altered between the groups (control: 3.75%, 2mM PQ: 2.53 %, 15mM PQ: 1.11 % and 30mM PQ: 3.75 %) (Figure S1B). We investigated if aberrant systemic induction of immune signaling cascades are seen upon PQ-mediated oxidative stress. For this, gene expression levels of different known immune activated genes were analyzed by qPCR such as *unpaired* (*upd*) cytokines (*upd1*, *upd2*, *upd3*), *Jak/STAT* down-stream targets (*Turandot-A* (*TotA*), *suppressor of cytokine signaling at 36E* (*Socs36E*)), stress induced JNK activation (*puckered* (*puc*)) (Figure 1B), as well as expression of *nuclear factor ‘kappa-light-chain-enhancer’ of activated B-cells* (*Nfkb)*-induced antimicrobial peptides (AMP) and further cytokines such as *eiger*, *dpp* and *dawdle* (*daw*) (Figure S1C-D). These analyses did not reveal significant differences for these gene groups on a systemic level. In contrast, genes associated with carbohydrate metabolism such as *insulin receptor* (*InR*), *phosphoenolpyruvate carboxykinase* (*Pepck*), and *Thor* were significantly upregulated upon PQ-feeding (Figure 1C). These findings point to an activation of *forkhead box, sub-group O* (*foxo)*-target genes and a potentially reduced insulin signaling, despite the lack of significant alterations in the expression of the Insulin-like peptides (*Ilps*) *Ilp2*, *Ilp3* and *Ilp5* (Figure 1C).

**Figure 1.**
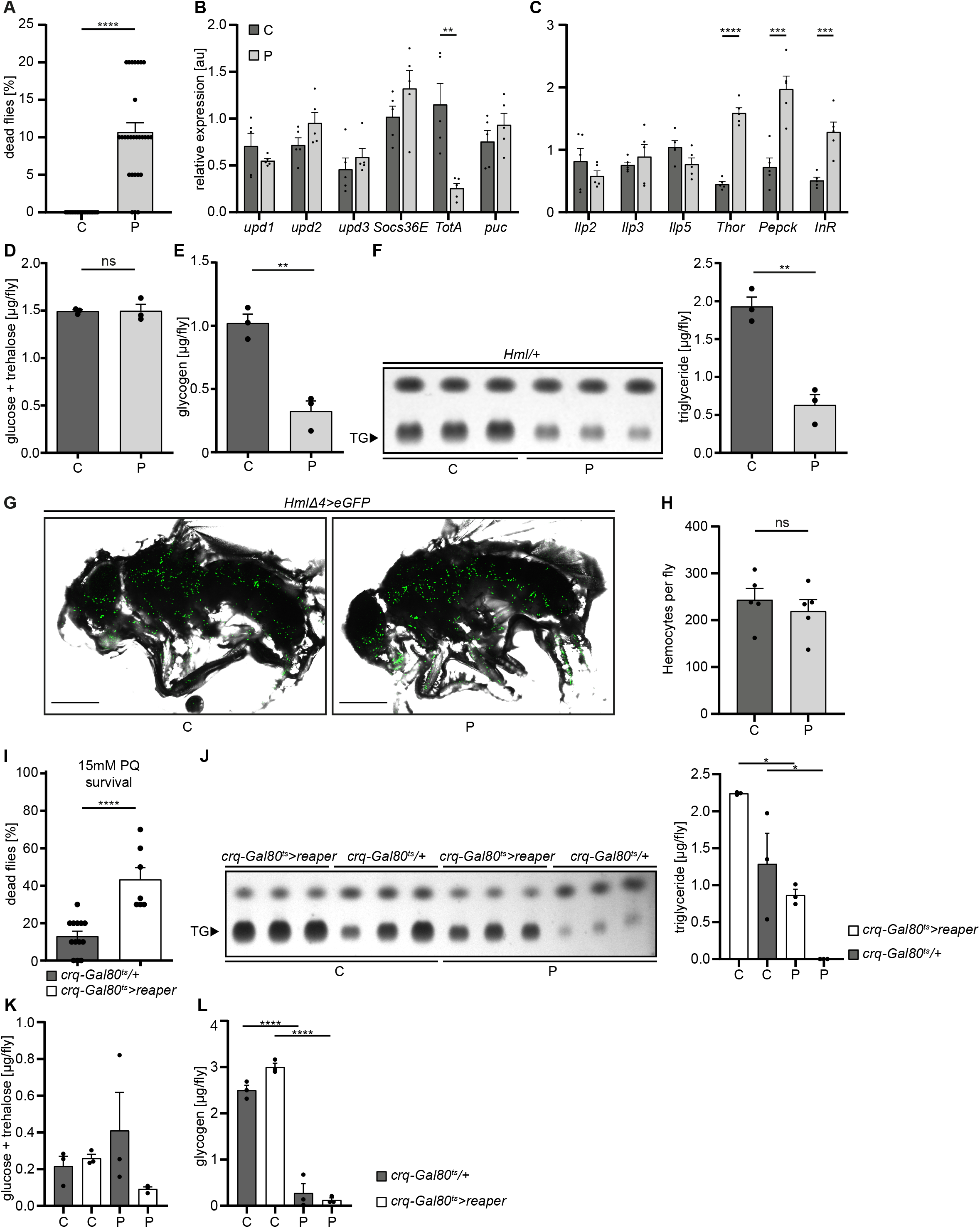
Loss of hemocytes increases the susceptibility of adult Drosophila to oxidative stress by Paraquat feeding. (A) Survival of *Hml/+* flies with control food (C) (n=20) and 15mM Paraquat (P) (n=28) at 29°C. Five independent experiments were performed. Student’s unpaired t-test: ****p<0.0001, each data point represents a sample with 10 flies. (B-C) RT-qPCR of whole flies were performed to investigate gene-expressions of the JAK-STAT-signaling pathway (B) and insulin-signaling pathway (C). All transcript levels were normalized to the expression of the loading control *Rpl1* and are shown in arbitrary units [au]. Each data point (n=5) represents a sample of three whole flies. Student’s unpaired t-tests were performed for each transcript to compare the gene-expression between control and treated flies: **p<0.01; ***p<0.001; ****p<0.0001. (D) Glucose and (E) glycogen levels of whole flies were measured in control and PQ-treated flies. Each dot (n=3) represents a sample containing three flies. Student’s unpaired t-test: **p<0.01. (F) Triglyceride (TG) amounts were determined via thin-layer chromatography (TLC). Left panel: Representative image of a TLC plate, each band represents a sample with ten flies. Right panel: Quantification thereof. n= 3 per group were analyzed. Student’s unpaired t-test: **p<0.01. (G) Representative images of 7 days old *HmlΔ4>eGFP* flies treated with 5% sucrose solution and 15mM PQ respectively. Scale bars = 500µm. (H) Hemocyte quantification of 7 days old *HmlΔ4>eGFP* flies treated with control food (5% sucrose solution) (n=5) or 15mM Paraquat (n=5). Each data point represents one fly. Statistical significance was tested using student’s unpaired t-test: p=0.508. (I) Survival of *crq-Gal80^ts^/+* control flies (n=13) and hemocyte depleted *crq-Gal80^ts^>reaper* flies (n=7) on control food (n=20) and 15mM Paraquat (n=28) at 29°C. Student’s unpaired t-test: ****p<0.0001, each data point represents a sample with 10 flies. (J) TG amounts in *crq-Gal80^ts^/+* control flies and hemocyte-deficient *crq-Gal80^ts^>reaper* flies on control food or Paraquat food. Left panel: Representative image of a TLC plate, each band represents a sample with ten flies. Right panel: Quantification thereof. n= 3 per group were analyzed. One-way ANOVA: *p<0.05. (K) Glucose and (L) Glycogen levels of whole flies were measured in *crq-Gal80^ts^/+* control flies and hemocyte-deficient *crq-Gal80^ts^>reaper* flies. Each dot (n= 3) represents a sample with three whole flies. One-way ANOVA: ****p<0.0001.

To test the consequences for energy mobilization and nutrient storage, we measured free glucose content, as well as glycogen and triglyceride levels in whole flies after 18 hrs on control or 15mM PQ. Although no changes in free glucose levels (control: 1.49±0.02µg and PQ: 1.50±0.07µg, ns p=0.954) (Figure 1D) were found, we observed a clear decrease in stored glycogen (control: 1.02±0.07µg and PQ: 0.33±0.08µg, \*\**p=0.003*) (Figure 1E) and stored triglycerides (Figure 1F) in PQ-treated flies compared to control. Next, we examined histological sections of flies to define which tissues might be affected. Hematoxylin and eosin (H&E) and Oil Red O (ORO) stainings showed a shrinkage in fat body cell size and stored lipid-filled vesicles in the abdominal fat body compared to flies kept under control conditions (Figure S1E-F). Fat body development, energy mobilization and storage have been recently connected to the presence, cell number and subsequent cross-talk of hemocytes with fat body^21^, therefore we quantified GFP^+^ hemocytes in *Hml/+* flies after 18 hrs on PQ or control food (Figure 1G-H). Our analysis did not reveal alterations in hemocyte numbers and especially no oxidative stress induced loss of hemocytes (control: 243.4±24.33 hemocytes per fly and PQ: 219.6±24.22 hemocytes per fly, ns *p=0.508*) (Figure 1G-H). To evaluate if hemocytes are essential contributors to the stress-mediated metabolic changes and susceptibility to oxidative stress, we used “hemocyte-deficient” *crq-Gal80^ts^>reaper* (*w;tub-Gal80ts/UAS-CD8-mCherry;crq-Gal4/UAS-reaper*) flies and tested their survival rate compared to control flies on PQ food (Figure 1I). Hemocyte-deficient flies displayed a significantly higher mortality rate on 15mM PQ (43.33±6.30%) compared to control flies (13.08±2.63%) (Figure 1I). Next, we analyzed triglycerides, as well as free glucose and glycogen levels in whole flies after 18 hrs on control or 15mM PQ. Interestingly, we found that hemocyte-deficient flies compared to the control genotype mobilized less triglycerides from the fat body during oxidative stress (Figure 1J). When we looked into free glucose levels, we found that these flies show a decrease in free glucose upon PQ feeding, whereas the control genotype showed similar free glucose levels as on normal food (Figure 1K). In contrast we did not detect any difference in glycogen levels between the two genotypes on control food and on PQ food (Figure 1L). Our findings show detrimental changes in energy mobilization upon oxidative stress in the absence of profound systemic overactivation of immune signaling cascades, however the presence or absence of hemocytes seem to be a key determinant of the susceptibility to oxidative stress and results in altered energy mobilization in adult *Drosophila*. This strongly points to an essential role for hemocytes in controlling stress response and organismal survival upon oxidative stress.

### Paraquat exposure generates diverse transcriptional states of adult hemocytes with a specific cell state associated with oxidative stress response

To gain further insights into the alterations induced in the adult fly upon oxidative stress and especially to the hemocyte compartment, we decided to take advantage of unbiased single nucleus RNA-sequencing (snRNA-seq). We isolated nuclei from control food and PQ-treated *Hml-DsRed.nuc* flies with a modified Frankenstein protocol^22^ and performed fluorescence activated nuclear sorting of DAPI^+^Draq7^+^ nuclei (Figure S2A). From two independent sorts we obtained a total of 29000 nuclei which were utilized for subsequent droplet-based barcoding of single nuclei, and generation of single nucleus cDNA libraries. We could successfully obtain gene expression data from 11839 nuclei (Figure S2B-D). Quality control analysis followed by unsupervised hierarchical clustering of all analyzed nuclei using Seurat package^23^ revealed 20 distinct nuclei clusters (C) with differential gene expression (Figure 2A). Cell identity was annotated according to the cell type-specific signatures determined by the Fly Cell Atlas consortium (Figure 2A)^24^. All annotated cell types were detected in PQ-treated and control flies (Figure 2B).

**Figure 2.**
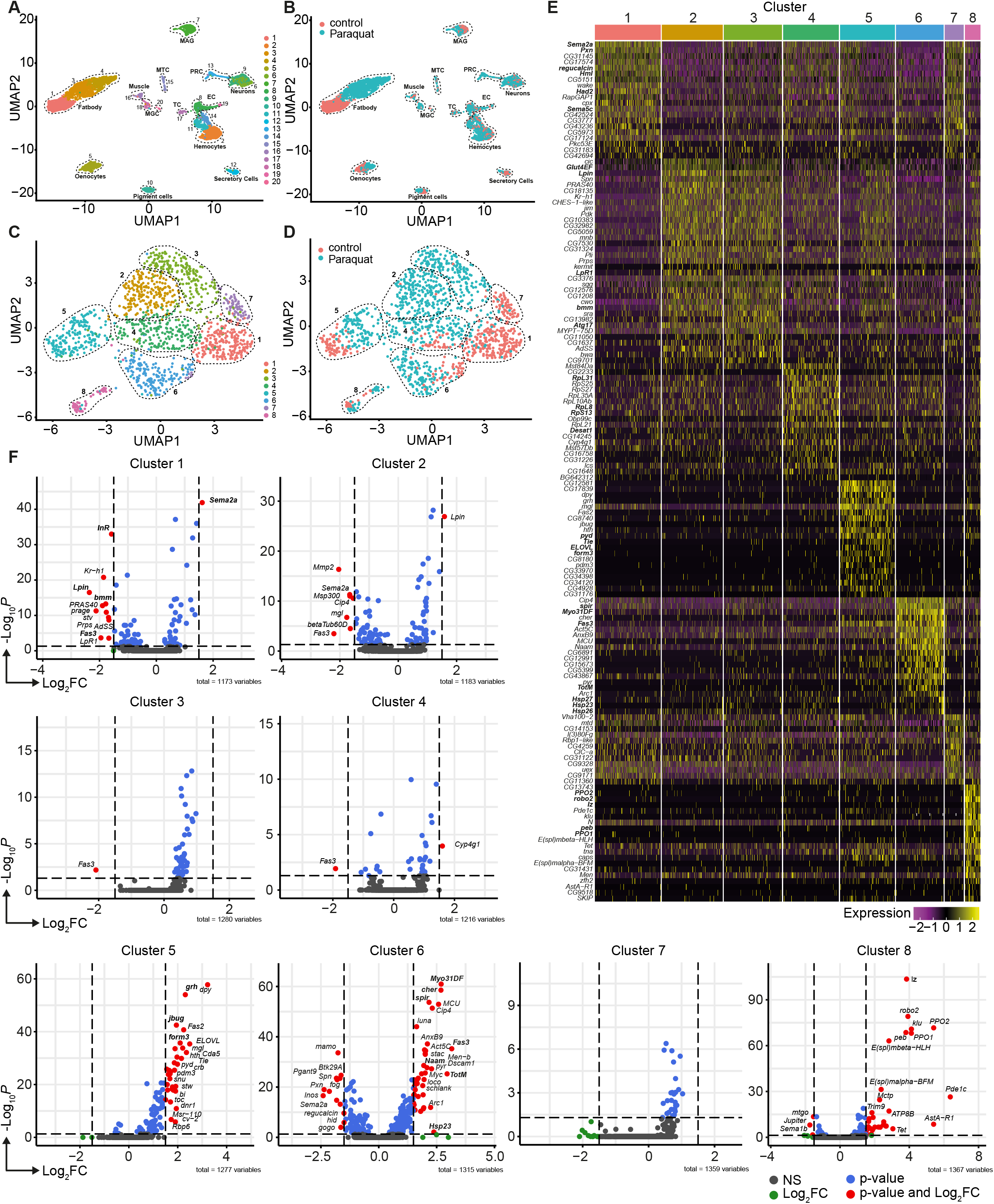
Unbiased single nuclei transcriptomic profiling identified diverse transcriptomic states of hemocytes associated with oxidative stress response. (A) Uniform Manifold Approximation and Projection (UMAP) visualization of single nucleus states from flies exposed to 15mM PQ containing or control food. Dashed lines indicate the broad cell types. Male accessory gland main cells (MAG), Malpighian tubule principal cells (MTC), male germline cells (MGC), outer photoreceptor cells (PRC), tracheolar cells (TC), epithelial cells (EC). Colors indicate distinct clusters. n=11839 nuclei are shown. (B) UMAP visualization of nuclei from **A** colored by treatment. Nuclei from control group are labeled in red and nuclei from PQ group are labeled in blue. Dashed lines indicate the broad cell type as **A**. (C) UMAP visualization of hemocytes from **A** after sub-setting and re-clustering. Colors and dashed lines indicate distinct clusters. n=1354 nuclei are shown. (D) UMAP visualization of hemocyte clusters colored by treatment. Nuclei from control group are labeled in red and nuclei from PQ group are labeled in blue. Dashed lines indicate distinct clusters. n=1354 nuclei are shown. (E) Heat map of top 20 DE genes for each hemocyte cluster. Gene names are indicated on the left. Colors in the heat map correspond to normalized scaled expression. (F) Volcano plots comparing pseudo bulk gene expression of individual hemocyte cluster vs. all hemocytes. The –log_10_-transformed adjusted P value (P adjusted, y-axis) is plotted against the log_2_-transformed fold change (FC) in expression between the indicated cluster vs remaining hemocytes (x-axis). Non-regulated genes are shown in grey, not significantly regulated genes are shown in green, significantly regulated genes with a FC<1.5 are shown in blue and significantly regulated genes with a FC>1.5 are shown in red.

Using cell type-specific signatures, three of the clusters were annotated as hemocytes (clusters: 2, 11 and 14) (Figure 2A-B). Re-scaling and clustering of all 1354 identified hemocyte nuclei revealed 8 distinct clusters (Figure 2C). We verified hemocyte-specific signature genes across these clusters including genes such as the transcription factor *Serpent* (*Srp*) and one or more of the following hemocyte signature genes: *Hemolectin* (*Hml*), *croquemort* (*crq*), *Nimrod C1* (*NimC1*), *eater,* and *Peroxidasin* (*Pxn*) (Figure S3A-L)^25,26^. Cluster 8 expressed various crystal cell-specific markers including *lozenge* (*lz*), *pebbled* (*peb*), *prophenoloxidase 1* (*PPO1*), and *prophenoloxidase* 2 (*PPO2*), whereas all other clusters seemed to be plasmatocytes (Figure S3A-L). In line with previous reports, the embryonic and larval hemocyte markers *Hemese* (*He*) (data not shown) and *singed (sn)* were not expressed across any of the identified clusters (Figure S3H). Interestingly, when we defined which hemocyte clusters were enriched in control flies or PQ-treated flies, the two plasmatocyte clusters C1 and C7 were primarily derived from control flies, whereas C2, C3, C4 and C6 are predominantly found in PQ-treated flies (Figure 2D and Figure S3M). The plasmatocyte cluster C5 as well as the crystal cell cluster C8 presented in both conditions to similar proportions (Figure 2D and Figure S3M).

Next, we analyzed the TOP20 differentially expressed (DE) genes across all hemocyte clusters. We found clear differences between the 7 distinct plasmatocyte clusters and the crystal cell cluster (Figure 2E and Table S1). The crystal cell cluster C8 showed a high expression of crystal cell specific genes including *lz*, *peb*, *PPO1* and *PPO2* (Figure 2E). Plasmatocyte cluster C1, which was mostly present in control conditions, had a remarkable expression of hemocyte specific marker genes such as *Hml* and *Pxn,* as well as migration-associated genes including the semaphorins *Sema2a and Sema5c* (Figure 2E). Yet this cluster also showed higher expression of genes contributing to metabolic regulations such as the L-gulonate 3-dehydrogenase *Had2* and the gluconolactonase *regucalcin* (Figure 2E). The cluster C7 and C1, mostly found in control flies, substantially shared gene expression profiles. In contrast, cluster C2 and C3 were nearly exclusively found in PQ-treated flies and showed expression of genes involved in metabolic adaptations. Due to this finding we verified that these cell clusters are *bona fide* hemocytes and not fat body cells. We analyzed the expression of characteristic fat body signature genes, such as *Trehalose-6-phosphate synthase 1* (*Tps1*), *CG16758*, *CG2233*, *CG4716*, *Ultrabithorax* (*Ubx*), *Calcium/calmodulin-dependent protein kinase I* (*CaMKI*), *Desaturase 1* (*Desat1*), *Phosphoribosyl pyrophosphate synthetase* (*Prps*), *CG34166* and *Apolipoprotein lipid transfer particle* (*Apoltp*) across all identified hemocyte clusters (Figure S4A). Metabolic adaptation within these clusters included high expression of the *glucose transporter 4 enhancer factor* (*Glut4EF*) or the *phosphatidate phosphatase Lipin* (*Lpin*) in C2 or the induced expression of the lipase *brummer* (*bmm*), the autophagy-associated gene *Atg17* or the *lipophorin receptor LpR1*, involved in uptake of neutral lipids from the hemolymph, in C3 (Figure 2E). Overall gene expression profiles in clusters C2 and C3 showed an induction of insulin signaling, increased autophagy and lipolysis pointing to adaptation of energy mobilization in these clusters upon PQ treatment (Figure 2E). Cluster C4 was characterized by a remarkable upregulation of genes involved in translation including many ribosomal proteins such as *RpL8*, *RpL31* or *RpS13* (Figure 2E). Furthermore, C4 included genes associated with unsaturated fatty acid metabolism such as *Desaturase 1 (Desat1)*. Similarly, cluster C5 also showed induction of genes involved in fatty acid metabolism such as *ELOVL fatty acid elongase 7* (*ELOVL*) (Figure 2E). Of note, we also found *Tie-like receptor tyrosine kinase* (*Tie*) specifically expressed in this cluster, which is described to bind PDGF– and VEGF-related factor (Pvf) ligands and contribute to cell survival and migration, as well as actin-associated binding proteins including *polychaetoid* (*pyd*) and *formin 3* (*form3*). All these gene expression changes are pointing to a plasmatocyte cluster with a gene expression profile associated with mobility and migration. Among all plasmatocyte clusters, we found cluster C6 which presented a strong immune activation phenotype with induction of *Jak/STAT* target genes including *Fasciclin 3* (*Fas3*) and *Turandot-M* (*TotM*) among its TOP20 DE genes, but also activation of oxidative stress response genes such as the heat shock proteins *Hsp23*, *Hsp26* and *Hsp27* or cytoskeletal genes including *myosin Myo31DF* or *spire* (*spir*) (Figure 2E).

Due to the partially shared expression profile of genes across hemocyte clusters, we wondered if there were genes exclusively expressed in individual clusters. However, cluster C1 showed that *Sema2a* seems to be exceptionally high expressed in these homeostatic plasmatocytes and genes associated with metabolic regulation (e.g. *InR*, *Lpin* or *bmm*) or Jak/STAT activation (e.g. *Fas3*), as seen in PQ-associated clusters, are downregulated (Figure 2F). Cluster C5 showed enrichment for genes such as the transcription factor *grainy head* (*grh*) but also cytoskeletal-associated genes such as *jitterbug* (*jbug*) and *form3,* suggesting a migratory potential of plasmatocytes from this cluster within their host tissues (Figure 2F). Interestingly, plasmatocyte cluster C6 presented with the most profound gene expression changes compared to all other clusters. Similar to our previous observations, we identified that genes specifically expressed in this cluster were associated with Jak/STAT pathway activation (e.g. *Fas3*, *TotM*), cytoskeletal rearrangements and migration (e.g. *Myo31DF*, *spir* or *cheerio* (*cher*)) and response to oxidative stress (e.g. *Nicotinamide amidase* (*Naam*) or *Hsp23*) (Figure 2F). We also included the crystal cell cluster C8 in this analysis and verified unique expression of crystal cell markers compared to the seven plasmatocyte clusters (Figure 2F). Our snRNA-seq analysis revealed that adult hemocytes respond differentially to oxidative stress with a specific cluster of plasmatocytes associated with immune activation and stress response. Despite our findings that on systemic level we did not detect a strong induction of immune activation upon oxidative stress, we now provide evidence that plasmatocyte cluster C6 showed a substantial immune response towards oxidative stress.

### Direct response to oxidative stress *in vitro* mimics the transcriptional changes seen in cluster C6 plasmatocytes *in vivo*

Plasmatocytes responded to PQ-mediated oxidative stress in different cellular states. We defined cluster C6 as a cell state which is associated with oxidative stress response and immune activation. Since cluster C6 showed gene expression associated with oxidative stress and immune activation we focused our further analysis on this cluster. To further substantiate our findings and address the signaling pathways that could lead to diverse transcriptional states, we inferred the transcription factor (TF) network based on gene expression data (Figure 3A). This analysis revealed that genes with binding sites for *Xbp1* were enriched. *Xbp1* is a transcription factor associated with stress response. In addition to this, the Nf-κB-related transcription factor *Relish* (*Rel*) and *kayak* (*kay*), a component of the *activator protein-1* (*AP-1*) transcription factor complex induced by JNK signaling, were also enriched in this cluster (Figure 3A). To determine if the gene expression signature of cluster C6 plasmatocytes reflects a functional state of plasmatocytes directly induced by oxidative stress, we treated S2 cells, a *Drosophila* plasmatocyte cell line, with different concentrations of PQ (15mM and 30mM) *in vitro* (Figure S5). First, we performed flow cytometric analysis of ROS levels via CellROX staining after 6 and 24 hours PQ treatment compared to controls (Figure S5A-B). We observed a dose-dependent increase in intracellular ROS at all time points analyzed with higher ROS levels at 6 hrs (0mM: 1903±64 median fluorescence intensity (MFI), 15mM: 6735±1096 MFI, and 30mM: 7435±237 MFI) compared to 24 hrs (0mM: 1236±135 MFI, 15mM: 4456±228 MFI, and 30mM: 5557±303 MFI) (Figure S5C). Increase in ROS and oxidative stress can lead to cellular damage such as damage in nuclear and mitochondrial DNA.

**Figure 3:**
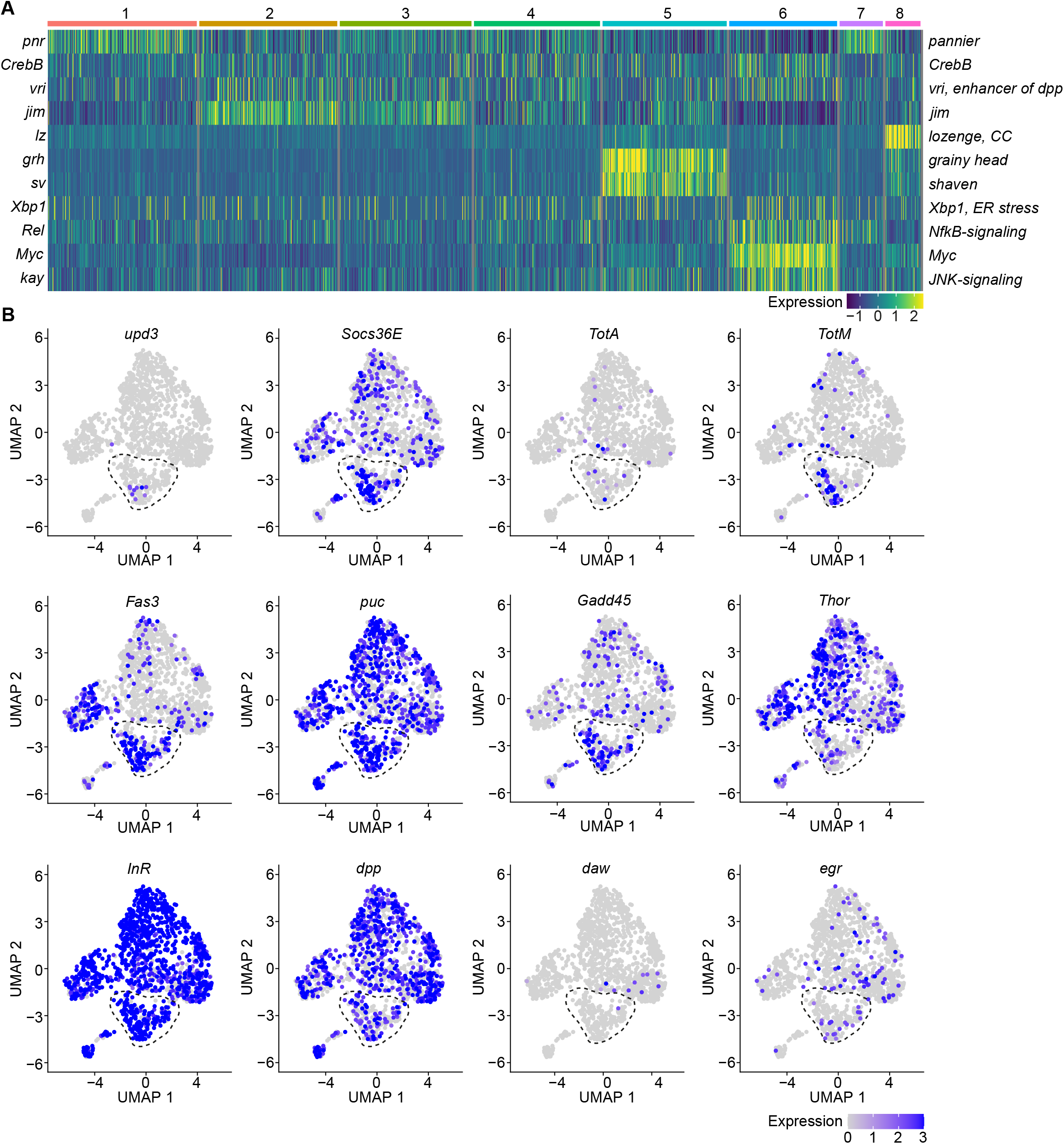
A specific cluster of plasmatocytes responds to oxidative stress with immune activation by Jak/STAT, DDR and JNK signaling. (A) Key transcription factors (TFs) regulating hemocytes states. Heat map of scaled TF activity at a single cell level in hemocytes states from Figure 2C. Colors in the heat map correspond to scaled expression. Numbers on top indicate the cluster identity. Labels on the left and right indicate the TFs. (B) Feature plots showing scaled expression of selected genes associated with JAK/STAT signaling (*upd3*, *Socs36E*, *TotA*, *TotM*, *Fas3*), JNK and DNA damage signaling (*puc*, *Gadd45*), insulin signaling (*Thor*, *InR*), TGFβ signaling (*dpp*, *daw*) and TNF signaling (*egr*).

We next tested if plasmatocytes directly exposed to oxidative stress might also experience this type of stress response by performing comet assays of treated and untreated S2 cells. Upon PQ treatment, S2 cells showed a remarkable and dose-dependent increase in DNA damage compared to untreated cells (0mM: 101.0±2.1, 5mM: 117.6±3.3, 10mM: 136.6±3.8, and 15mM: 147.8±5.5, data shown as olive tail moment) (Figure S5D). We were then wondering if PQ-treated S2 cells would resemble cluster C6 plasmatocytes *in vivo* and performed gene expression analysis of S2 cells treated with 15mM PQ after 6 and 24 hrs. Comparable to our results for cluster C6 *in vivo*, we found a profound induction of the Jak/STAT target genes by qRT-PCR. In S2 cells, we also detected a significant increase of the Jak/STAT cytokines *upd1*, *upd2* and *upd3* (Figure S5E), which we did not see before *in vivo*. Furthermore, we detected an increased expression of the JNK-target gene *puckered* (*puc*), but barely any induction of metabolic genes (Figure S5E). All these results further support our hypothesis that direct exposure of S2 cells with PQ *in vitro* mimics gene expression changes seen in a particular activated cell stage of plasmatocytes upon oxidative stress *in vivo*.

Because we found enrichment of genes associated with the Nf-κB-related transcription factor *Rel* in the immune-activated plasmatocyte cluster C6, we further verified if any other cytokines or AMPs are induced in PQ-treated S2 cells. We identified a slight but significant induction of the cytokine *daw*, but no other alterations in *dpp* or the AMPs *Drs*, *Dro* or *Mtk* (Figure S5F). The profound induction of Jak/STAT target genes together with the activation of JNK target genes and oxidative stress induced DNA damage in S2 cells *in vitro* highly support our hypothesis that cluster C6 plasmatocytes reflect a cellular state associated with the direct response to oxidative stress *in vivo*. Along this line, we went back to our snRNA-seq data and specifically searched for Jak/STAT target genes (*upd3*, *Socs36E*, *TotA*, *TotM* and *Fas3*) across all clusters (Figure 3B). Here we identified an enriched expression of these genes in cluster C6, and a unique expression of the pro-inflammatory cytokine *upd3* in cluster C6 which was not detected in any other hemocyte cluster (Figure 3B). Of note, we did not detect elevated *upd3* levels in gene expression profiling of whole flies (Figure 1B), but now unraveled elevated *upd3* induction in the particular hemocyte cell cluster C6 upon PQ treatment. Interestingly, we also identified that *Growth arrest and DNA damage-inducible 45* (*Gadd45*), a gene linking DNA damage signaling and stress induced JNK signaling, was enriched in plasmatocyte cluster C6 (Figure 3B). This could indicate that plasmatocytes within cluster C6 undergo oxidative stress mediated DNA damage, which results in activation of the DNA damage repair machinery but also JNK activation, similar to what we have seen in S2 cells *in vitro*. However within our snRNA-seq analysis, the JNK target gene *puc* was expressed across all hemocyte clusters analyzed *in vivo* without a specific induction in cluster C6 (Figure 3B). We looked again into the expression of metabolic genes including *Thor* and *InR* as well as other cytokines such as *dpp*, *daw* or *egr* but could not detect a specific enrichment in cluster C6 similar to our results from PQ-treated S2 cells *in vitro* (Figure 3B). These findings from the snRNA-seq analysis *in vivo* and S2 cells *in vitro* support the hypothesis that cluster C6 plasmatocytes represent a functional state of activated plasmatocytes directly responding to oxidative stress by JNK signaling and Jak/STAT pathway activation. Furthermore, we found evidence that cluster C6 specifically expresses the cytokine *upd3*, which could reflect an important mediator for oxidative stress response.

### Oxidative stress induces distinct cell states in the fat body, including the induction of a Jak/STAT responsive and the loss of an AMP-producing cell cluster

The diversification of plasmatocytes during oxidative stress and the defined cytokine producing immune-activated cell cluster C6 could play a key role in immune response to oxidative stress and also for interorgan communication. To further explore this hypothesis, we aimed to gain further insights into single-cell transcriptomic profiles of the *Drosophila* fat body during oxidative stress. Using cell type-specific signatures, three of the clusters were annotated as fat body cells (clusters: 1, 3 and 4) (Figure 2A-B). Re-scaling and clustering of all 3150 identified fat body nuclei revealed 8 distinct clusters (Figure 4A). First, we verified fat body-specific signature gene expression across these clusters (Figure S4B) and excluded expression of hemocyte-specific signature genes (Figure S4C). From the eight distinct fat body cell clusters, we identified that cluster C1, C6 and C8 were mainly enriched with cells obtained from control flies, whereas clusters C2-5 were almost exclusively derived from PQ treated flies (Figure 4B, Figure S4D). Only cluster C7 represents cells from both conditions (Figure 4B, Figure S4D). To define the transcriptional changes between these eight fat body cell clusters, we analyzed the TOP20 DE genes across all fat body cell clusters (Figure 4C, Table S2), as well as the unique regulated genes in each specific cluster compared to all others (Figure 4D). Here the cluster C1, mostly derived of fat body cells from control flies, showed an enrichment for gene induction associated with triglyceride storage, such as *Fatty acid synthase 1* (*FASN1*), *Glycerol-3-phosphate dehydrogenase 1* (*Gpdh1*) or *Acyl-CoA synthetase long-chain* (*Acsl*), as well as fat body signature genes (Figure 4C-D). Interestingly cluster C8, which mostly consisted of cells from control food, showed a distinct expression of AMP genes including *Dro*, *Drs* or *Diptericin-B* (*DptB*), however this cluster completely disappeared after PQ feeding (Figure 4C-D). In contrast, PQ treatment induced several fat body specific cell clusters which were not found in control conditions. For example, cluster C2, C4 and C5, which showed a strong gene induction of oxidative stress response genes, but also genes associated with Jak/STAT pathway (e.g. *Socs36E*, *upd3*, *Fas3*, *TotA*) and JNK activation (e.g. *puc*, *kay*) (Figure 4C-D). In all of these clusters, genes associated with triglyceride storage were significantly reduced. This is in line with our findings showing a reduction in triglyceride storage in the fat body upon oxidative stress (Figure 1F, Figure S1E-F). Overall our data revealed that the fat body has a unique heterogeneity upon oxidative stress with a clear separation of clusters with JNK and Jak/STAT activation signature, which could be either activated by the PQ-induced oxidative stress itself or by the cytokines released by immune-activated hemocytes, especially *upd3*.

**Figure 4:**
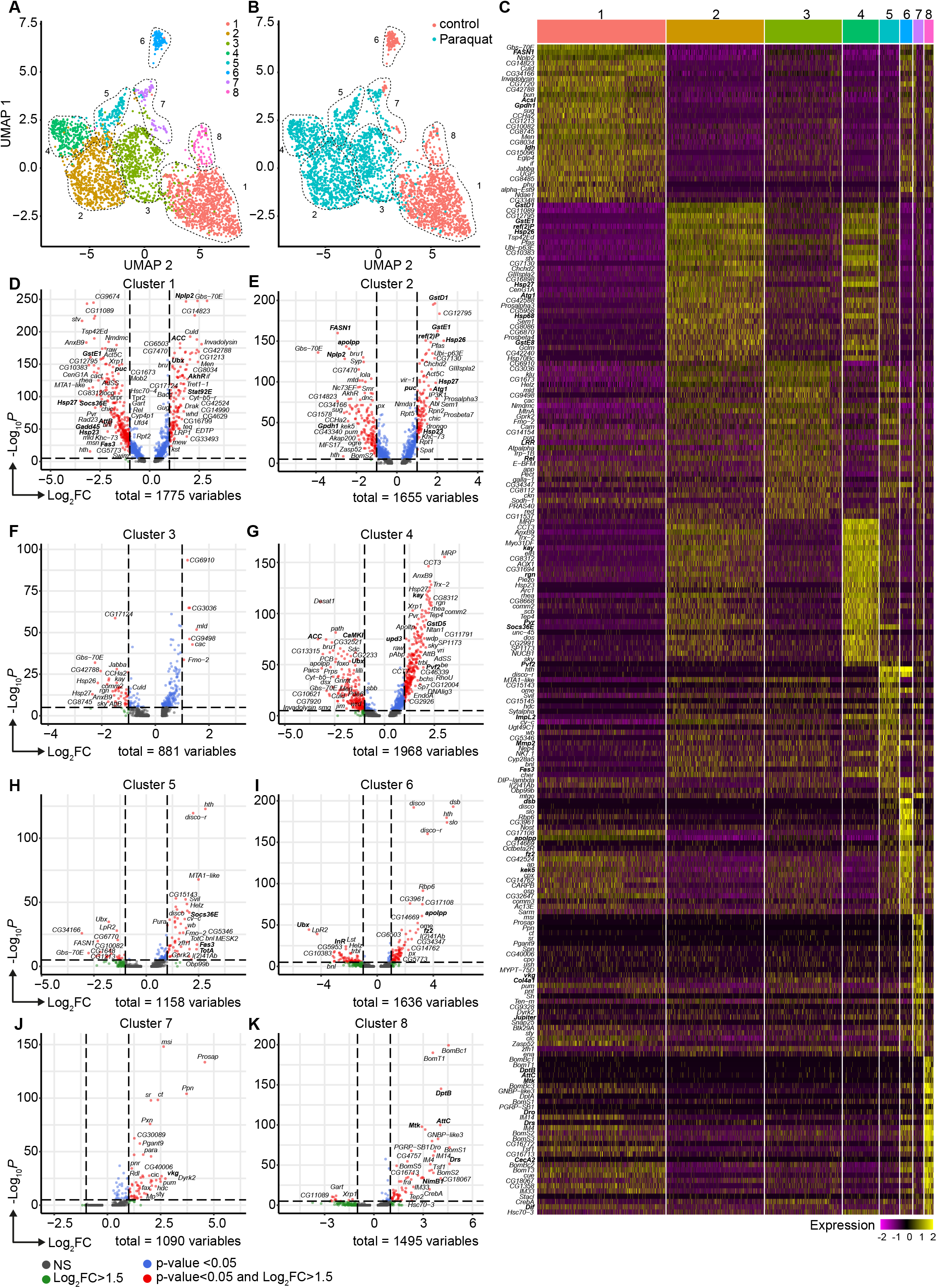
Oxidative stress induced different transcriptomic states in fat body cells, including cells with a distinct Jak/STAT activation signature. (A) UMAP visualization of fat body cells from Figure 2A by unsupervised clustering. Colors and dashed lines indicate different clusters. Each dot represents one nucleus. n=3150 nuclei are shown. (B) UMAP of fat body cell clusters split by treatment. Nuclei from control flies are labeled in red and nuclei from PQ treated flies are labeled in blue. Dashed lines indicate different clusters. Each dot represents one nucleus. n=3150 nuclei are shown. (C) Heat map of top 20 DE genes for each fat body cluster. Gene names indicated on the left. Fold change of gene expression is color coded as indicated in legend. (F) Volcano plots comparing pseudo bulk gene expression of individual fat body cluster vs. all fat body cells. The –log_10_-transformed adjusted P value (P adjusted, y-axis) is plotted against the log_2_-transformed fold change (FC) in expression between the indicated cluster vs remaining hemocytes (x-axis). Non-regulated genes are shown in grey, not significantly regulated genes are shown in green, significantly regulated genes with a FC<1.5 are shown in blue and significantly regulated genes with a FC>1.5 are shown in red.

### Loss of DNA damage signaling in adult hemocytes results in elevated *upd3* levels and a decreased survival on oxidative stress

To evaluate the role of immune activation in hemocytes during oxidative stress response, we aimed to specifically explore the implication of DNA damage signaling in hemocytes (Figure 1-3). Previously it has been observed that oxidative stress-induced DNA damage supports immune activation in macrophages via DNA damage signaling^27^. Based on our findings, we predict that also in adult hemocytes oxidative stress not only drives genome instability and activates repair, but plays a role in the induction and modulation of immune activation via specific gene expression changes in cluster C6 plasmatocytes. To test if DNA damage signaling is involved in the response of adult hemocytes to oxidative stress, we analyzed flies with a hemocyte-specific knock-down for different genes of the DNA damage repair (DDR) machinery including the DNA damage sensing kinases *telomere fusion* (*tefu*) and *meiotic 41* (*mei-41*) and the MRN complex protein *nbs*. Of note, by targeting adult hemocytes, mostly plasmatocytes are targeted which make up 95% of all hemocytes in the adult fly.

We generated *Hml>mei-41-IR* (*w;Hml-Gal4,UAS-2xeGFP/UAS-mei-41-IR*), *Hml>tefu-IR* (*w;Hml-Gal4,UAS-2xeGFP/UAS-tefu-IR*), *Hml>mei41-IR;tefu-IR* (*w;Hml-Gal4,UAS-2xeGFP/UAS-mei-41-IR;UAS-tefu-IR*) and *Hml>nbs-IR* (*w;Hml-Gal4,UAS-2xeGFP/UAS-nbs-IR*) flies (Figure S6A). First, we compared their susceptibility to PQ treatment after 18 hrs and found significantly increased susceptibility to the treatment across all DDR-deficient lines compared to the control genotype (*Hml/+*: 10.7±1.2%, *Hml>mei-41-IR*: 40.7±4.8%, *Hml>tefu-IR*: 20.7±4.3%, *Hml>mei41-IR;tefu-IR*: 37.5±4.8% and *Hml>nbs-IR*: 48.2±6.7%) (Figure 5A). In these flies, decreased survival after PQ did not correlate to an overall reduction in lifespan on normal food and a decrease in fitness (Figure S6B).

**Figure 5:**
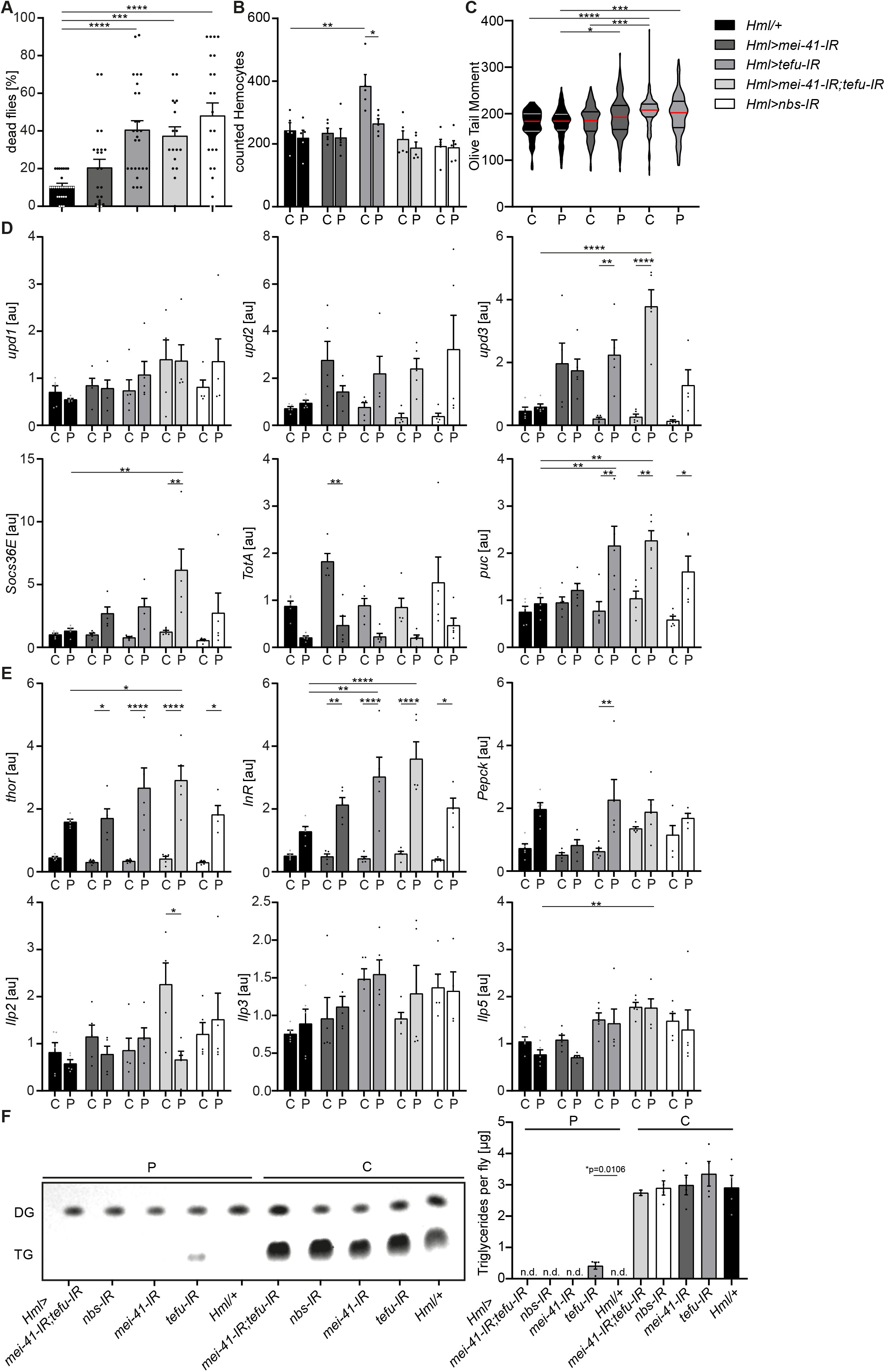
Loss of DNA damage signaling activity in hemocytes leads to an increase in systemic *upd3* levels and a higher susceptibility to oxidative stress. (A) Survival of *Hml/+* (n= 28), *Hml>mei41-IR (n*= 23); *Hml>tefu-IR (n*= 24); *Hml>mei41-IR,tefu-IR (n*= 16) and *Hml>nbs-IR* (n= 22) flies on control food and PQ. Each data point represents a sample of 10 flies. Mean ± SEM is shown. Three independent experiments were performed. One-way ANOVA: \*\*\**p<0.001*; \*\*\*\**p<0.0001*. (B) Hemocyte quantification of *Hml/+* control flies and DDR-knockdowns on control food (C) and 15mM Paraquat (P). Each data point represents one fly (n=5 for all groups). Mean ± SEM is shown. One way ANOVA: *Hml/+* (B)vs *Hml>tefu-IR* (C), \*\**p=0.0052*; *Hml>tefu-IR* (C) vs *Hml>tefu-IR* (P), \**p=0.03* (C) Comet assay of isolated hemocytes from *Hml/+* (C: n=73; P: n=129), *Hml>mei41-IR* (C: n=119; P: n=132) and *Hml>tefu-IR* (C: n=175; P: n=125) *-* flies on control food and PQ. Olive tail moment of comet assays of sorted hemocyte nuclei is shown. One-way ANOVA: *p<0.05; ***p<0.001; ****p<0.0001. (D) Gene expression analysis for *Jak/STAT* target genes via RT-qPCR in *Hml/+* control flies and DDR-knockdowns on control food (C) and 15mM Paraquat (P). All transcript levels were normalized to the expression of the loading control *Rpl1* and are shown in arbitrary units [au]. Each data point (n=5) represents a sample of three individual flies. Mean ± SEM is shown. Two-way ANOVA: \**p<0.05*; \*\**p<0.01*; *** *p<0.001*; \*\*\*\**p<0.0001*. (E) Gene expression analysis for insulin signaling target genes and *Ilps* via RT-qPCR in *Hml/+* control flies and DDR-knockdowns on control food (C) and 15mM Paraquat (P). All transcript levels were normalized to the expression of the loading control *Rpl1* and are shown in arbitrary units [au]. Each data point (n=5) represents a sample of three individual flies. Mean ± SEM is shown. Two-way ANOVA: \**p<0.05*; \*\**p<0.01*; *** *p<0.001*; \*\*\*\**p<0.0001*. (F) Triglyceride (TG) levels of *Hml/+*, *Hml>mei41-IR*, *Hml>tefu-IR, Hml>mei41-IR,tefu-IR* and *Hml>nbs-IR* flies on control food and PQ determined via thin-layer chromatography (TLC). One representative TLC is shown (left panel). Quantification thereof is shown on the right. Each data point (n=4 per group) represents a sample of ten individual flies. Mean ± SEM is shown. n.d. = not detectable. One-way ANOVA was performed for both groups (15mM PQ and control) separately: \**p=0.0106*.

Next, we inquired if DDR-deficiency in hemocytes might increase susceptibility of the fly to other stresses. For this we challenged all genotypes with starvation. Importantly, we did not observe a decreased survival of DDR-deficient flies on starvation compared to the control genotype, indicating that the observed susceptibility to oxidative stress of flies with a DDR-deficiency in hemocytes is specifically caused by oxidative stress due to PQ treatment (Figure S6C). As defects in the DDR machinery in hemocytes might already affect the development during embryonic and larval stages, we used a hemocyte-specific temperature-inducible knock-down of *mei-41*, *tefu* and *nbs1* which was induced by a temperature shift to 29°C directly after eclosion from the pupae to induce a knockdown in adult hemocytes only. We analyzed the lifespan of *Hml-Gal80^ts^>mei-41-IR* (*w;Hml-Gal4,UAS-2xeGFP/UAS-mei-41-IR;tub-Gal80*), *Hml-Gal80^ts^>tefu-IR* (*w;Hml-Gal4,UAS-2xeGFP/UAS-tefu-IR;tub-Gal80*), *Hml-Gal80^ts^>mei-41-IR;tefu-IR* (*w;Hml-Gal4,UAS-2xeGFP/UAS-mei-41-IR;tub-Gal80/UAS-tefu-IR*) and *Hml-Gal80^ts^>nbs-IR* (*w;Hml-Gal4,UAS-2xeGFP/UAS-nbs-IR;tub-Gal80*) compared to control flies *Hml-Gal80^ts^/+* (*w;Hml-Gal4,UAS-2xeGFP/+;tub-Gal80*) and again found no alterations in overall lifespan of all genotypes analyzed on control food except a slight decrease in lifespan of *Hml-Gal80^ts^>mei-41-IR;tefu-IR.* (Figure S6D). In contrast, we again found a higher susceptibility to oxidative stress via PQ-feeding in the DDR-deficient lines compared to control (*Hml-Gal80^ts^/+*: 30.3±5.2%, *Hml-Gal80^ts^>mei-41-IR*: 36.2±4.3%, *Hml-Gal80^ts^>tefu-IR*: 52.4±6.9%, *Hml-Gal80^ts^>mei-41-IR;tefu-IR*: 72.3±6.3% and *Hml Hml-Gal80^ts^>nbs-IR*: 36.2±4.3%) (Figure S6E), which directly confirms our previous results and conclusions (Figure 5A).

As we have seen before that loss of hemocytes is detrimental for the survival on oxidative stress, we wanted to exclude a reduction in hemocytes in the DDR deficient fly lines as a cause of the higher susceptibility to oxidative stress. We quantified GFP^+^ hemocytes in *Hml>mei-41-IR*, *Hml>tefu-IR*, *Hml>mei-41-IR;tefu-IR* and *Hml>nbs-IR* flies (Figure 5B, Figure S6A). We did not find alteration in hemocyte number across all fly lines on control food (with the exception of an increase in hemocytes in the *Hml>tefu-IR* line), nor did we observe any effect of PQ treatment on the number of hemocytes in DDR-deficient flies (Figure 5B, Figure S6A).

Next, we analyzed if the loss of the DDR signaling machinery via knockdown of *tefu* and *mei41* results in increased DNA damage in hemocytes. Comet assays of isolated hemocytes from DDR deficient flies and the control genotype on PQ and control food only indicated a slight increase in DNA damage in these flies, which demonstrates that the higher susceptibility of DDR deficient flies is not due to a loss of hemocyte numbers or a substantial increase in DNA damage, but rather due to changes in the downstream DDR-signaling events modulating immune activation of hemocytes (Figure 5C). Other studies demonstrated that upon tissue damage JNK-mediated secretion of the JAK/STAT cytokines upd1, upd2 and upd3 is a critical step for innate immune signaling^28,29^. Furthermore, DNA damage signaling is essential to induce such an immune response and to regulate insulin signaling^30^. To elucidate the implication of DDR signaling on hemocyte activation and oxidative stress resistance, we measured the gene expression levels of *upd* cytokines as well as JNK and Jak/STAT targets in DDR deficient flies and controls on PQ and control food. Loss of DNA damage signaling in hemocytes induced a significant induction of *upd3* expression upon PQ treatment in *Hml>tefu-IR* and *Hml>mei-41-IR;tefu-IR* flies (Figure 5D). In contrary, *upd1* and *upd2* were not significantly changed in DDR-deficient flies (Figure 5D). We measured the Jak/STAT target genes *Socs36E* and *TotA* on systemic level. A slight induction of *Socs36E* expression in all DDR-deficient genotypes on PQ, but only *Hml>mei-41-IR;tefu-IR* flies showed a significant induction of *Socs36E* upon PQ treatment compared to control food and *Hml/+* flies on PQ (Figure 5D). *TotA* expression tended to be reduced upon PQ treatment across all genotypes analyzed. However, when we checked the levels of *puc* as a *JNK signaling* target gene, we found a significant induction on PQ treatment in *Hml>tefu-IR*, *Hml>mei-41-IR;tefu-IR* and *Hml>nbs-IR* flies, which was not seen in control flies upon PQ treatment (Figure 5D). When we analyzed the gene expression of genes involved in metabolism and energy mobilization, we detected a significant induction and dysregulation of *Thor* and *InR* in all DDR-deficient genotypes upon PQ treatment compared to control flies (Figure 5E), as well as a slight induction of *Pepck1* (Figure 5E). To exclude that these changes in energy mobilization are due to altered expression of *Ilps*, we analyzed the levels of *Ilp*2, *Ilp3* and *Ilp5* in all genotypes on control and PQ-food, but could not detect remarkable alterations here (Figure 5E). To further exclude that the higher susceptibility of DDR-deficient flies and seen alterations in stress response are due to inefficient mobilization of triglycerides, we measured stored triglycerides across all genotypes and found efficient mobilization of triglyceride storage upon PQ treatment in all genotypes compared to control food (Figure 5F).

From our results we can conclude that DNA damage signaling in adult hemocytes, mostly consisting of plasmatocytes, essentially determines susceptibility to oxidative stress. Loss of DNA damage signaling in hemocytes results in upregulation of *JNK* signaling and elevated expression of the pro-inflammatory cytokine *upd3*, as well as transcriptional alterations in energy mobilization and insulin signaling. Based on our previous results on the role of hemocytes during oxidative stress, we can therefore assume that the DNA damage signaling machinery in hemocytes is activated by oxidative stress and plays an essential modulatory and rate-limiting role for subsequent immune activation.

### Adult hemocytes control susceptibility to oxidative stress by JNK signaling and *upd3* release

To this point, our data suggests that oxidative stress-induced DNA damage signaling in hemocytes limits immune activation and stress response upon oxidative stress. As we found that increased *upd3* levels correlate with a higher susceptibility to oxidative stress of flies with DDR-deficient hemocytes, we aimed to test if hemocyte-derived *upd3* is a major source to regulate susceptibility to PQ-induced oxidative stress. To do so, we either over-expressed *upd3* in hemocytes in *Hml>upd3* (*w;Hml-Gal4,UAS-2xeGFP/UAS-upd3*) flies or performed a hemocyte-specific knockdown of *upd3* in *Hml>upd3-IR* (*w;Hml-Gal4,UAS-2xeGFP/UAS-upd3-IR*) flies (Figure S6F). Both genotypes showed a normal overall lifespan compared to the *Hml/+* control (Figure S6G), but when we compared their susceptibility to PQ-induced oxidative stress, *Hml>upd3* flies showed a drastically decreased survival on PQ, whereas *Hml>upd3-IR* flies did not show a changed survival on PQ treatment after 18 hours (Figure 6A). We tested *upd3* expression levels on a whole fly level and found a slight but not significant induction in *Hml/+* flies after PQ treatment compared to control food, in line with our previous results (Figure 1B and 6B). As expected, *Hml>upd3* flies showed increased levels of *upd3* in both treatment groups compared to the other genotypes, but also showed a significant increase of *upd3* upon PQ compared to control food (Figure 6B). Surprisingly, *Hml>upd3-IR* flies still showed a slight induction of *upd3* upon PQ treatment, which could potentially be derived from another cellular source after eliminating hemocyte-derived *upd3*, for example fat body cells, as seen in our snRNA-seq analysis.

**Figure 6:**
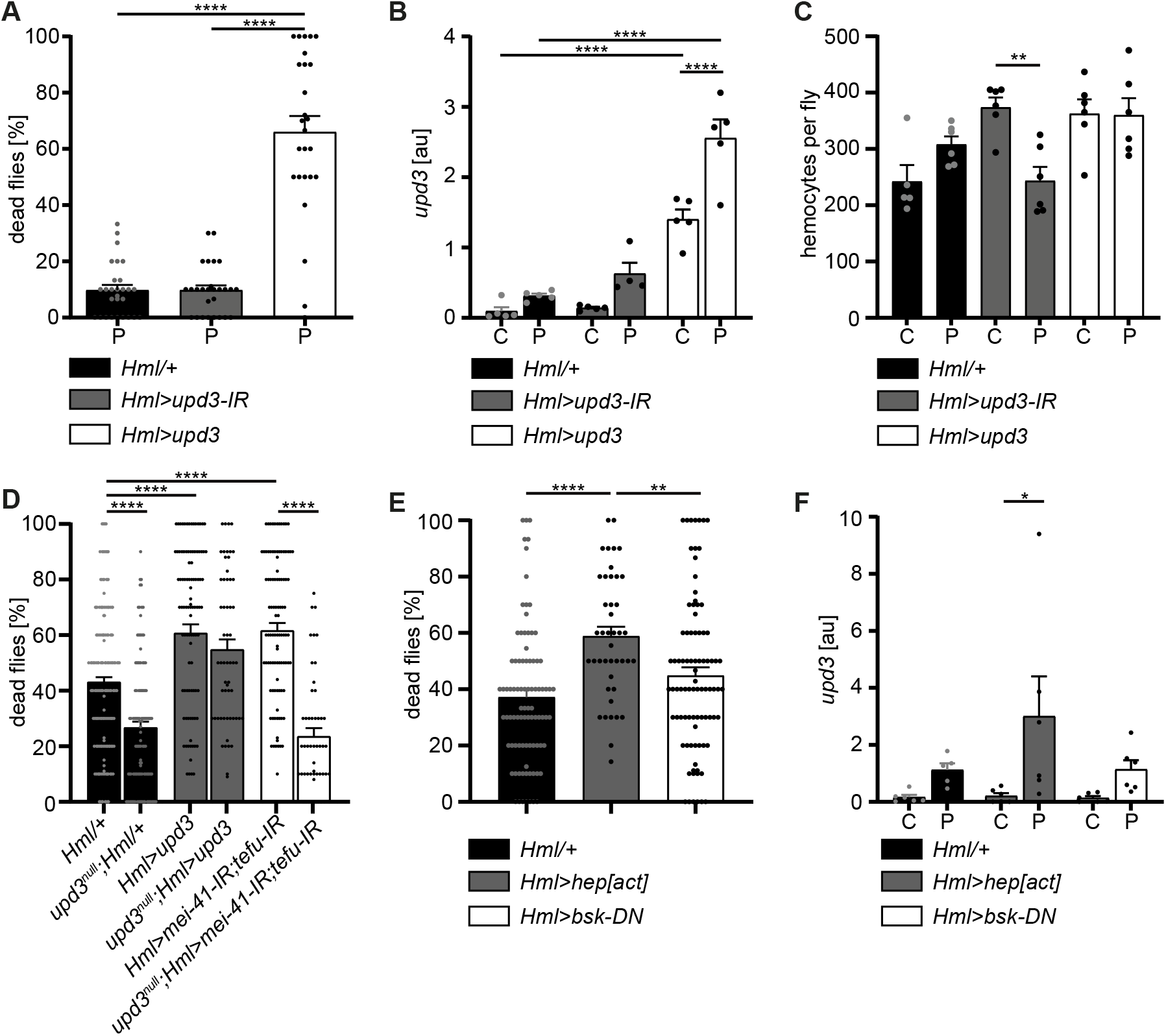
Hemocyte-derived *upd3* controls susceptibility to oxidative stress in adult *Drosophila*. (A) Survival of *Hml/+* (n=28), *Hml>upd3-IR* (n=27) and *Hml>upd3* (n=26) flies on 15mM PQ food. Each data point represents a sample of 10-15 flies. Mean ± SEM is shown. Three independent experiments were performed. One-way ANOVA: \*\*\*\**p<0.0001*. (B) Gene expression analysis for *upd3* via RT-qPCR in *Hml/+, Hml>upd3-IR* and *Hml>upd3* flies on control food (C) and 15mM Paraquat (P). Each data point (n=5-6) represents a sample of three individual flies. Mean ± SEM is shown. Two-way ANOVA: \*\*\*\**p<0.0001*. (C) Hemocyte quantifications in *Hml/+, Hml>upd3-IR* and *Hml>upd3* flies on control food and 15mM PQ food. Each data point represents one fly (n= 5-6). Mean ± SEM is shown. One-way ANOVA: **p<0.01. (D) Survival of *Hml/+* (n=241), *upd3^null^;Hml/+* (n=122), *Hml>upd3* (n=104), *upd3^null^;Hml>upd3* (n=57), *Hml>mei-41-IR;tefu-IR* (n=103) and *upd3^null^;Hml>mei-41-IR;tefu-IR* (n=50) flies on 15mM PQ food at 29°C. Each data point represents a sample with 10-15 flies. Mean ± SEM is shown. One-way ANOVA: \*\*\*\**p<0.0001*. (E) Survival of *Hml/+* (n=78), *Hml>hep[act]* (n=38) and *Hml>bsk-DN* (n=80) flies on 15mM PQ food. Each data point represents a sample with 10 flies. Mean ± SEM is shown. One-way ANOVA: \*\**p<0.01*; \*\*\*\**p<0.0001*. (F) Gene expression analysis for *upd3* via RT-qPCR of *Hml/+*, *Hml>hep[act]* and *Hml>bsk-DN* flies on control food (-) and 15mM Paraquat (+). Each data point (n=5-6) represents a sample of three individual flies. Mean ± SEM is shown. Two-way ANOVA: \**p<0.05*.

To identify if the higher susceptibility of *Hml>upd3* flies to oxidative stress was due to reduced hemocyte numbers, we quantified hemocytes across all three genotypes, but did not see any significant changes of hemocyte number in both genotypes compared to *Hml/+* control flies on control food and PQ (Figure 6C and Figure S6F). Only in the *Hml>upd3-IR* flies we found a reduction of hemocytes between flies on control food and PQ, but without consequences for their susceptibility to oxidative stress (Figure 6C and Figure S6G). Our data indicate that elevation of hemocyte-derived *upd3* is sufficient to increase the mortality of adult flies on PQ-mediated oxidative stress. Yet we could not exclude other cellular sources than hemocytes for *upd3* upon oxidative stress. Therefore, we wanted to probe if hemocyte-derived *upd3* is sufficient to increase the susceptibility to PQ treatment in the absence of other cellular sources. We backcrossed *Hml>upd3* flies to *upd3*-null mutants (*upd3^nul^*^l^) to generate *upd3^null^;Hml>upd3* flies (*upd3^null^;Hml-Gal4,UAS-2xeGFP/UAS-upd3)* and compared their survival on PQ to *Hml/+* and *upd3^null^* flies (Figure 6D). Of note, we found that *upd3^null^*mutants had a significant reduced susceptibility to oxidative stress compared to *Hml/+* flies, however we found a decreased survival in *upd3^null^;Hml>upd3* flies when hemocytes alone overexpressing *upd3* similar to the higher susceptibility of *Hml>upd3* flies (Figure 6D). We then wanted to test if loss of *upd3* would also rescue the higher susceptibility of flies with DDR-deficient hemocytes and backcrossed *Hml>mei41-IR;tefu-IR* to *upd3^null^*flies to generate *upd3^null^;Hml>mei41-IR;tefu-IR* flies (*upd3^null^;Hml-Gal4,UAS-2xeGFP/UAS-mei41-IR;UAS-tefu-IR*) (Figure 6D). Indeed we found a substantially decreased susceptibility of these flies to oxidative stress compared to *Hml>mei41-IR;tefu-IR* flies, supporting our hypothesis that loss of DNA damage signaling in hemocytes and subsequently increased levels of upd3 render adult flies more susceptible to oxidative stress.

Previous studies highlighted that JNK signaling can induce *upd3* expression in hemocytes^12^. In line with these reports, we found enrichment of genes with *kayak/AP-1* binding motifs in the plasmatocyte subset C6 which showed further upregulation of Jak/STAT target genes and also *upd3* expression (Figure 3). Therefore, we wanted to verify if *JNK* activation in hemocytes acts as a direct downstream signal activated by oxidative stress and subsequently induces the release of the pro-inflammatory cytokine *upd3*. We either overexpressed a constitutive active form of *JNKK hemipterous* (*hep*) in hemocytes using *Hml>hep[act]* (*w;Hml-Gal4,UAS-2xeGFP/UAS-hep[act]*) flies or overexpressed a dominant-negative version of the Jun kinase *basket* (*bsk*) in hemocytes using *Hml>bsk-DN* (*w;Hml-Gal4,UAS-2xeGFP/UAS-bsk-DN*). We compared their survival on PQ after 18 hrs and in line with our previous results, we found an increased susceptibility of *Hml>hep[act*] flies to PQ treatment, but not of *Hml>bsk-DN* flies compared to the control genotype (Figure 6E). We again analyzed the overall lifespan of these flies on normal food and could not find differences between the genotypes (Figure S6H). Furthermore, when we analyzed *upd3* expression in these flies upon PQ treatment, we found a significant induction of *upd3* expression after PQ treatment in *Hml>hep[act*] compared to control food, but only a slight increase in *Hml/+* and *Hml>bsk-DN* flies upon PQ feeding (Figure 6F). These findings support a substantial role of JNK-mediated induction of *upd3* in hemocytes upon oxidative stress.

In summary, our study revealed a defined immune-responsive subset of plasmatocytes during oxidative stress. We show that this oxidative stress-mediated immune activation seems to be regulated by DNA damage signaling in hemocytes, which controls JNK activation and subsequent *upd3* expression. Loss of DNA damage signaling and over-induction of JNK signaling or upd3 in hemocytes release renders the adult fly more susceptible to oxidative stress.

## Discussion

Similar to vertebrate macrophages, oxidative stress triggers immune activation in *Drosophila* hemocytes^18^. However, this immune activation was not obvious on a systemic level upon PQ-mediated oxidative stress. In contrast, localized oxidative stress-induced immune activation in damaged tissues is an essential step of the *Drosophila* immune response to infection or injury^19,31^. However, the precise contribution of *Drosophila* macrophages in this immune activation, in the organ-to-organ communication during stress response and the implication in the organism’s survival upon increased oxidative stress remained so far ill-defined. Overall, we describe the induction of diverse cellular states of adult hemocytes, namely plasmatocytes, during oxidative stress with a particular subset of plasmatocytes (cluster C6) which directly responds to oxidative stress by immune activation. We demonstrated that this immune activation seems to be at least partially controlled by the DNA damage signaling machinery in hemocytes. Loss of *nbs*, *tefu* or *mei-41* in hemocytes renders the organism more susceptible to oxidative stress without significantly affecting oxidative stress mediated DNA damage, but with an enhanced induction of JAK/STAT and JNK signaling, accompanied by changes in genes associated with energy mobilization and storage. In line with that, we describe substantial changes in the fat body on the transcriptomic level with diverse cell states, indicating decrease of triglyceride storage and induced response to oxidative stress and immune activation, but loss of AMP-producing cells. Finally, we showed that the control of hemocyte-derived *upd3* is key for the survival of the organism upon oxidative stress and that induction of the *JNK/upd3* signaling axis in hemocytes is sufficient to increase the susceptibility to oxidative stress. Hence, we postulate that hemocyte-derived *upd3* is most likely released by the activated plasmatocyte cluster C6 during oxidative stress *in vivo* and is subsequently controlling energy mobilization and subsequent tissue wasting upon oxidative stress.

*Drosophila* hemocytes have been described before to react to PQ-mediated or infection-mediated intestinal damage via release of cytokines to promote ISC proliferation in the gut^14,17^. PQ is a widely used model in *Drosophila* to induce oxidative stress via oral uptake^32^. In many reports the gut has been described to be the most affected organ and ISC proliferation is an essential step to regenerate the injured intestinal epithelium^33,34^. PQ treatment has been described to cause gut leakage and therefore this could be a potential cause of death for the fly^14^, though we did not find signs of gut leakage within the 18 hours PQ treatment with a concentration of 15mM. Aside from the gut, oral PQ treatment was shown to induce oxidative stress in many organs including hemocytes^35^, oenocytes^36^ and the brain^37^, indicating that the induced oxidative stress is not only localized to the gut tissue. PQ-induced oxidative stress was described before to cause neurodegeneration with parkinsonism-like symptoms, accompanied by strong immune activation and upregulation of *JNK* signaling in the CNS^37^, an activation profile similar to the identified activated plasmatocyte cluster C6 and the activated fat body clusters C2 and C4 in our snRNA-seq analysis. The snRNA-seq analysis further revealed distinct hemocyte clusters with seven plasmatocyte clusters and one crystal cell cluster. Whereas the plasmatocyte clusters C1 and C7 seem to represent cellular states found in steady state conditions only, we identified several other clusters which seemed to be exclusive or at least enriched under PQ treatment. Interestingly, we only identified one cluster, C6, which specifically responded to oxidative stress via immune activation and upregulation of oxidative stress response genes but also *Jak/STAT* and *JNK* target genes. We postulate that these cells might be directly exposed to the oxidative stress because their signature resembles the one of S2 cells directly treated with PQ *in vitro*. Even though oxidative DNA damage in hemocytes was followed by immune activation *in vitro* and *in vivo*, it remains to be determined if this activation is induced by cytokines derived from damaged organs during oxidative stress or by the oxidative stress directly. The hypothesis that this cluster is directly exposed to oxidative stress is further supported by an upregulation of genes such as *Hsps* in this cluster which are known to be directly induced by oxidative stress^38^. Furthermore, we found evidence that oxidative DNA damage and induced DNA damage signaling dynamically modulate immune activation cascades in cluster C6 plasmatocytes. This was supported by the increased DNA damage in S2 cells by PQ treatment but also the enrichment of DDR-associated genes such as *Gadd45* in cluster C6, which is associated with *JNK* activation and is further implicated in oxidative stress induced DNA damage^37^. Constant DNA damage signaling and repair is an essential step in all tissues to maintain tissue homeostasis^39^.

In our study, we point to a potential role of DNA damage signaling as a rate-limiting step for immune activation and response to oxidative stress in hemocytes. We did not find indications of canonical DNA damage response, such as induced apoptosis in hemocytes due to the oxidative DNA damage. Non-canonical DNA damage sensing has been implicated in immune responses in vertebrates and invertebrates^30,37,40,41^. It was demonstrated that vertebrate macrophages *in vitro* respond similarly to immune stimuli such as infection by synergistic activation of DDR and type I interferon signaling to induce downstream immune signaling cascades^27^. Here, we provide new evidence in *Drosophila* that potentially non-canonical DNA damage is used by immune cells *in vivo* to modulate and especially control their immune response including pro-inflammatory cytokine release upon oxidative stress. Localized DNA damage in larval epithelial cells has been shown to induce non-canonical DNA damage activity leading to the secretion of *upds* and the regulation of hemocyte expansion and activation, as well as subsequent hemocyte-mediated regulation of energy storage in the fat body^30^. In further support of our results, one report previously demonstrated that loss of DNA damage signaling machinery induced the release of *upd3* in hemocytes^14^. Most likely, the identified plasmatocyte cluster C6 is a central cellular source of this signal. However, we also detected *upd3* induction in the PQ-induced fat body cell cluster C4 as potential secondary source of upd3, which needs to be further determined in future studies. Furthermore, anatomical localization of this upd3-producing plasmatocyte cluster, as well as spatial distribution and cross-talk of the different plasmatocyte and fat body cell clusters would be of interest. We describe a strong connection between the control of *upd3* expression by hemocytes and the susceptibility of the adult fly to oxidative stress. Upon oxidative stress, we observe decreased triglycerides and glycogen stores and altered gene expression such as induced expression of lipolytic enzymes and other *foxo* target genes indicating energy mobilization from the adult fat body, which was further supported by the transcriptomic profiling of fat body cells. Accumulating evidence throughout our study points to an implication of hemocyte-derived *upd3* in the control of energy mobilization and tissue wasting during oxidative stress. Our data from hemocyte-deficient flies indicating defects in energy mobilization and maintenance of free glucose levels further support this assumption. Our presented findings are in line with our previous studies where we showed that upon high fat diet hemocytes produce high levels of *upd3* which induces insulin insensitivity in many tissues by activating *Jak/STAT signaling* in these organs and inhibiting the insulin signaling pathway^12^. Furthermore, we and others described a central role for hemocyte derived *upds* in the control of glucose metabolism in muscles during steady state and upon infection^13,42^. As shown before, hemocytes produce high levels of *upd3* upon infection with *Mycobacterium marinum* which also caused tissue wasting and extensive energy mobilization in a *foxo*-dependent manner^11,43^.

Our study provides further evidence that hemocytes serve as essential signaling hubs in “organ to organ communication” during oxidative stress, which is in line with previous studies in immune challenge as well as steady state^19,42,44^. Our findings in *Drosophila* highlight the role of macrophages for the survival on oxidative stress with an urgent need for a balanced immune response to overcome the tissue damage. We further unravel the diverse responses of hemocytes and fat body cells towards oxidative stress which can now serve as a baseline to further explore the spatiotemporal macrophage/hemocyte responses to oxidative stress and their communication with other host tissue cells such as the fat body. Future in depth analysis of the regulation of these cellular states and their coupling to tissue repair will allow to gain new insights for the treatment of diseases where tissue repair and tissue wasting needs to be balanced, including cancer or neurodegenerative diseases.

## Supporting information

Graphical Abstract

## Acknowledgements and Funding

The authors are very grateful to the members of the Kierdorf lab for their help and support. The authors would especially like to thank M. Oberle for her exceptional help with the maintenance of the Drosophila stocks and the management of the fly room. We would like to acknowledge the Lighthouse Core Facility and its staff, especially J. Bodinek-Wersing and U. Jagadeshwaran, for their assistance with the sorting of nuclei. The authors are especially grateful to D. Ganser and H. Fischer for the continuous support throughout the study. Graphical abstract was created with BioRender.com. KK was supported by project grants of the Fritz Thyssen Foundation and of the German Research Foundation (DFG). MSD was supported by Wellcome Trust (207467/Z/17/Z) and Biotechnology and Biological Sciences Research Council (BBSRC) (BB/W/001004/1). KK, MP, and OG were supported by the DFG through project grants within SFB/TRR167 (Project ID 259373024) and by project grants within CRC1479 (Project ID 441891347). KK, MP, OG, and AC were supported by the DFG under Germany’s Excellence Strategy (grant no. CIBSS—EXC-2189, Project ID 390939984). AC was further supported by the Boehringer Ingelheim Foundation (Plus3 Programme). OG was further supported by the European Research Council (ERC) through Starting Grant 337689 and Proof-of-Concept Grant 966687, and by the DFG through project grants in GRK2606 (Project ID 423813989), SFB1425 (Project ID 422681845), and SFB1160 (Project ID 256073931), as well as by the EU-H2020-MSCA-COFUND EURIdoc programme (No. 101034170). MP was further supported by the Novo Nordisk Foundation, the Ernst Jung Foundation, the DFG through project grants in SFB992 (Project ID 192904750), and SFB1160 (Project ID 256073931), a Reinhart Koselleck grant and the Gottfried Wilhelm Leibniz prize, the Ministry of Science, Research and Arts of the state of Baden-Wuerttemberg (Sonderlinie ‘Neuroinflammation’) and by the Alzheimer Forschung Initiative e.V. (AFI). GM was supported by a Marie Skłodowska-Curie postdoctoral fellowship.

## Author Contributions

FH contributed to project planning, performed all experiments, analyzed data, edited the manuscript and prepared the figures. TM and PW performed experiments, helped with the data analysis, and edited the manuscript. CC performed the snRNA-seq experiment and bioinformatics analysis of the hemocytes. GM performed data preprocessing and supported bioinformatics analysis. AC, MSD, KP, OG, and MP provided equipment and reagents, helped to analyze and interpret data, and edited the manuscript. KK planned and supervised the project, conceptualized the study and wrote the manuscript.

## Methods Section

### Drosophila melanogaster stocks

Flies were reared on a high yeast food containing 10% brewer’s yeast, 8% fructose, 2% polenta and 0.8% Agar. Propionic acid and nipagin were added to prevent bacterial or fungal growth. All crosses (except those containing the *tub-Gal80^ts^* construct or otherwise noted) were performed at 25°C with a 12 hours dark/light cycle. The crosses with *tub-Gal80^ts^*were performed at 18°C to ensure the inhibition of the Gal4-protein during developmental stages of the fly. The experimental male F1 flies were transferred to 29°C as soon as they hatched and were aged for six days. All transgenic lines used in this study are listed in the Table S3.

### Lifespan/Survival assays

Male flies were collected after eclosion and groups of 20 age-matched flies per genotype were housed in a food vial. The survival experiments were performed at 29°C. The vials were placed horizontal to avoid that flies fall into the food and become stuck. Dead flies were counted in a daily manner. The flies were transferred into a fresh food vial twice per week without CO_2_ anesthesia.

### Paraquat treatment

Flies were maintained at 29°C for six days prior to treatment. On day six they were starved for 6 hours. A filter paper soaked in 5% sucrose solution with or without 15mM Paraquat (PQ, Methyl viologen hydrate, Acros Organics) was added after starvation. Flies were fed for 18 hours on the PQ containing food in groups of ten. The PQ treatment was performed at 29°C in the dark^20^. To assess the survival rate the dead flies per vial were counted and the percentage was calculated. The living flies were further analyzed in other assays used in this study.

### Starvation experiments

10-20 age-matched male flies were kept in a vial containing 1% agar supplemented with 2% 1xPBS. The starvation experiments were performed at 25°C. Dead flies were counted every hour. Several individual experiments were performed and started on different daytimes to exclude diurnal derived artifacts. The data of the individual experiments were pooled and analyzed with GraphPad Prism 9.2.0.

### Paraffin sections and stainings

Anaesthetized flies were washed in 75% ethanol for 5-10 seconds and transferred into 4% PFA for 30 minutes at room temperature. The flies were washed for 1 hour in PBS on a shaker. Subsequently the flies were transferred into tissue cassettes, dehydrated and embedded in paraffin. The flies were cut in 7µm thick sections de-paraffinated and stained on slides with Hematoxylin-Gill (II) for five minutes. Subsequently the sections were blued in running water for ten minutes. It followed a 0.5% Eosin treatment for five minutes and a final ethanol treatment with increasing concentrations until 100%. The sections were transferred into Xylol and finally covered with synthetic resin and a cover slip.

### Cryo sections and Oil Red O staining

Anaesthetized flies were washed in 75% ethanol for 5-10 seconds and transferred into 4% PFA for 30 minutes at room temperature. Subsequently, the flies were dehydrated for 1h at 37°C in 30% sucrose solution. The dehydrated flies were placed in a Tissue-Tek cryo-mold and embedded in Tissue-Tek® O.C.T.™ Compound. The cryo-mold was placed on dry-ice which was embedded in a 100% ethanol bath until the whole block was completely frozen. The flies were cut in 10µm sections and stained on the slide. The slides were put on RT and dried. Subsequently, they were submerged in Oil red O solution for 30 minutes. The slides were rinsed two times in deionized water and subsequently stained for two minutes with hematoxylin to stain the nuclei. Finally, the slides were washed with water for 5 minutes and covered with a cover slip and glycerin.

### Semi-quantitative Real-Time PCR

Three flies were pooled and smashed in 100µl TRIzol to stabilize and isolate the RNA. Chloroform was used to extract the RNA followed by an isopropanol precipitation step. The RNA was cleaned with 70% ethanol and solubilized in water. A DNAse treatment was performed to digest potential genomic DNA contaminations. The purified and DNAse treated RNA was written into cDNA using RevertAid Reverse Transcriptase (Thermo Scientific) at 37°C for one hour. The reaction was stopped by incubating for 10 minutes at 70°C. The subsequent RTqPCR was done in SensiMix SYBR Green no-ROX (Meridian Bioscience) and was performed on a LightCycler 480 (Roche). The qPCR cycling started with a 10-minute 95°C step followed by 50 cycles with the following times and temperatures: 15s at 95°C, 15s at 60°C and 15s at 72°C. The gene expression levels were normalized to the value of the measured loading control gene Rpl1.

### Confocal microscopy

The flies were anesthetized with CO2, glued on a cover slip and imaged immediately. Confocal microscopy was done with a Leica SP8 microscope (Leica) and a 10×/NA0.4 Leica air objective. The images were acquired in a resolution of 512×512 with a scan speed of 600 Hz. Z-Stacks with a step size of 5µm and tile scans (2×2 or 2×3 images per fly) were acquired to quantify the hemocytes in whole flies. The tiled images were merged with the LAS-X software (Leica) and maximum projections were created using Fiji/ImageJ. The hemocytes were counted in the maximum projection images of whole flies.

### Smurf Assay

Smurf assays were performed to check the feeding behavior as well as the gut integrity of 7 days old, male flies upon PQ treatment. Brilliant Blue dye (FCF) was added into 5% sucrose solution in a concentration of 1% (w/v). PQ was added in concentrations of 2mM, 15mM and 30mM. Control food was 5% sucrose and 1% Brilliant Blue without PQ. The flies were fed with blue food according to the PQ-treatment protocol, as described above. Flies were analyzed after 18h of feeding on brilliant-blue food. Three kinds of flies were distinguished. Flies that did not eat at all were excluded. Flies with a blue gut or crop were classified as “non-smurf”. Flies which turned completely blue or showed distribution of blue dye outside the gut were classified as “smurf”.

### Thin Layer Chromatography (TLC)

Each sample in the TLC analysis contains 10 flies of the respective genotype. The flies were starved for one hour before they were anesthetized with CO2 and transferred into 100µl chloroform:methanol (3:1) on ice. The flies were centrifuged for 5 minutes at 15,000g at 4°C and subsequently homogenized with a pestle. The sample was centrifuged again with the same settings. Lard was dissolved in chloroform:methanol (3:1) and served as triglyceride control (standard 1). Standard 1 was used to produce a standard curve with decreasing triglyceride concentrations. The samples and the standard curve were loaded on a glass silica gel plate (Millipore). The mobile phase in the TLC chamber was made with hexane:diethylether (4:1). The plate was placed in the chamber until the solvent front reached the upper end of the plate. The plate was taken out, air dried and stained with ceric ammonium heptamolybdate (CAM). Subsequently the plate was incubated at 80°C for 2 hours to visualize the stained triglyceride bands. Images were taken with a gel documentation system (gelONE). Triglyceride density was measured and the concentrations were calculated according the pre-determined concentrations of the standard curve. Image analysis and calculations were done using ImageJ.

### Glucose Assay

Seven days old male flies were used for the analysis. They were treated for 18h with PQ or sucrose respectively. Subsequently the flies were starved for one hour to bring all flies into the same metabolic state. Three flies were pooled as one sample and manually homogenized in 75µl TE + 0.1% Triton X-100 (Sigma Aldrich). The samples were incubated for 20min at 75°C to inactivate any intrinsic enzymatic activity. 5µl of the samples were loaded into a flat-bottom 96-well plate. Each sample was measured four times to measure glucose, trehalose, glycogen and the fly background respectively. The fly background was determined by diluting the fly sample with water. Glucose reagent (Sentinel Diagnostics) was added to the fly sample to measure the glucose levels. Trehalase (Sigma Aldrich) or amyloglucosidase (Sigma Aldrich) was added to the glucose reagent to measure trehalose or glycogen respectively. Plates were incubated for 24h at 37°C before the colorimetric measurement was performed at a wavelength of 492nm on a Tecan® Spark plate reader. The concentrations were calculated according to standards loaded on the same plate.

### Nuclei FACS sorting

100 male *Hml-dsRed.nuc* flies per sample were homogenized in 500µl ice-cold EZ nuclei lysis buffer (Sigma). Another 500µl EZ lysis buffer was added and the nuclei were incubated for five minutes on ice. Following a pre-filtration step with a 70µm cell strainer (Miltenyi filters) the nuclei were centrifuged at 500g for six minutes at 4°C. The pellet was resuspended and incubated in 1ml of EZ lysis buffer for five minutes on ice following a second centrifugation step with previous used settings. After discarding the supernatant, 500µl of nuclei wash and resuspension buffer (1x PBS; 1% BSA; 0.2U/µl RNase inhibitor, NEB) was added without resuspending the pellet and incubated for 5 minutes on ice. Subsequently the pellet was resuspended with additional 500µl nuclei wash and resuspension buffer. The nuclei were centrifuged using previous settings and subsequently washed with 1ml of nuclei wash and resuspension buffer. After centrifugation the pellet was resuspended and incubated for 15 minutes with nuclei staining solution containing DAPI (1:1000) and DRAQ7 (1:100). The nuclei were washed once with 1ml nuclei wash and resuspension buffer. After centrifugation the pellet was dissolved in 200µl wash and resuspension buffer, filtered through a 40µm filter and sorted on a BD FACS ARIAIII. A duplet exclusion was performed by gating on singlets in FSC-A vs. FSC-W plot. Subsequently DR+DAPI+ events were sorted into nuclei wash and resuspension buffer.

### Single nucleus library preparation and sequencing

For single nucleus library preparation, nuclei were loaded onto a Chromium Single Cell 3′ G Chip (10X Genomics) to generate single-nucleus gel beads in emulsion (GEM) according to the manufacturer’s instructions without modifications using Chromium Next GEM Single Cell 3’ Reagents Kit v3.1 (10X Genomics). In order to multiplex the samples for sequencing, Single Index Kit T Set A (10x Genomics) was used for library preparation according to the manufacturer’s instructions. The cDNA content and size of post-sample index PCR samples was analyzed using a 2100 BioAnalyzer (Agilent). Library quantification was done using NEB Next® Library Quant Kit for Illumina® (New England Biolabs) following manufacturer’s instructions. Sequencing libraries were loaded on an Illumina Nextseq 550 flow cell, with sequencing settings according to the recommendations of 10× Genomics. Sample de-multiplexing was done using built-in BCL2FASTQ. Cell Ranger v6 software was implemented for gene alignment to the fly genome. The *Drosophila melanogaster* genome assembly and annotation file were downloaded from Ensembl (Release 104). The annotation files were filtered as suggested by 10x Genomics, to keep only the categories of interest (i.e. protein coding genes, long intergenic noncoding RNAs, antisense, pseudogenes). Cell Ranger v6 with the “include-intron” parameter was used for the generation of the count data.

### snRNA-seq data analysis

Downstream analysis implemented the SoupX v1 and Seurat v4 R-based packages. The matrix for each sample was loaded using the SoupX package^45^ and ambient RNA contamination was calculated using the default setting. The corrected matrix together with metadata on treatment were used to create an object in Seurat v4^23^. Doublet detection was done using the DoubletFinder package v2^46^. After doublet exclusion, a combined object was created from the list by using the “merge” function. Nuclei with < 5% of mitochondrial contamination and between 400 and 1,700 genes expressed were retained for further analysis. The filtered raw count matrix was log-normalized within each nucleus. The top 3,000 variable genes were calculated by Seurat using the variance stabilizing transformation selection method and data were scaled. The variable genes were used to perform principal component analysis (PCA) and the top 20 principal components were used for the unsupervised clustering. Seurat applies a graph-based clustering approach: the *“*FindNeighbors*”* function uses the number of principal components (dims = 20) to construct the k-nearest neighbors graph based on the Euclidean distance in PCA space, and the *“*FindClusters” function applies the Louvain algorithm to iteratively group nuclei (resolution = 1). We then used the PCA embedding to generate an UMAP for cluster visualization. For each cluster, differentially expressed genes (DEGs) were calculated using the *“*FindMarkers” and *“*FindAllMarkers” function with default parameters with adjusted P < 0.05 using Bonferroni correction. The DEGs were used to assign cell type identity to clusters based on known cell lineage markers described in previously published study^25^. To understand the impact of Paraquat treatment on each cluster, *“*FindMarkers” function was used to compare “control” vs “Paraquat” treated cells per cluster. Based on the DEGs obtained from this analysis, volcano plots were created using the EnhancedVolcano package v1^47^. UMAP visualizations, feature plots and dot plots for snRNA-seq data were generated using in-built plotting functionality of Seurat.

### Subclustering of hemocytes and further analysis

Based on the cell identity and known hemocyte marker genes from a previous study^25^, sub-clusterings were performed using the same unsupervised algorithm and downstream analysis. After removing non-hemocyte clusters, final sub-clustering was done with a modification that *“*FindNeighbors*”* function was run with dims = 10 and “FindClusters” function was run with resolution 0.8. Based on the DEGs found by the *“*FindAllMarkers” function (parameters: only.pos = T, test.use= “MAST”) a heat map was plotted for TOP 20 DE genes per clusters.

### Subclustering of fat body cells and further analysis

Based on the cell identity according to the FlyCell Atlas, sub-clustering was performed using the same unsupervised algorithm and downstream analysis as used for hemocytes. After removing non-fat body clusters, final sub-clustering was done with the *“*FindNeighbors*”* function which was run with dims = 10 and “FindClusters” function which was run with resolution 0.8. Based on the DEGs found by the *“*FindAllMarkers” function (parameters: only.pos = T, test.use= “MAST”) a heat map was plotted for TOP 20 DE genes per clusters.

### Regulation network analysis of hemocyte clusters

In order to determine the transcription factor activity, regulon analysis was performed using the R package SCENIC (v1.3.1)^48^. To remove noise, genes with low expression levels or low positive rates were filtered using the “geneFiltering” function with default settings to obtain a filtered matrix which was used to build co-expression network using the “runCorrelation” and “runGENIE3” functions. Gene regulatory networks (GRNs) were built and scored using the default parameters. Potential regulons based on DNA-motif analysis were selected by using RcisTarget and active gene networks were identified by AUCell. Regulon activity for each cell was calculated as the average normalized expression of putative target genes. Regulon activity matrix was exported which was used to create a new ‘AUC’ assay using “CreateAssayObject” function of Seurat. “DoHeatmap” function from Seurat was used to plot scaled TF activity in individual hemocytes.

### S2 Cell Culture, CellROX staining and flow cytometry

Drosophila S2 cells (Invitrogen™) were cultured in Schneider’s Drosophila medium (Gibco™) at 26°C and atmospheric oxygen and carbon dioxide conditions. The medium contained 10% FCS and 1% penicillin/streptomycin. PQ was added into the medium in concentrations of 15mM or 30mM to investigate the influence on the production of reactive oxygen species (ROS). Control cells remained untreated. To visualize ROS and oxidative stress, the cells were stained with CellROX™ Deep Red Reagent (Invitrogen) for 30 minutes directly in the medium in a concentration of 1:500 at 26°C. The cells were washed twice with 1xPBS containing 2mM EDTA. In order to stain dead cells DAPI was added to the samples in a concentration of 1:1000. Samples were acquired using a BD LSRFortessa™ (BD Biosciences) and analyzed with FlowJo analysis software.

### COMET assays

Comet assays were performed to analyze the amount of DNA damage in S2 cells treated with PQ. S2 cells were treated with 15mM or 30mM PQ for 24h. The cells were brought into a concentration of 1.8×10^5^ cells/ml. Low melting agarose (LMA) was boiled at 90°C and subsequently cooled down to 37°C. The cells were diluted in low melting agarose (LMA) at a ratio of 1:10 (cells:LMA) and transferred onto a CometSlide™ (Trevigen). To ensure proper attachment of the LMA, the slides were cooled for 30 minutes at 4°C. The lysis was performed for one hour at 4°C. Therefore, the slides were submerged in CometAssay® Lysis Solution (Trevigen). Subsequently the cells were submerged in alkaline unwinding solution (200mM NaOH, 1mM EDTA, pH 13) for 1h at 4°C. The electrophoresis was performed under alkaline conditions (pH13) for 30 minutes with a current of 20V (1V/cm). The slides were neutralized with ddH_2_O, dehydrated with 37% ethanol and dried properly at 37°C before they were stained with SYBR™Gold (Invitrogen) for 30 minutes. Comets were imaged with an Olympus BX61 fluorescence microscope. Comet data was analyzed via TriTrek CometScore 2.0.0.38 software.

### Statistical analysis and data handling

Statistical significance of real-time qPCR, Paraquat survival, hemocyte quantification, TLC, Glucose assay, CellROX and comet assay data was calculated with an unpaired t-test or one-way ANOVA as indicated in the figure legends. Lifespan/survival assays were analyzed using Log-Rank and Wilcoxon test. Significance digits indicate the following significance levels: \**p<0.05*, \*\**p<0.01*, \*\*\**p<0.001*, \*\*\*\**p<0.0001*. Significance tests were performed using GraphPad Prism 9 software. Data handling of snRNAseq data is described above.

## Data availability

snRNA-seq data will be available upon publication on the gene omnibus express (GEO) platform, GEO accession number: GSE24429. Please contact the lead contact for the token. Microscopy, flow cytometry and survival data reported in this paper will be shared by the lead contact upon request. Any additional information required to reanalyze the data reported in this paper is available from the lead contact upon request.

## Supplementary Figure Legends

**Figure S1:** Paraquat-induced oxidative stress does not cause gut leakage, but detrimental changes in triglyceride storage in the fat body. (A) Survival of *Hml/+* flies treated with control food (n=8), 2mM PQ (n=8), 15mM PQ (n=9) and 30mM PQ (n=8). Each data point represents a sample of 10 flies. Mean ± SEM is shown. One-way ANOVA: \*\**p<0.01*; \*\*\*\**p<0.0001*. (B) Smurf assay to test the gut integrity of 7 days old *Hml/+* flies treated with 2mM PQ (n=79), 15mM PQ (n=90), 30mM PQ (n=80) and controls (n=80). Data was pooled from two independent experiments. (C) Gene expression analysis for *AMP* genes via RT-qPCR in whole flies on control food (C) and 15mM Paraquat (P). All transcript levels were normalized to the expression of the loading control *Rpl1* and are shown in arbitrary units [au]. Each data point (n=5) represents a sample of three individual flies. Mean ± SEM is shown. (D) Gene expression analysis for *dpp*, *daw* and *eiger* via RT-qPCR in whole flies on control food (C) and 15mM Paraquat (P). All transcript levels were normalized to the expression of the loading control *Rpl1* and are shown in arbitrary units [au]. Each data point (n=5) represents a sample of three individual flies. Mean ± SEM is shown. (E) Representative light microscopic images of HE stained control flies (left panels) and flies treated with 15mM PQ (right panels). Inserts indicate location of higher magnification images below. Scale bar (overview) = 500 µm, scale bar (inserts) = 100 µm. (F) Representative light microscopic images of Oil Red O stained control flies (left panels) and flies treated with 15mM PQ (right panels). Arrows indicate stained fat droplets. Scale bar = 100 µm.

**Figure S2:** Gating strategy for the FACS isolation of nuclei from control and PQ-treated flies. (A) Flow cytometric sorting strategy of nuclei used for the snRNA-seq. Doublet exclusion was performed via gating on DR single positive nuclei. DR^+^DAPI^+^ double positive events were considered to be nuclei and were sorted for the snRNA-seq as previously described^49^. (B) Violin plot indicating number of genes identified in analyzed nuclei of the sorted samples. Color code is indicated in legend. (C) Violin plot indicating RNA counts of the analyzed nuclei of each sorted sample. Color code is indicated in legend. (D) Violin plot indicating percentage of mitochondrial genes detected in the nuclei of each sorted samples. Color code is indicated in legend.

**Figure S3:** Specific signature genes identify plasmatocyte and crystal cell clusters within the eight identified hemocyte clusters. (A) Dot plot of hemocyte signature gene expression according to Cattenoz et al^25^. Color code is indicated in legend. (B)-(L) Feature plots for expression levels of respective hemocyte signature genes across all hemocyte clusters. Color-code, indicating the expression levels of the respective genes, is shown on the right side of each graph. Gene names are indicated above plots shown. (M) Bar plot representing distribution of nuclei derived from flies under control condition (red) and flies treated with 15mM PQ (turquoise) across all eight identified hemocyte clusters. Each cluster is shown as one bar. Total number of hemocytes in the respective cluster is indicated below.

**Figure S4:** Fat body signature genes are highly expressed across the eight distinct fat body cell clusters and are absent in hemocyte cell clusters. (A) Dot plot of fat body signature gene expression according to the Fly Cell Atlas signature^24^ on the eight identified hemocytes clusters. Color code is indicated in legend. (B) Dot plot of fat body signature gene expression on the eight identified fat body clusters. Color code is indicated in the legend. (C) Dot plot of hemocyte signature gene expression according to Cattenoz et al.^25^ on the eight identified fat body clusters. Color code is indicated in the legend. (D) Bar plot representing distribution of nuclei derived from flies under control condition (red) and flies treated with 15mM PQ (turquoise) across all eight identified fat body clusters. Each cluster is shown as one bar. Total number of fat body cells in the respective cluster is indicated below.

**Figure S5:** PQ treatment of S2 cells *in vitro* induces reactive oxygen species, immune activation and DNA damage. (A) Gating strategy of flow cytometric analysis of control or PQ-treated S2-cells stained for reactive oxygen species (ROS) with CellROX. Control cells, 15mM PQ treated and 30mM PQ treated cells are shown in the first, second and third row respectively. After doublet exclusion, dead cells were excluded via DAPI staining. The amount of ROS was determined in the last gate were the CellROX staining is shown. (B) Representative FACS histograms of cells treated with PQ and analyzed via CellROX staining. Gates indicate the cut-off for determining CellROX^+^ cells. (C) Mean fluorescence intensity (MFI) of FACS analyzed, CellROX^+^ cells. Data were generated from three independent experiments (n=3). One-way ANOVA: *\*p<0.05; **p<0.01; ***p<0.001; ****p<0.0001*. (D) Comet assay of S2 cells treated without PQ (n=351), 5mM PQ (n=208), 10mM PQ (n=129) and 15mM PQ (n=71). Assay was performed in two independent experiments. Data of one representative assay are shown as violin plots with median (red), second and third quartile. One-way ANOVA: *\*\*\*p<0.001; ****p>0.0001*. (E-F) Gene expression analysis of genes associated with JAK-STAT signaling, JNK signaling, insulin signaling is shown in (E) and TGFβ signaling and AMPs is shown in (F). S2-cells treated with 15mM PQ for six or 24 hours. Controls were untreated. Each dot represents a sample containing RNA of 50.000 cells. One-way ANOVA: *\*p<0.05; **p<0.01; ***p<0.001; ****p<0.0001*.

**Figure S6:** Loss of DNA damage, upd3 or JNK signaling in hemocytes alters susceptibility to oxidative stress but not overall lifespan. (A) Representative images of *Hml/+* flies and DDR-Knockdown flies on control food (left panels) and treated with 15mM PQ (right panels). Five individual flies per group were analyzed. (B) Lifespan analysis of *Hml/+* (n=251)*, Hml>mei-41-IR* (n=231)*, Hml>tefu-IR* (n=219)*, Hml>mei-41-IR;tefu-IR* (n=144) and *Hml>nbs-IR* flies (n=299). *Hml/+* vs. *Hml>mei-41-IR*: Log-rank test \*\*\*\**p<0.0001*, Χ^2^=172.2; Wilcoxon test \*\*\*\**p<0.0001*, Χ^2^=155.9. *Hml/+* vs. *Hml>tefu-IR*: Log-rank test \*\*\*\**p<0.0001*, Χ^2^=56.09; Wilcoxon test \*\*\*\**p<0.0001*, Χ^2^=50.95. *Hml/+* vs. *Hml>mei-41-IR;tefu-IR*: Log-rank test ns, Χ^2^=3.1; Wilcoxon test \*\*\**p=0.0005*, Χ^2^=12.11. *Hml/+* vs *Hml>nbs-IR*: Log-rank test \*\*\*\**p<0.0001*, Χ^2^=60.90; Wilcoxon test \*\*\*\**p<0.0001*, Χ^2^=50.21. Data was obtained from three independent experiments. (C) Starvation survival of *Hml/+* (n=607)*, Hml>mei-41-IR* (n=421)*, Hml>tefu-IR* (n=480)*, Hml>mei-41-IR;tefu-IR* (n=266) and *Hml>nbs-IR* (n=341) flies. *Hml/+* vs. *Hml>mei-41-IR*: Log-rank test \*\**p=0.0052*, Χ^2^=7.81; Wilcoxon test ns, Χ^2^=0.02. *Hml/+* vs. *Hml>tefu-IR*: Log-rank test ns, Χ^2^=0.03; Wilcoxon test \**p=0.0153*, Χ^2^=5.88. *Hml/+* vs. *Hml>mei-41-IR;tefu-IR*: Log-rank test ns, Χ^2^=0.21; Wilcoxon test \**p=0.019*, Χ^2^=5.51. *Hml/+* vs *Hml>nbs-IR*: Log-rank test \*\*\**p=0.0003*, Χ^2^=13.30; Wilcoxon test ns, Χ^2^=1.24. Data was obtained from two independent experiments. (D) Lifespan analysis of *Hml-Gal80^ts^/+* (n=127)*, Hml-Gal80^ts^>mei-41-IR* (n=140)*, Hml-Gal80^ts^>tefu-IR* (n=164)*, Hml-Gal80^ts^>mei-41-IR;tefu-IR* (n=101) and *Hml-Gal80^ts^>nbs-IR* flies (n=157). *Hml-Gal80^ts^/+* vs. *Hml-Gal80^ts^>mei-41-IR*: Log-rank test \*\*\*\**p<0.0001*, chi^2^=21.80; Wilcoxon test \*\*\*\**p<0.0001*, Χ^2^=18.48. *Hml-Gal80^ts^/+* vs. *Hml-Gal80^ts^>tefu-IR*: Log-rank test \*\*\**p=0.0003*, Χ^2^=13.20; Wilcoxon test \**p=0.033*, Χ^2^=4.56. *Hml-Gal80^ts^/+* vs. *Hml-Gal80^ts^>mei-41-IR;tefu-IR*: Log-rank test \**p=0.0429*, Χ^2^=4.10; Wilcoxon test \*\*\**p=0.0001*, Χ^2^=14.95. *Hml-Gal80^ts^/+* vs *Hml-Gal80^ts^>nbs-IR*: Log-rank test \*\**p=0.0047*, Χ^2^=7.99; Wilcoxon test ns, Χ^2^=1.68. Data was obtained from five independent experiments. (E) Survival of *Hml-Gal80^ts^/+* (n=23)*, Hml-Gal80^ts^>mei-41-IR* (n=28)*, Hml-Gal80^ts^>tefu-IR* (n=18)*, Hml-Gal80^ts^>mei-41-IR;tefu-IR* (n=13) and *Hml-Gal80^ts^>nbs-IR* flies (n=31) on PQ food. Each dot represents a sample with ten flies. Mean ± SEM is shown. One-way ANOVA \**p<0.05*; \*\*\*\**p<0.0001*. Data was obtained from five independent experiments. (F) Representative images of *Hml/+, Hml>upd3-IR* and *Hml>upd3* flies on control food (left panels) and treated with 15mM PQ (right panels). Six individual flies per group were analyzed. (G) Lifespan analysis of *Hml/+ (n=266), Hml>upd3-IR (n=269),* and *Hml>upd3* flies (n=193). *Hml/+* vs. *Hml>upd3-IR*: Log-rank test \*\*\*\**p<0.0001*, Χ^2^=96.03; Wilcoxon test \*\*\*\**p<0.0001*, Χ^2^=81.16. *Hml/+* vs. *Hml>upd3*: Log-rank test \*\*\*\**p<0.0001*, Χ^2^=15.33; Wilcoxon test \**p=0.0124*, Χ^2^=6.26. Results are shown from two independent experiments. (H) Lifespan analysis of *Hml/+* (n=281), *Hml>hep[act]* (n=117), and *Hml>bsk-DN* (n=273) flies. *Hml/+* vs. *Hml>hep[act]*; Log-rank test: n.s.; p=0.69, Χ^2^=0.16; Wilcoxon test: n.s. p=0.18, Χ^2^=1.77. *Hml/+* vs. *Hml>bsk-DN*: Log-rank test: *\*\*\*\*p<0.0001*, Χ^2^=25.31; Wilcoxon test: *\*\*\*\*p<0.0001*, Χ^2^=23.61. Data was obtained from two independent experiments.

**Table S1:** Differentially expressed genes in hemocyte clusters C1-C8.

**Table S2:** Differentially expressed genes in fat body cell clusters C1-C8.

**Table S3:** Transgenic *Drosophila* lines and primers used in the study.

## References

1. Pham-Huy, L.A., He, H., and Pham-Huy, C. (2008). Free Radicals, Antioxidants in Disease and Health. Int J Biomed Sci 4, 89–96.

2. Sinenko, S.A., Shim, J., and Banerjee, U. (2011). Oxidative stress in the haematopoietic niche regulates the cellular immune response in Drosophila. EMBO Rep 13, 83–89. 10.1038/embor.2011.223.

3. Herb, M., and Schramm, M. (2021). Functions of ROS in Macrophages and Antimicrobial Immunity. Antioxidants (Basel) 10, 313. 10.3390/antiox10020313.

4. Nathan, C., and Cunningham-Bussel, A. (2013). Beyond oxidative stress: an immunologist’s guide to reactive oxygen species. Nat Rev Immunol 13, 349–361. 10.1038/nri3423.

5. Finkel, T., and Holbrook, N.J. (2000). Oxidants, oxidative stress and the biology of ageing. Nature 408, 239–247. 10.1038/35041687.

6. Moghadam, Z.M., Henneke, P., and Kolter, J. (2021). From Flies to Men: ROS and the NADPH Oxidase in Phagocytes. Frontiers in Cell and Developmental Biology 9, 618. 10.3389/fcell.2021.628991.

7. Meister, M., and Lagueux, M. (2003). Drosophila blood cells. Cell Microbiol 5, 573–580. 10.1046/j.1462-5822.2003.00302.x.

8. Banerjee, U., Girard, J.R., Goins, L.M., and Spratford, C.M. (2019). Drosophila as a Genetic Model for Hematopoiesis. Genetics 211, 367–417. 10.1534/genetics.118.300223.

9. Gold, K.S., and Brückner, K. (2014). Drosophila as a model for the two myeloid blood cell systems in vertebrates. Exp. Hematol. 42, 717–727. 10.1016/j.exphem.2014.06.002.

10. Holz, A., Bossinger, B., Strasser, T., Janning, W., and Klapper, R. (2003). The two origins of hemocytes in Drosophila. Development 130, 4955–4962. 10.1242/dev.00702.

11. Péan, C.B., Schiebler, M., Tan, S.W.S., Sharrock, J.A., Kierdorf, K., Brown, K.P., Maserumule, M.C., Menezes, S., Pilátová, M., Bronda, K., et al. (2017). Regulation of phagocyte triglyceride by a STAT-ATG2 pathway controls mycobacterial infection. Nat Commun 8, 14642. 10.1038/ncomms14642.

12. Woodcock, K.J., Kierdorf, K., Pouchelon, C.A., Vivancos, V., Dionne, M.S., and Geissmann, F. (2015). Macrophage-derived upd3 cytokine causes impaired glucose homeostasis and reduced lifespan in Drosophila fed a lipid-rich diet. Immunity 42, 133–144. 10.1016/j.immuni.2014.12.023.

13. Kierdorf, K., Hersperger, F., Sharrock, J., Vincent, C.M., Ustaoglu, P., Dou, J., Gyoergy, A., Groß, O., Siekhaus, D.E., and Dionne, M.S. (2020). Muscle function and homeostasis require cytokine inhibition of AKT activity in Drosophila. Elife 9, e51595. 10.7554/eLife.51595.

14. Ayyaz, A., Li, H., and Jasper, H. (2015). Haemocytes control stem cell activity in the Drosophila intestine. Nat. Cell Biol. 17, 736–748. 10.1038/ncb3174.

15. Myers, A.L., Harris, C.M., Choe, K.-M., and Brennan, C.A. (2018). Inflammatory production of reactive oxygen species by Drosophila hemocytes activates cellular immune defenses. Biochemical and Biophysical Research Communications 505, 726–732. 10.1016/j.bbrc.2018.09.126.

16. Fogarty, C.E., Diwanji, N., Lindblad, J.L., Tare, M., Amcheslavsky, A., Makhijani, K., Brückner, K., Fan, Y., and Bergmann, A. (2016). Extracellular Reactive Oxygen Species Drive Apoptosis-Induced Proliferation via Drosophila Macrophages. Current Biology 26, 575–584. 10.1016/j.cub.2015.12.064.

17. Chakrabarti, S., Dudzic, J.P., Li, X., Collas, E.J., Boquete, J.-P., and Lemaitre, B. (2016). Remote Control of Intestinal Stem Cell Activity by Haemocytes in Drosophila. PLoS Genet 12, e1006089. 10.1371/journal.pgen.1006089.

18. Chakrabarti, S., and Visweswariah, S.S. (2020). Intramacrophage ROS Primes the Innate Immune System via JAK/STAT and Toll Activation. Cell Reports 33, 108368. 10.1016/j.celrep.2020.108368.

19. Wu, S.-C., Liao, C.-W., Pan, R.-L., and Juang, J.-L. (2012). Infection-Induced Intestinal Oxidative Stress Triggers Organ-to-Organ Immunological Communication in Drosophila. Cell Host & Microbe 11, 410–417. 10.1016/j.chom.2012.03.004.

20. Wang, M.C., Bohmann, D., and Jasper, H. (2003). JNK Signaling Confers Tolerance to Oxidative Stress and Extends Lifespan in Drosophila. Developmental Cell 5, 811–816. 10.1016/S1534-5807(03)00323-X.

21. Cox, N., Crozet, L., Holtman, I.R., Loyher, P.-L., Lazarov, T., White, J.B., Mass, E., Stanley, E.R., Elemento, O., Glass, C.K., et al. (2021). Diet-regulated production of PDGFcc by macrophages controls energy storage. Science 373. 10.1126/science.abe9383.

22. Martelotto, L. (2020). ‘Frankenstein’ protocol for nuclei isolation from fresh and frozen tissue for snRNAseq.

23. Hao, Y., Hao, S., Andersen-Nissen, E., Mauck, W.M., Zheng, S., Butler, A., Lee, M.J., Wilk, A.J., Darby, C., Zager, M., et al. (2021). Integrated analysis of multimodal single-cell data. Cell 184, 3573–3587.e29. 10.1016/j.cell.2021.04.048.

24. Li, H., Janssens, J., De Waegeneer, M., Kolluru, S.S., Davie, K., Gardeux, V., Saelens, W., David, F.P.A., Brbić, M., Spanier, K., et al. (2022). Fly Cell Atlas: A single-nucleus transcriptomic atlas of the adult fruit fly. Science 375, eabk2432. 10.1126/science.abk2432.

25. Cattenoz, P.B., Sakr, R., Pavlidaki, A., Delaporte, C., Riba, A., Molina, N., Hariharan, N., Mukherjee, T., and Giangrande, A. (2020). Temporal specificity and heterogeneity of Drosophila immune cells. EMBO J 39, e104486. 10.15252/embj.2020104486.

26. Tattikota, S.G., Cho, B., Liu, Y., Hu, Y., Barrera, V., Steinbaugh, M.J., Yoon, S.-H., Comjean, A., Li, F., Dervis, F., et al. (2020). A single-cell survey of Drosophila blood. eLife 9, e54818. 10.7554/eLife.54818.

27. Morales, A.J., Carrero, J.A., Hung, P.J., Tubbs, A.T., Andrews, J.M., Edelson, B.T., Calderon, B., Innes, C.L., Paules, R.S., Payton, J.E., et al. (2017). A type I IFN-dependent DNA damage response regulates the genetic program and inflammasome activation in macrophages. eLife 6, e24655. 10.7554/eLife.24655.

28. Jiang, H., Patel, P.H., Kohlmaier, A., Grenley, M.O., McEwen, D.G., and Edgar, B.A. (2009). Cytokine/Jak/Stat Signaling Mediates Regeneration and Homeostasis in the Drosophila Midgut. Cell 137, 1343–1355. 10.1016/j.cell.2009.05.014.

29. Pastor-Pareja, J.C., Wu, M., and Xu, T. (2008). An innate immune response of blood cells to tumors and tissue damage in Drosophila. Dis Model Mech 1, 144–154; discussion 153. 10.1242/dmm.000950.

30. Karpac, J., Younger, A., and Jasper, H. (2011). Dynamic coordination of innate immune signaling and Insulin signaling regulates systemic responses to localized DNA damage. Dev Cell 20, 841– 854. 10.1016/j.devcel.2011.05.011.

31. Louradour, I., Sharma, A., Morin-Poulard, I., Letourneau, M., Vincent, A., Crozatier, M., and Vanzo, N. (2017). Reactive oxygen species-dependent Toll/NF-κB activation in the Drosophila hematopoietic niche confers resistance to wasp parasitism. eLife 6, e25496. 10.7554/eLife.25496.

32. Paraquat administration in Drosophila for use in metabolic studies of oxidative stress (2011). Analytical Biochemistry 419, 345–347. 10.1016/j.ab.2011.08.023.

33. Jiang, H., Grenley, M.O., Bravo, M.-J., Blumhagen, R.Z., and Edgar, B.A. (2011). EGFR/Ras/MAPK signaling mediates adult midgut epithelial homeostasis and regeneration in Drosophila. Cell Stem Cell 8, 84–95. 10.1016/j.stem.2010.11.026.

34. Patel, P.H., Pénalva, C., Kardorff, M., Roca, M., Pavlović, B., Thiel, A., Teleman, A.A., and Edgar, B.A. (2019). Damage sensing by a Nox-Ask1-MKK3-p38 signaling pathway mediates regeneration in the adult Drosophila midgut. Nat Commun 10, 4365. 10.1038/s41467-019-12336-w.

35. Azad, P., Ryu, J., and Haddad, G.G. (2011). Distinct role of Hsp70 in Drosophila hemocytes during severe hypoxia. Free Radic Biol Med 51, 530–538. 10.1016/j.freeradbiomed.2011.05.005.

36. Huang, K., Chen, W., Zhu, F., Li, P.W.-L., Kapahi, P., and Bai, H. (2019). RiboTag translatomic profiling of Drosophila oenocytes under aging and induced oxidative stress. BMC Genomics 20, 50. 10.1186/s12864-018-5404-4.

37. Maitra, U., Scaglione, M.N., Chtarbanova, S., and O’Donnell, J.M. (2019). Innate immune responses to paraquat exposure in a Drosophila model of Parkinson’s disease. Sci Rep 9, 12714. 10.1038/s41598-019-48977-6.

38. Donovan, M.R., and Marr, M.T. (2016). dFOXO Activates Large and Small Heat Shock Protein Genes in Response to Oxidative Stress to Maintain Proteostasis in Drosophila*. Journal of Biological Chemistry 291, 19042–19050. 10.1074/jbc.M116.723049.

39. Khan, C., Muliyil, S., and Rao, B.J. (2019). Genome Damage Sensing Leads to Tissue Homeostasis in Drosophila. Int Rev Cell Mol Biol 345, 173–224. 10.1016/bs.ircmb.2018.12.001.

40. Dunphy, G., Flannery, S.M., Almine, J.F., Connolly, D.J., Paulus, C., Jønsson, K.L., Jakobsen, M.R., Nevels, M.M., Bowie, A.G., and Unterholzner, L. (2018). Non-canonical Activation of the DNA Sensing Adaptor STING by ATM and IFI16 Mediates NF-κB Signaling after Nuclear DNA Damage. Mol Cell 71, 745–760.e5. 10.1016/j.molcel.2018.07.034.

41. Bednarski, J.J., and Sleckman, B.P. (2019). At the intersection of DNA damage and immune responses. Nat Rev Immunol 19, 231–242. 10.1038/s41577-019-0135-6.

42. Yang, H., Kronhamn, J., Ekström, J., Korkut, G.G., and Hultmark, D. (2015). JAK/STAT signaling in Drosophila muscles controls the cellular immune response against parasitoid infection. EMBO Rep 16, 1664–1672. 10.15252/embr.201540277.

43. Dionne, M.S., Pham, L.N., Shirasu-Hiza, M., and Schneider, D.S. (2006). Akt and FOXO dysregulation contribute to infection-induced wasting in Drosophila. Curr Biol 16, 1977–1985. 10.1016/j.cub.2006.08.052.

44. Yang, H., and Hultmark, D. (2016). Tissue communication in a systemic immune response of Drosophila. Fly (Austin) 10, 115–122. 10.1080/19336934.2016.1182269.

45. Young, M.D., and Behjati, S. (2020). SoupX removes ambient RNA contamination from droplet-based single-cell RNA sequencing data. Gigascience 9, giaa151. 10.1093/gigascience/giaa151.

46. McGinnis, C.S., Murrow, L.M., and Gartner, Z.J. (2019). DoubletFinder: Doublet Detection in Single-Cell RNA Sequencing Data Using Artificial Nearest Neighbors. Cell Syst 8, 329–337.e4. 10.1016/j.cels.2019.03.003.

47. Blighe, K. (2021). EnhancedVolcano: publication-ready volcano plots with enhanced colouring and labeling.

48. Aibar, S., González-Blas, C.B., Moerman, T., Huynh-Thu, V.A., Imrichova, H., Hulselmans, G., Rambow, F., Marine, J.-C., Geurts, P., Aerts, J., et al. (2017). SCENIC: single-cell regulatory network inference and clustering. Nat Methods 14, 1083–1086. 10.1038/nmeth.4463.

49. Hersperger, F., Kastl, M., Paeschke, K., and Kierdorf, K. (2024). Hemocyte Nuclei Isolation from Adult Drosophila melanogaster for snRNA-seq. Methods Mol Biol 2713, 71–79. 10.1007/978-1-0716-3437-0_4.

