## Supplementary figures and images for "DNA damage signaling in *Drosophila* macrophages modulates systemic cytokine levels in response to oxidative stress"

### Figure S1

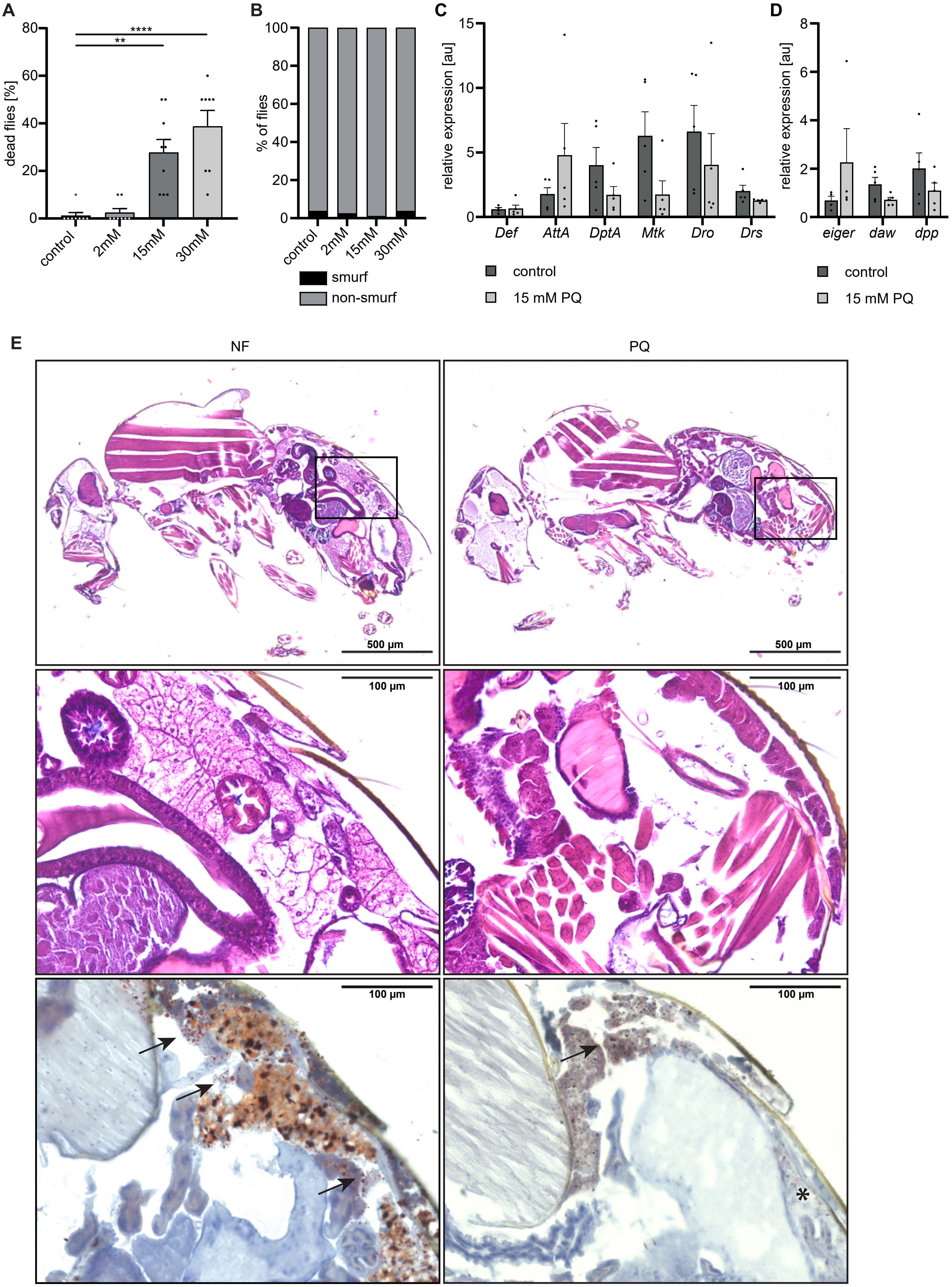

### Figure S2

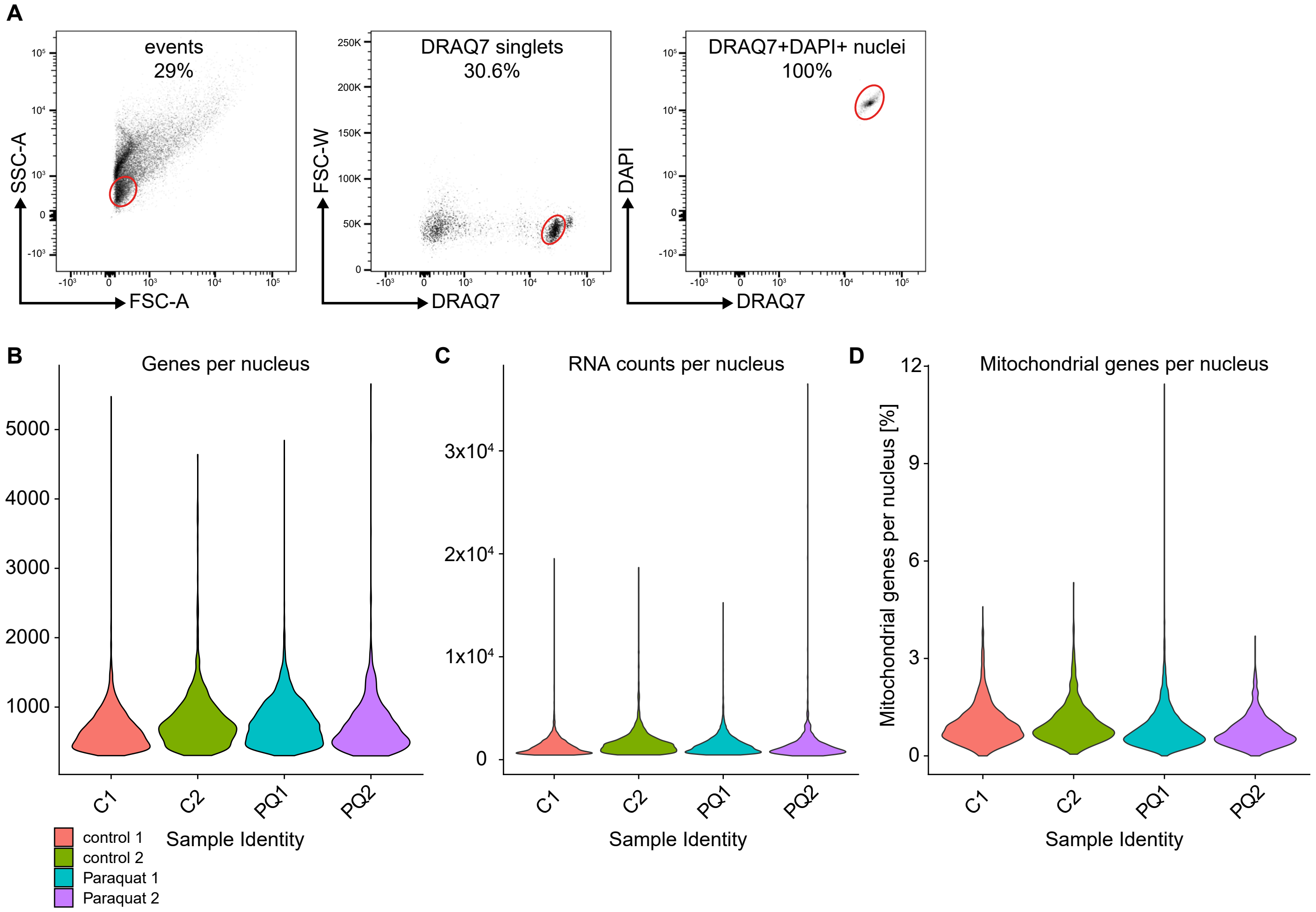

### Figure S3

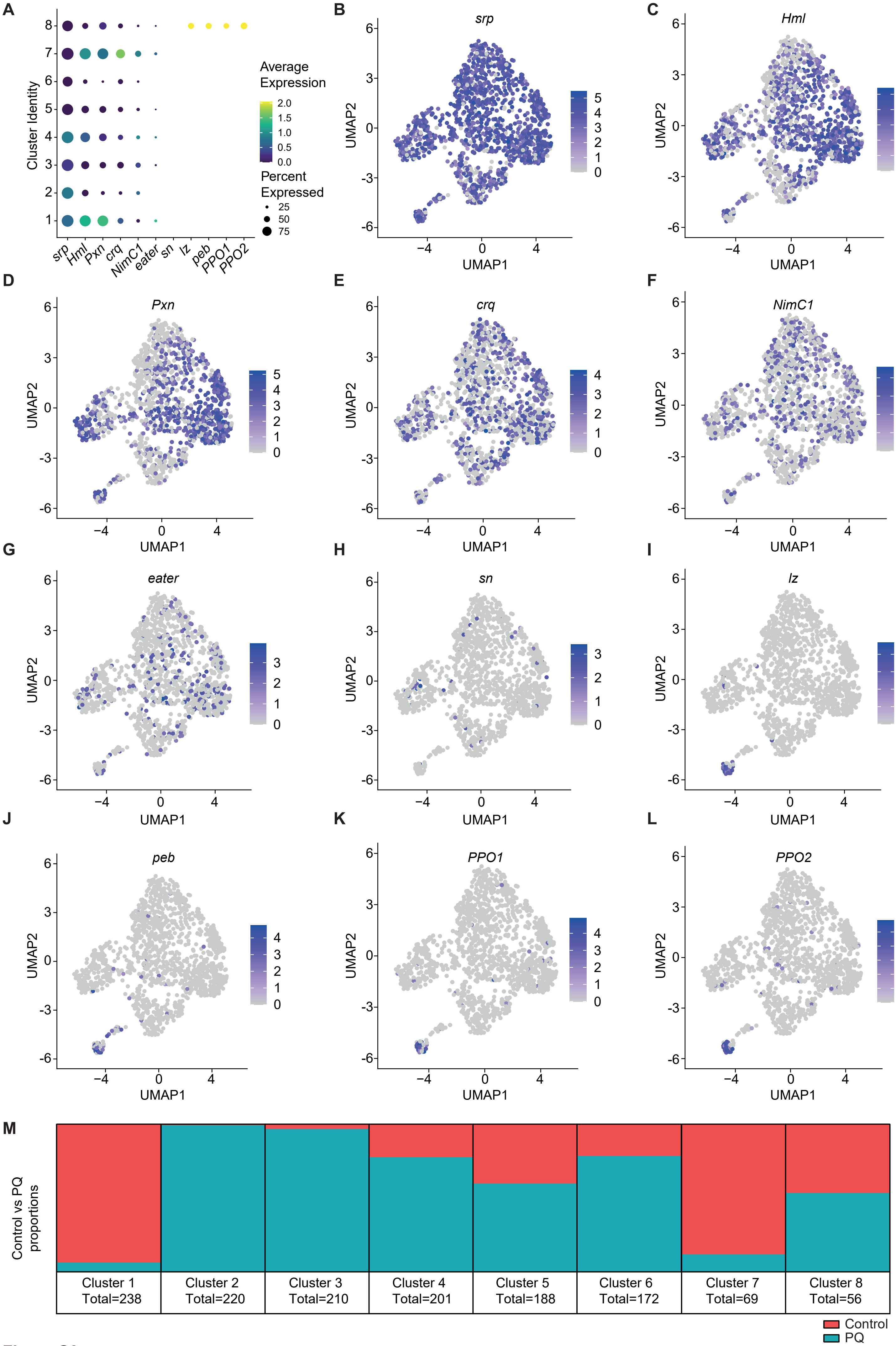

### Figure S4

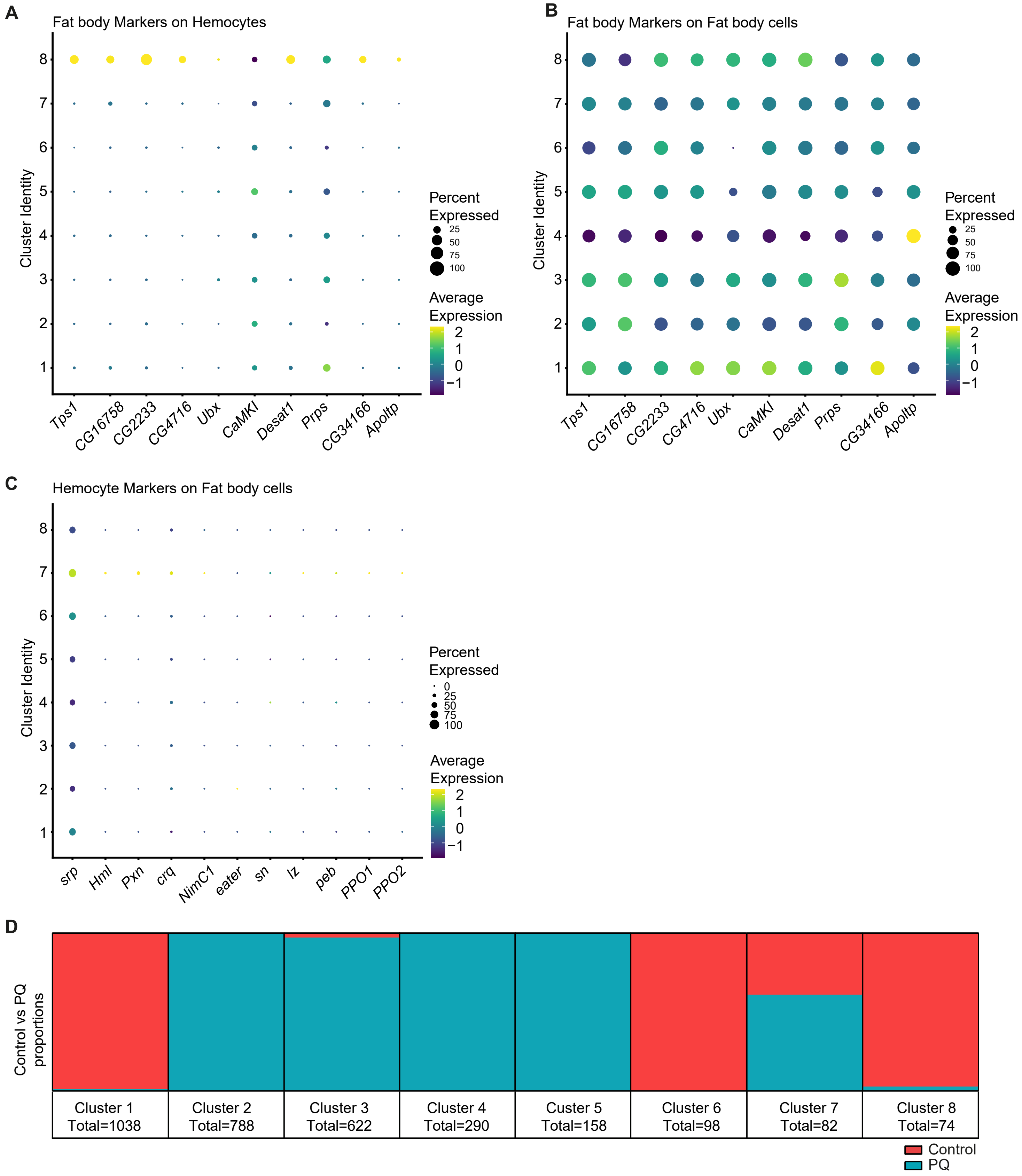

### Figure S5

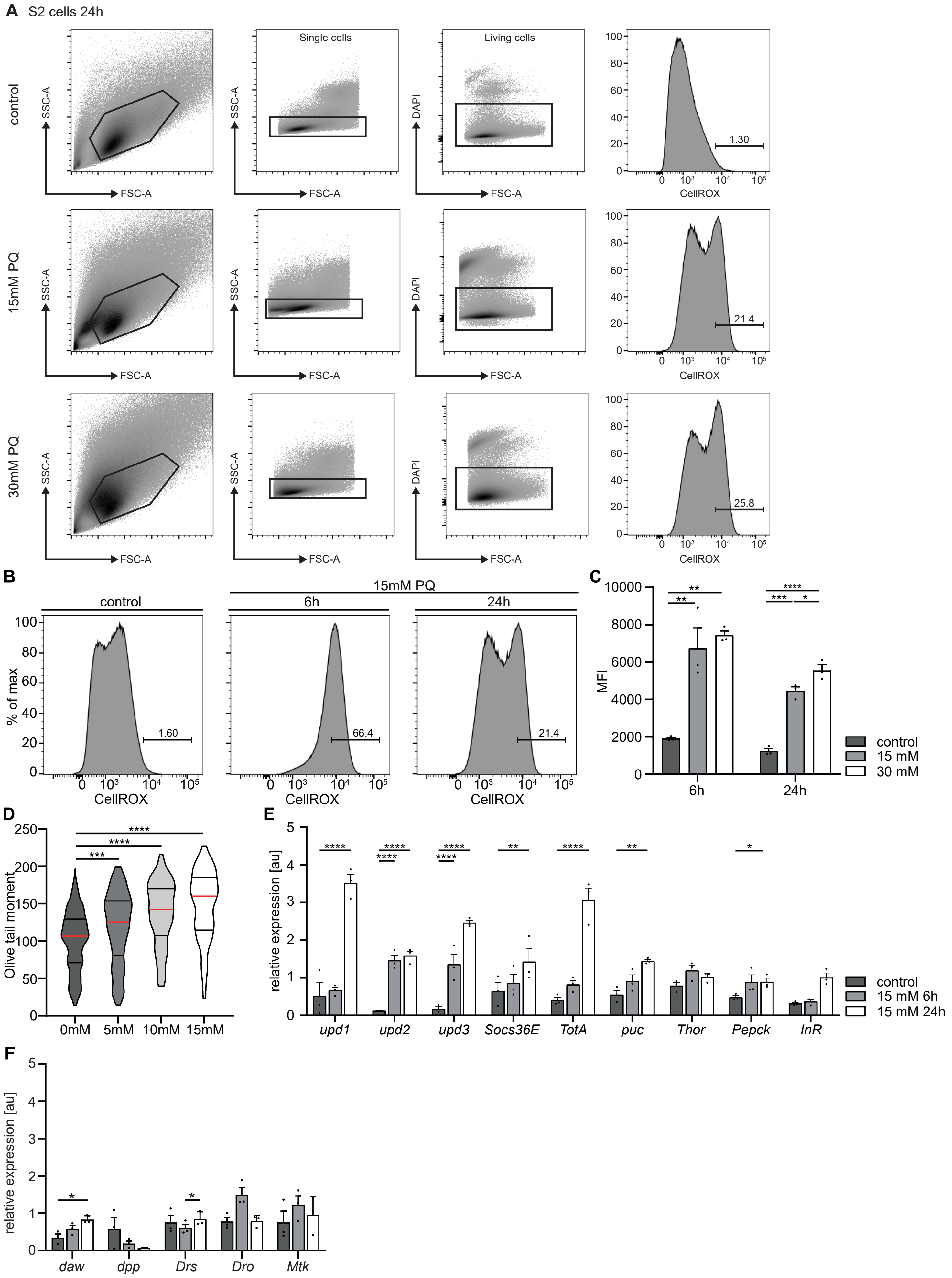

### Figure S6

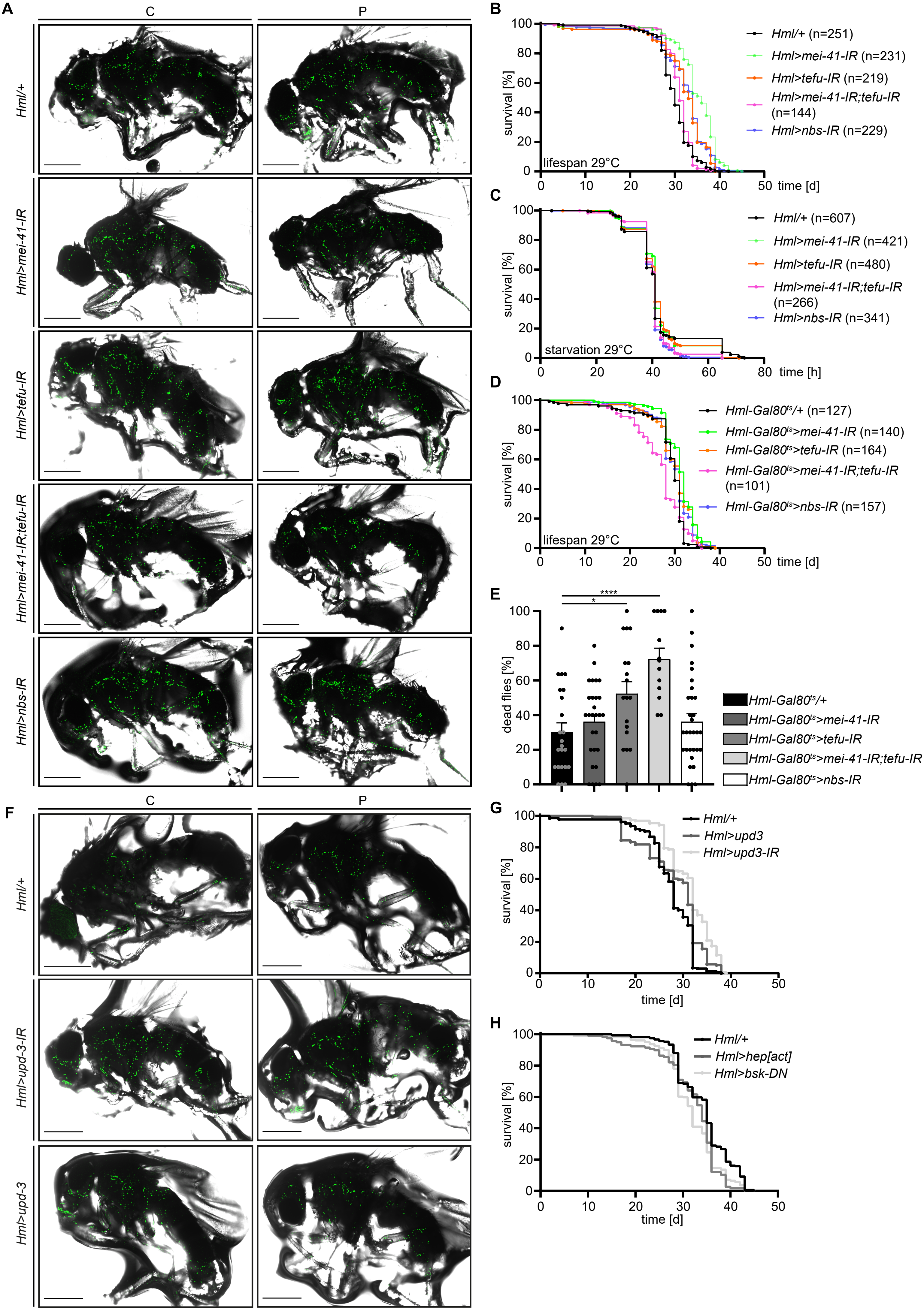

### Graphical Abstract

#### Graphical Abstract

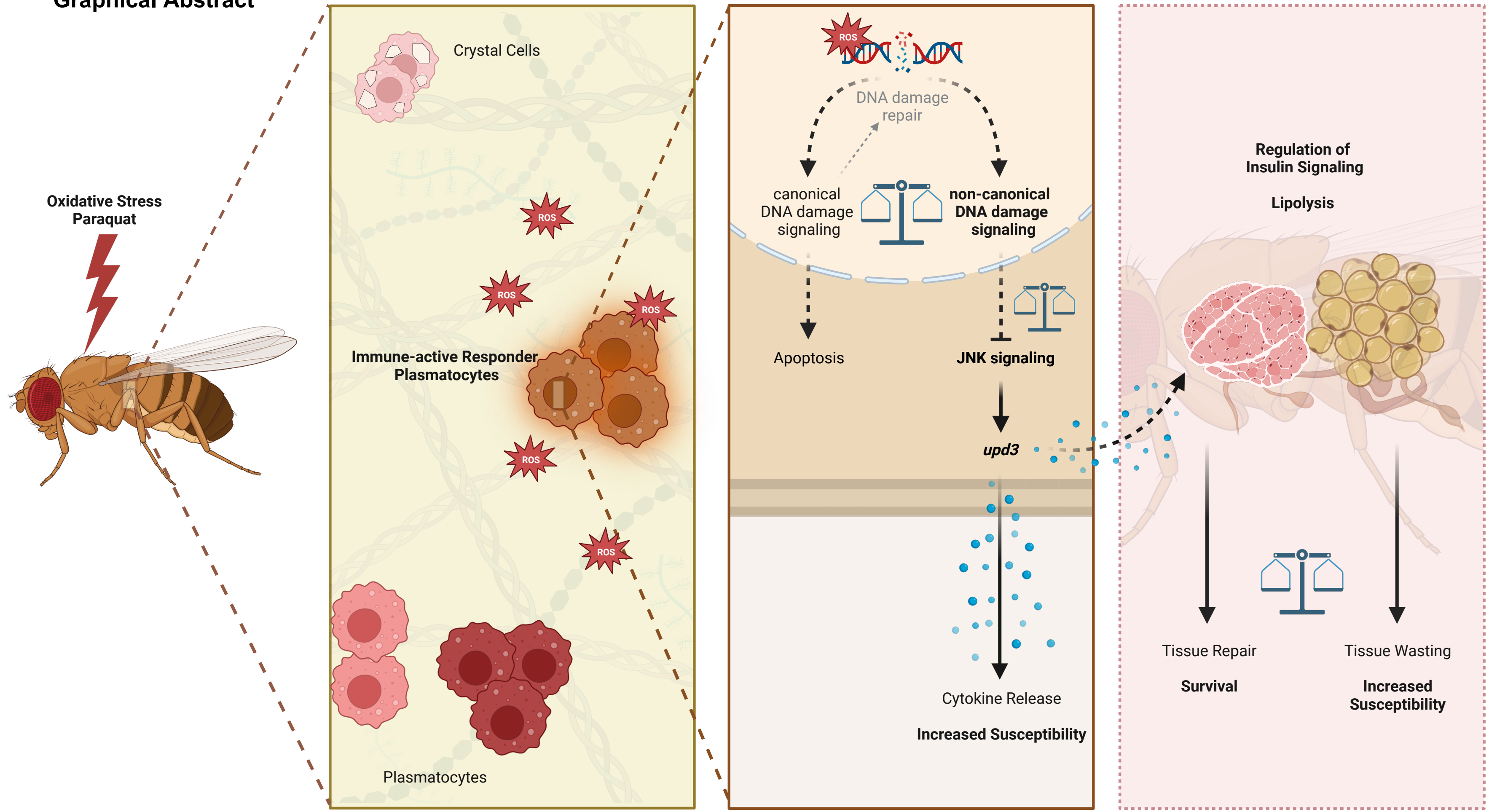
