## Supplementary material for "DNA damage signaling in *Drosophila* macrophages modulates systemic cytokine levels in response to oxidative stress": Table S1

| Gene name | p-value | Average Log2 Fold Chang | pct.1 | pct.2 | adjusted p-value | ter Ide |
|---|---|---|---|---|---|---|
| *Sema2a* | 1,72E-44 | 1,616288928 | 0,807 | 0,371 | 2,06E-40 | 1 |
| *Pxn* | 7,35E-41 | 1,418123886 | 0,861 | 0,475 | 8,77E-37 | 1 |
| *CG5953* | 1,68E-39 | -2,674606281 | 0,126 | 0,567 | 2,00E-35 | 1 |
| *CG31145* | 1,12E-37 | 1,12782434 | 0,929 | 0,706 | 1,34E-33 | 1 |
| *InR* | 1,28E-37 | -1,578699769 | 0,655 | 0,895 | 1,52E-33 | 1 |
| *CG17574* | 1,03E-36 | 1,249187276 | 0,857 | 0,578 | 1,23E-32 | 1 |
| *regucalcin* | 4,30E-33 | 1,279046828 | 0,819 | 0,425 | 5,12E-29 | 1 |
| *Ten-m* | 2,97E-31 | 0,673576668 | 1 | 0,995 | 3,55E-27 | 1 |
| *Hml* | 3,24E-28 | 1,073329431 | 0,891 | 0,6 | 3,87E-24 | 1 |
| *Kr-h1* | 5,16E-27 | -1,857482329 | 0,168 | 0,547 | 6,16E-23 | 1 |
| *Lpin* | 9,76E-27 | -2,359091913 | 0,055 | 0,363 | 1,16E-22 | 1 |
| *CG10383* | 2,93E-26 | -2,171414285 | 0,067 | 0,393 | 3,50E-22 | 1 |
| *pum* | 3,83E-26 | 0,56568374 | 1 | 0,993 | 4,57E-22 | 1 |
| *CG5151* | 4,40E-24 | 0,909497502 | 0,924 | 0,769 | 5,25E-20 | 1 |
| *cv-c* | 2,38E-23 | -1,021615167 | 0,824 | 0,942 | 2,83E-19 | 1 |
| *cwo* | 7,93E-23 | -1,413528543 | 0,345 | 0,688 | 9,47E-19 | 1 |
| *wake* | 3,34E-21 | 1,262195054 | 0,609 | 0,396 | 3,99E-17 | 1 |
| *bmm* | 1,72E-20 | -1,78085023 | 0,109 | 0,414 | 2,05E-16 | 1 |
| *stv* | 5,54E-20 | -2,122137453 | 0,034 | 0,282 | 6,60E-16 | 1 |
| *Had2* | 7,11E-20 | 1,209129541 | 0,534 | 0,28 | 8,48E-16 | 1 |
| *PRAS40* | 9,16E-20 | -1,896303462 | 0,105 | 0,402 | 1,09E-15 | 1 |
| *CG46385* | 9,82E-20 | -1,2771657 | 0,55 | 0,791 | 1,17E-15 | 1 |
| *AdSS* | 2,15E-19 | -1,683476971 | 0,013 | 0,227 | 2,56E-15 | 1 |
| *Ppn* | 6,90E-18 | 0,634534332 | 1 | 0,958 | 8,23E-14 | 1 |
| *CG31324* | 7,52E-18 | -1,866685747 | 0,126 | 0,402 | 8,97E-14 | 1 |
| *prage* | 1,06E-17 | -1,761450051 | 0,076 | 0,337 | 1,27E-13 | 1 |
| *RapGAP1* | 1,14E-17 | 0,867906202 | 0,866 | 0,673 | 1,36E-13 | 1 |
| *mamo* | 1,51E-17 | 0,710986212 | 0,95 | 0,828 | 1,80E-13 | 1 |
| *Pdk* | 1,78E-17 | -1,47359081 | 0,189 | 0,479 | 2,12E-13 | 1 |
| *cpx* | 2,63E-17 | 1,362378324 | 0,382 | 0,183 | 3,13E-13 | 1 |
| *psq* | 3,13E-16 | 0,672179839 | 0,916 | 0,799 | 3,73E-12 | 1 |
| *CG18135* | 1,27E-15 | -1,67122856 | 0,16 | 0,388 | 1,52E-11 | 1 |
| *pan* | 1,79E-15 | 0,83397023 | 0,84 | 0,712 | 2,14E-11 | 1 |
| *Prps* | 3,81E-15 | -1,669690425 | 0,16 | 0,392 | 4,54E-11 | 1 |
| *uex* | 1,03E-14 | 0,793185207 | 0,693 | 0,524 | 1,23E-10 | 1 |
| *CG32982* | 2,82E-14 | -1,639023983 | 0,231 | 0,48 | 3,37E-10 | 1 |
| *Prosap* | 5,43E-14 | 0,522203366 | 0,975 | 0,94 | 6,48E-10 | 1 |
| *mtd* | 7,81E-14 | 0,628844989 | 0,895 | 0,73 | 9,31E-10 | 1 |
| *sgg* | 8,95E-14 | -0,783173367 | 0,647 | 0,844 | 1,07E-09 | 1 |
| *Sema5c* | 1,34E-13 | 1,058550061 | 0,66 | 0,48 | 1,60E-09 | 1 |
| *CG42524* | 1,60E-13 | 0,91836379 | 0,529 | 0,331 | 1,91E-09 | 1 |
| *h* | 1,79E-12 | -1,128901694 | 0,176 | 0,425 | 2,13E-08 | 1 |
| *aay* | 3,17E-12 | -1,228569604 | 0,017 | 0,17 | 3,78E-08 | 1 |
| *cact* | 5,87E-12 | -1,009745124 | 0,227 | 0,465 | 7,00E-08 | 1 |
| *CG3777* | 2,00E-11 | 1,086859207 | 0,366 | 0,195 | 2,39E-07 | 1 |
| *hdc* | 2,07E-11 | 0,554701003 | 0,945 | 0,856 | 2,46E-07 | 1 |
| *Syp* | 2,17E-11 | 0,561237458 | 0,933 | 0,833 | 2,59E-07 | 1 |
| *puc* | 2,42E-11 | -1,08151243 | 0,353 | 0,591 | 2,89E-07 | 1 |
| *mbc* | 3,96E-11 | 0,824278803 | 0,567 | 0,464 | 4,73E-07 | 1 |
| *Xrp1* | 4,17E-11 | -0,899936681 | 0,525 | 0,747 | 4,97E-07 | 1 |
| *GstD1* | 6,85E-11 | -1,467784438 | 0,113 | 0,316 | 8,17E-07 | 1 |
| *LpR1* | 2,90E-10 | -1,67255392 | 0,05 | 0,196 | 3,46E-06 | 1 |
| *CG31637* | 3,71E-10 | 0,742259848 | 0,588 | 0,473 | 4,43E-06 | 1 |
| *MtnA* | 3,74E-10 | -0,996960575 | 0,13 | 0,329 | 4,46E-06 | 1 |
| *Thor* | 4,05E-10 | -1,110708788 | 0,223 | 0,44 | 4,84E-06 | 1 |
| *CG3961* | 7,23E-10 | 0,826903706 | 0,521 | 0,342 | 8,63E-06 | 1 |
| *CG43236* | 7,76E-10 | 1,246314055 | 0,256 | 0,113 | 9,26E-06 | 1 |
| *GlcAT-P* | 1,74E-09 | 0,679829365 | 0,672 | 0,502 | 2,07E-05 | 1 |
| *CG5973* | 2,00E-09 | 0,888571739 | 0,399 | 0,225 | 2,39E-05 | 1 |
| *CG41378* | 2,44E-09 | 0,589301057 | 0,681 | 0,591 | 2,91E-05 | 1 |

| | | | | | | |
|---|---|---|---|---|---|---|
| *CG9171* | 3,44E-09 | 0,780961632 | 0,454 | 0,334 | 4,10E-05 | 1 |
| *CG1208* | 4,23E-09 | -0,957227641 | 0,092 | 0,268 | 5,05E-05 | 1 |
| *CG18067* | 4,79E-09 | -1,09200724 | 0,08 | 0,143 | 5,72E-05 | 1 |
| *CG9328* | 5,04E-09 | 0,70928111 | 0,571 | 0,461 | 6,02E-05 | 1 |
| *Fas3* | 6,68E-09 | -1,950841579 | 0,084 | 0,25 | 7,97E-05 | 1 |
| *Pgant9* | 1,00E-08 | 0,471976961 | 0,832 | 0,678 | 0,000119287 | 1 |
| *CG17124* | 1,18E-08 | 0,836099956 | 0,529 | 0,406 | 0,000140309 | 1 |
| *Btk29A* | 1,38E-08 | 0,498342116 | 0,895 | 0,748 | 0,000165018 | 1 |
| *l(1)G0196* | 1,71E-08 | 0,667277642 | 0,534 | 0,377 | 0,000203535 | 1 |
| *Glut4EF* | 1,90E-08 | -0,762351701 | 0,815 | 0,909 | 0,000226383 | 1 |
| *Pkc53E* | 1,97E-08 | 1,015422954 | 0,185 | 0,092 | 0,000234952 | 1 |
| *CG17839* | 1,99E-08 | -1,750711986 | 0,021 | 0,139 | 0,000237652 | 1 |
| *pnr* | 2,15E-08 | 0,59943927 | 0,639 | 0,536 | 0,000255892 | 1 |
| *Mob2* | 2,40E-08 | -0,640082251 | 0,744 | 0,86 | 0,000286005 | 1 |
| *app* | 3,51E-08 | 0,686488669 | 0,454 | 0,375 | 0,000418646 | 1 |
| *CG7530* | 3,53E-08 | -1,203927735 | 0,042 | 0,176 | 0,000420906 | 1 |
| *CG11089* | 3,78E-08 | -0,78130758 | 0,021 | 0,134 | 0,00045053 | 1 |
| *mnb* | 3,84E-08 | -0,853724446 | 0,231 | 0,429 | 0,000457615 | 1 |
| *kuz* | 4,59E-08 | 0,487305344 | 0,891 | 0,806 | 0,000547975 | 1 |
| *shep* | 4,62E-08 | 0,298010782 | 1 | 0,983 | 0,000551691 | 1 |
| *sra* | 5,19E-08 | -0,866504549 | 0,092 | 0,245 | 0,00061942 | 1 |
| *GEFmeso* | 5,57E-08 | -0,775285947 | 0,269 | 0,471 | 0,000664115 | 1 |
| *Fer1HCH* | 5,98E-08 | -0,854784429 | 0,374 | 0,574 | 0,000713355 | 1 |
| *CG3376* | 6,54E-08 | -0,826736323 | 0,29 | 0,487 | 0,000780663 | 1 |
| *CG11050* | 6,93E-08 | -1,034669093 | 0,017 | 0,123 | 0,000826602 | 1 |
| *flw* | 7,20E-08 | 0,699439026 | 0,429 | 0,342 | 0,000858697 | 1 |
| *Fer2LCH* | 7,71E-08 | -0,846246193 | 0,399 | 0,593 | 0,000920095 | 1 |
| *Atg8a* | 7,90E-08 | -0,890773238 | 0,197 | 0,387 | 0,000942985 | 1 |
| *RpLP2* | 8,03E-08 | -0,914837371 | 0,101 | 0,223 | 0,000958246 | 1 |
| *MESR3* | 8,05E-08 | 0,476518466 | 0,777 | 0,737 | 0,000959961 | 1 |
| *CG1943* | 8,25E-08 | -0,891072688 | 0,08 | 0,228 | 0,000984106 | 1 |
| *MFS17* | 9,22E-08 | 0,50758145 | 0,739 | 0,676 | 0,001099731 | 1 |
| *kay* | 1,13E-07 | -0,878906222 | 0,252 | 0,449 | 0,00135366 | 1 |
| *Galphao* | 1,16E-07 | 0,705821803 | 0,378 | 0,289 | 0,001380046 | 1 |
| *jim* | 1,23E-07 | -0,758217961 | 0,29 | 0,49 | 0,001473108 | 1 |
| *CenG1A* | 1,36E-07 | -0,886976327 | 0,059 | 0,189 | 0,001620633 | 1 |
| *nkd* | 1,55E-07 | -1,131840514 | 0,071 | 0,22 | 0,001844877 | 1 |
| *Gadd45* | 1,60E-07 | -1,231228211 | 0,038 | 0,167 | 0,001905773 | 1 |
| *ham* | 1,60E-07 | 0,776409622 | 0,349 | 0,249 | 0,001909034 | 1 |
| *CG31183* | 1,72E-07 | 0,911780342 | 0,197 | 0,116 | 0,002051176 | 1 |
| *DIP-lambda* | 1,79E-07 | 0,63577876 | 0,609 | 0,481 | 0,0021355 | 1 |
| *Rala* | 1,85E-07 | 0,488000922 | 0,538 | 0,495 | 0,002202534 | 1 |
| *rl* | 1,92E-07 | 0,297637562 | 0,987 | 0,962 | 0,002286247 | 1 |
| *CG45050* | 2,14E-07 | -0,507568053 | 0,601 | 0,777 | 0,002553196 | 1 |
| *cnc* | 2,41E-07 | -0,5556084 | 0,546 | 0,724 | 0,002874899 | 1 |
| *ena* | 2,66E-07 | 0,618931724 | 0,689 | 0,562 | 0,003177817 | 1 |
| *CG42694* | 3,12E-07 | 0,869007203 | 0,172 | 0,086 | 0,003722247 | 1 |
| *Pect* | 3,14E-07 | -0,579660852 | 0,155 | 0,307 | 0,00375094 | 1 |
| *Nrg* | 3,39E-07 | -1,417852383 | 0,042 | 0,163 | 0,00404955 | 1 |
| *DIP-alpha* | 3,98E-07 | -0,346847707 | 0,479 | 0,667 | 0,004750651 | 1 |
| *RpL19* | 4,50E-07 | -1,005047921 | 0,168 | 0,324 | 0,00537241 | 1 |
| *fax* | 5,59E-07 | -0,702951604 | 0,466 | 0,646 | 0,006664092 | 1 |
| *anne* | 5,73E-07 | 0,44947108 | 0,294 | 0,269 | 0,006838497 | 1 |
| *Dbp80* | 5,88E-07 | 0,486783545 | 0,752 | 0,676 | 0,007015942 | 1 |
| *cv-2* | 6,37E-07 | -1,167564817 | 0,013 | 0,101 | 0,007601382 | 1 |
| *oys* | 6,66E-07 | -0,715235001 | 0,147 | 0,298 | 0,007944261 | 1 |
| *ush* | 6,70E-07 | 0,491086221 | 0,811 | 0,704 | 0,007986747 | 1 |
| *Atg18b* | 6,90E-07 | -0,808639549 | 0,126 | 0,277 | 0,008231556 | 1 |
| *CG3168* | 7,09E-07 | -1,160814111 | 0,025 | 0,126 | 0,008461654 | 1 |
| *SCaMC* | 7,70E-07 | -0,964803853 | 0,147 | 0,315 | 0,009187974 | 1 |
| *CG33144* | 8,07E-07 | 0,626423055 | 0,521 | 0,404 | 0,009623027 | 1 |

| | | | | | | |
|---|---|---|---|---|---|---|
| *unc-13* | 8,23E-07 | 0,504624298 | 0,664 | 0,566 | 0,009815376 | 1 |
| *unk* | 8,34E-07 | -0,850913581 | 0,097 | 0,244 | 0,009944936 | 1 |
| *caps* | 8,90E-07 | -1,276281859 | 0,021 | 0,121 | 0,010622334 | 1 |
| *ref(2)P* | 8,98E-07 | -0,69184126 | 0,063 | 0,19 | 0,010709409 | 1 |
| *spri* | 9,11E-07 | 0,413174863 | 0,798 | 0,745 | 0,010864451 | 1 |
| *Col4a1* | 9,39E-07 | 0,347456471 | 0,975 | 0,914 | 0,011204824 | 1 |
| *lbk* | 9,61E-07 | -0,66898761 | 0,45 | 0,611 | 0,011462034 | 1 |
| *CG13868* | 9,69E-07 | -0,800653207 | 0,244 | 0,431 | 0,011561222 | 1 |
| *RpS9* | 1,03E-06 | -0,777165015 | 0,13 | 0,272 | 0,012322372 | 1 |
| *Tgi* | 1,10E-06 | 0,700543226 | 0,298 | 0,205 | 0,013122325 | 1 |
| *Atg1* | 1,21E-06 | -0,821693401 | 0,16 | 0,321 | 0,014480078 | 1 |
| *Fim* | 1,22E-06 | -0,687396612 | 0,223 | 0,398 | 0,014497075 | 1 |
| *apolpp* | 1,23E-06 | 0,616339242 | 0,487 | 0,323 | 0,014709974 | 1 |
| *Gel* | 1,25E-06 | 0,577170649 | 0,353 | 0,286 | 0,01485513 | 1 |
| *Cals* | 1,46E-06 | 0,450055656 | 0,252 | 0,248 | 0,017371666 | 1 |
| *raskol* | 1,72E-06 | 0,557872487 | 0,248 | 0,214 | 0,020519203 | 1 |
| *CG15784* | 1,74E-06 | -0,874017661 | 0,025 | 0,116 | 0,020717094 | 1 |
| *NimC4* | 1,86E-06 | -1,232853492 | 0,017 | 0,11 | 0,022178172 | 1 |
| *Gyf* | 1,93E-06 | 0,436956095 | 0,336 | 0,319 | 0,022983415 | 1 |
| *DOR* | 1,97E-06 | 0,68273003 | 0,357 | 0,282 | 0,023546296 | 1 |
| *AnxB9* | 2,08E-06 | -0,869255501 | 0,189 | 0,354 | 0,024846927 | 1 |
| *NaCP60E* | 2,12E-06 | 0,709163779 | 0,319 | 0,226 | 0,025325588 | 1 |
| *ckn* | 2,27E-06 | 0,647770715 | 0,286 | 0,218 | 0,027108442 | 1 |
| *Hex-A* | 2,45E-06 | -0,744773546 | 0,155 | 0,314 | 0,02921733 | 1 |
| *Pde9* | 2,45E-06 | 0,592106944 | 0,681 | 0,576 | 0,029259145 | 1 |
| *Lmpt* | 2,47E-06 | -1,146657422 | 0,029 | 0,134 | 0,029506033 | 1 |
| *CG14207* | 2,48E-06 | -0,762754583 | 0,017 | 0,105 | 0,029579047 | 1 |
| *Hsc70Cb* | 2,64E-06 | -0,439465747 | 0,139 | 0,266 | 0,031441861 | 1 |
| *dl* | 2,85E-06 | -1,101914448 | 0,038 | 0,15 | 0,034044481 | 1 |
| *CG33298* | 3,01E-06 | 0,544607821 | 0,5 | 0,413 | 0,035909077 | 1 |
| *CG18628* | 3,46E-06 | -0,670396253 | 0,029 | 0,13 | 0,04122654 | 1 |
| *RhoGAP15B* | 3,50E-06 | -0,705549564 | 0,105 | 0,234 | 0,041768281 | 1 |
| *shot* | 3,60E-06 | 0,344190828 | 0,912 | 0,893 | 0,042948716 | 1 |
| *Ubi-p63E* | 3,68E-06 | -0,94550535 | 0,265 | 0,439 | 0,043839824 | 1 |
| *Mct1* | 3,78E-06 | -0,31805236 | 0,092 | 0,181 | 0,045051195 | 1 |
| *Pcyt1* | 3,97E-06 | -0,618343558 | 0,046 | 0,144 | 0,047376732 | 1 |
| *step* | 3,97E-06 | -0,29934965 | 0,298 | 0,449 | 0,04739655 | 1 |
| *cic* | 3,33E-32 | 1,195411682 | 0,991 | 0,949 | 3,98E-28 | 2 |
| *Glut4EF.1* | 2,39E-31 | 1,127211294 | 0,968 | 0,878 | 2,85E-27 | 2 |
| *Lpin.1* | 3,52E-29 | 1,58315817 | 0,605 | 0,253 | 4,20E-25 | 2 |
| *Spn.1* | 5,61E-24 | 1,028860096 | 0,855 | 0,637 | 6,69E-20 | 2 |
| *Mmp2.1* | 1,63E-22 | -2,042045334 | 0,168 | 0,517 | 1,94E-18 | 2 |
| *Mob2.1* | 6,65E-21 | 0,849549961 | 0,936 | 0,821 | 7,93E-17 | 2 |
| *sgg.1* | 5,73E-20 | 0,78487205 | 0,95 | 0,782 | 6,84E-16 | 2 |
| *PRAS40.1* | 6,18E-20 | 1,411203266 | 0,577 | 0,307 | 7,37E-16 | 2 |
| *CG18135.1* | 6,50E-19 | 1,282831868 | 0,541 | 0,311 | 7,75E-15 | 2 |
| *InR.1* | 4,29E-18 | 0,706142606 | 0,973 | 0,83 | 5,12E-14 | 2 |
| *Kr-h1.1* | 1,64E-17 | 1,052617263 | 0,7 | 0,439 | 1,96E-13 | 2 |
| *Pvr* | 9,81E-17 | 0,667890403 | 0,968 | 0,86 | 1,17E-12 | 2 |
| *Crtc.1* | 1,22E-16 | 0,928646068 | 0,7 | 0,5 | 1,45E-12 | 2 |
| *RhoGAP71E* | 2,33E-16 | -1,4354431 | 0,309 | 0,601 | 2,78E-12 | 2 |
| *Cip4* | 3,71E-16 | -1,549714887 | 0,286 | 0,58 | 4,42E-12 | 2 |
| *Sema2a.1* | 3,81E-16 | -1,659843544 | 0,209 | 0,493 | 4,54E-12 | 2 |
| *Msp300* | 4,28E-16 | -1,664260438 | 0,355 | 0,613 | 5,11E-12 | 2 |
| *CHES-1-like.1* | 5,81E-16 | 0,947789176 | 0,491 | 0,351 | 6,93E-12 | 2 |
| *jim.1* | 3,19E-15 | 0,947161863 | 0,641 | 0,42 | 3,81E-11 | 2 |
| *Pdk.1* | 1,07E-14 | 0,968747964 | 0,627 | 0,39 | 1,27E-10 | 2 |
| *CG12163* | 1,39E-14 | 0,834244537 | 0,691 | 0,537 | 1,65E-10 | 2 |
| *luna* | 1,43E-14 | -1,042325645 | 0,65 | 0,835 | 1,70E-10 | 2 |
| *Pxn.1* | 2,15E-14 | -1,356809894 | 0,309 | 0,587 | 2,56E-10 | 2 |
| *CG5953.1* | 3,20E-14 | 0,875665368 | 0,705 | 0,449 | 3,82E-10 | 2 |

| | | | | | | |
|---|---|---|---|---|---|---|
| *CG10383.1* | 3,84E-14 | 1,067552019 | 0,545 | 0,295 | 4,59E-10 | 2 |
| *CG32982.1* | 7,96E-14 | 1,174560637 | 0,614 | 0,402 | 9,50E-10 | 2 |
| *cwo.1* | 1,38E-13 | 0,871098429 | 0,759 | 0,603 | 1,64E-09 | 2 |
| *regucalcin.1* | 2,64E-13 | -0,991607534 | 0,259 | 0,539 | 3,15E-09 | 2 |
| *CG5059.1* | 3,21E-13 | 0,951045836 | 0,518 | 0,315 | 3,83E-09 | 2 |
| *egh.1* | 5,87E-13 | -1,115130537 | 0,195 | 0,465 | 7,01E-09 | 2 |
| *mnb.1* | 9,41E-13 | 0,950868218 | 0,582 | 0,358 | 1,12E-08 | 2 |
| *CG7530.1* | 9,77E-13 | 1,243891248 | 0,273 | 0,129 | 1,16E-08 | 2 |
| *Dyrk2* | 1,01E-12 | 0,853053473 | 0,714 | 0,535 | 1,20E-08 | 2 |
| *CG31324.1* | 1,26E-12 | 1,047283282 | 0,545 | 0,317 | 1,50E-08 | 2 |
| *Pli.1* | 1,63E-12 | 0,967006483 | 0,591 | 0,377 | 1,94E-08 | 2 |
| *hdc.1* | 2,04E-12 | -0,684330599 | 0,75 | 0,895 | 2,44E-08 | 2 |
| *Pax* | 3,87E-12 | -1,44562112 | 0,064 | 0,261 | 4,62E-08 | 2 |
| *CG13868.1* | 4,42E-12 | 0,920318409 | 0,555 | 0,368 | 5,27E-08 | 2 |
| *alphaTub84B* | 5,44E-12 | -1,1886492 | 0,073 | 0,269 | 6,49E-08 | 2 |
| *spir* | 5,92E-12 | -1,426507376 | 0,255 | 0,493 | 7,06E-08 | 2 |
| *CG10543* | 7,12E-12 | 0,678590209 | 0,809 | 0,722 | 8,50E-08 | 2 |
| *mgl* | 7,49E-12 | -1,7654156 | 0,123 | 0,355 | 8,93E-08 | 2 |
| *CG17574.1* | 1,54E-11 | -0,981551244 | 0,427 | 0,665 | 1,84E-07 | 2 |
| *gish* | 2,02E-11 | -0,980362013 | 0,277 | 0,542 | 2,41E-07 | 2 |
| *Rtnl1* | 4,17E-11 | -1,32707943 | 0,009 | 0,142 | 4,98E-07 | 2 |
| *RanBPM.1* | 4,42E-11 | 0,910915815 | 0,414 | 0,262 | 5,27E-07 | 2 |
| *flw.1* | 5,70E-11 | -1,059416747 | 0,164 | 0,394 | 6,80E-07 | 2 |
| *Moe* | 5,97E-11 | -0,780539784 | 0,264 | 0,511 | 7,12E-07 | 2 |
| *zip* | 6,38E-11 | -1,075563409 | 0,123 | 0,338 | 7,61E-07 | 2 |
| *Prps.1* | 9,54E-11 | 0,959630746 | 0,459 | 0,33 | 1,14E-06 | 2 |
| *aqz* | 1,49E-10 | 0,494790937 | 0,964 | 0,885 | 1,78E-06 | 2 |
| *Act42A.1* | 2,01E-10 | -1,275516841 | 0,036 | 0,199 | 2,40E-06 | 2 |
| *NimB4* | 2,83E-10 | 0,67490707 | 0,477 | 0,378 | 3,38E-06 | 2 |
| *CG3376.1* | 3,50E-10 | 0,864209488 | 0,605 | 0,423 | 4,18E-06 | 2 |
| *sr* | 4,38E-10 | 0,72135453 | 0,668 | 0,51 | 5,22E-06 | 2 |
| *CG32486* | 4,38E-10 | 0,634747292 | 0,632 | 0,523 | 5,23E-06 | 2 |
| *Atg1.1* | 5,04E-10 | 0,881977994 | 0,391 | 0,274 | 6,02E-06 | 2 |
| *Fas3.1* | 6,29E-10 | -2,207760271 | 0,082 | 0,247 | 7,51E-06 | 2 |
| *betaTub60D* | 7,38E-10 | -1,644408472 | 0,095 | 0,285 | 8,80E-06 | 2 |
| *CG43867* | 1,31E-09 | -1,450464877 | 0,036 | 0,189 | 1,56E-05 | 2 |
| *uex.1* | 1,44E-09 | -0,613377371 | 0,355 | 0,592 | 1,71E-05 | 2 |
| *bmm.1* | 2,00E-09 | 0,890654957 | 0,527 | 0,329 | 2,38E-05 | 2 |
| *MYPT-75D* | 2,81E-09 | 0,795699126 | 0,705 | 0,509 | 3,35E-05 | 2 |
| *AdSS.1* | 3,33E-09 | 0,94488504 | 0,309 | 0,167 | 3,97E-05 | 2 |
| *Thor.1* | 3,93E-09 | 0,885166446 | 0,545 | 0,374 | 4,68E-05 | 2 |
| *CG46385.1* | 4,45E-09 | 0,618168059 | 0,85 | 0,73 | 5,31E-05 | 2 |
| *CrebA.1* | 4,56E-09 | -1,251941755 | 0,095 | 0,288 | 5,45E-05 | 2 |
| *drpr* | 5,92E-09 | 0,571670516 | 0,855 | 0,733 | 7,06E-05 | 2 |
| *l(3)L1231* | 6,01E-09 | 0,499198967 | 0,773 | 0,719 | 7,17E-05 | 2 |
| *lbk.1* | 7,27E-09 | 0,621526784 | 0,673 | 0,566 | 8,67E-05 | 2 |
| *CG5888.1* | 1,05E-08 | -0,832188794 | 0,164 | 0,355 | 0,00012471 | 2 |
| *Atg17.1* | 1,08E-08 | 0,744417727 | 0,455 | 0,315 | 0,000129324 | 2 |
| *CG42524.1* | 1,10E-08 | -0,800872528 | 0,195 | 0,399 | 0,000130948 | 2 |
| *chic* | 1,24E-08 | -0,656245667 | 0,205 | 0,399 | 0,000147991 | 2 |
| *kermit* | 1,54E-08 | 0,965480807 | 0,136 | 0,051 | 0,000183995 | 2 |
| *app.1* | 1,70E-08 | -0,861512227 | 0,214 | 0,422 | 0,00020245 | 2 |
| *CG5973.1* | 1,82E-08 | -1,190474732 | 0,105 | 0,284 | 0,000217263 | 2 |
| *sra.1* | 1,87E-08 | 0,881504915 | 0,345 | 0,194 | 0,000223422 | 2 |
| *CG32767* | 2,07E-08 | 0,659562052 | 0,714 | 0,598 | 0,000247185 | 2 |
| *B4* | 2,21E-08 | -1,160317687 | 0,023 | 0,151 | 0,00026355 | 2 |
| *fs(1)h.1* | 2,22E-08 | 0,501816735 | 0,709 | 0,677 | 0,000264345 | 2 |
| *LpR1.1* | 2,84E-08 | 1,199901333 | 0,291 | 0,148 | 0,000338289 | 2 |
| *Smr* | 2,86E-08 | 0,478389654 | 0,886 | 0,851 | 0,000341489 | 2 |
| *scb* | 2,99E-08 | -1,119040681 | 0,036 | 0,171 | 0,000357161 | 2 |
| *Cam* | 3,21E-08 | -0,965880044 | 0,345 | 0,542 | 0,000382543 | 2 |

| | | | | | | |
|---|---|---|---|---|---|---|
| *foxo* | 3,29E-08 | 0,602016225 | 0,659 | 0,531 | 0,000392319 | 2 |
| *pigs.1* | 3,42E-08 | -0,7294682 | 0,391 | 0,62 | 0,000407823 | 2 |
| *CG4250.1* | 4,19E-08 | -0,917121773 | 0,191 | 0,385 | 0,000499927 | 2 |
| *LanA.1* | 5,38E-08 | -0,92759987 | 0,077 | 0,229 | 0,000642281 | 2 |
| *Had2.1* | 6,29E-08 | -1,000287285 | 0,164 | 0,355 | 0,000750539 | 2 |
| *CG4404.1* | 6,31E-08 | 0,900651281 | 0,264 | 0,147 | 0,00075297 | 2 |
| *Mkp3* | 6,55E-08 | -1,157353817 | 0,15 | 0,347 | 0,000781088 | 2 |
| *CG8679* | 9,27E-08 | 0,848005899 | 0,127 | 0,058 | 0,00110541 | 2 |
| *CG9005* | 9,39E-08 | 0,73031646 | 0,482 | 0,366 | 0,001119556 | 2 |
| *CG43236.1* | 1,02E-07 | -1,226345903 | 0,032 | 0,158 | 0,001220568 | 2 |
| *px* | 1,17E-07 | 0,469620742 | 0,805 | 0,77 | 0,001395568 | 2 |
| *crq* | 1,22E-07 | -0,384619421 | 0,282 | 0,465 | 0,001451022 | 2 |
| *Sap47* | 1,64E-07 | 0,66516154 | 0,495 | 0,408 | 0,001954341 | 2 |
| *Naam* | 1,80E-07 | -1,351695954 | 0,114 | 0,278 | 0,002151083 | 2 |
| *bbc* | 1,94E-07 | -0,882725593 | 0,095 | 0,25 | 0,00231971 | 2 |
| *Blimp-1* | 1,98E-07 | 0,517219209 | 0,614 | 0,553 | 0,002362464 | 2 |
| *Atg18b.1* | 2,03E-07 | 0,70809521 | 0,35 | 0,232 | 0,002421418 | 2 |
| *CG9003* | 2,43E-07 | 0,689013159 | 0,455 | 0,306 | 0,002903126 | 2 |
| *Frl* | 2,65E-07 | -0,910499008 | 0,236 | 0,426 | 0,003165308 | 2 |
| *norpA* | 2,92E-07 | -0,664092054 | 0,282 | 0,478 | 0,003488932 | 2 |
| *CG6191* | 2,96E-07 | 0,82946583 | 0,182 | 0,105 | 0,003525125 | 2 |
| *zfh1* | 3,07E-07 | 0,431924333 | 0,973 | 0,901 | 0,00365779 | 2 |
| *Gapvd1* | 3,14E-07 | 0,77456655 | 0,155 | 0,079 | 0,003749263 | 2 |
| *firl.1* | 3,31E-07 | -1,25016564 | 0,032 | 0,152 | 0,003943156 | 2 |
| *EDTP* | 3,43E-07 | 0,853396844 | 0,314 | 0,187 | 0,004086931 | 2 |
| *kst* | 3,99E-07 | -0,789578025 | 0,082 | 0,216 | 0,004757323 | 2 |
| *PDZ-GEF* | 4,02E-07 | 0,620671775 | 0,336 | 0,257 | 0,004791297 | 2 |
| *CG9674.1* | 4,13E-07 | 0,592728164 | 0,459 | 0,286 | 0,00492422 | 2 |
| *klar.1* | 4,36E-07 | -0,637603212 | 0,405 | 0,61 | 0,005206213 | 2 |
| *CG1208.1* | 4,61E-07 | 0,940504074 | 0,364 | 0,213 | 0,005498085 | 2 |
| *CG42674* | 4,87E-07 | -1,11975084 | 0,027 | 0,138 | 0,005809727 | 2 |
| *sick* | 5,29E-07 | -1,108740637 | 0,018 | 0,12 | 0,006312427 | 2 |
| *apolpp.1* | 5,65E-07 | -0,484966789 | 0,214 | 0,378 | 0,006736452 | 2 |
| *Sema2b.1* | 5,95E-07 | 0,701128334 | 0,323 | 0,26 | 0,007102391 | 2 |
| *dikar* | 6,05E-07 | 0,521892858 | 0,436 | 0,382 | 0,007221517 | 2 |
| *CG32264.1* | 6,08E-07 | -0,747078267 | 0,286 | 0,476 | 0,007248556 | 2 |
| *wake.1* | 6,16E-07 | -0,981227405 | 0,273 | 0,464 | 0,007351214 | 2 |
| *CG3328* | 6,20E-07 | 0,514715222 | 0,595 | 0,534 | 0,00739821 | 2 |
| *hfp* | 6,54E-07 | 0,632404435 | 0,318 | 0,24 | 0,007804761 | 2 |
| *sno* | 7,38E-07 | 0,718987771 | 0,209 | 0,142 | 0,008801617 | 2 |
| *Hml.1* | 7,41E-07 | -0,890346184 | 0,532 | 0,674 | 0,008844858 | 2 |
| *shep.1* | 7,50E-07 | -0,368158085 | 0,973 | 0,989 | 0,00894647 | 2 |
| *CG12065* | 8,60E-07 | 0,489267498 | 0,668 | 0,593 | 0,010258349 | 2 |
| *CG42668* | 9,33E-07 | -0,884936771 | 0,1 | 0,252 | 0,011133254 | 2 |
| *SCaMC.1* | 9,45E-07 | 0,626013497 | 0,323 | 0,279 | 0,011269576 | 2 |
| *CG12576.1* | 9,61E-07 | 0,753726059 | 0,373 | 0,254 | 0,011458242 | 2 |
| *cora.1* | 9,63E-07 | -0,611849604 | 0,323 | 0,517 | 0,011483531 | 2 |
| *Tlk.1* | 9,97E-07 | 0,422565927 | 0,686 | 0,679 | 0,011890186 | 2 |
| *eEF2.1* | 1,17E-06 | 0,269476638 | 0,268 | 0,323 | 0,013989008 | 2 |
| *unk.1* | 1,22E-06 | 0,821838941 | 0,323 | 0,198 | 0,01454296 | 2 |
| *mbc.1* | 1,31E-06 | -0,602637841 | 0,323 | 0,512 | 0,015624361 | 2 |
| *Cyp4ac3* | 1,33E-06 | 0,840257999 | 0,132 | 0,058 | 0,015902592 | 2 |
| *CG6891* | 1,42E-06 | -1,148447648 | 0,009 | 0,101 | 0,016946269 | 2 |
| *CG18171* | 1,59E-06 | 0,58107429 | 0,364 | 0,3 | 0,018933345 | 2 |
| *AdipoR* | 1,60E-06 | 0,567926158 | 0,182 | 0,163 | 0,019076946 | 2 |
| *CG3777.1* | 1,65E-06 | -1,006924065 | 0,095 | 0,25 | 0,019649085 | 2 |
| *ATP8A* | 1,81E-06 | -1,042495629 | 0,059 | 0,186 | 0,021597218 | 2 |
| *cdi* | 1,89E-06 | -0,798662699 | 0,159 | 0,327 | 0,022493836 | 2 |
| *CG6357* | 1,93E-06 | -0,62733401 | 0,027 | 0,116 | 0,023053887 | 2 |
| *bves* | 1,96E-06 | -0,783450625 | 0,114 | 0,257 | 0,02336286 | 2 |
| *Ziz.1* | 2,04E-06 | 0,761145298 | 0,441 | 0,313 | 0,024368734 | 2 |

| | | | | | | |
|---|---|---|---|---|---|---|
| *RhoGAP93B* | 2,27E-06 | -0,531962321 | 0,055 | 0,148 | 0,027119049 | 2 |
| *mtgo* | 2,50E-06 | 0,313741275 | 0,982 | 0,96 | 0,029830105 | 2 |
| *Socs16D* | 2,60E-06 | 0,674151711 | 0,164 | 0,12 | 0,031026729 | 2 |
| *bwa.1* | 2,63E-06 | 0,767085549 | 0,25 | 0,14 | 0,031340938 | 2 |
| *rudhira* | 2,68E-06 | 0,53424321 | 0,664 | 0,556 | 0,031914014 | 2 |
| *Ran* | 2,79E-06 | -0,530657886 | 0,027 | 0,107 | 0,03324765 | 2 |
| *tai* | 3,01E-06 | 0,42422894 | 0,877 | 0,809 | 0,035910174 | 2 |
| *Pdcd4.1* | 3,06E-06 | 0,623098869 | 0,168 | 0,13 | 0,036470868 | 2 |
| *Syp.1* | 3,10E-06 | -0,455943656 | 0,75 | 0,87 | 0,036983737 | 2 |
| *pan.1* | 3,12E-06 | -0,468904521 | 0,591 | 0,762 | 0,037205646 | 2 |
| *l(3)80Fg.1* | 3,13E-06 | -0,733575388 | 0,477 | 0,635 | 0,037387454 | 2 |
| *Usp15-31* | 4,03E-06 | -0,945177399 | 0,045 | 0,158 | 0,048045401 | 2 |
| *Kr-h1.2* | 2,16E-18 | 0,849086392 | 0,762 | 0,43 | 2,57E-14 | 3 |
| *CG10383.2* | 9,80E-17 | 0,6962719 | 0,586 | 0,29 | 1,17E-12 | 3 |
| *InR.2* | 1,53E-16 | 0,522828663 | 0,981 | 0,83 | 1,82E-12 | 3 |
| *CG3376.2* | 2,31E-16 | 0,813070251 | 0,714 | 0,405 | 2,75E-12 | 3 |
| *Pvr.1* | 1,83E-15 | 0,53802433 | 0,99 | 0,857 | 2,18E-11 | 3 |
| *sgg.2* | 3,00E-15 | 0,694113896 | 0,929 | 0,788 | 3,58E-11 | 3 |
| *CG12576.2* | 7,33E-15 | 0,687553816 | 0,49 | 0,234 | 8,75E-11 | 3 |
| *CG32982.2* | 8,72E-15 | 1,049622817 | 0,662 | 0,395 | 1,04E-10 | 3 |
| *CG1208.2* | 1,60E-14 | 0,754602377 | 0,448 | 0,199 | 1,91E-10 | 3 |
| *Lpin.2* | 1,70E-14 | 0,707664257 | 0,543 | 0,267 | 2,03E-10 | 3 |
| *cwo.2* | 1,92E-14 | 0,650525096 | 0,852 | 0,588 | 2,29E-10 | 3 |
| *bmm.2* | 2,65E-13 | 0,977922987 | 0,59 | 0,319 | 3,17E-09 | 3 |
| *sra.2* | 2,95E-13 | 0,855297818 | 0,414 | 0,183 | 3,52E-09 | 3 |
| *PDZ-GEF.1* | 4,99E-13 | 0,537590571 | 0,457 | 0,236 | 5,95E-09 | 3 |
| *CG13982* | 7,34E-13 | 1,01822441 | 0,371 | 0,137 | 8,76E-09 | 3 |
| *Atg17.2* | 8,70E-13 | 0,767421715 | 0,562 | 0,296 | 1,04E-08 | 3 |
| *Ser* | 4,70E-12 | 0,507807775 | 0,571 | 0,326 | 5,60E-08 | 3 |
| *uex.2* | 4,81E-12 | -0,514067781 | 0,576 | 0,549 | 5,74E-08 | 3 |
| *Pdk.2* | 5,79E-12 | 0,71299898 | 0,648 | 0,388 | 6,90E-08 | 3 |
| *CG5953.2* | 6,08E-12 | 0,309978405 | 0,695 | 0,453 | 7,26E-08 | 3 |
| *NimB4.1* | 6,65E-12 | 0,392947481 | 0,586 | 0,36 | 7,93E-08 | 3 |
| *MYPT-75D.1* | 6,93E-12 | 0,878228608 | 0,743 | 0,503 | 8,27E-08 | 3 |
| *CG32486.1* | 7,44E-12 | 0,448066824 | 0,752 | 0,502 | 8,88E-08 | 3 |
| *C1GalTA.2* | 3,93E-11 | 0,436919414 | 0,433 | 0,244 | 4,69E-07 | 3 |
| *CG1943.2* | 4,48E-11 | 0,383535887 | 0,352 | 0,175 | 5,34E-07 | 3 |
| *lbk.2* | 5,68E-11 | 0,629899469 | 0,776 | 0,548 | 6,77E-07 | 3 |
| *Hsc70-4.1* | 8,07E-11 | -0,575325679 | 0,314 | 0,289 | 9,62E-07 | 3 |
| *Pkn.1* | 8,95E-11 | 0,422685533 | 0,729 | 0,488 | 1,07E-06 | 3 |
| *Myc.1* | 1,13E-10 | -0,87801498 | 0,462 | 0,467 | 1,35E-06 | 3 |
| *Spn.2* | 1,58E-10 | 0,462651191 | 0,867 | 0,637 | 1,89E-06 | 3 |
| *flfl.2* | 1,68E-10 | 0,399352113 | 0,314 | 0,159 | 2,00E-06 | 3 |
| *CG12054* | 1,78E-10 | 0,289973211 | 0,41 | 0,237 | 2,12E-06 | 3 |
| *mam* | 2,16E-10 | 0,345877289 | 0,571 | 0,364 | 2,57E-06 | 3 |
| *hep* | 2,20E-10 | 0,260588722 | 0,424 | 0,263 | 2,62E-06 | 3 |
| *Fer2LCH.1* | 3,64E-10 | 0,522366239 | 0,752 | 0,524 | 4,34E-06 | 3 |
| *cv-c.2* | 3,69E-10 | 0,366434481 | 0,99 | 0,909 | 4,40E-06 | 3 |
| *CG12065.1* | 4,23E-10 | 0,383441963 | 0,8 | 0,569 | 5,05E-06 | 3 |
| *cpo.1* | 5,18E-10 | -0,370192429 | 0,776 | 0,688 | 6,18E-06 | 3 |
| *CG46385.2* | 5,31E-10 | 0,526170348 | 0,9 | 0,722 | 6,33E-06 | 3 |
| *CG12163.1* | 5,36E-10 | 0,448970024 | 0,757 | 0,526 | 6,39E-06 | 3 |
| *pcs.2* | 5,51E-10 | 0,272977131 | 0,324 | 0,181 | 6,57E-06 | 3 |
| *sbr.1* | 5,94E-10 | 0,258938558 | 0,405 | 0,25 | 7,09E-06 | 3 |
| *Pdp1.1* | 7,99E-10 | -0,543294913 | 0,595 | 0,6 | 9,53E-06 | 3 |
| *PRAS40.2* | 8,54E-10 | 0,412201036 | 0,529 | 0,318 | 1,02E-05 | 3 |
| *Crtc.2* | 8,65E-10 | 0,530938172 | 0,729 | 0,497 | 1,03E-05 | 3 |
| *Atg18b.2* | 9,25E-10 | 0,622293297 | 0,424 | 0,219 | 1,10E-05 | 3 |
| *Karl* | 1,38E-09 | 0,458184354 | 0,929 | 0,761 | 1,65E-05 | 3 |
| *CG5059.2* | 1,70E-09 | 0,370315415 | 0,519 | 0,316 | 2,03E-05 | 3 |
| *Mbs* | 2,22E-09 | 0,50828941 | 0,576 | 0,35 | 2,65E-05 | 3 |

| | | | | | | |
|---|---|---|---|---|---|---|
| *Msp300.1* | 2,30E-09 | -0,89848061 | 0,5 | 0,584 | 2,75E-05 | 3 |
| *CG12012.2* | 2,32E-09 | 0,392779927 | 0,319 | 0,166 | 2,76E-05 | 3 |
| *RpS26* | 2,54E-09 | -0,398812929 | 0,295 | 0,241 | 3,03E-05 | 3 |
| *CG6966* | 2,56E-09 | 0,333949808 | 0,424 | 0,264 | 3,06E-05 | 3 |
| *hid.1* | 2,97E-09 | 0,488209505 | 0,557 | 0,338 | 3,54E-05 | 3 |
| *CG4404.2* | 3,00E-09 | 0,516892476 | 0,29 | 0,143 | 3,58E-05 | 3 |
| *REPTOR.2* | 3,01E-09 | 0,431159125 | 0,786 | 0,562 | 3,59E-05 | 3 |
| *kay.1* | 4,09E-09 | -0,260598608 | 0,495 | 0,4 | 4,87E-05 | 3 |
| *Mob2.2* | 4,20E-09 | 0,565266403 | 0,938 | 0,821 | 5,00E-05 | 3 |
| *CG11050.1* | 4,39E-09 | 0,680356184 | 0,229 | 0,081 | 5,24E-05 | 3 |
| *Sema2b.2* | 4,68E-09 | 0,627136011 | 0,443 | 0,239 | 5,59E-05 | 3 |
| *Eip74EF.2* | 4,70E-09 | 0,46015826 | 0,762 | 0,55 | 5,60E-05 | 3 |
| *Yeti* | 6,92E-09 | -0,330125724 | 0,362 | 0,328 | 8,25E-05 | 3 |
| *Exn.1* | 7,51E-09 | 0,540975006 | 0,495 | 0,282 | 8,96E-05 | 3 |
| *Trpm.1* | 8,01E-09 | 0,293946504 | 0,619 | 0,426 | 9,56E-05 | 3 |
| *Glut4EF.2* | 8,16E-09 | 0,384225575 | 0,976 | 0,877 | 9,74E-05 | 3 |
| *CG17574.2* | 8,34E-09 | -0,572717798 | 0,595 | 0,633 | 9,95E-05 | 3 |
| *CG32091.1* | 9,38E-09 | 0,45849747 | 0,833 | 0,64 | 0,000111945 | 3 |
| *Socs36E.1* | 1,02E-08 | -0,801027323 | 0,243 | 0,24 | 0,000121524 | 3 |
| *CG31324.2* | 1,17E-08 | 0,646469084 | 0,543 | 0,32 | 0,000139731 | 3 |
| *cpx.2* | 1,20E-08 | -1,066095292 | 0,171 | 0,226 | 0,000142889 | 3 |
| *Tis11.1* | 1,20E-08 | 0,31159179 | 0,938 | 0,78 | 0,000143178 | 3 |
| *SCaMC.2* | 1,30E-08 | 0,368790725 | 0,438 | 0,258 | 0,000154849 | 3 |
| *Sik3* | 1,67E-08 | 0,376075208 | 0,381 | 0,225 | 0,000198718 | 3 |
| *Sox14* | 1,76E-08 | 0,478436311 | 0,281 | 0,142 | 0,000210019 | 3 |
| *sqd.1* | 1,82E-08 | -0,338752782 | 0,638 | 0,57 | 0,000217616 | 3 |
| *ced-6.1* | 1,84E-08 | 0,373435496 | 0,652 | 0,448 | 0,000219852 | 3 |
| *Khc-73.1* | 1,96E-08 | 0,34213538 | 0,452 | 0,283 | 0,000233617 | 3 |
| *Cip4.1* | 2,08E-08 | -0,73901957 | 0,529 | 0,533 | 0,000248423 | 3 |
| *CG31689.1* | 2,21E-08 | 0,512482986 | 0,629 | 0,413 | 0,000263986 | 3 |
| *Naprt.2* | 2,21E-08 | 0,334169019 | 0,29 | 0,165 | 0,000264065 | 3 |
| *gogo.1* | 2,26E-08 | 0,623079095 | 0,348 | 0,178 | 0,000270168 | 3 |
| *g.2* | 2,47E-08 | 0,480548386 | 0,314 | 0,158 | 0,000295087 | 3 |
| *Jupiter* | 2,73E-08 | 0,284955352 | 0,962 | 0,819 | 0,000325886 | 3 |
| *RpL28.1* | 2,76E-08 | -0,484008636 | 0,371 | 0,328 | 0,00032969 | 3 |
| *Cyp6v1* | 2,76E-08 | 0,302721799 | 0,233 | 0,127 | 0,000329738 | 3 |
| *CG5151.2* | 2,83E-08 | -0,519198062 | 0,81 | 0,794 | 0,000338096 | 3 |
| *Fas3.2* | 2,98E-08 | -2,084238733 | 0,095 | 0,244 | 0,000355079 | 3 |
| *RpL27A.1* | 3,11E-08 | -0,936221333 | 0,319 | 0,389 | 0,000370751 | 3 |
| *Plp.1* | 3,18E-08 | -0,27181222 | 0,252 | 0,209 | 0,000378822 | 3 |
| *RpL10Ab.1* | 3,93E-08 | -0,530402756 | 0,281 | 0,256 | 0,000468672 | 3 |
| *14-3-3zeta* | 4,03E-08 | -0,353735239 | 0,39 | 0,362 | 0,000480448 | 3 |
| *RpS17.1* | 4,05E-08 | -0,566154032 | 0,243 | 0,213 | 0,000482707 | 3 |
| *Dscam1.1* | 4,89E-08 | -0,39569565 | 0,29 | 0,253 | 0,00058351 | 3 |
| *mnb.2* | 5,68E-08 | 0,622393756 | 0,581 | 0,36 | 0,000677318 | 3 |
| *Sap-r.1* | 5,73E-08 | 0,46000899 | 0,614 | 0,403 | 0,0006836 | 3 |
| *drpr.1* | 5,73E-08 | 0,38993865 | 0,886 | 0,729 | 0,000684018 | 3 |
| *pigs.2* | 6,21E-08 | -0,269236986 | 0,652 | 0,57 | 0,000741248 | 3 |
| *Prps.2* | 6,66E-08 | 0,387119495 | 0,514 | 0,321 | 0,000794501 | 3 |
| *Mondo.1* | 7,30E-08 | 0,250336447 | 0,305 | 0,183 | 0,000870234 | 3 |
| *scrib* | 7,61E-08 | -0,705655212 | 0,257 | 0,319 | 0,0009075 | 3 |
| *Fer1HCH.1* | 8,47E-08 | 0,293939081 | 0,705 | 0,509 | 0,001010453 | 3 |
| *alc.1* | 8,70E-08 | 0,349162526 | 0,414 | 0,256 | 0,001037285 | 3 |
| *CG1637* | 8,90E-08 | 0,66530987 | 0,31 | 0,146 | 0,00106171 | 3 |
| *rhea.1* | 8,99E-08 | -0,283865877 | 0,424 | 0,36 | 0,001071832 | 3 |
| *AdSS.2* | 9,10E-08 | 0,710755943 | 0,333 | 0,164 | 0,00108572 | 3 |
| *CycG.1* | 9,66E-08 | 0,376727738 | 0,886 | 0,72 | 0,001152791 | 3 |
| *Svil.1* | 9,91E-08 | -0,578130857 | 0,424 | 0,432 | 0,001182479 | 3 |
| *step.1* | 1,01E-07 | 0,434981172 | 0,595 | 0,392 | 0,001201962 | 3 |
| *cno.2* | 1,05E-07 | 0,569723464 | 0,524 | 0,318 | 0,001256844 | 3 |
| *GstD1.1* | 1,13E-07 | -0,678042302 | 0,305 | 0,276 | 0,001342606 | 3 |

| | | | | | | |
|---|---|---|---|---|---|---|
| *bip2* | 1,14E-07 | 0,414509562 | 0,59 | 0,398 | 0,001362813 | 3 |
| *RpS7.1* | 1,21E-07 | -0,824659573 | 0,252 | 0,308 | 0,001441827 | 3 |
| *RpL31* | 1,24E-07 | -0,659107482 | 0,233 | 0,258 | 0,001473721 | 3 |
| *Rab7.2* | 1,24E-07 | 0,393586754 | 0,238 | 0,116 | 0,001477537 | 3 |
| *sta* | 1,33E-07 | -0,590671466 | 0,267 | 0,277 | 0,001589844 | 3 |
| *spin.1* | 1,36E-07 | 0,267709939 | 0,557 | 0,38 | 0,001627272 | 3 |
| *RpS25* | 1,42E-07 | -0,527509482 | 0,248 | 0,233 | 0,001688011 | 3 |
| *mt:CoI.1* | 1,42E-07 | -0,740829073 | 0,171 | 0,202 | 0,001698856 | 3 |
| *scyl* | 1,52E-07 | 0,257798356 | 0,676 | 0,485 | 0,001808865 | 3 |
| *bwa.2* | 1,56E-07 | 0,695129008 | 0,286 | 0,134 | 0,001858693 | 3 |
| *Pxn.2* | 1,58E-07 | -0,669537258 | 0,514 | 0,548 | 0,001888616 | 3 |
| *RpL32.1* | 1,60E-07 | -0,851860973 | 0,233 | 0,291 | 0,001906776 | 3 |
| *bin3* | 1,61E-07 | 0,272371084 | 0,3 | 0,178 | 0,001919596 | 3 |
| *Mapmodulin.2* | 1,74E-07 | 0,286289595 | 0,286 | 0,164 | 0,002070732 | 3 |
| *vir-1.2* | 1,90E-07 | -0,805144702 | 0,324 | 0,302 | 0,00226278 | 3 |
| *Su(z)2* | 1,97E-07 | 0,498668076 | 0,143 | 0,054 | 0,002353109 | 3 |
| *CG30046* | 2,00E-07 | 0,361917918 | 0,281 | 0,161 | 0,002383386 | 3 |
| *CG13907* | 2,01E-07 | 0,298592823 | 0,314 | 0,195 | 0,002401174 | 3 |
| *fax.1* | 2,02E-07 | 0,33717329 | 0,776 | 0,585 | 0,002405951 | 3 |
| *jim.2* | 2,13E-07 | 0,542289033 | 0,633 | 0,423 | 0,002539232 | 3 |
| *RhoGAP71E.1* | 2,15E-07 | -0,447852935 | 0,557 | 0,554 | 0,002570632 | 3 |
| *CG42542* | 2,38E-07 | 0,4786366 | 0,429 | 0,249 | 0,002845054 | 3 |
| *ABCD* | 2,56E-07 | 0,32650145 | 0,357 | 0,215 | 0,003053469 | 3 |
| *CG2233* | 2,59E-07 | -0,962829874 | 0,014 | 0,117 | 0,00308763 | 3 |
| *Arc1.1* | 2,95E-07 | -1,158327713 | 0,129 | 0,105 | 0,003513686 | 3 |
| *RhoGAP15B.1* | 3,11E-07 | 0,412188795 | 0,338 | 0,189 | 0,003705783 | 3 |
| *unk.2* | 3,21E-07 | 0,351699302 | 0,333 | 0,198 | 0,003826573 | 3 |
| *CDase* | 3,29E-07 | 0,435904002 | 0,119 | 0,041 | 0,003921575 | 3 |
| *Paip2.2* | 3,37E-07 | 0,572028661 | 0,386 | 0,213 | 0,004024608 | 3 |
| *RpL24* | 3,52E-07 | -0,781852244 | 0,205 | 0,284 | 0,004203981 | 3 |
| *mgl.1* | 3,77E-07 | -1,338008859 | 0,219 | 0,335 | 0,004500938 | 3 |
| *RpS18.1* | 3,99E-07 | -0,339401166 | 0,367 | 0,306 | 0,004762964 | 3 |
| *l(3)L1231.1* | 4,48E-07 | 0,329946637 | 0,867 | 0,702 | 0,005344374 | 3 |
| *RpL23* | 4,50E-07 | -0,561894308 | 0,324 | 0,332 | 0,005368458 | 3 |
| *CG18135.2* | 4,64E-07 | 0,617394661 | 0,514 | 0,318 | 0,005533936 | 3 |
| *RpS9.1* | 4,67E-07 | -0,353985176 | 0,29 | 0,239 | 0,00556507 | 3 |
| *RpS28b.1* | 4,70E-07 | -0,446749364 | 0,229 | 0,224 | 0,005605798 | 3 |
| *RpS8.1* | 5,01E-07 | -0,405661031 | 0,376 | 0,321 | 0,005980835 | 3 |
| *CG42524.2* | 5,02E-07 | -0,475024691 | 0,343 | 0,37 | 0,005984371 | 3 |
| *CG30089.2* | 5,24E-07 | 0,261249323 | 0,538 | 0,378 | 0,006248248 | 3 |
| *SREBP.2* | 5,93E-07 | 0,307792327 | 0,281 | 0,162 | 0,007069217 | 3 |
| *Drat.2* | 5,94E-07 | 0,310626576 | 0,305 | 0,175 | 0,007080032 | 3 |
| *Dmtn* | 6,31E-07 | 0,32781401 | 0,476 | 0,307 | 0,00753285 | 3 |
| *Mur2B* | 6,39E-07 | 0,256812393 | 0,643 | 0,461 | 0,007618884 | 3 |
| *RpL23A* | 6,90E-07 | -0,485206057 | 0,19 | 0,172 | 0,008233482 | 3 |
| *CHES-1-like.2* | 7,96E-07 | 0,304930364 | 0,505 | 0,35 | 0,009492422 | 3 |
| *CG15309* | 8,33E-07 | 0,316732491 | 0,21 | 0,112 | 0,009942661 | 3 |
| *CG34331* | 8,34E-07 | 0,272484699 | 0,214 | 0,115 | 0,009951239 | 3 |
| *lap.1* | 9,65E-07 | 0,291217242 | 0,571 | 0,406 | 0,011514374 | 3 |
| *Act5C.2* | 1,05E-06 | -0,718492463 | 0,319 | 0,373 | 0,012537271 | 3 |
| *Haspin.1* | 1,08E-06 | -0,263881478 | 0,252 | 0,227 | 0,012897317 | 3 |
| *wake.2* | 1,16E-06 | -0,717902304 | 0,395 | 0,44 | 0,013823409 | 3 |
| *CG17754.1* | 1,22E-06 | 0,352825258 | 0,252 | 0,147 | 0,0145409 | 3 |
| *Idh.1* | 1,32E-06 | 0,470714938 | 0,257 | 0,127 | 0,015770237 | 3 |
| *MCPH1.1* | 1,44E-06 | 0,299773997 | 0,248 | 0,139 | 0,017154003 | 3 |
| *CG1578.1* | 1,50E-06 | 0,303397703 | 0,395 | 0,25 | 0,017909168 | 3 |
| *Vha100-2.2* | 1,63E-06 | -0,290233275 | 0,243 | 0,221 | 0,019473197 | 3 |
| *CG4080* | 1,73E-06 | 0,453273173 | 0,643 | 0,455 | 0,020685573 | 3 |
| *spg.2* | 1,78E-06 | 0,279329205 | 0,224 | 0,127 | 0,021236808 | 3 |
| *slpr.1* | 1,79E-06 | 0,482027791 | 0,229 | 0,114 | 0,021391307 | 3 |
| *CG4911* | 1,81E-06 | 0,308791239 | 0,238 | 0,133 | 0,021551724 | 3 |

| | | | | | | |
|---|---|---|---|---|---|---|
| *Hipk* | 1,97E-06 | 0,341370287 | 0,714 | 0,526 | 0,023515042 | 3 |
| *Btk29A.2* | 2,01E-06 | 0,30851103 | 0,89 | 0,752 | 0,023925959 | 3 |
| *CG5080.2* | 2,17E-06 | 0,529087035 | 0,676 | 0,492 | 0,025842777 | 3 |
| *Drak.2* | 2,20E-06 | -0,481048868 | 0,495 | 0,51 | 0,026269132 | 3 |
| *galla-1.1* | 2,28E-06 | 0,319554021 | 0,2 | 0,101 | 0,027192668 | 3 |
| *hzg.1* | 2,38E-06 | -0,369598537 | 0,176 | 0,168 | 0,028357972 | 3 |
| *Ald1.1* | 2,55E-06 | -0,509622021 | 0,352 | 0,333 | 0,030423364 | 3 |
| *Gdh.1* | 2,62E-06 | 0,448072268 | 0,176 | 0,069 | 0,031216061 | 3 |
| *Vha100-1* | 2,63E-06 | 0,253451677 | 0,2 | 0,111 | 0,03136123 | 3 |
| *RpL34b* | 2,76E-06 | -0,577258432 | 0,21 | 0,219 | 0,032889168 | 3 |
| *px.1* | 2,89E-06 | 0,25084561 | 0,9 | 0,753 | 0,034481171 | 3 |
| *nmo* | 2,89E-06 | -0,269589399 | 0,305 | 0,256 | 0,034530402 | 3 |
| *CG10082.1* | 3,14E-06 | 0,375694724 | 0,41 | 0,254 | 0,037503277 | 3 |
| *spir.1* | 3,16E-06 | -0,411063913 | 0,495 | 0,447 | 0,037666398 | 3 |
| *cdi.1* | 3,24E-06 | -0,506935839 | 0,3 | 0,3 | 0,038691618 | 3 |
| *CG9701.2* | 3,39E-06 | 0,77095913 | 0,2 | 0,083 | 0,040464799 | 3 |
| *sm* | 3,46E-06 | -0,335147193 | 0,276 | 0,236 | 0,041224208 | 3 |
| *Drs* | 3,86E-06 | -1,05291109 | 0,095 | 0,12 | 0,046011558 | 3 |
| *CG2201* | 3,96E-06 | -0,454333141 | 0,124 | 0,133 | 0,047226743 | 3 |
| *COX6B* | 3,99E-06 | -0,365549399 | 0,129 | 0,117 | 0,047582435 | 3 |
| *Dyrk2.1* | 4,05E-06 | 0,451670796 | 0,719 | 0,536 | 0,048321063 | 3 |
| *sky.1* | 4,10E-06 | 0,276816346 | 0,786 | 0,62 | 0,048867861 | 3 |
| *flw.2* | 4,15E-06 | -0,69729512 | 0,319 | 0,364 | 0,049510253 | 3 |
| *eEF1alpha1.2* | 4,87E-15 | 0,936512974 | 0,582 | 0,512 | 5,81E-11 | 4 |
| *Mst84Da.2* | 2,47E-14 | 1,401788323 | 0,328 | 0,132 | 2,95E-10 | 4 |
| *CG2233.1* | 1,04E-13 | 1,709212823 | 0,269 | 0,072 | 1,24E-09 | 4 |
| *Ppn.1* | 1,75E-13 | 0,573812859 | 1 | 0,96 | 2,09E-09 | 4 |
| *RpL31.1* | 7,31E-13 | 1,026028645 | 0,353 | 0,237 | 8,72E-09 | 4 |
| *RpS25.1* | 8,20E-12 | 1,230388253 | 0,398 | 0,207 | 9,78E-08 | 4 |
| *RpS27.2* | 8,96E-12 | 1,046730952 | 0,433 | 0,291 | 1,07E-07 | 4 |
| *Inos* | 2,30E-11 | 0,934065384 | 0,647 | 0,496 | 2,74E-07 | 4 |
| *RpL35A.2* | 3,19E-11 | 1,065097098 | 0,348 | 0,197 | 3,80E-07 | 4 |
| *RpL10Ab.2* | 3,99E-11 | 1,041831409 | 0,398 | 0,236 | 4,76E-07 | 4 |
| *RpL8.1* | 1,17E-10 | 1,002209877 | 0,388 | 0,25 | 1,40E-06 | 4 |
| *RpS13* | 2,40E-10 | 1,003557742 | 0,343 | 0,223 | 2,86E-06 | 4 |
| *Eip93F* | 2,60E-10 | -0,418309412 | 0,985 | 0,994 | 3,10E-06 | 4 |
| *Obp99c.1* | 5,75E-10 | 1,191257836 | 0,219 | 0,101 | 6,86E-06 | 4 |
| *eEF2.3* | 1,36E-09 | 0,86496703 | 0,438 | 0,293 | 1,62E-05 | 4 |
| *RpL21.3* | 1,76E-09 | 0,980992585 | 0,373 | 0,258 | 2,10E-05 | 4 |
| *InR.3* | 1,90E-09 | -0,753364691 | 0,786 | 0,865 | 2,26E-05 | 4 |
| *cathD* | 1,93E-09 | 0,94248068 | 0,284 | 0,206 | 2,30E-05 | 4 |
| *RpL28.2* | 2,10E-09 | 0,848639585 | 0,393 | 0,325 | 2,51E-05 | 4 |
| *lilli* | 4,84E-09 | -0,40183275 | 0,522 | 0,745 | 5,77E-05 | 4 |
| *CG42788.1* | 5,48E-09 | 0,560930526 | 0,249 | 0,273 | 6,53E-05 | 4 |
| *RpL36.1* | 8,43E-09 | 0,833488732 | 0,338 | 0,27 | 0,000100505 | 4 |
| *RpL27A.2* | 9,82E-09 | 0,951674394 | 0,488 | 0,359 | 0,000117198 | 4 |
| *Desat1* | 1,10E-08 | 1,2592209 | 0,299 | 0,123 | 0,000131004 | 4 |
| *CG14245.2* | 1,28E-08 | 1,488610459 | 0,184 | 0,122 | 0,00015314 | 4 |
| *Cyp4g1* | 1,54E-08 | 1,601959274 | 0,388 | 0,216 | 0,000183888 | 4 |
| *mt:CoI.2* | 1,98E-08 | 0,89575401 | 0,269 | 0,185 | 0,000235964 | 4 |
| *RpL7.1* | 2,21E-08 | 0,787699614 | 0,284 | 0,237 | 0,000263771 | 4 |
| *RpL14.1* | 3,07E-08 | 0,950569662 | 0,393 | 0,293 | 0,000365762 | 4 |
| *RpL26.1* | 3,63E-08 | 0,825926687 | 0,358 | 0,258 | 0,000432496 | 4 |
| *RpL11.1* | 3,82E-08 | 0,915399562 | 0,284 | 0,181 | 0,000455449 | 4 |
| *RpS18.2* | 3,85E-08 | 0,835421003 | 0,418 | 0,298 | 0,000459092 | 4 |
| *RpLP2.1* | 4,09E-08 | 0,933668705 | 0,299 | 0,185 | 0,000487371 | 4 |
| *RpS17.2* | 4,53E-08 | 0,956449931 | 0,274 | 0,208 | 0,000540378 | 4 |
| *RpS4.1* | 4,57E-08 | 0,930326548 | 0,299 | 0,194 | 0,000544654 | 4 |
| *RpL32.2* | 5,43E-08 | 0,800309995 | 0,353 | 0,27 | 0,000648259 | 4 |
| *RpS28b.2* | 5,75E-08 | 0,78926605 | 0,294 | 0,213 | 0,000685838 | 4 |
| *RpS9.2* | 6,47E-08 | 0,866319937 | 0,353 | 0,229 | 0,000771755 | 4 |

| | | | | | | |
|---|---|---|---|---|---|---|
| *Mst57Db.1* | 6,94E-08 | 0,975357685 | 0,239 | 0,132 | 0,000827384 | 4 |
| *RpL37A.2* | 7,45E-08 | 0,905939483 | 0,264 | 0,183 | 0,000888746 | 4 |
| *CG7766.1* | 1,00E-07 | 0,461148894 | 0,159 | 0,151 | 0,001192983 | 4 |
| *CG16758.2* | 1,37E-07 | 0,965261139 | 0,224 | 0,083 | 0,001632759 | 4 |
| *CG3662* | 1,43E-07 | 0,469142021 | 0,348 | 0,342 | 0,001709552 | 4 |
| *RpL15.2* | 1,48E-07 | 0,782977793 | 0,383 | 0,301 | 0,001764137 | 4 |
| *CG3124* | 1,59E-07 | 0,900602917 | 0,134 | 0,041 | 0,001891063 | 4 |
| *RpL35.2* | 1,66E-07 | 0,885003988 | 0,303 | 0,207 | 0,001985999 | 4 |
| *awd.1* | 1,80E-07 | 0,749329366 | 0,154 | 0,11 | 0,002148476 | 4 |
| *CG31226* | 1,89E-07 | 1,036031632 | 0,154 | 0,07 | 0,002250025 | 4 |
| *CG46059* | 2,00E-07 | 0,924620677 | 0,129 | 0,058 | 0,002381479 | 4 |
| *lcs.1* | 2,00E-07 | 1,106962608 | 0,299 | 0,179 | 0,002389681 | 4 |
| *RpL27.2* | 2,08E-07 | 0,822916658 | 0,323 | 0,255 | 0,00247693 | 4 |
| *hppy.1* | 2,32E-07 | -0,44001817 | 0,418 | 0,612 | 0,002771563 | 4 |
| *RpS10b.1* | 2,64E-07 | 0,926697541 | 0,358 | 0,225 | 0,003153123 | 4 |
| *RpS29.1* | 2,79E-07 | 0,95724928 | 0,303 | 0,187 | 0,003333555 | 4 |
| *Got2.1* | 2,89E-07 | 0,854938545 | 0,149 | 0,081 | 0,00345061 | 4 |
| *CG1648.2* | 3,17E-07 | 1,331856316 | 0,144 | 0,131 | 0,003784458 | 4 |
| *Fas3.3* | 3,23E-07 | -1,91053723 | 0,09 | 0,243 | 0,003857505 | 4 |
| *CG6503.1* | 3,76E-07 | 0,603863348 | 0,129 | 0,03 | 0,004480998 | 4 |
| *NimB2* | 4,77E-07 | 0,583198471 | 0,274 | 0,236 | 0,005685198 | 4 |
| *RpL41* | 4,88E-07 | 0,862004489 | 0,428 | 0,266 | 0,005815804 | 4 |
| *RpS21.2* | 4,91E-07 | 0,955580163 | 0,373 | 0,229 | 0,005855867 | 4 |
| *CG42674.2* | 5,44E-07 | -1,260180019 | 0,025 | 0,137 | 0,006493488 | 4 |
| *RpLP1.2* | 6,10E-07 | 0,957366302 | 0,338 | 0,221 | 0,007274115 | 4 |
| *CG5080.3* | 7,81E-07 | 0,647080378 | 0,567 | 0,512 | 0,009312511 | 4 |
| *GEFmeso.2* | 8,33E-07 | -0,728258227 | 0,279 | 0,463 | 0,00994009 | 4 |
| *RpL13* | 9,82E-07 | 0,734919036 | 0,313 | 0,24 | 0,01171055 | 4 |
| *SPARC.1* | 1,18E-06 | 0,539156077 | 0,612 | 0,504 | 0,014040481 | 4 |
| *RpL39* | 1,33E-06 | 0,905872766 | 0,299 | 0,21 | 0,015816014 | 4 |
| *lectin-24Db.2* | 1,34E-06 | 0,590311077 | 0,154 | 0,13 | 0,01602226 | 4 |
| *RpS2* | 1,39E-06 | 0,78599725 | 0,294 | 0,179 | 0,016555312 | 4 |
| *spir.2* | 1,44E-06 | -0,927837057 | 0,289 | 0,483 | 0,017204582 | 4 |
| *RpS30.2* | 1,45E-06 | 0,856448823 | 0,333 | 0,218 | 0,01725427 | 4 |
| *hid.2* | 1,80E-06 | -0,816190021 | 0,219 | 0,398 | 0,021421529 | 4 |
| *RpS3.1* | 1,80E-06 | 0,805111844 | 0,264 | 0,176 | 0,021473939 | 4 |
| *ocn* | 1,81E-06 | 0,958231089 | 0,129 | 0,059 | 0,021629814 | 4 |
| *ATPsynD* | 1,88E-06 | 0,745081338 | 0,129 | 0,069 | 0,022391308 | 4 |
| *BG642312* | 2,08E-06 | 1,00667529 | 0,139 | 0,061 | 0,024801509 | 4 |
| *bun.2* | 2,21E-06 | -0,543297859 | 0,741 | 0,856 | 0,026411749 | 4 |
| *Mob2.3* | 2,33E-06 | -0,561323333 | 0,736 | 0,857 | 0,027737766 | 4 |
| *Fim.1* | 2,34E-06 | -0,785863754 | 0,219 | 0,393 | 0,027865876 | 4 |
| *prage.3* | 2,44E-06 | -1,069539974 | 0,149 | 0,316 | 0,029140816 | 4 |
| *ACC.1* | 2,45E-06 | 0,906031095 | 0,189 | 0,128 | 0,029275228 | 4 |
| *RpL3.2* | 2,60E-06 | 0,720432257 | 0,328 | 0,25 | 0,030968562 | 4 |
| *msi.1* | 2,80E-06 | -0,455198302 | 0,786 | 0,866 | 0,033454938 | 4 |
| *eEF1beta* | 2,99E-06 | 0,781371077 | 0,189 | 0,115 | 0,035638356 | 4 |
| *Hml.3* | 3,02E-06 | 0,490049497 | 0,776 | 0,629 | 0,036082622 | 4 |
| *MtnA.2* | 3,15E-06 | 0,802305348 | 0,388 | 0,278 | 0,037541547 | 4 |
| *Ance-5.1* | 3,23E-06 | 0,67878823 | 0,204 | 0,142 | 0,038561644 | 4 |
| *Diap1* | 3,57E-06 | -0,673537839 | 0,308 | 0,491 | 0,04254256 | 4 |
| *RpL7A* | 3,64E-06 | 0,619031393 | 0,323 | 0,261 | 0,043446559 | 4 |
| *Abl* | 3,76E-06 | -0,642103574 | 0,542 | 0,718 | 0,044882442 | 4 |
| *CG12581* | 4,11E-64 | 2,820735668 | 0,449 | 0,018 | 4,90E-60 | 5 |
| *CG17839.4* | 2,68E-58 | 3,196780833 | 0,52 | 0,05 | 3,20E-54 | 5 |
| *dpy* | 2,73E-38 | 3,222347954 | 0,313 | 0,015 | 3,25E-34 | 5 |
| *grh* | 9,70E-38 | 2,310710282 | 0,313 | 0,018 | 1,16E-33 | 5 |
| *mgl.3* | 2,11E-37 | 2,202476509 | 0,667 | 0,258 | 2,51E-33 | 5 |
| *Fas2* | 4,54E-30 | 2,243656548 | 0,308 | 0,031 | 5,41E-26 | 5 |
| *CG8740.2* | 9,09E-30 | 1,987496589 | 0,348 | 0,045 | 1,08E-25 | 5 |
| *jbug* | 2,32E-28 | 1,944398065 | 0,212 | 0,006 | 2,77E-24 | 5 |

| | | | | | | |
|---|---|---|---|---|---|---|
| *hth.1* | 4,79E-27 | 2,356175316 | 0,323 | 0,051 | 5,72E-23 | 5 |
| *pyd.3* | 8,07E-26 | 2,151956151 | 0,333 | 0,059 | 9,62E-22 | 5 |
| *Tie* | 3,04E-25 | 1,970866864 | 0,217 | 0,018 | 3,62E-21 | 5 |
| *ELOVL* | 6,37E-25 | 2,490571738 | 0,202 | 0,009 | 7,60E-21 | 5 |
| *CG11073* | 8,20E-25 | 1,621623962 | 0,187 | 0,008 | 9,78E-21 | 5 |
| *Cda5* | 1,23E-24 | 1,8891144 | 0,207 | 0,012 | 1,47E-20 | 5 |
| *form3* | 1,80E-24 | 2,100592589 | 0,212 | 0,011 | 2,15E-20 | 5 |
| *ed.3* | 7,62E-24 | 1,707170905 | 0,505 | 0,166 | 9,09E-20 | 5 |
| *CG8180* | 1,56E-23 | 1,959366841 | 0,192 | 0,007 | 1,86E-19 | 5 |
| *siz.2* | 7,35E-23 | 1,614571572 | 0,424 | 0,112 | 8,76E-19 | 5 |
| *Dys.4* | 1,11E-22 | 1,68648377 | 0,439 | 0,124 | 1,32E-18 | 5 |
| *Fas3.4* | 1,40E-21 | 0,740040422 | 0,51 | 0,172 | 1,67E-17 | 5 |
| *crb* | 1,92E-21 | 1,861254103 | 0,247 | 0,029 | 2,30E-17 | 5 |
| *pdm3* | 7,41E-21 | 1,989308548 | 0,172 | 0,009 | 8,84E-17 | 5 |
| *vvl* | 5,18E-20 | 1,648844527 | 0,227 | 0,027 | 6,18E-16 | 5 |
| *CG33970* | 7,52E-20 | 2,47638364 | 0,177 | 0,009 | 8,97E-16 | 5 |
| *CG30460* | 8,12E-20 | 1,798936202 | 0,232 | 0,031 | 9,69E-16 | 5 |
| *CG6040* | 2,38E-19 | 1,637680787 | 0,227 | 0,027 | 2,84E-15 | 5 |
| *snu* | 6,47E-19 | 1,568963489 | 0,202 | 0,027 | 7,71E-15 | 5 |
| *CG6959* | 9,00E-19 | 1,464014854 | 0,237 | 0,036 | 1,07E-14 | 5 |
| *Nost* | 9,12E-19 | 1,681069502 | 0,258 | 0,039 | 1,09E-14 | 5 |
| *CG34398* | 1,38E-18 | 2,009958074 | 0,187 | 0,015 | 1,65E-14 | 5 |
| *ASPP* | 1,71E-18 | 1,550090348 | 0,187 | 0,015 | 2,04E-14 | 5 |
| *CG34120* | 1,73E-18 | 2,117890915 | 0,177 | 0,015 | 2,06E-14 | 5 |
| *CG33110* | 1,84E-18 | 1,828932137 | 0,167 | 0,009 | 2,19E-14 | 5 |
| *RhoGEF64C* | 1,01E-17 | 1,86715077 | 0,146 | 0,006 | 1,20E-13 | 5 |
| *stw* | 1,39E-17 | 1,678946777 | 0,197 | 0,022 | 1,66E-13 | 5 |
| *CG32521.1* | 2,48E-17 | 1,669529396 | 0,308 | 0,075 | 2,95E-13 | 5 |
| *Lmpt.3* | 4,80E-17 | 1,784518672 | 0,323 | 0,081 | 5,73E-13 | 5 |
| *CG12814* | 1,72E-16 | 1,528915655 | 0,187 | 0,018 | 2,05E-12 | 5 |
| *sick.2* | 2,07E-16 | 1,771741696 | 0,298 | 0,07 | 2,47E-12 | 5 |
| *Hpd* | 2,19E-16 | 1,917124966 | 0,172 | 0,019 | 2,62E-12 | 5 |
| *CG9628* | 2,82E-16 | 1,311581215 | 0,192 | 0,026 | 3,36E-12 | 5 |
| *bru2* | 2,84E-16 | 1,58655365 | 0,237 | 0,045 | 3,38E-12 | 5 |
| *Wdr62.1* | 5,03E-16 | 1,055578991 | 0,631 | 0,414 | 6,00E-12 | 5 |
| *CG4928* | 6,79E-16 | 1,950232456 | 0,222 | 0,06 | 8,10E-12 | 5 |
| *nahoda* | 8,30E-16 | 1,32407829 | 0,131 | 0,007 | 9,90E-12 | 5 |
| *CG32694* | 1,38E-15 | 1,662928245 | 0,141 | 0,013 | 1,64E-11 | 5 |
| *CG3655* | 1,54E-15 | 1,526406363 | 0,187 | 0,022 | 1,83E-11 | 5 |
| *ovo* | 2,11E-15 | 1,427456673 | 0,131 | 0,007 | 2,51E-11 | 5 |
| *CG41520.1* | 2,31E-15 | 1,603843767 | 0,424 | 0,175 | 2,75E-11 | 5 |
| *sv* | 1,13E-14 | 1,82047943 | 0,116 | 0,006 | 1,34E-10 | 5 |
| *toc.4* | 1,72E-14 | 1,506517537 | 0,268 | 0,069 | 2,05E-10 | 5 |
| *rdgA.1* | 2,10E-14 | 1,263252265 | 0,359 | 0,119 | 2,51E-10 | 5 |
| *CG32137* | 2,59E-14 | 1,340054332 | 0,177 | 0,021 | 3,09E-10 | 5 |
| *CG43759* | 4,04E-14 | 1,490443879 | 0,146 | 0,015 | 4,81E-10 | 5 |
| *sprt* | 4,20E-14 | 1,380496503 | 0,146 | 0,013 | 5,01E-10 | 5 |
| *Best2* | 4,97E-14 | 1,130914971 | 0,157 | 0,017 | 5,92E-10 | 5 |
| *Msr-110* | 5,15E-14 | 1,942179981 | 0,227 | 0,061 | 6,15E-10 | 5 |
| *bi* | 7,70E-14 | 1,910417498 | 0,106 | 0,004 | 9,19E-10 | 5 |
| *Ets98B* | 7,97E-14 | 1,261886994 | 0,162 | 0,019 | 9,51E-10 | 5 |
| *dnr1* | 1,01E-13 | 1,703237066 | 0,182 | 0,033 | 1,20E-09 | 5 |
| *CG2841* | 1,14E-13 | 1,512236628 | 0,141 | 0,013 | 1,36E-09 | 5 |
| *Kank* | 1,24E-13 | 1,380862294 | 0,167 | 0,025 | 1,48E-09 | 5 |
| *sdt* | 1,86E-13 | 1,348767019 | 0,242 | 0,052 | 2,22E-09 | 5 |
| *CG31176* | 4,56E-13 | 1,980790717 | 0,121 | 0,009 | 5,44E-09 | 5 |
| *Gs2.1* | 6,60E-13 | 1,369576129 | 0,348 | 0,124 | 7,88E-09 | 5 |
| *Ptp10D* | 2,14E-12 | 1,366681661 | 0,232 | 0,055 | 2,55E-08 | 5 |
| *CG14830* | 2,32E-12 | 1,094748334 | 0,101 | 0,006 | 2,77E-08 | 5 |
| *pio* | 2,70E-12 | 1,187307427 | 0,116 | 0,009 | 3,22E-08 | 5 |
| *bowl* | 2,87E-12 | 1,08745391 | 0,121 | 0,009 | 3,42E-08 | 5 |

| | | | | | | |
|---|---|---|---|---|---|---|
| *Sp1* | 4,05E-12 | 1,246415514 | 0,106 | 0,007 | 4,84E-08 | 5 |
| *CG17834* | 4,64E-12 | 1,553799971 | 0,141 | 0,02 | 5,54E-08 | 5 |
| *Amph* | 1,60E-11 | 1,369946937 | 0,232 | 0,057 | 1,90E-07 | 5 |
| *CG6398* | 1,79E-11 | 1,158220388 | 0,126 | 0,014 | 2,14E-07 | 5 |
| *CG3168.1* | 2,05E-11 | 1,324109091 | 0,253 | 0,084 | 2,44E-07 | 5 |
| *osp* | 2,73E-11 | 1,259990747 | 0,212 | 0,052 | 3,25E-07 | 5 |
| *CG6409.1* | 5,27E-11 | 1,289225968 | 0,192 | 0,077 | 6,29E-07 | 5 |
| *caps.3* | 6,65E-11 | 1,126163554 | 0,263 | 0,076 | 7,93E-07 | 5 |
| *CG32264.3* | 1,14E-10 | 0,854978198 | 0,652 | 0,411 | 1,36E-06 | 5 |
| *Rbp6* | 1,61E-10 | 1,90262144 | 0,182 | 0,057 | 1,92E-06 | 5 |
| *kmr* | 1,79E-10 | 1,060546354 | 0,136 | 0,021 | 2,13E-06 | 5 |
| *sog* | 2,01E-10 | 1,35303971 | 0,187 | 0,039 | 2,40E-06 | 5 |
| *cpo.2* | 3,42E-10 | 0,735925897 | 0,833 | 0,679 | 4,08E-06 | 5 |
| *CG8312.1* | 5,79E-10 | 1,004215697 | 0,616 | 0,414 | 6,91E-06 | 5 |
| *Fas1* | 7,27E-10 | 0,926020008 | 0,182 | 0,039 | 8,67E-06 | 5 |
| *GstT4* | 1,40E-09 | 1,131293286 | 0,202 | 0,051 | 1,68E-05 | 5 |
| *CG42327* | 1,51E-09 | 1,148655012 | 0,141 | 0,024 | 1,81E-05 | 5 |
| *zormin* | 1,86E-09 | 1,016244893 | 0,136 | 0,024 | 2,22E-05 | 5 |
| *CG11147* | 2,09E-09 | 1,063824878 | 0,126 | 0,018 | 2,49E-05 | 5 |
| *cv-2.1* | 2,35E-09 | 1,733463687 | 0,207 | 0,065 | 2,80E-05 | 5 |
| *chas* | 2,49E-09 | 1,024964751 | 0,111 | 0,011 | 2,97E-05 | 5 |
| *bbg* | 3,82E-09 | 1,272227912 | 0,121 | 0,021 | 4,56E-05 | 5 |
| *scrib.1* | 4,60E-09 | 0,921579175 | 0,48 | 0,28 | 5,49E-05 | 5 |
| *CG42673* | 4,67E-09 | 1,348947893 | 0,111 | 0,015 | 5,57E-05 | 5 |
| *Nrg.4* | 6,95E-09 | 0,703995135 | 0,293 | 0,116 | 8,29E-05 | 5 |
| *Msp300.2* | 1,38E-08 | 0,923032504 | 0,697 | 0,55 | 0,000164154 | 5 |
| *nrm* | 1,41E-08 | 1,025077759 | 0,101 | 0,013 | 0,000167719 | 5 |
| *Pura* | 1,55E-08 | 1,06190429 | 0,106 | 0,014 | 0,000184436 | 5 |
| *CenG1A.1* | 2,10E-08 | 0,993542868 | 0,293 | 0,145 | 0,000250835 | 5 |
| *Ddc* | 2,23E-08 | 0,940642614 | 0,106 | 0,014 | 0,000265468 | 5 |
| *EbpIII* | 2,58E-08 | 1,134771014 | 0,177 | 0,07 | 0,000307952 | 5 |
| *fng* | 3,50E-08 | 1,083925652 | 0,101 | 0,016 | 0,00041797 | 5 |
| *CG7378* | 3,83E-08 | 1,130543457 | 0,116 | 0,019 | 0,000457385 | 5 |
| *csw* | 3,92E-08 | 0,971280149 | 0,394 | 0,234 | 0,000467264 | 5 |
| *bab2* | 5,09E-08 | 1,261697264 | 0,116 | 0,027 | 0,000607774 | 5 |
| *SPoCk* | 5,57E-08 | 1,068200327 | 0,172 | 0,057 | 0,000664679 | 5 |
| *sima* | 6,38E-08 | 0,568787721 | 0,859 | 0,78 | 0,00076144 | 5 |
| *Gp150* | 7,72E-08 | 0,853500306 | 0,187 | 0,061 | 0,000920972 | 5 |
| *emp* | 8,53E-08 | 0,966751704 | 0,131 | 0,033 | 0,001017768 | 5 |
| *pum.1* | 2,00E-07 | -0,339908236 | 0,99 | 0,995 | 0,002384149 | 5 |
| *fus* | 2,07E-07 | 0,747376236 | 0,141 | 0,035 | 0,002467823 | 5 |
| *Egfr* | 2,23E-07 | 0,983594981 | 0,182 | 0,053 | 0,002661975 | 5 |
| *Hex-A.1* | 2,50E-07 | 0,864695726 | 0,439 | 0,261 | 0,002984054 | 5 |
| *T48* | 2,70E-07 | 1,042520134 | 0,167 | 0,046 | 0,00322472 | 5 |
| *how.1* | 2,98E-07 | 1,015062793 | 0,434 | 0,257 | 0,003558093 | 5 |
| *plx* | 3,26E-07 | 0,924243454 | 0,121 | 0,026 | 0,003890956 | 5 |
| *dally* | 4,99E-07 | 1,339826318 | 0,136 | 0,04 | 0,005950141 | 5 |
| *sm.1* | 5,99E-07 | 0,984188743 | 0,359 | 0,222 | 0,007150274 | 5 |
| *Bx* | 7,94E-07 | 0,994245487 | 0,157 | 0,043 | 0,009467499 | 5 |
| *rols* | 1,23E-06 | 0,998404471 | 0,101 | 0,022 | 0,014654189 | 5 |
| *jing.2* | 1,33E-06 | 0,856601261 | 0,343 | 0,172 | 0,015885738 | 5 |
| *msn.1* | 2,20E-06 | -0,527843197 | 0,813 | 0,895 | 0,026276645 | 5 |
| *tna.3* | 2,48E-06 | 0,501800058 | 0,374 | 0,196 | 0,029554658 | 5 |
| *Eaat1* | 3,07E-06 | 0,831789715 | 0,111 | 0,028 | 0,036646938 | 5 |
| *ex* | 3,10E-06 | 0,873113255 | 0,172 | 0,067 | 0,036995504 | 5 |
| *ps* | 3,69E-06 | 0,338216164 | 0,98 | 0,913 | 0,044024018 | 5 |
| *Cip4.4* | 1,53E-79 | 2,311988719 | 0,89 | 0,481 | 1,83E-75 | 6 |
| *luna.3* | 4,60E-61 | 1,635969959 | 0,983 | 0,779 | 5,49E-57 | 6 |
| *spir.3* | 1,25E-60 | 2,172087296 | 0,919 | 0,388 | 1,50E-56 | 6 |
| *Myo31DF.4* | 2,54E-53 | 2,694650499 | 0,692 | 0,154 | 3,02E-49 | 6 |
| *mamo.1* | 4,08E-42 | -1,761354046 | 0,488 | 0,901 | 4,86E-38 | 6 |

| | | | | | | |
|---|---|---|---|---|---|---|
| *Myc.2* | 7,18E-42 | 1,942823141 | 0,767 | 0,423 | 8,56E-38 | 6 |
| *cher.3* | 1,42E-41 | 2,686033838 | 0,465 | 0,052 | 1,70E-37 | 6 |
| *Fas3.5* | 6,39E-41 | 3,153263902 | 0,576 | 0,169 | 7,63E-37 | 6 |
| *Act5C.3* | 1,60E-40 | 2,024191504 | 0,733 | 0,312 | 1,91E-36 | 6 |
| *AnxB9.3* | 4,91E-40 | 2,101255046 | 0,721 | 0,268 | 5,86E-36 | 6 |
| *MCU.3* | 1,11E-38 | 2,58288177 | 0,442 | 0,053 | 1,32E-34 | 6 |
| *Dscam1.4* | 3,02E-34 | 2,007325192 | 0,628 | 0,206 | 3,61E-30 | 6 |
| *Ppn.2* | 5,51E-32 | -1,174418807 | 0,884 | 0,977 | 6,57E-28 | 6 |
| *RhoGAP71E.3* | 1,19E-31 | 1,532628926 | 0,814 | 0,517 | 1,42E-27 | 6 |
| *fog.3* | 7,24E-31 | -1,799856927 | 0,262 | 0,697 | 8,63E-27 | 6 |
| *Col4a1.2* | 1,37E-30 | -1,103232545 | 0,797 | 0,943 | 1,63E-26 | 6 |
| *Pgant9.1* | 1,64E-30 | -1,67633168 | 0,349 | 0,756 | 1,96E-26 | 6 |
| *Spn.3* | 2,15E-30 | -1,82906011 | 0,302 | 0,726 | 2,56E-26 | 6 |
| *Btk29A.4* | 3,77E-30 | -1,627639208 | 0,453 | 0,82 | 4,50E-26 | 6 |
| *Naam.2* | 1,50E-29 | 2,083952506 | 0,587 | 0,203 | 1,79E-25 | 6 |
| *CG6891.1* | 7,78E-29 | 2,06918031 | 0,355 | 0,048 | 9,28E-25 | 6 |
| *Mmp2.4* | 4,24E-27 | 1,652042499 | 0,779 | 0,415 | 5,06E-23 | 6 |
| *CG12991.1* | 7,25E-27 | 2,092732408 | 0,343 | 0,043 | 8,65E-23 | 6 |
| *Pxn.3* | 8,63E-27 | -2,362552497 | 0,198 | 0,592 | 1,03E-22 | 6 |
| *GEFmeso.3* | 1,65E-26 | 1,430382634 | 0,75 | 0,39 | 1,97E-22 | 6 |
| *MYPT-75D.2* | 2,89E-26 | -2,134124054 | 0,18 | 0,592 | 3,45E-22 | 6 |
| *mys.2* | 5,01E-26 | 1,497545668 | 0,698 | 0,355 | 5,97E-22 | 6 |
| *CG3726.1* | 5,75E-26 | -1,561559912 | 0,279 | 0,693 | 6,86E-22 | 6 |
| *CG15673* | 6,29E-26 | 2,010981844 | 0,32 | 0,034 | 7,50E-22 | 6 |
| *Pvr.3* | 9,35E-26 | -1,069579314 | 0,663 | 0,909 | 1,12E-21 | 6 |
| *stac* | 2,67E-25 | 1,977013772 | 0,331 | 0,044 | 3,18E-21 | 6 |
| *aqz.1* | 2,80E-25 | -1,230774301 | 0,738 | 0,92 | 3,34E-21 | 6 |
| *Men-b.3* | 3,26E-25 | 2,00983158 | 0,378 | 0,063 | 3,89E-21 | 6 |
| *CG5399* | 7,03E-25 | 2,204576199 | 0,337 | 0,044 | 8,38E-21 | 6 |
| *px.2* | 7,41E-25 | -1,310978382 | 0,541 | 0,81 | 8,84E-21 | 6 |
| *mtd.2* | 8,33E-25 | -1,394092992 | 0,517 | 0,794 | 9,93E-21 | 6 |
| *Sema2a.3* | 1,38E-24 | -2,402163204 | 0,11 | 0,496 | 1,65E-20 | 6 |
| *rhea.3* | 1,61E-24 | 1,535077338 | 0,64 | 0,331 | 1,92E-20 | 6 |
| *Ae2.2* | 2,62E-24 | 1,957159601 | 0,494 | 0,158 | 3,13E-20 | 6 |
| *Pax.2* | 1,85E-23 | 1,704524125 | 0,541 | 0,185 | 2,20E-19 | 6 |
| *CG31145.2* | 1,06E-22 | -1,357797425 | 0,488 | 0,782 | 1,27E-18 | 6 |
| *Fim.3* | 1,16E-22 | 1,523974884 | 0,68 | 0,322 | 1,39E-18 | 6 |
| *loco.2* | 1,81E-22 | 2,007866929 | 0,39 | 0,086 | 2,15E-18 | 6 |
| *CG43867.2* | 1,92E-22 | 2,017918855 | 0,448 | 0,123 | 2,28E-18 | 6 |
| *Xrp1.2* | 2,33E-22 | 1,338928157 | 0,919 | 0,678 | 2,78E-18 | 6 |
| *schlank.2* | 4,32E-22 | 1,851069708 | 0,407 | 0,097 | 5,15E-18 | 6 |
| *Prosap.1* | 1,21E-21 | -0,590106749 | 0,808 | 0,966 | 1,45E-17 | 6 |
| *Inos.2* | 2,62E-21 | -1,819669962 | 0,186 | 0,566 | 3,12E-17 | 6 |
| *wge.3* | 6,65E-21 | 1,590089516 | 0,512 | 0,172 | 7,94E-17 | 6 |
| *pyr* | 9,66E-21 | 2,281959675 | 0,262 | 0,033 | 1,15E-16 | 6 |
| *sr.1* | 1,49E-20 | -1,66502435 | 0,25 | 0,576 | 1,78E-16 | 6 |
| *cic.1* | 2,03E-20 | -1,061478089 | 0,884 | 0,966 | 2,43E-16 | 6 |
| *vir-1.3* | 3,73E-20 | 1,916328329 | 0,581 | 0,266 | 4,45E-16 | 6 |
| *lola* | 5,86E-20 | -0,797333821 | 0,843 | 0,956 | 6,99E-16 | 6 |
| *CG40006.1* | 6,00E-20 | -1,257106806 | 0,308 | 0,663 | 7,15E-16 | 6 |
| *vkg.2* | 7,86E-20 | -1,169821686 | 0,57 | 0,845 | 9,38E-16 | 6 |
| *CG5004.3* | 7,98E-20 | 1,709094105 | 0,331 | 0,062 | 9,52E-16 | 6 |
| *Hml.4* | 1,05E-19 | -1,459276939 | 0,355 | 0,694 | 1,25E-15 | 6 |
| *TotM* | 4,15E-19 | 2,951844451 | 0,227 | 0,025 | 4,95E-15 | 6 |
| *fax.3* | 5,99E-19 | 1,141003835 | 0,843 | 0,581 | 7,14E-15 | 6 |
| *betaTub60D.4* | 6,20E-19 | 1,700812767 | 0,529 | 0,215 | 7,40E-15 | 6 |
| *Cam.2* | 9,25E-19 | 1,365393578 | 0,733 | 0,478 | 1,10E-14 | 6 |
| *scb.2* | 1,26E-18 | 1,661723428 | 0,413 | 0,112 | 1,51E-14 | 6 |
| *Socs36E.2* | 1,53E-18 | 1,808496009 | 0,448 | 0,211 | 1,83E-14 | 6 |
| *Gadd45.2* | 3,38E-18 | 1,899719155 | 0,395 | 0,108 | 4,03E-14 | 6 |
| *cdi.2* | 1,20E-17 | 1,35350464 | 0,576 | 0,26 | 1,43E-13 | 6 |

| | | | | | | |
|---|---|---|---|---|---|---|
| *gce.2* | 1,57E-17 | -1,451650325 | 0,262 | 0,583 | 1,87E-13 | 6 |
| *CD98hc.1* | 1,65E-17 | 1,534977643 | 0,297 | 0,06 | 1,97E-13 | 6 |
| *Sr-CI.2* | 1,76E-17 | -1,438404488 | 0,297 | 0,619 | 2,10E-13 | 6 |
| *Rtnl1.2* | 3,83E-17 | 1,57605777 | 0,349 | 0,087 | 4,57E-13 | 6 |
| *kek1.1* | 5,08E-17 | 1,926378557 | 0,471 | 0,212 | 6,06E-13 | 6 |
| *CG9328.1* | 5,47E-17 | -1,643720292 | 0,192 | 0,522 | 6,52E-13 | 6 |
| *CG17760.1* | 8,51E-17 | 1,817328714 | 0,297 | 0,064 | 1,01E-12 | 6 |
| *Eb1.2* | 1,05E-16 | 1,281315073 | 0,57 | 0,288 | 1,25E-12 | 6 |
| *CG32486.3* | 1,11E-16 | -1,366354391 | 0,273 | 0,579 | 1,32E-12 | 6 |
| *CG42674.4* | 2,54E-16 | 1,702589318 | 0,331 | 0,09 | 3,04E-12 | 6 |
| *Abl.2* | 2,63E-16 | 1,12286753 | 0,826 | 0,673 | 3,14E-12 | 6 |
| *CG1572* | 4,95E-16 | 1,466613231 | 0,291 | 0,057 | 5,90E-12 | 6 |
| *Ac76E.2* | 6,88E-16 | 1,482263244 | 0,424 | 0,144 | 8,20E-12 | 6 |
| *Hsp83.2* | 1,14E-15 | 1,566035243 | 0,663 | 0,369 | 1,36E-11 | 6 |
| *bbc.2* | 1,37E-15 | 1,46141217 | 0,471 | 0,19 | 1,63E-11 | 6 |
| *Nrg.5* | 1,90E-15 | 1,943668522 | 0,337 | 0,113 | 2,27E-11 | 6 |
| *regucalcin.5* | 2,46E-15 | -1,503064963 | 0,221 | 0,534 | 2,94E-11 | 6 |
| *Myo61F* | 3,44E-15 | 1,511234504 | 0,244 | 0,046 | 4,11E-11 | 6 |
| *alphaTub84B.2* | 4,24E-15 | 1,501192712 | 0,488 | 0,201 | 5,06E-11 | 6 |
| *CG12163.2* | 6,89E-15 | -1,274202562 | 0,297 | 0,6 | 8,22E-11 | 6 |
| *chic.1* | 7,51E-15 | 1,042396388 | 0,64 | 0,328 | 8,96E-11 | 6 |
| *Frl.1* | 1,19E-14 | 1,193886322 | 0,587 | 0,367 | 1,42E-10 | 6 |
| *foxo.1* | 1,45E-14 | -1,220764507 | 0,297 | 0,589 | 1,73E-10 | 6 |
| *tai.2* | 2,10E-14 | -0,827653499 | 0,692 | 0,839 | 2,51E-10 | 6 |
| *CG6006.2* | 3,83E-14 | 1,624388193 | 0,349 | 0,125 | 4,57E-10 | 6 |
| *srp.1* | 3,88E-14 | -0,585436407 | 0,797 | 0,93 | 4,63E-10 | 6 |
| *Shrm.1* | 5,35E-14 | 0,941584015 | 0,227 | 0,049 | 6,38E-10 | 6 |
| *lectin-28C.3* | 6,08E-14 | -1,213224792 | 0,384 | 0,666 | 7,25E-10 | 6 |
| *Imp.3* | 6,94E-14 | 0,980014071 | 0,709 | 0,414 | 8,28E-10 | 6 |
| *Arc1.2* | 7,76E-14 | 2,239687029 | 0,285 | 0,083 | 9,26E-10 | 6 |
| *Galphai.2* | 9,20E-14 | 1,248712118 | 0,488 | 0,227 | 1,10E-09 | 6 |
| *GlcAT-P.2* | 1,07E-13 | -1,199676109 | 0,262 | 0,57 | 1,28E-09 | 6 |
| *Ten-m.3* | 1,66E-13 | -0,507922569 | 0,994 | 0,996 | 1,98E-09 | 6 |
| *CG7120.3* | 2,05E-13 | -1,40931443 | 0,209 | 0,497 | 2,45E-09 | 6 |
| *capu.2* | 4,10E-13 | 1,48943507 | 0,413 | 0,169 | 4,89E-09 | 6 |
| *CG31523.2* | 4,10E-13 | 1,322041765 | 0,413 | 0,148 | 4,89E-09 | 6 |
| *zfh1.2* | 4,21E-13 | -0,637997512 | 0,773 | 0,933 | 5,03E-09 | 6 |
| *LamC* | 4,81E-13 | 1,203151915 | 0,244 | 0,049 | 5,74E-09 | 6 |
| *fz2.1* | 1,02E-12 | -1,197668029 | 0,273 | 0,555 | 1,21E-08 | 6 |
| *RhoGAP18B.1* | 1,13E-12 | 1,163871789 | 0,599 | 0,362 | 1,35E-08 | 6 |
| *KrT95D.3* | 1,25E-12 | 1,346826214 | 0,506 | 0,279 | 1,49E-08 | 6 |
| *Sema5c.2* | 1,67E-12 | -1,425581685 | 0,285 | 0,544 | 1,99E-08 | 6 |
| *CG10543.1* | 2,13E-12 | -0,976747993 | 0,57 | 0,76 | 2,54E-08 | 6 |
| *Pkn.2* | 2,18E-12 | -1,171534454 | 0,297 | 0,558 | 2,60E-08 | 6 |
| *LRR.1* | 2,62E-12 | 0,978015628 | 0,738 | 0,482 | 3,13E-08 | 6 |
| *psq.2* | 3,00E-12 | -0,738036199 | 0,657 | 0,843 | 3,57E-08 | 6 |
| *Pde9.1* | 3,06E-12 | -1,242795535 | 0,395 | 0,623 | 3,66E-08 | 6 |
| *CG5953.4* | 3,70E-12 | 1,248160739 | 0,709 | 0,459 | 4,41E-08 | 6 |
| *pan.2* | 4,26E-12 | -0,889092524 | 0,669 | 0,744 | 5,08E-08 | 6 |
| *spin.2* | 5,06E-12 | 0,934308541 | 0,663 | 0,371 | 6,03E-08 | 6 |
| *rudhira.2* | 5,43E-12 | -1,093585806 | 0,343 | 0,607 | 6,48E-08 | 6 |
| *CG10126* | 5,56E-12 | 1,174334219 | 0,227 | 0,047 | 6,63E-08 | 6 |
| *Fit1.1* | 6,46E-12 | 1,383010976 | 0,273 | 0,077 | 7,71E-08 | 6 |
| *l(3)L1231.2* | 7,55E-12 | -0,938199202 | 0,552 | 0,753 | 9,00E-08 | 6 |
| *mew.2* | 7,55E-12 | -0,687276393 | 0,686 | 0,859 | 9,00E-08 | 6 |
| *norpA.1* | 8,78E-12 | 0,954879911 | 0,692 | 0,411 | 1,05E-07 | 6 |
| *Lk6.2* | 9,20E-12 | -0,960188824 | 0,593 | 0,742 | 1,10E-07 | 6 |
| *CG32091.3* | 1,89E-11 | -0,937135715 | 0,483 | 0,696 | 2,25E-07 | 6 |
| *Glut4EF.4* | 2,04E-11 | -0,843889224 | 0,797 | 0,906 | 2,43E-07 | 6 |
| *hid.3* | 2,05E-11 | -1,501510635 | 0,157 | 0,403 | 2,44E-07 | 6 |
| *sdk.3* | 2,87E-11 | 1,112948051 | 0,61 | 0,378 | 3,43E-07 | 6 |

| | | | | | | |
|---|---|---|---|---|---|---|
| *Snoo* | 3,23E-11 | -0,977819891 | 0,419 | 0,647 | 3,85E-07 | 6 |
| *CG15145* | 3,42E-11 | 1,444345633 | 0,203 | 0,037 | 4,08E-07 | 6 |
| *r-l* | 3,73E-11 | 1,062070019 | 0,128 | 0,009 | 4,45E-07 | 6 |
| *CCT2* | 3,81E-11 | 1,055575202 | 0,221 | 0,049 | 4,54E-07 | 6 |
| *pain* | 3,85E-11 | 1,430776602 | 0,203 | 0,038 | 4,59E-07 | 6 |
| *Mmp1* | 4,10E-11 | 1,526664701 | 0,186 | 0,032 | 4,89E-07 | 6 |
| *GstD3* | 4,55E-11 | 1,548140168 | 0,227 | 0,055 | 5,43E-07 | 6 |
| *CG5346.1* | 4,92E-11 | 0,886308909 | 0,215 | 0,053 | 5,87E-07 | 6 |
| *jim.3* | 5,12E-11 | -1,21888525 | 0,262 | 0,483 | 6,11E-07 | 6 |
| *gpp.1* | 5,75E-11 | -0,747446151 | 0,599 | 0,789 | 6,86E-07 | 6 |
| *Baldspot.1* | 1,05E-10 | 1,202017441 | 0,343 | 0,117 | 1,25E-06 | 6 |
| *puc.2* | 1,22E-10 | 0,888059261 | 0,721 | 0,524 | 1,45E-06 | 6 |
| *Doa.2* | 1,23E-10 | 1,085188737 | 0,628 | 0,448 | 1,46E-06 | 6 |
| *Smr.1* | 1,55E-10 | -0,659198327 | 0,738 | 0,873 | 1,84E-06 | 6 |
| *pum.2* | 1,68E-10 | -0,360255142 | 0,977 | 0,997 | 2,00E-06 | 6 |
| *coro* | 1,83E-10 | 1,269551948 | 0,238 | 0,063 | 2,19E-06 | 6 |
| *l(2)41Ab.1* | 2,24E-10 | -1,03651271 | 0,471 | 0,689 | 2,67E-06 | 6 |
| *CG9253* | 2,59E-10 | 0,781440443 | 0,14 | 0,02 | 3,09E-06 | 6 |
| *dl.3* | 2,79E-10 | 1,375158479 | 0,314 | 0,104 | 3,32E-06 | 6 |
| *msk* | 2,83E-10 | 0,967967209 | 0,227 | 0,055 | 3,37E-06 | 6 |
| *CG44325.2* | 2,96E-10 | 1,102533079 | 0,419 | 0,171 | 3,53E-06 | 6 |
| *gogo.2* | 3,05E-10 | -1,639517467 | 0,041 | 0,227 | 3,64E-06 | 6 |
| *Hsp27.1* | 3,23E-10 | 3,007936702 | 0,192 | 0,105 | 3,85E-06 | 6 |
| *CG31955* | 5,03E-10 | 0,972652873 | 0,134 | 0,016 | 6,00E-06 | 6 |
| *vari.2* | 5,78E-10 | 1,184418389 | 0,395 | 0,157 | 6,89E-06 | 6 |
| *GstD1.2* | 6,63E-10 | 1,414572098 | 0,407 | 0,263 | 7,91E-06 | 6 |
| *Act42A.3* | 8,13E-10 | 1,079754951 | 0,372 | 0,144 | 9,70E-06 | 6 |
| *Nop60B* | 1,02E-09 | 1,104540284 | 0,192 | 0,041 | 1,21E-05 | 6 |
| *fru* | 1,04E-09 | 1,093142433 | 0,203 | 0,049 | 1,24E-05 | 6 |
| *RhoL* | 1,11E-09 | 1,081231325 | 0,302 | 0,105 | 1,32E-05 | 6 |
| *CG3961.2* | 1,47E-09 | -1,194459452 | 0,163 | 0,404 | 1,75E-05 | 6 |
| *CG13384.2* | 1,52E-09 | 1,037996356 | 0,407 | 0,194 | 1,82E-05 | 6 |
| *CG17646.2* | 2,82E-09 | 1,095951891 | 0,529 | 0,333 | 3,36E-05 | 6 |
| *ced-6.2* | 2,91E-09 | -0,98886765 | 0,256 | 0,512 | 3,48E-05 | 6 |
| *Traf4.1* | 3,04E-09 | -1,350646933 | 0,093 | 0,299 | 3,62E-05 | 6 |
| *EDTP.3* | 3,05E-09 | -1,418680858 | 0,058 | 0,229 | 3,63E-05 | 6 |
| *Ets21C* | 3,34E-09 | 1,278344444 | 0,267 | 0,087 | 3,98E-05 | 6 |
| *Drp1* | 3,53E-09 | 0,754899835 | 0,151 | 0,03 | 4,21E-05 | 6 |
| *sqd.2* | 4,79E-09 | 0,755851179 | 0,738 | 0,558 | 5,72E-05 | 6 |
| *CG6770* | 4,84E-09 | -0,926763455 | 0,43 | 0,657 | 5,77E-05 | 6 |
| *Lac.1* | 4,99E-09 | 1,089877742 | 0,238 | 0,07 | 5,95E-05 | 6 |
| *CG15144* | 5,30E-09 | 1,265051389 | 0,169 | 0,036 | 6,33E-05 | 6 |
| *Rel.1* | 5,64E-09 | 1,337779711 | 0,343 | 0,137 | 6,73E-05 | 6 |
| *CG5853* | 5,72E-09 | 1,358995108 | 0,151 | 0,031 | 6,83E-05 | 6 |
| *Xbp1* | 5,87E-09 | 0,653104571 | 0,419 | 0,203 | 7,00E-05 | 6 |
| *rl.1* | 6,30E-09 | -0,448138402 | 0,919 | 0,973 | 7,51E-05 | 6 |
| *stv.2* | 6,47E-09 | 1,354697786 | 0,43 | 0,211 | 7,71E-05 | 6 |
| *NimB4.2* | 6,50E-09 | -1,022636617 | 0,186 | 0,424 | 7,75E-05 | 6 |
| *CG9005.2* | 6,51E-09 | -1,124213236 | 0,203 | 0,411 | 7,77E-05 | 6 |
| *Gclm.1* | 6,62E-09 | 1,234904849 | 0,285 | 0,102 | 7,89E-05 | 6 |
| *Atf3.2* | 7,08E-09 | 1,106443037 | 0,39 | 0,177 | 8,45E-05 | 6 |
| *Arpc2* | 7,91E-09 | 0,888447528 | 0,233 | 0,072 | 9,44E-05 | 6 |
| *spz* | 9,34E-09 | 1,131375887 | 0,157 | 0,029 | 0,000111475 | 6 |
| *Nmda1.1* | 9,91E-09 | 1,099687848 | 0,267 | 0,089 | 0,000118163 | 6 |
| *CG9003.3* | 1,13E-08 | -1,114834178 | 0,157 | 0,355 | 0,000134975 | 6 |
| *Itgbn* | 1,15E-08 | 1,063941601 | 0,145 | 0,023 | 0,000137309 | 6 |
| *mam.1* | 1,18E-08 | -1,174287715 | 0,203 | 0,424 | 0,000141114 | 6 |
| *ifc* | 1,20E-08 | 0,956946699 | 0,128 | 0,021 | 0,000142633 | 6 |
| *sgg.3* | 1,28E-08 | -0,65874411 | 0,733 | 0,82 | 0,000152938 | 6 |
| *RasGAP1.1* | 1,41E-08 | 1,050905939 | 0,483 | 0,275 | 0,000167852 | 6 |
| *eIF2beta.1* | 1,44E-08 | 1,090512021 | 0,331 | 0,135 | 0,000171196 | 6 |

| | | | | | | |
|---|---|---|---|---|---|---|
| *sbb* | 1,49E-08 | -0,668665945 | 0,645 | 0,81 | 0,000178238 | 6 |
| *gish.1* | 1,54E-08 | 0,676625865 | 0,703 | 0,47 | 0,000183602 | 6 |
| *CG15143* | 1,60E-08 | 1,361528195 | 0,174 | 0,045 | 0,00019129 | 6 |
| *cindr.1* | 1,67E-08 | 0,897480619 | 0,535 | 0,295 | 0,000198832 | 6 |
| *prage.4* | 1,91E-08 | 0,921117987 | 0,494 | 0,262 | 0,000228108 | 6 |
| *Dmtn.1* | 1,93E-08 | -0,989248918 | 0,151 | 0,359 | 0,000229754 | 6 |
| *CG13117* | 1,94E-08 | 0,933874234 | 0,151 | 0,027 | 0,000231936 | 6 |
| *scyl.1* | 1,97E-08 | -0,906231673 | 0,355 | 0,538 | 0,000234802 | 6 |
| *Nhe3* | 2,00E-08 | 1,048524956 | 0,267 | 0,088 | 0,000238813 | 6 |
| *h.1* | 2,04E-08 | 1,01204801 | 0,57 | 0,354 | 0,00024372 | 6 |
| *SNF4Agamma.1* | 2,25E-08 | 0,729876454 | 0,831 | 0,707 | 0,000268487 | 6 |
| *PAPLA1.1* | 2,48E-08 | 1,188486338 | 0,378 | 0,191 | 0,000296381 | 6 |
| *Tm1.1* | 2,63E-08 | 0,848254832 | 0,564 | 0,359 | 0,00031382 | 6 |
| *CG13185* | 2,85E-08 | 0,921840327 | 0,174 | 0,042 | 0,000340017 | 6 |
| *Drs.2* | 3,24E-08 | 1,902785563 | 0,192 | 0,106 | 0,000386247 | 6 |
| *CG31122.1* | 3,25E-08 | -1,242221355 | 0,029 | 0,182 | 0,000388264 | 6 |
| *Sur-8* | 3,29E-08 | 1,067954863 | 0,169 | 0,039 | 0,000392867 | 6 |
| *DOR.3* | 3,44E-08 | -1,177623174 | 0,116 | 0,321 | 0,000409997 | 6 |
| *CG11791.1* | 3,62E-08 | 1,025329497 | 0,39 | 0,18 | 0,000431449 | 6 |
| *Tsp42Ed.1* | 3,73E-08 | 1,122308775 | 0,326 | 0,135 | 0,000444632 | 6 |
| *14-3-3zeta.1* | 3,96E-08 | 0,774063728 | 0,558 | 0,339 | 0,000472505 | 6 |
| *Ntf-2* | 4,69E-08 | 0,805071186 | 0,203 | 0,06 | 0,000559596 | 6 |
| *CG6231* | 4,84E-08 | 0,8970855 | 0,122 | 0,018 | 0,000577625 | 6 |
| *Ser.3* | 4,84E-08 | -1,127483607 | 0,18 | 0,39 | 0,000577882 | 6 |
| *Rac2* | 4,94E-08 | 0,863544485 | 0,558 | 0,32 | 0,000589819 | 6 |
| *Dhap-at* | 5,09E-08 | 0,998684564 | 0,134 | 0,023 | 0,000607141 | 6 |
| *CG42668.2* | 5,16E-08 | 0,864456919 | 0,413 | 0,201 | 0,000615873 | 6 |
| *SPARC.3* | 5,51E-08 | -0,796006599 | 0,326 | 0,548 | 0,000657715 | 6 |
| *plum.1* | 6,17E-08 | 1,39404565 | 0,267 | 0,115 | 0,00073556 | 6 |
| *Hsp23* | 6,27E-08 | 2,379192275 | 0,134 | 0,044 | 0,000748454 | 6 |
| *pigs.4* | 6,51E-08 | 0,780671352 | 0,674 | 0,57 | 0,000776845 | 6 |
| *Npc2a.1* | 7,03E-08 | 1,082682472 | 0,273 | 0,096 | 0,000838476 | 6 |
| *Irc.1* | 7,11E-08 | 0,907132597 | 0,314 | 0,126 | 0,000848149 | 6 |
| *Hsc70-4.3* | 8,85E-08 | 0,941748734 | 0,465 | 0,268 | 0,001055152 | 6 |
| *CG32425* | 1,00E-07 | 1,002421795 | 0,343 | 0,152 | 0,001198093 | 6 |
| *CG42588* | 1,18E-07 | 0,971877137 | 0,203 | 0,061 | 0,001402657 | 6 |
| *Pmp70* | 1,43E-07 | 1,085045705 | 0,157 | 0,037 | 0,001704145 | 6 |
| *Ziz.3* | 1,55E-07 | -1,171327141 | 0,157 | 0,359 | 0,001845873 | 6 |
| *CG10311* | 1,58E-07 | 1,11228308 | 0,209 | 0,06 | 0,001883809 | 6 |
| *Ist1.2* | 1,67E-07 | 0,902985569 | 0,314 | 0,133 | 0,001990627 | 6 |
| *CG5059.3* | 1,68E-07 | -1,053146752 | 0,221 | 0,366 | 0,002007459 | 6 |
| *CG17739* | 1,93E-07 | 0,79679555 | 0,105 | 0,012 | 0,002304611 | 6 |
| *bves.2* | 2,03E-07 | 0,98099254 | 0,413 | 0,208 | 0,002420176 | 6 |
| *Diap1.2* | 2,06E-07 | 0,746469571 | 0,57 | 0,449 | 0,002456325 | 6 |
| *Mvl.2* | 2,26E-07 | 0,904109027 | 0,535 | 0,311 | 0,002700972 | 6 |
| *tsr.2* | 2,30E-07 | 0,661033065 | 0,262 | 0,102 | 0,002743273 | 6 |
| *Tomosyn* | 2,31E-07 | 0,649250432 | 0,587 | 0,378 | 0,002759205 | 6 |
| *kay.2* | 2,59E-07 | 1,03935901 | 0,57 | 0,393 | 0,003084264 | 6 |
| *PDZ-GEF.2* | 2,78E-07 | -0,988606202 | 0,105 | 0,294 | 0,00331083 | 6 |
| *Vinc.1* | 2,93E-07 | 0,833651885 | 0,244 | 0,093 | 0,003493656 | 6 |
| *Fib* | 3,18E-07 | 0,858008716 | 0,105 | 0,015 | 0,003790113 | 6 |
| *egh.4* | 3,25E-07 | 0,645327664 | 0,61 | 0,394 | 0,0038823 | 6 |
| *Spn88Ea* | 3,37E-07 | 1,006877601 | 0,157 | 0,04 | 0,00402076 | 6 |
| *Hers.1* | 3,52E-07 | -0,7626553 | 0,43 | 0,621 | 0,004202523 | 6 |
| *pod1.1* | 4,44E-07 | 0,922917057 | 0,395 | 0,194 | 0,005292714 | 6 |
| *Pdp1.3* | 4,79E-07 | 0,586667325 | 0,767 | 0,575 | 0,005714628 | 6 |
| *PRAS40.4* | 5,19E-07 | -1,037006603 | 0,18 | 0,375 | 0,006192007 | 6 |
| *Hel89B.1* | 5,87E-07 | 0,85322026 | 0,512 | 0,299 | 0,0070017 | 6 |
| *Dyrk2.3* | 6,08E-07 | -0,78708965 | 0,39 | 0,589 | 0,007251419 | 6 |
| *alc.2* | 6,30E-07 | -1,012641539 | 0,116 | 0,304 | 0,007513409 | 6 |
| *Sirup.1* | 7,25E-07 | -1,091642604 | 0,087 | 0,259 | 0,008648754 | 6 |

| | | | | | | |
|---|---|---|---|---|---|---|
| *pnt.5* | 7,27E-07 | -0,449388475 | 0,82 | 0,909 | 0,008676865 | 6 |
| *Chd64.3* | 8,45E-07 | 0,788949727 | 0,64 | 0,481 | 0,010081745 | 6 |
| *dnc.1* | 8,65E-07 | -1,007453678 | 0,308 | 0,501 | 0,010321447 | 6 |
| *ATP8A.2* | 9,64E-07 | 1,005781807 | 0,308 | 0,145 | 0,011493795 | 6 |
| *RhoU* | 9,76E-07 | 1,256679819 | 0,169 | 0,049 | 0,011637333 | 6 |
| *CARPB* | 9,89E-07 | 0,701568924 | 0,116 | 0,023 | 0,011799643 | 6 |
| *cib.1* | 1,04E-06 | 1,048596131 | 0,262 | 0,117 | 0,012425877 | 6 |
| *CG10365.1* | 1,09E-06 | -1,123212558 | 0,035 | 0,136 | 0,012966155 | 6 |
| *CG6357.1* | 1,10E-06 | 1,001532293 | 0,238 | 0,082 | 0,013088814 | 6 |
| *CG12576.3* | 1,13E-06 | -1,024442911 | 0,122 | 0,295 | 0,01353936 | 6 |
| *Esyt2.1* | 1,14E-06 | 0,971846906 | 0,401 | 0,206 | 0,013588387 | 6 |
| *CG5151.3* | 1,21E-06 | -0,50595011 | 0,651 | 0,817 | 0,01438153 | 6 |
| *Corin.1* | 1,23E-06 | -1,06027059 | 0,105 | 0,257 | 0,01469616 | 6 |
| *GlcT* | 1,24E-06 | 0,872637799 | 0,128 | 0,028 | 0,014802625 | 6 |
| *Moe.2* | 1,35E-06 | 0,648957806 | 0,657 | 0,445 | 0,01612283 | 6 |
| *slmb.1* | 1,44E-06 | 0,825444144 | 0,244 | 0,103 | 0,017235533 | 6 |
| *Ac13E.1* | 1,45E-06 | -0,738837417 | 0,291 | 0,459 | 0,017330069 | 6 |
| *Thor.3* | 1,61E-06 | -1,173573705 | 0,244 | 0,424 | 0,01926434 | 6 |
| *Msp300.3* | 1,77E-06 | 0,84412639 | 0,669 | 0,557 | 0,021076562 | 6 |
| *caz.1* | 1,80E-06 | 0,790651412 | 0,302 | 0,137 | 0,021436832 | 6 |
| *NKAIN* | 1,80E-06 | 0,931814278 | 0,355 | 0,175 | 0,021458311 | 6 |
| *NimB1.1* | 1,92E-06 | -0,997034375 | 0,163 | 0,325 | 0,022882628 | 6 |
| *NFAT.1* | 2,02E-06 | -0,737718563 | 0,395 | 0,524 | 0,024103863 | 6 |
| *mask* | 2,04E-06 | 0,810105921 | 0,541 | 0,402 | 0,024313152 | 6 |
| *eIF4A.2* | 2,41E-06 | 0,759245639 | 0,419 | 0,228 | 0,02870877 | 6 |
| *DIP-epsilon.1* | 2,44E-06 | -1,077814713 | 0,105 | 0,276 | 0,029091002 | 6 |
| *Sarm.3* | 2,47E-06 | 0,704239408 | 0,709 | 0,608 | 0,029496716 | 6 |
| *ZnT63C.1* | 2,74E-06 | 0,836560867 | 0,372 | 0,188 | 0,032651561 | 6 |
| *RapGAP1.2* | 2,74E-06 | -0,610192224 | 0,57 | 0,727 | 0,032680069 | 6 |
| *apt* | 2,76E-06 | 0,987565767 | 0,14 | 0,039 | 0,032950179 | 6 |
| *sd.1* | 2,85E-06 | 0,75544349 | 0,413 | 0,218 | 0,034043485 | 6 |
| *Tsp42Ee.1* | 2,86E-06 | 0,947586884 | 0,256 | 0,112 | 0,034081377 | 6 |
| *CG6051.1* | 2,91E-06 | 0,712888412 | 0,552 | 0,351 | 0,034733572 | 6 |
| *Tre1.1* | 2,98E-06 | 0,934720158 | 0,244 | 0,095 | 0,035538707 | 6 |
| *Timp* | 2,99E-06 | 1,137420457 | 0,128 | 0,031 | 0,035722421 | 6 |
| *eEF1alpha1.3* | 3,08E-06 | 0,57993048 | 0,68 | 0,499 | 0,036684631 | 6 |
| *srl* | 3,10E-06 | 0,94455201 | 0,116 | 0,023 | 0,037003246 | 6 |
| *mnb.4* | 3,11E-06 | -0,805067432 | 0,285 | 0,41 | 0,037088499 | 6 |
| *Had2.3* | 3,22E-06 | -1,017899742 | 0,169 | 0,346 | 0,038416735 | 6 |
| *Sf3b3* | 3,25E-06 | 0,784888102 | 0,157 | 0,044 | 0,038807729 | 6 |
| *Ubi-p63E.1* | 3,30E-06 | 1,004664721 | 0,541 | 0,389 | 0,039308484 | 6 |
| *Crtc.3* | 3,42E-06 | -0,792411266 | 0,372 | 0,555 | 0,040767364 | 6 |
| *CG31637.1* | 3,53E-06 | -0,916162561 | 0,349 | 0,514 | 0,042164578 | 6 |
| *CG7029* | 3,66E-06 | 0,828729548 | 0,36 | 0,221 | 0,043669465 | 6 |
| *CG32066.1* | 3,69E-06 | 0,669457659 | 0,686 | 0,555 | 0,043972475 | 6 |
| *Tpr2.1* | 3,71E-06 | 0,965332832 | 0,413 | 0,238 | 0,044256452 | 6 |
| *Hsp26.1* | 4,03E-06 | 2,493337664 | 0,238 | 0,119 | 0,048041117 | 6 |
| *Arpc3B* | 4,12E-06 | 0,617568184 | 0,11 | 0,027 | 0,049167976 | 6 |
| *CG31145.3* | 4,60E-13 | 0,911557957 | 1 | 0,731 | 5,49E-09 | 7 |
| *Vha100-2.4* | 1,31E-10 | 0,958372425 | 0,565 | 0,206 | 1,57E-06 | 7 |
| *cwo.3* | 1,83E-10 | -1,205292646 | 0,594 | 0,63 | 2,18E-06 | 7 |
| *blot.1* | 2,89E-10 | -1,203597006 | 0,551 | 0,532 | 3,45E-06 | 7 |
| *DOR.4* | 3,12E-10 | 0,373873765 | 0,58 | 0,28 | 3,72E-06 | 7 |
| *CrebA.5* | 5,13E-10 | 0,276923635 | 0,536 | 0,242 | 6,12E-06 | 7 |
| *CaMKI.1* | 5,75E-10 | -0,257045617 | 0,493 | 0,297 | 6,86E-06 | 7 |
| *RpL10Ab.3* | 1,30E-09 | -1,454094 | 0,145 | 0,266 | 1,55E-05 | 7 |
| *Cals.2* | 1,85E-09 | 0,471293938 | 0,522 | 0,234 | 2,21E-05 | 7 |
| *Pp1alpha-96A* | 2,92E-09 | -0,522246444 | 0,203 | 0,143 | 3,48E-05 | 7 |
| *eEF2.4* | 4,57E-09 | -0,798242043 | 0,362 | 0,312 | 5,46E-05 | 7 |
| *mtd.3* | 5,57E-09 | 0,756477849 | 0,942 | 0,749 | 6,65E-05 | 7 |
| *DIP-epsilon.2* | 6,79E-09 | 0,600439784 | 0,536 | 0,239 | 8,10E-05 | 7 |

| | | | | | | |
|---|---|---|---|---|---|---|
| *app.3* | 6,88E-09 | 0,613235389 | 0,725 | 0,371 | 8,20E-05 | 7 |
| *Caper* | 9,68E-09 | -0,378670942 | 0,362 | 0,275 | 0,000115467 | 7 |
| *Best1.1* | 1,38E-08 | 0,63901194 | 0,551 | 0,25 | 0,000164699 | 7 |
| *sug* | 1,44E-08 | 0,70642019 | 0,188 | 0,031 | 0,000171802 | 7 |
| *CG3777.3* | 1,72E-08 | 1,009332387 | 0,536 | 0,208 | 0,0002051 | 7 |
| *l(1)G0196.3* | 2,74E-08 | 0,612972945 | 0,725 | 0,387 | 0,000326316 | 7 |
| *pum.3* | 2,95E-08 | 0,470488238 | 1 | 0,994 | 0,000351716 | 7 |
| *eIF4A.3* | 3,00E-08 | -0,945090427 | 0,246 | 0,253 | 0,000358367 | 7 |
| *Pxn.4* | 3,11E-08 | 0,715547704 | 0,87 | 0,525 | 0,000371257 | 7 |
| *CG43236.2* | 3,44E-08 | 1,082107295 | 0,406 | 0,124 | 0,000409942 | 7 |
| *Had2.4* | 4,27E-08 | 0,911002664 | 0,652 | 0,307 | 0,000509638 | 7 |
| *CG5151.4* | 4,58E-08 | 0,748702068 | 0,957 | 0,788 | 0,00054666 | 7 |
| *CG3961.3* | 4,96E-08 | 0,722330295 | 0,71 | 0,355 | 0,00059161 | 7 |
| *Pdk.4* | 5,58E-08 | -1,743106509 | 0,203 | 0,44 | 0,000665088 | 7 |
| *Ten-m.4* | 5,72E-08 | 0,529774137 | 1 | 0,995 | 0,000682378 | 7 |
| *Prps.3* | 6,20E-08 | -1,827973938 | 0,159 | 0,361 | 0,0007396 | 7 |
| *RpL4.2* | 6,84E-08 | -1,157818444 | 0,145 | 0,202 | 0,00081543 | 7 |
| *CG14153.1* | 7,00E-08 | 0,876663029 | 0,304 | 0,083 | 0,000835135 | 7 |
| *hppy.2* | 7,56E-08 | 0,437835198 | 0,884 | 0,568 | 0,000902061 | 7 |
| *Adar.1* | 7,83E-08 | 0,502206501 | 0,464 | 0,213 | 0,000934348 | 7 |
| *l(3)80Fg.2* | 9,48E-08 | 0,768664562 | 0,899 | 0,594 | 0,001131438 | 7 |
| *Rbp1-like.2* | 9,60E-08 | 0,793698637 | 0,522 | 0,225 | 0,001144975 | 7 |
| *wrd.1* | 1,03E-07 | 0,325438449 | 0,391 | 0,182 | 0,001229976 | 7 |
| *PGRP-LC.3* | 1,10E-07 | 0,662945426 | 0,638 | 0,324 | 0,001306317 | 7 |
| *Ten-a.1* | 1,12E-07 | -1,307066332 | 0,159 | 0,219 | 0,001340564 | 7 |
| *RpL21.4* | 1,13E-07 | -1,165351416 | 0,217 | 0,278 | 0,001343552 | 7 |
| *Idgf2* | 1,17E-07 | 0,258676856 | 0,348 | 0,168 | 0,001398724 | 7 |
| *jim.4* | 1,19E-07 | -0,282603348 | 0,594 | 0,448 | 0,001420889 | 7 |
| *CG42694.1* | 1,21E-07 | 0,596757708 | 0,261 | 0,093 | 0,001440688 | 7 |
| *Lpin.4* | 1,37E-07 | -2,013663414 | 0,101 | 0,32 | 0,001633716 | 7 |
| *Nc73EF.3* | 1,49E-07 | -0,632095107 | 0,203 | 0,177 | 0,001776768 | 7 |
| *Cat* | 1,52E-07 | -0,534170314 | 0,174 | 0,128 | 0,00180844 | 7 |
| *ninaE.1* | 1,53E-07 | -1,269981636 | 0,217 | 0,242 | 0,001819898 | 7 |
| *CG3164.2* | 1,59E-07 | 0,726581053 | 0,507 | 0,222 | 0,001893257 | 7 |
| *Culd* | 1,89E-07 | -0,420608899 | 0,145 | 0,1 | 0,002256574 | 7 |
| *CG4259* | 2,08E-07 | 1,168164573 | 0,391 | 0,118 | 0,002480693 | 7 |
| *Pcf11.1* | 2,21E-07 | -0,282414065 | 0,391 | 0,28 | 0,002639083 | 7 |
| *larp.1* | 2,23E-07 | -0,744855143 | 0,275 | 0,276 | 0,002656793 | 7 |
| *Sema2a.4* | 2,29E-07 | 0,816868916 | 0,768 | 0,43 | 0,002726598 | 7 |
| *RpL3.4* | 2,46E-07 | -1,546188252 | 0,101 | 0,27 | 0,002931161 | 7 |
| *Indy.1* | 2,53E-07 | 0,556254699 | 0,435 | 0,189 | 0,003018594 | 7 |
| *Pde9.2* | 2,62E-07 | 0,322826352 | 0,87 | 0,58 | 0,003125209 | 7 |
| *CG15695.4* | 2,64E-07 | 0,46979004 | 0,246 | 0,083 | 0,003154514 | 7 |
| *ush.4* | 2,68E-07 | 0,67853958 | 0,942 | 0,711 | 0,003200527 | 7 |
| *Ank* | 2,75E-07 | -0,459548673 | 0,406 | 0,354 | 0,003276291 | 7 |
| *bves.3* | 3,18E-07 | 0,471703924 | 0,449 | 0,222 | 0,003798064 | 7 |
| *Btk29A.5* | 4,05E-07 | 0,693016379 | 0,957 | 0,764 | 0,004826919 | 7 |
| *Mad* | 4,41E-07 | 0,399012439 | 0,435 | 0,215 | 0,005265941 | 7 |
| *ClC-a.1* | 4,48E-07 | 0,805839499 | 0,391 | 0,144 | 0,00534152 | 7 |
| *CG31122.2* | 4,50E-07 | 0,831901167 | 0,391 | 0,151 | 0,005364306 | 7 |
| *Sema5c.3* | 4,99E-07 | 1,030934417 | 0,754 | 0,498 | 0,005954593 | 7 |
| *egh.5* | 5,26E-07 | 0,690901476 | 0,739 | 0,405 | 0,006273216 | 7 |
| *fl(2)d* | 5,64E-07 | 0,301770331 | 0,377 | 0,18 | 0,006731563 | 7 |
| *rhea.4* | 5,96E-07 | -0,513394218 | 0,435 | 0,367 | 0,007104708 | 7 |
| *eEF1alpha1.4* | 6,15E-07 | -0,926147961 | 0,507 | 0,523 | 0,007334346 | 7 |
| *Rga.1* | 6,33E-07 | 0,535222136 | 0,333 | 0,143 | 0,007553775 | 7 |
| *RhoGEF2* | 6,57E-07 | 0,512636649 | 0,681 | 0,381 | 0,007839174 | 7 |
| *CG31211.2* | 6,72E-07 | 0,402083545 | 0,319 | 0,147 | 0,008012816 | 7 |
| *Thd1.2* | 6,73E-07 | 0,376344259 | 0,609 | 0,358 | 0,008032423 | 7 |
| *eEF1beta.1* | 8,42E-07 | -0,720617881 | 0,116 | 0,127 | 0,010047459 | 7 |
| *CG17124.2* | 1,01E-06 | 0,636757481 | 0,739 | 0,411 | 0,011993632 | 7 |

| | | | | | | |
|---|---|---|---|---|---|---|
| *pAbp.4* | 1,07E-06 | -0,832879954 | 0,42 | 0,492 | 0,012738004 | 7 |
| *blw.1* | 1,14E-06 | -0,672633094 | 0,13 | 0,135 | 0,013643782 | 7 |
| *CG9328.2* | 1,24E-06 | 0,817300542 | 0,783 | 0,464 | 0,014821554 | 7 |
| *CG41099* | 1,30E-06 | 0,511486233 | 0,348 | 0,152 | 0,015470696 | 7 |
| *NimB2.2* | 1,31E-06 | -0,586356537 | 0,261 | 0,24 | 0,015663339 | 7 |
| *upSET* | 1,48E-06 | 0,465643371 | 0,725 | 0,44 | 0,017624562 | 7 |
| *Srrm234* | 1,50E-06 | 0,30747616 | 0,377 | 0,195 | 0,017910763 | 7 |
| *uex.4* | 1,60E-06 | 0,820612548 | 0,826 | 0,539 | 0,01911205 | 7 |
| *Cyp6w1* | 1,62E-06 | 0,305983696 | 0,188 | 0,064 | 0,019310307 | 7 |
| *drongo.1* | 1,62E-06 | 0,506436527 | 0,783 | 0,48 | 0,019316812 | 7 |
| *RpS30.3* | 1,71E-06 | -0,813224983 | 0,232 | 0,235 | 0,020437683 | 7 |
| *UQCR-Q.1* | 1,81E-06 | -0,355886011 | 0,145 | 0,088 | 0,021604479 | 7 |
| *GstE12.1* | 1,93E-06 | 0,272313061 | 0,232 | 0,08 | 0,022994405 | 7 |
| *Atf6.1* | 2,01E-06 | 0,713607523 | 0,594 | 0,294 | 0,023956654 | 7 |
| *Saf-B.2* | 2,08E-06 | 0,301613491 | 0,29 | 0,139 | 0,024853085 | 7 |
| *l(3)L1231.3* | 2,10E-06 | 0,343390737 | 0,957 | 0,715 | 0,025073017 | 7 |
| *Bacc.2* | 2,76E-06 | -0,468186681 | 0,464 | 0,388 | 0,032953523 | 7 |
| *RpL23.2* | 2,77E-06 | -0,980105584 | 0,275 | 0,334 | 0,033097127 | 7 |
| *InR.5* | 2,86E-06 | -0,993041751 | 0,812 | 0,856 | 0,034172333 | 7 |
| *sta.3* | 3,05E-06 | -0,760325988 | 0,246 | 0,277 | 0,036423482 | 7 |
| *CG9171.4* | 3,31E-06 | 0,799132227 | 0,652 | 0,339 | 0,039435066 | 7 |
| *Evi5.1* | 3,34E-06 | 0,460184043 | 0,42 | 0,208 | 0,039896868 | 7 |
| *Syp.3* | 3,35E-06 | 0,570230479 | 0,971 | 0,844 | 0,039956161 | 7 |
| *SCaMC.3* | 3,54E-06 | -1,651676999 | 0,087 | 0,297 | 0,042187792 | 7 |
| *crq.1* | 3,63E-06 | 0,709927603 | 0,725 | 0,42 | 0,04329871 | 7 |
| *CG3726.2* | 3,66E-06 | 0,385607869 | 0,899 | 0,627 | 0,043659863 | 7 |
| *RpL7.2* | 3,72E-06 | -0,964301376 | 0,188 | 0,247 | 0,044334333 | 7 |
| *Hml.5* | 3,74E-06 | 0,730232787 | 0,899 | 0,638 | 0,044591317 | 7 |
| *RpS15.3* | 3,75E-06 | -1,453278673 | 0,072 | 0,328 | 0,044748805 | 7 |
| *CG11360.2* | 3,76E-06 | 0,764327916 | 0,319 | 0,111 | 0,0448111 | 7 |
| *CG31635.1* | 4,00E-06 | 0,596939043 | 0,609 | 0,324 | 0,047714761 | 7 |
| *sesB.2* | 4,10E-06 | -1,176935054 | 0,13 | 0,212 | 0,048885115 | 7 |
| *RpS27.3* | 4,18E-06 | -1,5598844 | 0,13 | 0,322 | 0,049905029 | 7 |
| *rl.2* | 4,19E-06 | 0,443673766 | 1 | 0,964 | 0,049962938 | 7 |
| *CG13743* | 2,30E-36 | 5,053771763 | 0,518 | 0,019 | 2,75E-32 | 8 |
| *PPO2* | 3,98E-35 | 5,408811201 | 0,536 | 0,024 | 4,74E-31 | 8 |
| *robo2* | 6,91E-35 | 3,921896649 | 0,643 | 0,033 | 8,25E-31 | 8 |
| *lz* | 2,65E-32 | 3,843451278 | 0,482 | 0,006 | 3,16E-28 | 8 |
| *Pde1c* | 6,51E-32 | 6,368113964 | 0,464 | 0,062 | 7,77E-28 | 8 |
| *klu* | 1,52E-30 | 4,124946793 | 0,536 | 0,024 | 1,82E-26 | 8 |
| *N.3* | 1,58E-29 | 2,830268982 | 0,786 | 0,288 | 1,88E-25 | 8 |
| *peb* | 9,30E-28 | 3,805778578 | 0,482 | 0,018 | 1,11E-23 | 8 |
| *Ten-m.5* | 1,41E-27 | -1,353538709 | 0,929 | 0,998 | 1,68E-23 | 8 |
| *PPO1* | 6,49E-27 | 4,119930235 | 0,446 | 0,015 | 7,74E-23 | 8 |
| *E(spl)mbeta-HLH* | 2,94E-26 | 2,825689357 | 0,518 | 0,024 | 3,51E-22 | 8 |
| *Tet.2* | 1,78E-24 | 3,051853875 | 0,482 | 0,196 | 2,13E-20 | 8 |
| *ct.3* | 2,76E-23 | 1,808075378 | 0,786 | 0,764 | 3,30E-19 | 8 |
| *mtgo.1* | 2,18E-20 | -1,532172696 | 0,804 | 0,97 | 2,60E-16 | 8 |
| *tna.5* | 2,96E-20 | 2,667440852 | 0,536 | 0,208 | 3,54E-16 | 8 |
| *CG43187* | 3,49E-15 | 2,051995704 | 0,214 | 0,001 | 4,16E-11 | 8 |
| *caps.5* | 3,61E-15 | 2,643640629 | 0,357 | 0,093 | 4,31E-11 | 8 |
| *E(spl)malpha-BFM* | 1,20E-14 | 2,375337586 | 0,321 | 0,02 | 1,43E-10 | 8 |
| *Jupiter.2* | 1,55E-13 | -1,741863725 | 0,536 | 0,854 | 1,85E-09 | 8 |
| *CG31431* | 2,51E-13 | 2,412144802 | 0,375 | 0,047 | 2,99E-09 | 8 |
| *Ppn.3* | 2,08E-12 | -1,169080747 | 0,768 | 0,974 | 2,48E-08 | 8 |
| *cpo.4* | 7,67E-12 | 1,326492564 | 0,893 | 0,693 | 9,15E-08 | 8 |
| *Mctp* | 1,14E-11 | 2,287285796 | 0,232 | 0,012 | 1,36E-07 | 8 |
| *Men.3* | 1,93E-11 | 2,320647444 | 0,554 | 0,216 | 2,31E-07 | 8 |
| *CG9932.1* | 4,58E-11 | -1,199003042 | 0,768 | 0,94 | 5,46E-07 | 8 |
| *zfh2* | 1,18E-10 | 2,524518907 | 0,25 | 0,036 | 1,41E-06 | 8 |
| *CG13252* | 1,30E-10 | 1,498091984 | 0,214 | 0,01 | 1,56E-06 | 8 |

| | | | | | | |
|---|---|---|---|---|---|---|
| *Cirl.1* | 1,77E-10 | 1,492940806 | 0,607 | 0,405 | 2,11E-06 | 8 |
| *Ndae1* | 3,08E-10 | 2,21968017 | 0,375 | 0,1 | 3,68E-06 | 8 |
| *ATP8B.1* | 6,13E-10 | 1,907987213 | 0,357 | 0,06 | 7,31E-06 | 8 |
| *CG5828* | 4,44E-09 | 1,551893003 | 0,179 | 0,008 | 5,29E-05 | 8 |
| *Prosap.3* | 5,32E-09 | -1,215066021 | 0,821 | 0,952 | 6,34E-05 | 8 |
| *kst.2* | 6,26E-09 | 2,011108786 | 0,518 | 0,18 | 7,47E-05 | 8 |
| *Trim9* | 1,07E-08 | 1,841805013 | 0,179 | 0,011 | 0,000128167 | 8 |
| *drpr.3* | 3,65E-08 | -1,264288371 | 0,482 | 0,765 | 0,000435214 | 8 |
| *Pde9.3* | 1,10E-07 | 1,153604446 | 0,821 | 0,585 | 0,001315127 | 8 |
| *bru2.1* | 1,44E-07 | 1,325493599 | 0,304 | 0,063 | 0,001723721 | 8 |
| *fkh* | 1,51E-07 | 1,200260333 | 0,143 | 0,008 | 0,001801489 | 8 |
| *Rtnl1.4* | 1,76E-07 | 1,252315272 | 0,411 | 0,108 | 0,002099742 | 8 |
| *CG32264.6* | 1,95E-07 | 1,336807392 | 0,696 | 0,435 | 0,002322574 | 8 |
| *sm.3* | 2,33E-07 | 1,417934997 | 0,536 | 0,229 | 0,002780002 | 8 |
| *CAP* | 2,50E-07 | 1,63227155 | 0,304 | 0,064 | 0,002982203 | 8 |
| *Ctr1A.1* | 2,51E-07 | 1,81889674 | 0,286 | 0,1 | 0,002994819 | 8 |
| *Elk* | 3,28E-07 | 1,342676286 | 0,143 | 0,005 | 0,003917684 | 8 |
| *Col4a1.3* | 3,35E-07 | -0,815942915 | 0,714 | 0,933 | 0,003996657 | 8 |
| *fok.4* | 3,54E-07 | 1,784485958 | 0,411 | 0,176 | 0,004218029 | 8 |
| *AstA-R1* | 3,97E-07 | 5,406423927 | 0,143 | 0,012 | 0,004734589 | 8 |
| *Sema1b.2* | 3,99E-07 | -1,58154195 | 0,304 | 0,556 | 0,004754495 | 8 |
| *kek5.2* | 4,35E-07 | 1,245954082 | 0,589 | 0,38 | 0,0051875 | 8 |
| *Fili* | 4,38E-07 | 1,449785134 | 0,161 | 0,011 | 0,005219015 | 8 |
| *CG9518* | 4,39E-07 | 2,391078925 | 0,125 | 0,023 | 0,005232487 | 8 |
| *SKIP* | 4,63E-07 | 2,508831604 | 0,214 | 0,03 | 0,005520929 | 8 |
| *Pvr.4* | 4,76E-07 | -1,055349535 | 0,732 | 0,884 | 0,005674541 | 8 |
| *aqz.2* | 4,85E-07 | 1,064216746 | 0,964 | 0,894 | 0,005790403 | 8 |
| *CG9743* | 5,16E-07 | 1,694602721 | 0,214 | 0,026 | 0,006150506 | 8 |
| *sty.4* | 6,52E-07 | -1,098343802 | 0,643 | 0,859 | 0,007780424 | 8 |
| *CG8468* | 7,65E-07 | 1,606394327 | 0,196 | 0,029 | 0,009129008 | 8 |
| *CAH2* | 7,83E-07 | 1,064536415 | 0,143 | 0,008 | 0,009339616 | 8 |
| *LpR2.4* | 8,22E-07 | -0,982033725 | 0,768 | 0,908 | 0,009806558 | 8 |
| *Mvl.4* | 9,90E-07 | -1,882622023 | 0,054 | 0,352 | 0,01181276 | 8 |
| *CG43693* | 1,01E-06 | 1,273620679 | 0,196 | 0,023 | 0,012094857 | 8 |
| *cv-c.6* | 1,20E-06 | 0,844524068 | 0,982 | 0,919 | 0,014311523 | 8 |
| *Egfr.1* | 1,27E-06 | 1,730482324 | 0,286 | 0,063 | 0,015167881 | 8 |
| *Dl* | 2,42E-06 | 1,681662426 | 0,125 | 0,006 | 0,028866383 | 8 |
| *Had2.5* | 2,69E-06 | -1,859418209 | 0,054 | 0,336 | 0,032056858 | 8 |
| *nudC* | 3,09E-06 | 1,14376621 | 0,196 | 0,028 | 0,03690003 | 8 |
| *Ncc69* | 3,27E-06 | 1,65321262 | 0,161 | 0,013 | 0,03897657 | 8 |
| *CG31145.4* | 3,49E-06 | -1,1183218 | 0,5 | 0,755 | 0,041587152 | 8 |
| *Mmp2.5* | 3,50E-06 | -2,069633931 | 0,179 | 0,473 | 0,041757379 | 8 |

ntity
