## Supplementary material for "DNA damage signaling in *Drosophila* macrophages modulates systemic cytokine levels in response to oxidative stress": Table S2

| Gene name | p-value | Average Log2 Fold Change | pct.1 | pct.2 | adjusted p-value | ter Iden |
|---|---|---|---|---|---|---|
| *Gbs-70E* | 0 | 2,896368201 | 0,956 | 0,32 | 0 | 1 |
| *FASN1* | 0 | 2,412893799 | 0,993 | 0,692 | 0 | 1 |
| *Nplp2* | 1,83E-251 | 1,834846643 | 0,968 | 0,52 | 2,18E-247 | 1 |
| *CG9674* | 1,10E-249 | -2,83187681 | 0,241 | 0,826 | 1,31E-245 | 1 |
| *GstD1* | 4,13E-248 | -3,159999252 | 0,274 | 0,84 | 4,92E-244 | 1 |
| *CG14823* | 5,38E-230 | 2,27517809 | 0,713 | 0,165 | 6,42E-226 | 1 |
| *CG11089* | 4,95E-229 | -2,755218747 | 0,294 | 0,84 | 5,91E-225 | 1 |
| *ref(2)P* | 3,04E-225 | -2,802596089 | 0,274 | 0,815 | 3,62E-221 | 1 |
| *stv* | 1,43E-221 | -3,409918205 | 0,244 | 0,803 | 1,71E-217 | 1 |
| *Culd* | 2,43E-197 | 2,044099788 | 0,679 | 0,179 | 2,90E-193 | 1 |
| *Tsp42Ed* | 1,71E-194 | -2,790859636 | 0,053 | 0,625 | 2,04E-190 | 1 |
| *CG34166* | 7,28E-193 | 2,383013481 | 0,954 | 0,728 | 8,68E-189 | 1 |
| *Nmdmc* | 2,44E-189 | -2,067661189 | 0,292 | 0,817 | 2,91E-185 | 1 |
| *AnxB9* | 3,28E-188 | -3,193625219 | 0,127 | 0,678 | 3,91E-184 | 1 |
| *ACC* | 7,15E-187 | 1,534764509 | 0,947 | 0,585 | 8,53E-183 | 1 |
| *Invadolysin* | 1,10E-185 | 2,621709746 | 0,583 | 0,121 | 1,31E-181 | 1 |
| *InR* | 2,23E-184 | -1,667163042 | 0,818 | 0,959 | 2,65E-180 | 1 |
| *CG6503* | 7,74E-180 | 1,536688962 | 0,965 | 0,719 | 9,23E-176 | 1 |
| *raw* | 3,47E-179 | -2,231633591 | 0,325 | 0,786 | 4,14E-175 | 1 |
| *Pfas* | 1,17E-178 | -2,393082303 | 0,223 | 0,734 | 1,39E-174 | 1 |
| *CG7720* | 7,48E-177 | 2,386192092 | 0,674 | 0,199 | 8,93E-173 | 1 |
| *CG42788* | 4,81E-174 | 2,471849449 | 0,771 | 0,34 | 5,74E-170 | 1 |
| *bun* | 6,92E-174 | 1,812011993 | 0,964 | 0,784 | 8,25E-170 | 1 |
| *Acsl* | 7,44E-173 | 1,791833491 | 0,827 | 0,423 | 8,88E-169 | 1 |
| *Gpdh1* | 1,27E-172 | 2,138803904 | 0,668 | 0,213 | 1,52E-168 | 1 |
| *CG7470* | 3,84E-172 | 1,258541255 | 0,956 | 0,616 | 4,58E-168 | 1 |
| *sug* | 6,52E-172 | 2,335559474 | 0,639 | 0,17 | 7,77E-168 | 1 |
| *CCHa2* | 6,21E-171 | 2,052475931 | 0,564 | 0,114 | 7,40E-167 | 1 |
| *GstE1* | 1,64E-165 | -2,98721268 | 0,034 | 0,545 | 1,96E-161 | 1 |
| *CG14207* | 5,40E-165 | -2,331066787 | 0,265 | 0,745 | 6,44E-161 | 1 |
| *Ubx* | 3,02E-164 | 1,190324941 | 0,986 | 0,793 | 3,60E-160 | 1 |
| *CG5953* | 2,02E-163 | -2,381228188 | 0,418 | 0,808 | 2,41E-159 | 1 |
| *AcCoAS* | 5,37E-163 | 1,629352671 | 0,889 | 0,583 | 6,41E-159 | 1 |
| *eIF2beta* | 1,99E-162 | -2,466070653 | 0,207 | 0,686 | 2,37E-158 | 1 |
| *CG1213* | 4,18E-162 | 2,033728296 | 0,645 | 0,191 | 4,98E-158 | 1 |
| *CaMKI* | 5,04E-161 | 0,904942942 | 0,994 | 0,946 | 6,01E-157 | 1 |
| *CG10082* | 1,10E-158 | 1,671279874 | 0,846 | 0,485 | 1,31E-154 | 1 |
| *CG8745* | 2,58E-158 | 2,434027011 | 0,66 | 0,223 | 3,07E-154 | 1 |
| *Act5C* | 3,54E-155 | -2,183841477 | 0,225 | 0,704 | 4,22E-151 | 1 |
| *Xrp1* | 7,47E-155 | -1,393167977 | 0,828 | 0,952 | 8,91E-151 | 1 |
| *Ubi-p63E* | 7,47E-155 | -2,227207753 | 0,327 | 0,753 | 8,91E-151 | 1 |
| *Men* | 5,19E-154 | 1,88198364 | 0,868 | 0,576 | 6,19E-150 | 1 |
| *vir-1* | 1,40E-153 | -2,025323696 | 0,517 | 0,847 | 1,67E-149 | 1 |
| *CG12795* | 2,31E-153 | -3,06555317 | 0,026 | 0,509 | 2,76E-149 | 1 |
| *CG45050* | 3,75E-153 | -1,532756464 | 0,814 | 0,946 | 4,48E-149 | 1 |
| *Abl* | 4,22E-153 | -2,805991287 | 0,047 | 0,532 | 5,03E-149 | 1 |
| *bru1* | 1,92E-152 | 1,151993629 | 0,954 | 0,724 | 2,29E-148 | 1 |
| *CG13315* | 9,92E-151 | 1,556077475 | 0,956 | 0,756 | 1,18E-146 | 1 |
| *cv-c* | 1,60E-150 | -1,845739098 | 0,356 | 0,795 | 1,90E-146 | 1 |
| *Syp* | 7,16E-150 | 1,162891412 | 0,958 | 0,744 | 8,54E-146 | 1 |
| *CG10383* | 2,97E-148 | -2,582648287 | 0,146 | 0,631 | 3,54E-144 | 1 |
| *Egfr* | 1,38E-147 | 1,258631841 | 0,944 | 0,69 | 1,65E-143 | 1 |
| *CG8034* | 9,06E-147 | 1,850377493 | 0,826 | 0,496 | 1,08E-142 | 1 |
| *Idh* | 2,31E-146 | 1,800118448 | 0,697 | 0,283 | 2,76E-142 | 1 |
| *PhKgamma* | 1,57E-144 | 1,461591623 | 0,834 | 0,492 | 1,87E-140 | 1 |
| *Nc73EF* | 6,56E-143 | 1,415172331 | 0,864 | 0,549 | 7,82E-139 | 1 |
| *CenG1A* | 1,09E-142 | -2,80401486 | 0,25 | 0,694 | 1,30E-138 | 1 |
| *puc* | 4,36E-142 | -1,878952684 | 0,442 | 0,817 | 5,20E-138 | 1 |
| *oys* | 2,91E-141 | -2,651198532 | 0,026 | 0,482 | 3,47E-137 | 1 |
| *Helz* | 3,96E-140 | -1,574693743 | 0,842 | 0,942 | 4,73E-136 | 1 |

| | | | | | | |
|---|---|---|---|---|---|---|
| *MtnA* | 1,92E-139 | -2,145205107 | 0,259 | 0,719 | 2,29E-135 | 1 |
| *AkhR* | 6,28E-138 | 1,668387235 | 0,6 | 0,181 | 7,50E-134 | 1 |
| *apolpp* | 1,57E-137 | 0,921326065 | 0,964 | 0,725 | 1,87E-133 | 1 |
| *CG1648* | 2,31E-137 | 1,626411909 | 0,844 | 0,496 | 2,76E-133 | 1 |
| *CG15096* | 9,13E-137 | 1,669542815 | 0,52 | 0,121 | 1,09E-132 | 1 |
| *CG1673* | 1,42E-136 | -1,853976912 | 0,303 | 0,725 | 1,70E-132 | 1 |
| *CG17841* | 8,22E-136 | 1,534251041 | 0,737 | 0,348 | 9,80E-132 | 1 |
| *CG7766* | 2,59E-135 | 1,383149312 | 0,81 | 0,456 | 3,09E-131 | 1 |
| *AdipoR* | 2,53E-134 | 1,418191611 | 0,792 | 0,413 | 3,02E-130 | 1 |
| *mamo* | 3,75E-134 | 1,435381842 | 0,813 | 0,429 | 4,47E-130 | 1 |
| *Eglp4* | 5,46E-134 | 1,747461472 | 0,648 | 0,233 | 6,51E-130 | 1 |
| *CG16898* | 6,33E-133 | -2,392832022 | 0,133 | 0,586 | 7,55E-129 | 1 |
| *Frl* | 7,06E-133 | -2,166028302 | 0,223 | 0,657 | 8,42E-129 | 1 |
| *LRR* | 1,03E-132 | -2,082973495 | 0,224 | 0,67 | 1,23E-128 | 1 |
| *if* | 5,44E-132 | 1,905660683 | 0,62 | 0,214 | 6,48E-128 | 1 |
| *cact* | 4,70E-131 | -1,808506883 | 0,382 | 0,768 | 5,60E-127 | 1 |
| *IP3K1* | 4,79E-131 | -2,0535559 | 0,276 | 0,698 | 5,71E-127 | 1 |
| *rhea* | 7,45E-130 | -2,584365263 | 0,215 | 0,631 | 8,89E-126 | 1 |
| *Trxr-1* | 2,34E-128 | -2,172396702 | 0,095 | 0,536 | 2,79E-124 | 1 |
| *GEFmeso* | 1,75E-127 | -2,142933928 | 0,145 | 0,583 | 2,09E-123 | 1 |
| *Hers* | 2,41E-127 | -1,717270723 | 0,366 | 0,749 | 2,88E-123 | 1 |
| *Chd64* | 3,30E-127 | -2,178789318 | 0,166 | 0,598 | 3,94E-123 | 1 |
| *kay* | 7,00E-127 | -2,220735567 | 0,241 | 0,653 | 8,35E-123 | 1 |
| *Tret1-1* | 3,83E-125 | 1,57240269 | 0,791 | 0,449 | 4,57E-121 | 1 |
| *AdSS* | 1,13E-124 | -2,116676248 | 0,121 | 0,561 | 1,35E-120 | 1 |
| *Jabba* | 2,06E-124 | 1,740273796 | 0,542 | 0,157 | 2,45E-120 | 1 |
| *UGP* | 2,76E-124 | 1,689822468 | 0,681 | 0,295 | 3,30E-120 | 1 |
| *Snoo* | 5,74E-124 | 1,084721287 | 0,955 | 0,784 | 6,84E-120 | 1 |
| *CG3036* | 4,43E-123 | -1,632652382 | 0,367 | 0,791 | 5,28E-119 | 1 |
| *Fer1HCH* | 6,26E-122 | -1,381967341 | 0,624 | 0,89 | 7,47E-118 | 1 |
| *CG17124* | 1,57E-119 | 1,270308751 | 0,896 | 0,676 | 1,87E-115 | 1 |
| *Mob2* | 1,59E-119 | -1,647374215 | 0,579 | 0,838 | 1,89E-115 | 1 |
| *mtd* | 1,65E-119 | 1,358104619 | 0,878 | 0,6 | 1,96E-115 | 1 |
| *MTA1-like* | 3,30E-119 | -2,86523508 | 0,165 | 0,565 | 3,94E-115 | 1 |
| *GIIIspla2* | 4,31E-119 | -2,189108673 | 0,129 | 0,558 | 5,14E-115 | 1 |
| *TER94* | 4,26E-118 | -1,861588223 | 0,218 | 0,627 | 5,08E-114 | 1 |
| *Chchd2* | 5,27E-118 | -1,704363791 | 0,365 | 0,742 | 6,29E-114 | 1 |
| *Hsp26* | 1,70E-117 | -3,05830312 | 0,138 | 0,56 | 2,03E-113 | 1 |
| *CG8485* | 2,05E-117 | 1,789829669 | 0,484 | 0,131 | 2,45E-113 | 1 |
| *lola* | 8,84E-116 | 0,917388744 | 0,964 | 0,814 | 1,05E-111 | 1 |
| *mbf1* | 5,44E-115 | -2,168431937 | 0,074 | 0,486 | 6,49E-111 | 1 |
| *CG17108* | 6,77E-115 | 1,060532064 | 0,68 | 0,264 | 8,08E-111 | 1 |
| *spoon* | 1,07E-114 | 1,455230229 | 0,671 | 0,312 | 1,27E-110 | 1 |
| *CG42588* | 1,42E-114 | -2,243616442 | 0,048 | 0,451 | 1,69E-110 | 1 |
| *Stat92E* | 2,52E-114 | 1,480932693 | 0,801 | 0,486 | 3,01E-110 | 1 |
| *Hsc70-4* | 5,16E-114 | -1,444622723 | 0,552 | 0,832 | 6,15E-110 | 1 |
| *loco* | 8,62E-114 | -2,46566665 | 0,015 | 0,402 | 1,03E-109 | 1 |
| *phu* | 2,20E-113 | 1,740645346 | 0,328 | 0,036 | 2,63E-109 | 1 |
| *Akap200* | 3,61E-113 | 1,553187866 | 0,757 | 0,455 | 4,31E-109 | 1 |
| *trbl* | 5,27E-112 | -1,762898009 | 0,245 | 0,644 | 6,28E-108 | 1 |
| *CG7130* | 4,28E-111 | -2,482493606 | 0,015 | 0,396 | 5,10E-107 | 1 |
| *Pde6* | 1,90E-110 | 1,326059462 | 0,749 | 0,395 | 2,27E-106 | 1 |
| *CG6770* | 3,02E-110 | 0,941314839 | 0,987 | 0,886 | 3,61E-106 | 1 |
| *CG1578* | 3,83E-110 | 1,51910213 | 0,631 | 0,259 | 4,56E-106 | 1 |
| *BomT3* | 4,26E-110 | 1,126068767 | 0,695 | 0,295 | 5,08E-106 | 1 |
| *Pka-C1* | 9,02E-110 | -1,544778142 | 0,441 | 0,778 | 1,08E-105 | 1 |
| *CG4716* | 1,64E-109 | 0,990136683 | 0,952 | 0,811 | 1,96E-105 | 1 |
| *Hsp83* | 2,02E-109 | -1,760521093 | 0,532 | 0,834 | 2,42E-105 | 1 |
| *Bacc* | 2,70E-108 | 1,06621647 | 0,902 | 0,683 | 3,22E-104 | 1 |
| *CG8312* | 2,53E-107 | -2,660739209 | 0,067 | 0,454 | 3,01E-103 | 1 |
| *drpr* | 4,16E-107 | -1,91440328 | 0,195 | 0,585 | 4,96E-103 | 1 |

| | | | | | | |
|---|---|---|---|---|---|---|
| *CG13607* | 7,61E-107 | 1,461451944 | 0,382 | 0,07 | 9,07E-103 | 1 |
| *cher* | 5,83E-106 | -2,459319133 | 0,04 | 0,415 | 6,96E-102 | 1 |
| *Src64B* | 2,17E-105 | -2,068032023 | 0,298 | 0,651 | 2,59E-101 | 1 |
| *CG32521* | 2,20E-105 | 1,079664962 | 0,925 | 0,694 | 2,62E-101 | 1 |
| *Cyt-b5-r* | 3,14E-103 | 1,229204451 | 0,793 | 0,452 | 3,74E-99 | 1 |
| *alpha-Est9* | 9,80E-102 | 1,701170385 | 0,39 | 0,08 | 1,17E-97 | 1 |
| *LpR2* | 2,63E-101 | 1,051398825 | 0,957 | 0,793 | 3,14E-97 | 1 |
| *CG32647* | 4,39E-101 | 1,332867242 | 0,783 | 0,456 | 5,24E-97 | 1 |
| *Socs36E* | 6,61E-101 | -2,411777925 | 0,072 | 0,446 | 7,89E-97 | 1 |
| *CG8086* | 7,17E-99 | -2,123788283 | 0,075 | 0,447 | 8,56E-95 | 1 |
| *Hsp27* | 3,89E-98 | -3,500920991 | 0,096 | 0,462 | 4,64E-94 | 1 |
| *CG42524* | 1,44E-97 | 1,204600516 | 0,704 | 0,356 | 1,72E-93 | 1 |
| *AOX1* | 2,40E-97 | -2,100525165 | 0,088 | 0,455 | 2,86E-93 | 1 |
| *Tpr2* | 1,21E-96 | -1,491800853 | 0,36 | 0,714 | 1,44E-92 | 1 |
| *Rpn6* | 2,04E-96 | -1,668626128 | 0,134 | 0,515 | 2,43E-92 | 1 |
| *Fer2LCH* | 7,54E-96 | -1,283181341 | 0,569 | 0,831 | 8,99E-92 | 1 |
| *Sox102F* | 9,57E-96 | 0,939174608 | 0,901 | 0,672 | 1,14E-91 | 1 |
| *Vps13* | 1,88E-95 | -1,800618405 | 0,113 | 0,485 | 2,25E-91 | 1 |
| *Smg5* | 3,50E-95 | -1,777390761 | 0,246 | 0,603 | 4,17E-91 | 1 |
| *Hsc70Cb* | 3,95E-95 | -1,855306021 | 0,295 | 0,643 | 4,72E-91 | 1 |
| *Nmda1* | 5,00E-95 | -1,738064413 | 0,104 | 0,473 | 5,96E-91 | 1 |
| *ps* | 1,65E-93 | 0,701052748 | 0,996 | 0,978 | 1,97E-89 | 1 |
| *Drak* | 3,11E-92 | 1,55685793 | 0,79 | 0,539 | 3,71E-88 | 1 |
| *Gart* | 1,10E-90 | -1,325888371 | 0,464 | 0,765 | 1,31E-86 | 1 |
| *Srr* | 1,47E-90 | 1,585774031 | 0,408 | 0,107 | 1,76E-86 | 1 |
| *Pvr* | 7,52E-90 | -2,454220779 | 0,204 | 0,551 | 8,97E-86 | 1 |
| *gce* | 1,18E-89 | 1,326338449 | 0,687 | 0,383 | 1,41E-85 | 1 |
| *Gug* | 2,97E-89 | 1,048655902 | 0,816 | 0,557 | 3,54E-85 | 1 |
| *CG5151* | 7,17E-89 | 1,49820848 | 0,806 | 0,577 | 8,56E-85 | 1 |
| *Pde9* | 7,47E-89 | 1,045453538 | 0,892 | 0,673 | 8,91E-85 | 1 |
| *kdn* | 4,52E-88 | 1,471737637 | 0,518 | 0,19 | 5,39E-84 | 1 |
| *retm* | 4,69E-88 | 1,463938889 | 0,496 | 0,183 | 5,60E-84 | 1 |
| *CG44325* | 6,71E-88 | -1,749942608 | 0,086 | 0,432 | 8,01E-84 | 1 |
| *Hsp68* | 1,00E-87 | -2,744761721 | 0,071 | 0,411 | 1,20E-83 | 1 |
| *fbp* | 1,45E-87 | 1,33279003 | 0,531 | 0,211 | 1,73E-83 | 1 |
| *MFS14* | 9,30E-87 | 1,421430786 | 0,617 | 0,295 | 1,11E-82 | 1 |
| *bmm* | 5,83E-86 | -0,966698125 | 0,564 | 0,86 | 6,96E-82 | 1 |
| *chic* | 6,53E-86 | -1,771984967 | 0,19 | 0,53 | 7,79E-82 | 1 |
| *sra* | 7,41E-86 | -1,846289122 | 0,049 | 0,379 | 8,84E-82 | 1 |
| *elB* | 4,49E-85 | -1,966157599 | 0,085 | 0,423 | 5,36E-81 | 1 |
| *Rpn2* | 1,35E-84 | -1,637671708 | 0,08 | 0,419 | 1,61E-80 | 1 |
| *PKD* | 2,16E-84 | 1,430272025 | 0,501 | 0,188 | 2,58E-80 | 1 |
| *Piezo* | 6,68E-84 | -1,797886041 | 0,17 | 0,52 | 7,97E-80 | 1 |
| *CG14990* | 4,50E-83 | 1,399776395 | 0,351 | 0,082 | 5,37E-79 | 1 |
| *Tsf1* | 7,13E-83 | 1,223072438 | 0,512 | 0,187 | 8,50E-79 | 1 |
| *PCB* | 7,48E-83 | 1,015717712 | 0,814 | 0,512 | 8,93E-79 | 1 |
| *Ndae1* | 3,95E-82 | 1,722268773 | 0,539 | 0,24 | 4,71E-78 | 1 |
| *Pmp70* | 1,04E-81 | 1,43307853 | 0,474 | 0,171 | 1,24E-77 | 1 |
| *pyr* | 1,36E-81 | -1,939060272 | 0,077 | 0,405 | 1,62E-77 | 1 |
| *Rel* | 3,52E-81 | -1,303683836 | 0,307 | 0,65 | 4,20E-77 | 1 |
| *CG7920* | 3,67E-81 | 1,269886576 | 0,506 | 0,193 | 4,38E-77 | 1 |
| *Gpo1* | 4,26E-81 | 1,380275419 | 0,487 | 0,177 | 5,08E-77 | 1 |
| *CG16926* | 1,37E-80 | 1,052542271 | 0,8 | 0,536 | 1,63E-76 | 1 |
| *pcs* | 2,65E-80 | -1,964721586 | 0,204 | 0,532 | 3,17E-76 | 1 |
| *klu* | 3,03E-79 | -1,416161577 | 0,258 | 0,603 | 3,62E-75 | 1 |
| *Prosalpha3* | 3,59E-79 | -1,760234278 | 0,06 | 0,371 | 4,29E-75 | 1 |
| *eff* | 5,48E-79 | -1,350263173 | 0,391 | 0,688 | 6,54E-75 | 1 |
| *dnc* | 1,70E-78 | 0,909603446 | 0,893 | 0,683 | 2,03E-74 | 1 |
| *AspRS* | 3,30E-78 | -1,808215951 | 0,029 | 0,325 | 3,93E-74 | 1 |
| *whd* | 1,29E-77 | 1,181796032 | 0,72 | 0,417 | 1,54E-73 | 1 |
| *Droj2* | 2,63E-77 | -1,410324972 | 0,192 | 0,525 | 3,14E-73 | 1 |

| | | | | | | |
|---|---|---|---|---|---|---|
| *CCT3* | 3,84E-77 | -1,906064675 | 0,041 | 0,34 | 4,58E-73 | 1 |
| *Atg1* | 1,36E-76 | -1,670699124 | 0,293 | 0,608 | 1,62E-72 | 1 |
| *cic* | 3,82E-76 | 0,932152248 | 0,819 | 0,589 | 4,55E-72 | 1 |
| *IM4* | 6,30E-76 | 0,81334175 | 0,525 | 0,2 | 7,51E-72 | 1 |
| *Rpn5* | 7,62E-76 | -1,613432609 | 0,061 | 0,366 | 9,10E-72 | 1 |
| *Cyp4p1* | 5,00E-75 | -1,609787789 | 0,097 | 0,415 | 5,97E-71 | 1 |
| *zfh1* | 9,98E-75 | -1,588387415 | 0,164 | 0,494 | 1,19E-70 | 1 |
| *CG5955* | 1,32E-74 | -1,722562168 | 0,015 | 0,295 | 1,58E-70 | 1 |
| *Gdap2* | 2,03E-74 | -1,516191629 | 0,189 | 0,514 | 2,42E-70 | 1 |
| *CG13360* | 3,30E-74 | 1,365741669 | 0,317 | 0,071 | 3,93E-70 | 1 |
| *Trx-2* | 8,76E-74 | -1,834948938 | 0,085 | 0,39 | 1,05E-69 | 1 |
| *Lsd-1* | 6,51E-73 | 1,46588597 | 0,507 | 0,211 | 7,77E-69 | 1 |
| *scyl* | 6,87E-73 | -1,127432466 | 0,672 | 0,875 | 8,20E-69 | 1 |
| *ImpL2* | 8,43E-73 | -1,801143257 | 0,051 | 0,344 | 1,01E-68 | 1 |
| *CG42668* | 1,46E-72 | -1,246428909 | 0,401 | 0,696 | 1,74E-68 | 1 |
| *CG4629* | 1,50E-72 | 1,491765795 | 0,437 | 0,162 | 1,79E-68 | 1 |
| *Rpn10* | 2,68E-72 | -1,537023305 | 0,052 | 0,346 | 3,19E-68 | 1 |
| *Pdha* | 6,34E-72 | 1,217383454 | 0,422 | 0,146 | 7,57E-68 | 1 |
| *Prosbeta7* | 1,31E-71 | -1,549706343 | 0,056 | 0,345 | 1,57E-67 | 1 |
| *Apoltp* | 2,44E-71 | -1,065078201 | 0,658 | 0,86 | 2,91E-67 | 1 |
| *CG17646* | 2,74E-71 | 0,953744812 | 0,93 | 0,813 | 3,27E-67 | 1 |
| *Idgf1* | 3,54E-71 | -1,562721902 | 0,083 | 0,384 | 4,23E-67 | 1 |
| *CG8389* | 3,42E-70 | 1,6264232 | 0,307 | 0,075 | 4,08E-66 | 1 |
| *pAbp* | 7,00E-70 | -0,980461135 | 0,685 | 0,86 | 8,35E-66 | 1 |
| *Rad23* | 9,91E-70 | -1,39850126 | 0,123 | 0,435 | 1,18E-65 | 1 |
| *Dmtn* | 1,62E-69 | 0,944921139 | 0,865 | 0,692 | 1,93E-65 | 1 |
| *cwo* | 1,77E-69 | 0,832666033 | 0,938 | 0,85 | 2,11E-65 | 1 |
| *CG32369* | 4,49E-69 | -1,265362343 | 0,5 | 0,758 | 5,35E-65 | 1 |
| *Hipk* | 5,64E-69 | -1,062459934 | 0,528 | 0,765 | 6,73E-65 | 1 |
| *CG18135* | 1,72E-68 | 0,854640532 | 0,868 | 0,649 | 2,05E-64 | 1 |
| *Prosalpha6* | 2,26E-68 | -1,558320892 | 0,042 | 0,316 | 2,70E-64 | 1 |
| *CG11791* | 5,63E-68 | -1,509851003 | 0,227 | 0,534 | 6,71E-64 | 1 |
| *Prosbeta4* | 1,06E-67 | -1,515628769 | 0,054 | 0,335 | 1,26E-63 | 1 |
| *Xbp1* | 3,56E-67 | -1,127081095 | 0,378 | 0,67 | 4,25E-63 | 1 |
| *CG16799* | 4,12E-67 | 1,340493222 | 0,356 | 0,108 | 4,92E-63 | 1 |
| *c11.1* | 9,74E-67 | -1,486324649 | 0,059 | 0,339 | 1,16E-62 | 1 |
| *Hsp70Bc* | 1,36E-66 | -1,946290834 | 0,011 | 0,262 | 1,62E-62 | 1 |
| *NFAT* | 2,93E-66 | 0,905175831 | 0,83 | 0,646 | 3,49E-62 | 1 |
| *AttB* | 5,41E-66 | -2,364325882 | 0,021 | 0,277 | 6,46E-62 | 1 |
| *ppl* | 8,40E-66 | 1,277401385 | 0,401 | 0,143 | 1,00E-61 | 1 |
| *DOR* | 9,90E-66 | 1,17395365 | 0,76 | 0,534 | 1,18E-61 | 1 |
| *dsx* | 1,30E-65 | 0,823473301 | 0,732 | 0,434 | 1,55E-61 | 1 |
| *cue* | 2,76E-65 | 1,25842065 | 0,321 | 0,085 | 3,29E-61 | 1 |
| *atk* | 4,47E-65 | -1,603766051 | 0,009 | 0,254 | 5,33E-61 | 1 |
| *Myo31DF* | 4,87E-65 | -2,064370498 | 0,05 | 0,318 | 5,81E-61 | 1 |
| *frma* | 6,71E-65 | 1,180736813 | 0,462 | 0,195 | 8,00E-61 | 1 |
| *HDAC6* | 1,39E-64 | 1,078743101 | 0,526 | 0,25 | 1,66E-60 | 1 |
| *Ufd4* | 1,69E-64 | -1,382634595 | 0,149 | 0,442 | 2,01E-60 | 1 |
| *cbt* | 5,03E-64 | 1,202211445 | 0,459 | 0,188 | 6,00E-60 | 1 |
| *l(1)G0196* | 2,16E-63 | 1,025118531 | 0,618 | 0,342 | 2,58E-59 | 1 |
| *Spn88Eb* | 3,65E-63 | -1,451939737 | 0,03 | 0,286 | 4,35E-59 | 1 |
| *Rpn13* | 5,30E-63 | -1,335902641 | 0,087 | 0,367 | 6,32E-59 | 1 |
| *Rpt5* | 9,05E-63 | -1,430504115 | 0,039 | 0,297 | 1,08E-58 | 1 |
| *CG41378* | 1,39E-62 | 0,963656451 | 0,716 | 0,46 | 1,66E-58 | 1 |
| *teq* | 5,53E-62 | 1,061599224 | 0,427 | 0,163 | 6,60E-58 | 1 |
| *CG6040* | 5,97E-62 | 1,341273213 | 0,308 | 0,082 | 7,12E-58 | 1 |
| *LamC* | 6,45E-62 | -1,560277748 | 0,082 | 0,356 | 7,70E-58 | 1 |
| *Sem1* | 1,19E-61 | -1,469327833 | 0,045 | 0,3 | 1,42E-57 | 1 |
| *MRP* | 1,19E-61 | -2,016198016 | 0,241 | 0,519 | 1,42E-57 | 1 |
| *px* | 1,70E-61 | 0,826955348 | 0,854 | 0,672 | 2,03E-57 | 1 |
| *CG10527* | 2,71E-61 | -1,548490534 | 0,047 | 0,303 | 3,24E-57 | 1 |

| | | | | | | |
|---|---|---|---|---|---|---|
| *CG9399* | 4,10E-61 | 1,175965022 | 0,285 | 0,072 | 4,89E-57 | 1 |
| *CG31635* | 4,58E-61 | 1,017093477 | 0,717 | 0,47 | 5,46E-57 | 1 |
| *nkd* | 5,18E-61 | -1,465853495 | 0,252 | 0,544 | 6,18E-57 | 1 |
| *CG1358* | 5,99E-61 | 1,384560014 | 0,368 | 0,125 | 7,15E-57 | 1 |
| *AdSL* | 6,11E-61 | -1,427306087 | 0,048 | 0,308 | 7,29E-57 | 1 |
| *pst* | 7,05E-61 | -1,114429655 | 0,402 | 0,673 | 8,41E-57 | 1 |
| *Ork1* | 8,82E-61 | 1,186890244 | 0,52 | 0,247 | 1,05E-56 | 1 |
| *CG3764* | 1,42E-60 | 1,221274181 | 0,474 | 0,212 | 1,70E-56 | 1 |
| *Svil* | 2,24E-60 | -1,379211528 | 0,377 | 0,633 | 2,67E-56 | 1 |
| *bnl* | 2,48E-60 | -2,213555209 | 0,139 | 0,421 | 2,96E-56 | 1 |
| *baf* | 3,39E-60 | -1,703342924 | 0,039 | 0,287 | 4,04E-56 | 1 |
| *DnaJ-1* | 3,75E-60 | -1,233264207 | 0,341 | 0,629 | 4,48E-56 | 1 |
| *CG6067* | 4,32E-60 | 1,077503901 | 0,515 | 0,241 | 5,16E-56 | 1 |
| *baz* | 6,12E-60 | -1,48968969 | 0,063 | 0,321 | 7,30E-56 | 1 |
| *CAH7* | 2,69E-59 | 1,380965271 | 0,315 | 0,093 | 3,21E-55 | 1 |
| *AGBE* | 5,19E-59 | 1,212689729 | 0,257 | 0,059 | 6,19E-55 | 1 |
| *Gadd45* | 8,33E-59 | -1,496498249 | 0,06 | 0,317 | 9,93E-55 | 1 |
| *CG31678* | 1,37E-58 | -1,494847498 | 0,05 | 0,304 | 1,64E-54 | 1 |
| *Atg18b* | 1,64E-58 | -1,322556665 | 0,25 | 0,539 | 1,95E-54 | 1 |
| *CG5958* | 2,50E-58 | -1,471762652 | 0,225 | 0,505 | 2,99E-54 | 1 |
| *Lsd-2* | 2,97E-58 | 0,79955559 | 0,899 | 0,779 | 3,54E-54 | 1 |
| *l(3)80Fg* | 6,74E-58 | 0,839139228 | 0,762 | 0,537 | 8,04E-54 | 1 |
| *Blimp-1* | 9,57E-58 | 0,86532605 | 0,823 | 0,638 | 1,14E-53 | 1 |
| *hdc* | 2,26E-57 | -1,728423472 | 0,035 | 0,272 | 2,70E-53 | 1 |
| *Cyp6d5* | 2,39E-57 | 1,241046362 | 0,519 | 0,262 | 2,85E-53 | 1 |
| *Cyp28a5* | 2,68E-57 | -1,434157498 | 0,024 | 0,257 | 3,20E-53 | 1 |
| *Rpn1* | 4,56E-57 | -1,306302651 | 0,064 | 0,32 | 5,43E-53 | 1 |
| *Atg8a* | 6,78E-57 | -0,990161714 | 0,557 | 0,761 | 8,09E-53 | 1 |
| *jvl* | 1,04E-56 | -0,979535072 | 0,627 | 0,81 | 1,25E-52 | 1 |
| *EDTP* | 1,09E-56 | 1,144943318 | 0,508 | 0,256 | 1,30E-52 | 1 |
| *wrd* | 1,45E-56 | 0,936213937 | 0,609 | 0,357 | 1,73E-52 | 1 |
| *Gdh* | 1,73E-56 | -1,436537464 | 0,197 | 0,477 | 2,06E-52 | 1 |
| *GstE8* | 4,71E-56 | -1,507122984 | 0,041 | 0,282 | 5,62E-52 | 1 |
| *Lk6* | 5,79E-56 | -0,757797897 | 0,842 | 0,923 | 6,91E-52 | 1 |
| *Desat1* | 6,29E-56 | 0,605400256 | 0,971 | 0,854 | 7,50E-52 | 1 |
| *mdy* | 1,43E-55 | 1,149848409 | 0,309 | 0,091 | 1,70E-51 | 1 |
| *AdamTS-A* | 1,69E-55 | 1,45256095 | 0,436 | 0,19 | 2,02E-51 | 1 |
| *CycG* | 2,08E-55 | -0,64175777 | 0,922 | 0,96 | 2,48E-51 | 1 |
| *CG8665* | 2,33E-55 | 1,505086512 | 0,359 | 0,13 | 2,78E-51 | 1 |
| *CG5910* | 2,43E-55 | 1,006819173 | 0,573 | 0,312 | 2,90E-51 | 1 |
| *Gprk2* | 9,24E-55 | -1,057752298 | 0,566 | 0,76 | 1,10E-50 | 1 |
| *CG15293* | 9,73E-55 | 0,986602265 | 0,616 | 0,364 | 1,16E-50 | 1 |
| *Cyp6w1* | 1,78E-54 | 1,227103176 | 0,371 | 0,14 | 2,13E-50 | 1 |
| *Cam* | 2,02E-54 | -0,836845204 | 0,697 | 0,881 | 2,41E-50 | 1 |
| *Irc* | 5,59E-54 | -1,059160048 | 0,335 | 0,604 | 6,67E-50 | 1 |
| *Gk2* | 7,79E-54 | 1,155979278 | 0,323 | 0,107 | 9,30E-50 | 1 |
| *Exn* | 7,90E-54 | -1,131625762 | 0,332 | 0,593 | 9,43E-50 | 1 |
| *Cyp309a1* | 2,54E-53 | -1,21938218 | 0,071 | 0,321 | 3,02E-49 | 1 |
| *NUCB1* | 2,62E-53 | -1,525123344 | 0,066 | 0,304 | 3,13E-49 | 1 |
| *Lip4* | 2,83E-53 | -1,266445212 | 0,221 | 0,5 | 3,38E-49 | 1 |
| *CG32767* | 3,17E-53 | 0,978640894 | 0,556 | 0,304 | 3,78E-49 | 1 |
| *CG3168* | 4,66E-53 | -1,344642882 | 0,12 | 0,38 | 5,55E-49 | 1 |
| *Pdfr* | 4,68E-53 | 0,912273625 | 0,649 | 0,393 | 5,58E-49 | 1 |
| *CG17549* | 5,65E-53 | 1,057202343 | 0,537 | 0,292 | 6,73E-49 | 1 |
| *Prosalpha2* | 8,71E-53 | -1,335894669 | 0,067 | 0,306 | 1,04E-48 | 1 |
| *aay* | 1,00E-52 | -0,947099312 | 0,364 | 0,659 | 1,20E-48 | 1 |
| *shrb* | 1,36E-52 | -1,248343757 | 0,117 | 0,375 | 1,63E-48 | 1 |
| *CG1416* | 1,88E-52 | -1,361416473 | 0,035 | 0,257 | 2,25E-48 | 1 |
| *CG6870* | 2,03E-52 | -1,591386798 | 0,049 | 0,281 | 2,42E-48 | 1 |
| *Smr* | 3,95E-52 | 0,692062762 | 0,882 | 0,683 | 4,72E-48 | 1 |
| *CG9932* | 4,32E-52 | 0,578146859 | 0,987 | 0,929 | 5,15E-48 | 1 |

| | | | | | | |
|---|---|---|---|---|---|---|
| *Prosbeta5* | 9,09E-52 | -1,297796929 | 0,06 | 0,294 | 1,08E-47 | 1 |
| *Prosalpha7* | 3,16E-51 | -1,229543286 | 0,063 | 0,295 | 3,77E-47 | 1 |
| *Prosalpha5* | 3,27E-51 | -1,275268586 | 0,042 | 0,265 | 3,90E-47 | 1 |
| *Smg6* | 3,78E-51 | -1,31762885 | 0,066 | 0,302 | 4,51E-47 | 1 |
| *Rpn3* | 5,94E-51 | -1,224101978 | 0,05 | 0,278 | 7,09E-47 | 1 |
| *CG6115* | 1,04E-50 | -1,03212371 | 0,243 | 0,52 | 1,24E-46 | 1 |
| *Msp300* | 2,58E-50 | 0,591945103 | 0,952 | 0,847 | 3,08E-46 | 1 |
| *Rpn9* | 5,35E-50 | -1,301851958 | 0,039 | 0,255 | 6,39E-46 | 1 |
| *dos* | 5,94E-50 | -1,4369856 | 0,04 | 0,259 | 7,09E-46 | 1 |
| *fus* | 8,65E-50 | 0,918778122 | 0,557 | 0,322 | 1,03E-45 | 1 |
| *lilli* | 9,84E-50 | 0,658831902 | 0,853 | 0,671 | 1,17E-45 | 1 |
| *rgn* | 1,21E-49 | -1,69766611 | 0,302 | 0,538 | 1,45E-45 | 1 |
| *LRP1* | 1,24E-49 | 1,100019689 | 0,587 | 0,346 | 1,47E-45 | 1 |
| *Acbp2* | 1,93E-49 | 1,063989615 | 0,476 | 0,233 | 2,30E-45 | 1 |
| *GstE7* | 2,02E-49 | -1,306119868 | 0,038 | 0,255 | 2,41E-45 | 1 |
| *tkv* | 2,10E-49 | 1,108830196 | 0,545 | 0,318 | 2,50E-45 | 1 |
| *CG16756* | 2,51E-49 | 1,031091475 | 0,299 | 0,096 | 2,99E-45 | 1 |
| *Ctl2* | 3,12E-49 | -1,282472918 | 0,041 | 0,259 | 3,73E-45 | 1 |
| *Rpt3* | 3,15E-49 | -1,194762813 | 0,053 | 0,279 | 3,76E-45 | 1 |
| *alt* | 4,14E-49 | -1,267836728 | 0,188 | 0,442 | 4,94E-45 | 1 |
| *CG2233* | 1,09E-48 | 0,537517223 | 0,971 | 0,887 | 1,30E-44 | 1 |
| *Galk* | 1,30E-48 | 0,968684152 | 0,651 | 0,429 | 1,55E-44 | 1 |
| *Ist1* | 1,37E-48 | -1,142786428 | 0,13 | 0,377 | 1,64E-44 | 1 |
| *Tlk* | 1,38E-48 | 0,777282921 | 0,716 | 0,491 | 1,65E-44 | 1 |
| *Nadsyn* | 1,83E-48 | 1,07556035 | 0,435 | 0,213 | 2,18E-44 | 1 |
| *mld* | 3,03E-48 | -1,413269463 | 0,138 | 0,391 | 3,61E-44 | 1 |
| *myo* | 3,08E-48 | 0,925229827 | 0,547 | 0,312 | 3,68E-44 | 1 |
| *msn* | 3,87E-48 | -1,010250229 | 0,434 | 0,66 | 4,61E-44 | 1 |
| *Ire1* | 4,52E-48 | -1,112978897 | 0,146 | 0,401 | 5,39E-44 | 1 |
| *DppIII* | 4,78E-48 | -1,220594614 | 0,05 | 0,265 | 5,71E-44 | 1 |
| *CG9691* | 7,71E-48 | 0,89225392 | 0,507 | 0,263 | 9,20E-44 | 1 |
| *LPCAT* | 7,90E-48 | -1,262571634 | 0,056 | 0,278 | 9,42E-44 | 1 |
| *CG11594* | 1,45E-47 | 1,087320898 | 0,346 | 0,135 | 1,73E-43 | 1 |
| *MCPH1* | 1,48E-47 | -1,198431267 | 0,149 | 0,397 | 1,76E-43 | 1 |
| *CG31704* | 2,26E-47 | -1,317917933 | 0,071 | 0,295 | 2,70E-43 | 1 |
| *Rpn8* | 2,33E-47 | -1,174543538 | 0,045 | 0,257 | 2,77E-43 | 1 |
| *Ntan1* | 3,00E-47 | -1,004361358 | 0,45 | 0,682 | 3,57E-43 | 1 |
| *CG15099* | 3,27E-47 | -1,144457751 | 0,132 | 0,378 | 3,90E-43 | 1 |
| *Hsc70-5* | 3,99E-47 | -1,235195095 | 0,119 | 0,362 | 4,76E-43 | 1 |
| *ALiX* | 5,24E-47 | -1,129751514 | 0,072 | 0,3 | 6,25E-43 | 1 |
| *sima* | 6,11E-47 | 0,593626941 | 0,967 | 0,907 | 7,29E-43 | 1 |
| *SCaMC* | 1,42E-46 | -0,764493477 | 0,649 | 0,822 | 1,69E-42 | 1 |
| *h* | 1,53E-46 | -1,069181652 | 0,486 | 0,707 | 1,82E-42 | 1 |
| *CG32486* | 1,54E-46 | 0,924312876 | 0,618 | 0,406 | 1,84E-42 | 1 |
| *CG43340* | 4,40E-46 | 0,974722891 | 0,563 | 0,342 | 5,25E-42 | 1 |
| *CG12290* | 1,11E-45 | -1,445179445 | 0,119 | 0,355 | 1,33E-41 | 1 |
| *chb* | 1,14E-45 | -1,27499194 | 0,135 | 0,372 | 1,36E-41 | 1 |
| *MCU* | 1,18E-45 | -1,243015666 | 0,126 | 0,364 | 1,41E-41 | 1 |
| *Ilp6* | 1,36E-45 | 1,158660816 | 0,461 | 0,236 | 1,62E-41 | 1 |
| *caps* | 1,48E-45 | -1,036063527 | 0,334 | 0,583 | 1,76E-41 | 1 |
| *CG17278* | 3,09E-45 | -1,183129249 | 0,132 | 0,37 | 3,69E-41 | 1 |
| *Rpt6* | 5,37E-45 | -1,15815792 | 0,058 | 0,269 | 6,40E-41 | 1 |
| *CG5966* | 2,13E-44 | -1,067004642 | 0,28 | 0,536 | 2,55E-40 | 1 |
| *wdp* | 4,27E-44 | -1,232135557 | 0,151 | 0,393 | 5,09E-40 | 1 |
| *Arf79F* | 8,12E-44 | -0,984542777 | 0,33 | 0,56 | 9,69E-40 | 1 |
| *CG6330* | 9,61E-44 | -1,174798113 | 0,135 | 0,374 | 1,15E-39 | 1 |
| *CHES-1-like* | 1,40E-43 | 0,89905754 | 0,619 | 0,415 | 1,67E-39 | 1 |
| *Jafrac1* | 1,46E-43 | -1,001890097 | 0,227 | 0,473 | 1,74E-39 | 1 |
| *Chmp1* | 1,90E-43 | -1,304587169 | 0,079 | 0,291 | 2,26E-39 | 1 |
| *CG33493* | 2,04E-43 | 1,058562046 | 0,364 | 0,155 | 2,44E-39 | 1 |
| *step* | 2,08E-43 | -1,014793166 | 0,27 | 0,518 | 2,48E-39 | 1 |

| | | | | | | |
|---|---|---|---|---|---|---|
| *RhoGAP71E* | 2,34E-43 | -1,175411622 | 0,267 | 0,505 | 2,79E-39 | 1 |
| *Caper* | 2,53E-43 | 0,755296948 | 0,781 | 0,617 | 3,02E-39 | 1 |
| *CG13868* | 2,56E-43 | 0,67170554 | 0,841 | 0,7 | 3,06E-39 | 1 |
| *CG3348* | 2,63E-43 | 1,874859949 | 0,296 | 0,11 | 3,14E-39 | 1 |
| *pum* | 2,79E-43 | 0,596405061 | 0,694 | 0,461 | 3,33E-39 | 1 |
| *Cyp18a1* | 3,49E-43 | 1,105219663 | 0,313 | 0,118 | 4,17E-39 | 1 |
| *ScsbetaA* | 4,10E-43 | 0,798373487 | 0,627 | 0,41 | 4,89E-39 | 1 |
| *Hsp60A* | 7,95E-43 | -1,129070009 | 0,095 | 0,314 | 9,49E-39 | 1 |
| *Pisd* | 1,05E-42 | 0,988127047 | 0,269 | 0,088 | 1,25E-38 | 1 |
| *red* | 1,25E-42 | -1,317835147 | 0,066 | 0,273 | 1,49E-38 | 1 |
| *poe* | 1,28E-42 | -1,070597838 | 0,179 | 0,419 | 1,53E-38 | 1 |
| *Cyp6g1* | 2,20E-42 | 0,971998861 | 0,544 | 0,333 | 2,62E-38 | 1 |
| *CG7945* | 2,58E-42 | -1,419481583 | 0,095 | 0,308 | 3,08E-38 | 1 |
| *aralar1* | 2,65E-42 | -0,775246332 | 0,489 | 0,706 | 3,16E-38 | 1 |
| *Spn43Ab* | 4,99E-42 | 0,896175817 | 0,511 | 0,287 | 5,95E-38 | 1 |
| *Agpat3* | 5,84E-42 | 0,966381486 | 0,312 | 0,12 | 6,97E-38 | 1 |
| *SP1173* | 7,20E-42 | -1,376993095 | 0,058 | 0,259 | 8,59E-38 | 1 |
| *GstE12* | 8,05E-42 | 0,873896336 | 0,395 | 0,183 | 9,60E-38 | 1 |
| *CCT2* | 9,15E-42 | -1,163827442 | 0,11 | 0,329 | 1,09E-37 | 1 |
| *CG15120* | 1,05E-41 | 1,020966288 | 0,258 | 0,084 | 1,26E-37 | 1 |
| *Gclm* | 1,34E-41 | -1,332194464 | 0,206 | 0,438 | 1,60E-37 | 1 |
| *Rbfox1* | 2,15E-41 | 0,792773921 | 0,658 | 0,446 | 2,56E-37 | 1 |
| *MFS17* | 3,16E-41 | 0,625057647 | 0,812 | 0,638 | 3,77E-37 | 1 |
| *CG32425* | 3,50E-41 | 0,950204745 | 0,475 | 0,262 | 4,17E-37 | 1 |
| *CG34054* | 6,19E-41 | -1,101706314 | 0,055 | 0,256 | 7,38E-37 | 1 |
| *uex* | 8,76E-41 | 0,652846262 | 0,786 | 0,616 | 1,05E-36 | 1 |
| *ATPCL* | 1,20E-40 | 0,993070135 | 0,553 | 0,34 | 1,43E-36 | 1 |
| *EcR* | 1,34E-40 | 0,855660611 | 0,606 | 0,403 | 1,60E-36 | 1 |
| *zormin* | 1,56E-40 | -1,036349097 | 0,251 | 0,489 | 1,86E-36 | 1 |
| *Plod* | 2,78E-40 | -1,143605066 | 0,135 | 0,356 | 3,31E-36 | 1 |
| *Zasp52* | 2,99E-40 | 1,250713218 | 0,296 | 0,116 | 3,57E-36 | 1 |
| *Nep4* | 3,07E-40 | -1,207388482 | 0,08 | 0,288 | 3,66E-36 | 1 |
| *Pak3* | 3,10E-40 | -1,097786547 | 0,141 | 0,361 | 3,70E-36 | 1 |
| *GstE9* | 4,75E-40 | -1,131932718 | 0,109 | 0,325 | 5,66E-36 | 1 |
| *Swip-1* | 5,95E-40 | -0,98027016 | 0,238 | 0,481 | 7,09E-36 | 1 |
| *RasGAP1* | 6,72E-40 | -1,135617071 | 0,131 | 0,35 | 8,01E-36 | 1 |
| *larp* | 7,39E-40 | 0,876779274 | 0,697 | 0,532 | 8,81E-36 | 1 |
| *maf-S* | 4,67E-39 | -1,22702453 | 0,07 | 0,266 | 5,57E-35 | 1 |
| *goe* | 1,31E-38 | 1,043976855 | 0,408 | 0,204 | 1,57E-34 | 1 |
| *pes* | 2,01E-38 | -1,104102283 | 0,193 | 0,414 | 2,40E-34 | 1 |
| *shf* | 2,04E-38 | 1,002751014 | 0,276 | 0,101 | 2,44E-34 | 1 |
| *mew* | 3,72E-38 | 1,21035711 | 0,52 | 0,325 | 4,44E-34 | 1 |
| *Lapsyn* | 3,75E-38 | 0,918711995 | 0,297 | 0,117 | 4,47E-34 | 1 |
| *Khc-73* | 4,89E-38 | -1,132902883 | 0,224 | 0,446 | 5,84E-34 | 1 |
| *tsr* | 6,18E-38 | -1,085686547 | 0,114 | 0,317 | 7,37E-34 | 1 |
| *CG31694* | 2,23E-37 | -1,321408935 | 0,233 | 0,446 | 2,66E-33 | 1 |
| *Cyp28d1* | 4,96E-37 | -0,972731711 | 0,084 | 0,285 | 5,91E-33 | 1 |
| *Map205* | 5,89E-37 | 1,056922877 | 0,638 | 0,493 | 7,03E-33 | 1 |
| *Ubc6* | 6,87E-37 | -0,998562197 | 0,13 | 0,339 | 8,20E-33 | 1 |
| *ctp* | 3,44E-36 | -1,227984095 | 0,156 | 0,359 | 4,10E-32 | 1 |
| *Gbp3* | 3,78E-36 | 0,904059481 | 0,398 | 0,204 | 4,50E-32 | 1 |
| *CG9449* | 4,08E-36 | 0,951869202 | 0,34 | 0,155 | 4,87E-32 | 1 |
| *hzg* | 4,86E-36 | 0,860827038 | 0,433 | 0,238 | 5,80E-32 | 1 |
| *Sam-S* | 5,06E-36 | 0,990710115 | 0,721 | 0,561 | 6,04E-32 | 1 |
| *blot* | 9,48E-36 | -0,975785856 | 0,245 | 0,473 | 1,13E-31 | 1 |
| *Wbp2* | 1,22E-35 | -0,901210868 | 0,188 | 0,409 | 1,46E-31 | 1 |
| *NK7.1* | 1,25E-35 | -1,04487904 | 0,094 | 0,295 | 1,49E-31 | 1 |
| *CG10621* | 1,53E-35 | 1,038056932 | 0,402 | 0,203 | 1,82E-31 | 1 |
| *GramD1B* | 2,07E-35 | -0,790905081 | 0,408 | 0,623 | 2,47E-31 | 1 |
| *Ubqn* | 3,92E-35 | -0,957668828 | 0,121 | 0,322 | 4,67E-31 | 1 |
| *Fmo-2* | 4,09E-35 | -1,188390459 | 0,102 | 0,295 | 4,88E-31 | 1 |

| | | | | | | |
|---|---|---|---|---|---|---|
| *robo2* | 5,77E-35 | -1,110748198 | 0,28 | 0,492 | 6,88E-31 | 1 |
| *Pfrx* | 8,10E-35 | 0,852517035 | 0,429 | 0,232 | 9,66E-31 | 1 |
| *IM14* | 1,39E-34 | 0,458128099 | 0,337 | 0,146 | 1,66E-30 | 1 |
| *Aldh-III* | 2,51E-34 | 0,810160844 | 0,276 | 0,11 | 2,99E-30 | 1 |
| *CG6428* | 8,97E-34 | -1,015599234 | 0,138 | 0,338 | 1,07E-29 | 1 |
| *S6k* | 1,01E-33 | 0,751876748 | 0,6 | 0,421 | 1,21E-29 | 1 |
| *Pp1alpha-96A* | 1,57E-33 | 0,696088295 | 0,703 | 0,531 | 1,87E-29 | 1 |
| *CG12065* | 3,72E-33 | -1,018115887 | 0,159 | 0,362 | 4,44E-29 | 1 |
| *mura* | 5,73E-33 | 0,772172668 | 0,616 | 0,446 | 6,83E-29 | 1 |
| *Tl* | 6,32E-33 | -0,751647762 | 0,454 | 0,659 | 7,54E-29 | 1 |
| *IM33* | 1,03E-32 | 0,77012771 | 0,501 | 0,303 | 1,23E-28 | 1 |
| *eIF4A* | 1,12E-32 | -0,74780695 | 0,484 | 0,666 | 1,34E-28 | 1 |
| *Oatp30B* | 1,46E-32 | 0,751117016 | 0,69 | 0,536 | 1,74E-28 | 1 |
| *Pdp1* | 1,84E-32 | 0,352432199 | 0,999 | 0,99 | 2,19E-28 | 1 |
| *glec* | 1,95E-32 | -0,979243674 | 0,221 | 0,432 | 2,32E-28 | 1 |
| *Atg13* | 2,15E-32 | 0,921137188 | 0,321 | 0,153 | 2,57E-28 | 1 |
| *cac* | 2,17E-32 | -1,212155907 | 0,317 | 0,523 | 2,58E-28 | 1 |
| *Yeti* | 2,27E-32 | -0,744326368 | 0,477 | 0,682 | 2,71E-28 | 1 |
| *crc* | 2,84E-32 | -0,872010465 | 0,232 | 0,443 | 3,38E-28 | 1 |
| *l(1)G0289* | 3,88E-32 | -0,780831951 | 0,328 | 0,545 | 4,62E-28 | 1 |
| *Pcyt1* | 4,04E-32 | -0,795033824 | 0,335 | 0,552 | 4,82E-28 | 1 |
| *CG15784* | 4,28E-32 | -0,953928645 | 0,108 | 0,298 | 5,10E-28 | 1 |
| *Prat2* | 4,41E-32 | -0,72817786 | 0,624 | 0,779 | 5,27E-28 | 1 |
| *sgg* | 4,82E-32 | 0,478009326 | 0,962 | 0,942 | 5,75E-28 | 1 |
| *nrv1* | 5,17E-32 | -1,05125864 | 0,124 | 0,311 | 6,16E-28 | 1 |
| *ogre* | 8,72E-32 | 0,796116204 | 0,387 | 0,201 | 1,04E-27 | 1 |
| *CG8679* | 8,91E-32 | -1,014710295 | 0,081 | 0,257 | 1,06E-27 | 1 |
| *corto* | 9,00E-32 | 0,86321488 | 0,599 | 0,432 | 1,07E-27 | 1 |
| *GstE3* | 1,01E-31 | -1,043129177 | 0,088 | 0,266 | 1,21E-27 | 1 |
| *cnn* | 1,31E-31 | -0,983034798 | 0,149 | 0,339 | 1,56E-27 | 1 |
| *path* | 1,38E-31 | 0,628906229 | 0,919 | 0,872 | 1,65E-27 | 1 |
| *CG2991* | 1,83E-31 | -1,107634901 | 0,082 | 0,255 | 2,19E-27 | 1 |
| *PRL-1* | 2,35E-31 | -0,969485356 | 0,169 | 0,364 | 2,81E-27 | 1 |
| *Rab11* | 2,48E-31 | -0,878951953 | 0,228 | 0,434 | 2,96E-27 | 1 |
| *CG12012* | 3,97E-31 | -1,004877687 | 0,148 | 0,339 | 4,73E-27 | 1 |
| *Ahcy* | 6,67E-31 | -0,837294126 | 0,161 | 0,359 | 7,96E-27 | 1 |
| *CG11267* | 1,40E-30 | -0,982670141 | 0,083 | 0,253 | 1,67E-26 | 1 |
| *KrT95D* | 1,52E-30 | -0,842214849 | 0,597 | 0,727 | 1,81E-26 | 1 |
| *jim* | 2,12E-30 | 0,661266761 | 0,632 | 0,459 | 2,53E-26 | 1 |
| *lqf* | 2,73E-30 | -0,836559576 | 0,446 | 0,615 | 3,26E-26 | 1 |
| *Cka* | 3,01E-30 | 0,815484027 | 0,439 | 0,259 | 3,60E-26 | 1 |
| *Dic1* | 3,14E-30 | 0,971121697 | 0,302 | 0,14 | 3,74E-26 | 1 |
| *Tab2* | 3,41E-30 | -0,807264826 | 0,268 | 0,472 | 4,06E-26 | 1 |
| *Cyp4g1* | 4,73E-30 | -0,694628402 | 0,224 | 0,441 | 5,64E-26 | 1 |
| *ssp7* | 1,03E-29 | 0,686844547 | 0,588 | 0,394 | 1,23E-25 | 1 |
| *ninaE* | 1,80E-29 | -0,858306087 | 0,122 | 0,306 | 2,14E-25 | 1 |
| *siz* | 2,08E-29 | -0,878915735 | 0,343 | 0,533 | 2,48E-25 | 1 |
| *ckn* | 3,01E-29 | -0,841163943 | 0,21 | 0,409 | 3,59E-25 | 1 |
| *Hex-C* | 3,44E-29 | 0,708085275 | 0,566 | 0,386 | 4,11E-25 | 1 |
| *nmo* | 3,53E-29 | -0,686473411 | 0,44 | 0,644 | 4,21E-25 | 1 |
| *Obp99c* | 3,95E-29 | 0,475688653 | 0,844 | 0,708 | 4,71E-25 | 1 |
| *cdi* | 4,32E-29 | -0,878420227 | 0,264 | 0,469 | 5,16E-25 | 1 |
| *CG42663* | 4,99E-29 | 0,866667193 | 0,394 | 0,22 | 5,95E-25 | 1 |
| *nej* | 8,50E-29 | 0,646304701 | 0,626 | 0,459 | 1,01E-24 | 1 |
| *kst* | 8,89E-29 | 1,093951831 | 0,397 | 0,233 | 1,06E-24 | 1 |
| *CG1640* | 1,08E-28 | 0,812622016 | 0,385 | 0,211 | 1,29E-24 | 1 |
| *Rab5* | 1,29E-28 | -0,828043528 | 0,217 | 0,411 | 1,54E-24 | 1 |
| *kibra* | 1,48E-28 | -0,833279482 | 0,307 | 0,513 | 1,77E-24 | 1 |
| *Odc1* | 1,77E-28 | 0,775458264 | 0,264 | 0,114 | 2,12E-24 | 1 |
| *scb* | 1,80E-28 | -1,087378158 | 0,086 | 0,251 | 2,15E-24 | 1 |
| *hts* | 1,99E-28 | -0,692429107 | 0,354 | 0,554 | 2,37E-24 | 1 |

| | | | | | | |
|---|---|---|---|---|---|---|
| *Npl4* | 2,28E-28 | -0,798082858 | 0,1 | 0,272 | 2,71E-24 | 1 |
| *CG11400* | 2,33E-28 | 0,694676485 | 0,508 | 0,325 | 2,78E-24 | 1 |
| *shep* | 3,10E-28 | 0,472923384 | 0,83 | 0,708 | 3,70E-24 | 1 |
| *l(3)L1231* | 4,00E-28 | 0,638840804 | 0,683 | 0,545 | 4,77E-24 | 1 |
| *Npc2g* | 4,11E-28 | 0,764100133 | 0,464 | 0,288 | 4,90E-24 | 1 |
| *comm2* | 4,76E-28 | -1,078026201 | 0,145 | 0,321 | 5,68E-24 | 1 |
| *Nha2* | 5,62E-28 | 0,810028405 | 0,26 | 0,113 | 6,70E-24 | 1 |
| *CG31523* | 6,51E-28 | -0,846056851 | 0,146 | 0,321 | 7,77E-24 | 1 |
| *Zip99C* | 8,24E-28 | -0,903066594 | 0,087 | 0,252 | 9,83E-24 | 1 |
| *lbk* | 8,76E-28 | -0,956335943 | 0,147 | 0,326 | 1,04E-23 | 1 |
| *Idgf4* | 1,03E-27 | -0,892324907 | 0,17 | 0,347 | 1,23E-23 | 1 |
| *yuri* | 1,10E-27 | -0,832150978 | 0,182 | 0,37 | 1,32E-23 | 1 |
| *kirre* | 1,28E-27 | 0,787882539 | 0,297 | 0,139 | 1,53E-23 | 1 |
| *eEF1alpha1* | 1,93E-27 | -0,422686336 | 0,858 | 0,934 | 2,30E-23 | 1 |
| *CG5773* | 2,15E-27 | -1,170925058 | 0,118 | 0,287 | 2,56E-23 | 1 |
| *CG3902* | 2,45E-27 | 0,967285583 | 0,418 | 0,253 | 2,92E-23 | 1 |
| *CG12004* | 3,08E-27 | -0,877214511 | 0,149 | 0,328 | 3,67E-23 | 1 |
| *CG31705* | 3,28E-27 | -0,625031692 | 0,447 | 0,631 | 3,91E-23 | 1 |
| *CG14762* | 3,31E-27 | 0,720937246 | 0,513 | 0,335 | 3,95E-23 | 1 |
| *CG43078* | 4,05E-27 | 0,802963959 | 0,285 | 0,134 | 4,84E-23 | 1 |
| *CG32066* | 7,81E-27 | -0,72954388 | 0,307 | 0,501 | 9,31E-23 | 1 |
| *CG12116* | 1,12E-26 | 0,79879335 | 0,519 | 0,36 | 1,34E-22 | 1 |
| *Uba1* | 1,47E-26 | -0,874158567 | 0,093 | 0,253 | 1,76E-22 | 1 |
| *Galt* | 1,99E-26 | 0,757846647 | 0,326 | 0,165 | 2,37E-22 | 1 |
| *D2hgdh* | 2,15E-26 | 0,746784493 | 0,419 | 0,25 | 2,56E-22 | 1 |
| *Dif* | 2,50E-26 | 0,580503271 | 0,511 | 0,336 | 2,98E-22 | 1 |
| *Parp* | 2,74E-26 | 0,517704536 | 0,773 | 0,633 | 3,27E-22 | 1 |
| *IscU* | 3,00E-26 | -0,67304661 | 0,289 | 0,488 | 3,58E-22 | 1 |
| *Nup358* | 3,03E-26 | -0,876451652 | 0,122 | 0,292 | 3,62E-22 | 1 |
| *CG42672* | 4,20E-26 | 0,769402541 | 0,314 | 0,16 | 5,01E-22 | 1 |
| *pyd* | 4,20E-26 | 0,716503644 | 0,667 | 0,519 | 5,01E-22 | 1 |
| *CG4297* | 4,46E-26 | 0,735149737 | 0,343 | 0,185 | 5,32E-22 | 1 |
| *scu* | 5,54E-26 | 0,723143258 | 0,317 | 0,161 | 6,61E-22 | 1 |
| *puml* | 6,82E-26 | -0,813056255 | 0,102 | 0,265 | 8,13E-22 | 1 |
| *eIF3c* | 6,87E-26 | -0,781110915 | 0,111 | 0,274 | 8,20E-22 | 1 |
| *CG11537* | 7,38E-26 | -0,910744556 | 0,097 | 0,256 | 8,81E-22 | 1 |
| *Lis-1* | 8,21E-26 | -0,759796098 | 0,239 | 0,427 | 9,79E-22 | 1 |
| *ATPsynC* | 8,98E-26 | 0,666270822 | 0,568 | 0,405 | 1,07E-21 | 1 |
| *psq* | 9,79E-26 | 0,590678297 | 0,623 | 0,454 | 1,17E-21 | 1 |
| *eEF2* | 1,26E-25 | -0,49821719 | 0,675 | 0,796 | 1,50E-21 | 1 |
| *GstT4* | 1,27E-25 | 0,988542937 | 0,444 | 0,29 | 1,51E-21 | 1 |
| *Rab7* | 1,34E-25 | -0,76080827 | 0,178 | 0,359 | 1,60E-21 | 1 |
| *gw* | 1,35E-25 | 0,59950131 | 0,65 | 0,515 | 1,61E-21 | 1 |
| *CG18659* | 1,37E-25 | -0,863623307 | 0,145 | 0,315 | 1,63E-21 | 1 |
| *Nop17l* | 2,56E-25 | -0,886099317 | 0,294 | 0,473 | 3,06E-21 | 1 |
| *bgm* | 2,63E-25 | 0,674773559 | 0,737 | 0,586 | 3,14E-21 | 1 |
| *Letm1* | 2,68E-25 | -0,865400342 | 0,147 | 0,318 | 3,19E-21 | 1 |
| *trx* | 3,20E-25 | 0,630785699 | 0,601 | 0,45 | 3,81E-21 | 1 |
| *CG44008* | 3,53E-25 | -0,789458349 | 0,143 | 0,312 | 4,21E-21 | 1 |
| *Sodh-1* | 3,77E-25 | 0,737414884 | 0,408 | 0,241 | 4,50E-21 | 1 |
| *osp* | 4,50E-25 | 0,659083289 | 0,435 | 0,26 | 5,37E-21 | 1 |
| *Csk* | 4,94E-25 | 0,68738064 | 0,585 | 0,429 | 5,89E-21 | 1 |
| *kek5* | 5,10E-25 | 0,419522605 | 0,716 | 0,545 | 6,09E-21 | 1 |
| *Oatp74D* | 7,81E-25 | 0,663997133 | 0,476 | 0,304 | 9,32E-21 | 1 |
| *Mnt* | 8,83E-25 | 0,572960883 | 0,632 | 0,47 | 1,05E-20 | 1 |
| *CG10600* | 1,10E-24 | -0,814123799 | 0,109 | 0,27 | 1,31E-20 | 1 |
| *Pur-alpha* | 1,18E-24 | 0,612677415 | 0,556 | 0,397 | 1,41E-20 | 1 |
| *CG6051* | 1,27E-24 | -0,516021547 | 0,414 | 0,609 | 1,52E-20 | 1 |
| *Syx1A* | 1,85E-24 | -0,753849811 | 0,436 | 0,603 | 2,21E-20 | 1 |
| *srp* | 2,31E-24 | 0,592208761 | 0,71 | 0,593 | 2,75E-20 | 1 |
| *CG34417* | 3,20E-24 | -0,527078974 | 0,49 | 0,677 | 3,81E-20 | 1 |

| | | | | | | |
|---|---|---|---|---|---|---|
| *Lmpt* | 4,67E-24 | -0,735582815 | 0,334 | 0,51 | 5,57E-20 | 1 |
| *Eb1* | 5,03E-24 | -0,774482218 | 0,138 | 0,303 | 6,00E-20 | 1 |
| *flw* | 5,15E-24 | -0,682822673 | 0,294 | 0,482 | 6,15E-20 | 1 |
| *su(w[a])* | 7,07E-24 | 0,751304171 | 0,251 | 0,119 | 8,43E-20 | 1 |
| *PAPLA1* | 1,42E-23 | -0,682420246 | 0,375 | 0,552 | 1,70E-19 | 1 |
| *Akr1B* | 1,68E-23 | -0,812126926 | 0,133 | 0,294 | 2,00E-19 | 1 |
| *ced-6* | 1,74E-23 | 0,643989489 | 0,578 | 0,429 | 2,08E-19 | 1 |
| *CG10960* | 1,85E-23 | 0,649822047 | 0,857 | 0,821 | 2,21E-19 | 1 |
| *tsl* | 2,04E-23 | 0,807138393 | 0,25 | 0,117 | 2,43E-19 | 1 |
| *CG42238* | 2,24E-23 | 0,690309879 | 0,489 | 0,334 | 2,68E-19 | 1 |
| *Efa6* | 2,61E-23 | -0,795686272 | 0,179 | 0,346 | 3,12E-19 | 1 |
| *Idgf6* | 2,83E-23 | 0,747788223 | 0,302 | 0,159 | 3,37E-19 | 1 |
| *CG10433* | 3,31E-23 | 0,539600012 | 0,741 | 0,645 | 3,95E-19 | 1 |
| *Mst84Da* | 3,69E-23 | -0,883089388 | 0,207 | 0,376 | 4,40E-19 | 1 |
| *CrebA* | 4,35E-23 | -1,245519613 | 0,148 | 0,304 | 5,19E-19 | 1 |
| *rl* | 6,37E-23 | 0,322932565 | 0,946 | 0,874 | 7,60E-19 | 1 |
| *Cyp6v1* | 9,66E-23 | 0,771921234 | 0,264 | 0,128 | 1,15E-18 | 1 |
| *Ald1* | 1,09E-22 | -0,522062927 | 0,842 | 0,902 | 1,30E-18 | 1 |
| *pan* | 1,31E-22 | 0,653422788 | 0,657 | 0,534 | 1,57E-18 | 1 |
| *PMCA* | 1,32E-22 | 0,4770069 | 0,685 | 0,532 | 1,57E-18 | 1 |
| *Kr-h1* | 1,35E-22 | -0,639126568 | 0,351 | 0,528 | 1,61E-18 | 1 |
| *BomBc2* | 1,43E-22 | 0,359734581 | 0,661 | 0,507 | 1,71E-18 | 1 |
| *Tps1* | 1,44E-22 | 0,341803891 | 0,967 | 0,93 | 1,71E-18 | 1 |
| *mys* | 3,47E-22 | -0,771440206 | 0,278 | 0,448 | 4,14E-18 | 1 |
| *Gel* | 3,50E-22 | 0,647411856 | 0,557 | 0,414 | 4,17E-18 | 1 |
| *RpL23* | 4,32E-22 | -0,622803427 | 0,595 | 0,741 | 5,15E-18 | 1 |
| *ens* | 4,87E-22 | -0,622165609 | 0,38 | 0,543 | 5,81E-18 | 1 |
| *Vha16-1* | 5,57E-22 | -0,499421288 | 0,685 | 0,809 | 6,65E-18 | 1 |
| *Bruce* | 5,88E-22 | -0,668562514 | 0,259 | 0,426 | 7,02E-18 | 1 |
| *Gnmt* | 7,03E-22 | 0,58396989 | 0,728 | 0,605 | 8,39E-18 | 1 |
| *Syb* | 7,53E-22 | -0,724792067 | 0,207 | 0,373 | 8,98E-18 | 1 |
| *Pli* | 7,61E-22 | 0,752640144 | 0,645 | 0,525 | 9,08E-18 | 1 |
| *Cf2* | 9,40E-22 | 0,651306644 | 0,448 | 0,301 | 1,12E-17 | 1 |
| *foxo* | 9,48E-22 | 0,452192534 | 0,815 | 0,701 | 1,13E-17 | 1 |
| *CAH1* | 1,02E-21 | 0,675219246 | 0,361 | 0,208 | 1,22E-17 | 1 |
| *MSBP* | 1,07E-21 | -0,689955229 | 0,179 | 0,338 | 1,28E-17 | 1 |
| *CG3638* | 1,08E-21 | 0,650088894 | 0,575 | 0,429 | 1,28E-17 | 1 |
| *scrib* | 1,27E-21 | 0,688559255 | 0,471 | 0,33 | 1,51E-17 | 1 |
| *Clk* | 1,29E-21 | 0,683226449 | 0,331 | 0,187 | 1,54E-17 | 1 |
| *BomS3* | 1,76E-21 | 0,282535061 | 0,544 | 0,378 | 2,10E-17 | 1 |
| *CG6910* | 2,40E-21 | -0,746557014 | 0,697 | 0,789 | 2,86E-17 | 1 |
| *Ets98B* | 3,30E-21 | 0,539290539 | 0,609 | 0,467 | 3,94E-17 | 1 |
| *Hnf4* | 3,70E-21 | 0,587978615 | 0,718 | 0,608 | 4,41E-17 | 1 |
| *Golgin245* | 3,86E-21 | -0,700300199 | 0,12 | 0,269 | 4,60E-17 | 1 |
| *CG42369* | 5,08E-21 | 0,692654989 | 0,319 | 0,179 | 6,05E-17 | 1 |
| *Kank* | 7,21E-21 | 0,60063689 | 0,25 | 0,121 | 8,60E-17 | 1 |
| *PyK* | 8,23E-21 | 0,732511571 | 0,396 | 0,258 | 9,82E-17 | 1 |
| *shn* | 1,07E-20 | -0,564667675 | 0,522 | 0,664 | 1,28E-16 | 1 |
| *mei-P26* | 1,25E-20 | 0,576763541 | 0,538 | 0,387 | 1,49E-16 | 1 |
| *sqd* | 1,30E-20 | 0,388669381 | 0,895 | 0,803 | 1,55E-16 | 1 |
| *chrb* | 1,30E-20 | -0,99025678 | 0,14 | 0,281 | 1,55E-16 | 1 |
| *Flo2* | 2,28E-20 | -0,760123176 | 0,193 | 0,355 | 2,73E-16 | 1 |
| *CD98hc* | 2,47E-20 | -0,756275711 | 0,125 | 0,27 | 2,95E-16 | 1 |
| *CG15098* | 3,72E-20 | -0,676013966 | 0,28 | 0,441 | 4,44E-16 | 1 |
| *Gbs-76A* | 3,92E-20 | 0,60140068 | 0,515 | 0,363 | 4,67E-16 | 1 |
| *vri* | 4,26E-20 | -0,747779882 | 0,4 | 0,55 | 5,08E-16 | 1 |
| *swm* | 4,56E-20 | -0,786711855 | 0,224 | 0,38 | 5,44E-16 | 1 |
| *kraken* | 5,09E-20 | -0,706423602 | 0,137 | 0,283 | 6,07E-16 | 1 |
| *Nedd4* | 5,34E-20 | -0,673979775 | 0,224 | 0,385 | 6,37E-16 | 1 |
| *CG14073* | 6,31E-20 | -0,644034517 | 0,29 | 0,451 | 7,52E-16 | 1 |
| *drongo* | 6,95E-20 | -0,762767271 | 0,491 | 0,624 | 8,29E-16 | 1 |

| | | | | | | |
|---|---|---|---|---|---|---|
| *sws* | 7,28E-20 | 0,712838171 | 0,38 | 0,24 | 8,68E-16 | 1 |
| *stx* | 9,68E-20 | 0,689842385 | 0,487 | 0,351 | 1,15E-15 | 1 |
| *CG4199* | 1,40E-19 | -0,64326871 | 0,142 | 0,291 | 1,67E-15 | 1 |
| *CG1943* | 1,60E-19 | -0,714461223 | 0,118 | 0,258 | 1,91E-15 | 1 |
| *CG6145* | 1,87E-19 | 0,829328806 | 0,527 | 0,401 | 2,23E-15 | 1 |
| *lost* | 1,94E-19 | -0,64838529 | 0,281 | 0,446 | 2,31E-15 | 1 |
| *kis* | 2,22E-19 | 0,43360415 | 0,769 | 0,664 | 2,65E-15 | 1 |
| *Set1* | 2,25E-19 | -0,647482801 | 0,243 | 0,402 | 2,68E-15 | 1 |
| *ap* | 2,38E-19 | 0,694912622 | 0,436 | 0,297 | 2,84E-15 | 1 |
| *Pdk1* | 2,44E-19 | -0,59951566 | 0,686 | 0,754 | 2,92E-15 | 1 |
| *CG42324* | 3,83E-19 | 0,447658394 | 0,724 | 0,599 | 4,57E-15 | 1 |
| *Sin3A* | 4,02E-19 | 0,540873465 | 0,555 | 0,423 | 4,80E-15 | 1 |
| *Gclc* | 4,30E-19 | -0,746864452 | 0,425 | 0,582 | 5,13E-15 | 1 |
| *sky* | 8,46E-19 | -1,002325496 | 0,151 | 0,288 | 1,01E-14 | 1 |
| *Shmt* | 8,98E-19 | -0,527492397 | 0,509 | 0,666 | 1,07E-14 | 1 |
| *RpL21* | 9,56E-19 | -0,511481536 | 0,513 | 0,666 | 1,14E-14 | 1 |
| *for* | 1,14E-18 | 0,449814184 | 0,793 | 0,71 | 1,36E-14 | 1 |
| *tay* | 1,29E-18 | 0,544529807 | 0,508 | 0,368 | 1,53E-14 | 1 |
| *CG7530* | 1,41E-18 | 0,739251624 | 0,388 | 0,26 | 1,69E-14 | 1 |
| *CG17691* | 1,51E-18 | 0,557363533 | 0,537 | 0,409 | 1,80E-14 | 1 |
| *PHGPx* | 1,86E-18 | -0,680329569 | 0,273 | 0,428 | 2,22E-14 | 1 |
| *Galphao* | 1,95E-18 | 0,476546088 | 0,649 | 0,514 | 2,32E-14 | 1 |
| *Mmp2* | 2,27E-18 | -0,892818072 | 0,126 | 0,258 | 2,71E-14 | 1 |
| *anne* | 3,01E-18 | 0,669125586 | 0,346 | 0,219 | 3,59E-14 | 1 |
| *CG9451* | 3,03E-18 | -0,904650712 | 0,14 | 0,276 | 3,61E-14 | 1 |
| *upSET* | 3,61E-18 | 0,581186493 | 0,502 | 0,377 | 4,31E-14 | 1 |
| *CG43736* | 3,87E-18 | 0,615821767 | 0,419 | 0,282 | 4,61E-14 | 1 |
| *RpL17* | 4,25E-18 | -0,553799697 | 0,584 | 0,722 | 5,07E-14 | 1 |
| *fs(1)h* | 4,51E-18 | 0,43008876 | 0,789 | 0,693 | 5,38E-14 | 1 |
| *Atf3* | 4,71E-18 | -0,57236961 | 0,193 | 0,344 | 5,61E-14 | 1 |
| *Rbp1-like* | 5,13E-18 | 0,641686515 | 0,35 | 0,224 | 6,12E-14 | 1 |
| *PlexA* | 1,02E-17 | -0,546918483 | 0,333 | 0,489 | 1,21E-13 | 1 |
| *daw* | 1,06E-17 | 0,63618248 | 0,482 | 0,344 | 1,27E-13 | 1 |
| *RhoGAP15B* | 1,07E-17 | -0,688473931 | 0,136 | 0,274 | 1,28E-13 | 1 |
| *Lpin* | 1,08E-17 | -0,718998839 | 0,381 | 0,531 | 1,29E-13 | 1 |
| *spir* | 1,27E-17 | -0,597426687 | 0,272 | 0,419 | 1,51E-13 | 1 |
| *mts* | 1,41E-17 | -0,556903536 | 0,268 | 0,421 | 1,69E-13 | 1 |
| *CG8036* | 1,68E-17 | 0,607948979 | 0,447 | 0,309 | 2,01E-13 | 1 |
| *Mcad* | 2,12E-17 | 0,610211671 | 0,294 | 0,166 | 2,52E-13 | 1 |
| *Socs16D* | 2,81E-17 | 0,570257722 | 0,256 | 0,137 | 3,35E-13 | 1 |
| *Rac2* | 4,41E-17 | -0,621267946 | 0,212 | 0,357 | 5,26E-13 | 1 |
| *tim* | 4,44E-17 | 0,480106772 | 0,693 | 0,597 | 5,30E-13 | 1 |
| *Tango1* | 4,53E-17 | -0,588731716 | 0,125 | 0,256 | 5,41E-13 | 1 |
| *CG9044* | 7,04E-17 | 0,636877033 | 0,359 | 0,235 | 8,40E-13 | 1 |
| *arg* | 8,27E-17 | 0,714169587 | 0,27 | 0,155 | 9,87E-13 | 1 |
| *Sh3beta* | 9,65E-17 | -0,597517726 | 0,12 | 0,25 | 1,15E-12 | 1 |
| *sowah* | 1,18E-16 | -0,655856004 | 0,23 | 0,377 | 1,41E-12 | 1 |
| *bic* | 1,25E-16 | -0,558400494 | 0,295 | 0,442 | 1,49E-12 | 1 |
| *wdb* | 1,40E-16 | 0,433837723 | 0,724 | 0,616 | 1,67E-12 | 1 |
| *Rab1* | 1,60E-16 | -0,565491348 | 0,214 | 0,358 | 1,91E-12 | 1 |
| *RpL3* | 1,83E-16 | -0,454991408 | 0,55 | 0,688 | 2,18E-12 | 1 |
| *Rap1* | 2,36E-16 | 0,506503514 | 0,58 | 0,455 | 2,82E-12 | 1 |
| *CG12163* | 2,47E-16 | 0,577329951 | 0,572 | 0,469 | 2,95E-12 | 1 |
| *Cat* | 2,79E-16 | -0,625306884 | 0,255 | 0,395 | 3,33E-12 | 1 |
| *Stam* | 3,46E-16 | -0,556927568 | 0,18 | 0,321 | 4,12E-12 | 1 |
| *CG32276* | 4,94E-16 | -0,584649537 | 0,198 | 0,336 | 5,90E-12 | 1 |
| *CG14478* | 6,24E-16 | -0,510555955 | 0,415 | 0,56 | 7,44E-12 | 1 |
| *CG31689* | 7,58E-16 | 0,433644833 | 0,739 | 0,647 | 9,04E-12 | 1 |
| *eas* | 9,09E-16 | 0,496874764 | 0,556 | 0,428 | 1,08E-11 | 1 |
| *CARPB* | 9,24E-16 | 0,617923722 | 0,28 | 0,163 | 1,10E-11 | 1 |
| *CG31145* | 1,05E-15 | 0,413424649 | 0,784 | 0,709 | 1,26E-11 | 1 |

| | | | | | | |
|---|---|---|---|---|---|---|
| *alpha-Man-Ia* | 1,32E-15 | 0,564020514 | 0,443 | 0,32 | 1,57E-11 | 1 |
| *CG1979* | 1,46E-15 | 0,89839275 | 0,26 | 0,154 | 1,74E-11 | 1 |
| *mam* | 2,16E-15 | 0,52714633 | 0,473 | 0,346 | 2,58E-11 | 1 |
| *AGO1* | 2,33E-15 | 0,416739299 | 0,682 | 0,577 | 2,79E-11 | 1 |
| *kra* | 2,48E-15 | -0,436186338 | 0,445 | 0,598 | 2,96E-11 | 1 |
| *Mitf* | 2,85E-15 | 0,498674663 | 0,38 | 0,256 | 3,40E-11 | 1 |
| *CG42514* | 3,39E-15 | 0,620521406 | 0,27 | 0,158 | 4,05E-11 | 1 |
| *RpL18A* | 3,75E-15 | -0,532005205 | 0,539 | 0,661 | 4,47E-11 | 1 |
| *CG7220* | 4,28E-15 | 0,618458486 | 0,306 | 0,191 | 5,10E-11 | 1 |
| *rdx* | 5,53E-15 | 0,425486407 | 0,664 | 0,564 | 6,60E-11 | 1 |
| *Ubi-p5E* | 5,79E-15 | -0,577968413 | 0,181 | 0,311 | 6,90E-11 | 1 |
| *Su(var)2-HP2* | 6,66E-15 | -0,659052758 | 0,159 | 0,282 | 7,95E-11 | 1 |
| *RhoGAP68F* | 7,12E-15 | -0,584218087 | 0,14 | 0,262 | 8,50E-11 | 1 |
| *Diap1* | 8,76E-15 | -0,498084615 | 0,7 | 0,771 | 1,04E-10 | 1 |
| *slmb* | 8,89E-15 | -0,582284467 | 0,13 | 0,251 | 1,06E-10 | 1 |
| *noc* | 9,34E-15 | -0,702902847 | 0,137 | 0,253 | 1,11E-10 | 1 |
| *Mpcp2* | 1,13E-14 | 0,55371309 | 0,293 | 0,182 | 1,34E-10 | 1 |
| *Not1* | 1,21E-14 | -0,379444721 | 0,679 | 0,776 | 1,44E-10 | 1 |
| *ci* | 1,52E-14 | 0,582107925 | 0,26 | 0,152 | 1,81E-10 | 1 |
| *olf186-M* | 1,70E-14 | 0,464651696 | 0,501 | 0,363 | 2,03E-10 | 1 |
| *RpS27* | 1,74E-14 | -0,49223589 | 0,587 | 0,689 | 2,08E-10 | 1 |
| *CG12091* | 1,84E-14 | 0,560166814 | 0,268 | 0,161 | 2,19E-10 | 1 |
| *Trp1* | 2,08E-14 | -0,633299231 | 0,154 | 0,275 | 2,48E-10 | 1 |
| *BuGZ* | 2,63E-14 | 0,556966072 | 0,264 | 0,158 | 3,13E-10 | 1 |
| *l(3)05822* | 2,82E-14 | -0,623622615 | 0,131 | 0,25 | 3,36E-10 | 1 |
| *RpS11* | 3,52E-14 | -0,507690343 | 0,539 | 0,67 | 4,20E-10 | 1 |
| *MESK2* | 3,79E-14 | -0,593732588 | 0,263 | 0,393 | 4,52E-10 | 1 |
| *lolal* | 3,97E-14 | 0,596968004 | 0,379 | 0,271 | 4,74E-10 | 1 |
| *UK114* | 4,25E-14 | 0,540921373 | 0,496 | 0,383 | 5,07E-10 | 1 |
| *eEF5* | 5,23E-14 | -0,535069957 | 0,449 | 0,587 | 6,24E-10 | 1 |
| *18w* | 6,68E-14 | 0,623314313 | 0,276 | 0,17 | 7,96E-10 | 1 |
| *tai* | 6,83E-14 | 0,316810694 | 0,874 | 0,835 | 8,14E-10 | 1 |
| *dco* | 7,51E-14 | -0,509617998 | 0,207 | 0,341 | 8,96E-10 | 1 |
| *CG12054* | 7,94E-14 | 0,474788252 | 0,465 | 0,351 | 9,47E-10 | 1 |
| *RpL38* | 1,14E-13 | -0,573430012 | 0,3 | 0,433 | 1,36E-09 | 1 |
| *RpS10b* | 1,28E-13 | -0,473794167 | 0,561 | 0,681 | 1,53E-09 | 1 |
| *RpL37a* | 1,53E-13 | -0,490239523 | 0,502 | 0,63 | 1,83E-09 | 1 |
| *AGO2* | 1,81E-13 | -0,528214125 | 0,155 | 0,275 | 2,16E-09 | 1 |
| *RpL39* | 1,83E-13 | -0,576251047 | 0,405 | 0,527 | 2,19E-09 | 1 |
| *Ptpmeg* | 1,94E-13 | 0,540603264 | 0,604 | 0,509 | 2,31E-09 | 1 |
| *Tob* | 1,95E-13 | 0,60375899 | 0,251 | 0,151 | 2,32E-09 | 1 |
| *CG30015* | 2,22E-13 | -0,32450773 | 0,726 | 0,82 | 2,64E-09 | 1 |
| *RN-tre* | 2,28E-13 | -0,519048881 | 0,137 | 0,253 | 2,71E-09 | 1 |
| *RpS21* | 3,15E-13 | -0,56332254 | 0,496 | 0,607 | 3,76E-09 | 1 |
| *REPTOR* | 4,55E-13 | -0,475526105 | 0,648 | 0,751 | 5,43E-09 | 1 |
| *Imp* | 5,29E-13 | 0,437309616 | 0,666 | 0,576 | 6,32E-09 | 1 |
| *sbb* | 5,93E-13 | 0,307054164 | 0,671 | 0,548 | 7,08E-09 | 1 |
| *p23* | 5,99E-13 | -0,533071728 | 0,167 | 0,279 | 7,14E-09 | 1 |
| *Hrb27C* | 6,44E-13 | 0,355533451 | 0,802 | 0,729 | 7,69E-09 | 1 |
| *Fis1* | 6,98E-13 | -0,579750024 | 0,152 | 0,26 | 8,32E-09 | 1 |
| *SPARC* | 7,25E-13 | 0,460258566 | 0,388 | 0,266 | 8,65E-09 | 1 |
| *NAT1* | 8,34E-13 | -0,39214771 | 0,625 | 0,729 | 9,95E-09 | 1 |
| *Mkp3* | 8,89E-13 | 0,574952087 | 0,474 | 0,365 | 1,06E-08 | 1 |
| *sdk* | 9,75E-13 | 0,547486869 | 0,271 | 0,17 | 1,16E-08 | 1 |
| *Pcf11* | 1,12E-12 | 0,500339161 | 0,497 | 0,401 | 1,34E-08 | 1 |
| *Treh* | 1,15E-12 | 0,793197707 | 0,27 | 0,176 | 1,37E-08 | 1 |
| *RpL10* | 1,22E-12 | -0,410983001 | 0,462 | 0,591 | 1,46E-08 | 1 |
| *CG34136* | 1,58E-12 | -0,594767136 | 0,193 | 0,312 | 1,89E-08 | 1 |
| *RpL28* | 1,59E-12 | -0,446826638 | 0,673 | 0,754 | 1,89E-08 | 1 |
| *RpL34b* | 1,69E-12 | -0,496820784 | 0,46 | 0,574 | 2,02E-08 | 1 |
| *Ten-m* | 1,77E-12 | -0,732380181 | 0,521 | 0,601 | 2,11E-08 | 1 |

| | | | | | | |
|---|---|---|---|---|---|---|
| *blw* | 1,78E-12 | 0,503619802 | 0,488 | 0,384 | 2,12E-08 | 1 |
| *RpL31* | 2,26E-12 | -0,51998498 | 0,51 | 0,62 | 2,70E-08 | 1 |
| *Pkn* | 2,26E-12 | 0,439241088 | 0,632 | 0,552 | 2,70E-08 | 1 |
| *Pect* | 2,50E-12 | 0,401540316 | 0,636 | 0,522 | 2,98E-08 | 1 |
| *CG10077* | 2,54E-12 | 0,370990985 | 0,68 | 0,6 | 3,03E-08 | 1 |
| *Cerk* | 3,43E-12 | -0,459744694 | 0,235 | 0,362 | 4,09E-08 | 1 |
| *RpL27* | 3,57E-12 | -0,486265078 | 0,525 | 0,65 | 4,26E-08 | 1 |
| *ctrip* | 3,58E-12 | -0,453166767 | 0,294 | 0,418 | 4,27E-08 | 1 |
| *Gs1* | 3,63E-12 | -0,457712118 | 0,204 | 0,327 | 4,34E-08 | 1 |
| *lap* | 3,95E-12 | 0,524702239 | 0,374 | 0,272 | 4,72E-08 | 1 |
| *Tm1* | 4,17E-12 | 0,332663844 | 0,697 | 0,594 | 4,98E-08 | 1 |
| *Oda* | 5,64E-12 | -0,410860801 | 0,785 | 0,841 | 6,73E-08 | 1 |
| *CG5946* | 5,75E-12 | 0,565298655 | 0,254 | 0,16 | 6,86E-08 | 1 |
| *Paip2* | 6,57E-12 | 0,655242488 | 0,437 | 0,341 | 7,84E-08 | 1 |
| *Sap-r* | 6,78E-12 | -0,430423681 | 0,48 | 0,589 | 8,09E-08 | 1 |
| *Itpr* | 6,81E-12 | 0,496288194 | 0,392 | 0,288 | 8,12E-08 | 1 |
| *mtgo* | 7,43E-12 | -0,863499334 | 0,193 | 0,3 | 8,86E-08 | 1 |
| *RpS12* | 7,50E-12 | -0,465460572 | 0,517 | 0,639 | 8,95E-08 | 1 |
| *eIF4H1* | 7,53E-12 | 0,570069076 | 0,27 | 0,175 | 8,98E-08 | 1 |
| *RpS30* | 7,90E-12 | -0,454817228 | 0,443 | 0,565 | 9,43E-08 | 1 |
| *spi* | 8,23E-12 | 0,560138881 | 0,276 | 0,181 | 9,82E-08 | 1 |
| *Sfxn1-3* | 8,45E-12 | -0,47038685 | 0,278 | 0,405 | 1,01E-07 | 1 |
| *RpL7* | 8,50E-12 | -0,440080807 | 0,521 | 0,648 | 1,01E-07 | 1 |
| *Mapmodulin* | 1,00E-11 | 0,507347726 | 0,313 | 0,214 | 1,19E-07 | 1 |
| *RpS25* | 1,10E-11 | -0,515276401 | 0,499 | 0,613 | 1,31E-07 | 1 |
| *Gale* | 1,16E-11 | 0,459552056 | 0,307 | 0,205 | 1,38E-07 | 1 |
| *SREBP* | 1,18E-11 | -0,466645514 | 0,335 | 0,458 | 1,40E-07 | 1 |
| *RpLP1* | 1,23E-11 | -0,462361538 | 0,551 | 0,669 | 1,47E-07 | 1 |
| *Vha100-1* | 1,30E-11 | 0,521156246 | 0,279 | 0,184 | 1,56E-07 | 1 |
| *DIP-alpha* | 1,42E-11 | 0,516523628 | 0,914 | 0,892 | 1,69E-07 | 1 |
| *spin* | 1,67E-11 | 0,426351437 | 0,69 | 0,599 | 2,00E-07 | 1 |
| *Cals* | 1,94E-11 | 0,442863778 | 0,348 | 0,242 | 2,32E-07 | 1 |
| *mt:CoI* | 2,36E-11 | -0,49586873 | 0,179 | 0,286 | 2,82E-07 | 1 |
| *RpS15* | 2,46E-11 | -0,458481536 | 0,616 | 0,712 | 2,93E-07 | 1 |
| *bip2* | 2,54E-11 | 0,42617959 | 0,538 | 0,438 | 3,03E-07 | 1 |
| *Paics* | 2,96E-11 | 0,363409903 | 0,817 | 0,768 | 3,53E-07 | 1 |
| *fry* | 3,10E-11 | -0,538070644 | 0,149 | 0,251 | 3,70E-07 | 1 |
| *bbg* | 4,27E-11 | 0,436900782 | 0,409 | 0,303 | 5,09E-07 | 1 |
| *Cyp1* | 4,43E-11 | -0,416326515 | 0,312 | 0,431 | 5,29E-07 | 1 |
| *SERCA* | 4,56E-11 | -0,431436831 | 0,228 | 0,34 | 5,44E-07 | 1 |
| *RpS5a* | 4,72E-11 | -0,402962033 | 0,437 | 0,554 | 5,63E-07 | 1 |
| *CG33494* | 4,74E-11 | -0,51225449 | 0,246 | 0,356 | 5,65E-07 | 1 |
| *cu* | 4,84E-11 | -0,41663015 | 0,408 | 0,528 | 5,78E-07 | 1 |
| *Pfk* | 4,93E-11 | 0,457246297 | 0,293 | 0,197 | 5,88E-07 | 1 |
| *GstS1* | 5,07E-11 | -0,530309257 | 0,159 | 0,262 | 6,05E-07 | 1 |
| *RpL32* | 5,57E-11 | -0,383666797 | 0,556 | 0,667 | 6,65E-07 | 1 |
| *CG3829* | 6,10E-11 | 0,497717735 | 0,389 | 0,293 | 7,27E-07 | 1 |
| *CG44774* | 6,32E-11 | -0,570774307 | 0,187 | 0,291 | 7,54E-07 | 1 |
| *wb* | 6,36E-11 | -0,66559631 | 0,266 | 0,375 | 7,59E-07 | 1 |
| *sxc* | 6,50E-11 | 0,480621071 | 0,408 | 0,308 | 7,76E-07 | 1 |
| *RpL14* | 6,78E-11 | -0,41087034 | 0,6 | 0,705 | 8,09E-07 | 1 |
| *CG17734* | 6,96E-11 | 0,494149533 | 0,317 | 0,223 | 8,31E-07 | 1 |
| *CG30069* | 8,47E-11 | -0,594739155 | 0,253 | 0,366 | 1,01E-06 | 1 |
| *Nipped-B* | 8,81E-11 | -0,352689817 | 0,698 | 0,776 | 1,05E-06 | 1 |
| *rdgB* | 9,69E-11 | 0,568478791 | 0,275 | 0,183 | 1,16E-06 | 1 |
| *RpL9* | 1,07E-10 | -0,392546862 | 0,476 | 0,586 | 1,27E-06 | 1 |
| *milt* | 1,15E-10 | 0,451914617 | 0,481 | 0,388 | 1,37E-06 | 1 |
| *CG4080* | 1,25E-10 | -0,512076591 | 0,2 | 0,307 | 1,49E-06 | 1 |
| *Marf* | 1,27E-10 | 0,506376313 | 0,371 | 0,282 | 1,51E-06 | 1 |
| *sno* | 1,34E-10 | -0,450699243 | 0,171 | 0,28 | 1,59E-06 | 1 |
| *l(3)80Fj* | 1,41E-10 | 0,326225623 | 0,795 | 0,733 | 1,68E-06 | 1 |

| | | | | | | |
|---|---|---|---|---|---|---|
| *Hsc70-3* | 1,46E-10 | -0,461467814 | 0,33 | 0,451 | 1,74E-06 | 1 |
| *Scamp* | 1,47E-10 | -0,420667382 | 0,18 | 0,289 | 1,75E-06 | 1 |
| *CG10799* | 1,49E-10 | -0,420650444 | 0,422 | 0,543 | 1,78E-06 | 1 |
| *CG42268* | 1,57E-10 | 0,480209055 | 0,522 | 0,445 | 1,87E-06 | 1 |
| *CG14435* | 1,61E-10 | 0,516411883 | 0,273 | 0,181 | 1,92E-06 | 1 |
| *RpS3A* | 1,97E-10 | -0,444530658 | 0,5 | 0,605 | 2,35E-06 | 1 |
| *RpL5* | 2,14E-10 | -0,386516582 | 0,527 | 0,631 | 2,55E-06 | 1 |
| *Hmgs* | 2,14E-10 | 0,526676532 | 0,296 | 0,207 | 2,55E-06 | 1 |
| *CG10680* | 2,82E-10 | 0,389703566 | 0,479 | 0,387 | 3,37E-06 | 1 |
| *Dr* | 2,93E-10 | -0,435687232 | 0,166 | 0,273 | 3,49E-06 | 1 |
| *RpS28b* | 3,41E-10 | -0,424854137 | 0,422 | 0,533 | 4,06E-06 | 1 |
| *Ssdp* | 3,58E-10 | 0,502414907 | 0,282 | 0,192 | 4,27E-06 | 1 |
| *Usp10* | 3,59E-10 | -0,442599613 | 0,306 | 0,418 | 4,28E-06 | 1 |
| *Eip63E* | 3,65E-10 | 0,489447964 | 0,667 | 0,623 | 4,35E-06 | 1 |
| *conu* | 3,68E-10 | 0,351077419 | 0,566 | 0,474 | 4,39E-06 | 1 |
| *hfp* | 3,84E-10 | 0,470782103 | 0,363 | 0,272 | 4,58E-06 | 1 |
| *Ank* | 4,47E-10 | 0,38768432 | 0,547 | 0,463 | 5,34E-06 | 1 |
| *dally* | 4,52E-10 | 0,56908273 | 0,328 | 0,239 | 5,40E-06 | 1 |
| *RpL37A* | 4,54E-10 | -0,411544635 | 0,408 | 0,526 | 5,42E-06 | 1 |
| *CaMKII* | 5,53E-10 | 0,505773742 | 0,375 | 0,287 | 6,59E-06 | 1 |
| *PRAS40* | 5,55E-10 | -0,47900233 | 0,214 | 0,326 | 6,62E-06 | 1 |
| *Mef2* | 5,56E-10 | 0,380363064 | 0,605 | 0,514 | 6,63E-06 | 1 |
| *RpL15* | 5,63E-10 | -0,407385426 | 0,582 | 0,68 | 6,72E-06 | 1 |
| *unc-13* | 5,98E-10 | 0,470696654 | 0,515 | 0,432 | 7,14E-06 | 1 |
| *Thd1* | 6,39E-10 | 0,387280339 | 0,513 | 0,426 | 7,62E-06 | 1 |
| *Nipped-A* | 6,70E-10 | 0,460844094 | 0,348 | 0,256 | 7,99E-06 | 1 |
| *Thor* | 6,78E-10 | 0,336863765 | 0,585 | 0,513 | 8,09E-06 | 1 |
| *fon* | 8,06E-10 | 0,424126332 | 0,381 | 0,279 | 9,61E-06 | 1 |
| *kuz* | 8,53E-10 | -0,495676726 | 0,235 | 0,34 | 1,02E-05 | 1 |
| *14-3-3zeta* | 8,70E-10 | -0,323513781 | 0,502 | 0,605 | 1,04E-05 | 1 |
| *Mical* | 8,94E-10 | 0,566686042 | 0,369 | 0,277 | 1,07E-05 | 1 |
| *Gprk1* | 9,12E-10 | 0,468753228 | 0,405 | 0,318 | 1,09E-05 | 1 |
| *RhoGAPp190* | 9,31E-10 | 0,477456352 | 0,271 | 0,185 | 1,11E-05 | 1 |
| *RpL29* | 9,49E-10 | -0,435859455 | 0,352 | 0,463 | 1,13E-05 | 1 |
| *CG4538* | 9,78E-10 | -0,416280527 | 0,216 | 0,323 | 1,17E-05 | 1 |
| *luna* | 9,83E-10 | 0,276422609 | 0,769 | 0,695 | 1,17E-05 | 1 |
| *CG2852* | 1,02E-09 | -0,459369269 | 0,199 | 0,297 | 1,22E-05 | 1 |
| *Eaat1* | 1,08E-09 | -0,548625491 | 0,267 | 0,369 | 1,29E-05 | 1 |
| *Got2* | 1,10E-09 | -0,33458748 | 0,469 | 0,589 | 1,31E-05 | 1 |
| *RpL35* | 1,37E-09 | -0,409100952 | 0,47 | 0,579 | 1,64E-05 | 1 |
| *CG13784* | 1,39E-09 | 0,435482392 | 0,674 | 0,59 | 1,66E-05 | 1 |
| *Tet* | 1,71E-09 | -0,486448669 | 0,5 | 0,598 | 2,04E-05 | 1 |
| *RpL24* | 1,80E-09 | -0,36563303 | 0,535 | 0,64 | 2,14E-05 | 1 |
| *spri* | 2,04E-09 | 0,36290923 | 0,682 | 0,616 | 2,44E-05 | 1 |
| *alpha-Spec* | 2,27E-09 | -0,416994599 | 0,176 | 0,273 | 2,71E-05 | 1 |
| *Mct1* | 2,36E-09 | 0,423109012 | 0,274 | 0,187 | 2,81E-05 | 1 |
| *CG33229* | 2,63E-09 | -0,347602426 | 0,621 | 0,712 | 3,14E-05 | 1 |
| *aqz* | 2,82E-09 | -0,326377993 | 0,935 | 0,954 | 3,36E-05 | 1 |
| *SNF4Agamma* | 2,85E-09 | -0,254473554 | 0,866 | 0,904 | 3,40E-05 | 1 |
| *CG3597* | 3,14E-09 | 0,491535656 | 0,303 | 0,216 | 3,74E-05 | 1 |
| *E(bx)* | 3,28E-09 | 0,487337096 | 0,323 | 0,241 | 3,92E-05 | 1 |
| *RpS4* | 4,74E-09 | -0,387588882 | 0,513 | 0,607 | 5,65E-05 | 1 |
| *Saf-B* | 5,42E-09 | 0,427566683 | 0,273 | 0,189 | 6,46E-05 | 1 |
| *CG34376* | 5,58E-09 | 0,464966603 | 0,566 | 0,491 | 6,66E-05 | 1 |
| *CG1468* | 5,66E-09 | 0,340754609 | 0,413 | 0,314 | 6,76E-05 | 1 |
| *CG3164* | 5,73E-09 | 0,359453108 | 0,569 | 0,495 | 6,84E-05 | 1 |
| *Sik3* | 6,09E-09 | 0,394700843 | 0,554 | 0,476 | 7,26E-05 | 1 |
| *RpS20* | 8,49E-09 | -0,364930213 | 0,502 | 0,598 | 0,000101227 | 1 |
| *RpL30* | 8,93E-09 | -0,39086962 | 0,219 | 0,32 | 0,000106494 | 1 |
| *Mur2B* | 9,13E-09 | 0,28656387 | 0,508 | 0,4 | 0,000108909 | 1 |
| *RpL12* | 9,15E-09 | -0,357574086 | 0,532 | 0,626 | 0,000109122 | 1 |

| | | | | | | |
|---|---|---|---|---|---|---|
| *cathD* | 9,21E-09 | -0,439815828 | 0,219 | 0,31 | 0,000109916 | 1 |
| *Wdr62* | 9,74E-09 | -0,500960534 | 0,559 | 0,65 | 0,000116154 | 1 |
| *CG6701* | 9,85E-09 | -0,488763845 | 0,202 | 0,298 | 0,000117547 | 1 |
| *CG8671* | 1,08E-08 | -0,364903708 | 0,24 | 0,345 | 0,000129034 | 1 |
| *CG6966* | 1,11E-08 | -0,469459722 | 0,382 | 0,481 | 0,000132255 | 1 |
| *RpL27A* | 1,23E-08 | -0,410593867 | 0,674 | 0,75 | 0,000146913 | 1 |
| *cta* | 1,26E-08 | -0,380044198 | 0,38 | 0,485 | 0,000149927 | 1 |
| *sesB* | 1,78E-08 | 0,358562124 | 0,541 | 0,467 | 0,000212923 | 1 |
| *Irp-1B* | 1,93E-08 | -0,309896689 | 0,211 | 0,315 | 0,000230295 | 1 |
| *RpL19* | 2,17E-08 | -0,303801534 | 0,618 | 0,716 | 0,000259123 | 1 |
| *Ttd14* | 2,36E-08 | 0,465462877 | 0,333 | 0,255 | 0,000281468 | 1 |
| *ftz-f1* | 2,55E-08 | 0,296012026 | 0,664 | 0,591 | 0,000304213 | 1 |
| *santa-maria* | 2,72E-08 | 0,444754656 | 0,319 | 0,234 | 0,000324111 | 1 |
| *Hn* | 2,91E-08 | -0,384671807 | 0,333 | 0,435 | 0,00034664 | 1 |
| *Rala* | 3,24E-08 | -0,3871667 | 0,428 | 0,523 | 0,00038654 | 1 |
| *Doa* | 3,33E-08 | 0,319939916 | 0,571 | 0,496 | 0,000396996 | 1 |
| *RpL35A* | 3,37E-08 | -0,333641845 | 0,491 | 0,587 | 0,000401807 | 1 |
| *Cirl* | 3,44E-08 | -0,415292197 | 0,303 | 0,4 | 0,000409913 | 1 |
| *Dad* | 3,72E-08 | -0,438203569 | 0,41 | 0,5 | 0,000443236 | 1 |
| *Ac13E* | 4,00E-08 | 0,376153396 | 0,319 | 0,232 | 0,000476864 | 1 |
| *Bx* | 4,11E-08 | 0,565487205 | 0,251 | 0,176 | 0,000490651 | 1 |
| *CG17508* | 5,09E-08 | 0,402335304 | 0,36 | 0,283 | 0,000607566 | 1 |
| *Fit1* | 5,24E-08 | -0,435340137 | 0,17 | 0,258 | 0,000624506 | 1 |
| *CG9005* | 5,38E-08 | 0,361913761 | 0,425 | 0,343 | 0,000642246 | 1 |
| *RpL4* | 6,24E-08 | -0,346162912 | 0,487 | 0,577 | 0,000744278 | 1 |
| *Smox* | 6,75E-08 | 0,399626024 | 0,263 | 0,186 | 0,000804665 | 1 |
| *ltl* | 7,38E-08 | -0,44683712 | 0,167 | 0,251 | 0,000879852 | 1 |
| *svp* | 7,53E-08 | 0,329437541 | 0,582 | 0,501 | 0,000898761 | 1 |
| *JIL-1* | 8,13E-08 | 0,426792024 | 0,28 | 0,204 | 0,000969238 | 1 |
| *RpL41* | 8,32E-08 | -0,394387675 | 0,528 | 0,611 | 0,000992338 | 1 |
| *Akt1* | 8,60E-08 | -0,332292827 | 0,307 | 0,411 | 0,001026439 | 1 |
| *Atf6* | 8,88E-08 | -0,355824467 | 0,539 | 0,623 | 0,00105887 | 1 |
| *CrebB* | 9,77E-08 | -0,427826617 | 0,199 | 0,289 | 0,001165962 | 1 |
| *Myd88* | 1,13E-07 | -0,394770409 | 0,298 | 0,395 | 0,001346572 | 1 |
| *Pka-R2* | 1,16E-07 | 0,388332523 | 0,352 | 0,268 | 0,001387091 | 1 |
| *RpS23* | 1,25E-07 | -0,400552755 | 0,625 | 0,703 | 0,001496072 | 1 |
| *krz* | 1,54E-07 | -0,349274194 | 0,17 | 0,252 | 0,00183366 | 1 |
| *Nuak1* | 1,69E-07 | -0,387517359 | 0,222 | 0,309 | 0,002010919 | 1 |
| *Trpm* | 1,83E-07 | 0,297750618 | 0,39 | 0,304 | 0,0021798 | 1 |
| *aop* | 1,91E-07 | -0,273910117 | 0,521 | 0,627 | 0,00227322 | 1 |
| *lp259* | 1,97E-07 | -0,345533711 | 0,183 | 0,267 | 0,002351415 | 1 |
| *nuf* | 2,16E-07 | -0,332614413 | 0,487 | 0,578 | 0,002582133 | 1 |
| *CG2765* | 2,53E-07 | 0,362550897 | 0,294 | 0,214 | 0,00301994 | 1 |
| *CG3376* | 2,64E-07 | 0,347867948 | 0,611 | 0,557 | 0,003154934 | 1 |
| *mrva* | 2,72E-07 | -0,434394806 | 0,242 | 0,327 | 0,003244311 | 1 |
| *B4* | 2,83E-07 | -0,269033163 | 0,418 | 0,517 | 0,003371266 | 1 |
| *RpL11* | 3,22E-07 | -0,330937139 | 0,447 | 0,544 | 0,003846759 | 1 |
| *CG32687* | 3,29E-07 | 0,347894311 | 0,541 | 0,465 | 0,003925337 | 1 |
| *Vha13* | 3,62E-07 | -0,488464967 | 0,191 | 0,267 | 0,004321499 | 1 |
| *kmr* | 3,89E-07 | -0,457795632 | 0,184 | 0,264 | 0,004642018 | 1 |
| *Nacalpha* | 4,20E-07 | -0,375796715 | 0,3 | 0,387 | 0,005014665 | 1 |
| *Cys* | 4,30E-07 | -0,464006809 | 0,244 | 0,321 | 0,005130034 | 1 |
| *B52* | 4,37E-07 | 0,381226853 | 0,362 | 0,296 | 0,005213421 | 1 |
| *eEF1beta* | 4,47E-07 | -0,330490794 | 0,313 | 0,408 | 0,0053291 | 1 |
| *RpL40* | 4,65E-07 | -0,316427229 | 0,408 | 0,497 | 0,005552505 | 1 |
| *NKAIN* | 4,67E-07 | -0,352609521 | 0,25 | 0,34 | 0,005566093 | 1 |
| *Nap1* | 4,68E-07 | -0,328702408 | 0,2 | 0,283 | 0,005580124 | 1 |
| *Pzl* | 4,84E-07 | 0,454105346 | 0,308 | 0,238 | 0,005776235 | 1 |
| *Cbs* | 5,02E-07 | 0,447443844 | 0,308 | 0,234 | 0,005984384 | 1 |
| *Dys* | 5,32E-07 | 0,268247433 | 0,616 | 0,541 | 0,006351352 | 1 |
| *RpS7* | 5,57E-07 | -0,311719215 | 0,624 | 0,707 | 0,006640648 | 1 |

| | | | | | | |
|---|---|---|---|---|---|---|
| *RhoGAP19D* | 5,71E-07 | 0,342834008 | 0,525 | 0,46 | 0,006811567 | 1 |
| *RpL13A* | 7,30E-07 | -0,342477029 | 0,602 | 0,672 | 0,008702516 | 1 |
| *CG17574* | 7,77E-07 | -0,45863775 | 0,198 | 0,273 | 0,009269994 | 1 |
| *Pde11* | 8,11E-07 | 0,301931793 | 0,62 | 0,575 | 0,009676773 | 1 |
| *Lst* | 8,55E-07 | -0,386301533 | 0,767 | 0,806 | 0,010201935 | 1 |
| *BomS6* | 8,69E-07 | 0,374810601 | 0,259 | 0,188 | 0,010361971 | 1 |
| *mub* | 1,19E-06 | 0,417982149 | 0,268 | 0,203 | 0,0141984 | 1 |
| *wnd* | 1,25E-06 | -0,359594012 | 0,272 | 0,361 | 0,014910199 | 1 |
| *RpS29* | 1,27E-06 | -0,380023019 | 0,517 | 0,602 | 0,015175703 | 1 |
| *CG1677* | 1,39E-06 | 0,368136071 | 0,336 | 0,265 | 0,016581122 | 1 |
| *Eno* | 1,48E-06 | -0,35243106 | 0,446 | 0,525 | 0,01764759 | 1 |
| *Mbs* | 1,66E-06 | -0,341051292 | 0,272 | 0,358 | 0,019750493 | 1 |
| *Ref1* | 1,69E-06 | -0,283569136 | 0,202 | 0,288 | 0,020103676 | 1 |
| *akirin* | 1,78E-06 | -0,291070136 | 0,261 | 0,35 | 0,021174701 | 1 |
| *His3.3B* | 2,04E-06 | 0,300630196 | 0,443 | 0,373 | 0,024327432 | 1 |
| *Zyx* | 2,07E-06 | 0,397624338 | 0,265 | 0,199 | 0,02469959 | 1 |
| *RapGAP1* | 2,08E-06 | 0,427794803 | 0,299 | 0,231 | 0,024792069 | 1 |
| *mim* | 2,37E-06 | 0,324845983 | 0,437 | 0,363 | 0,028309897 | 1 |
| *dnr1* | 2,42E-06 | -0,312296065 | 0,193 | 0,272 | 0,028827634 | 1 |
| *Unr* | 2,54E-06 | -0,350351816 | 0,714 | 0,744 | 0,030335351 | 1 |
| *tweek* | 2,76E-06 | -0,313493718 | 0,281 | 0,365 | 0,032932381 | 1 |
| *RpS24* | 3,01E-06 | -0,352493827 | 0,341 | 0,43 | 0,035852268 | 1 |
| *RpS8* | 3,09E-06 | -0,286840302 | 0,664 | 0,737 | 0,036870645 | 1 |
| *hep* | 3,33E-06 | -0,342579454 | 0,265 | 0,344 | 0,039741546 | 1 |
| *RpL36A* | 3,69E-06 | -0,334619269 | 0,521 | 0,598 | 0,044027454 | 1 |
| *dlg1* | 3,83E-06 | 0,326065918 | 0,441 | 0,379 | 0,045678004 | 1 |
| *RpL8* | 3,98E-06 | -0,254392323 | 0,618 | 0,692 | 0,04743632 | 1 |
| *Trf4-1* | 3,99E-06 | 0,39680346 | 0,263 | 0,201 | 0,047591163 | 1 |
| *Cdep* | 4,00E-06 | 0,386834322 | 0,409 | 0,345 | 0,047659993 | 1 |
| *Mrtf* | 4,72E-06 | -0,357061775 | 0,224 | 0,303 | 0,056313787 | 1 |
| *RpS27A* | 4,82E-06 | -0,346054353 | 0,609 | 0,677 | 0,057475812 | 1 |
| *Tango5* | 4,89E-06 | -0,308524155 | 0,3 | 0,384 | 0,058314009 | 1 |
| *alc* | 5,38E-06 | 0,307799897 | 0,272 | 0,207 | 0,064213677 | 1 |
| *CkIIbeta* | 5,43E-06 | 0,36861174 | 0,342 | 0,277 | 0,064785385 | 1 |
| *eIF4G2* | 5,78E-06 | 0,311668864 | 0,38 | 0,311 | 0,069004403 | 1 |
| *RpL26* | 6,54E-06 | -0,250232707 | 0,547 | 0,637 | 0,078012535 | 1 |
| *COX4* | 6,64E-06 | 0,291748843 | 0,351 | 0,279 | 0,079206102 | 1 |
| *sfl* | 7,00E-06 | -0,262038209 | 0,458 | 0,541 | 0,083475366 | 1 |
| *Vha26* | 7,20E-06 | -0,307787573 | 0,211 | 0,286 | 0,085927563 | 1 |
| *ABCD* | 9,15E-06 | 0,336043141 | 0,26 | 0,2 | 0,109145774 | 1 |
| *p120ctn* | 1,02E-05 | 0,325651752 | 0,396 | 0,334 | 0,121959624 | 1 |
| *Opa1* | 1,19E-05 | -0,253273769 | 0,188 | 0,262 | 0,142344068 | 1 |
| *gpp* | 1,20E-05 | 0,278815529 | 0,649 | 0,603 | 0,143559948 | 1 |
| *Mlc-c* | 1,54E-05 | -0,308803255 | 0,287 | 0,364 | 0,183697837 | 1 |
| *Mdh1* | 1,62E-05 | 0,308209895 | 0,42 | 0,351 | 0,193810263 | 1 |
| *pnt* | 1,65E-05 | -0,373186607 | 0,236 | 0,308 | 0,196253548 | 1 |
| *Rim2* | 1,65E-05 | 0,315253496 | 0,29 | 0,226 | 0,197282037 | 1 |
| *glob1* | 1,80E-05 | -0,277073142 | 0,699 | 0,762 | 0,215136029 | 1 |
| *E(Pc)* | 1,80E-05 | -0,270601078 | 0,216 | 0,291 | 0,215257516 | 1 |
| *bol* | 1,91E-05 | 0,449306505 | 0,303 | 0,245 | 0,227924569 | 1 |
| *CG11360* | 2,08E-05 | 0,330864559 | 0,258 | 0,198 | 0,248574737 | 1 |
| *Synd* | 2,34E-05 | -0,250626843 | 0,224 | 0,3 | 0,27966127 | 1 |
| *Spat* | 2,49E-05 | -0,549860557 | 0,425 | 0,482 | 0,29734827 | 1 |
| *HDAC4* | 2,49E-05 | -0,307914915 | 0,299 | 0,376 | 0,297499293 | 1 |
| *Stlk* | 2,50E-05 | 0,330595037 | 0,335 | 0,275 | 0,29835873 | 1 |
| *Tomosyn* | 2,51E-05 | -0,317255276 | 0,312 | 0,386 | 0,29900585 | 1 |
| *RpL7A* | 2,51E-05 | -0,333544147 | 0,583 | 0,643 | 0,299289887 | 1 |
| *galectin* | 2,56E-05 | 0,30607574 | 0,306 | 0,247 | 0,305637596 | 1 |
| *Gmap* | 2,65E-05 | 0,290694503 | 0,51 | 0,461 | 0,316300645 | 1 |
| *vsg* | 2,67E-05 | -0,298403149 | 0,25 | 0,321 | 0,318808219 | 1 |
| *CG32850* | 2,69E-05 | 0,551602396 | 0,26 | 0,205 | 0,321169678 | 1 |

| | | | | | | |
|---|---|---|---|---|---|---|
| *Gbeta13F* | 2,73E-05 | 0,342478791 | 0,556 | 0,518 | 0,32614005 | 1 |
| *Rox8* | 2,83E-05 | 0,294323801 | 0,323 | 0,262 | 0,337977393 | 1 |
| *fog* | 3,21E-05 | 0,344474579 | 0,268 | 0,21 | 0,383197835 | 1 |
| *CG34347* | 3,38E-05 | -0,446513051 | 0,329 | 0,402 | 0,402789686 | 1 |
| *CG30197* | 3,44E-05 | 0,273796211 | 0,255 | 0,195 | 0,409951472 | 1 |
| *Chc* | 3,86E-05 | -0,273366279 | 0,254 | 0,327 | 0,460845989 | 1 |
| *RpS9* | 3,89E-05 | -0,263242458 | 0,545 | 0,615 | 0,463711979 | 1 |
| *Rm62* | 4,69E-05 | 0,253160988 | 0,727 | 0,702 | 0,559206102 | 1 |
| *Nak* | 4,91E-05 | -0,26665959 | 0,183 | 0,253 | 0,585528101 | 1 |
| *Pgm1* | 5,52E-05 | 0,328054188 | 0,292 | 0,236 | 0,658536481 | 1 |
| *RpL10Ab* | 5,72E-05 | -0,256777277 | 0,54 | 0,611 | 0,682035431 | 1 |
| *Top1* | 5,73E-05 | 0,29367145 | 0,366 | 0,307 | 0,683031765 | 1 |
| *CG2865* | 6,10E-05 | -0,379156318 | 0,285 | 0,357 | 0,727079289 | 1 |
| *CG6700* | 6,10E-05 | 0,291782412 | 0,446 | 0,399 | 0,727858918 | 1 |
| *CG42674* | 7,14E-05 | 0,278186998 | 0,439 | 0,38 | 0,851443708 | 1 |
| *RpS17* | 7,35E-05 | -0,250765547 | 0,491 | 0,564 | 0,877202577 | 1 |
| *RpL18* | 7,97E-05 | -0,267642652 | 0,502 | 0,569 | 0,951164914 | 1 |
| *Mal-B2* | 9,36E-05 | -0,320728635 | 0,48 | 0,547 | 1 | 1 |
| *eEF1gamma* | 0,000113836 | -0,26064547 | 0,272 | 0,342 | 1 | 1 |
| *C3G* | 0,000120025 | 0,32305355 | 0,267 | 0,214 | 1 | 1 |
| *RpS6* | 0,000127983 | -0,267742756 | 0,449 | 0,513 | 1 | 1 |
| *CASK* | 0,000130045 | 0,309251159 | 0,364 | 0,312 | 1 | 1 |
| *CG8108* | 0,000140251 | -0,28691848 | 0,336 | 0,407 | 1 | 1 |
| *CG5381* | 0,000145172 | 0,380597036 | 0,277 | 0,23 | 1 | 1 |
| *sm* | 0,000146679 | 0,328979071 | 0,338 | 0,286 | 1 | 1 |
| *Lamp1* | 0,000151957 | 0,299128216 | 0,402 | 0,35 | 1 | 1 |
| *UbcE2H* | 0,000176781 | 0,271672382 | 0,393 | 0,34 | 1 | 1 |
| *alphaTub84B* | 0,000190798 | -0,288955875 | 0,308 | 0,377 | 1 | 1 |
| *S* | 0,000209045 | -0,336287617 | 0,214 | 0,274 | 1 | 1 |
| *Pgant5* | 0,000220332 | 0,289088581 | 0,303 | 0,25 | 1 | 1 |
| *app* | 0,000222643 | -0,268737587 | 0,342 | 0,413 | 1 | 1 |
| *gus* | 0,000228234 | 0,309040243 | 0,263 | 0,214 | 1 | 1 |
| *CG12567* | 0,000252242 | 0,299836863 | 0,338 | 0,282 | 1 | 1 |
| *arr* | 0,000269868 | 0,366339031 | 0,349 | 0,303 | 1 | 1 |
| *Pax* | 0,000325281 | 0,287778687 | 0,254 | 0,204 | 1 | 1 |
| *flfl* | 0,000433369 | 0,260450627 | 0,333 | 0,283 | 1 | 1 |
| *RpS15Aa* | 0,00049827 | -0,256455341 | 0,446 | 0,508 | 1 | 1 |
| *Taldo* | 0,000544999 | 0,417756856 | 0,33 | 0,286 | 1 | 1 |
| *Sarm* | 0,000624775 | -0,336783054 | 0,539 | 0,589 | 1 | 1 |
| *Mad* | 0,000653588 | -0,25774926 | 0,285 | 0,349 | 1 | 1 |
| *ATPsynF* | 0,000735972 | 0,282183622 | 0,285 | 0,235 | 1 | 1 |
| *His4r* | 0,00078145 | -0,274099731 | 0,223 | 0,277 | 1 | 1 |
| *east* | 0,00086783 | 0,312948936 | 0,269 | 0,223 | 1 | 1 |
| *Gdi* | 0,000877062 | -0,252200141 | 0,276 | 0,335 | 1 | 1 |
| *Cp1* | 0,000954897 | 0,434479886 | 0,768 | 0,785 | 1 | 1 |
| *crp* | 0,000955505 | 0,306014706 | 0,355 | 0,312 | 1 | 1 |
| *RpL36* | 0,001380807 | -0,280443133 | 0,571 | 0,625 | 1 | 1 |
| *AP-1gamma* | 0,001654238 | -0,265552428 | 0,234 | 0,287 | 1 | 1 |
| *Pkc98E* | 0,002446697 | 0,27353619 | 0,391 | 0,354 | 1 | 1 |
| *smg* | 0,004034808 | 0,275829704 | 0,325 | 0,285 | 1 | 1 |
| *Naprt* | 0,005956441 | -0,321445988 | 0,334 | 0,379 | 1 | 1 |
| *Xe7* | 0,009310429 | 0,30963977 | 0,325 | 0,299 | 1 | 1 |
| *GstD1.1* | 4,16E-201 | 1,882121653 | 0,956 | 0,552 | 4,96E-197 | 2 |
| *CG11089.1* | 1,10E-199 | 1,794891738 | 0,962 | 0,56 | 1,31E-195 | 2 |
| *CG12795.1* | 1,60E-188 | 2,10593452 | 0,749 | 0,216 | 1,91E-184 | 2 |
| *GstE1.1* | 4,32E-167 | 2,01560537 | 0,753 | 0,251 | 5,15E-163 | 2 |
| *FASN1.1* | 9,37E-165 | -2,929609447 | 0,569 | 0,866 | 1,12E-160 | 2 |
| *ref(2)P.1* | 1,45E-155 | 1,426830234 | 0,937 | 0,536 | 1,73E-151 | 2 |
| *Hsp26.1* | 2,41E-155 | 2,31789737 | 0,778 | 0,302 | 2,87E-151 | 2 |
| *Tsp42Ed.1* | 2,28E-153 | 1,673759149 | 0,793 | 0,318 | 2,72E-149 | 2 |
| *apolpp.1* | 2,96E-148 | -2,503861425 | 0,553 | 0,887 | 3,53E-144 | 2 |

| | | | | | | |
|---|---|---|---|---|---|---|
| *ACC.1* | 1,68E-145 | -2,360769191 | 0,414 | 0,801 | 2,01E-141 | 2 |
| *Pfas.1* | 3,96E-144 | 1,587849805 | 0,874 | 0,463 | 4,72E-140 | 2 |
| *Gbs-70E.1* | 7,59E-141 | -3,896395003 | 0,16 | 0,653 | 9,05E-137 | 2 |
| *Ubi-p63E.1* | 6,95E-140 | 1,464552552 | 0,89 | 0,52 | 8,30E-136 | 2 |
| *CG10383.1* | 1,03E-139 | 1,765845118 | 0,805 | 0,36 | 1,23E-135 | 2 |
| *stv.1* | 1,84E-138 | 1,413195221 | 0,914 | 0,52 | 2,20E-134 | 2 |
| *bru1.1* | 7,15E-135 | -1,665366597 | 0,569 | 0,877 | 8,53E-131 | 2 |
| *CG7130.1* | 8,23E-134 | 1,86734049 | 0,598 | 0,162 | 9,82E-130 | 2 |
| *Nplp2.1* | 4,18E-133 | -2,579257629 | 0,354 | 0,773 | 4,99E-129 | 2 |
| *AnxB9.1* | 1,96E-131 | 1,27828583 | 0,846 | 0,379 | 2,33E-127 | 2 |
| *CG9674.1* | 2,19E-131 | 1,249139322 | 0,924 | 0,536 | 2,61E-127 | 2 |
| *Chchd2.1* | 1,89E-129 | 1,467346535 | 0,883 | 0,529 | 2,25E-125 | 2 |
| *Syp.1* | 3,26E-129 | -1,64252 | 0,603 | 0,885 | 3,88E-125 | 2 |
| *InR.1* | 2,45E-127 | 1,123251872 | 0,99 | 0,887 | 2,92E-123 | 2 |
| *GIIIspla2.1* | 1,67E-120 | 1,711205416 | 0,723 | 0,315 | 2,00E-116 | 2 |
| *CG7470.1* | 5,89E-120 | -1,884210726 | 0,489 | 0,807 | 7,03E-116 | 2 |
| *raw.1* | 2,16E-117 | 1,234644891 | 0,905 | 0,544 | 2,58E-113 | 2 |
| *Act5C.1* | 1,85E-112 | 1,34870786 | 0,834 | 0,45 | 2,20E-108 | 2 |
| *CG14207.1* | 1,62E-111 | 1,314245886 | 0,864 | 0,494 | 1,93E-107 | 2 |
| *CG16898.1* | 1,81E-110 | 1,476967655 | 0,745 | 0,334 | 2,15E-106 | 2 |
| *lola.1* | 9,78E-109 | -1,266948597 | 0,694 | 0,92 | 1,17E-104 | 2 |
| *vir-1.1* | 6,58E-107 | 1,125923006 | 0,944 | 0,67 | 7,85E-103 | 2 |
| *CG32521.1* | 1,87E-106 | -1,707251551 | 0,565 | 0,838 | 2,24E-102 | 2 |
| *CG6503.1* | 1,92E-106 | -1,844355691 | 0,621 | 0,86 | 2,29E-102 | 2 |
| *Frl.1* | 5,79E-105 | 1,344313139 | 0,799 | 0,418 | 6,90E-101 | 2 |
| *Egfr.1* | 8,13E-105 | -1,557655227 | 0,574 | 0,841 | 9,70E-101 | 2 |
| *Hsc70-4.1* | 1,79E-104 | 1,134597541 | 0,905 | 0,685 | 2,14E-100 | 2 |
| *Hsp27.1* | 1,44E-103 | 1,921398134 | 0,635 | 0,243 | 1,71E-99 | 2 |
| *CenG1A.1* | 4,51E-100 | 1,570522071 | 0,797 | 0,464 | 5,38E-96 | 2 |
| *mtd.1* | 1,22E-98 | -1,821213404 | 0,445 | 0,773 | 1,46E-94 | 2 |
| *mamo.1* | 3,92E-98 | -1,910349193 | 0,259 | 0,655 | 4,68E-94 | 2 |
| *Atg1.1* | 1,64E-95 | 1,6737444 | 0,741 | 0,425 | 1,96E-91 | 2 |
| *CG42588.1* | 9,40E-95 | 1,353998473 | 0,603 | 0,223 | 1,12E-90 | 2 |
| *puc.1* | 6,00E-94 | 1,033044812 | 0,918 | 0,619 | 7,16E-90 | 2 |
| *AdSS.1* | 1,68E-91 | 1,300658452 | 0,693 | 0,323 | 2,00E-87 | 2 |
| *CG5953.1* | 3,16E-91 | 1,148265146 | 0,891 | 0,609 | 3,76E-87 | 2 |
| *Nc73EF.1* | 3,60E-91 | -1,585393147 | 0,404 | 0,736 | 4,30E-87 | 2 |
| *Sox102F.1* | 5,74E-91 | -1,284626943 | 0,514 | 0,826 | 6,85E-87 | 2 |
| *Snoo.1* | 2,96E-90 | -1,200941909 | 0,681 | 0,893 | 3,54E-86 | 2 |
| *oys.1* | 4,35E-90 | 1,274894369 | 0,618 | 0,237 | 5,19E-86 | 2 |
| *IP3K1.1* | 2,76E-89 | 1,202598201 | 0,822 | 0,471 | 3,29E-85 | 2 |
| *Nmdmc.1* | 8,49E-89 | 0,928114903 | 0,915 | 0,553 | 1,01E-84 | 2 |
| *Gart.1* | 2,52E-85 | 1,111923406 | 0,869 | 0,598 | 3,01E-81 | 2 |
| *Prosalpha3.1* | 4,05E-85 | 1,504816849 | 0,514 | 0,186 | 4,83E-81 | 2 |
| *CG5958.1* | 5,50E-85 | 1,455083319 | 0,666 | 0,328 | 6,56E-81 | 2 |
| *Hsp68.1* | 3,87E-84 | 1,978688547 | 0,548 | 0,216 | 4,62E-80 | 2 |
| *CG14823.1* | 4,11E-84 | -2,267162976 | 0,062 | 0,44 | 4,90E-80 | 2 |
| *CG45050.1* | 9,99E-84 | 0,9062794 | 0,966 | 0,881 | 1,19E-79 | 2 |
| *CG17108.1* | 1,30E-83 | -2,377838746 | 0,112 | 0,497 | 1,56E-79 | 2 |
| *Trxr-1.1* | 8,04E-83 | 1,130340181 | 0,668 | 0,299 | 9,59E-79 | 2 |
| *Acsl.1* | 9,50E-83 | -1,815122604 | 0,293 | 0,644 | 1,13E-78 | 2 |
| *Smr.1* | 4,58E-82 | -1,204055762 | 0,529 | 0,822 | 5,46E-78 | 2 |
| *Hsp83.1* | 6,71E-82 | 1,02271271 | 0,891 | 0,682 | 8,01E-78 | 2 |
| *Atg8a.1* | 1,54E-81 | 1,044031467 | 0,86 | 0,638 | 1,84E-77 | 2 |
| *pcs.1* | 1,78E-81 | 1,346163964 | 0,681 | 0,338 | 2,12E-77 | 2 |
| *eIF2beta.1* | 2,30E-81 | 1,054796966 | 0,794 | 0,439 | 2,74E-77 | 2 |
| *Abl.1* | 4,82E-81 | 1,187276788 | 0,646 | 0,281 | 5,75E-77 | 2 |
| *CG7720.1* | 6,16E-81 | -2,93350349 | 0,082 | 0,447 | 7,35E-77 | 2 |
| *Atg18b.1* | 1,53E-79 | 1,206489448 | 0,698 | 0,359 | 1,83E-75 | 2 |
| *AspRS.1* | 3,99E-79 | 1,336982621 | 0,466 | 0,148 | 4,75E-75 | 2 |
| *dnc.1* | 1,15E-76 | -1,295207703 | 0,567 | 0,814 | 1,37E-72 | 2 |

| | | | | | | |
|---|---|---|---|---|---|---|
| *Sem1.1* | 3,04E-76 | 1,380327422 | 0,437 | 0,143 | 3,63E-72 | 2 |
| *CG17841.1* | 3,22E-76 | -1,686467718 | 0,211 | 0,565 | 3,84E-72 | 2 |
| *CG34166.1* | 3,64E-76 | -2,33714137 | 0,668 | 0,847 | 4,35E-72 | 2 |
| *dsx.1* | 3,68E-76 | -1,769710145 | 0,277 | 0,617 | 4,39E-72 | 2 |
| *CG42524.1* | 5,23E-76 | -1,963233905 | 0,213 | 0,557 | 6,24E-72 | 2 |
| *CG8086.1* | 4,68E-75 | 1,4454412 | 0,571 | 0,242 | 5,58E-71 | 2 |
| *Hsc70Cb.1* | 1,46E-74 | 1,074773037 | 0,773 | 0,446 | 1,74E-70 | 2 |
| *Culd.1* | 1,77E-74 | -2,047937706 | 0,084 | 0,431 | 2,11E-70 | 2 |
| *BomT3.1* | 7,37E-74 | -2,292205093 | 0,162 | 0,515 | 8,79E-70 | 2 |
| *CG6870.1* | 9,69E-74 | 1,621253088 | 0,419 | 0,133 | 1,16E-69 | 2 |
| *PhKgamma.1* | 7,00E-73 | -1,417774221 | 0,364 | 0,685 | 8,36E-69 | 2 |
| *Src64B.1* | 7,17E-73 | 1,326383423 | 0,739 | 0,467 | 8,56E-69 | 2 |
| *Xrp1.1* | 9,20E-73 | 0,651797387 | 0,986 | 0,886 | 1,10E-68 | 2 |
| *kay.1* | 1,87E-72 | 0,917474725 | 0,769 | 0,434 | 2,22E-68 | 2 |
| *Eglp4.1* | 2,20E-72 | -2,208035139 | 0,112 | 0,456 | 2,63E-68 | 2 |
| *Rpn2.1* | 4,30E-72 | 1,225333563 | 0,543 | 0,228 | 5,13E-68 | 2 |
| *rhea.1* | 8,17E-72 | 1,032835118 | 0,732 | 0,414 | 9,75E-68 | 2 |
| *trbl.1* | 1,26E-71 | 1,008346711 | 0,758 | 0,431 | 1,50E-67 | 2 |
| *Prosalpha2.1* | 1,53E-71 | 1,268749601 | 0,447 | 0,155 | 1,82E-67 | 2 |
| *AkhR.1* | 3,56E-71 | -1,89599814 | 0,062 | 0,405 | 4,25E-67 | 2 |
| *CG1213.1* | 5,20E-71 | -2,214644735 | 0,089 | 0,425 | 6,20E-67 | 2 |
| *PCB.1* | 6,10E-71 | -1,474852559 | 0,393 | 0,685 | 7,28E-67 | 2 |
| *GEFmeso.1* | 9,42E-71 | 1,0489921 | 0,695 | 0,353 | 1,12E-66 | 2 |
| *mbf1.1* | 2,98E-70 | 1,078342003 | 0,603 | 0,266 | 3,55E-66 | 2 |
| *Prosbeta4.1* | 3,73E-70 | 1,361021222 | 0,461 | 0,17 | 4,44E-66 | 2 |
| *CG41378.1* | 1,74E-69 | -1,473976852 | 0,299 | 0,626 | 2,07E-65 | 2 |
| *CG44325.1* | 1,98E-69 | 1,154653828 | 0,556 | 0,238 | 2,36E-65 | 2 |
| *Pka-C1.1* | 5,92E-69 | 0,954940952 | 0,853 | 0,605 | 7,06E-65 | 2 |
| *CG32647.1* | 2,79E-68 | -1,893741271 | 0,343 | 0,638 | 3,33E-64 | 2 |
| *Pvr.1* | 2,99E-68 | 1,318977082 | 0,675 | 0,357 | 3,57E-64 | 2 |
| *CG1673.1* | 3,54E-68 | 0,98592952 | 0,802 | 0,514 | 4,22E-64 | 2 |
| *UGP.1* | 3,55E-68 | -1,870976754 | 0,168 | 0,507 | 4,23E-64 | 2 |
| *CG42788.1* | 5,86E-68 | -2,480528069 | 0,241 | 0,562 | 6,99E-64 | 2 |
| *Prosbeta7.1* | 6,44E-68 | 1,196309451 | 0,468 | 0,177 | 7,69E-64 | 2 |
| *Irc.1* | 9,30E-68 | 1,056470478 | 0,735 | 0,442 | 1,11E-63 | 2 |
| *AcCoAS.1* | 9,46E-68 | -1,487742919 | 0,499 | 0,746 | 1,13E-63 | 2 |
| *Rpn10.1* | 1,23E-67 | 1,279638935 | 0,466 | 0,177 | 1,47E-63 | 2 |
| *px.1* | 1,25E-67 | -1,241890425 | 0,553 | 0,791 | 1,49E-63 | 2 |
| *sug.1* | 9,83E-67 | -2,606460407 | 0,081 | 0,406 | 1,17E-62 | 2 |
| *CG15096.1* | 3,22E-66 | -1,999500451 | 0,023 | 0,329 | 3,85E-62 | 2 |
| *IM4.1* | 1,41E-65 | -2,338274084 | 0,067 | 0,387 | 1,68E-61 | 2 |
| *GstE8.1* | 3,42E-65 | 1,44408142 | 0,404 | 0,135 | 4,08E-61 | 2 |
| *baf.1* | 4,95E-65 | 1,248133766 | 0,412 | 0,136 | 5,91E-61 | 2 |
| *Smg5.1* | 6,39E-65 | 1,05902143 | 0,704 | 0,412 | 7,62E-61 | 2 |
| *Pde9.1* | 9,24E-65 | -1,232304716 | 0,582 | 0,799 | 1,10E-60 | 2 |
| *CaMKI.1* | 1,02E-64 | -0,723872227 | 0,93 | 0,972 | 1,21E-60 | 2 |
| *Vps13.1* | 1,69E-64 | 1,177024273 | 0,595 | 0,285 | 2,02E-60 | 2 |
| *Rad23.1* | 4,46E-64 | 1,065387667 | 0,56 | 0,257 | 5,32E-60 | 2 |
| *CG9932.1* | 6,67E-64 | -0,792679163 | 0,885 | 0,97 | 7,95E-60 | 2 |
| *CG7766.1* | 8,79E-64 | -1,36064878 | 0,362 | 0,644 | 1,05E-59 | 2 |
| *chic.1* | 1,18E-63 | 1,045805774 | 0,648 | 0,341 | 1,41E-59 | 2 |
| *CG1578.1* | 1,94E-63 | -1,764132963 | 0,141 | 0,462 | 2,31E-59 | 2 |
| *AdipoR.1* | 2,31E-63 | -1,415929165 | 0,303 | 0,616 | 2,75E-59 | 2 |
| *Ist1.1* | 3,35E-63 | 1,152006896 | 0,515 | 0,223 | 3,99E-59 | 2 |
| *Gclm.1* | 3,54E-63 | 1,351175432 | 0,584 | 0,287 | 4,23E-59 | 2 |
| *Rpn6.1* | 4,55E-63 | 1,07995522 | 0,614 | 0,314 | 5,43E-59 | 2 |
| *AdSL.1* | 1,53E-62 | 1,294461844 | 0,424 | 0,155 | 1,83E-58 | 2 |
| *aay.1* | 1,04E-61 | 1,181408799 | 0,751 | 0,498 | 1,24E-57 | 2 |
| *NFAT.1* | 1,87E-60 | -1,16588653 | 0,532 | 0,765 | 2,23E-56 | 2 |
| *bun.1* | 1,91E-60 | -1,477553965 | 0,744 | 0,877 | 2,28E-56 | 2 |
| *Idh.1* | 4,07E-60 | -1,725387261 | 0,19 | 0,495 | 4,85E-56 | 2 |

| | | | | | | |
|---|---|---|---|---|---|---|
| *Tret1-1.1* | 5,74E-60 | -1,443183559 | 0,34 | 0,636 | 6,84E-56 | 2 |
| *Tpr2.1* | 8,54E-60 | 0,967380696 | 0,791 | 0,533 | 1,02E-55 | 2 |
| *if.1* | 1,34E-59 | -2,04638687 | 0,118 | 0,425 | 1,60E-55 | 2 |
| *TER94.1* | 1,38E-59 | 0,918944408 | 0,709 | 0,42 | 1,64E-55 | 2 |
| *CCHa2.1* | 3,24E-59 | -2,060154081 | 0,046 | 0,334 | 3,87E-55 | 2 |
| *Msp300.1* | 7,46E-59 | -0,861689236 | 0,782 | 0,915 | 8,90E-55 | 2 |
| *Nmda1.1* | 1,17E-58 | 1,076121433 | 0,57 | 0,279 | 1,39E-54 | 2 |
| *Prosbeta5.1* | 1,94E-58 | 1,241494229 | 0,409 | 0,152 | 2,32E-54 | 2 |
| *MFS17.1* | 7,26E-58 | -1,075757865 | 0,503 | 0,76 | 8,66E-54 | 2 |
| *CG42240* | 7,53E-58 | 1,379250527 | 0,339 | 0,105 | 8,98E-54 | 2 |
| *Men.1* | 1,22E-57 | -1,601455423 | 0,496 | 0,731 | 1,45E-53 | 2 |
| *spoon.1* | 1,67E-57 | -1,484673743 | 0,204 | 0,506 | 1,99E-53 | 2 |
| *Jra* | 1,73E-57 | 1,209590527 | 0,34 | 0,103 | 2,06E-53 | 2 |
| *drpr.1* | 3,16E-57 | 0,984508432 | 0,68 | 0,382 | 3,77E-53 | 2 |
| *CG8034.1* | 7,13E-57 | -1,607974271 | 0,421 | 0,666 | 8,50E-53 | 2 |
| *l(1)G0196.1* | 1,12E-56 | -1,349469929 | 0,202 | 0,511 | 1,34E-52 | 2 |
| *Invadolysin.1* | 1,49E-56 | -2,411398176 | 0,06 | 0,344 | 1,77E-52 | 2 |
| *Cyt-b5-r.1* | 9,76E-56 | -1,212754408 | 0,344 | 0,638 | 1,16E-51 | 2 |
| *CG13315.1* | 2,74E-55 | -1,429019267 | 0,707 | 0,86 | 3,27E-51 | 2 |
| *Tlk.1* | 4,80E-55 | -1,144171451 | 0,345 | 0,638 | 5,73E-51 | 2 |
| *drongo.1* | 1,40E-54 | 1,136273532 | 0,754 | 0,522 | 1,67E-50 | 2 |
| *CG5966.1* | 1,76E-54 | 1,025904138 | 0,671 | 0,378 | 2,10E-50 | 2 |
| *sra.1* | 3,21E-54 | 1,162361736 | 0,471 | 0,203 | 3,83E-50 | 2 |
| *Pdfr.1* | 3,86E-54 | -1,391393251 | 0,26 | 0,55 | 4,60E-50 | 2 |
| *SCaMC.1* | 4,13E-54 | 0,787930526 | 0,886 | 0,725 | 4,93E-50 | 2 |
| *Desat1.1* | 4,48E-54 | -0,88697329 | 0,843 | 0,909 | 5,35E-50 | 2 |
| *Gpdh1.1* | 6,68E-54 | -1,953118912 | 0,152 | 0,433 | 7,97E-50 | 2 |
| *DnaJ-1.1* | 7,81E-54 | 1,041589554 | 0,717 | 0,473 | 9,32E-50 | 2 |
| *AttB.1* | 1,05E-53 | 1,256718399 | 0,378 | 0,131 | 1,26E-49 | 2 |
| *kdn.1* | 3,11E-53 | -1,89259384 | 0,089 | 0,368 | 3,71E-49 | 2 |
| *Hsp70Bc.1* | 3,53E-53 | 1,497680919 | 0,354 | 0,121 | 4,22E-49 | 2 |
| *Rpn5.1* | 2,14E-52 | 1,053727412 | 0,466 | 0,198 | 2,56E-48 | 2 |
| *ps.1* | 2,29E-52 | -0,642592084 | 0,961 | 0,992 | 2,74E-48 | 2 |
| *Spn88Eb.1* | 9,27E-52 | 1,137465088 | 0,381 | 0,142 | 1,11E-47 | 2 |
| *cv-c.1* | 2,22E-51 | 0,781571987 | 0,844 | 0,586 | 2,65E-47 | 2 |
| *Stat92E.1* | 5,43E-51 | -1,530036188 | 0,4 | 0,653 | 6,48E-47 | 2 |
| *Chmp1.1* | 7,63E-51 | 1,227831211 | 0,401 | 0,161 | 9,10E-47 | 2 |
| *lilli.1* | 1,04E-50 | -0,926425112 | 0,577 | 0,782 | 1,25E-46 | 2 |
| *kek5.1* | 1,84E-50 | -1,100440671 | 0,396 | 0,67 | 2,19E-46 | 2 |
| *sima.1* | 1,98E-50 | -0,709810296 | 0,862 | 0,949 | 2,36E-46 | 2 |
| *loco.1* | 2,91E-50 | 0,850123216 | 0,486 | 0,204 | 3,47E-46 | 2 |
| *Chd64.1* | 3,25E-50 | 0,876434055 | 0,67 | 0,384 | 3,88E-46 | 2 |
| *Hsp23* | 3,96E-50 | 1,180416168 | 0,293 | 0,081 | 4,72E-46 | 2 |
| *Rpn1.1* | 4,01E-50 | 1,117457323 | 0,419 | 0,174 | 4,79E-46 | 2 |
| *sbb.1* | 9,53E-50 | -1,174698236 | 0,402 | 0,651 | 1,14E-45 | 2 |
| *Xbp1.1* | 9,88E-50 | 0,869927377 | 0,747 | 0,516 | 1,18E-45 | 2 |
| *Gdap2.1* | 1,52E-49 | 1,035183848 | 0,602 | 0,342 | 1,81E-45 | 2 |
| *Rpn8.1* | 3,28E-49 | 1,020756153 | 0,359 | 0,13 | 3,91E-45 | 2 |
| *Idgf1.1* | 1,10E-48 | 1,037835883 | 0,477 | 0,22 | 1,31E-44 | 2 |
| *CG3638.1* | 1,14E-48 | -1,230770853 | 0,272 | 0,546 | 1,36E-44 | 2 |
| *PSMG1* | 1,18E-48 | 1,081252709 | 0,265 | 0,074 | 1,41E-44 | 2 |
| *Prosalpha5.1* | 2,12E-48 | 1,109500691 | 0,359 | 0,135 | 2,52E-44 | 2 |
| *CG8312.1* | 7,20E-48 | 0,869397659 | 0,541 | 0,255 | 8,58E-44 | 2 |
| *Atpalpha* | 8,53E-48 | -1,005474594 | 0,58 | 0,789 | 1,02E-43 | 2 |
| *CCT3.1* | 9,54E-48 | 0,974018765 | 0,431 | 0,178 | 1,14E-43 | 2 |
| *l(3)80Fg.1* | 1,03E-47 | -1,066237285 | 0,423 | 0,674 | 1,23E-43 | 2 |
| *maf-S.1* | 1,89E-47 | 1,170371705 | 0,372 | 0,145 | 2,25E-43 | 2 |
| *Prosalpha6.1* | 4,48E-47 | 1,116927722 | 0,398 | 0,168 | 5,34E-43 | 2 |
| *CG6428.1* | 7,33E-47 | 1,053624605 | 0,458 | 0,21 | 8,75E-43 | 2 |
| *CG31704.1* | 8,84E-47 | 1,214647865 | 0,392 | 0,164 | 1,05E-42 | 2 |
| *Hsp60A.1* | 9,51E-47 | 0,988874454 | 0,424 | 0,181 | 1,13E-42 | 2 |

| | | | | | | |
|---|---|---|---|---|---|---|
| *jvl.1* | 9,87E-47 | 0,826921267 | 0,855 | 0,714 | 1,18E-42 | 2 |
| *CG5151.1* | 1,63E-46 | -1,63792629 | 0,509 | 0,701 | 1,95E-42 | 2 |
| *pum.1* | 2,85E-46 | -1,285578106 | 0,343 | 0,603 | 3,40E-42 | 2 |
| *Fer1HCH.1* | 3,00E-46 | 0,598566043 | 0,918 | 0,764 | 3,57E-42 | 2 |
| *Rpt5.1* | 4,38E-46 | 1,04796085 | 0,385 | 0,155 | 5,22E-42 | 2 |
| *DppIII.1* | 4,95E-46 | 1,078035683 | 0,359 | 0,139 | 5,90E-42 | 2 |
| *Hers.1* | 7,37E-46 | 0,780669129 | 0,793 | 0,566 | 8,80E-42 | 2 |
| *Rpn9.1* | 8,67E-46 | 1,062068546 | 0,346 | 0,129 | 1,03E-41 | 2 |
| *Ork1.1* | 8,91E-46 | -1,587936919 | 0,141 | 0,403 | 1,06E-41 | 2 |
| *CG8679.1* | 1,31E-45 | 1,084593209 | 0,368 | 0,143 | 1,56E-41 | 2 |
| *nkd.1* | 1,67E-45 | 1,186549646 | 0,629 | 0,387 | 2,00E-41 | 2 |
| *Rpn11* | 1,75E-45 | 1,051746609 | 0,339 | 0,123 | 2,09E-41 | 2 |
| *bmm.1* | 1,86E-45 | 0,606109965 | 0,91 | 0,713 | 2,22E-41 | 2 |
| *Rpn13.1* | 2,31E-45 | 0,968769714 | 0,457 | 0,214 | 2,76E-41 | 2 |
| *CG1416.1* | 3,56E-45 | 1,069791108 | 0,345 | 0,13 | 4,25E-41 | 2 |
| *Lsd-2.1* | 4,39E-45 | -0,869066274 | 0,695 | 0,859 | 5,23E-41 | 2 |
| *pyd.1* | 6,08E-45 | -1,185325419 | 0,378 | 0,631 | 7,25E-41 | 2 |
| *Rpn3.1* | 1,44E-44 | 1,014948639 | 0,369 | 0,148 | 1,72E-40 | 2 |
| *Myo31DF.1* | 1,80E-44 | 1,039779948 | 0,414 | 0,168 | 2,14E-40 | 2 |
| *Tsf1.1* | 9,00E-44 | -1,616475805 | 0,105 | 0,357 | 1,07E-39 | 2 |
| *CG33969* | 1,13E-43 | 1,181889382 | 0,292 | 0,096 | 1,35E-39 | 2 |
| *Mob2.1* | 1,60E-43 | 0,683366167 | 0,864 | 0,715 | 1,91E-39 | 2 |
| *cic.1* | 1,68E-43 | -1,118550254 | 0,506 | 0,717 | 2,00E-39 | 2 |
| *Rpt3.1* | 1,70E-43 | 0,997008989 | 0,372 | 0,149 | 2,03E-39 | 2 |
| *CG10527.1* | 1,94E-43 | 0,930155503 | 0,39 | 0,161 | 2,31E-39 | 2 |
| *CG3036.1* | 2,32E-43 | 0,410781237 | 0,869 | 0,578 | 2,77E-39 | 2 |
| *shep.1* | 3,92E-43 | -0,862544037 | 0,603 | 0,797 | 4,67E-39 | 2 |
| *CG5955.1* | 7,27E-43 | 1,003513247 | 0,371 | 0,147 | 8,67E-39 | 2 |
| *Gpo1.1* | 1,59E-42 | -1,723983628 | 0,096 | 0,34 | 1,90E-38 | 2 |
| *PKD.1* | 1,89E-42 | -1,585763582 | 0,109 | 0,351 | 2,25E-38 | 2 |
| *Prosbeta2* | 2,60E-42 | 0,883685984 | 0,331 | 0,122 | 3,11E-38 | 2 |
| *CG5493* | 4,40E-42 | 1,090165144 | 0,258 | 0,077 | 5,25E-38 | 2 |
| *CG43340.1* | 5,09E-42 | -1,245739874 | 0,228 | 0,477 | 6,07E-38 | 2 |
| *Khc-73.1* | 6,43E-42 | 1,076287402 | 0,551 | 0,314 | 7,67E-38 | 2 |
| *Ubx.1* | 7,93E-42 | -0,897430786 | 0,86 | 0,855 | 9,46E-38 | 2 |
| *CG5910.1* | 2,57E-41 | -1,285743094 | 0,209 | 0,461 | 3,06E-37 | 2 |
| *Prosalpha7.1* | 3,36E-41 | 0,944730003 | 0,381 | 0,165 | 4,01E-37 | 2 |
| *HDAC6.1* | 7,56E-41 | -1,191428517 | 0,151 | 0,404 | 9,02E-37 | 2 |
| *MtnA.1* | 7,72E-41 | 0,801951495 | 0,766 | 0,501 | 9,21E-37 | 2 |
| *Pomp* | 8,27E-41 | 0,982226921 | 0,31 | 0,113 | 9,86E-37 | 2 |
| *Parp.1* | 1,23E-40 | -0,853693711 | 0,518 | 0,733 | 1,47E-36 | 2 |
| *CG1648.1* | 1,42E-40 | -1,166278733 | 0,439 | 0,668 | 1,70E-36 | 2 |
| *LamC.1* | 1,46E-40 | 0,882491142 | 0,443 | 0,207 | 1,74E-36 | 2 |
| *CG1358.1* | 1,91E-40 | -1,762723799 | 0,042 | 0,259 | 2,27E-36 | 2 |
| *Drak.1* | 3,50E-40 | -1,511124059 | 0,481 | 0,669 | 4,17E-36 | 2 |
| *atk.1* | 5,85E-40 | 1,004036318 | 0,325 | 0,122 | 6,98E-36 | 2 |
| *Pmp70.1* | 3,79E-39 | -1,382057479 | 0,096 | 0,329 | 4,53E-35 | 2 |
| *Droj2.1* | 3,92E-39 | 0,825518808 | 0,59 | 0,356 | 4,68E-35 | 2 |
| *ssp7.1* | 4,01E-39 | -1,191328103 | 0,269 | 0,521 | 4,79E-35 | 2 |
| *Prat2.1* | 4,90E-39 | 0,713120837 | 0,843 | 0,69 | 5,85E-35 | 2 |
| *CG8485.1* | 6,48E-39 | -1,392399708 | 0,076 | 0,304 | 7,73E-35 | 2 |
| *CG13784.1* | 7,11E-39 | -1,031255886 | 0,449 | 0,674 | 8,48E-35 | 2 |
| *CG7920.1* | 8,03E-39 | -1,239229231 | 0,118 | 0,355 | 9,57E-35 | 2 |
| *AdamTS-A.1* | 1,09E-38 | -1,912474941 | 0,1 | 0,328 | 1,30E-34 | 2 |
| *rgn.1* | 1,26E-38 | 0,7776282 | 0,641 | 0,4 | 1,51E-34 | 2 |
| *Jabba.1* | 1,34E-38 | -1,441550333 | 0,108 | 0,343 | 1,59E-34 | 2 |
| *Rpt6.1* | 1,36E-38 | 0,996522738 | 0,35 | 0,149 | 1,62E-34 | 2 |
| *PMCA.1* | 1,41E-38 | -0,875903353 | 0,397 | 0,644 | 1,68E-34 | 2 |
| *CHES-1-like.1* | 1,82E-38 | -1,028971853 | 0,303 | 0,542 | 2,17E-34 | 2 |
| *Lsd-1.1* | 2,17E-38 | -1,337758178 | 0,123 | 0,37 | 2,59E-34 | 2 |
| *tsr.1* | 2,80E-38 | 0,936643599 | 0,409 | 0,197 | 3,34E-34 | 2 |

| | | | | | | |
|---|---|---|---|---|---|---|
| *CG2233.1* | 3,14E-38 | -0,614599014 | 0,849 | 0,937 | 3,75E-34 | 2 |
| *Exn.1* | 3,55E-38 | 0,828640353 | 0,662 | 0,455 | 4,24E-34 | 2 |
| *Rpt1* | 8,26E-38 | 1,022245587 | 0,315 | 0,127 | 9,85E-34 | 2 |
| *CG32369.1* | 1,44E-37 | 0,584932683 | 0,822 | 0,623 | 1,72E-33 | 2 |
| *shrb.1* | 1,95E-37 | 0,902302959 | 0,457 | 0,234 | 2,33E-33 | 2 |
| *CCT1* | 2,27E-37 | 0,960266275 | 0,311 | 0,119 | 2,70E-33 | 2 |
| *PRL-1.1* | 2,68E-37 | 0,970578323 | 0,463 | 0,245 | 3,20E-33 | 2 |
| *Akap200.1* | 3,24E-37 | -1,17649815 | 0,395 | 0,608 | 3,87E-33 | 2 |
| *c11.1.1* | 3,54E-37 | 0,860967341 | 0,407 | 0,193 | 4,22E-33 | 2 |
| *Yeti.1* | 3,82E-37 | 0,807812611 | 0,747 | 0,57 | 4,55E-33 | 2 |
| *eas.1* | 5,57E-37 | -1,160189123 | 0,297 | 0,528 | 6,64E-33 | 2 |
| *CG11791.1* | 5,79E-37 | 0,694733858 | 0,615 | 0,372 | 6,91E-33 | 2 |
| *Stip1* | 1,07E-36 | 0,978371584 | 0,275 | 0,099 | 1,27E-32 | 2 |
| *Sdc* | 3,66E-36 | -0,560866753 | 0,916 | 0,96 | 4,36E-32 | 2 |
| *Usp14* | 6,18E-36 | 0,841164483 | 0,315 | 0,124 | 7,37E-32 | 2 |
| *IM14.1* | 7,88E-36 | -1,640859687 | 0,055 | 0,26 | 9,40E-32 | 2 |
| *Rpt2* | 7,92E-36 | 0,899416961 | 0,298 | 0,116 | 9,45E-32 | 2 |
| *uex.1* | 1,18E-35 | -0,776816914 | 0,518 | 0,724 | 1,40E-31 | 2 |
| *Prosbeta3* | 1,41E-35 | 1,006897194 | 0,277 | 0,102 | 1,68E-31 | 2 |
| *Tsp42Ea* | 1,71E-35 | 0,983430851 | 0,286 | 0,109 | 2,04E-31 | 2 |
| *Cyp6w1.1* | 2,03E-35 | -1,408850212 | 0,062 | 0,268 | 2,42E-31 | 2 |
| *teq.1* | 2,20E-35 | -1,184846339 | 0,088 | 0,304 | 2,62E-31 | 2 |
| *Pde6.1* | 2,93E-35 | -1,053736352 | 0,341 | 0,569 | 3,49E-31 | 2 |
| *tai.1* | 3,15E-35 | -0,633477221 | 0,74 | 0,884 | 3,76E-31 | 2 |
| *psq.1* | 3,82E-35 | -0,950415581 | 0,336 | 0,567 | 4,56E-31 | 2 |
| *Bet1* | 4,26E-35 | 0,911431391 | 0,273 | 0,099 | 5,09E-31 | 2 |
| *CG14762.1* | 5,98E-35 | -1,287910493 | 0,227 | 0,449 | 7,13E-31 | 2 |
| *Rpn12* | 1,02E-34 | 0,945638256 | 0,253 | 0,089 | 1,22E-30 | 2 |
| *pyr.1* | 1,49E-34 | 0,902358263 | 0,467 | 0,24 | 1,78E-30 | 2 |
| *wrd.1* | 1,57E-34 | -0,992247285 | 0,273 | 0,496 | 1,87E-30 | 2 |
| *ap.1* | 1,74E-34 | -1,444542162 | 0,176 | 0,399 | 2,08E-30 | 2 |
| *whd.1* | 1,81E-34 | -1,059890217 | 0,34 | 0,576 | 2,16E-30 | 2 |
| *Rbfox1.1* | 3,86E-34 | -0,94994481 | 0,344 | 0,573 | 4,60E-30 | 2 |
| *Hsp70Ab* | 4,20E-34 | 1,016564198 | 0,327 | 0,138 | 5,02E-30 | 2 |
| *Pdp1.1* | 5,14E-34 | -0,484208218 | 0,986 | 0,995 | 6,13E-30 | 2 |
| *EDTP.1* | 6,08E-34 | -1,217422856 | 0,173 | 0,394 | 7,26E-30 | 2 |
| *l(2)41Ab* | 7,83E-34 | -1,866943507 | 0,201 | 0,42 | 9,34E-30 | 2 |
| *Fer2LCH.1* | 8,38E-34 | 0,585842428 | 0,849 | 0,71 | 1,00E-29 | 2 |
| *CG11594.1* | 8,53E-34 | -1,248927879 | 0,056 | 0,254 | 1,02E-29 | 2 |
| *Sodh-1.1* | 1,03E-33 | -1,404643709 | 0,138 | 0,348 | 1,23E-29 | 2 |
| *CCT5* | 1,06E-33 | 0,947279487 | 0,284 | 0,11 | 1,26E-29 | 2 |
| *Lip4.1* | 1,10E-33 | 0,945997289 | 0,57 | 0,354 | 1,31E-29 | 2 |
| *cbt.1* | 1,75E-33 | -1,215387794 | 0,117 | 0,331 | 2,09E-29 | 2 |
| *CG32767.1* | 1,84E-33 | -1,02034085 | 0,213 | 0,445 | 2,19E-29 | 2 |
| *LRP1.1* | 2,80E-33 | -1,209554789 | 0,254 | 0,483 | 3,35E-29 | 2 |
| *cher.1* | 4,50E-33 | 0,663987702 | 0,47 | 0,232 | 5,37E-29 | 2 |
| *sqh* | 4,88E-33 | 0,94015674 | 0,293 | 0,121 | 5,82E-29 | 2 |
| *CG32486.1* | 7,13E-33 | -1,008916571 | 0,31 | 0,532 | 8,51E-29 | 2 |
| *Dmtn.1* | 9,15E-33 | -0,888820198 | 0,641 | 0,785 | 1,09E-28 | 2 |
| *Ets98B.1* | 1,16E-32 | -0,873534265 | 0,345 | 0,57 | 1,39E-28 | 2 |
| *CG17646.1* | 1,48E-32 | -0,874834891 | 0,791 | 0,872 | 1,77E-28 | 2 |
| *CG15099.1* | 1,54E-32 | 0,783241869 | 0,456 | 0,244 | 1,84E-28 | 2 |
| *Tep4* | 1,80E-32 | 0,888704234 | 0,278 | 0,105 | 2,15E-28 | 2 |
| *CG6067.1* | 1,87E-32 | -1,122128429 | 0,169 | 0,386 | 2,24E-28 | 2 |
| *CG32425.1* | 3,13E-32 | -1,195577012 | 0,168 | 0,387 | 3,74E-28 | 2 |
| *kraken.1* | 3,43E-32 | 0,882753313 | 0,379 | 0,186 | 4,09E-28 | 2 |
| *cact.1* | 4,09E-32 | 0,567590834 | 0,78 | 0,595 | 4,87E-28 | 2 |
| *rl.1* | 4,68E-32 | -0,47906998 | 0,811 | 0,926 | 5,59E-28 | 2 |
| *CCT7* | 7,87E-32 | 0,856277754 | 0,264 | 0,099 | 9,39E-28 | 2 |
| *Pur-alpha.1* | 1,02E-31 | -0,909096381 | 0,284 | 0,504 | 1,22E-27 | 2 |
| *Cyp4p1.1* | 1,14E-31 | 0,803683639 | 0,472 | 0,256 | 1,36E-27 | 2 |

| | | | | | | |
|---|---|---|---|---|---|---|
| *Pak3.1* | 1,14E-31 | 0,77088256 | 0,444 | 0,237 | 1,36E-27 | 2 |
| *CG12012.1* | 1,24E-31 | 0,952288168 | 0,424 | 0,227 | 1,48E-27 | 2 |
| *MCPH1.1* | 1,45E-31 | 0,91897799 | 0,462 | 0,266 | 1,73E-27 | 2 |
| *Fur1* | 1,47E-31 | -0,650909647 | 0,577 | 0,765 | 1,75E-27 | 2 |
| *pAbp.1* | 1,54E-31 | 0,530589836 | 0,882 | 0,776 | 1,84E-27 | 2 |
| *Ndae1.1* | 1,83E-31 | -1,481627941 | 0,181 | 0,391 | 2,18E-27 | 2 |
| *pan.1* | 2,48E-31 | -0,859228621 | 0,421 | 0,625 | 2,95E-27 | 2 |
| *CG11267.1* | 2,56E-31 | 0,94313783 | 0,331 | 0,152 | 3,06E-27 | 2 |
| *Ufd4.1* | 3,58E-31 | 0,769991227 | 0,503 | 0,293 | 4,27E-27 | 2 |
| *ALiX.1* | 5,79E-31 | 0,843231793 | 0,367 | 0,178 | 6,90E-27 | 2 |
| *Rab7.1* | 5,93E-31 | 0,827060875 | 0,448 | 0,25 | 7,08E-27 | 2 |
| *Arf79F.1* | 8,20E-31 | 0,75896169 | 0,624 | 0,437 | 9,79E-27 | 2 |
| *Prosalpha4* | 8,28E-31 | 0,827891466 | 0,291 | 0,123 | 9,88E-27 | 2 |
| *Wdr62.1* | 1,05E-30 | 0,870904082 | 0,725 | 0,585 | 1,26E-26 | 2 |
| *LpR2.1* | 1,20E-30 | -0,996204033 | 0,825 | 0,854 | 1,43E-26 | 2 |
| *Gdh.1* | 1,30E-30 | 0,888081726 | 0,538 | 0,334 | 1,55E-26 | 2 |
| *eff.1* | 1,50E-30 | 0,708808718 | 0,73 | 0,544 | 1,79E-26 | 2 |
| *myo.1* | 1,64E-30 | -0,966113214 | 0,23 | 0,442 | 1,96E-26 | 2 |
| *gce.1* | 1,64E-30 | -1,051738575 | 0,338 | 0,531 | 1,96E-26 | 2 |
| *Ald1.1* | 3,94E-30 | 0,57957216 | 0,915 | 0,871 | 4,70E-26 | 2 |
| *CG46385* | 6,70E-30 | -0,532208343 | 0,912 | 0,971 | 7,99E-26 | 2 |
| *vri.1* | 7,69E-30 | 0,621182583 | 0,656 | 0,448 | 9,17E-26 | 2 |
| *CG7945.1* | 1,50E-29 | 0,9145722 | 0,376 | 0,192 | 1,78E-25 | 2 |
| *nej.1* | 1,64E-29 | -0,816448241 | 0,354 | 0,567 | 1,95E-25 | 2 |
| *CG10621.1* | 2,05E-29 | -1,544405865 | 0,124 | 0,316 | 2,44E-25 | 2 |
| *mam.1* | 2,16E-29 | -0,99106932 | 0,23 | 0,441 | 2,58E-25 | 2 |
| *fus.1* | 2,62E-29 | -0,838138354 | 0,24 | 0,453 | 3,12E-25 | 2 |
| *elB.1* | 2,65E-29 | 0,632073329 | 0,478 | 0,256 | 3,16E-25 | 2 |
| *Ilp6.1* | 3,05E-29 | -1,272653936 | 0,159 | 0,361 | 3,64E-25 | 2 |
| *CG4629.1* | 4,69E-29 | -1,243216626 | 0,108 | 0,301 | 5,59E-25 | 2 |
| *bnl.1* | 6,84E-29 | 0,492944105 | 0,492 | 0,273 | 8,16E-25 | 2 |
| *ogre.1* | 1,22E-28 | -1,05474074 | 0,114 | 0,312 | 1,46E-24 | 2 |
| *GstE7.1* | 1,66E-28 | 0,886873809 | 0,311 | 0,141 | 1,98E-24 | 2 |
| *aralar1.1* | 2,12E-28 | 0,613708004 | 0,754 | 0,595 | 2,53E-24 | 2 |
| *Spn43Ab.1* | 4,47E-28 | -0,864729265 | 0,202 | 0,414 | 5,33E-24 | 2 |
| *CG17549.1* | 4,53E-28 | -0,970835321 | 0,221 | 0,423 | 5,41E-24 | 2 |
| *Ahcy.1* | 5,33E-28 | 0,801645999 | 0,434 | 0,247 | 6,35E-24 | 2 |
| *CG9934* | 6,03E-28 | 0,841712824 | 0,272 | 0,116 | 7,20E-24 | 2 |
| *CG34423* | 8,95E-28 | 0,857465699 | 0,256 | 0,104 | 1,07E-23 | 2 |
| *Helz.1* | 9,83E-28 | 0,347347416 | 0,953 | 0,894 | 1,17E-23 | 2 |
| *MFS14.1* | 1,36E-27 | -1,237018768 | 0,259 | 0,448 | 1,63E-23 | 2 |
| *Prosbeta1* | 1,37E-27 | 0,888450576 | 0,284 | 0,127 | 1,63E-23 | 2 |
| *Wbp2.1* | 1,64E-27 | 0,751336389 | 0,48 | 0,288 | 1,96E-23 | 2 |
| *BomS3.1* | 2,54E-27 | -1,24054696 | 0,288 | 0,481 | 3,02E-23 | 2 |
| *osp.1* | 3,07E-27 | -1,109559552 | 0,168 | 0,368 | 3,66E-23 | 2 |
| *CG6115.1* | 3,36E-27 | 0,707078322 | 0,574 | 0,381 | 4,00E-23 | 2 |
| *CG42674.1* | 3,58E-27 | -0,910823138 | 0,247 | 0,45 | 4,27E-23 | 2 |
| *Pdk1.1* | 3,79E-27 | 0,648965404 | 0,798 | 0,71 | 4,52E-23 | 2 |
| *Gadd45.1* | 3,87E-27 | 0,751551279 | 0,371 | 0,186 | 4,62E-23 | 2 |
| *sqd.1* | 5,13E-27 | -0,5161759 | 0,72 | 0,871 | 6,11E-23 | 2 |
| *Syb.1* | 5,39E-27 | 0,836927431 | 0,454 | 0,273 | 6,43E-23 | 2 |
| *AOX1.1* | 7,06E-27 | 0,664312924 | 0,482 | 0,285 | 8,42E-23 | 2 |
| *Lpin.1* | 8,29E-27 | 0,873720455 | 0,613 | 0,438 | 9,89E-23 | 2 |
| *Rel.1* | 9,07E-27 | 0,580911149 | 0,69 | 0,486 | 1,08E-22 | 2 |
| *frma.1* | 9,54E-27 | -0,98527036 | 0,141 | 0,331 | 1,14E-22 | 2 |
| *yuri.1* | 4,11E-26 | 0,754198969 | 0,447 | 0,262 | 4,90E-22 | 2 |
| *CG42663.1* | 4,17E-26 | -1,08027899 | 0,137 | 0,324 | 4,97E-22 | 2 |
| *ATPCL.1* | 4,25E-26 | -1,028079602 | 0,264 | 0,459 | 5,08E-22 | 2 |
| *retm.1* | 4,53E-26 | -1,165914047 | 0,151 | 0,331 | 5,41E-22 | 2 |
| *CG10960.1* | 5,89E-26 | -0,951930226 | 0,816 | 0,838 | 7,02E-22 | 2 |
| *CG16758* | 8,59E-26 | 0,401336392 | 0,981 | 0,901 | 1,03E-21 | 2 |

| | | | | | | |
|---|---|---|---|---|---|---|
| *Nadsyn.1* | 9,82E-26 | -1,050965032 | 0,15 | 0,332 | 1,17E-21 | 2 |
| *CG10082.1* | 1,07E-25 | -1,048549732 | 0,504 | 0,638 | 1,28E-21 | 2 |
| *CG42238.1* | 1,69E-25 | -0,897038692 | 0,236 | 0,435 | 2,02E-21 | 2 |
| *Prosbeta6* | 2,00E-25 | 0,834513217 | 0,269 | 0,12 | 2,39E-21 | 2 |
| *RhoGAP15B.1* | 2,63E-25 | 0,809707264 | 0,358 | 0,185 | 3,14E-21 | 2 |
| *aqz.1* | 3,11E-25 | 0,538967541 | 0,972 | 0,94 | 3,71E-21 | 2 |
| *upSET.1* | 4,10E-25 | -0,81018331 | 0,274 | 0,467 | 4,89E-21 | 2 |
| *ValRS* | 6,51E-25 | 0,776060335 | 0,278 | 0,125 | 7,77E-21 | 2 |
| *Pdha.1* | 6,70E-25 | -0,880983662 | 0,103 | 0,282 | 8,00E-21 | 2 |
| *CG44008.1* | 6,76E-25 | 0,783185186 | 0,383 | 0,213 | 8,07E-21 | 2 |
| *nrv1.1* | 7,15E-25 | 0,779023443 | 0,381 | 0,206 | 8,53E-21 | 2 |
| *tamo* | 8,66E-25 | 0,777743716 | 0,284 | 0,13 | 1,03E-20 | 2 |
| *Gug.1* | 9,21E-25 | -0,706596996 | 0,523 | 0,682 | 1,10E-20 | 2 |
| *Cyp6d5.1* | 9,54E-25 | -1,054609907 | 0,203 | 0,395 | 1,14E-20 | 2 |
| *GstE3.1* | 1,06E-24 | 0,767922763 | 0,332 | 0,165 | 1,26E-20 | 2 |
| *Shmt.1* | 1,19E-24 | 0,659932248 | 0,727 | 0,576 | 1,42E-20 | 2 |
| *ftz-f1.1* | 1,36E-24 | -0,67179162 | 0,476 | 0,662 | 1,62E-20 | 2 |
| *Trx-2.1* | 1,65E-24 | 0,579125761 | 0,433 | 0,241 | 1,96E-20 | 2 |
| *fbp.1* | 2,42E-24 | -0,966174275 | 0,181 | 0,362 | 2,88E-20 | 2 |
| *MTA1-like.1* | 2,45E-24 | 0,372531229 | 0,589 | 0,381 | 2,92E-20 | 2 |
| *CIAPIN1* | 2,48E-24 | 0,736934615 | 0,27 | 0,121 | 2,96E-20 | 2 |
| *Pepck2* | 2,56E-24 | 0,882234388 | 0,321 | 0,163 | 3,06E-20 | 2 |
| *Oatp74D.1* | 2,68E-24 | -0,848726107 | 0,217 | 0,409 | 3,20E-20 | 2 |
| *BomBc2.1* | 2,89E-24 | -0,988059111 | 0,43 | 0,6 | 3,45E-20 | 2 |
| *Tis11* | 3,72E-24 | 0,447132259 | 0,942 | 0,901 | 4,43E-20 | 2 |
| *EcR.1* | 4,15E-24 | -0,842441076 | 0,338 | 0,514 | 4,95E-20 | 2 |
| *Cyp309a1.1* | 4,42E-24 | 0,732039244 | 0,367 | 0,196 | 5,27E-20 | 2 |
| *REPTOR.1* | 5,72E-24 | 0,59983796 | 0,797 | 0,691 | 6,82E-20 | 2 |
| *DnaJ-H* | 7,26E-24 | 0,746786521 | 0,28 | 0,131 | 8,66E-20 | 2 |
| *scrib.1* | 8,64E-24 | -0,918439087 | 0,241 | 0,421 | 1,03E-19 | 2 |
| *CG8036.1* | 9,34E-24 | -0,875903961 | 0,212 | 0,402 | 1,11E-19 | 2 |
| *Rab11.1* | 1,06E-23 | 0,750565289 | 0,496 | 0,323 | 1,27E-19 | 2 |
| *CG3764.1* | 1,42E-23 | -1,069219979 | 0,169 | 0,341 | 1,70E-19 | 2 |
| *Pfrx.1* | 1,50E-23 | -0,910194392 | 0,166 | 0,34 | 1,79E-19 | 2 |
| *CCT2.1* | 1,87E-23 | 0,76233383 | 0,382 | 0,215 | 2,23E-19 | 2 |
| *Cyp28a5.1* | 3,02E-23 | 0,722609178 | 0,296 | 0,142 | 3,61E-19 | 2 |
| *CG8745.1* | 3,12E-23 | -1,618977653 | 0,24 | 0,41 | 3,72E-19 | 2 |
| *CAH1.1* | 3,81E-23 | -1,108962077 | 0,129 | 0,301 | 4,55E-19 | 2 |
| *DIP-lambda* | 4,64E-23 | -1,558977589 | 0,148 | 0,318 | 5,54E-19 | 2 |
| *Mur2B.1* | 4,80E-23 | -0,765952838 | 0,291 | 0,483 | 5,72E-19 | 2 |
| *CG4716.1* | 5,84E-23 | -0,585289327 | 0,793 | 0,879 | 6,96E-19 | 2 |
| *step.1* | 6,38E-23 | 0,659314237 | 0,561 | 0,394 | 7,62E-19 | 2 |
| *CG15293.1* | 8,73E-23 | -0,777739144 | 0,305 | 0,494 | 1,04E-18 | 2 |
| *CG9005.1* | 9,03E-23 | -0,826564996 | 0,237 | 0,414 | 1,08E-18 | 2 |
| *mop* | 1,09E-22 | 0,695638108 | 0,298 | 0,147 | 1,30E-18 | 2 |
| *GstE9.1* | 1,51E-22 | 0,82900382 | 0,374 | 0,213 | 1,81E-18 | 2 |
| *goe.1* | 1,60E-22 | -1,067402967 | 0,145 | 0,313 | 1,91E-18 | 2 |
| *Acbp2.1* | 1,80E-22 | -0,903844568 | 0,178 | 0,359 | 2,14E-18 | 2 |
| *Piezo.1* | 2,03E-22 | 0,383793474 | 0,556 | 0,354 | 2,43E-18 | 2 |
| *CG30423* | 2,36E-22 | 0,746623366 | 0,25 | 0,113 | 2,82E-18 | 2 |
| *Hipk.1* | 2,65E-22 | 0,561469836 | 0,769 | 0,659 | 3,16E-18 | 2 |
| *CG1640.1* | 2,71E-22 | -0,940073472 | 0,142 | 0,31 | 3,23E-18 | 2 |
| *jim.1* | 4,13E-22 | -0,747888262 | 0,391 | 0,558 | 4,93E-18 | 2 |
| *Lis-1.1* | 4,27E-22 | 0,6688689 | 0,494 | 0,322 | 5,09E-18 | 2 |
| *kis.1* | 4,32E-22 | -0,536870818 | 0,577 | 0,739 | 5,15E-18 | 2 |
| *cl* | 4,37E-22 | 0,760608129 | 0,25 | 0,116 | 5,21E-18 | 2 |
| *Spat.1* | 6,14E-22 | 1,029149896 | 0,565 | 0,429 | 7,32E-18 | 2 |
| *Smg6.1* | 6,48E-22 | 0,798433851 | 0,34 | 0,185 | 7,73E-18 | 2 |
| *CG16926.1* | 6,66E-22 | -0,703087912 | 0,505 | 0,662 | 7,94E-18 | 2 |
| *Mnt.1* | 6,82E-22 | -0,711706363 | 0,398 | 0,565 | 8,13E-18 | 2 |
| *hzg.1* | 6,90E-22 | -0,794310407 | 0,17 | 0,346 | 8,23E-18 | 2 |

| | | | | | | |
|---|---|---|---|---|---|---|
| *Rbp1-like.1* | 1,28E-21 | -0,907942041 | 0,143 | 0,307 | 1,53E-17 | 2 |
| *hebe* | 1,58E-21 | 0,754163495 | 0,287 | 0,144 | 1,89E-17 | 2 |
| *Bacc.1* | 2,11E-21 | -0,557940969 | 0,661 | 0,787 | 2,52E-17 | 2 |
| *spict* | 2,16E-21 | 0,736588927 | 0,258 | 0,122 | 2,58E-17 | 2 |
| *Tet.1* | 2,44E-21 | 0,789152172 | 0,666 | 0,533 | 2,91E-17 | 2 |
| *NAT1.1* | 2,50E-21 | 0,524215671 | 0,78 | 0,666 | 2,99E-17 | 2 |
| *Hsc70-5.1* | 3,03E-21 | 0,659548719 | 0,409 | 0,24 | 3,61E-17 | 2 |
| *Pdk* | 3,06E-21 | 0,477165195 | 0,873 | 0,859 | 3,65E-17 | 2 |
| *foxo.1* | 3,10E-21 | -0,62278028 | 0,652 | 0,767 | 3,70E-17 | 2 |
| *Nop17l.1* | 3,61E-21 | 0,593543235 | 0,543 | 0,371 | 4,31E-17 | 2 |
| *CG9691.1* | 4,46E-21 | -0,794689125 | 0,211 | 0,387 | 5,32E-17 | 2 |
| *zormin.1* | 5,73E-21 | 0,645344902 | 0,542 | 0,367 | 6,83E-17 | 2 |
| *cdi.1* | 6,46E-21 | 0,642349996 | 0,527 | 0,36 | 7,70E-17 | 2 |
| *Clc* | 8,27E-21 | 0,710921648 | 0,259 | 0,124 | 9,87E-17 | 2 |
| *lqf.1* | 1,07E-20 | 0,677046304 | 0,646 | 0,53 | 1,27E-16 | 2 |
| *Ttd14.1* | 1,13E-20 | -0,873244457 | 0,156 | 0,323 | 1,35E-16 | 2 |
| *p47* | 1,46E-20 | 0,752851652 | 0,255 | 0,122 | 1,74E-16 | 2 |
| *CG17124.1* | 1,66E-20 | -0,875414679 | 0,688 | 0,769 | 1,98E-16 | 2 |
| *Hex-C.1* | 1,68E-20 | -0,756305151 | 0,316 | 0,489 | 2,00E-16 | 2 |
| *Swip-1.1* | 1,99E-20 | 0,537835123 | 0,544 | 0,353 | 2,38E-16 | 2 |
| *srp.1* | 2,16E-20 | -0,705108311 | 0,527 | 0,666 | 2,58E-16 | 2 |
| *RhoGAP68F.1* | 2,45E-20 | 0,745523156 | 0,331 | 0,185 | 2,93E-16 | 2 |
| *Oda.1* | 3,08E-20 | 0,501478317 | 0,873 | 0,806 | 3,68E-16 | 2 |
| *Cyp6g1.1* | 3,74E-20 | -0,847027342 | 0,28 | 0,444 | 4,46E-16 | 2 |
| *PyK.1* | 4,58E-20 | -0,794038924 | 0,179 | 0,345 | 5,46E-16 | 2 |
| *gpp.1* | 4,66E-20 | -0,669507337 | 0,515 | 0,652 | 5,56E-16 | 2 |
| *RpL23.1* | 5,81E-20 | 0,617677395 | 0,774 | 0,666 | 6,93E-16 | 2 |
| *Socs36E.1* | 6,28E-20 | 0,372027944 | 0,466 | 0,276 | 7,49E-16 | 2 |
| *gw.1* | 6,87E-20 | -0,575317639 | 0,43 | 0,602 | 8,19E-16 | 2 |
| *fon.1* | 7,42E-20 | -0,937439494 | 0,192 | 0,353 | 8,85E-16 | 2 |
| *SPARC.1* | 7,63E-20 | -0,758607103 | 0,178 | 0,349 | 9,10E-16 | 2 |
| *Efa6.1* | 7,77E-20 | 0,724157716 | 0,405 | 0,253 | 9,27E-16 | 2 |
| *unc-13.1* | 9,87E-20 | -0,823921911 | 0,34 | 0,5 | 1,18E-15 | 2 |
| *Dif.1* | 9,94E-20 | -0,98173207 | 0,279 | 0,431 | 1,19E-15 | 2 |
| *bchs* | 1,50E-19 | 0,673923295 | 0,279 | 0,141 | 1,79E-15 | 2 |
| *LRR.1* | 1,79E-19 | 0,335509506 | 0,669 | 0,475 | 2,13E-15 | 2 |
| *tkv.1* | 1,87E-19 | -0,910420925 | 0,278 | 0,431 | 2,23E-15 | 2 |
| *scyl.1* | 2,04E-19 | 0,358663534 | 0,904 | 0,776 | 2,43E-15 | 2 |
| *RpL21.1* | 2,13E-19 | 0,605078689 | 0,712 | 0,583 | 2,54E-15 | 2 |
| *Dbp80* | 2,33E-19 | -0,444910982 | 0,529 | 0,7 | 2,78E-15 | 2 |
| *CG6330.1* | 2,65E-19 | 0,639198724 | 0,423 | 0,253 | 3,17E-15 | 2 |
| *CG31145.1* | 3,00E-19 | -0,559629277 | 0,638 | 0,765 | 3,58E-15 | 2 |
| *Caper.1* | 3,03E-19 | -0,587532864 | 0,569 | 0,705 | 3,61E-15 | 2 |
| *DIP-alpha.1* | 3,43E-19 | -0,837955223 | 0,855 | 0,914 | 4,09E-15 | 2 |
| *daw.1* | 4,31E-19 | -0,805940605 | 0,263 | 0,431 | 5,14E-15 | 2 |
| *dos.1* | 4,51E-19 | 0,575805758 | 0,294 | 0,151 | 5,38E-15 | 2 |
| *CG6966.1* | 6,77E-19 | 0,803179103 | 0,558 | 0,412 | 8,08E-15 | 2 |
| *Npc2g.1* | 7,15E-19 | -0,776124118 | 0,222 | 0,388 | 8,53E-15 | 2 |
| *Ac13E.1* | 7,30E-19 | -0,9248575 | 0,146 | 0,298 | 8,71E-15 | 2 |
| *Dys.1* | 7,57E-19 | -0,619410916 | 0,442 | 0,607 | 9,04E-15 | 2 |
| *CaMKII.1* | 7,87E-19 | -0,737428393 | 0,195 | 0,356 | 9,39E-15 | 2 |
| *Lst.1* | 8,28E-19 | 0,5084929 | 0,857 | 0,772 | 9,87E-15 | 2 |
| *S6k.1* | 8,99E-19 | -0,65694317 | 0,36 | 0,52 | 1,07E-14 | 2 |
| *RpL37a.1* | 1,29E-18 | 0,678018948 | 0,68 | 0,557 | 1,54E-14 | 2 |
| *cpx* | 1,35E-18 | -0,946249684 | 0,22 | 0,372 | 1,61E-14 | 2 |
| *l(1)G0289.1* | 1,40E-18 | 0,521961896 | 0,603 | 0,43 | 1,67E-14 | 2 |
| *CG31635.1* | 1,60E-18 | -0,718826992 | 0,438 | 0,589 | 1,90E-14 | 2 |
| *l(3)80Fj.1* | 1,96E-18 | -0,482790646 | 0,656 | 0,786 | 2,33E-14 | 2 |
| *RhoGAP19D.1* | 2,23E-18 | -0,939836395 | 0,373 | 0,518 | 2,66E-14 | 2 |
| *CG17278.1* | 2,52E-18 | 0,586122107 | 0,409 | 0,252 | 3,00E-14 | 2 |
| *CG34054.1* | 2,78E-18 | 0,743202335 | 0,292 | 0,156 | 3,32E-14 | 2 |

| | | | | | | |
|---|---|---|---|---|---|---|
| *alpha-Man-Ia.1* | 3,19E-18 | -0,829344279 | 0,244 | 0,399 | 3,81E-14 | 2 |
| *anne.1* | 3,21E-18 | -0,794734963 | 0,151 | 0,298 | 3,83E-14 | 2 |
| *Mthfs* | 3,43E-18 | 0,788496472 | 0,297 | 0,166 | 4,09E-14 | 2 |
| *RhoGAP71E.1* | 3,51E-18 | 0,604946721 | 0,53 | 0,392 | 4,19E-14 | 2 |
| *CG33493.1* | 3,80E-18 | -0,853738527 | 0,114 | 0,26 | 4,53E-14 | 2 |
| *stx.1* | 3,83E-18 | -0,827480746 | 0,278 | 0,435 | 4,57E-14 | 2 |
| *ScsbetaA.1* | 6,93E-18 | -0,505668811 | 0,35 | 0,525 | 8,26E-14 | 2 |
| *CG43736.1* | 8,47E-18 | -0,7339015 | 0,208 | 0,367 | 1,01E-13 | 2 |
| *MCU.1* | 9,88E-18 | 0,656363695 | 0,4 | 0,247 | 1,18E-13 | 2 |
| *larp.1* | 1,08E-17 | -0,655639562 | 0,485 | 0,62 | 1,29E-13 | 2 |
| *Pcyt1.1* | 1,08E-17 | 0,520305034 | 0,599 | 0,441 | 1,29E-13 | 2 |
| *His4r.1* | 1,35E-17 | 0,762811402 | 0,36 | 0,225 | 1,61E-13 | 2 |
| *CLIP-190* | 1,37E-17 | -0,434921303 | 0,558 | 0,713 | 1,63E-13 | 2 |
| *CG30015.1* | 1,38E-17 | 0,381995304 | 0,867 | 0,763 | 1,64E-13 | 2 |
| *RpL28.1* | 1,66E-17 | 0,623395605 | 0,793 | 0,706 | 1,98E-13 | 2 |
| *rdx.1* | 1,91E-17 | -0,565318423 | 0,497 | 0,63 | 2,28E-13 | 2 |
| *RpL27A.1* | 1,95E-17 | 0,66221834 | 0,783 | 0,706 | 2,33E-13 | 2 |
| *mew.1* | 1,98E-17 | -1,090031618 | 0,28 | 0,425 | 2,36E-13 | 2 |
| *CG10600.1* | 2,76E-17 | 0,643549665 | 0,32 | 0,182 | 3,29E-13 | 2 |
| *RpS11.1* | 3,75E-17 | 0,678393099 | 0,714 | 0,597 | 4,48E-13 | 2 |
| *Pfk.1* | 5,00E-17 | -0,665759039 | 0,122 | 0,264 | 5,97E-13 | 2 |
| *NUCB1.1* | 5,35E-17 | 0,577706639 | 0,331 | 0,19 | 6,38E-13 | 2 |
| *milt.1* | 5,47E-17 | -0,677998348 | 0,303 | 0,457 | 6,53E-13 | 2 |
| *Ire1.1* | 5,60E-17 | 0,592952893 | 0,428 | 0,28 | 6,68E-13 | 2 |
| *Lac* | 5,72E-17 | 0,654402756 | 0,259 | 0,134 | 6,82E-13 | 2 |
| *CG14154* | 6,20E-17 | -0,593714567 | 0,34 | 0,503 | 7,40E-13 | 2 |
| *l(3)L1231.1* | 6,66E-17 | -0,561750212 | 0,49 | 0,624 | 7,95E-13 | 2 |
| *CG12054.1* | 7,32E-17 | -0,661781698 | 0,27 | 0,428 | 8,73E-13 | 2 |
| *conu.1* | 7,93E-17 | -0,575469347 | 0,395 | 0,541 | 9,46E-13 | 2 |
| *Fis1.1* | 7,99E-17 | 0,735306758 | 0,32 | 0,193 | 9,54E-13 | 2 |
| *Rab1.1* | 8,03E-17 | 0,579776571 | 0,423 | 0,273 | 9,58E-13 | 2 |
| *Obp99c.1* | 8,92E-17 | -0,377658563 | 0,648 | 0,788 | 1,06E-12 | 2 |
| *Plc21C* | 9,54E-17 | -0,563447922 | 0,463 | 0,609 | 1,14E-12 | 2 |
| *RpL14.1* | 9,99E-17 | 0,578388394 | 0,746 | 0,645 | 1,19E-12 | 2 |
| *Tab2.1* | 1,05E-16 | 0,539945162 | 0,52 | 0,366 | 1,25E-12 | 2 |
| *RasGAP1.1* | 1,13E-16 | 0,638618693 | 0,386 | 0,242 | 1,35E-12 | 2 |
| *CG30026* | 1,18E-16 | 0,770126567 | 0,264 | 0,14 | 1,41E-12 | 2 |
| *mys.1* | 1,33E-16 | 0,593697064 | 0,503 | 0,356 | 1,59E-12 | 2 |
| *RpS15.1* | 1,43E-16 | 0,634819468 | 0,759 | 0,654 | 1,71E-12 | 2 |
| *CG6512* | 1,71E-16 | 0,673520185 | 0,266 | 0,143 | 2,04E-12 | 2 |
| *cwo.1* | 1,79E-16 | -0,466969136 | 0,819 | 0,899 | 2,14E-12 | 2 |
| *svp.1* | 2,08E-16 | -0,596096182 | 0,409 | 0,567 | 2,49E-12 | 2 |
| *mrva.1* | 2,90E-16 | 0,717466906 | 0,397 | 0,266 | 3,45E-12 | 2 |
| *lolal.1* | 3,21E-16 | -0,668419323 | 0,194 | 0,344 | 3,83E-12 | 2 |
| *Mitf.1* | 3,28E-16 | -0,7047943 | 0,192 | 0,332 | 3,91E-12 | 2 |
| *CG13887* | 3,93E-16 | 0,632191316 | 0,25 | 0,133 | 4,69E-12 | 2 |
| *poe.1* | 4,37E-16 | 0,540654542 | 0,453 | 0,302 | 5,21E-12 | 2 |
| *Sirt1* | 4,59E-16 | 0,714929988 | 0,317 | 0,19 | 5,47E-12 | 2 |
| *baz.1* | 4,60E-16 | 0,624997058 | 0,336 | 0,202 | 5,48E-12 | 2 |
| *kst.1* | 5,90E-16 | -0,950626474 | 0,184 | 0,322 | 7,04E-12 | 2 |
| *CG12065.1* | 6,37E-16 | 0,702925074 | 0,398 | 0,261 | 7,60E-12 | 2 |
| *Galk.1* | 6,60E-16 | -0,718265353 | 0,402 | 0,535 | 7,87E-12 | 2 |
| *Diap1.1* | 6,83E-16 | 0,466171413 | 0,801 | 0,73 | 8,15E-12 | 2 |
| *Snx6* | 6,99E-16 | 0,612424091 | 0,288 | 0,162 | 8,34E-12 | 2 |
| *Galt.1* | 8,08E-16 | -0,732817158 | 0,117 | 0,251 | 9,63E-12 | 2 |
| *PHGPx.1* | 8,28E-16 | 0,664400957 | 0,478 | 0,343 | 9,87E-12 | 2 |
| *Clk.1* | 9,20E-16 | -0,769035532 | 0,135 | 0,268 | 1,10E-11 | 2 |
| *Cyp4g1.1* | 9,43E-16 | 0,886537139 | 0,473 | 0,335 | 1,12E-11 | 2 |
| *Gprk1.1* | 9,87E-16 | -0,645624913 | 0,236 | 0,384 | 1,18E-11 | 2 |
| *Rac2.1* | 1,04E-15 | 0,569593253 | 0,412 | 0,274 | 1,24E-11 | 2 |
| *IM33.1* | 1,24E-15 | -0,852345357 | 0,255 | 0,406 | 1,48E-11 | 2 |

| | | | | | | |
|---|---|---|---|---|---|---|
| *santa-maria.1* | 1,30E-15 | -0,658077144 | 0,154 | 0,298 | 1,55E-11 | 2 |
| *Rab5.1* | 1,52E-15 | 0,596777988 | 0,445 | 0,314 | 1,82E-11 | 2 |
| *Cf2.1* | 1,55E-15 | -0,645158341 | 0,242 | 0,385 | 1,85E-11 | 2 |
| *Bsg* | 1,56E-15 | 0,276081548 | 0,977 | 0,984 | 1,86E-11 | 2 |
| *kibra.1* | 1,66E-15 | 0,582321498 | 0,547 | 0,411 | 1,98E-11 | 2 |
| *Pli.1* | 1,71E-15 | -0,841761599 | 0,477 | 0,594 | 2,04E-11 | 2 |
| *SP1173.1* | 1,71E-15 | 0,523913869 | 0,288 | 0,16 | 2,04E-11 | 2 |
| *E(spl)malpha-BFM* | 1,84E-15 | -0,585211723 | 0,231 | 0,385 | 2,19E-11 | 2 |
| *Jafrac1.1* | 1,93E-15 | 0,539525303 | 0,5 | 0,356 | 2,30E-11 | 2 |
| *Syx1A.1* | 2,42E-15 | 0,527104267 | 0,637 | 0,519 | 2,89E-11 | 2 |
| *CG3829.1* | 2,68E-15 | -0,66294992 | 0,218 | 0,36 | 3,20E-11 | 2 |
| *fz2* | 2,72E-15 | -0,896321832 | 0,176 | 0,316 | 3,24E-11 | 2 |
| *hfp.1* | 2,80E-15 | -0,698398086 | 0,202 | 0,336 | 3,34E-11 | 2 |
| *trx.1* | 2,82E-15 | -0,517582284 | 0,383 | 0,539 | 3,36E-11 | 2 |
| *LPCAT.1* | 3,29E-15 | 0,602095107 | 0,298 | 0,174 | 3,93E-11 | 2 |
| *Mical.1* | 3,89E-15 | -0,790410184 | 0,201 | 0,343 | 4,65E-11 | 2 |
| *eIF4A.1* | 4,04E-15 | 0,494737152 | 0,687 | 0,579 | 4,82E-11 | 2 |
| *fs(1)h.1* | 7,03E-15 | -0,434161693 | 0,633 | 0,755 | 8,39E-11 | 2 |
| *Stlk.1* | 7,13E-15 | -0,673425523 | 0,194 | 0,329 | 8,51E-11 | 2 |
| *Gbp3.1* | 7,34E-15 | -0,704404724 | 0,168 | 0,301 | 8,75E-11 | 2 |
| *blw.1* | 9,55E-15 | -0,612021381 | 0,307 | 0,456 | 1,14E-10 | 2 |
| *Imp.1* | 1,28E-14 | -0,508643864 | 0,504 | 0,64 | 1,53E-10 | 2 |
| *Ubqn.1* | 1,35E-14 | 0,575825655 | 0,353 | 0,224 | 1,62E-10 | 2 |
| *mub.1* | 1,58E-14 | -0,655280748 | 0,128 | 0,256 | 1,89E-10 | 2 |
| *pst.1* | 1,69E-14 | 0,494516861 | 0,661 | 0,558 | 2,01E-10 | 2 |
| *RpL27.1* | 1,70E-14 | 0,685071655 | 0,671 | 0,588 | 2,02E-10 | 2 |
| *Lk6.1* | 2,38E-14 | 0,31556202 | 0,928 | 0,886 | 2,83E-10 | 2 |
| *ppl.1* | 2,49E-14 | -0,683638942 | 0,131 | 0,261 | 2,97E-10 | 2 |
| *Lmpt.1* | 4,04E-14 | 0,5321543 | 0,553 | 0,419 | 4,82E-10 | 2 |
| *wun* | 4,06E-14 | 0,585155786 | 0,273 | 0,155 | 4,85E-10 | 2 |
| *Sam-S.1* | 4,59E-14 | -0,75254929 | 0,516 | 0,646 | 5,48E-10 | 2 |
| *CG42324.1* | 4,75E-14 | -0,486271501 | 0,553 | 0,669 | 5,67E-10 | 2 |
| *Btk29A* | 4,95E-14 | -0,781246237 | 0,14 | 0,266 | 5,91E-10 | 2 |
| *CG6910.1* | 5,10E-14 | 0,456076735 | 0,839 | 0,732 | 6,08E-10 | 2 |
| *BomS2* | 5,43E-14 | -1,039415433 | 0,296 | 0,428 | 6,48E-10 | 2 |
| *RpL34b.1* | 5,99E-14 | 0,603391209 | 0,61 | 0,512 | 7,15E-10 | 2 |
| *bgm.1* | 6,54E-14 | -0,684951612 | 0,562 | 0,66 | 7,80E-10 | 2 |
| *Eb1.1* | 6,55E-14 | 0,522373618 | 0,345 | 0,216 | 7,81E-10 | 2 |
| *iPLA2-VIA* | 6,68E-14 | 0,537085928 | 0,269 | 0,154 | 7,97E-10 | 2 |
| *Cka.1* | 7,00E-14 | -0,610099575 | 0,216 | 0,353 | 8,36E-10 | 2 |
| *Vha13.1* | 8,84E-14 | 0,7412294 | 0,327 | 0,213 | 1,05E-09 | 2 |
| *ens.1* | 9,55E-14 | 0,529274796 | 0,567 | 0,463 | 1,14E-09 | 2 |
| *CG17691.1* | 9,78E-14 | -0,53475233 | 0,346 | 0,486 | 1,17E-09 | 2 |
| *CG34136.1* | 9,84E-14 | 0,606838778 | 0,371 | 0,24 | 1,17E-09 | 2 |
| *CG3168.1* | 1,12E-13 | 0,561783856 | 0,392 | 0,262 | 1,34E-09 | 2 |
| *bic.1* | 1,21E-13 | 0,530515799 | 0,484 | 0,363 | 1,45E-09 | 2 |
| *sxc.1* | 1,32E-13 | -0,604987428 | 0,24 | 0,375 | 1,57E-09 | 2 |
| *ATPsynC.1* | 1,50E-13 | -0,498194627 | 0,349 | 0,495 | 1,80E-09 | 2 |
| *Myc* | 1,65E-13 | -0,646258433 | 0,586 | 0,682 | 1,97E-09 | 2 |
| *Sh3beta.1* | 2,14E-13 | 0,526912946 | 0,297 | 0,177 | 2,56E-09 | 2 |
| *CG4297.1* | 2,86E-13 | -0,611744416 | 0,143 | 0,268 | 3,41E-09 | 2 |
| *Hn.1* | 2,98E-13 | 0,486327776 | 0,505 | 0,367 | 3,55E-09 | 2 |
| *GstE12.1* | 3,10E-13 | -0,558727085 | 0,156 | 0,285 | 3,69E-09 | 2 |
| *Cdep.1* | 3,28E-13 | -0,641752065 | 0,266 | 0,4 | 3,92E-09 | 2 |
| *Not1.1* | 3,53E-13 | 0,328919366 | 0,806 | 0,723 | 4,21E-09 | 2 |
| *puml.1* | 3,73E-13 | 0,550802041 | 0,299 | 0,182 | 4,45E-09 | 2 |
| *mura.1* | 4,03E-13 | -0,540065542 | 0,406 | 0,534 | 4,81E-09 | 2 |
| *CG10680.1* | 4,98E-13 | -0,557366246 | 0,315 | 0,452 | 5,94E-09 | 2 |
| *Gel.1* | 6,30E-13 | -0,553758027 | 0,36 | 0,495 | 7,51E-09 | 2 |
| *RpS25.1* | 6,69E-13 | 0,65225059 | 0,646 | 0,552 | 7,98E-09 | 2 |
| *RpS10b.1* | 6,86E-13 | 0,603637887 | 0,694 | 0,624 | 8,18E-09 | 2 |

| | | | | | | |
|---|---|---|---|---|---|---|
| *IscU.1* | 6,98E-13 | 0,50437283 | 0,516 | 0,391 | 8,33E-09 | 2 |
| *Fit1.1* | 7,09E-13 | 0,655717307 | 0,315 | 0,2 | 8,46E-09 | 2 |
| *RpL5.1* | 7,90E-13 | 0,49355071 | 0,662 | 0,575 | 9,42E-09 | 2 |
| *RpL31.1* | 8,53E-13 | 0,586627371 | 0,659 | 0,558 | 1,02E-08 | 2 |
| *Sap-r.1* | 8,97E-13 | 0,503659423 | 0,629 | 0,528 | 1,07E-08 | 2 |
| *CG44774.1* | 9,80E-13 | 0,566615425 | 0,345 | 0,227 | 1,17E-08 | 2 |
| *tay.1* | 1,02E-12 | -0,518537633 | 0,312 | 0,448 | 1,22E-08 | 2 |
| *Mal-B2.1* | 1,03E-12 | 0,479483982 | 0,618 | 0,494 | 1,22E-08 | 2 |
| *Pi3K92E* | 1,06E-12 | 0,561246779 | 0,283 | 0,172 | 1,27E-08 | 2 |
| *Nipped-A.1* | 1,08E-12 | -0,541016714 | 0,188 | 0,319 | 1,29E-08 | 2 |
| *RpL35.1* | 1,10E-12 | 0,58983081 | 0,622 | 0,517 | 1,31E-08 | 2 |
| *CG3902.1* | 1,16E-12 | -0,831310398 | 0,217 | 0,338 | 1,38E-08 | 2 |
| *robo2.1* | 1,16E-12 | 0,498486608 | 0,522 | 0,389 | 1,38E-08 | 2 |
| *RpL12.1* | 1,18E-12 | 0,568041798 | 0,656 | 0,575 | 1,41E-08 | 2 |
| *CG3597.1* | 1,20E-12 | -0,634929138 | 0,155 | 0,275 | 1,43E-08 | 2 |
| *RpS3A.1* | 1,22E-12 | 0,475744067 | 0,648 | 0,544 | 1,46E-08 | 2 |
| *RpS28b.1* | 1,23E-12 | 0,687589907 | 0,565 | 0,474 | 1,46E-08 | 2 |
| *Nrg* | 1,53E-12 | -0,718952687 | 0,176 | 0,299 | 1,83E-08 | 2 |
| *RpL8.1* | 1,54E-12 | 0,480363615 | 0,74 | 0,643 | 1,83E-08 | 2 |
| *Synd.1* | 1,55E-12 | 0,558311423 | 0,364 | 0,245 | 1,84E-08 | 2 |
| *RpLP1.1* | 1,69E-12 | 0,527908491 | 0,708 | 0,604 | 2,02E-08 | 2 |
| *CG9451.1* | 1,77E-12 | 0,601800974 | 0,319 | 0,202 | 2,12E-08 | 2 |
| *CG18659.1* | 1,85E-12 | 0,635686954 | 0,345 | 0,23 | 2,20E-08 | 2 |
| *MESR3* | 1,89E-12 | 0,660394526 | 0,301 | 0,186 | 2,25E-08 | 2 |
| *comm2.1* | 1,97E-12 | 0,368964612 | 0,364 | 0,229 | 2,35E-08 | 2 |
| *PNUTS* | 2,02E-12 | -0,44744698 | 0,522 | 0,639 | 2,41E-08 | 2 |
| *KrT95D.1* | 2,47E-12 | 0,431264869 | 0,755 | 0,661 | 2,95E-08 | 2 |
| *RpS27A.1* | 2,59E-12 | 0,588193699 | 0,708 | 0,636 | 3,09E-08 | 2 |
| *SREBP.1* | 2,91E-12 | 0,479171507 | 0,504 | 0,389 | 3,47E-08 | 2 |
| *Vinc* | 3,12E-12 | 0,525160414 | 0,263 | 0,156 | 3,72E-08 | 2 |
| *Doa.1* | 3,24E-12 | -0,479395918 | 0,423 | 0,553 | 3,87E-08 | 2 |
| *Cyp28d1.1* | 3,42E-12 | 0,484848045 | 0,306 | 0,189 | 4,08E-08 | 2 |
| *CG42668.1* | 3,68E-12 | 0,356465476 | 0,694 | 0,567 | 4,39E-08 | 2 |
| *ImpL2.1* | 3,79E-12 | 0,377577075 | 0,341 | 0,216 | 4,53E-08 | 2 |
| *CG8369* | 4,01E-12 | 0,591119819 | 0,25 | 0,148 | 4,79E-08 | 2 |
| *CdGAPr* | 4,26E-12 | 0,624635567 | 0,269 | 0,165 | 5,08E-08 | 2 |
| *Mlf* | 4,77E-12 | 0,573322719 | 0,253 | 0,15 | 5,69E-08 | 2 |
| *Pitslre* | 4,89E-12 | -0,391351479 | 0,31 | 0,449 | 5,84E-08 | 2 |
| *GstT1* | 5,23E-12 | 0,586161414 | 0,26 | 0,154 | 6,24E-08 | 2 |
| *Ctl2.1* | 5,57E-12 | 0,575997453 | 0,264 | 0,162 | 6,65E-08 | 2 |
| *RpL24.1* | 6,58E-12 | 0,593287065 | 0,678 | 0,581 | 7,85E-08 | 2 |
| *CG31694.1* | 6,66E-12 | 0,385575525 | 0,475 | 0,343 | 7,94E-08 | 2 |
| *Thd1.1* | 6,68E-12 | -0,487076758 | 0,357 | 0,487 | 7,96E-08 | 2 |
| *CG31183* | 7,06E-12 | -0,488523555 | 0,473 | 0,591 | 8,42E-08 | 2 |
| *corto.1* | 7,32E-12 | -0,644671402 | 0,402 | 0,515 | 8,74E-08 | 2 |
| *CG6287* | 8,27E-12 | 0,4443816 | 0,641 | 0,521 | 9,86E-08 | 2 |
| *CG6700.1* | 1,15E-11 | -0,462040739 | 0,312 | 0,449 | 1,37E-07 | 2 |
| *CG15784.1* | 1,26E-11 | 0,609840017 | 0,319 | 0,208 | 1,50E-07 | 2 |
| *sws.1* | 1,36E-11 | -0,577341789 | 0,194 | 0,316 | 1,62E-07 | 2 |
| *Col4a1* | 1,59E-11 | -0,655930375 | 0,142 | 0,258 | 1,89E-07 | 2 |
| *IP3K2* | 1,77E-11 | -0,478118244 | 0,589 | 0,682 | 2,11E-07 | 2 |
| *CG31705.1* | 1,90E-11 | 0,392273768 | 0,646 | 0,545 | 2,27E-07 | 2 |
| *lap.1* | 1,97E-11 | -0,599043794 | 0,22 | 0,334 | 2,34E-07 | 2 |
| *Sik3.1* | 2,07E-11 | -0,44114693 | 0,405 | 0,534 | 2,47E-07 | 2 |
| *CtBP* | 2,18E-11 | -0,397826184 | 0,519 | 0,638 | 2,60E-07 | 2 |
| *PAPLA1.1* | 2,24E-11 | 0,353176372 | 0,593 | 0,461 | 2,67E-07 | 2 |
| *RpL35A.1* | 2,24E-11 | 0,529063693 | 0,622 | 0,533 | 2,67E-07 | 2 |
| *CG42369.1* | 2,31E-11 | -0,620637516 | 0,145 | 0,252 | 2,76E-07 | 2 |
| *CtsB1* | 2,32E-11 | 0,50962598 | 0,585 | 0,478 | 2,77E-07 | 2 |
| *RpS5a.1* | 2,32E-11 | 0,522330098 | 0,586 | 0,492 | 2,77E-07 | 2 |
| *Apoltp.1* | 3,79E-11 | 0,257670347 | 0,855 | 0,773 | 4,53E-07 | 2 |

| | | | | | | |
|---|---|---|---|---|---|---|
| *RpL17.1* | 4,02E-11 | 0,500224753 | 0,737 | 0,656 | 4,79E-07 | 2 |
| *cu.1* | 4,25E-11 | 0,465998729 | 0,572 | 0,461 | 5,07E-07 | 2 |
| *CG44014* | 4,82E-11 | 0,539736633 | 0,253 | 0,153 | 5,75E-07 | 2 |
| *Cbs.1* | 5,36E-11 | -0,629612674 | 0,175 | 0,287 | 6,39E-07 | 2 |
| *CG7220.1* | 5,73E-11 | -0,695928158 | 0,151 | 0,255 | 6,83E-07 | 2 |
| *Sin3A.1* | 5,90E-11 | -0,512545728 | 0,382 | 0,494 | 7,04E-07 | 2 |
| *CG34347.1* | 7,12E-11 | -0,8646885 | 0,299 | 0,404 | 8,49E-07 | 2 |
| *CG33494.1* | 7,13E-11 | 0,463857152 | 0,411 | 0,289 | 8,51E-07 | 2 |
| *RpS23.1* | 7,49E-11 | 0,534974344 | 0,734 | 0,658 | 8,94E-07 | 2 |
| *bip2.1* | 8,71E-11 | -0,462603134 | 0,381 | 0,501 | 1,04E-06 | 2 |
| *E(bx).1* | 9,94E-11 | -0,587716336 | 0,188 | 0,295 | 1,19E-06 | 2 |
| *Letm1.1* | 1,12E-10 | 0,618865764 | 0,338 | 0,237 | 1,33E-06 | 2 |
| *Ptpmeg2* | 1,14E-10 | -0,643889287 | 0,151 | 0,255 | 1,36E-06 | 2 |
| *mts.1* | 1,15E-10 | 0,458127097 | 0,451 | 0,344 | 1,38E-06 | 2 |
| *Blimp-1.1* | 1,16E-10 | -0,409583288 | 0,619 | 0,725 | 1,38E-06 | 2 |
| *Trpm.1* | 1,26E-10 | -0,458388319 | 0,242 | 0,363 | 1,50E-06 | 2 |
| *RpL18A.1* | 1,29E-10 | 0,554353494 | 0,671 | 0,604 | 1,54E-06 | 2 |
| *p23.1* | 1,30E-10 | 0,545563469 | 0,319 | 0,216 | 1,55E-06 | 2 |
| *Mvl* | 1,45E-10 | 0,496146021 | 0,251 | 0,155 | 1,73E-06 | 2 |
| *eEF5.1* | 1,46E-10 | 0,602733189 | 0,608 | 0,519 | 1,74E-06 | 2 |
| *Aldh* | 1,87E-10 | -0,502216192 | 0,459 | 0,571 | 2,24E-06 | 2 |
| *CG31678.1* | 2,14E-10 | 0,427161237 | 0,302 | 0,193 | 2,56E-06 | 2 |
| *RpS29.1* | 2,44E-10 | 0,586853614 | 0,636 | 0,554 | 2,91E-06 | 2 |
| *Tl.1* | 2,60E-10 | 0,426591464 | 0,657 | 0,569 | 3,10E-06 | 2 |
| *RpS7.1* | 2,90E-10 | 0,481203245 | 0,735 | 0,661 | 3,46E-06 | 2 |
| *mei-P26.1* | 3,12E-10 | -0,440240933 | 0,341 | 0,469 | 3,72E-06 | 2 |
| *swm.1* | 4,00E-10 | 0,482826348 | 0,412 | 0,301 | 4,77E-06 | 2 |
| *CG6051.1* | 4,15E-10 | 0,325568093 | 0,631 | 0,517 | 4,94E-06 | 2 |
| *CG1943.1* | 4,36E-10 | 0,522200183 | 0,286 | 0,188 | 5,20E-06 | 2 |
| *RpL7.1* | 4,77E-10 | 0,503379617 | 0,665 | 0,587 | 5,69E-06 | 2 |
| *CG8206* | 4,96E-10 | 0,528753479 | 0,288 | 0,188 | 5,92E-06 | 2 |
| *RpS12.1* | 5,13E-10 | 0,606399424 | 0,651 | 0,582 | 6,12E-06 | 2 |
| *eEF2.1* | 5,19E-10 | 0,314178729 | 0,802 | 0,741 | 6,19E-06 | 2 |
| *Gbs-76A.1* | 6,01E-10 | -0,460411925 | 0,321 | 0,444 | 7,17E-06 | 2 |
| *cv-2* | 6,19E-10 | -0,590973858 | 0,401 | 0,5 | 7,38E-06 | 2 |
| *alt.1* | 7,18E-10 | 0,516433967 | 0,431 | 0,334 | 8,57E-06 | 2 |
| *fog.1* | 7,83E-10 | -0,577042305 | 0,154 | 0,254 | 9,34E-06 | 2 |
| *Hel89B* | 8,21E-10 | 0,517689524 | 0,272 | 0,174 | 9,79E-06 | 2 |
| *Mst84Da.1* | 8,30E-10 | 0,664681339 | 0,395 | 0,296 | 9,90E-06 | 2 |
| *CG4538.1* | 8,52E-10 | 0,461611388 | 0,365 | 0,262 | 1,02E-05 | 2 |
| *Set1.1* | 8,72E-10 | 0,42557502 | 0,433 | 0,321 | 1,04E-05 | 2 |
| *CG12290.1* | 8,80E-10 | 0,400678933 | 0,364 | 0,248 | 1,05E-05 | 2 |
| *l(3)05822.1* | 9,30E-10 | 0,473007208 | 0,284 | 0,186 | 1,11E-05 | 2 |
| *Rab2* | 9,73E-10 | 0,534561654 | 0,268 | 0,175 | 1,16E-05 | 2 |
| *Sfxn1-3.1* | 1,02E-09 | 0,468369054 | 0,439 | 0,338 | 1,22E-05 | 2 |
| *wdb.1* | 1,13E-09 | -0,376810969 | 0,584 | 0,674 | 1,35E-05 | 2 |
| *DOR.1* | 1,19E-09 | -0,634567256 | 0,551 | 0,628 | 1,42E-05 | 2 |
| *RpL15.1* | 1,29E-09 | 0,529931056 | 0,701 | 0,63 | 1,53E-05 | 2 |
| *mbc* | 1,42E-09 | -0,454391583 | 0,178 | 0,289 | 1,70E-05 | 2 |
| *CG6770.1* | 1,46E-09 | -0,314870661 | 0,904 | 0,924 | 1,75E-05 | 2 |
| *caps.1* | 1,48E-09 | 0,424253524 | 0,577 | 0,476 | 1,76E-05 | 2 |
| *CG17734.1* | 1,62E-09 | -0,520604654 | 0,178 | 0,28 | 1,93E-05 | 2 |
| *CG16721* | 1,67E-09 | 0,457040514 | 0,284 | 0,185 | 1,99E-05 | 2 |
| *Gmap.1* | 1,79E-09 | -0,314432215 | 0,379 | 0,509 | 2,14E-05 | 2 |
| *Unc-76* | 1,95E-09 | 0,46962764 | 0,315 | 0,212 | 2,33E-05 | 2 |
| *CG7530.1* | 2,25E-09 | -0,612605885 | 0,226 | 0,328 | 2,68E-05 | 2 |
| *CG10543* | 2,45E-09 | -0,539608628 | 0,245 | 0,346 | 2,92E-05 | 2 |
| *Gale.1* | 2,49E-09 | -0,598903643 | 0,166 | 0,262 | 2,97E-05 | 2 |
| *Crag* | 2,59E-09 | 0,514696463 | 0,284 | 0,188 | 3,08E-05 | 2 |
| *east.1* | 2,82E-09 | -0,537926896 | 0,164 | 0,263 | 3,36E-05 | 2 |
| *CG10799.1* | 2,94E-09 | 0,363185737 | 0,579 | 0,478 | 3,50E-05 | 2 |

| | | | | | | |
|---|---|---|---|---|---|---|
| *RpS27.1* | 3,70E-09 | 0,451577723 | 0,699 | 0,641 | 4,41E-05 | 2 |
| *CG32549* | 3,82E-09 | -0,433519202 | 0,607 | 0,701 | 4,56E-05 | 2 |
| *CG8963* | 4,11E-09 | 0,432394542 | 0,273 | 0,177 | 4,91E-05 | 2 |
| *Pzl.1* | 4,23E-09 | -0,497197892 | 0,184 | 0,287 | 5,04E-05 | 2 |
| *Trp1.1* | 4,25E-09 | 0,494500053 | 0,306 | 0,212 | 5,07E-05 | 2 |
| *Ubc6.1* | 4,64E-09 | 0,432314889 | 0,345 | 0,245 | 5,54E-05 | 2 |
| *jing* | 4,74E-09 | -0,472109612 | 0,157 | 0,257 | 5,66E-05 | 2 |
| *D2hgdh.1* | 5,13E-09 | -0,531951626 | 0,228 | 0,331 | 6,11E-05 | 2 |
| *ttk* | 5,39E-09 | -0,514564979 | 0,176 | 0,278 | 6,43E-05 | 2 |
| *MSBP.1* | 6,59E-09 | 0,489794658 | 0,357 | 0,262 | 7,87E-05 | 2 |
| *how* | 8,01E-09 | -0,362791733 | 0,467 | 0,567 | 9,55E-05 | 2 |
| *Pect.1* | 9,07E-09 | -0,610480613 | 0,504 | 0,578 | 0,000108231 | 2 |
| *Cyp1.1* | 9,93E-09 | 0,446331859 | 0,458 | 0,37 | 0,000118453 | 2 |
| *prage* | 1,00E-08 | 0,322218421 | 0,801 | 0,718 | 0,000119468 | 2 |
| *kra.1* | 1,00E-08 | 0,342100399 | 0,613 | 0,525 | 0,000119536 | 2 |
| *RpL32.1* | 1,19E-08 | 0,484084515 | 0,673 | 0,616 | 0,000141798 | 2 |
| *sesB.1* | 1,21E-08 | -0,400801983 | 0,414 | 0,518 | 0,000144438 | 2 |
| *CG2991.1* | 1,26E-08 | 0,334249918 | 0,268 | 0,175 | 0,000149882 | 2 |
| *CG13868.1* | 1,28E-08 | -0,370291814 | 0,702 | 0,761 | 0,000153282 | 2 |
| *Ank.1* | 1,48E-08 | -0,392575053 | 0,418 | 0,515 | 0,000176167 | 2 |
| *mnb* | 1,61E-08 | -0,528289941 | 0,339 | 0,429 | 0,000192504 | 2 |
| *siz.1* | 1,77E-08 | 0,425370843 | 0,543 | 0,446 | 0,000210844 | 2 |
| *Galphao.1* | 2,03E-08 | -0,357609094 | 0,482 | 0,584 | 0,000242735 | 2 |
| *Stam.1* | 2,15E-08 | 0,408968393 | 0,348 | 0,251 | 0,000256293 | 2 |
| *RpL13A.1* | 2,21E-08 | 0,436760489 | 0,692 | 0,635 | 0,000263388 | 2 |
| *eEF1beta.1* | 2,59E-08 | 0,486808763 | 0,442 | 0,355 | 0,000308866 | 2 |
| *hep.1* | 2,61E-08 | 0,526291871 | 0,386 | 0,296 | 0,000310857 | 2 |
| *Uba1.1* | 2,63E-08 | 0,357867688 | 0,269 | 0,177 | 0,000314161 | 2 |
| *app.1* | 2,69E-08 | -0,454544381 | 0,312 | 0,415 | 0,000320455 | 2 |
| *RpS13* | 2,83E-08 | 0,416336603 | 0,654 | 0,584 | 0,000337793 | 2 |
| *hang* | 2,93E-08 | -0,42942864 | 0,222 | 0,326 | 0,0003495 | 2 |
| *CkIIbeta.1* | 2,98E-08 | -0,444892749 | 0,225 | 0,323 | 0,000355292 | 2 |
| *Atg4a* | 3,07E-08 | 0,494765374 | 0,273 | 0,187 | 0,000365998 | 2 |
| *RpS8.1* | 3,18E-08 | 0,469550432 | 0,749 | 0,701 | 0,000379206 | 2 |
| *Mapmodulin.1* | 3,43E-08 | -0,46722391 | 0,174 | 0,271 | 0,000409482 | 2 |
| *CG14073.1* | 3,66E-08 | 0,39054142 | 0,473 | 0,373 | 0,000436016 | 2 |
| *RpL24-like* | 3,74E-08 | 0,521046124 | 0,254 | 0,172 | 0,000445622 | 2 |
| *Hrb27C.1* | 3,99E-08 | -0,306347093 | 0,692 | 0,773 | 0,000475955 | 2 |
| *CG33307* | 4,38E-08 | 0,570333935 | 0,272 | 0,187 | 0,000522825 | 2 |
| *RpL4.1* | 4,79E-08 | 0,450463349 | 0,602 | 0,53 | 0,000571411 | 2 |
| *Map205.1* | 4,84E-08 | -0,592290794 | 0,477 | 0,562 | 0,00057766 | 2 |
| *Pkn.1* | 4,87E-08 | -0,404993243 | 0,511 | 0,601 | 0,000581025 | 2 |
| *Got2.1* | 5,11E-08 | 0,376920033 | 0,619 | 0,526 | 0,00060983 | 2 |
| *lbk.1* | 5,81E-08 | 0,425171364 | 0,335 | 0,245 | 0,000693669 | 2 |
| *ttv* | 5,90E-08 | -0,434895789 | 0,239 | 0,337 | 0,000703886 | 2 |
| *CG4199.1* | 6,00E-08 | 0,353138482 | 0,313 | 0,218 | 0,000716326 | 2 |
| *Pgm1.1* | 7,53E-08 | -0,416090577 | 0,185 | 0,278 | 0,000898489 | 2 |
| *alpha-Cat* | 7,61E-08 | -0,448241371 | 0,173 | 0,264 | 0,000907881 | 2 |
| *flfl.1* | 7,69E-08 | -0,448901732 | 0,228 | 0,323 | 0,000917203 | 2 |
| *Tango1.1* | 7,84E-08 | 0,402377234 | 0,277 | 0,192 | 0,000934879 | 2 |
| *CG12567.1* | 8,86E-08 | -0,414572684 | 0,226 | 0,326 | 0,001056794 | 2 |
| *CG42268.1* | 8,88E-08 | -0,412008623 | 0,397 | 0,494 | 0,001059661 | 2 |
| *Rim2.1* | 9,37E-08 | -0,533472972 | 0,179 | 0,27 | 0,00111781 | 2 |
| *Xe7.1* | 1,13E-07 | -0,411645666 | 0,236 | 0,331 | 0,001345041 | 2 |
| *par-1* | 1,13E-07 | -0,25234047 | 0,934 | 0,944 | 0,001352867 | 2 |
| *RpL26.1* | 1,33E-07 | 0,458292363 | 0,661 | 0,59 | 0,001588439 | 2 |
| *RpL9.1* | 1,36E-07 | 0,451399679 | 0,609 | 0,53 | 0,001626668 | 2 |
| *RpL10.1* | 1,48E-07 | 0,428417056 | 0,602 | 0,531 | 0,001762498 | 2 |
| *RN-tre.1* | 1,49E-07 | 0,467200288 | 0,277 | 0,194 | 0,001775451 | 2 |
| *RpL19.1* | 1,50E-07 | 0,446278214 | 0,731 | 0,669 | 0,00179479 | 2 |
| *Golgin245.1* | 1,55E-07 | 0,461249461 | 0,282 | 0,2 | 0,00184335 | 2 |

| | | | | | | |
|---|---|---|---|---|---|---|
| *Ntan1.1* | 1,55E-07 | 0,258257017 | 0,669 | 0,585 | 0,00184415 | 2 |
| *RpL18.1* | 1,79E-07 | 0,368921266 | 0,605 | 0,528 | 0,002132549 | 2 |
| *CG32276.1* | 1,96E-07 | 0,478914181 | 0,354 | 0,27 | 0,00233689 | 2 |
| *klar* | 2,08E-07 | -0,409254541 | 0,4 | 0,5 | 0,002478214 | 2 |
| *RpL38.1* | 2,16E-07 | 0,523707479 | 0,452 | 0,368 | 0,002573206 | 2 |
| *Chc.1* | 2,17E-07 | 0,40700487 | 0,365 | 0,282 | 0,002591958 | 2 |
| *Akt1.1* | 2,32E-07 | 0,378772586 | 0,445 | 0,354 | 0,002773056 | 2 |
| *RapGAP1.1* | 2,37E-07 | -0,506019025 | 0,188 | 0,275 | 0,002825713 | 2 |
| *crc.1* | 2,38E-07 | 0,361675896 | 0,445 | 0,349 | 0,002842328 | 2 |
| *RpL3.1* | 2,45E-07 | 0,354985251 | 0,687 | 0,628 | 0,002924717 | 2 |
| *ctp.1* | 2,58E-07 | 0,378082215 | 0,358 | 0,271 | 0,003077537 | 2 |
| *Thor.1* | 2,75E-07 | 0,500436181 | 0,599 | 0,516 | 0,00327991 | 2 |
| *sowah.1* | 2,80E-07 | 0,475100702 | 0,395 | 0,307 | 0,003341006 | 2 |
| *RpS18* | 2,81E-07 | 0,377600801 | 0,758 | 0,71 | 0,003351456 | 2 |
| *RpL11.1* | 3,07E-07 | 0,473712664 | 0,558 | 0,497 | 0,003659321 | 2 |
| *NKAIN.1* | 3,42E-07 | 0,405446816 | 0,373 | 0,29 | 0,004079765 | 2 |
| *luna.1* | 3,67E-07 | -0,307695824 | 0,67 | 0,736 | 0,004376049 | 2 |
| *hyd* | 3,68E-07 | 0,371243 | 0,256 | 0,175 | 0,004384444 | 2 |
| *Top1.1* | 3,79E-07 | -0,412319698 | 0,258 | 0,349 | 0,004526426 | 2 |
| *cnn.1* | 3,84E-07 | 0,338827038 | 0,345 | 0,254 | 0,004578601 | 2 |
| *Bruce.1* | 4,23E-07 | 0,371494589 | 0,433 | 0,35 | 0,005051194 | 2 |
| *CG15771* | 5,77E-07 | -0,461395669 | 0,223 | 0,312 | 0,006888232 | 2 |
| *RpS4.1* | 5,82E-07 | 0,421230536 | 0,618 | 0,562 | 0,006945688 | 2 |
| *bol.1* | 5,98E-07 | -0,559939305 | 0,204 | 0,285 | 0,007134125 | 2 |
| *CG11400.1* | 6,15E-07 | -0,372772079 | 0,311 | 0,41 | 0,007332404 | 2 |
| *dally.1* | 6,32E-07 | -0,584418667 | 0,209 | 0,287 | 0,007536415 | 2 |
| *CG32695* | 6,58E-07 | 0,475701944 | 0,426 | 0,347 | 0,007853326 | 2 |
| *CG34126* | 6,67E-07 | 0,427091282 | 0,26 | 0,183 | 0,007956742 | 2 |
| *CG3164.1* | 6,95E-07 | -0,280577639 | 0,449 | 0,543 | 0,008289998 | 2 |
| *Nap1.1* | 6,99E-07 | 0,407297029 | 0,316 | 0,235 | 0,00833929 | 2 |
| *slmb.1* | 7,18E-07 | 0,378113967 | 0,272 | 0,191 | 0,008570195 | 2 |
| *tim.1* | 7,54E-07 | -0,356889483 | 0,565 | 0,649 | 0,008991007 | 2 |
| *csw* | 8,15E-07 | 0,485846335 | 0,509 | 0,451 | 0,009724812 | 2 |
| *RpS30.1* | 8,80E-07 | 0,474492325 | 0,574 | 0,508 | 0,010496425 | 2 |
| *Ubi-p5E.1* | 8,93E-07 | 0,384521798 | 0,33 | 0,248 | 0,010652043 | 2 |
| *Ac76E* | 9,46E-07 | -0,326983682 | 0,297 | 0,398 | 0,011281963 | 2 |
| *Nacalpha.1* | 9,73E-07 | 0,414457494 | 0,414 | 0,34 | 0,011601799 | 2 |
| *Pgant5.1* | 9,83E-07 | -0,400359972 | 0,203 | 0,289 | 0,011726169 | 2 |
| *rudhira* | 1,01E-06 | -0,333205747 | 0,627 | 0,702 | 0,012000073 | 2 |
| *ltl.1* | 1,17E-06 | 0,429010304 | 0,283 | 0,204 | 0,013937104 | 2 |
| *Slik* | 1,18E-06 | 0,373465596 | 0,475 | 0,403 | 0,014084993 | 2 |
| *RpL6* | 1,28E-06 | 0,331543151 | 0,628 | 0,569 | 0,01523955 | 2 |
| *CG10077.1* | 1,29E-06 | -0,25189037 | 0,56 | 0,649 | 0,015400928 | 2 |
| *Atx2* | 1,34E-06 | -0,352982529 | 0,183 | 0,268 | 0,016011204 | 2 |
| *eIF4G2.1* | 1,40E-06 | -0,385858202 | 0,266 | 0,356 | 0,016660354 | 2 |
| *CG12163.1* | 1,54E-06 | -0,447856897 | 0,449 | 0,521 | 0,018369379 | 2 |
| *sm.1* | 1,57E-06 | -0,450687429 | 0,24 | 0,325 | 0,018785882 | 2 |
| *B52.1* | 1,60E-06 | -0,30464091 | 0,25 | 0,34 | 0,019054469 | 2 |
| *CG2852.1* | 1,62E-06 | 0,466711261 | 0,32 | 0,247 | 0,019353395 | 2 |
| *Cat.1* | 1,79E-06 | 0,407428525 | 0,41 | 0,329 | 0,021338823 | 2 |
| *Gyf* | 1,93E-06 | -0,369329292 | 0,358 | 0,441 | 0,023029342 | 2 |
| *RpL37A.1* | 2,23E-06 | 0,455459916 | 0,533 | 0,472 | 0,026572728 | 2 |
| *rin* | 2,51E-06 | -0,319086035 | 0,316 | 0,415 | 0,029893519 | 2 |
| *Nup153* | 2,51E-06 | -0,336188362 | 0,251 | 0,341 | 0,029912776 | 2 |
| *CASK.1* | 3,05E-06 | -0,329540693 | 0,264 | 0,351 | 0,036431979 | 2 |
| *Cerk.1* | 3,10E-06 | 0,399065942 | 0,379 | 0,3 | 0,036984387 | 2 |
| *Npl4.1* | 3,17E-06 | 0,351937534 | 0,27 | 0,197 | 0,037808281 | 2 |
| *p120ctn.1* | 3,19E-06 | -0,361379966 | 0,291 | 0,376 | 0,038026581 | 2 |
| *lost.1* | 3,38E-06 | 0,353789535 | 0,453 | 0,371 | 0,040304103 | 2 |
| *CG5773.1* | 3,43E-06 | 0,334832284 | 0,293 | 0,21 | 0,040889108 | 2 |
| *Oatp30B.1* | 4,03E-06 | -0,361635738 | 0,522 | 0,609 | 0,048031111 | 2 |

| | | | | | | |
|---|---|---|---|---|---|---|
| *Scamp.1* | 4,24E-06 | 0,309194744 | 0,313 | 0,233 | 0,050587535 | 2 |
| *chb.1* | 4,35E-06 | 0,370280075 | 0,352 | 0,275 | 0,05186704 | 2 |
| *RpS21.1* | 4,97E-06 | 0,484284668 | 0,605 | 0,559 | 0,059306099 | 2 |
| *Mad.1* | 6,02E-06 | 0,42630173 | 0,386 | 0,309 | 0,071818199 | 2 |
| *C3G.1* | 6,12E-06 | -0,374481407 | 0,174 | 0,251 | 0,07297919 | 2 |
| *CG32066.1* | 6,15E-06 | 0,296715145 | 0,497 | 0,417 | 0,073359203 | 2 |
| *CG9331* | 6,26E-06 | -0,390039396 | 0,24 | 0,322 | 0,074631341 | 2 |
| *sno.1* | 7,29E-06 | 0,332858361 | 0,299 | 0,226 | 0,087021624 | 2 |
| *CG15098.1* | 7,30E-06 | 0,428109195 | 0,44 | 0,37 | 0,087042607 | 2 |
| *mim.1* | 8,48E-06 | -0,32840762 | 0,327 | 0,408 | 0,101145557 | 2 |
| *Cys.1* | 9,00E-06 | 0,512055434 | 0,343 | 0,28 | 0,107417545 | 2 |
| *Fkbp14* | 9,36E-06 | -0,384495834 | 0,217 | 0,292 | 0,111659781 | 2 |
| *Unr.1* | 9,41E-06 | 0,342936373 | 0,747 | 0,73 | 0,112268864 | 2 |
| *Pde11.1* | 9,53E-06 | -0,314372723 | 0,547 | 0,605 | 0,113637695 | 2 |
| *CG32512* | 9,58E-06 | -0,36625384 | 0,184 | 0,259 | 0,114239258 | 2 |
| *CG1677.1* | 1,00E-05 | -0,338258244 | 0,231 | 0,307 | 0,119634585 | 2 |
| *nmo.1* | 1,18E-05 | 0,279705153 | 0,638 | 0,557 | 0,140311286 | 2 |
| *Gs2* | 1,28E-05 | -0,561432054 | 0,274 | 0,345 | 0,152292614 | 2 |
| *GILT2* | 1,39E-05 | 0,454365326 | 0,278 | 0,217 | 0,165706621 | 2 |
| *Tango5.1* | 1,42E-05 | 0,415710795 | 0,405 | 0,34 | 0,168937183 | 2 |
| *Cirl.1* | 1,53E-05 | 0,365281605 | 0,42 | 0,351 | 0,182348898 | 2 |
| *Plod.1* | 1,53E-05 | 0,296185558 | 0,344 | 0,263 | 0,182937042 | 2 |
| *Cals.1* | 1,55E-05 | -0,370172301 | 0,225 | 0,294 | 0,184603511 | 2 |
| *cathD.1* | 1,57E-05 | 0,345664532 | 0,334 | 0,262 | 0,187811688 | 2 |
| *RpL36.1* | 1,63E-05 | 0,400128477 | 0,646 | 0,594 | 0,194552676 | 2 |
| *chrb.1* | 1,65E-05 | 0,285153274 | 0,292 | 0,215 | 0,197136766 | 2 |
| *Dyrk3* | 1,70E-05 | -0,261051634 | 0,445 | 0,529 | 0,203096271 | 2 |
| *Naprt.1* | 1,77E-05 | 0,431429059 | 0,415 | 0,347 | 0,211423807 | 2 |
| *RpS9.1* | 1,78E-05 | 0,362757219 | 0,631 | 0,579 | 0,211883889 | 2 |
| *Itpr.1* | 1,90E-05 | -0,307609674 | 0,265 | 0,342 | 0,226535824 | 2 |
| *spir.1* | 1,95E-05 | 0,26214016 | 0,426 | 0,352 | 0,233081648 | 2 |
| *Mrtf.1* | 2,03E-05 | 0,437048941 | 0,327 | 0,26 | 0,24228033 | 2 |
| *RpL7A.1* | 2,06E-05 | 0,379160153 | 0,647 | 0,616 | 0,245186802 | 2 |
| *gro* | 2,43E-05 | -0,321743849 | 0,231 | 0,307 | 0,289777326 | 2 |
| *Tep2* | 2,59E-05 | -0,309626182 | 0,234 | 0,312 | 0,308692854 | 2 |
| *RpL36A.1* | 2,61E-05 | 0,43111419 | 0,61 | 0,561 | 0,310943899 | 2 |
| *dlg1.1* | 3,26E-05 | -0,307419878 | 0,341 | 0,419 | 0,388339206 | 2 |
| *Nedd4.1* | 3,31E-05 | 0,338889402 | 0,387 | 0,313 | 0,395207679 | 2 |
| *RpS19a* | 3,84E-05 | 0,343743344 | 0,522 | 0,469 | 0,4576311 | 2 |
| *Sar1* | 3,90E-05 | -0,388572244 | 0,294 | 0,361 | 0,464754437 | 2 |
| *dco.1* | 3,92E-05 | 0,255083722 | 0,355 | 0,277 | 0,468140497 | 2 |
| *Sod1* | 4,14E-05 | 0,433650104 | 0,273 | 0,213 | 0,493756371 | 2 |
| *CG12004.1* | 4,41E-05 | 0,343965909 | 0,321 | 0,252 | 0,525608783 | 2 |
| *noc.1* | 4,65E-05 | 0,26155983 | 0,264 | 0,198 | 0,55467301 | 2 |
| *CG31523.1* | 4,84E-05 | 0,342120992 | 0,312 | 0,248 | 0,57683277 | 2 |
| *UK114.1* | 4,89E-05 | -0,295224348 | 0,363 | 0,439 | 0,582920718 | 2 |
| *AGO3* | 5,59E-05 | -0,466038132 | 0,19 | 0,252 | 0,666375386 | 2 |
| *Rox8.1* | 5,81E-05 | -0,339272216 | 0,228 | 0,3 | 0,693517188 | 2 |
| *Mondo* | 7,03E-05 | -0,328525034 | 0,61 | 0,67 | 0,8384693 | 2 |
| *Non1* | 7,73E-05 | 0,31952501 | 0,398 | 0,334 | 0,921593637 | 2 |
| *Atg17* | 7,87E-05 | 0,278653813 | 0,533 | 0,467 | 0,938404566 | 2 |
| *Gclc.1* | 8,25E-05 | 0,319182475 | 0,58 | 0,514 | 0,98450984 | 2 |
| *Nep4.1* | 8,48E-05 | 0,291036122 | 0,268 | 0,203 | 1 | 2 |
| *Csk.1* | 8,93E-05 | -0,352806228 | 0,43 | 0,497 | 1 | 2 |
| *AGO2.1* | 9,02E-05 | 0,309328358 | 0,284 | 0,219 | 1 | 2 |
| *CG6145.1* | 9,12E-05 | -0,46894544 | 0,396 | 0,458 | 1 | 2 |
| *CG12116.1* | 9,15E-05 | -0,372138789 | 0,359 | 0,431 | 1 | 2 |
| *Vha26.1* | 9,49E-05 | 0,351080491 | 0,308 | 0,246 | 1 | 2 |
| *RpS17.1* | 9,51E-05 | 0,342563064 | 0,58 | 0,527 | 1 | 2 |
| *Crtc* | 0,000107469 | -0,258988977 | 0,454 | 0,522 | 1 | 2 |
| *simj* | 0,000108299 | -0,330689652 | 0,225 | 0,293 | 1 | 2 |

| | | | | | | |
|---|---|---|---|---|---|---|
| *Marf.1* | 0,000112351 | -0,298307109 | 0,26 | 0,328 | 1 | 2 |
| *CG4080.1* | 0,000121232 | 0,407325315 | 0,316 | 0,257 | 1 | 2 |
| *RpS15Aa.1* | 0,000131704 | 0,339646903 | 0,525 | 0,475 | 1 | 2 |
| *pnt.1* | 0,000136986 | -0,304586734 | 0,232 | 0,302 | 1 | 2 |
| *CG5853* | 0,000138928 | -0,377526706 | 0,247 | 0,313 | 1 | 2 |
| *nuf.1* | 0,000150844 | 0,256260688 | 0,594 | 0,533 | 1 | 2 |
| *Dad.1* | 0,000161441 | 0,344053517 | 0,51 | 0,457 | 1 | 2 |
| *ced-6.1* | 0,000171069 | -0,332158641 | 0,435 | 0,492 | 1 | 2 |
| *tara* | 0,000183486 | -0,272530805 | 0,334 | 0,403 | 1 | 2 |
| *path.1* | 0,000189388 | -0,40318908 | 0,925 | 0,875 | 1 | 2 |
| *Rab14* | 0,000192196 | 0,30674985 | 0,263 | 0,204 | 1 | 2 |
| *RpLP2* | 0,000192215 | 0,418469573 | 0,676 | 0,648 | 1 | 2 |
| *pico* | 0,000192865 | -0,348057555 | 0,202 | 0,265 | 1 | 2 |
| *Flo2.1* | 0,000212684 | 0,285719555 | 0,353 | 0,285 | 1 | 2 |
| *Hnf4.1* | 0,000218297 | -0,283312341 | 0,599 | 0,66 | 1 | 2 |
| *CrebA.1* | 0,000223052 | -0,627848691 | 0,204 | 0,269 | 1 | 2 |
| *glec.1* | 0,000239171 | 0,3227838 | 0,411 | 0,346 | 1 | 2 |
| *CG8079* | 0,000251764 | -0,309867838 | 0,246 | 0,306 | 1 | 2 |
| *Pka-R2.1* | 0,00025757 | -0,260196697 | 0,242 | 0,314 | 1 | 2 |
| *Taldo.1* | 0,000267279 | -0,381351338 | 0,255 | 0,316 | 1 | 2 |
| *RpS20.1* | 0,000276325 | 0,374692991 | 0,596 | 0,556 | 1 | 2 |
| *eIF1* | 0,000325806 | 0,3688621 | 0,254 | 0,201 | 1 | 2 |
| *Cyt-b5* | 0,000367558 | 0,363209421 | 0,514 | 0,48 | 1 | 2 |
| *mt:CoIII* | 0,000422856 | 0,432434786 | 0,275 | 0,22 | 1 | 2 |
| *spri.1* | 0,00044779 | -0,276097532 | 0,613 | 0,646 | 1 | 2 |
| *CG5381.1* | 0,000462624 | -0,301377342 | 0,203 | 0,26 | 1 | 2 |
| *galectin.1* | 0,000463891 | -0,271697335 | 0,222 | 0,281 | 1 | 2 |
| *eEF1gamma.1* | 0,00055828 | 0,332793474 | 0,362 | 0,305 | 1 | 2 |
| *RpL13* | 0,000595891 | 0,342806052 | 0,595 | 0,558 | 1 | 2 |
| *wcy* | 0,000629927 | -0,261114953 | 0,208 | 0,263 | 1 | 2 |
| *CG10512* | 0,000656448 | 0,338817233 | 0,255 | 0,198 | 1 | 2 |
| *CNBP* | 0,000683327 | -0,338202904 | 0,212 | 0,267 | 1 | 2 |
| *rdgA* | 0,000807854 | -0,350572757 | 0,195 | 0,251 | 1 | 2 |
| *jar* | 0,000824299 | -0,258264745 | 0,368 | 0,429 | 1 | 2 |
| *Pcf11.1* | 0,000958191 | -0,256147484 | 0,385 | 0,449 | 1 | 2 |
| *RpS16* | 0,001005222 | 0,283901955 | 0,575 | 0,55 | 1 | 2 |
| *Idgf4.1* | 0,001019654 | 0,284164748 | 0,327 | 0,276 | 1 | 2 |
| *Atf3.1* | 0,001145517 | -0,372958871 | 0,254 | 0,308 | 1 | 2 |
| *Sap47* | 0,001310395 | -0,310557665 | 0,319 | 0,378 | 1 | 2 |
| *RpS6.1* | 0,0013716 | 0,30535764 | 0,514 | 0,485 | 1 | 2 |
| *CanA-14F* | 0,001388579 | -0,276859291 | 0,393 | 0,447 | 1 | 2 |
| *mt:CoI.1* | 0,001428926 | 0,319354675 | 0,287 | 0,238 | 1 | 2 |
| *Dp1* | 0,001439262 | -0,280812168 | 0,206 | 0,257 | 1 | 2 |
| *CG33144* | 0,001492649 | -0,268799506 | 0,197 | 0,251 | 1 | 2 |
| *RpS24.1* | 0,001528906 | 0,410726706 | 0,433 | 0,39 | 1 | 2 |
| *UbcE2H.1* | 0,001645288 | -0,318690954 | 0,32 | 0,37 | 1 | 2 |
| *CD98hc.1* | 0,001655977 | 0,259169124 | 0,261 | 0,209 | 1 | 2 |
| *CG18067* | 0,001979431 | -0,850148278 | 0,439 | 0,497 | 1 | 2 |
| *crp.1* | 0,00233069 | -0,311489799 | 0,291 | 0,339 | 1 | 2 |
| *Mlc-c.1* | 0,003329124 | 0,254363456 | 0,373 | 0,327 | 1 | 2 |
| *RpL29.1* | 0,003766553 | 0,26873667 | 0,459 | 0,415 | 1 | 2 |
| *CG7324* | 0,003817764 | -0,250606281 | 0,237 | 0,287 | 1 | 2 |
| *Tctp* | 0,004295222 | 0,263675972 | 0,433 | 0,392 | 1 | 2 |
| *eIF3c.1* | 0,007300731 | 0,285444783 | 0,25 | 0,21 | 1 | 2 |
| *mino* | 0,008627233 | -0,255424776 | 0,665 | 0,678 | 1 | 2 |
| *Nuak1.1* | 0,008837873 | -0,327446657 | 0,246 | 0,291 | 1 | 2 |
| *Eaat1.1* | 0,009623504 | -0,305462171 | 0,298 | 0,348 | 1 | 2 |
| *CG6910.2* | 2,39E-98 | 1,200805164 | 0,963 | 0,708 | 2,85E-94 | 3 |
| *CG3036.2* | 1,07E-69 | 1,285733179 | 0,868 | 0,598 | 1,28E-65 | 3 |
| *klu.1* | 1,17E-69 | 1,243256548 | 0,749 | 0,425 | 1,39E-65 | 3 |
| *Pdp1.2* | 6,19E-66 | 0,602398452 | 0,997 | 0,992 | 7,38E-62 | 3 |

| | | | | | | |
|---|---|---|---|---|---|---|
| *CG17124.2* | 1,69E-63 | -1,500345044 | 0,558 | 0,795 | 2,02E-59 | 3 |
| *CG1673.2* | 3,73E-59 | 0,890585385 | 0,836 | 0,524 | 4,45E-55 | 3 |
| *Helz.2* | 9,97E-59 | 0,944160989 | 0,965 | 0,895 | 1,19E-54 | 3 |
| *mld.1* | 1,40E-56 | 1,538707797 | 0,547 | 0,248 | 1,67E-52 | 3 |
| *CG9498* | 5,85E-51 | 1,272098157 | 0,314 | 0,091 | 6,98E-47 | 3 |
| *Sdc.1* | 1,00E-50 | 0,585329148 | 0,99 | 0,939 | 1,19E-46 | 3 |
| *Prps* | 1,93E-50 | 0,659531545 | 0,987 | 0,909 | 2,30E-46 | 3 |
| *cac.1* | 1,23E-47 | 1,384486136 | 0,669 | 0,403 | 1,47E-43 | 3 |
| *CycG.1* | 2,30E-47 | 0,579822058 | 0,976 | 0,94 | 2,74E-43 | 3 |
| *Nmdmc.2* | 3,60E-47 | 0,8676258 | 0,859 | 0,591 | 4,30E-43 | 3 |
| *CG6770.2* | 3,96E-46 | -0,854162533 | 0,844 | 0,938 | 4,72E-42 | 3 |
| *CG10960.2* | 1,96E-44 | 0,723789445 | 0,937 | 0,807 | 2,34E-40 | 3 |
| *MtnA.2* | 1,05E-42 | 0,938913535 | 0,786 | 0,514 | 1,25E-38 | 3 |
| *Gprk2.1* | 6,82E-42 | 0,858005852 | 0,839 | 0,661 | 8,14E-38 | 3 |
| *Fmo-2.1* | 1,96E-38 | 1,063451623 | 0,42 | 0,185 | 2,34E-34 | 3 |
| *Cam.1* | 1,44E-37 | 0,812681569 | 0,929 | 0,794 | 1,71E-33 | 3 |
| *Gbs-70E.2* | 1,00E-36 | -2,251666332 | 0,383 | 0,566 | 1,20E-32 | 3 |
| *Desat1.2* | 2,70E-36 | 0,577169147 | 0,976 | 0,872 | 3,22E-32 | 3 |
| *CG46385.1* | 2,06E-34 | 0,607598545 | 0,986 | 0,949 | 2,46E-30 | 3 |
| *IP3K2.1* | 7,50E-34 | 0,747387912 | 0,815 | 0,62 | 8,95E-30 | 3 |
| *CG14154.1* | 4,13E-33 | 0,769324455 | 0,664 | 0,413 | 4,92E-29 | 3 |
| *par-1.1* | 5,34E-33 | 0,529623034 | 0,974 | 0,934 | 6,36E-29 | 3 |
| *Jabba.2* | 7,35E-33 | -1,438586284 | 0,095 | 0,33 | 8,76E-29 | 3 |
| *pug* | 1,28E-32 | 0,806986479 | 0,749 | 0,523 | 1,53E-28 | 3 |
| *LRR.2* | 2,15E-32 | 0,899661151 | 0,699 | 0,48 | 2,57E-28 | 3 |
| *mino.1* | 6,42E-32 | 0,698577164 | 0,836 | 0,635 | 7,65E-28 | 3 |
| *Invadolysin.2* | 1,14E-31 | -2,043104515 | 0,096 | 0,316 | 1,36E-27 | 3 |
| *Pfas.2* | 1,70E-31 | 0,49314027 | 0,797 | 0,509 | 2,03E-27 | 3 |
| *Mob2.2* | 1,93E-31 | 0,765682132 | 0,871 | 0,723 | 2,30E-27 | 3 |
| *Atpalpha.1* | 4,53E-31 | 0,88383341 | 0,842 | 0,71 | 5,40E-27 | 3 |
| *CG5151.2* | 6,62E-31 | -1,01574294 | 0,489 | 0,693 | 7,90E-27 | 3 |
| *Irp-1B.1* | 5,11E-30 | 0,847069424 | 0,458 | 0,237 | 6,09E-26 | 3 |
| *Rel.2* | 5,91E-30 | 0,796071134 | 0,704 | 0,496 | 7,05E-26 | 3 |
| *E(spl)malpha-BFM.* | 1,38E-28 | 0,819556969 | 0,526 | 0,302 | 1,65E-24 | 3 |
| *app.2* | 5,58E-28 | 0,817071942 | 0,564 | 0,347 | 6,66E-24 | 3 |
| *Pect.2* | 5,77E-28 | 0,791977182 | 0,706 | 0,523 | 6,88E-24 | 3 |
| *spin.1* | 3,29E-27 | -0,904210602 | 0,495 | 0,662 | 3,93E-23 | 3 |
| *CG42788.2* | 3,86E-27 | -1,729682792 | 0,326 | 0,52 | 4,60E-23 | 3 |
| *Lst.2* | 5,03E-27 | 0,596486592 | 0,905 | 0,766 | 6,01E-23 | 3 |
| *Myo31DF.2* | 4,07E-26 | -1,742317447 | 0,077 | 0,267 | 4,86E-22 | 3 |
| *galla-1* | 6,94E-26 | 0,877084722 | 0,26 | 0,103 | 8,28E-22 | 3 |
| *CG8468* | 1,35E-25 | 0,709827156 | 0,802 | 0,643 | 1,61E-21 | 3 |
| *CrebA.2* | 3,68E-25 | 0,600618238 | 0,408 | 0,214 | 4,40E-21 | 3 |
| *Hipk.2* | 5,12E-25 | 0,536840811 | 0,825 | 0,653 | 6,11E-21 | 3 |
| *CCHa2.2* | 1,36E-24 | -1,441389986 | 0,109 | 0,3 | 1,62E-20 | 3 |
| *Hsp26.2* | 1,53E-24 | -2,147017937 | 0,291 | 0,453 | 1,83E-20 | 3 |
| *foxo.2* | 2,83E-24 | 0,538630203 | 0,865 | 0,707 | 3,37E-20 | 3 |
| *bmm.2* | 7,07E-24 | 0,54536253 | 0,9 | 0,728 | 8,44E-20 | 3 |
| *CG8079.1* | 2,67E-23 | 0,747729306 | 0,449 | 0,252 | 3,18E-19 | 3 |
| *ACC.2* | 4,61E-23 | 0,383976356 | 0,892 | 0,658 | 5,50E-19 | 3 |
| *CG31678.2* | 1,61E-22 | 0,763961753 | 0,363 | 0,185 | 1,91E-18 | 3 |
| *CG34347.2* | 2,63E-22 | 0,820275118 | 0,529 | 0,341 | 3,13E-18 | 3 |
| *comm2.2* | 5,78E-22 | -1,282486211 | 0,121 | 0,298 | 6,90E-18 | 3 |
| *Gclm.2* | 9,00E-22 | -1,344583948 | 0,225 | 0,395 | 1,07E-17 | 3 |
| *eas.2* | 1,34E-21 | 0,59700938 | 0,624 | 0,432 | 1,59E-17 | 3 |
| *Mal-B2.2* | 3,21E-21 | 0,666757317 | 0,666 | 0,491 | 3,83E-17 | 3 |
| *CG10082.2* | 4,06E-21 | -0,924039592 | 0,487 | 0,633 | 4,85E-17 | 3 |
| *CG8112* | 5,63E-21 | 0,786893803 | 0,289 | 0,136 | 6,71E-17 | 3 |
| *ckn.1* | 7,26E-21 | 0,904496633 | 0,484 | 0,309 | 8,66E-17 | 3 |
| *Stat92E.2* | 1,52E-20 | -1,079868277 | 0,471 | 0,619 | 1,81E-16 | 3 |
| *sug.2* | 1,70E-20 | -1,472239583 | 0,182 | 0,36 | 2,03E-16 | 3 |

| | | | | | | |
|---|---|---|---|---|---|---|
| *Sodh-1.2* | 1,83E-20 | 0,782810823 | 0,437 | 0,261 | 2,18E-16 | 3 |
| *rgn.2* | 1,00E-19 | -1,497260132 | 0,341 | 0,49 | 1,20E-15 | 3 |
| *PRAS40.1* | 1,04E-19 | 0,799329054 | 0,431 | 0,254 | 1,24E-15 | 3 |
| *CG5773.2* | 1,22E-19 | -1,368150641 | 0,101 | 0,263 | 1,46E-15 | 3 |
| *fok* | 1,55E-19 | 0,762553554 | 0,661 | 0,487 | 1,84E-15 | 3 |
| *Pvr.2* | 1,56E-19 | -1,591262692 | 0,322 | 0,465 | 1,86E-15 | 3 |
| *Map205.2* | 2,60E-19 | -0,959740424 | 0,432 | 0,568 | 3,10E-15 | 3 |
| *vri.2* | 6,62E-19 | -0,95302956 | 0,384 | 0,529 | 7,90E-15 | 3 |
| *Gpdh1.2* | 1,18E-18 | -1,261436478 | 0,238 | 0,393 | 1,41E-14 | 3 |
| *zfh1.1* | 1,31E-18 | 0,5539962 | 0,542 | 0,347 | 1,56E-14 | 3 |
| *Culd.2* | 1,54E-18 | -1,104924073 | 0,219 | 0,375 | 1,83E-14 | 3 |
| *MTA1-like.2* | 2,23E-18 | 0,74417684 | 0,563 | 0,402 | 2,66E-14 | 3 |
| *Dys.2* | 2,52E-18 | 0,538714154 | 0,707 | 0,531 | 3,01E-14 | 3 |
| *Got1* | 3,72E-18 | 0,70354726 | 0,349 | 0,195 | 4,44E-14 | 3 |
| *PGRP-LB* | 4,18E-18 | 0,612651506 | 0,304 | 0,152 | 4,98E-14 | 3 |
| *luna.2* | 4,26E-18 | -0,668260775 | 0,643 | 0,738 | 5,08E-14 | 3 |
| *Rtnl1* | 4,96E-18 | -1,098157407 | 0,219 | 0,382 | 5,92E-14 | 3 |
| *LPCAT.2* | 7,57E-18 | 0,602817238 | 0,334 | 0,173 | 9,03E-14 | 3 |
| *Hsp27.2* | 9,61E-18 | -2,34909621 | 0,233 | 0,368 | 1,15E-13 | 3 |
| *red.1* | 1,35E-17 | 0,882844105 | 0,325 | 0,176 | 1,61E-13 | 3 |
| *PAPLA1.2* | 1,40E-17 | 0,69047977 | 0,621 | 0,462 | 1,67E-13 | 3 |
| *Svil.1* | 2,49E-17 | 0,63913945 | 0,672 | 0,518 | 2,97E-13 | 3 |
| *Glut4EF* | 2,73E-17 | 0,555418371 | 0,891 | 0,804 | 3,25E-13 | 3 |
| *CG7130.2* | 4,79E-17 | -1,537986338 | 0,159 | 0,298 | 5,72E-13 | 3 |
| *rudhira.1* | 5,19E-17 | 0,493689195 | 0,807 | 0,653 | 6,19E-13 | 3 |
| *CG33494.2* | 5,40E-17 | -0,806893989 | 0,188 | 0,352 | 6,44E-13 | 3 |
| *CG14478.1* | 7,29E-17 | 0,574923422 | 0,637 | 0,481 | 8,70E-13 | 3 |
| *Hsc70Cb.2* | 1,07E-16 | -1,032154424 | 0,432 | 0,551 | 1,28E-12 | 3 |
| *CG8745.2* | 1,60E-16 | -1,289760618 | 0,241 | 0,398 | 1,90E-12 | 3 |
| *CG10621.2* | 1,62E-16 | 0,694858896 | 0,392 | 0,238 | 1,94E-12 | 3 |
| *CG11537.1* | 1,87E-16 | 0,829305548 | 0,317 | 0,176 | 2,23E-12 | 3 |
| *pst.2* | 3,47E-16 | 0,457512663 | 0,72 | 0,55 | 4,14E-12 | 3 |
| *AspRS.2* | 4,42E-16 | -1,087222059 | 0,114 | 0,256 | 5,28E-12 | 3 |
| *Akap200.2* | 7,21E-16 | -0,893255308 | 0,46 | 0,578 | 8,60E-12 | 3 |
| *LRP1.2* | 1,08E-15 | -0,977484847 | 0,307 | 0,455 | 1,28E-11 | 3 |
| *CG5955.2* | 1,10E-15 | 0,648150108 | 0,317 | 0,175 | 1,31E-11 | 3 |
| *ninaE.1* | 1,15E-15 | 0,733804799 | 0,371 | 0,215 | 1,37E-11 | 3 |
| *eEF2.2* | 1,34E-15 | 0,39675652 | 0,841 | 0,736 | 1,59E-11 | 3 |
| *bnl.2* | 1,56E-15 | 0,648246136 | 0,449 | 0,298 | 1,86E-11 | 3 |
| *Thor.2* | 1,71E-15 | -0,704843175 | 0,423 | 0,565 | 2,04E-11 | 3 |
| *Gp150* | 2,33E-15 | 0,494962394 | 0,653 | 0,492 | 2,78E-11 | 3 |
| *Tab2.2* | 2,36E-15 | 0,562734852 | 0,531 | 0,374 | 2,82E-11 | 3 |
| *CCT3.2* | 2,81E-15 | -1,163257456 | 0,133 | 0,268 | 3,36E-11 | 3 |
| *CG8312.2* | 3,46E-15 | -1,370874659 | 0,217 | 0,354 | 4,13E-11 | 3 |
| *Got2.2* | 4,26E-15 | 0,439839674 | 0,682 | 0,517 | 5,08E-11 | 3 |
| *CG14153* | 6,40E-15 | 0,652600433 | 0,259 | 0,134 | 7,63E-11 | 3 |
| *Hex-C.2* | 7,35E-15 | 0,506011784 | 0,576 | 0,413 | 8,76E-11 | 3 |
| *Fer2LCH.2* | 8,28E-15 | 0,432543939 | 0,846 | 0,72 | 9,88E-11 | 3 |
| *Flo2.2* | 1,10E-14 | 0,689784551 | 0,421 | 0,272 | 1,31E-10 | 3 |
| *stas* | 1,50E-14 | 0,668055898 | 0,41 | 0,265 | 1,79E-10 | 3 |
| *Bacc.2* | 1,55E-14 | -0,60770677 | 0,709 | 0,767 | 1,84E-10 | 3 |
| *RpL41.1* | 1,66E-14 | 0,503734239 | 0,711 | 0,552 | 1,98E-10 | 3 |
| *pcs.2* | 3,31E-14 | 0,750429485 | 0,537 | 0,396 | 3,95E-10 | 3 |
| *kay.2* | 4,37E-14 | -1,108392226 | 0,452 | 0,534 | 5,21E-10 | 3 |
| *RpS2* | 6,38E-14 | 0,498350739 | 0,654 | 0,514 | 7,61E-10 | 3 |
| *Plod.2* | 7,11E-14 | 0,558618504 | 0,402 | 0,254 | 8,48E-10 | 3 |
| *SCaMC.2* | 7,82E-14 | 0,33414662 | 0,865 | 0,741 | 9,32E-10 | 3 |
| *RpL10Ab.1* | 1,57E-13 | 0,428576789 | 0,712 | 0,557 | 1,88E-09 | 3 |
| *Mdh2* | 1,59E-13 | 0,492493162 | 0,659 | 0,512 | 1,90E-09 | 3 |
| *dsx.2* | 1,73E-13 | 0,692052056 | 0,646 | 0,504 | 2,06E-09 | 3 |
| *CG8547* | 1,82E-13 | 0,668298627 | 0,268 | 0,148 | 2,17E-09 | 3 |

| | | | | | | |
|---|---|---|---|---|---|---|
| *RpL18A.2* | 1,85E-13 | 0,423269377 | 0,735 | 0,593 | 2,21E-09 | 3 |
| *bun.2* | 1,98E-13 | -0,913845115 | 0,815 | 0,85 | 2,36E-09 | 3 |
| *AnxB9.2* | 2,18E-13 | -1,490552776 | 0,46 | 0,505 | 2,60E-09 | 3 |
| *Mkp3.1* | 2,53E-13 | -0,751742132 | 0,289 | 0,428 | 3,02E-09 | 3 |
| *Hsp68.2* | 2,93E-13 | -1,758813389 | 0,207 | 0,322 | 3,50E-09 | 3 |
| *Mondo.1* | 3,32E-13 | 0,52673078 | 0,754 | 0,631 | 3,96E-09 | 3 |
| *pHCl-2* | 5,28E-13 | 0,51193016 | 0,265 | 0,143 | 6,30E-09 | 3 |
| *cact.2* | 6,12E-13 | 0,355983959 | 0,775 | 0,608 | 7,30E-09 | 3 |
| *Tsf1.2* | 6,77E-13 | -0,978209058 | 0,188 | 0,32 | 8,07E-09 | 3 |
| *tara.1* | 7,99E-13 | 0,526215782 | 0,503 | 0,357 | 9,53E-09 | 3 |
| *RpL37A.2* | 1,13E-12 | 0,392927919 | 0,621 | 0,454 | 1,35E-08 | 3 |
| *CG32687.1* | 1,27E-12 | 0,428129545 | 0,611 | 0,46 | 1,51E-08 | 3 |
| *CrebB.1* | 1,31E-12 | 0,615004115 | 0,365 | 0,234 | 1,57E-08 | 3 |
| *RpL39.1* | 1,43E-12 | 0,483811685 | 0,603 | 0,458 | 1,70E-08 | 3 |
| *Shmt.2* | 1,57E-12 | 0,346496913 | 0,744 | 0,582 | 1,87E-08 | 3 |
| *CG9005.2* | 1,73E-12 | 0,473123644 | 0,49 | 0,341 | 2,07E-08 | 3 |
| *Paics.1* | 1,94E-12 | 0,324281417 | 0,887 | 0,758 | 2,32E-08 | 3 |
| *Trx-2.2* | 2,12E-12 | -1,128028875 | 0,196 | 0,312 | 2,52E-08 | 3 |
| *krz.1* | 2,16E-12 | 0,51509347 | 0,33 | 0,199 | 2,57E-08 | 3 |
| *CG9674.2* | 2,23E-12 | 0,377535513 | 0,775 | 0,598 | 2,66E-08 | 3 |
| *CRMP* | 2,34E-12 | 0,552959053 | 0,259 | 0,143 | 2,79E-08 | 3 |
| *Khc-73.2* | 2,71E-12 | -0,857063276 | 0,278 | 0,397 | 3,23E-08 | 3 |
| *CG2865.1* | 2,81E-12 | 0,751393113 | 0,436 | 0,309 | 3,35E-08 | 3 |
| *chic.2* | 2,81E-12 | -0,980010788 | 0,333 | 0,439 | 3,35E-08 | 3 |
| *dnc.2* | 3,17E-12 | 0,25593238 | 0,878 | 0,721 | 3,78E-08 | 3 |
| *CG7720.2* | 4,66E-12 | -1,406330041 | 0,262 | 0,379 | 5,56E-08 | 3 |
| *SNF4Agamma.1* | 4,68E-12 | 0,297123511 | 0,942 | 0,879 | 5,58E-08 | 3 |
| *Gbs-76A.2* | 5,02E-12 | -0,63968421 | 0,314 | 0,438 | 5,99E-08 | 3 |
| *Col4a1.1* | 5,50E-12 | 0,354202586 | 0,336 | 0,203 | 6,56E-08 | 3 |
| *eEF1alpha1.1* | 5,60E-12 | 0,274070333 | 0,949 | 0,9 | 6,68E-08 | 3 |
| *blot.1* | 6,09E-12 | 0,383229886 | 0,526 | 0,366 | 7,27E-08 | 3 |
| *Ndae1.2* | 6,36E-12 | -1,08583983 | 0,241 | 0,362 | 7,59E-08 | 3 |
| *nes* | 6,40E-12 | 0,492009015 | 0,272 | 0,152 | 7,63E-08 | 3 |
| *RpS21.2* | 6,41E-12 | 0,398701963 | 0,686 | 0,542 | 7,64E-08 | 3 |
| *IP3K1.2* | 6,95E-12 | 0,394104985 | 0,691 | 0,526 | 8,29E-08 | 3 |
| *CG3246* | 7,34E-12 | 0,557234678 | 0,262 | 0,148 | 8,76E-08 | 3 |
| *Pgm1.2* | 7,94E-12 | 0,477442109 | 0,362 | 0,228 | 9,47E-08 | 3 |
| *Smr.2* | 8,18E-12 | 0,288746293 | 0,844 | 0,725 | 9,76E-08 | 3 |
| *smash* | 1,01E-11 | 0,527056428 | 0,744 | 0,614 | 1,20E-07 | 3 |
| *Parp.2* | 1,02E-11 | 0,339737454 | 0,781 | 0,654 | 1,22E-07 | 3 |
| *CG2233.2* | 1,15E-11 | 0,257586015 | 0,965 | 0,903 | 1,37E-07 | 3 |
| *RpL17.2* | 1,50E-11 | 0,339740673 | 0,77 | 0,653 | 1,80E-07 | 3 |
| *CG31778* | 2,04E-11 | 0,624744829 | 0,302 | 0,184 | 2,43E-07 | 3 |
| *CG16758.1* | 2,31E-11 | 0,322950438 | 0,957 | 0,913 | 2,76E-07 | 3 |
| *CenG1A.2* | 2,81E-11 | 0,571309805 | 0,672 | 0,517 | 3,35E-07 | 3 |
| *sky.1* | 4,60E-11 | -1,116186135 | 0,154 | 0,265 | 5,49E-07 | 3 |
| *Mur2B.2* | 5,99E-11 | 0,445975037 | 0,551 | 0,407 | 7,15E-07 | 3 |
| *Snp* | 6,44E-11 | 0,554939328 | 0,503 | 0,373 | 7,68E-07 | 3 |
| *CG3829.2* | 6,64E-11 | 0,393833798 | 0,436 | 0,297 | 7,92E-07 | 3 |
| *path.2* | 7,27E-11 | 0,281878486 | 0,945 | 0,873 | 8,68E-07 | 3 |
| *Frl.2* | 7,37E-11 | -0,938629721 | 0,466 | 0,525 | 8,80E-07 | 3 |
| *CG14823.2* | 7,91E-11 | -0,936219916 | 0,259 | 0,367 | 9,44E-07 | 3 |
| *CG2765.1* | 9,11E-11 | -0,802407439 | 0,151 | 0,263 | 1,09E-06 | 3 |
| *Hers.2* | 9,32E-11 | 0,37979004 | 0,723 | 0,598 | 1,11E-06 | 3 |
| *eIF4G1* | 1,01E-10 | 0,419099881 | 0,688 | 0,561 | 1,21E-06 | 3 |
| *RpS26* | 1,22E-10 | 0,399363519 | 0,656 | 0,518 | 1,45E-06 | 3 |
| *bbg.1* | 1,26E-10 | -0,620382413 | 0,233 | 0,364 | 1,50E-06 | 3 |
| *Syx1A.2* | 1,27E-10 | -0,648895852 | 0,476 | 0,566 | 1,52E-06 | 3 |
| *gish* | 1,51E-10 | 0,337996843 | 0,809 | 0,709 | 1,80E-06 | 3 |
| *Myc.1* | 1,59E-10 | 0,518505869 | 0,746 | 0,637 | 1,89E-06 | 3 |
| *RpL9.2* | 1,88E-10 | 0,324190068 | 0,651 | 0,525 | 2,24E-06 | 3 |

| | | | | | | |
|---|---|---|---|---|---|---|
| *Cdep.2* | 1,88E-10 | 0,415130983 | 0,474 | 0,34 | 2,24E-06 | 3 |
| *RpLP2.1* | 2,05E-10 | 0,32279153 | 0,759 | 0,629 | 2,45E-06 | 3 |
| *CG6175* | 2,07E-10 | 0,634677583 | 0,268 | 0,162 | 2,47E-06 | 3 |
| *PH4alphaEFB* | 2,09E-10 | 0,425104072 | 0,264 | 0,155 | 2,49E-06 | 3 |
| *CG4629.2* | 2,35E-10 | -0,949091098 | 0,167 | 0,274 | 2,80E-06 | 3 |
| *Sxl* | 2,39E-10 | 0,374543046 | 0,863 | 0,786 | 2,86E-06 | 3 |
| *Dmtn.2* | 2,70E-10 | -0,447607173 | 0,691 | 0,763 | 3,23E-06 | 3 |
| *PCB.2* | 2,74E-10 | 0,300984917 | 0,725 | 0,584 | 3,26E-06 | 3 |
| *cbt.2* | 2,76E-10 | -0,687370938 | 0,183 | 0,301 | 3,29E-06 | 3 |
| *sbb.2* | 3,40E-10 | 0,382256462 | 0,685 | 0,565 | 4,06E-06 | 3 |
| *RhoGAP15B.2* | 3,48E-10 | -0,674316418 | 0,14 | 0,25 | 4,16E-06 | 3 |
| *flfl.2* | 4,04E-10 | 0,419984416 | 0,404 | 0,274 | 4,82E-06 | 3 |
| *ced-6.2* | 4,14E-10 | -0,546911505 | 0,387 | 0,5 | 4,94E-06 | 3 |
| *RpL27.2* | 4,43E-10 | 0,284163369 | 0,728 | 0,58 | 5,28E-06 | 3 |
| *Cdc7* | 4,77E-10 | 0,393262133 | 0,265 | 0,156 | 5,69E-06 | 3 |
| *Caper.2* | 5,64E-10 | -0,508897102 | 0,63 | 0,681 | 6,72E-06 | 3 |
| *CG8485.2* | 5,86E-10 | -0,889339916 | 0,164 | 0,267 | 6,99E-06 | 3 |
| *CG32549.1* | 5,89E-10 | 0,384975032 | 0,775 | 0,653 | 7,03E-06 | 3 |
| *RpL40.1* | 6,70E-10 | 0,355086149 | 0,574 | 0,442 | 7,99E-06 | 3 |
| *IM4.2* | 6,75E-10 | -1,035088925 | 0,214 | 0,33 | 8,05E-06 | 3 |
| *Cyp309a1.2* | 7,07E-10 | 0,382144973 | 0,338 | 0,214 | 8,43E-06 | 3 |
| *Gs2.1* | 7,80E-10 | 0,719612762 | 0,418 | 0,305 | 9,30E-06 | 3 |
| *AkhR.2* | 8,56E-10 | -0,760979295 | 0,233 | 0,341 | 1,02E-05 | 3 |
| *B4.1* | 9,30E-10 | 0,411575966 | 0,574 | 0,462 | 1,11E-05 | 3 |
| *mim.2* | 9,51E-10 | 0,41471447 | 0,49 | 0,362 | 1,13E-05 | 3 |
| *LamC.2* | 1,00E-09 | -0,771345428 | 0,18 | 0,287 | 1,20E-05 | 3 |
| *Mst84Da.2* | 1,06E-09 | 0,402349799 | 0,42 | 0,296 | 1,27E-05 | 3 |
| *CG2201* | 1,45E-09 | 0,437037027 | 0,46 | 0,335 | 1,73E-05 | 3 |
| *Eip63E.1* | 1,56E-09 | -0,490999381 | 0,572 | 0,653 | 1,87E-05 | 3 |
| *PMCA.2* | 1,71E-09 | 0,359712895 | 0,675 | 0,559 | 2,04E-05 | 3 |
| *Lk6.2* | 1,78E-09 | 0,298734322 | 0,941 | 0,885 | 2,12E-05 | 3 |
| *conu.2* | 2,02E-09 | 0,33191072 | 0,611 | 0,478 | 2,41E-05 | 3 |
| *Hsp83.2* | 2,03E-09 | -0,893695151 | 0,777 | 0,724 | 2,42E-05 | 3 |
| *CG12795.2* | 2,06E-09 | -1,253350784 | 0,291 | 0,364 | 2,46E-05 | 3 |
| *Gs1.1* | 2,44E-09 | 0,41205431 | 0,383 | 0,263 | 2,92E-05 | 3 |
| *Plc21C.1* | 2,46E-09 | 0,346457268 | 0,672 | 0,548 | 2,93E-05 | 3 |
| *CG43658* | 2,47E-09 | 0,326884523 | 0,765 | 0,653 | 2,94E-05 | 3 |
| *CG6115.2* | 2,63E-09 | 0,33731485 | 0,547 | 0,4 | 3,14E-05 | 3 |
| *RpS30.2* | 3,27E-09 | 0,332112041 | 0,629 | 0,499 | 3,90E-05 | 3 |
| *fray* | 3,34E-09 | -0,663315383 | 0,182 | 0,287 | 3,98E-05 | 3 |
| *Kr-h1.1* | 3,63E-09 | 0,449877983 | 0,566 | 0,446 | 4,33E-05 | 3 |
| *Abl.2* | 4,43E-09 | -0,974221102 | 0,31 | 0,387 | 5,29E-05 | 3 |
| *vir-1.2* | 4,50E-09 | -0,651466339 | 0,743 | 0,737 | 5,37E-05 | 3 |
| *RpL13A.2* | 6,27E-09 | 0,296065238 | 0,746 | 0,625 | 7,48E-05 | 3 |
| *RpS15.2* | 6,46E-09 | 0,309289378 | 0,77 | 0,658 | 7,70E-05 | 3 |
| *fz2.1* | 6,70E-09 | -0,881338169 | 0,199 | 0,301 | 7,99E-05 | 3 |
| *RpLP1.2* | 6,75E-09 | 0,339921253 | 0,72 | 0,608 | 8,05E-05 | 3 |
| *RpL29.2* | 7,31E-09 | 0,403257396 | 0,524 | 0,402 | 8,72E-05 | 3 |
| *elB.2* | 7,35E-09 | -0,824597618 | 0,233 | 0,331 | 8,77E-05 | 3 |
| *kst.2* | 7,39E-09 | -0,829951692 | 0,207 | 0,307 | 8,81E-05 | 3 |
| *gce.2* | 7,46E-09 | -0,676604746 | 0,413 | 0,5 | 8,90E-05 | 3 |
| *CG1943.2* | 7,65E-09 | 0,423889537 | 0,297 | 0,191 | 9,12E-05 | 3 |
| *Gale.2* | 9,10E-09 | 0,560851248 | 0,32 | 0,218 | 0,000108549 | 3 |
| *Ntan1.2* | 1,15E-08 | -0,620995243 | 0,559 | 0,617 | 0,000137032 | 3 |
| *RpS18.1* | 1,24E-08 | 0,25406737 | 0,809 | 0,701 | 0,000148391 | 3 |
| *Eip74EF* | 1,29E-08 | 0,349969133 | 0,723 | 0,603 | 0,000153501 | 3 |
| *CG11791.2* | 1,54E-08 | -0,738535371 | 0,368 | 0,449 | 0,000183248 | 3 |
| *RpS4.2* | 1,95E-08 | 0,272093818 | 0,683 | 0,549 | 0,000232985 | 3 |
| *CG3597.2* | 2,05E-08 | 0,374249015 | 0,333 | 0,223 | 0,000244707 | 3 |
| *CG34417.1* | 2,27E-08 | 0,400487805 | 0,719 | 0,59 | 0,000270668 | 3 |
| *Diap1.2* | 2,31E-08 | -0,374999581 | 0,717 | 0,756 | 0,0002759 | 3 |

| | | | | | | |
|---|---|---|---|---|---|---|
| *Gart.2* | 2,43E-08 | 0,26551907 | 0,772 | 0,64 | 0,000289785 | 3 |
| *Blimp-1.2* | 2,51E-08 | -0,428305293 | 0,651 | 0,71 | 0,000299994 | 3 |
| *DnaJ-1.2* | 3,05E-08 | -0,713889425 | 0,502 | 0,542 | 0,00036361 | 3 |
| *Socs36E.2* | 3,08E-08 | -0,906742328 | 0,251 | 0,341 | 0,00036757 | 3 |
| *BomT3.2* | 3,33E-08 | -1,003043354 | 0,354 | 0,445 | 0,000397736 | 3 |
| *Shrm* | 4,01E-08 | -0,591872054 | 0,241 | 0,346 | 0,000477878 | 3 |
| *Idh.2* | 4,07E-08 | -0,649938686 | 0,342 | 0,438 | 0,000484928 | 3 |
| *Mad.2* | 4,14E-08 | -0,611458379 | 0,252 | 0,347 | 0,000493666 | 3 |
| *CG31694.2* | 4,32E-08 | -0,789975353 | 0,312 | 0,392 | 0,000515415 | 3 |
| *vig* | 4,32E-08 | 0,355186688 | 0,27 | 0,172 | 0,000515429 | 3 |
| *CG7945.2* | 4,44E-08 | -0,933807007 | 0,167 | 0,255 | 0,000529427 | 3 |
| *eIF3c.2* | 5,68E-08 | 0,328939159 | 0,304 | 0,2 | 0,000677425 | 3 |
| *CG43394* | 6,55E-08 | 0,419992083 | 0,273 | 0,178 | 0,000781352 | 3 |
| *MRP.1* | 7,68E-08 | -1,166643718 | 0,373 | 0,441 | 0,00091641 | 3 |
| *CG8034.2* | 8,92E-08 | -0,852215826 | 0,568 | 0,614 | 0,001063578 | 3 |
| *Nak.1* | 9,93E-08 | 0,375280522 | 0,31 | 0,21 | 0,001185095 | 3 |
| *Stim* | 1,06E-07 | 0,384817821 | 0,272 | 0,18 | 0,001259546 | 3 |
| *CG13868.2* | 1,15E-07 | -0,304204929 | 0,696 | 0,759 | 0,001377487 | 3 |
| *RpL7A.2* | 1,25E-07 | 0,30264764 | 0,712 | 0,602 | 0,001488577 | 3 |
| *wnd.1* | 1,27E-07 | 0,440295083 | 0,42 | 0,31 | 0,001519191 | 3 |
| *KrT95D.2* | 1,35E-07 | -0,500715244 | 0,65 | 0,693 | 0,001606914 | 3 |
| *RpL38.2* | 1,38E-07 | 0,347905341 | 0,479 | 0,367 | 0,001652105 | 3 |
| *retm.2* | 1,47E-07 | -0,663169626 | 0,214 | 0,304 | 0,001748563 | 3 |
| *CG10433.1* | 1,53E-07 | -0,423289242 | 0,637 | 0,686 | 0,001827614 | 3 |
| *eEF5.2* | 1,59E-07 | 0,255066572 | 0,643 | 0,517 | 0,001895955 | 3 |
| *Cbs.2* | 1,66E-07 | 0,383638855 | 0,338 | 0,239 | 0,001977279 | 3 |
| *sfl.1* | 1,66E-07 | 0,315836468 | 0,606 | 0,491 | 0,001985436 | 3 |
| *CG3638.2* | 1,67E-07 | 0,259175699 | 0,574 | 0,453 | 0,001991276 | 3 |
| *CG4538.2* | 1,67E-07 | 0,349820511 | 0,375 | 0,266 | 0,00199555 | 3 |
| *RpL23A* | 1,79E-07 | 0,317864713 | 0,559 | 0,45 | 0,00213604 | 3 |
| *goe.2* | 2,02E-07 | 0,312761616 | 0,359 | 0,25 | 0,002405836 | 3 |
| *RpS29.2* | 2,05E-07 | 0,282467639 | 0,669 | 0,551 | 0,002447105 | 3 |
| *ftz-f1.2* | 2,19E-07 | 0,280200471 | 0,707 | 0,593 | 0,002615324 | 3 |
| *Swip-1.2* | 2,29E-07 | -0,509792753 | 0,323 | 0,42 | 0,00273692 | 3 |
| *Nop17l.2* | 2,30E-07 | -0,683521465 | 0,35 | 0,43 | 0,002739065 | 3 |
| *Tret1-1.2* | 2,42E-07 | -0,645994016 | 0,508 | 0,575 | 0,002890045 | 3 |
| *Pdha.2* | 2,55E-07 | -0,618673764 | 0,166 | 0,254 | 0,003044483 | 3 |
| *Cyp28d1.2* | 2,65E-07 | 0,420241159 | 0,293 | 0,2 | 0,003156865 | 3 |
| *CG6330.2* | 2,87E-07 | 0,413070071 | 0,375 | 0,276 | 0,003419475 | 3 |
| *Lsd-1.2* | 3,14E-07 | -0,776899263 | 0,236 | 0,326 | 0,003742725 | 3 |
| *RpS3* | 3,46E-07 | 0,270315082 | 0,587 | 0,469 | 0,004129136 | 3 |
| *RpL23.2* | 3,84E-07 | 0,282334328 | 0,773 | 0,673 | 0,00457577 | 3 |
| *Pfrx.2* | 4,45E-07 | 0,321119066 | 0,379 | 0,276 | 0,005304133 | 3 |
| *E2f1* | 4,57E-07 | 0,388078795 | 0,526 | 0,416 | 0,005448767 | 3 |
| *rhea.2* | 4,57E-07 | -1,146970486 | 0,482 | 0,497 | 0,005456989 | 3 |
| *RpL11.2* | 4,95E-07 | 0,250700068 | 0,608 | 0,489 | 0,00590013 | 3 |
| *Pcyt1.2* | 5,66E-07 | 0,370135015 | 0,572 | 0,458 | 0,006747201 | 3 |
| *tsr.2* | 5,96E-07 | -0,59020513 | 0,182 | 0,267 | 0,007114058 | 3 |
| *drongo.2* | 6,32E-07 | -0,470063631 | 0,518 | 0,596 | 0,007538503 | 3 |
| *DOR.2* | 7,22E-07 | -0,465663372 | 0,548 | 0,623 | 0,008614244 | 3 |
| *RpS24.2* | 7,31E-07 | 0,274424079 | 0,492 | 0,378 | 0,008714875 | 3 |
| *RpL3.2* | 7,67E-07 | 0,279686239 | 0,728 | 0,622 | 0,009151789 | 3 |
| *Dr.1* | 7,87E-07 | 0,403359675 | 0,317 | 0,218 | 0,00939266 | 3 |
| *IM33.2* | 8,34E-07 | -0,708365098 | 0,304 | 0,384 | 0,009946352 | 3 |
| *CG5958.2* | 8,70E-07 | -0,687479718 | 0,352 | 0,428 | 0,010383743 | 3 |
| *CG5910.2* | 9,13E-07 | 0,344313225 | 0,489 | 0,376 | 0,010895517 | 3 |
| *lp259.1* | 9,33E-07 | 0,310993367 | 0,314 | 0,221 | 0,011132508 | 3 |
| *MSBP.2* | 9,79E-07 | 0,314127134 | 0,365 | 0,266 | 0,011675411 | 3 |
| *yuri.2* | 9,81E-07 | 0,378094057 | 0,389 | 0,288 | 0,011698191 | 3 |
| *Meltrin* | 9,91E-07 | 0,370311014 | 0,341 | 0,245 | 0,011826104 | 3 |
| *Dbp80.1* | 1,22E-06 | 0,278819829 | 0,748 | 0,635 | 0,01459467 | 3 |

| | | | | | | |
|---|---|---|---|---|---|---|
| *Pmp70.2* | 1,28E-06 | -0,615918498 | 0,203 | 0,288 | 0,015210536 | 3 |
| *fry.1* | 1,28E-06 | 0,334526804 | 0,291 | 0,199 | 0,015220481 | 3 |
| *RpL34b.2* | 1,35E-06 | 0,280808687 | 0,625 | 0,515 | 0,016109706 | 3 |
| *Gdi.1* | 1,40E-06 | 0,359613006 | 0,394 | 0,296 | 0,016696215 | 3 |
| *CtBP.1* | 1,55E-06 | 0,2714354 | 0,691 | 0,588 | 0,018435125 | 3 |
| *Nplp2.2* | 1,55E-06 | -0,640889602 | 0,664 | 0,669 | 0,018441821 | 3 |
| *Doa.2* | 1,60E-06 | 0,29269115 | 0,609 | 0,499 | 0,019045804 | 3 |
| *Hsc70-4.2* | 1,64E-06 | -0,554791316 | 0,768 | 0,733 | 0,019619349 | 3 |
| *CG17108.2* | 1,88E-06 | -1,032435399 | 0,344 | 0,415 | 0,022439785 | 3 |
| *RpS14a* | 1,89E-06 | 0,268853245 | 0,468 | 0,367 | 0,022542806 | 3 |
| *oys.2* | 2,11E-06 | -0,843917201 | 0,285 | 0,344 | 0,025190898 | 3 |
| *Kdm4B* | 2,19E-06 | 0,326486805 | 0,347 | 0,25 | 0,026140643 | 3 |
| *Lpin.2* | 3,11E-06 | 0,316791682 | 0,564 | 0,461 | 0,037157972 | 3 |
| *CG31183.1* | 3,56E-06 | 0,255918717 | 0,646 | 0,541 | 0,042461391 | 3 |
| *Crtc.1* | 3,68E-06 | 0,34760451 | 0,588 | 0,485 | 0,043904355 | 3 |
| *RpS20.2* | 3,79E-06 | 0,258712911 | 0,64 | 0,548 | 0,04526162 | 3 |
| *Piezo.2* | 4,22E-06 | -0,71750913 | 0,357 | 0,417 | 0,05030233 | 3 |
| *for.1* | 4,30E-06 | -0,364802626 | 0,707 | 0,744 | 0,051301044 | 3 |
| *RpS6.2* | 4,70E-06 | 0,286625538 | 0,571 | 0,473 | 0,056047691 | 3 |
| *SERCA.1* | 5,73E-06 | 0,289225965 | 0,381 | 0,284 | 0,06833516 | 3 |
| *CAP* | 6,01E-06 | -0,482872806 | 0,249 | 0,329 | 0,071669473 | 3 |
| *CG11400.2* | 6,05E-06 | -0,430003228 | 0,32 | 0,402 | 0,072154836 | 3 |
| *Ahcy.2* | 6,17E-06 | 0,278764125 | 0,37 | 0,275 | 0,073652997 | 3 |
| *CG11899* | 6,32E-06 | 0,375360113 | 0,378 | 0,287 | 0,075413832 | 3 |
| *Mmp2.1* | 6,56E-06 | 0,333134093 | 0,281 | 0,198 | 0,078242334 | 3 |
| *vsg.1* | 6,98E-06 | 0,374348805 | 0,367 | 0,28 | 0,083281595 | 3 |
| *Indy* | 8,24E-06 | 0,321500859 | 0,695 | 0,621 | 0,098344094 | 3 |
| *NAT1.2* | 9,23E-06 | -0,305243571 | 0,664 | 0,703 | 0,110157309 | 3 |
| *CG9008* | 9,34E-06 | 0,259719775 | 0,27 | 0,188 | 0,111410435 | 3 |
| *Rack1* | 9,72E-06 | 0,275252606 | 0,569 | 0,46 | 0,115984869 | 3 |
| *CNBP.1* | 9,79E-06 | 0,276797699 | 0,325 | 0,236 | 0,116740639 | 3 |
| *Npl4.2* | 1,27E-05 | 0,298606572 | 0,281 | 0,199 | 0,15173172 | 3 |
| *kmr.1* | 1,27E-05 | -0,492684229 | 0,177 | 0,253 | 0,151953112 | 3 |
| *Rip11* | 1,34E-05 | 0,26065027 | 0,37 | 0,282 | 0,159680181 | 3 |
| *Hnf4.2* | 1,41E-05 | -0,312574402 | 0,593 | 0,657 | 0,16794343 | 3 |
| *CG12699* | 1,47E-05 | 0,395609291 | 0,272 | 0,194 | 0,175948315 | 3 |
| *Pak3.2* | 1,59E-05 | -0,536577805 | 0,232 | 0,303 | 0,189488644 | 3 |
| *Atf6.1* | 1,74E-05 | 0,29813024 | 0,654 | 0,581 | 0,207308922 | 3 |
| *CG17278.2* | 1,88E-05 | 0,366850906 | 0,362 | 0,274 | 0,223869263 | 3 |
| *CG15771.1* | 2,00E-05 | 0,284997135 | 0,363 | 0,272 | 0,238948867 | 3 |
| *Atg17.1* | 2,00E-05 | -0,367343298 | 0,424 | 0,498 | 0,238970361 | 3 |
| *Nipped-B.1* | 2,33E-05 | 0,266155465 | 0,801 | 0,738 | 0,278032258 | 3 |
| *Eip75B* | 2,34E-05 | -0,270481072 | 0,846 | 0,851 | 0,279639466 | 3 |
| *olf186-M.1* | 2,39E-05 | -0,432308737 | 0,341 | 0,425 | 0,284759957 | 3 |
| *mys.2* | 2,46E-05 | -0,438030888 | 0,336 | 0,406 | 0,293888406 | 3 |
| *cindr* | 2,52E-05 | -0,397283802 | 0,468 | 0,544 | 0,300869573 | 3 |
| *sm.2* | 2,72E-05 | 0,323452426 | 0,373 | 0,286 | 0,324573924 | 3 |
| *Atet* | 2,85E-05 | 0,343431079 | 0,264 | 0,188 | 0,340408922 | 3 |
| *drpr.2* | 2,91E-05 | -0,616049349 | 0,42 | 0,466 | 0,346796837 | 3 |
| *Pino* | 3,13E-05 | 0,250707294 | 0,82 | 0,754 | 0,37278347 | 3 |
| *mnb.1* | 3,13E-05 | 0,354460885 | 0,474 | 0,39 | 0,37335681 | 3 |
| *CG42524.2* | 3,38E-05 | -0,461342273 | 0,418 | 0,484 | 0,403670939 | 3 |
| *RpS15Aa.2* | 3,44E-05 | 0,266964264 | 0,561 | 0,469 | 0,410347913 | 3 |
| *ttk.1* | 3,57E-05 | -0,531424474 | 0,196 | 0,266 | 0,425346389 | 3 |
| *RpL24-like.1* | 3,57E-05 | 0,334821237 | 0,251 | 0,178 | 0,42628977 | 3 |
| *CG30069.1* | 3,63E-05 | -0,511464265 | 0,275 | 0,342 | 0,433098543 | 3 |
| *RhoGAP93B* | 4,41E-05 | 0,308801849 | 0,299 | 0,221 | 0,526316751 | 3 |
| *Scsalpha1* | 4,46E-05 | 0,264705672 | 0,466 | 0,376 | 0,531516105 | 3 |
| *Stlk.2* | 4,66E-05 | 0,264252489 | 0,365 | 0,278 | 0,555930234 | 3 |
| *CG42668.2* | 5,10E-05 | 0,331424751 | 0,654 | 0,585 | 0,608036047 | 3 |
| *Pkn.2* | 5,19E-05 | -0,331358848 | 0,551 | 0,585 | 0,619654856 | 3 |

| | | | | | | |
|---|---|---|---|---|---|---|
| *CG9451.2* | 6,48E-05 | 0,425639896 | 0,289 | 0,216 | 0,773064266 | 3 |
| *CG8108.1* | 6,54E-05 | -0,37267065 | 0,326 | 0,398 | 0,780710228 | 3 |
| *AGO3.1* | 6,87E-05 | 0,315289936 | 0,296 | 0,222 | 0,819860863 | 3 |
| *Paip2.1* | 7,21E-05 | -0,494939886 | 0,32 | 0,386 | 0,86026112 | 3 |
| *SPARC.2* | 7,45E-05 | 0,295924341 | 0,367 | 0,292 | 0,888705191 | 3 |
| *CG32264* | 7,56E-05 | 0,294419677 | 0,635 | 0,546 | 0,901538507 | 3 |
| *CG7920.2* | 8,50E-05 | -0,442159941 | 0,24 | 0,31 | 1 | 3 |
| *PRL-1.2* | 8,92E-05 | -0,485766121 | 0,249 | 0,312 | 1 | 3 |
| *mbc.1* | 8,94E-05 | -0,457331599 | 0,209 | 0,274 | 1 | 3 |
| *Pka-R1* | 0,000115085 | 0,270980703 | 0,344 | 0,263 | 1 | 3 |
| *Irc.2* | 0,000118415 | -0,458739506 | 0,495 | 0,52 | 1 | 3 |
| *babo* | 0,00011902 | 0,277151155 | 0,362 | 0,281 | 1 | 3 |
| *CG15293.2* | 0,000125327 | -0,401348264 | 0,399 | 0,458 | 1 | 3 |
| *Haspin* | 0,000125575 | 0,345576915 | 0,288 | 0,216 | 1 | 3 |
| *Rho1* | 0,000127317 | -0,292300304 | 0,551 | 0,589 | 1 | 3 |
| *Rab11.2* | 0,000142359 | -0,412320041 | 0,322 | 0,377 | 1 | 3 |
| *Chchd2.2* | 0,000146932 | -0,594159503 | 0,638 | 0,613 | 1 | 3 |
| *nuf.2* | 0,000151984 | -0,326238554 | 0,523 | 0,555 | 1 | 3 |
| *Sap47.1* | 0,000161636 | 0,341993034 | 0,423 | 0,348 | 1 | 3 |
| *bip2.2* | 0,000163964 | 0,280575391 | 0,531 | 0,456 | 1 | 3 |
| *CG12116.2* | 0,000164515 | -0,38128275 | 0,363 | 0,425 | 1 | 3 |
| *fon.2* | 0,000168074 | 0,284767412 | 0,373 | 0,298 | 1 | 3 |
| *rdgA.1* | 0,000206863 | 0,286385792 | 0,294 | 0,223 | 1 | 3 |
| *shep.2* | 0,000207649 | -0,35566265 | 0,744 | 0,75 | 1 | 3 |
| *CG32369.2* | 0,000212496 | -0,428065521 | 0,672 | 0,673 | 1 | 3 |
| *CG6966.2* | 0,000218891 | -0,429537086 | 0,399 | 0,46 | 1 | 3 |
| *Naprt.2* | 0,000221527 | -0,413234797 | 0,314 | 0,377 | 1 | 3 |
| *CG9331.1* | 0,000239857 | 0,38858692 | 0,359 | 0,287 | 1 | 3 |
| *Tep2.1* | 0,000243382 | -0,47268097 | 0,241 | 0,305 | 1 | 3 |
| *Lis-1.2* | 0,000267061 | -0,405271531 | 0,315 | 0,377 | 1 | 3 |
| *Dp1.1* | 0,000298432 | 0,250605106 | 0,301 | 0,23 | 1 | 3 |
| *Cys.2* | 0,000306256 | -0,467219213 | 0,246 | 0,308 | 1 | 3 |
| *Hsp60A.2* | 0,000372396 | -0,386116428 | 0,193 | 0,254 | 1 | 3 |
| *Atg4a.1* | 0,000383362 | 0,263135007 | 0,26 | 0,196 | 1 | 3 |
| *CG6051.2* | 0,000383862 | 0,294305051 | 0,616 | 0,528 | 1 | 3 |
| *puc.2* | 0,000386412 | -0,540184633 | 0,728 | 0,685 | 1 | 3 |
| *CG5059* | 0,000393733 | -0,291238691 | 0,675 | 0,699 | 1 | 3 |
| *CG31635.2* | 0,000418217 | -0,374145382 | 0,513 | 0,561 | 1 | 3 |
| *if.2* | 0,000419525 | -0,607065395 | 0,307 | 0,358 | 1 | 3 |
| *Trp1.2* | 0,000423133 | 0,284682868 | 0,289 | 0,222 | 1 | 3 |
| *puml.2* | 0,000443398 | 0,289840363 | 0,264 | 0,199 | 1 | 3 |
| *Pep* | 0,000479016 | -0,300810981 | 0,286 | 0,35 | 1 | 3 |
| *ImpL2.2* | 0,000492578 | -0,642767718 | 0,206 | 0,258 | 1 | 3 |
| *UbcE2H.2* | 0,000512701 | 0,29525468 | 0,416 | 0,343 | 1 | 3 |
| *sty* | 0,000530134 | -0,55976462 | 0,201 | 0,253 | 1 | 3 |
| *Moe* | 0,000553314 | -0,361692726 | 0,275 | 0,336 | 1 | 3 |
| *swm.2* | 0,000597014 | -0,483943063 | 0,289 | 0,338 | 1 | 3 |
| *Acbp2.2* | 0,000626775 | -0,354055455 | 0,259 | 0,327 | 1 | 3 |
| *RhoGAP18B* | 0,000637516 | -0,254703376 | 0,469 | 0,518 | 1 | 3 |
| *CG1648.2* | 0,000685509 | -0,41730703 | 0,58 | 0,618 | 1 | 3 |
| *Ubi-p63E.2* | 0,000711954 | -0,672039172 | 0,654 | 0,602 | 1 | 3 |
| *Pde6.2* | 0,000714899 | -0,362401228 | 0,486 | 0,518 | 1 | 3 |
| *Oatp30B.2* | 0,00082144 | -0,336005395 | 0,561 | 0,593 | 1 | 3 |
| *AdamTS-A.2* | 0,000830998 | 0,289822577 | 0,323 | 0,258 | 1 | 3 |
| *fwd* | 0,000841593 | -0,35105349 | 0,309 | 0,36 | 1 | 3 |
| *mamo.2* | 0,00086041 | -0,48988197 | 0,54 | 0,559 | 1 | 3 |
| *teq.2* | 0,00086223 | -0,361648289 | 0,206 | 0,261 | 1 | 3 |
| *CG3164.2* | 0,000862626 | -0,32430544 | 0,482 | 0,529 | 1 | 3 |
| *fbp.2* | 0,001027715 | -0,346213131 | 0,272 | 0,328 | 1 | 3 |
| *CLIP-190.1* | 0,001077997 | 0,265796806 | 0,719 | 0,664 | 1 | 3 |
| *Gug.2* | 0,001119297 | -0,320863797 | 0,629 | 0,646 | 1 | 3 |

| | | | | | | |
|---|---|---|---|---|---|---|
| *Act5C.2* | 0,001130462 | -0,580626623 | 0,556 | 0,544 | 1 | 3 |
| *glob1.1* | 0,001172469 | 0,382048575 | 0,78 | 0,732 | 1 | 3 |
| *l(1)G0289.2* | 0,001204219 | 0,287049594 | 0,523 | 0,461 | 1 | 3 |
| *flw.1* | 0,001278246 | 0,297443535 | 0,474 | 0,407 | 1 | 3 |
| *trx.2* | 0,001395495 | -0,295056926 | 0,471 | 0,507 | 1 | 3 |
| *RapGAP1.2* | 0,001398666 | -0,366646799 | 0,207 | 0,264 | 1 | 3 |
| *CG15099.2* | 0,001739074 | -0,457147764 | 0,262 | 0,306 | 1 | 3 |
| *Trxr-1.2* | 0,001857395 | -0,508014349 | 0,371 | 0,396 | 1 | 3 |
| *Nmda1.2* | 0,002020728 | -0,469052518 | 0,326 | 0,358 | 1 | 3 |
| *CG4716.2* | 0,002209979 | -0,303265986 | 0,899 | 0,847 | 1 | 3 |
| *cic.2* | 0,002471203 | -0,400175007 | 0,67 | 0,663 | 1 | 3 |
| *csw.1* | 0,002478593 | 0,269134536 | 0,524 | 0,451 | 1 | 3 |
| *Itpr.2* | 0,002485492 | -0,314186509 | 0,281 | 0,333 | 1 | 3 |
| *CG32767.2* | 0,002514094 | -0,365961502 | 0,354 | 0,395 | 1 | 3 |
| *CG7220.2* | 0,002617841 | 0,250533229 | 0,275 | 0,218 | 1 | 3 |
| *milt.2* | 0,002970653 | -0,30787491 | 0,383 | 0,428 | 1 | 3 |
| *Rad23.2* | 0,002986069 | -0,398817871 | 0,304 | 0,339 | 1 | 3 |
| *CG7029* | 0,003329642 | -0,337931407 | 0,36 | 0,4 | 1 | 3 |
| *Gdh.2* | 0,003836758 | 0,349571046 | 0,437 | 0,372 | 1 | 3 |
| *CG44325.2* | 0,004096855 | -0,369378456 | 0,28 | 0,327 | 1 | 3 |
| *Eglp4.2* | 0,004122478 | -0,466701257 | 0,338 | 0,378 | 1 | 3 |
| *PKD.2* | 0,004190932 | -0,279815404 | 0,248 | 0,301 | 1 | 3 |
| *mbf1.2* | 0,004234666 | -0,678984679 | 0,338 | 0,353 | 1 | 3 |
| *GstE12.2* | 0,00558921 | -0,277521397 | 0,214 | 0,263 | 1 | 3 |
| *Yeti.2* | 0,005619906 | -0,32779105 | 0,617 | 0,614 | 1 | 3 |
| *RasGAP1.2* | 0,005831794 | -0,403896011 | 0,244 | 0,286 | 1 | 3 |
| *CG15096.2* | 0,007473116 | -0,289965439 | 0,217 | 0,261 | 1 | 3 |
| *spoon.2* | 0,00802478 | -0,356159069 | 0,412 | 0,435 | 1 | 3 |
| *betaTub97EF* | 0,008119502 | -0,325535128 | 0,248 | 0,289 | 1 | 3 |
| *Cip4* | 0,008283555 | -0,292461161 | 0,233 | 0,277 | 1 | 3 |
| *CG10680.2* | 0,008419848 | 0,276429425 | 0,465 | 0,406 | 1 | 3 |
| *CCT2.2* | 0,008788581 | -0,475292345 | 0,23 | 0,263 | 1 | 3 |
| *RhoGAP19D.2* | 0,009393814 | 0,375643573 | 0,523 | 0,472 | 1 | 3 |
| *osp.2* | 0,009424185 | -0,337531891 | 0,28 | 0,328 | 1 | 3 |
| *MRP.2* | 3,81E-160 | 3,007647843 | 0,966 | 0,373 | 4,54E-156 | 4 |
| *CCT3.3* | 4,21E-151 | 2,206254614 | 0,824 | 0,183 | 5,02E-147 | 4 |
| *AnxB9.3* | 2,41E-136 | 2,282605913 | 0,99 | 0,446 | 2,88E-132 | 4 |
| *Trx-2.3* | 5,54E-133 | 2,341622865 | 0,824 | 0,235 | 6,60E-129 | 4 |
| *Myo31DF.3* | 3,23E-130 | 2,270284148 | 0,759 | 0,176 | 3,85E-126 | 4 |
| *Hsp27.3* | 2,72E-127 | 2,252343802 | 0,903 | 0,284 | 3,24E-123 | 4 |
| *stv.2* | 3,66E-126 | 2,168763275 | 0,993 | 0,581 | 4,36E-122 | 4 |
| *kay.3* | 3,42E-123 | 2,107569546 | 0,966 | 0,472 | 4,08E-119 | 4 |
| *elB.3* | 2,44E-121 | 2,16728648 | 0,831 | 0,259 | 2,91E-117 | 4 |
| *CG8312.3* | 1,49E-119 | 2,26361055 | 0,848 | 0,274 | 1,78E-115 | 4 |
| *AOX1.2* | 1,36E-118 | 2,120166198 | 0,859 | 0,281 | 1,62E-114 | 4 |
| *CG31694.3* | 2,49E-117 | 2,23630049 | 0,866 | 0,327 | 2,97E-113 | 4 |
| *Desat1.3* | 2,70E-116 | -3,189107459 | 0,483 | 0,934 | 3,22E-112 | 4 |
| *rgn.3* | 1,01E-115 | 2,346898204 | 0,928 | 0,413 | 1,20E-111 | 4 |
| *Piezo.3* | 4,62E-115 | 2,252087514 | 0,9 | 0,355 | 5,51E-111 | 4 |
| *Hsp23.1* | 4,13E-113 | 2,299056168 | 0,559 | 0,091 | 4,92E-109 | 4 |
| *Arc1* | 1,33E-112 | 2,360308751 | 0,528 | 0,086 | 1,58E-108 | 4 |
| *rhea.3* | 1,90E-109 | 2,233565655 | 0,914 | 0,451 | 2,26E-105 | 4 |
| *Hsp83.3* | 5,81E-108 | 1,892515505 | 0,986 | 0,709 | 6,92E-104 | 4 |
| *Xrp1.2* | 9,38E-108 | 1,415314715 | 0,997 | 0,902 | 1,12E-103 | 4 |
| *CG8668* | 2,47E-107 | 2,050733452 | 0,562 | 0,105 | 2,94E-103 | 4 |
| *comm2.3* | 1,70E-105 | 2,093241693 | 0,741 | 0,214 | 2,03E-101 | 4 |
| *scb.1* | 1,90E-105 | 2,214837192 | 0,645 | 0,151 | 2,27E-101 | 4 |
| *Ssrp* | 6,16E-104 | 1,862129393 | 0,41 | 0,05 | 7,35E-100 | 4 |
| *Tep4.1* | 4,18E-102 | 1,984796887 | 0,566 | 0,106 | 4,98E-98 | 4 |
| *Hsc70Cb.3* | 3,27E-101 | 1,861281303 | 0,938 | 0,486 | 3,90E-97 | 4 |
| *CG5953.2* | 1,53E-99 | 1,708979312 | 0,997 | 0,647 | 1,83E-95 | 4 |

| | | | | | | |
|---|---|---|---|---|---|---|
| *Pvr.3* | 1,97E-98 | 2,080517518 | 0,897 | 0,39 | 2,35E-94 | 4 |
| *Socs36E.3* | 3,54E-97 | 1,934535421 | 0,81 | 0,274 | 4,23E-93 | 4 |
| *GstD5* | 2,14E-94 | 1,647746809 | 0,297 | 0,024 | 2,56E-90 | 4 |
| *unc-45* | 2,27E-93 | 1,99343746 | 0,362 | 0,042 | 2,70E-89 | 4 |
| *Hsp26.3* | 6,83E-92 | 1,475856997 | 0,934 | 0,369 | 8,15E-88 | 4 |
| *dos.2* | 1,42E-91 | 2,023466504 | 0,597 | 0,145 | 1,69E-87 | 4 |
| *chic.3* | 2,88E-91 | 1,824152046 | 0,855 | 0,373 | 3,43E-87 | 4 |
| *Apoltp.2* | 6,41E-91 | 1,558724737 | 0,972 | 0,776 | 7,65E-87 | 4 |
| *vir-1.3* | 3,47E-90 | 1,56142859 | 0,979 | 0,714 | 4,14E-86 | 4 |
| *Mocs1* | 1,73E-89 | 1,866048425 | 0,424 | 0,065 | 2,07E-85 | 4 |
| *Ntan1.3* | 3,36E-87 | 1,642986412 | 0,941 | 0,572 | 4,01E-83 | 4 |
| *Hsp68.3* | 9,53E-87 | 1,670100066 | 0,772 | 0,251 | 1,14E-82 | 4 |
| *path.3* | 2,83E-86 | -2,449368366 | 0,59 | 0,918 | 3,38E-82 | 4 |
| *puc.3* | 4,38E-85 | 1,489980007 | 0,976 | 0,665 | 5,23E-81 | 4 |
| *kug* | 7,42E-85 | 1,900022027 | 0,503 | 0,106 | 8,85E-81 | 4 |
| *Abl.3* | 2,58E-84 | 1,648906302 | 0,834 | 0,325 | 3,07E-80 | 4 |
| *CG11791.3* | 3,49E-83 | 1,846352569 | 0,848 | 0,391 | 4,16E-79 | 4 |
| *CG32369.3* | 3,74E-83 | 1,703011617 | 0,955 | 0,644 | 4,46E-79 | 4 |
| *oys.3* | 3,85E-83 | 1,714835867 | 0,79 | 0,286 | 4,59E-79 | 4 |
| *Ubi-p63E.3* | 6,06E-82 | 1,498950146 | 0,945 | 0,579 | 7,23E-78 | 4 |
| *mbf1.3* | 2,17E-81 | 1,822697268 | 0,783 | 0,306 | 2,59E-77 | 4 |
| *GstD3* | 3,57E-81 | 1,781792909 | 0,376 | 0,055 | 4,25E-77 | 4 |
| *Ets21C* | 1,30E-80 | 1,676165003 | 0,428 | 0,073 | 1,55E-76 | 4 |
| *GstD10* | 1,33E-80 | 1,559965219 | 0,314 | 0,036 | 1,59E-76 | 4 |
| *CG2991.2* | 1,68E-80 | 2,002351893 | 0,586 | 0,159 | 2,00E-76 | 4 |
| *ACC.3* | 9,33E-80 | -3,334109691 | 0,21 | 0,754 | 1,11E-75 | 4 |
| *CaMKI.2* | 1,59E-79 | -1,493934571 | 0,869 | 0,971 | 1,90E-75 | 4 |
| *CG7470.2* | 2,40E-79 | -2,928565848 | 0,252 | 0,776 | 2,87E-75 | 4 |
| *wdp.1* | 1,93E-78 | 1,84591776 | 0,745 | 0,27 | 2,31E-74 | 4 |
| *CG45050.2* | 2,61E-78 | 1,221843956 | 0,993 | 0,893 | 3,11E-74 | 4 |
| *CG7130.3* | 3,96E-78 | 1,570288754 | 0,707 | 0,227 | 4,72E-74 | 4 |
| *LamC.3* | 3,29E-77 | 1,737615037 | 0,686 | 0,223 | 3,93E-73 | 4 |
| *CG7945.3* | 1,58E-76 | 1,855039088 | 0,645 | 0,197 | 1,88E-72 | 4 |
| *CG32521.2* | 1,69E-76 | -2,63472589 | 0,369 | 0,81 | 2,02E-72 | 4 |
| *pyr.2* | 2,32E-76 | 1,627702635 | 0,734 | 0,253 | 2,76E-72 | 4 |
| *Trxr-1.3* | 8,48E-76 | 1,601254175 | 0,824 | 0,347 | 1,01E-71 | 4 |
| *SP1173.2* | 5,50E-75 | 1,938595464 | 0,572 | 0,154 | 6,56E-71 | 4 |
| *Fer1HCH.2* | 1,44E-74 | 1,384266672 | 0,976 | 0,785 | 1,72E-70 | 4 |
| *NUCB1.2* | 9,69E-74 | 1,946885597 | 0,607 | 0,187 | 1,16E-69 | 4 |
| *baf.2* | 3,70E-72 | 1,820523967 | 0,583 | 0,167 | 4,41E-68 | 4 |
| *sky.2* | 1,45E-71 | 2,071921005 | 0,621 | 0,205 | 1,73E-67 | 4 |
| *bru1.2* | 1,18E-70 | -2,219705924 | 0,497 | 0,83 | 1,41E-66 | 4 |
| *raw.2* | 1,43E-70 | 1,266786992 | 0,959 | 0,601 | 1,70E-66 | 4 |
| *GstD2* | 3,05E-69 | 1,676769985 | 0,403 | 0,076 | 3,64E-65 | 4 |
| *CG18135.1* | 1,15E-68 | -2,574558877 | 0,303 | 0,764 | 1,37E-64 | 4 |
| *vri.3* | 2,24E-68 | 1,594209536 | 0,848 | 0,465 | 2,67E-64 | 4 |
| *Glut4EF.1* | 8,51E-68 | -2,048636693 | 0,521 | 0,852 | 1,02E-63 | 4 |
| *CCT8* | 9,22E-68 | 1,590778007 | 0,431 | 0,091 | 1,10E-63 | 4 |
| *cora* | 2,03E-67 | 1,776848535 | 0,528 | 0,145 | 2,42E-63 | 4 |
| *CG12290.2* | 4,89E-67 | 1,845851165 | 0,662 | 0,238 | 5,83E-63 | 4 |
| *Lac.1* | 7,64E-67 | 1,613401367 | 0,514 | 0,13 | 9,12E-63 | 4 |
| *CG13315.2* | 9,43E-67 | -2,74131542 | 0,566 | 0,848 | 1,13E-62 | 4 |
| *Sdc.2* | 1,50E-66 | -1,452203245 | 0,845 | 0,959 | 1,79E-62 | 4 |
| *TER94.2* | 1,07E-65 | 1,518837971 | 0,848 | 0,456 | 1,28E-61 | 4 |
| *scyl.2* | 1,48E-65 | 1,357798989 | 0,99 | 0,79 | 1,77E-61 | 4 |
| *Pde9.2* | 1,57E-65 | -2,322061421 | 0,362 | 0,784 | 1,88E-61 | 4 |
| *ref(2)P.2* | 3,04E-65 | 1,410083163 | 0,955 | 0,604 | 3,62E-61 | 4 |
| *FASN1.2* | 3,79E-65 | -3,09304287 | 0,49 | 0,822 | 4,52E-61 | 4 |
| *pAbp.2* | 7,16E-65 | 1,22286871 | 0,969 | 0,786 | 8,54E-61 | 4 |
| *CG4716.3* | 7,34E-65 | -1,869090693 | 0,603 | 0,883 | 8,76E-61 | 4 |
| *drpr.3* | 1,35E-64 | 1,455879678 | 0,838 | 0,418 | 1,61E-60 | 4 |

| | | | | | | |
|---|---|---|---|---|---|---|
| *Gclm.3* | 3,27E-64 | 1,427579066 | 0,769 | 0,32 | 3,90E-60 | 4 |
| *Smg6.2* | 3,88E-64 | 1,429692628 | 0,603 | 0,186 | 4,63E-60 | 4 |
| *Frl.3* | 5,65E-64 | 1,354158469 | 0,883 | 0,476 | 6,74E-60 | 4 |
| *CG10527.2* | 6,11E-64 | 1,72139519 | 0,579 | 0,182 | 7,29E-60 | 4 |
| *AspRS.3* | 1,13E-63 | 1,450100678 | 0,607 | 0,189 | 1,35E-59 | 4 |
| *eIF2beta.2* | 1,79E-63 | 1,302995675 | 0,91 | 0,489 | 2,13E-59 | 4 |
| *AttB.2* | 2,89E-63 | 1,754150855 | 0,552 | 0,157 | 3,45E-59 | 4 |
| *CG2233.3* | 3,72E-63 | -1,688901511 | 0,738 | 0,933 | 4,44E-59 | 4 |
| *Nop17l.3* | 4,10E-63 | 1,645049803 | 0,79 | 0,376 | 4,89E-59 | 4 |
| *Egfr.2* | 4,16E-63 | -2,150408022 | 0,407 | 0,811 | 4,97E-59 | 4 |
| *Hsc70-4.3* | 2,39E-62 | 1,167600491 | 0,955 | 0,718 | 2,86E-58 | 4 |
| *CG12795.3* | 1,16E-61 | 1,366989745 | 0,769 | 0,307 | 1,38E-57 | 4 |
| *neur* | 1,59E-61 | 1,795668957 | 0,462 | 0,117 | 1,89E-57 | 4 |
| *dnc.3* | 1,26E-60 | -1,999355918 | 0,383 | 0,79 | 1,50E-56 | 4 |
| *CG6503.2* | 2,41E-60 | -2,431856295 | 0,528 | 0,828 | 2,88E-56 | 4 |
| *stc* | 2,65E-60 | 1,645556714 | 0,403 | 0,089 | 3,16E-56 | 4 |
| *Tps1.1* | 2,79E-60 | -1,367192026 | 0,793 | 0,958 | 3,33E-56 | 4 |
| *cher.2* | 3,08E-60 | 1,519502713 | 0,676 | 0,253 | 3,67E-56 | 4 |
| *Rtnl1.1* | 3,31E-60 | 1,591459485 | 0,731 | 0,311 | 3,95E-56 | 4 |
| *AdSS.2* | 7,53E-60 | 1,431675689 | 0,807 | 0,376 | 8,99E-56 | 4 |
| *poe.2* | 1,01E-59 | 1,445883547 | 0,71 | 0,302 | 1,21E-55 | 4 |
| *KrT95D.3* | 4,46E-59 | 1,259686264 | 0,921 | 0,66 | 5,32E-55 | 4 |
| *pes.1* | 1,34E-58 | 1,578084587 | 0,703 | 0,304 | 1,60E-54 | 4 |
| *Ubx.2* | 4,83E-58 | -1,740632833 | 0,721 | 0,87 | 5,76E-54 | 4 |
| *PCB.3* | 1,86E-57 | -2,456162468 | 0,176 | 0,656 | 2,21E-53 | 4 |
| *GstD9* | 2,36E-57 | 1,313853585 | 0,39 | 0,085 | 2,81E-53 | 4 |
| *eff.2* | 4,51E-57 | 1,334403383 | 0,852 | 0,564 | 5,38E-53 | 4 |
| *CG30069.2* | 4,72E-57 | 1,611155277 | 0,686 | 0,292 | 5,64E-53 | 4 |
| *Chd64.2* | 7,68E-57 | 1,373072664 | 0,821 | 0,419 | 9,16E-53 | 4 |
| *Nup358.1* | 8,42E-57 | 1,576862568 | 0,576 | 0,201 | 1,00E-52 | 4 |
| *trbl.2* | 1,58E-56 | 1,324980608 | 0,845 | 0,479 | 1,88E-52 | 4 |
| *Zip99C.1* | 1,75E-56 | 1,594045853 | 0,528 | 0,164 | 2,09E-52 | 4 |
| *comm* | 5,54E-56 | 1,39730619 | 0,403 | 0,091 | 6,61E-52 | 4 |
| *DnaJ-1.3* | 5,63E-56 | 1,277575553 | 0,879 | 0,499 | 6,72E-52 | 4 |
| *Act5C.3* | 7,25E-56 | 1,245821254 | 0,872 | 0,513 | 8,65E-52 | 4 |
| *CG32195* | 3,34E-55 | 1,351359991 | 0,29 | 0,048 | 3,99E-51 | 4 |
| *Hsc70-5.2* | 1,29E-54 | 1,504318113 | 0,631 | 0,247 | 1,53E-50 | 4 |
| *Prps.1* | 4,64E-54 | -1,46506359 | 0,814 | 0,936 | 5,54E-50 | 4 |
| *CG5290* | 1,12E-53 | 1,249705746 | 0,3 | 0,052 | 1,33E-49 | 4 |
| *be* | 3,08E-53 | 1,573275714 | 0,445 | 0,119 | 3,68E-49 | 4 |
| *CG5773.3* | 1,20E-52 | 1,619964273 | 0,566 | 0,197 | 1,43E-48 | 4 |
| *apolpp.2* | 2,43E-52 | -2,704739164 | 0,603 | 0,824 | 2,90E-48 | 4 |
| *h.1* | 2,97E-52 | 1,449987574 | 0,91 | 0,606 | 3,55E-48 | 4 |
| *Naam* | 3,20E-52 | 1,491027858 | 0,269 | 0,043 | 3,82E-48 | 4 |
| *CCT1.1* | 3,98E-52 | 1,326613294 | 0,476 | 0,136 | 4,74E-48 | 4 |
| *Pvf2* | 4,57E-52 | 1,948505735 | 0,259 | 0,04 | 5,45E-48 | 4 |
| *Ctl2.2* | 8,65E-52 | 1,415217899 | 0,507 | 0,155 | 1,03E-47 | 4 |
| *CG16898.2* | 1,77E-51 | 1,230058728 | 0,814 | 0,399 | 2,11E-47 | 4 |
| *Nmda1.3* | 2,03E-51 | 1,280530771 | 0,71 | 0,315 | 2,42E-47 | 4 |
| *edl* | 2,31E-51 | 1,629260449 | 0,445 | 0,125 | 2,75E-47 | 4 |
| *CG34166.2* | 2,91E-51 | -3,148702299 | 0,569 | 0,826 | 3,47E-47 | 4 |
| *chrb.2* | 7,28E-51 | 1,683874262 | 0,555 | 0,202 | 8,69E-47 | 4 |
| *Smg5.2* | 2,13E-50 | 1,284010524 | 0,834 | 0,45 | 2,54E-46 | 4 |
| *RpA-70* | 2,16E-50 | 1,229920827 | 0,276 | 0,047 | 2,58E-46 | 4 |
| *CG6910.3* | 2,70E-50 | -2,140093003 | 0,479 | 0,787 | 3,22E-46 | 4 |
| *ctp.2* | 2,85E-50 | 1,449010233 | 0,645 | 0,257 | 3,40E-46 | 4 |
| *CG5004* | 4,55E-50 | 1,403569299 | 0,372 | 0,088 | 5,43E-46 | 4 |
| *Paics.2* | 4,95E-50 | -1,767954358 | 0,497 | 0,813 | 5,91E-46 | 4 |
| *CCT7.1* | 1,57E-49 | 1,42047967 | 0,421 | 0,112 | 1,87E-45 | 4 |
| *loco.2* | 2,34E-49 | 1,231561698 | 0,641 | 0,237 | 2,80E-45 | 4 |
| *Snoo.2* | 2,66E-49 | -1,526308951 | 0,655 | 0,859 | 3,17E-45 | 4 |

| | | | | | | |
|---|---|---|---|---|---|---|
| *CG10737* | 4,61E-49 | 1,488312705 | 0,49 | 0,156 | 5,51E-45 | 4 |
| *Gadd45.2* | 5,51E-49 | 1,481293503 | 0,555 | 0,199 | 6,58E-45 | 4 |
| *CG2064* | 1,26E-48 | 1,449758567 | 0,39 | 0,099 | 1,50E-44 | 4 |
| *Clbn* | 1,95E-48 | 1,254523426 | 0,324 | 0,068 | 2,32E-44 | 4 |
| *trio* | 3,94E-48 | 1,510331692 | 0,476 | 0,149 | 4,70E-44 | 4 |
| *Lst.3* | 5,90E-48 | -1,749744458 | 0,576 | 0,815 | 7,03E-44 | 4 |
| *Hsp70Bc.2* | 6,58E-48 | 1,354624224 | 0,486 | 0,148 | 7,85E-44 | 4 |
| *GstE7.2* | 9,48E-48 | 1,351129283 | 0,486 | 0,153 | 1,13E-43 | 4 |
| *GEFmeso.2* | 1,04E-47 | 1,117831138 | 0,81 | 0,401 | 1,24E-43 | 4 |
| *ImpL2.3* | 1,08E-47 | 1,364859533 | 0,579 | 0,214 | 1,29E-43 | 4 |
| *CG46339* | 4,01E-46 | 1,482711534 | 0,417 | 0,116 | 4,78E-42 | 4 |
| *foxo.3* | 4,12E-46 | -1,556185394 | 0,434 | 0,769 | 4,91E-42 | 4 |
| *CG7766.2* | 3,32E-45 | -2,04049013 | 0,2 | 0,611 | 3,96E-41 | 4 |
| *CG10960.3* | 4,58E-45 | -1,827040051 | 0,655 | 0,851 | 5,46E-41 | 4 |
| *Droj2.2* | 6,23E-45 | 1,226584565 | 0,738 | 0,382 | 7,43E-41 | 4 |
| *MFS17.2* | 1,14E-44 | -1,712717525 | 0,397 | 0,726 | 1,36E-40 | 4 |
| *Syx1A.3* | 2,34E-44 | 1,251555973 | 0,845 | 0,518 | 2,80E-40 | 4 |
| *Cyt-b5-r.2* | 1,53E-43 | -2,305499321 | 0,21 | 0,6 | 1,82E-39 | 4 |
| *GstS1.1* | 1,69E-43 | 1,389141851 | 0,534 | 0,197 | 2,01E-39 | 4 |
| *noc.2* | 1,73E-43 | 1,468083147 | 0,51 | 0,185 | 2,07E-39 | 4 |
| *cnn.2* | 4,05E-43 | 1,300365441 | 0,593 | 0,245 | 4,83E-39 | 4 |
| *Nplp2.3* | 6,02E-43 | -2,57576115 | 0,376 | 0,698 | 7,18E-39 | 4 |
| *RhoU* | 6,70E-43 | 1,341122646 | 0,483 | 0,166 | 8,00E-39 | 4 |
| *Acsl.2* | 9,52E-43 | -2,342095018 | 0,207 | 0,592 | 1,14E-38 | 4 |
| *Xpc* | 1,20E-42 | 1,335271118 | 0,397 | 0,116 | 1,43E-38 | 4 |
| *bark* | 1,23E-42 | 1,522903175 | 0,31 | 0,071 | 1,47E-38 | 4 |
| *CG16758.2* | 2,62E-42 | -1,320567567 | 0,859 | 0,928 | 3,13E-38 | 4 |
| *Gclc.2* | 3,48E-42 | 1,134513282 | 0,831 | 0,5 | 4,15E-38 | 4 |
| *AdipoR.2* | 4,54E-42 | -2,077331591 | 0,179 | 0,574 | 5,41E-38 | 4 |
| *MCU.2* | 4,59E-42 | 1,22740976 | 0,607 | 0,253 | 5,48E-38 | 4 |
| *Khc-73.3* | 6,79E-42 | 1,155732422 | 0,693 | 0,341 | 8,10E-38 | 4 |
| *PEK* | 7,04E-42 | 1,215978649 | 0,379 | 0,105 | 8,40E-38 | 4 |
| *Best1* | 7,71E-42 | 1,37217732 | 0,362 | 0,099 | 9,20E-38 | 4 |
| *CG4797* | 8,08E-42 | 1,443371503 | 0,445 | 0,144 | 9,64E-38 | 4 |
| *GstE3.2* | 9,07E-42 | 1,327040613 | 0,497 | 0,178 | 1,08E-37 | 4 |
| *CG10638* | 9,70E-42 | 1,410661058 | 0,303 | 0,07 | 1,16E-37 | 4 |
| *mfas* | 1,20E-41 | 1,315551256 | 0,369 | 0,101 | 1,43E-37 | 4 |
| *smash.1* | 5,17E-41 | -1,841435613 | 0,29 | 0,675 | 6,16E-37 | 4 |
| *subdued* | 8,84E-41 | 1,325640744 | 0,328 | 0,082 | 1,05E-36 | 4 |
| *CG3168.2* | 1,06E-40 | 1,396266568 | 0,6 | 0,264 | 1,27E-36 | 4 |
| *Hsp70Ab.1* | 1,53E-40 | 1,309416934 | 0,462 | 0,157 | 1,83E-36 | 4 |
| *Psc* | 1,82E-40 | 1,356321173 | 0,3 | 0,069 | 2,17E-36 | 4 |
| *mtd.2* | 3,42E-40 | -1,821078156 | 0,421 | 0,719 | 4,08E-36 | 4 |
| *Pak3.3* | 3,67E-40 | 1,313551271 | 0,59 | 0,258 | 4,38E-36 | 4 |
| *CG15099.3* | 4,85E-40 | 1,274758775 | 0,6 | 0,266 | 5,78E-36 | 4 |
| *CG1416.2* | 5,54E-40 | 1,250394157 | 0,459 | 0,156 | 6,61E-36 | 4 |
| *lola.2* | 6,47E-40 | -1,264101016 | 0,741 | 0,876 | 7,71E-36 | 4 |
| *Sox102F.2* | 7,33E-40 | -1,477045079 | 0,507 | 0,772 | 8,74E-36 | 4 |
| *bchs.1* | 1,12E-39 | 1,35878473 | 0,441 | 0,149 | 1,33E-35 | 4 |
| *CCT2.3* | 1,34E-39 | 1,310394771 | 0,552 | 0,227 | 1,60E-35 | 4 |
| *Vps13.2* | 1,84E-39 | 1,149573453 | 0,676 | 0,33 | 2,20E-35 | 4 |
| *CG6357* | 2,24E-39 | 1,666103882 | 0,486 | 0,192 | 2,67E-35 | 4 |
| *Smr.3* | 2,75E-39 | -1,407354416 | 0,5 | 0,774 | 3,28E-35 | 4 |
| *CG10383.2* | 1,10E-38 | 1,058566739 | 0,81 | 0,437 | 1,32E-34 | 4 |
| *Gnmt.1* | 1,23E-38 | -1,856713998 | 0,355 | 0,675 | 1,46E-34 | 4 |
| *Swip-1.3* | 1,91E-38 | 1,099308792 | 0,714 | 0,369 | 2,27E-34 | 4 |
| *px.2* | 3,26E-38 | -1,551359683 | 0,479 | 0,757 | 3,89E-34 | 4 |
| *RASSF8* | 5,47E-38 | 1,333365161 | 0,417 | 0,138 | 6,53E-34 | 4 |
| *Ntf-2* | 6,29E-38 | 1,312240557 | 0,428 | 0,144 | 7,50E-34 | 4 |
| *Mrp4* | 7,00E-38 | 1,284315266 | 0,438 | 0,147 | 8,35E-34 | 4 |
| *CG33494.3* | 7,62E-38 | 1,134460256 | 0,621 | 0,289 | 9,09E-34 | 4 |

| | | | | | | |
|---|---|---|---|---|---|---|
| *CG42588.2* | 7,81E-38 | 1,241269677 | 0,631 | 0,286 | 9,32E-34 | 4 |
| *Apc* | 3,56E-37 | 1,172462098 | 0,334 | 0,09 | 4,25E-33 | 4 |
| *MESR4* | 5,51E-37 | 1,252383929 | 0,41 | 0,136 | 6,57E-33 | 4 |
| *stac* | 9,56E-37 | 1,275744306 | 0,348 | 0,102 | 1,14E-32 | 4 |
| *Hrs* | 1,72E-36 | 1,123312259 | 0,362 | 0,107 | 2,05E-32 | 4 |
| *CG12004.2* | 2,05E-36 | 1,188886364 | 0,562 | 0,24 | 2,45E-32 | 4 |
| *Pect.3* | 2,21E-36 | -1,91481585 | 0,238 | 0,592 | 2,64E-32 | 4 |
| *CG41378.2* | 2,31E-36 | -1,847793219 | 0,217 | 0,577 | 2,76E-32 | 4 |
| *Pdp1.3* | 2,95E-36 | -0,795666503 | 0,986 | 0,994 | 3,52E-32 | 4 |
| *eIF4A.2* | 4,25E-36 | 0,968637035 | 0,841 | 0,582 | 5,07E-32 | 4 |
| *swm.3* | 4,86E-36 | 1,31763556 | 0,607 | 0,3 | 5,79E-32 | 4 |
| *Parp.3* | 6,19E-36 | -1,448607057 | 0,41 | 0,706 | 7,39E-32 | 4 |
| *CG15611* | 7,02E-36 | 1,256639102 | 0,376 | 0,116 | 8,37E-32 | 4 |
| *CCT5.1* | 8,80E-36 | 1,178844241 | 0,397 | 0,129 | 1,05E-31 | 4 |
| *lilli.2* | 2,45E-35 | -1,280686742 | 0,493 | 0,755 | 2,92E-31 | 4 |
| *Su(var)2-HP2.1* | 5,63E-35 | 1,311552977 | 0,507 | 0,214 | 6,71E-31 | 4 |
| *GstE1.2* | 5,88E-35 | 0,936267454 | 0,69 | 0,345 | 7,01E-31 | 4 |
| *robo2.2* | 7,18E-35 | 1,164715861 | 0,707 | 0,393 | 8,57E-31 | 4 |
| *AcCoAS.2* | 9,06E-35 | -1,804157758 | 0,441 | 0,708 | 1,08E-30 | 4 |
| *InR.2* | 1,74E-34 | 0,68792667 | 0,993 | 0,905 | 2,07E-30 | 4 |
| *Diap1.3* | 2,49E-34 | 0,885311114 | 0,903 | 0,732 | 2,97E-30 | 4 |
| *Usp1* | 3,07E-34 | 1,274709473 | 0,434 | 0,162 | 3,66E-30 | 4 |
| *dsx.3* | 3,16E-34 | -1,956055092 | 0,21 | 0,565 | 3,77E-30 | 4 |
| *Rad23.3* | 5,57E-34 | 1,074093376 | 0,624 | 0,303 | 6,64E-30 | 4 |
| *Swim* | 5,69E-34 | 1,549268064 | 0,303 | 0,083 | 6,79E-30 | 4 |
| *CG32647.2* | 8,51E-34 | -2,143108322 | 0,269 | 0,594 | 1,01E-29 | 4 |
| *Rpn6.2* | 1,34E-33 | 1,079094646 | 0,676 | 0,36 | 1,60E-29 | 4 |
| *LpR2.2* | 1,85E-33 | -1,473494074 | 0,748 | 0,857 | 2,20E-29 | 4 |
| *siz.2* | 2,24E-33 | 1,166416455 | 0,738 | 0,443 | 2,67E-29 | 4 |
| *CG8468.1* | 2,51E-33 | -1,64354616 | 0,414 | 0,701 | 2,99E-29 | 4 |
| *NFAT.2* | 3,17E-33 | -1,436944946 | 0,469 | 0,731 | 3,79E-29 | 4 |
| *Galk.2* | 4,32E-33 | -1,829836582 | 0,183 | 0,534 | 5,16E-29 | 4 |
| *capt* | 6,83E-33 | 1,151994148 | 0,279 | 0,072 | 8,14E-29 | 4 |
| *CG17108.3* | 7,84E-33 | -2,600601558 | 0,079 | 0,434 | 9,35E-29 | 4 |
| *CG1129* | 8,64E-33 | 1,163332432 | 0,372 | 0,122 | 1,03E-28 | 4 |
| *GramD1B.1* | 9,83E-33 | 1,003167778 | 0,807 | 0,526 | 1,17E-28 | 4 |
| *kmr.2* | 2,03E-32 | 1,241538508 | 0,5 | 0,211 | 2,42E-28 | 4 |
| *Stip1.1* | 2,42E-32 | 1,072621514 | 0,369 | 0,12 | 2,89E-28 | 4 |
| *lqf.2* | 2,79E-32 | 1,003958297 | 0,81 | 0,534 | 3,33E-28 | 4 |
| *spict.1* | 3,52E-32 | 1,13266447 | 0,39 | 0,132 | 4,20E-28 | 4 |
| *Ada2b* | 4,03E-32 | 1,121759546 | 0,352 | 0,11 | 4,81E-28 | 4 |
| *mamo.3* | 5,45E-32 | -1,786025026 | 0,255 | 0,586 | 6,50E-28 | 4 |
| *Tpr2.2* | 5,57E-32 | 1,0061215 | 0,828 | 0,574 | 6,64E-28 | 4 |
| *tsr.3* | 5,86E-32 | 1,16692119 | 0,514 | 0,223 | 6,99E-28 | 4 |
| *RasGAP1.3* | 8,62E-32 | 1,221821761 | 0,541 | 0,251 | 1,03E-27 | 4 |
| *CG17841.2* | 9,01E-32 | -1,889852093 | 0,176 | 0,507 | 1,07E-27 | 4 |
| *cpa* | 9,62E-32 | 1,047623792 | 0,31 | 0,088 | 1,15E-27 | 4 |
| *c11.1.2* | 1,31E-31 | 1,124140634 | 0,51 | 0,22 | 1,56E-27 | 4 |
| *kibra.2* | 4,63E-31 | 1,125159598 | 0,717 | 0,417 | 5,53E-27 | 4 |
| *Men.2* | 9,64E-31 | -2,040353202 | 0,462 | 0,694 | 1,15E-26 | 4 |
| *Chchd2.3* | 9,88E-31 | 0,827449555 | 0,845 | 0,595 | 1,18E-26 | 4 |
| *rdog* | 1,05E-30 | 1,325501542 | 0,345 | 0,113 | 1,25E-26 | 4 |
| *Hex-C.3* | 1,58E-30 | -1,779486368 | 0,148 | 0,476 | 1,88E-26 | 4 |
| *baz.2* | 1,65E-30 | 1,144728962 | 0,497 | 0,209 | 1,97E-26 | 4 |
| *BomBc2.2* | 2,11E-30 | -2,207638768 | 0,297 | 0,584 | 2,52E-26 | 4 |
| *Prosalpha6.2* | 2,92E-30 | 1,041347388 | 0,483 | 0,2 | 3,48E-26 | 4 |
| *CG11267.2* | 3,29E-30 | 1,070750227 | 0,441 | 0,172 | 3,92E-26 | 4 |
| *CG1578.2* | 5,51E-30 | -2,117703199 | 0,079 | 0,412 | 6,57E-26 | 4 |
| *Hsc70-3.1* | 5,96E-30 | 1,042006009 | 0,69 | 0,383 | 7,11E-26 | 4 |
| *CG34376.1* | 8,36E-30 | -1,618976511 | 0,217 | 0,546 | 9,98E-26 | 4 |
| *PhKgamma.2* | 8,58E-30 | -1,401733657 | 0,324 | 0,633 | 1,02E-25 | 4 |

| | | | | | | |
|---|---|---|---|---|---|---|
| *GstD1.2* | 1,20E-29 | 0,738367982 | 0,928 | 0,626 | 1,43E-25 | 4 |
| *rl.2* | 1,27E-29 | -0,882980802 | 0,776 | 0,91 | 1,51E-25 | 4 |
| *CG18171* | 1,56E-29 | 0,989904271 | 0,355 | 0,117 | 1,86E-25 | 4 |
| *sbb.3* | 1,73E-29 | -1,513460942 | 0,307 | 0,617 | 2,06E-25 | 4 |
| *CG14823.3* | 1,95E-29 | -2,262583277 | 0,048 | 0,376 | 2,33E-25 | 4 |
| *betaTub56D* | 2,63E-29 | 1,219263037 | 0,383 | 0,14 | 3,14E-25 | 4 |
| *CG44325.3* | 2,75E-29 | 0,918426687 | 0,603 | 0,289 | 3,28E-25 | 4 |
| *Akr1B.1* | 3,03E-29 | 1,131550862 | 0,49 | 0,215 | 3,61E-25 | 4 |
| *DnaJ-H.1* | 3,14E-29 | 1,030183875 | 0,397 | 0,145 | 3,74E-25 | 4 |
| *Gbs-70E.3* | 3,29E-29 | -2,905365873 | 0,269 | 0,556 | 3,92E-25 | 4 |
| *Pdfr.2* | 4,81E-29 | -1,765429338 | 0,186 | 0,507 | 5,74E-25 | 4 |
| *Ref1.1* | 5,03E-29 | 1,083826293 | 0,514 | 0,234 | 6,00E-25 | 4 |
| *sra.2* | 8,68E-29 | 1,018886297 | 0,534 | 0,243 | 1,04E-24 | 4 |
| *maf-S.2* | 1,35E-28 | 1,099207682 | 0,434 | 0,178 | 1,61E-24 | 4 |
| *chb.2* | 1,53E-28 | 1,058601129 | 0,562 | 0,267 | 1,82E-24 | 4 |
| *CCT4* | 1,95E-28 | 1,037166119 | 0,331 | 0,109 | 2,33E-24 | 4 |
| *Ptpmeg.1* | 2,21E-28 | -1,510773059 | 0,248 | 0,57 | 2,64E-24 | 4 |
| *rad50* | 2,27E-28 | 1,379831758 | 0,321 | 0,108 | 2,71E-24 | 4 |
| *Cat.2* | 2,60E-28 | 1,137129226 | 0,61 | 0,322 | 3,10E-24 | 4 |
| *cnc* | 3,14E-28 | 0,541389017 | 0,997 | 0,959 | 3,75E-24 | 4 |
| *tamo.1* | 4,55E-28 | 0,946541464 | 0,397 | 0,145 | 5,42E-24 | 4 |
| *mino.2* | 6,60E-28 | -1,322481119 | 0,448 | 0,698 | 7,87E-24 | 4 |
| *AnxB11* | 1,57E-27 | 1,012006546 | 0,324 | 0,107 | 1,88E-23 | 4 |
| *CG46385.2* | 1,79E-27 | -0,84306927 | 0,931 | 0,959 | 2,13E-23 | 4 |
| *CG31689.1* | 2,12E-27 | -1,331730597 | 0,438 | 0,701 | 2,53E-23 | 4 |
| *CG42668.3* | 2,55E-27 | 0,791627321 | 0,866 | 0,572 | 3,04E-23 | 4 |
| *Rab11.3* | 3,92E-27 | 0,950735839 | 0,624 | 0,34 | 4,68E-23 | 4 |
| *Eglp4.3* | 4,60E-27 | -2,312602843 | 0,093 | 0,398 | 5,49E-23 | 4 |
| *eas.3* | 4,85E-27 | -1,761268176 | 0,197 | 0,498 | 5,79E-23 | 4 |
| *Culd.3* | 6,29E-27 | -2,114229257 | 0,069 | 0,372 | 7,50E-23 | 4 |
| *Uba1.2* | 8,25E-27 | 1,237477451 | 0,421 | 0,178 | 9,84E-23 | 4 |
| *GstE8.2* | 1,01E-26 | 0,945319886 | 0,438 | 0,179 | 1,20E-22 | 4 |
| *wun.1* | 1,07E-26 | 1,017306526 | 0,41 | 0,162 | 1,28E-22 | 4 |
| *inaE* | 1,27E-26 | 1,043019197 | 0,355 | 0,127 | 1,52E-22 | 4 |
| *ttk.2* | 1,35E-26 | 1,258647458 | 0,486 | 0,229 | 1,61E-22 | 4 |
| *BomT3.3* | 1,94E-26 | -2,316435814 | 0,152 | 0,455 | 2,31E-22 | 4 |
| *kek5.2* | 2,74E-26 | -1,439018187 | 0,352 | 0,627 | 3,27E-22 | 4 |
| *luna.3* | 2,77E-26 | 0,752211482 | 0,893 | 0,702 | 3,31E-22 | 4 |
| *CG7720.3* | 6,01E-26 | -2,771810912 | 0,083 | 0,383 | 7,16E-22 | 4 |
| *GlcT* | 6,04E-26 | 0,974254295 | 0,279 | 0,085 | 7,21E-22 | 4 |
| *Rpn5.2* | 6,56E-26 | 0,981013491 | 0,51 | 0,24 | 7,82E-22 | 4 |
| *EndoA* | 7,03E-26 | 1,179275991 | 0,293 | 0,095 | 8,38E-22 | 4 |
| *Drak.2* | 7,44E-26 | -1,977212501 | 0,421 | 0,642 | 8,88E-22 | 4 |
| *CIAPIN1.1* | 8,50E-26 | 0,995450577 | 0,366 | 0,137 | 1,01E-21 | 4 |
| *Pde6.3* | 1,46E-25 | -1,486789028 | 0,234 | 0,54 | 1,75E-21 | 4 |
| *Dys.3* | 1,52E-25 | -1,305577429 | 0,3 | 0,593 | 1,81E-21 | 4 |
| *Ptp61F* | 1,89E-25 | 0,961280402 | 0,328 | 0,113 | 2,26E-21 | 4 |
| *MFS14.2* | 2,01E-25 | -1,863338197 | 0,128 | 0,428 | 2,40E-21 | 4 |
| *koi* | 2,72E-25 | 1,05669231 | 0,266 | 0,08 | 3,25E-21 | 4 |
| *bgm.2* | 3,10E-25 | -1,555150991 | 0,407 | 0,659 | 3,70E-21 | 4 |
| *NK7.1.1* | 5,13E-25 | 1,051775226 | 0,459 | 0,206 | 6,12E-21 | 4 |
| *ScsbetaA.2* | 5,36E-25 | -1,531761092 | 0,228 | 0,507 | 6,39E-21 | 4 |
| *sni* | 6,47E-25 | 1,002191291 | 0,283 | 0,09 | 7,72E-21 | 4 |
| *rudhira.2* | 8,01E-25 | -1,120136981 | 0,462 | 0,706 | 9,56E-21 | 4 |
| *bun.3* | 8,69E-25 | -1,394063142 | 0,731 | 0,855 | 1,04E-20 | 4 |
| *CG17646.2* | 1,10E-24 | -1,131648651 | 0,741 | 0,863 | 1,32E-20 | 4 |
| *Rpn2.2* | 1,26E-24 | 0,950574929 | 0,548 | 0,283 | 1,51E-20 | 4 |
| *CG10082.3* | 1,27E-24 | -1,551560954 | 0,376 | 0,627 | 1,52E-20 | 4 |
| *Ork1.2* | 2,15E-24 | -1,943659126 | 0,069 | 0,364 | 2,57E-20 | 4 |
| *Pino.1* | 2,18E-24 | -1,191888645 | 0,617 | 0,782 | 2,60E-20 | 4 |
| *Mal-B2.3* | 2,23E-24 | -1,461037485 | 0,283 | 0,55 | 2,66E-20 | 4 |

| | | | | | | |
|---|---|---|---|---|---|---|
| *CG8745.3* | 2,36E-24 | -2,747164027 | 0,11 | 0,393 | 2,81E-20 | 4 |
| *Rpn9.2* | 2,60E-24 | 1,020576743 | 0,397 | 0,162 | 3,10E-20 | 4 |
| *if.3* | 2,84E-24 | -2,131521882 | 0,083 | 0,375 | 3,39E-20 | 4 |
| *Plod.3* | 4,64E-24 | 1,000413534 | 0,521 | 0,259 | 5,53E-20 | 4 |
| *CG1213.2* | 5,83E-24 | -2,092785088 | 0,083 | 0,367 | 6,96E-20 | 4 |
| *MsrA* | 2,51E-23 | 1,081633229 | 0,279 | 0,094 | 3,00E-19 | 4 |
| *Spat.2* | 3,68E-23 | -1,630205717 | 0,217 | 0,488 | 4,39E-19 | 4 |
| *mys.3* | 4,20E-23 | 0,917416032 | 0,624 | 0,369 | 5,01E-19 | 4 |
| *CG31704.2* | 5,14E-23 | 0,965200955 | 0,438 | 0,199 | 6,13E-19 | 4 |
| *l(3)80Fg.2* | 5,70E-23 | -1,242429722 | 0,397 | 0,633 | 6,80E-19 | 4 |
| *RhoGAP15B.3* | 5,71E-23 | 0,91805485 | 0,455 | 0,205 | 6,82E-19 | 4 |
| *Ufd4.2* | 6,35E-23 | 0,903925382 | 0,583 | 0,321 | 7,57E-19 | 4 |
| *Stoml2* | 6,93E-23 | 0,902599519 | 0,279 | 0,091 | 8,27E-19 | 4 |
| *alphaTub84B.1* | 7,18E-23 | 1,010777254 | 0,597 | 0,33 | 8,57E-19 | 4 |
| *Ubc6.2* | 7,88E-23 | 0,99106886 | 0,486 | 0,248 | 9,41E-19 | 4 |
| *fray.1* | 9,38E-23 | 1,012718448 | 0,49 | 0,243 | 1,12E-18 | 4 |
| *Capr* | 1,02E-22 | 1,00311207 | 0,29 | 0,101 | 1,22E-18 | 4 |
| *GATAd* | 1,04E-22 | 1,025036656 | 0,328 | 0,125 | 1,24E-18 | 4 |
| *Ire1.2* | 1,39E-22 | 0,8728088 | 0,555 | 0,293 | 1,66E-18 | 4 |
| *Shrm.1* | 1,86E-22 | 0,880193162 | 0,572 | 0,3 | 2,22E-18 | 4 |
| *Syp.2* | 2,05E-22 | -1,060333664 | 0,717 | 0,824 | 2,45E-18 | 4 |
| *par-1.2* | 2,25E-22 | -0,692734924 | 0,893 | 0,947 | 2,68E-18 | 4 |
| *wrd.2* | 2,43E-22 | -1,358832376 | 0,183 | 0,466 | 2,90E-18 | 4 |
| *CG15784.2* | 2,76E-22 | 0,948548599 | 0,459 | 0,213 | 3,30E-18 | 4 |
| *Nedd4.2* | 3,71E-22 | 0,903949706 | 0,559 | 0,309 | 4,42E-18 | 4 |
| *CG8108.2* | 3,99E-22 | 0,972610121 | 0,607 | 0,361 | 4,77E-18 | 4 |
| *tkv.2* | 4,00E-22 | -1,450591686 | 0,138 | 0,419 | 4,77E-18 | 4 |
| *CG6686* | 4,29E-22 | 0,862290528 | 0,255 | 0,08 | 5,12E-18 | 4 |
| *zormin.2* | 4,38E-22 | 0,927471819 | 0,641 | 0,387 | 5,23E-18 | 4 |
| *CG18659.2* | 4,78E-22 | 0,941464278 | 0,479 | 0,237 | 5,70E-18 | 4 |
| *Lis-1.3* | 4,84E-22 | 0,907720031 | 0,607 | 0,34 | 5,77E-18 | 4 |
| *cact.3* | 4,96E-22 | 0,624838873 | 0,859 | 0,619 | 5,91E-18 | 4 |
| *jar.1* | 1,33E-21 | 0,884646147 | 0,634 | 0,391 | 1,59E-17 | 4 |
| *Xbp1.2* | 1,53E-21 | 0,752336299 | 0,79 | 0,552 | 1,82E-17 | 4 |
| *CG2926* | 1,80E-21 | 1,038536145 | 0,462 | 0,233 | 2,15E-17 | 4 |
| *CG3764.2* | 1,88E-21 | -1,640684254 | 0,055 | 0,323 | 2,25E-17 | 4 |
| *Atpalpha.2* | 2,07E-21 | -1,038248059 | 0,566 | 0,754 | 2,47E-17 | 4 |
| *alt.2* | 2,54E-21 | 0,869818693 | 0,597 | 0,334 | 3,04E-17 | 4 |
| *Hel25E* | 3,24E-21 | 0,928162458 | 0,269 | 0,091 | 3,87E-17 | 4 |
| *shrb.2* | 3,30E-21 | 0,934797207 | 0,51 | 0,267 | 3,94E-17 | 4 |
| *CG31705.2* | 3,30E-21 | 0,750789957 | 0,772 | 0,55 | 3,94E-17 | 4 |
| *Yeti.3* | 3,60E-21 | 0,748201269 | 0,824 | 0,593 | 4,29E-17 | 4 |
| *Rpt5.2* | 3,95E-21 | 0,88926284 | 0,421 | 0,191 | 4,72E-17 | 4 |
| *Pur-alpha.2* | 4,11E-21 | -1,394671509 | 0,224 | 0,472 | 4,90E-17 | 4 |
| *CG43340.2* | 4,67E-21 | -1,394504504 | 0,169 | 0,44 | 5,57E-17 | 4 |
| *Eip75B.1* | 4,73E-21 | 0,636134514 | 0,938 | 0,841 | 5,64E-17 | 4 |
| *Fer2LCH.3* | 5,22E-21 | 0,752190945 | 0,89 | 0,73 | 6,23E-17 | 4 |
| *CG10433.2* | 5,25E-21 | -1,065824253 | 0,476 | 0,697 | 6,26E-17 | 4 |
| *Tlk.2* | 5,50E-21 | -1,157265906 | 0,348 | 0,587 | 6,57E-17 | 4 |
| *Vha16-1.1* | 6,02E-21 | 0,692104713 | 0,903 | 0,754 | 7,18E-17 | 4 |
| *CG32687.2* | 6,98E-21 | -1,251293923 | 0,245 | 0,515 | 8,32E-17 | 4 |
| *Cyp4p1.2* | 1,05E-20 | 0,972726838 | 0,531 | 0,288 | 1,26E-16 | 4 |
| *Fim* | 1,15E-20 | 0,958124384 | 0,341 | 0,141 | 1,38E-16 | 4 |
| *Sam-S.2* | 1,24E-20 | -1,511415841 | 0,431 | 0,632 | 1,47E-16 | 4 |
| *Usp14.1* | 1,44E-20 | 0,950943669 | 0,359 | 0,152 | 1,72E-16 | 4 |
| *gce.3* | 1,51E-20 | -1,243503525 | 0,231 | 0,508 | 1,80E-16 | 4 |
| *Gdap2.2* | 1,75E-20 | 0,864668005 | 0,631 | 0,384 | 2,09E-16 | 4 |
| *IP3K2.2* | 1,88E-20 | -1,033648517 | 0,448 | 0,68 | 2,24E-16 | 4 |
| *Hsp60A.3* | 1,97E-20 | 0,864345996 | 0,455 | 0,22 | 2,35E-16 | 4 |
| *ncm* | 2,82E-20 | 0,933559437 | 0,259 | 0,088 | 3,37E-16 | 4 |
| *HDAC6.2* | 4,22E-20 | -1,459627862 | 0,114 | 0,364 | 5,04E-16 | 4 |

| | | | | | | |
|---|---|---|---|---|---|---|
| *Arpc2* | 4,45E-20 | 0,869624938 | 0,283 | 0,103 | 5,31E-16 | 4 |
| *CG14207.2* | 4,67E-20 | 0,576145053 | 0,841 | 0,561 | 5,57E-16 | 4 |
| *Arf79F.2* | 5,75E-20 | 0,83502706 | 0,683 | 0,464 | 6,86E-16 | 4 |
| *Hel89B.1* | 6,01E-20 | 1,017992136 | 0,39 | 0,179 | 7,17E-16 | 4 |
| *Fas3* | 6,14E-20 | 1,211196083 | 0,328 | 0,136 | 7,32E-16 | 4 |
| *dl* | 6,40E-20 | 0,811873218 | 0,272 | 0,095 | 7,64E-16 | 4 |
| *EDTP.2* | 6,64E-20 | -1,557572512 | 0,11 | 0,362 | 7,92E-16 | 4 |
| *out* | 6,83E-20 | 0,992575282 | 0,314 | 0,123 | 8,15E-16 | 4 |
| *Gdh.3* | 6,96E-20 | 0,643995752 | 0,634 | 0,36 | 8,30E-16 | 4 |
| *PRL-1.3* | 7,72E-20 | 0,894925249 | 0,51 | 0,278 | 9,21E-16 | 4 |
| *CG10621.3* | 8,33E-20 | -2,063584096 | 0,048 | 0,291 | 9,94E-16 | 4 |
| *pyd.2* | 8,61E-20 | -1,310023195 | 0,359 | 0,589 | 1,03E-15 | 4 |
| *Prosalpha3.2* | 9,68E-20 | 0,75367711 | 0,493 | 0,245 | 1,15E-15 | 4 |
| *cpx.1* | 9,91E-20 | -1,722291186 | 0,11 | 0,357 | 1,18E-15 | 4 |
| *Pp1alpha-96A.1* | 1,00E-19 | -1,273192403 | 0,376 | 0,609 | 1,19E-15 | 4 |
| *Ube3a* | 1,21E-19 | 0,91098821 | 0,259 | 0,092 | 1,44E-15 | 4 |
| *whd.2* | 1,24E-19 | -1,441039794 | 0,3 | 0,539 | 1,48E-15 | 4 |
| *Dhap-at* | 1,56E-19 | 0,812364955 | 0,255 | 0,087 | 1,86E-15 | 4 |
| *Ilp6.2* | 1,57E-19 | -1,667792306 | 0,083 | 0,333 | 1,88E-15 | 4 |
| *CG15096.3* | 1,59E-19 | -1,70056305 | 0,034 | 0,274 | 1,89E-15 | 4 |
| *Mur2B.3* | 1,61E-19 | -1,285962471 | 0,197 | 0,459 | 1,91E-15 | 4 |
| *cic.3* | 1,78E-19 | -1,234492162 | 0,493 | 0,682 | 2,13E-15 | 4 |
| *CG32982* | 2,22E-19 | 0,918990985 | 0,279 | 0,102 | 2,65E-15 | 4 |
| *AkhR.3* | 2,45E-19 | -1,622285504 | 0,097 | 0,342 | 2,92E-15 | 4 |
| *pan.2* | 2,73E-19 | -1,10660911 | 0,362 | 0,596 | 3,26E-15 | 4 |
| *Not1.2* | 2,81E-19 | 0,664127358 | 0,855 | 0,733 | 3,35E-15 | 4 |
| *slmb.2* | 3,18E-19 | 0,949691168 | 0,4 | 0,192 | 3,79E-15 | 4 |
| *Lk6.3* | 3,18E-19 | 0,552382703 | 0,938 | 0,892 | 3,79E-15 | 4 |
| *oaf* | 3,43E-19 | 1,023564682 | 0,314 | 0,129 | 4,09E-15 | 4 |
| *sug.3* | 3,49E-19 | -2,35831113 | 0,103 | 0,347 | 4,16E-15 | 4 |
| *CCT6* | 3,62E-19 | 0,859018925 | 0,303 | 0,12 | 4,32E-15 | 4 |
| *l(2)41Ab.1* | 4,75E-19 | -2,337982587 | 0,141 | 0,388 | 5,67E-15 | 4 |
| *Pfrx.3* | 5,48E-19 | -1,412944146 | 0,076 | 0,319 | 6,54E-15 | 4 |
| *mew.2* | 6,54E-19 | -1,84875433 | 0,172 | 0,411 | 7,80E-15 | 4 |
| *pum.2* | 6,62E-19 | -1,384892393 | 0,328 | 0,559 | 7,90E-15 | 4 |
| *glec.2* | 7,03E-19 | 0,826895054 | 0,597 | 0,338 | 8,39E-15 | 4 |
| *Sc2* | 1,05E-18 | 1,030714119 | 0,283 | 0,109 | 1,25E-14 | 4 |
| *CG34347.3* | 1,05E-18 | -1,866447729 | 0,159 | 0,4 | 1,26E-14 | 4 |
| *CG12163.2* | 1,15E-18 | -1,351144438 | 0,303 | 0,523 | 1,37E-14 | 4 |
| *Dad.2* | 1,21E-18 | 0,825165581 | 0,676 | 0,45 | 1,44E-14 | 4 |
| *PMCA.3* | 1,45E-18 | -0,986926199 | 0,383 | 0,602 | 1,73E-14 | 4 |
| *CG42524.3* | 1,82E-18 | -1,536742695 | 0,259 | 0,492 | 2,17E-14 | 4 |
| *fbp.3* | 1,93E-18 | -1,421747171 | 0,097 | 0,339 | 2,31E-14 | 4 |
| *jim.2* | 2,03E-18 | -1,098951552 | 0,307 | 0,537 | 2,42E-14 | 4 |
| *Eaat1.2* | 2,17E-18 | 0,816829397 | 0,555 | 0,313 | 2,58E-14 | 4 |
| *Smurf* | 2,35E-18 | 0,827742741 | 0,341 | 0,147 | 2,81E-14 | 4 |
| *nrv1.2* | 2,37E-18 | 0,929346182 | 0,445 | 0,23 | 2,82E-14 | 4 |
| *Tsp42Ed.2* | 2,40E-18 | 0,542913947 | 0,7 | 0,41 | 2,86E-14 | 4 |
| *CG5910.3* | 3,08E-18 | -1,363860474 | 0,169 | 0,421 | 3,68E-14 | 4 |
| *mim.3* | 3,51E-18 | -1,249781884 | 0,166 | 0,41 | 4,19E-14 | 4 |
| *UGP.2* | 3,79E-18 | -1,612535189 | 0,214 | 0,443 | 4,52E-14 | 4 |
| *CG4607* | 4,29E-18 | 0,982566314 | 0,414 | 0,206 | 5,11E-14 | 4 |
| *csw.2* | 4,36E-18 | -1,186765094 | 0,241 | 0,488 | 5,20E-14 | 4 |
| *Esyt2* | 4,62E-18 | 0,911727889 | 0,297 | 0,121 | 5,51E-14 | 4 |
| *CG11899.1* | 5,98E-18 | -1,376689025 | 0,09 | 0,327 | 7,13E-14 | 4 |
| *Ubr1* | 8,04E-18 | 0,872987945 | 0,528 | 0,31 | 9,59E-14 | 4 |
| *dnr1.1* | 8,21E-18 | 0,785290395 | 0,448 | 0,225 | 9,80E-14 | 4 |
| *Fur1.1* | 8,33E-18 | 0,703024933 | 0,866 | 0,703 | 9,94E-14 | 4 |
| *eRF3* | 9,09E-18 | 0,791799725 | 0,317 | 0,134 | 1,08E-13 | 4 |
| *caps.2* | 9,12E-18 | 0,715708193 | 0,703 | 0,481 | 1,09E-13 | 4 |
| *Gel.2* | 1,03E-17 | -1,128878776 | 0,245 | 0,483 | 1,22E-13 | 4 |

| | | | | | | |
|---|---|---|---|---|---|---|
| *Kdm2* | 1,27E-17 | 0,899472468 | 0,283 | 0,114 | 1,51E-13 | 4 |
| *CG3597.3* | 1,29E-17 | -1,421279485 | 0,045 | 0,265 | 1,54E-13 | 4 |
| *CG10311* | 1,31E-17 | 0,952763004 | 0,307 | 0,128 | 1,56E-13 | 4 |
| *Gbp3.2* | 1,57E-17 | -1,30588758 | 0,059 | 0,289 | 1,87E-13 | 4 |
| *conu.3* | 2,08E-17 | -1,028728519 | 0,279 | 0,527 | 2,48E-13 | 4 |
| *CG8079.2* | 2,41E-17 | -1,380145874 | 0,083 | 0,312 | 2,87E-13 | 4 |
| *CG6067.2* | 2,51E-17 | -1,388400053 | 0,121 | 0,353 | 2,99E-13 | 4 |
| *CG14762.2* | 2,54E-17 | -1,496264441 | 0,183 | 0,415 | 3,03E-13 | 4 |
| *Gpo1.2* | 3,11E-17 | -1,799453606 | 0,076 | 0,299 | 3,71E-13 | 4 |
| *bel* | 3,43E-17 | 0,699833093 | 0,779 | 0,583 | 4,09E-13 | 4 |
| *CG34133* | 3,60E-17 | 0,781272049 | 0,3 | 0,124 | 4,29E-13 | 4 |
| *retm.3* | 3,65E-17 | -1,442298475 | 0,079 | 0,307 | 4,35E-13 | 4 |
| *pug.1* | 3,72E-17 | -1,062896493 | 0,372 | 0,587 | 4,44E-13 | 4 |
| *sgg.1* | 3,90E-17 | -0,609109344 | 0,893 | 0,954 | 4,66E-13 | 4 |
| *Prosbeta2.1* | 3,97E-17 | 0,758514227 | 0,348 | 0,157 | 4,74E-13 | 4 |
| *cv-2.1* | 4,30E-17 | 0,880626174 | 0,662 | 0,456 | 5,13E-13 | 4 |
| *Dgp-1* | 4,97E-17 | 0,786770494 | 0,252 | 0,093 | 5,92E-13 | 4 |
| *ap.2* | 5,11E-17 | -1,621022619 | 0,134 | 0,364 | 6,10E-13 | 4 |
| *Aldh.1* | 5,21E-17 | 0,881920706 | 0,703 | 0,527 | 6,21E-13 | 4 |
| *Gpdh1.3* | 5,82E-17 | -1,800010857 | 0,162 | 0,383 | 6,95E-13 | 4 |
| *Wnk* | 8,92E-17 | 0,924263684 | 0,372 | 0,179 | 1,06E-12 | 4 |
| *CCHa2.3* | 9,64E-17 | -1,685864417 | 0,062 | 0,283 | 1,15E-12 | 4 |
| *Dmtn.3* | 1,13E-16 | -0,980423114 | 0,614 | 0,763 | 1,34E-12 | 4 |
| *Tsf1.3* | 1,15E-16 | -1,701145085 | 0,093 | 0,314 | 1,38E-12 | 4 |
| *Cdc37* | 1,56E-16 | 0,908963661 | 0,286 | 0,12 | 1,86E-12 | 4 |
| *spoon.3* | 1,75E-16 | -1,317699576 | 0,238 | 0,45 | 2,09E-12 | 4 |
| *EcR.2* | 1,76E-16 | -1,114002858 | 0,259 | 0,492 | 2,09E-12 | 4 |
| *rdx.2* | 2,37E-16 | -0,889758892 | 0,414 | 0,615 | 2,82E-12 | 4 |
| *CG4080.2* | 2,56E-16 | 0,912237716 | 0,462 | 0,253 | 3,06E-12 | 4 |
| *Crtc.2* | 2,59E-16 | -1,034475402 | 0,286 | 0,528 | 3,09E-12 | 4 |
| *ssp7.2* | 2,59E-16 | -1,4155737 | 0,276 | 0,476 | 3,09E-12 | 4 |
| *NTPase* | 2,68E-16 | 0,849445273 | 0,293 | 0,123 | 3,20E-12 | 4 |
| *myo.2* | 3,30E-16 | -1,136286743 | 0,186 | 0,41 | 3,94E-12 | 4 |
| *Jafrac1.2* | 3,35E-16 | 0,763721451 | 0,579 | 0,373 | 3,99E-12 | 4 |
| *Pde11.2* | 3,61E-16 | -0,932565421 | 0,414 | 0,608 | 4,31E-12 | 4 |
| *Moe.1* | 3,78E-16 | 0,82593981 | 0,517 | 0,305 | 4,51E-12 | 4 |
| *vari* | 4,60E-16 | 0,981466308 | 0,321 | 0,145 | 5,49E-12 | 4 |
| *Drp1* | 4,67E-16 | 0,749824813 | 0,283 | 0,116 | 5,58E-12 | 4 |
| *wdb.2* | 6,28E-16 | -0,77611784 | 0,479 | 0,669 | 7,50E-12 | 4 |
| *coro* | 7,04E-16 | 0,701972415 | 0,252 | 0,094 | 8,39E-12 | 4 |
| *CG6701.1* | 7,40E-16 | 0,935316817 | 0,445 | 0,248 | 8,83E-12 | 4 |
| *l(3)80Fj.2* | 8,67E-16 | -0,758467449 | 0,624 | 0,767 | 1,03E-11 | 4 |
| *CG2765.2* | 8,93E-16 | 0,872982806 | 0,417 | 0,223 | 1,06E-11 | 4 |
| *CG6512.1* | 9,57E-16 | 0,69914375 | 0,341 | 0,157 | 1,14E-11 | 4 |
| *AGO2.2* | 9,70E-16 | 0,749788099 | 0,417 | 0,217 | 1,16E-11 | 4 |
| *ps.2* | 1,19E-15 | -0,509557524 | 0,969 | 0,985 | 1,42E-11 | 4 |
| *Shmt.3* | 1,28E-15 | -0,946748082 | 0,445 | 0,631 | 1,53E-11 | 4 |
| *Jra.1* | 1,39E-15 | 0,668545076 | 0,328 | 0,146 | 1,66E-11 | 4 |
| *l(1)G0196.2* | 1,89E-15 | -1,162812249 | 0,238 | 0,453 | 2,25E-11 | 4 |
| *Mad.3* | 2,22E-15 | 0,69853796 | 0,528 | 0,308 | 2,65E-11 | 4 |
| *Nadsyn.2* | 2,41E-15 | -1,230697256 | 0,09 | 0,306 | 2,87E-11 | 4 |
| *spin.2* | 2,70E-15 | 0,562789967 | 0,783 | 0,614 | 3,22E-11 | 4 |
| *D2hgdh.2* | 3,01E-15 | -1,194794499 | 0,11 | 0,325 | 3,59E-11 | 4 |
| *Mef2.1* | 3,30E-15 | -0,951581888 | 0,352 | 0,563 | 3,93E-11 | 4 |
| *S6k.2* | 3,53E-15 | -0,95765544 | 0,293 | 0,499 | 4,21E-11 | 4 |
| *CG7920.3* | 4,45E-15 | -1,286697207 | 0,103 | 0,315 | 5,31E-11 | 4 |
| *uex.2* | 4,84E-15 | -0,833779631 | 0,524 | 0,687 | 5,77E-11 | 4 |
| *crc.2* | 5,80E-15 | 0,725887307 | 0,566 | 0,354 | 6,92E-11 | 4 |
| *Cyp6g1.2* | 6,25E-15 | -1,221910271 | 0,214 | 0,422 | 7,46E-11 | 4 |
| *CG9674.3* | 6,70E-15 | 0,367763341 | 0,883 | 0,608 | 7,99E-11 | 4 |
| *Rala.1* | 8,58E-15 | 0,809896158 | 0,645 | 0,476 | 1,02E-10 | 4 |

| | | | | | | |
|---|---|---|---|---|---|---|
| *NAT1.3* | 8,79E-15 | 0,558058253 | 0,855 | 0,679 | 1,05E-10 | 4 |
| *CHKov1* | 1,03E-14 | 0,884000974 | 0,338 | 0,164 | 1,23E-10 | 4 |
| *CG5355* | 1,19E-14 | 0,728710506 | 0,259 | 0,106 | 1,41E-10 | 4 |
| *Prosbeta6.1* | 1,22E-14 | 0,708713468 | 0,314 | 0,142 | 1,45E-10 | 4 |
| *Hers.3* | 1,36E-14 | 0,555102341 | 0,81 | 0,603 | 1,63E-10 | 4 |
| *RhoGAP18B.1* | 1,39E-14 | 0,687938388 | 0,693 | 0,49 | 1,66E-10 | 4 |
| *spri.2* | 1,47E-14 | -0,969010025 | 0,472 | 0,655 | 1,75E-10 | 4 |
| *AOX3* | 1,51E-14 | 0,876250658 | 0,29 | 0,129 | 1,80E-10 | 4 |
| *CG1648.3* | 1,72E-14 | -1,299372356 | 0,445 | 0,628 | 2,05E-10 | 4 |
| *Khc* | 2,02E-14 | 0,761754726 | 0,338 | 0,16 | 2,42E-10 | 4 |
| *Spt5* | 2,08E-14 | 0,84233842 | 0,262 | 0,111 | 2,48E-10 | 4 |
| *Sodh-1.3* | 2,31E-14 | -1,414065347 | 0,114 | 0,314 | 2,76E-10 | 4 |
| *Ac76E.1* | 2,57E-14 | 0,773352859 | 0,562 | 0,353 | 3,07E-10 | 4 |
| *stas.1* | 2,65E-14 | -1,166442753 | 0,107 | 0,312 | 3,16E-10 | 4 |
| *fus.2* | 2,67E-14 | -1,167869748 | 0,228 | 0,417 | 3,18E-10 | 4 |
| *CG3829.3* | 2,97E-14 | -1,078256711 | 0,134 | 0,344 | 3,54E-10 | 4 |
| *PKD.3* | 4,09E-14 | -1,436265864 | 0,11 | 0,309 | 4,88E-10 | 4 |
| *frma.2* | 4,21E-14 | -1,12148597 | 0,097 | 0,302 | 5,03E-10 | 4 |
| *akirin.1* | 5,51E-14 | 0,674226149 | 0,493 | 0,303 | 6,57E-10 | 4 |
| *DOR.3* | 6,36E-14 | -1,12867505 | 0,455 | 0,624 | 7,58E-10 | 4 |
| *ninaE.2* | 6,74E-14 | 0,668665113 | 0,421 | 0,228 | 8,04E-10 | 4 |
| *Eb1.2* | 6,79E-14 | 0,918580347 | 0,41 | 0,232 | 8,09E-10 | 4 |
| *CtsB1.1* | 7,96E-14 | -1,026592693 | 0,334 | 0,522 | 9,50E-10 | 4 |
| *Idh.3* | 8,27E-14 | -1,263316659 | 0,241 | 0,437 | 9,87E-10 | 4 |
| *Eno.1* | 8,52E-14 | 0,756869935 | 0,669 | 0,482 | 1,02E-09 | 4 |
| *kdn.2* | 9,18E-14 | -1,584110776 | 0,124 | 0,316 | 1,10E-09 | 4 |
| *Dif.2* | 9,74E-14 | -1,222871927 | 0,207 | 0,412 | 1,16E-09 | 4 |
| *E(spl)malpha-BFM.* | 9,85E-14 | -1,109626892 | 0,162 | 0,365 | 1,17E-09 | 4 |
| *SPARC.3* | 1,05E-13 | -1,099140235 | 0,124 | 0,325 | 1,26E-09 | 4 |
| *Rpn1.2* | 1,15E-13 | 0,704191682 | 0,407 | 0,218 | 1,37E-09 | 4 |
| *msn.1* | 1,19E-13 | 0,637017082 | 0,741 | 0,57 | 1,42E-09 | 4 |
| *Vta1* | 1,93E-13 | 0,752276705 | 0,272 | 0,121 | 2,30E-09 | 4 |
| *DPCoAC* | 1,99E-13 | 0,710170024 | 0,328 | 0,16 | 2,37E-09 | 4 |
| *CG3638.3* | 2,30E-13 | -1,049968755 | 0,307 | 0,494 | 2,75E-09 | 4 |
| *RhoGDI* | 2,38E-13 | 0,639153348 | 0,259 | 0,112 | 2,84E-09 | 4 |
| *Cyp6d5.2* | 2,50E-13 | -1,392130962 | 0,176 | 0,364 | 2,98E-09 | 4 |
| *Gbs-76A.3* | 2,75E-13 | 0,618353319 | 0,6 | 0,394 | 3,28E-09 | 4 |
| *CD98hc.2* | 2,81E-13 | 0,726696835 | 0,39 | 0,205 | 3,36E-09 | 4 |
| *CG10600.2* | 3,27E-13 | 0,713240728 | 0,379 | 0,2 | 3,90E-09 | 4 |
| *shot* | 3,28E-13 | 0,41983978 | 0,972 | 0,926 | 3,91E-09 | 4 |
| *Mnt.2* | 3,31E-13 | -0,875651389 | 0,341 | 0,542 | 3,95E-09 | 4 |
| *CG9005.3* | 3,65E-13 | -1,117541298 | 0,193 | 0,388 | 4,35E-09 | 4 |
| *Spn43Ab.2* | 4,21E-13 | -1,113839373 | 0,186 | 0,379 | 5,02E-09 | 4 |
| *MtnA.3* | 4,93E-13 | 0,49056644 | 0,769 | 0,547 | 5,89E-09 | 4 |
| *Irc.3* | 5,81E-13 | 0,650817004 | 0,666 | 0,5 | 6,93E-09 | 4 |
| *teq.3* | 6,18E-13 | -1,140158425 | 0,079 | 0,267 | 7,37E-09 | 4 |
| *Tl.2* | 6,22E-13 | 0,589262328 | 0,755 | 0,575 | 7,42E-09 | 4 |
| *Rbp1-like.2* | 6,82E-13 | -1,156737225 | 0,1 | 0,283 | 8,13E-09 | 4 |
| *hyd.1* | 7,72E-13 | 0,769380661 | 0,345 | 0,18 | 9,21E-09 | 4 |
| *da* | 8,19E-13 | 0,67138893 | 0,386 | 0,21 | 9,76E-09 | 4 |
| *Cul3* | 8,74E-13 | 0,749603315 | 0,31 | 0,155 | 1,04E-08 | 4 |
| *Prosbeta4.2* | 9,69E-13 | 0,596496265 | 0,414 | 0,225 | 1,16E-08 | 4 |
| *psq.2* | 9,89E-13 | -0,91610993 | 0,345 | 0,526 | 1,18E-08 | 4 |
| *IM4.3* | 1,02E-12 | -1,586180416 | 0,134 | 0,324 | 1,22E-08 | 4 |
| *CG8963.1* | 1,34E-12 | 0,699600947 | 0,355 | 0,186 | 1,60E-08 | 4 |
| *PNUTS.1* | 1,50E-12 | 0,662794546 | 0,741 | 0,597 | 1,79E-08 | 4 |
| *gw.2* | 1,51E-12 | -0,774382871 | 0,4 | 0,576 | 1,80E-08 | 4 |
| *Mondo.2* | 1,52E-12 | -0,862968552 | 0,538 | 0,667 | 1,81E-08 | 4 |
| *CG3902.2* | 1,62E-12 | -1,285475543 | 0,138 | 0,325 | 1,94E-08 | 4 |
| *Prosbeta7.2* | 1,79E-12 | 0,699062993 | 0,414 | 0,233 | 2,14E-08 | 4 |
| *l(3)L1231.2* | 2,12E-12 | -0,782093323 | 0,431 | 0,607 | 2,53E-08 | 4 |

| | | | | | | |
|---|---|---|---|---|---|---|
| *Npc2g.2* | 2,36E-12 | -1,087910573 | 0,179 | 0,363 | 2,81E-08 | 4 |
| *nuf.3* | 2,75E-12 | 0,623274381 | 0,697 | 0,533 | 3,28E-08 | 4 |
| *Src64B.2* | 3,19E-12 | 0,495383741 | 0,703 | 0,517 | 3,81E-08 | 4 |
| *fok.1* | 3,27E-12 | -1,070811617 | 0,359 | 0,537 | 3,90E-08 | 4 |
| *jvl.2* | 3,39E-12 | 0,418470334 | 0,862 | 0,738 | 4,05E-08 | 4 |
| *Oda.2* | 3,44E-12 | 0,538816937 | 0,9 | 0,815 | 4,10E-08 | 4 |
| *RpL27A.2* | 3,46E-12 | -0,826044434 | 0,586 | 0,739 | 4,12E-08 | 4 |
| *Rap1.1* | 3,57E-12 | -0,833599364 | 0,317 | 0,515 | 4,26E-08 | 4 |
| *blot.2* | 3,79E-12 | 0,752474939 | 0,572 | 0,38 | 4,52E-08 | 4 |
| *Ldh* | 3,81E-12 | 0,641552528 | 0,286 | 0,133 | 4,54E-08 | 4 |
| *Ubi-p5E.2* | 3,88E-12 | 0,711168838 | 0,428 | 0,252 | 4,63E-08 | 4 |
| *Pi3K92E.1* | 3,94E-12 | 0,698729963 | 0,352 | 0,184 | 4,70E-08 | 4 |
| *CG32486.2* | 4,67E-12 | -0,947752652 | 0,321 | 0,492 | 5,57E-08 | 4 |
| *RhoGAP71E.2* | 5,10E-12 | 0,712811919 | 0,593 | 0,41 | 6,08E-08 | 4 |
| *CdGAPr.1* | 5,22E-12 | 0,619723384 | 0,348 | 0,175 | 6,23E-08 | 4 |
| *CG17691.2* | 5,54E-12 | -0,928330135 | 0,297 | 0,467 | 6,60E-08 | 4 |
| *emb* | 5,65E-12 | 0,631620159 | 0,331 | 0,166 | 6,74E-08 | 4 |
| *bip2.3* | 5,80E-12 | -0,890224426 | 0,307 | 0,488 | 6,92E-08 | 4 |
| *svp.2* | 6,72E-12 | -0,860061464 | 0,376 | 0,543 | 8,01E-08 | 4 |
| *Gprk2.2* | 6,78E-12 | -0,928058515 | 0,614 | 0,705 | 8,09E-08 | 4 |
| *cv-c.2* | 6,90E-12 | 0,386311898 | 0,852 | 0,63 | 8,24E-08 | 4 |
| *CG34136.2* | 7,62E-12 | 0,601512721 | 0,441 | 0,256 | 9,09E-08 | 4 |
| *Invadolysin.3* | 8,04E-12 | -1,902690259 | 0,117 | 0,289 | 9,59E-08 | 4 |
| *RpS27A.2* | 9,17E-12 | -0,767351536 | 0,517 | 0,668 | 1,09E-07 | 4 |
| *CG17124.3* | 9,64E-12 | -0,986319213 | 0,69 | 0,755 | 1,15E-07 | 4 |
| *Atf3.2* | 9,81E-12 | 0,567444898 | 0,472 | 0,276 | 1,17E-07 | 4 |
| *RpLP2.2* | 9,93E-12 | -0,835287113 | 0,517 | 0,669 | 1,18E-07 | 4 |
| *fon.3* | 1,15E-11 | -1,220840856 | 0,152 | 0,329 | 1,37E-07 | 4 |
| *smg.1* | 1,28E-11 | -1,050755556 | 0,134 | 0,314 | 1,53E-07 | 4 |
| *Csk.2* | 1,29E-11 | -0,96945402 | 0,324 | 0,496 | 1,53E-07 | 4 |
| *cwo.2* | 1,30E-11 | -0,652938721 | 0,862 | 0,881 | 1,55E-07 | 4 |
| *IM33.3* | 1,44E-11 | -1,181230775 | 0,193 | 0,386 | 1,71E-07 | 4 |
| *Obp99c.2* | 1,44E-11 | -0,762361701 | 0,652 | 0,763 | 1,72E-07 | 4 |
| *Oatp30B.3* | 1,49E-11 | -0,839412544 | 0,445 | 0,601 | 1,78E-07 | 4 |
| *Indy.1* | 1,64E-11 | -0,881878626 | 0,514 | 0,648 | 1,96E-07 | 4 |
| *CG11089.2* | 1,68E-11 | 0,339108638 | 0,872 | 0,639 | 2,01E-07 | 4 |
| *flfl.3* | 1,72E-11 | -1,006581755 | 0,138 | 0,316 | 2,05E-07 | 4 |
| *CG7668* | 1,85E-11 | 0,589313994 | 0,29 | 0,141 | 2,21E-07 | 4 |
| *iPLA2-VIA.1* | 1,94E-11 | 0,590021027 | 0,328 | 0,168 | 2,31E-07 | 4 |
| *CG17734.2* | 2,28E-11 | -0,938751446 | 0,097 | 0,27 | 2,72E-07 | 4 |
| *Tet.2* | 2,32E-11 | -0,816249751 | 0,39 | 0,584 | 2,77E-07 | 4 |
| *hdc.1* | 2,36E-11 | 0,576509083 | 0,341 | 0,179 | 2,81E-07 | 4 |
| *RpS2.1* | 2,56E-11 | -0,797473813 | 0,379 | 0,558 | 3,05E-07 | 4 |
| *Doa.3* | 2,65E-11 | -0,762456928 | 0,366 | 0,536 | 3,17E-07 | 4 |
| *eEF1alpha1.2* | 2,77E-11 | 0,363172631 | 0,955 | 0,905 | 3,31E-07 | 4 |
| *Rbfox1.2* | 2,85E-11 | -0,874034068 | 0,362 | 0,531 | 3,41E-07 | 4 |
| *CG17508.1* | 3,30E-11 | -0,939088306 | 0,148 | 0,324 | 3,94E-07 | 4 |
| *kst.3* | 3,34E-11 | 0,562341385 | 0,452 | 0,271 | 3,99E-07 | 4 |
| *Mbs.1* | 3,50E-11 | 0,578846928 | 0,497 | 0,313 | 4,17E-07 | 4 |
| *CG3036.3* | 3,52E-11 | -1,302246998 | 0,61 | 0,655 | 4,20E-07 | 4 |
| *CG34417.2* | 3,58E-11 | 0,37670616 | 0,779 | 0,599 | 4,28E-07 | 4 |
| *nmo.2* | 4,48E-11 | 0,434138881 | 0,752 | 0,559 | 5,35E-07 | 4 |
| *Gale.3* | 4,51E-11 | -1,083327309 | 0,09 | 0,253 | 5,38E-07 | 4 |
| *Rpn3.2* | 4,72E-11 | 0,556318253 | 0,352 | 0,188 | 5,63E-07 | 4 |
| *Nc73EF.2* | 5,04E-11 | -0,891208053 | 0,569 | 0,662 | 6,01E-07 | 4 |
| *CG31145.2* | 5,11E-11 | -0,679435415 | 0,655 | 0,742 | 6,10E-07 | 4 |
| *Sh3beta.2* | 5,39E-11 | 0,703839227 | 0,352 | 0,193 | 6,43E-07 | 4 |
| *CG14073.2* | 5,89E-11 | 0,690022605 | 0,541 | 0,384 | 7,02E-07 | 4 |
| *RpL18.2* | 6,03E-11 | -0,83783175 | 0,414 | 0,56 | 7,19E-07 | 4 |
| *goe.3* | 6,19E-11 | -1,107302262 | 0,11 | 0,287 | 7,38E-07 | 4 |
| *app.3* | 6,51E-11 | -1,036459444 | 0,234 | 0,405 | 7,76E-07 | 4 |

| | | | | | | |
|---|---|---|---|---|---|---|
| *Atg18b.2* | 8,38E-11 | 0,443550703 | 0,628 | 0,425 | 1,00E-06 | 4 |
| *CG31635.3* | 8,90E-11 | -0,849959184 | 0,397 | 0,567 | 1,06E-06 | 4 |
| *GstE12.3* | 9,08E-11 | -1,036902661 | 0,107 | 0,268 | 1,08E-06 | 4 |
| *sima.2* | 1,02E-10 | -0,528423658 | 0,893 | 0,93 | 1,22E-06 | 4 |
| *Ran* | 1,18E-10 | 0,702606622 | 0,403 | 0,241 | 1,41E-06 | 4 |
| *CG9691.2* | 1,19E-10 | -0,97921891 | 0,183 | 0,359 | 1,42E-06 | 4 |
| *IP3K1.3* | 1,24E-10 | 0,4280674 | 0,741 | 0,54 | 1,48E-06 | 4 |
| *alpha-Man-Ia.2* | 1,29E-10 | -0,831490654 | 0,193 | 0,377 | 1,54E-06 | 4 |
| *Rab14.1* | 1,58E-10 | 0,560590882 | 0,369 | 0,203 | 1,88E-06 | 4 |
| *Pdk.1* | 1,63E-10 | 0,406563073 | 0,924 | 0,856 | 1,95E-06 | 4 |
| *AGO1.1* | 1,73E-10 | -0,654311056 | 0,462 | 0,627 | 2,07E-06 | 4 |
| *step.2* | 1,92E-10 | 0,588620831 | 0,593 | 0,42 | 2,29E-06 | 4 |
| *ATPCL.2* | 1,96E-10 | -1,075371591 | 0,259 | 0,426 | 2,33E-06 | 4 |
| *Marf.2* | 2,20E-10 | -0,859111319 | 0,159 | 0,327 | 2,62E-06 | 4 |
| *RpS29.3* | 2,22E-10 | -0,907890595 | 0,438 | 0,588 | 2,65E-06 | 4 |
| *ALiX.2* | 2,24E-10 | 0,548523792 | 0,372 | 0,21 | 2,67E-06 | 4 |
| *Gp150.1* | 2,26E-10 | -0,753806046 | 0,359 | 0,541 | 2,70E-06 | 4 |
| *RpLP1.3* | 2,34E-10 | -0,806658151 | 0,49 | 0,644 | 2,79E-06 | 4 |
| *emc* | 2,46E-10 | 0,667623367 | 0,334 | 0,184 | 2,93E-06 | 4 |
| *Nuak1.2* | 2,48E-10 | 0,620772212 | 0,428 | 0,265 | 2,96E-06 | 4 |
| *RpS7.2* | 2,52E-10 | -0,761705533 | 0,562 | 0,692 | 3,00E-06 | 4 |
| *Prosalpha5.2* | 2,61E-10 | 0,633587923 | 0,328 | 0,178 | 3,12E-06 | 4 |
| *Chmp1.2* | 2,69E-10 | 0,558507706 | 0,369 | 0,206 | 3,20E-06 | 4 |
| *CG14154.2* | 2,77E-10 | -0,832727829 | 0,317 | 0,477 | 3,30E-06 | 4 |
| *CycG.2* | 2,82E-10 | -0,4623922 | 0,921 | 0,95 | 3,37E-06 | 4 |
| *Prosalpha2.2* | 3,31E-10 | 0,768570767 | 0,366 | 0,214 | 3,94E-06 | 4 |
| *Mitf.2* | 3,33E-10 | -0,807871556 | 0,141 | 0,313 | 3,98E-06 | 4 |
| *Cp1.1* | 3,66E-10 | -0,809228452 | 0,703 | 0,787 | 4,36E-06 | 4 |
| *UK114.2* | 4,02E-10 | -0,886407203 | 0,266 | 0,436 | 4,79E-06 | 4 |
| *mura.2* | 4,03E-10 | -0,768629263 | 0,352 | 0,517 | 4,81E-06 | 4 |
| *Galphai* | 4,18E-10 | 0,640000649 | 0,462 | 0,297 | 4,99E-06 | 4 |
| *Rpt3.2* | 4,19E-10 | 0,626805258 | 0,345 | 0,191 | 4,99E-06 | 4 |
| *Ank.2* | 5,09E-10 | -0,676112277 | 0,331 | 0,507 | 6,08E-06 | 4 |
| *Naprt.3* | 5,74E-10 | 0,799093309 | 0,507 | 0,35 | 6,85E-06 | 4 |
| *Cdep.3* | 5,88E-10 | -0,953504524 | 0,221 | 0,381 | 7,01E-06 | 4 |
| *Rpt6.2* | 6,44E-10 | 0,601720367 | 0,334 | 0,186 | 7,68E-06 | 4 |
| *CG3376.1* | 6,49E-10 | -0,743716437 | 0,445 | 0,588 | 7,74E-06 | 4 |
| *CG33969.1* | 6,86E-10 | 0,421547626 | 0,269 | 0,133 | 8,18E-06 | 4 |
| *GstT4.1* | 7,11E-10 | -1,288576967 | 0,2 | 0,355 | 8,48E-06 | 4 |
| *Stat92E.3* | 7,66E-10 | -1,108913145 | 0,483 | 0,6 | 9,14E-06 | 4 |
| *ens.2* | 7,78E-10 | 0,52749827 | 0,641 | 0,474 | 9,28E-06 | 4 |
| *kra.2* | 7,88E-10 | 0,513641942 | 0,679 | 0,534 | 9,40E-06 | 4 |
| *velo* | 8,17E-10 | 0,585626829 | 0,297 | 0,157 | 9,75E-06 | 4 |
| *RpL10Ab.2* | 8,56E-10 | -0,720643961 | 0,448 | 0,602 | 1,02E-05 | 4 |
| *CG8034.3* | 8,70E-10 | -1,104607796 | 0,507 | 0,615 | 1,04E-05 | 4 |
| *eIF3c.3* | 8,72E-10 | 0,521634225 | 0,362 | 0,206 | 1,04E-05 | 4 |
| *Mes2* | 9,01E-10 | 0,679814608 | 0,3 | 0,162 | 1,07E-05 | 4 |
| *Idgf4.2* | 9,40E-10 | 0,624847664 | 0,434 | 0,274 | 1,12E-05 | 4 |
| *Hn.2* | 1,00E-09 | -0,776034264 | 0,245 | 0,417 | 1,20E-05 | 4 |
| *AGO3.2* | 1,05E-09 | -0,983364873 | 0,093 | 0,251 | 1,25E-05 | 4 |
| *CG44014.1* | 1,08E-09 | 0,623680347 | 0,307 | 0,165 | 1,28E-05 | 4 |
| *CASK.2* | 1,10E-09 | -0,846928489 | 0,183 | 0,344 | 1,31E-05 | 4 |
| *unc-13.2* | 1,17E-09 | -0,890547347 | 0,321 | 0,474 | 1,40E-05 | 4 |
| *osp.3* | 1,23E-09 | -1,02847964 | 0,169 | 0,333 | 1,47E-05 | 4 |
| *CG42324.2* | 1,42E-09 | -0,64015887 | 0,5 | 0,654 | 1,69E-05 | 4 |
| *Baldspot* | 1,49E-09 | 0,662794541 | 0,262 | 0,135 | 1,77E-05 | 4 |
| *CG5958.3* | 1,58E-09 | 0,399522563 | 0,586 | 0,395 | 1,88E-05 | 4 |
| *RhoGAP19D.3* | 1,61E-09 | -1,070984413 | 0,362 | 0,494 | 1,92E-05 | 4 |
| *Gug.3* | 1,62E-09 | -0,602833887 | 0,514 | 0,655 | 1,93E-05 | 4 |
| *sqh.1* | 1,63E-09 | 0,562560927 | 0,29 | 0,151 | 1,95E-05 | 4 |
| *CHES-1-like.2* | 1,64E-09 | -0,773950488 | 0,341 | 0,497 | 1,96E-05 | 4 |

| | | | | | | |
|---|---|---|---|---|---|---|
| *tna* | 1,66E-09 | 0,508595829 | 0,745 | 0,593 | 1,98E-05 | 4 |
| *Slmap* | 1,71E-09 | 0,538227845 | 0,283 | 0,147 | 2,04E-05 | 4 |
| *Myc.2* | 1,72E-09 | 0,518239077 | 0,772 | 0,647 | 2,05E-05 | 4 |
| *aop.1* | 1,74E-09 | 0,453320003 | 0,731 | 0,578 | 2,07E-05 | 4 |
| *for.2* | 1,83E-09 | -0,567129691 | 0,617 | 0,749 | 2,19E-05 | 4 |
| *Prosbeta5.2* | 1,94E-09 | 0,535109378 | 0,355 | 0,202 | 2,31E-05 | 4 |
| *Cyp1.2* | 2,01E-09 | 0,557901914 | 0,538 | 0,377 | 2,40E-05 | 4 |
| *CG5059.1* | 2,01E-09 | -0,590948311 | 0,6 | 0,703 | 2,40E-05 | 4 |
| *Prosalpha4.1* | 2,35E-09 | 0,537997769 | 0,29 | 0,152 | 2,80E-05 | 4 |
| *Dbp80.2* | 2,55E-09 | -0,525549864 | 0,514 | 0,672 | 3,04E-05 | 4 |
| *Unr.2* | 2,66E-09 | 0,491792044 | 0,8 | 0,728 | 3,17E-05 | 4 |
| *betaTub97EF.1* | 2,72E-09 | 0,602062212 | 0,428 | 0,266 | 3,24E-05 | 4 |
| *Ac13E.2* | 2,77E-09 | -1,076596912 | 0,124 | 0,274 | 3,31E-05 | 4 |
| *RpS18.2* | 2,84E-09 | -0,673177203 | 0,614 | 0,733 | 3,39E-05 | 4 |
| *nkd.2* | 2,92E-09 | 0,457708992 | 0,614 | 0,431 | 3,49E-05 | 4 |
| *Prosbeta1.1* | 3,13E-09 | 0,511076634 | 0,29 | 0,154 | 3,73E-05 | 4 |
| *UbcE2H.3* | 3,15E-09 | -0,782400006 | 0,207 | 0,373 | 3,75E-05 | 4 |
| *Cbs.3* | 3,53E-09 | -0,994139675 | 0,124 | 0,272 | 4,21E-05 | 4 |
| *spen* | 3,61E-09 | 0,373634374 | 0,979 | 0,958 | 4,31E-05 | 4 |
| *zip* | 3,63E-09 | 0,733299116 | 0,31 | 0,182 | 4,33E-05 | 4 |
| *Atg8a.2* | 3,95E-09 | 0,414343499 | 0,807 | 0,683 | 4,72E-05 | 4 |
| *Vinc.1* | 4,01E-09 | 0,505912303 | 0,314 | 0,17 | 4,78E-05 | 4 |
| *mbc.2* | 4,60E-09 | 0,717964251 | 0,397 | 0,247 | 5,49E-05 | 4 |
| *Sap-r.2* | 5,25E-09 | 0,399867437 | 0,683 | 0,54 | 6,27E-05 | 4 |
| *DppIII.2* | 5,39E-09 | 0,528993852 | 0,321 | 0,181 | 6,43E-05 | 4 |
| *SCaMC.3* | 5,67E-09 | -0,524458861 | 0,683 | 0,773 | 6,76E-05 | 4 |
| *CG31729* | 5,77E-09 | 0,634867055 | 0,269 | 0,147 | 6,88E-05 | 4 |
| *Pli.2* | 6,38E-09 | -1,061904749 | 0,472 | 0,574 | 7,61E-05 | 4 |
| *CG10680.3* | 6,46E-09 | -0,814049431 | 0,279 | 0,431 | 7,71E-05 | 4 |
| *lost.2* | 9,43E-09 | 0,577260086 | 0,531 | 0,378 | 0,000112537 | 4 |
| *sm.3* | 9,66E-09 | -0,952490145 | 0,176 | 0,316 | 0,000115205 | 4 |
| *nej.2* | 1,04E-08 | -0,686187831 | 0,393 | 0,526 | 0,000124583 | 4 |
| *MCPH1.2* | 1,09E-08 | 0,500458237 | 0,462 | 0,3 | 0,000129876 | 4 |
| *cac.2* | 1,09E-08 | -1,450912344 | 0,334 | 0,467 | 0,000130176 | 4 |
| *CG43402* | 1,24E-08 | 0,468409403 | 0,345 | 0,198 | 0,000147531 | 4 |
| *srp.2* | 1,32E-08 | -0,69390374 | 0,548 | 0,64 | 0,000156948 | 4 |
| *RpL36.2* | 1,42E-08 | -0,700213548 | 0,5 | 0,618 | 0,000169858 | 4 |
| *Tis11.1* | 1,47E-08 | -0,517045231 | 0,862 | 0,916 | 0,00017554 | 4 |
| *BomS3.2* | 1,62E-08 | -1,090006674 | 0,297 | 0,447 | 0,000192908 | 4 |
| *Idgf3* | 1,67E-08 | 0,520672347 | 0,286 | 0,154 | 0,000199288 | 4 |
| *Got2.3* | 1,70E-08 | -0,707087371 | 0,431 | 0,562 | 0,000202588 | 4 |
| *CaMKII.2* | 1,81E-08 | -0,902222495 | 0,19 | 0,329 | 0,000215454 | 4 |
| *Jupiter* | 2,08E-08 | 0,481014963 | 0,283 | 0,156 | 0,000247771 | 4 |
| *Wbp2.2* | 2,19E-08 | 0,509485518 | 0,479 | 0,321 | 0,000261632 | 4 |
| *Scsalpha1.1* | 2,89E-08 | -0,835904168 | 0,276 | 0,406 | 0,000345298 | 4 |
| *scrib.2* | 3,09E-08 | -0,788171209 | 0,245 | 0,39 | 0,000368221 | 4 |
| *sws.2* | 3,13E-08 | -0,845366198 | 0,159 | 0,299 | 0,000372979 | 4 |
| *Golgin245.2* | 3,22E-08 | 0,523086382 | 0,345 | 0,208 | 0,000384662 | 4 |
| *RpL7A.3* | 3,26E-08 | -0,598902911 | 0,507 | 0,635 | 0,000389006 | 4 |
| *CG9281* | 3,39E-08 | 0,556962541 | 0,266 | 0,143 | 0,000404553 | 4 |
| *Opa1.1* | 3,76E-08 | 0,50322163 | 0,366 | 0,224 | 0,000448397 | 4 |
| *Oatp74D.2* | 3,83E-08 | -0,628395742 | 0,217 | 0,376 | 0,000456665 | 4 |
| *CG1468.1* | 4,01E-08 | -0,737683056 | 0,21 | 0,36 | 0,000478151 | 4 |
| *Mlf.1* | 4,10E-08 | 0,622873571 | 0,286 | 0,164 | 0,000489619 | 4 |
| *CG8369.1* | 4,26E-08 | 0,415560992 | 0,29 | 0,162 | 0,000508676 | 4 |
| *Pdi* | 4,32E-08 | 0,455627719 | 0,383 | 0,24 | 0,000515628 | 4 |
| *Rpn10.2* | 4,54E-08 | 0,489082955 | 0,383 | 0,235 | 0,000541297 | 4 |
| *Eip93F* | 4,55E-08 | 0,307320668 | 0,993 | 0,987 | 0,000542929 | 4 |
| *l(3)05822.2* | 4,56E-08 | 0,580497097 | 0,331 | 0,198 | 0,000544525 | 4 |
| *CG8485.3* | 4,84E-08 | -1,077846181 | 0,128 | 0,259 | 0,000577098 | 4 |
| *corto.2* | 5,07E-08 | -0,790353296 | 0,359 | 0,5 | 0,000605038 | 4 |

| | | | | | | |
|---|---|---|---|---|---|---|
| *RpL27.3* | 5,57E-08 | -0,787553641 | 0,524 | 0,617 | 0,000664701 | 4 |
| *Pdk1.2* | 5,58E-08 | 0,345202214 | 0,831 | 0,722 | 0,000665078 | 4 |
| *CG2201.1* | 5,81E-08 | -0,75265796 | 0,234 | 0,372 | 0,000692695 | 4 |
| *CG33158* | 5,90E-08 | 0,731566716 | 0,307 | 0,187 | 0,00070351 | 4 |
| *spir.2* | 5,91E-08 | 0,50203934 | 0,507 | 0,357 | 0,000705547 | 4 |
| *RpS3.1* | 5,93E-08 | -0,697438631 | 0,376 | 0,504 | 0,000707269 | 4 |
| *PRAS40.2* | 5,99E-08 | -0,935770758 | 0,162 | 0,302 | 0,000714595 | 4 |
| *mts.2* | 6,39E-08 | 0,512047737 | 0,51 | 0,357 | 0,000762091 | 4 |
| *tok* | 6,52E-08 | -0,869105546 | 0,293 | 0,433 | 0,000778061 | 4 |
| *Prosalpha7.2* | 6,55E-08 | 0,532605182 | 0,341 | 0,206 | 0,000781376 | 4 |
| *CG6707* | 6,87E-08 | 0,631073498 | 0,283 | 0,164 | 0,000819659 | 4 |
| *tara.2* | 6,97E-08 | -0,700555175 | 0,255 | 0,399 | 0,000831608 | 4 |
| *RpL35A.2* | 6,98E-08 | -0,717977666 | 0,448 | 0,566 | 0,000832276 | 4 |
| *RpL18A.3* | 7,45E-08 | -0,774527156 | 0,534 | 0,629 | 0,000888727 | 4 |
| *aralar1.2* | 7,82E-08 | 0,361808847 | 0,762 | 0,622 | 0,000933304 | 4 |
| *CG15293.3* | 8,01E-08 | -0,781129334 | 0,324 | 0,459 | 0,000955816 | 4 |
| *Rpt2.1* | 8,08E-08 | 0,452019618 | 0,272 | 0,15 | 0,000964046 | 4 |
| *Atg17.2* | 8,19E-08 | 0,492132544 | 0,607 | 0,471 | 0,000977034 | 4 |
| *CG7530.2* | 8,39E-08 | -0,837709758 | 0,176 | 0,315 | 0,001001129 | 4 |
| *RpS11.2* | 8,41E-08 | -0,619648837 | 0,5 | 0,64 | 0,001003613 | 4 |
| *sqd.2* | 8,48E-08 | -0,448594952 | 0,79 | 0,837 | 0,001011718 | 4 |
| *CG16721.1* | 8,68E-08 | 0,648897042 | 0,321 | 0,199 | 0,001035274 | 4 |
| *Rack1.1* | 9,01E-08 | -0,746452437 | 0,376 | 0,493 | 0,001074809 | 4 |
| *CG4629.3* | 9,20E-08 | -1,040732089 | 0,131 | 0,265 | 0,001097982 | 4 |
| *Akap200.3* | 9,34E-08 | -0,884603286 | 0,445 | 0,565 | 0,001113868 | 4 |
| *RpS15.3* | 9,45E-08 | -0,625021329 | 0,576 | 0,691 | 0,001127732 | 4 |
| *CG12065.2* | 9,63E-08 | 0,469224634 | 0,428 | 0,282 | 0,00114863 | 4 |
| *pcs.3* | 9,72E-08 | -1,122519543 | 0,31 | 0,436 | 0,001159008 | 4 |
| *CG12054.2* | 1,04E-07 | -0,705156051 | 0,266 | 0,401 | 0,001234759 | 4 |
| *RpS15Aa.3* | 1,17E-07 | -0,584153229 | 0,359 | 0,5 | 0,001400084 | 4 |
| *Cals.2* | 1,23E-07 | -0,711706361 | 0,148 | 0,29 | 0,001464177 | 4 |
| *Cys.3* | 1,26E-07 | 0,598547813 | 0,421 | 0,283 | 0,001497294 | 4 |
| *CG8086.2* | 1,26E-07 | 0,504109692 | 0,459 | 0,311 | 0,00150022 | 4 |
| *sd* | 1,28E-07 | 0,552010837 | 0,279 | 0,158 | 0,001527738 | 4 |
| *Hsf* | 1,31E-07 | 0,629691174 | 0,255 | 0,14 | 0,001562446 | 4 |
| *RpL12.2* | 1,52E-07 | -0,577226985 | 0,476 | 0,607 | 0,001814347 | 4 |
| *stx.2* | 1,65E-07 | -0,861081756 | 0,276 | 0,408 | 0,001972051 | 4 |
| *Sar1.1* | 1,76E-07 | -0,717957029 | 0,217 | 0,357 | 0,002101564 | 4 |
| *Tnpo* | 1,80E-07 | 0,490322247 | 0,372 | 0,239 | 0,002145889 | 4 |
| *Irp-1B.2* | 2,00E-07 | -0,898168578 | 0,166 | 0,292 | 0,002380781 | 4 |
| *RpS4.3* | 2,09E-07 | -0,630360433 | 0,455 | 0,588 | 0,002491298 | 4 |
| *sowah.2* | 2,25E-07 | 0,465127458 | 0,459 | 0,315 | 0,002679744 | 4 |
| *E(Pc).1* | 2,25E-07 | 0,568909201 | 0,386 | 0,254 | 0,002685472 | 4 |
| *Kr-h1.2* | 2,29E-07 | 0,437546354 | 0,61 | 0,456 | 0,002729609 | 4 |
| *Pomp.1* | 2,56E-07 | 0,584345396 | 0,266 | 0,151 | 0,003056958 | 4 |
| *REPTOR.2* | 2,63E-07 | 0,369231225 | 0,834 | 0,706 | 0,003141737 | 4 |
| *AdamTS-A.3* | 2,75E-07 | -1,456775795 | 0,159 | 0,283 | 0,003277526 | 4 |
| *RpS8.2* | 2,84E-07 | -0,57118488 | 0,614 | 0,723 | 0,003392574 | 4 |
| *CG30015.2* | 2,90E-07 | 0,288895664 | 0,883 | 0,779 | 0,003455047 | 4 |
| *Pka-C1.2* | 2,93E-07 | 0,39612251 | 0,779 | 0,656 | 0,003494459 | 4 |
| *p47.1* | 3,01E-07 | 0,504332963 | 0,259 | 0,145 | 0,003593848 | 4 |
| *RpS26.1* | 3,17E-07 | -0,704974332 | 0,441 | 0,556 | 0,003782958 | 4 |
| *RpL6.1* | 3,68E-07 | -0,55857573 | 0,476 | 0,594 | 0,004393451 | 4 |
| *ctrip.1* | 3,82E-07 | 0,488478915 | 0,503 | 0,364 | 0,004560354 | 4 |
| *Pmp70.3* | 3,87E-07 | -0,951166519 | 0,159 | 0,282 | 0,004616145 | 4 |
| *S.1* | 4,07E-07 | 0,751284006 | 0,369 | 0,242 | 0,004856369 | 4 |
| *Rpn11.1* | 4,29E-07 | 0,529819544 | 0,283 | 0,166 | 0,00511287 | 4 |
| *RpL7.2* | 4,49E-07 | -0,626418158 | 0,531 | 0,614 | 0,005353036 | 4 |
| *Wdr62.2* | 4,64E-07 | -0,733830854 | 0,545 | 0,628 | 0,005533844 | 4 |
| *Spn42Da* | 4,90E-07 | 0,65994545 | 0,255 | 0,149 | 0,005847122 | 4 |
| *CG12116.3* | 5,25E-07 | -0,808152803 | 0,307 | 0,423 | 0,006260264 | 4 |

| | | | | | | |
|---|---|---|---|---|---|---|
| *RpL37A.3* | 5,91E-07 | -0,643794439 | 0,372 | 0,499 | 0,007050217 | 4 |
| *lap.2* | 5,97E-07 | -0,562772295 | 0,183 | 0,318 | 0,007126838 | 4 |
| *Rab7.2* | 6,02E-07 | 0,407696832 | 0,428 | 0,286 | 0,007186044 | 4 |
| *ftz-f1.3* | 6,12E-07 | -0,576220508 | 0,538 | 0,623 | 0,007304398 | 4 |
| *Ugt301D1* | 6,32E-07 | 0,540660501 | 0,29 | 0,17 | 0,007534268 | 4 |
| *Stam.2* | 6,74E-07 | 0,492220003 | 0,4 | 0,262 | 0,00804213 | 4 |
| *shn.1* | 7,00E-07 | 0,383209844 | 0,724 | 0,606 | 0,008344842 | 4 |
| *CG1640.2* | 7,20E-07 | -0,803919205 | 0,155 | 0,28 | 0,008589996 | 4 |
| *cv-d* | 7,28E-07 | 0,600115888 | 0,293 | 0,176 | 0,008679419 | 4 |
| *sfl.2* | 7,72E-07 | -0,684425026 | 0,424 | 0,522 | 0,009209336 | 4 |
| *Rpn13.2* | 8,01E-07 | 0,438728003 | 0,4 | 0,262 | 0,009557112 | 4 |
| *CG32767.3* | 8,70E-07 | -0,630215627 | 0,276 | 0,398 | 0,010381604 | 4 |
| *Npl4.3* | 9,31E-07 | 0,443683948 | 0,331 | 0,204 | 0,011110664 | 4 |
| *CG8547.1* | 9,69E-07 | 0,449233053 | 0,276 | 0,161 | 0,011557482 | 4 |
| *Tango1.2* | 9,70E-07 | 0,478623846 | 0,324 | 0,202 | 0,011568155 | 4 |
| *mask* | 1,03E-06 | 0,502568788 | 0,648 | 0,55 | 0,012256784 | 4 |
| *PyK.2* | 1,04E-06 | -0,707191973 | 0,19 | 0,315 | 0,012397257 | 4 |
| *CG17549.2* | 1,06E-06 | -0,652528105 | 0,252 | 0,385 | 0,012682471 | 4 |
| *CG32425.2* | 1,07E-06 | -0,897931685 | 0,224 | 0,343 | 0,012759873 | 4 |
| *BicD* | 1,08E-06 | 0,440983624 | 0,314 | 0,194 | 0,012879937 | 4 |
| *RpL8.2* | 1,09E-06 | -0,575596029 | 0,583 | 0,676 | 0,012985064 | 4 |
| *Mdh2.1* | 1,18E-06 | -0,550992995 | 0,424 | 0,553 | 0,014022344 | 4 |
| *CG31523.2* | 1,22E-06 | 0,451137586 | 0,386 | 0,251 | 0,014528001 | 4 |
| *CAP.1* | 1,33E-06 | 0,469300891 | 0,434 | 0,301 | 0,015881639 | 4 |
| *Rpn8.2* | 1,33E-06 | 0,461159944 | 0,293 | 0,177 | 0,015919151 | 4 |
| *RpS14a.1* | 1,36E-06 | -0,65220477 | 0,276 | 0,398 | 0,016232682 | 4 |
| *CG5853.1* | 1,37E-06 | -1,002960533 | 0,19 | 0,308 | 0,016333318 | 4 |
| *Stlk.3* | 1,45E-06 | -0,648260273 | 0,179 | 0,307 | 0,017245648 | 4 |
| *Sem1.2* | 1,46E-06 | 0,540103988 | 0,321 | 0,206 | 0,017404027 | 4 |
| *red.2* | 1,56E-06 | 0,342919176 | 0,317 | 0,194 | 0,018608757 | 4 |
| *dally.2* | 1,58E-06 | -0,718038057 | 0,152 | 0,28 | 0,018894362 | 4 |
| *Gbp2* | 1,62E-06 | -0,643982931 | 0,197 | 0,323 | 0,019345819 | 4 |
| *CG16926.2* | 1,67E-06 | -0,629402116 | 0,548 | 0,63 | 0,019925213 | 4 |
| *CG6330.3* | 1,72E-06 | 0,453374681 | 0,421 | 0,283 | 0,020543709 | 4 |
| *RpL28.2* | 1,79E-06 | -0,478560444 | 0,61 | 0,74 | 0,021370791 | 4 |
| *Col4a1.2* | 1,88E-06 | 0,561923406 | 0,331 | 0,219 | 0,022385079 | 4 |
| *RpL14.2* | 1,99E-06 | -0,565871264 | 0,597 | 0,678 | 0,023717095 | 4 |
| *CG13887.1* | 1,99E-06 | 0,566810235 | 0,255 | 0,153 | 0,023741009 | 4 |
| *Nup153.1* | 2,05E-06 | 0,49175029 | 0,434 | 0,307 | 0,024479774 | 4 |
| *CG10365* | 2,16E-06 | -0,750895992 | 0,134 | 0,251 | 0,025788104 | 4 |
| *Msp300.2* | 2,17E-06 | -0,443337377 | 0,869 | 0,883 | 0,025858788 | 4 |
| *RpL24.2* | 2,34E-06 | -0,63848909 | 0,507 | 0,615 | 0,027868596 | 4 |
| *SNF4Agamma.2* | 2,36E-06 | -0,398010874 | 0,866 | 0,894 | 0,02820106 | 4 |
| *Ubqn.2* | 2,61E-06 | 0,457557658 | 0,366 | 0,245 | 0,031142745 | 4 |
| *RpS12.2* | 2,86E-06 | -0,596967682 | 0,514 | 0,608 | 0,034151808 | 4 |
| *CG44774.2* | 2,99E-06 | 0,551125974 | 0,366 | 0,245 | 0,035715493 | 4 |
| *RpL37a.2* | 3,13E-06 | -0,637688556 | 0,503 | 0,596 | 0,037332077 | 4 |
| *Myd88.1* | 3,18E-06 | 0,491179482 | 0,479 | 0,351 | 0,037922228 | 4 |
| *CG3726* | 3,47E-06 | -1,096260081 | 0,19 | 0,304 | 0,041345776 | 4 |
| *Gprk1.2* | 3,52E-06 | -0,713810397 | 0,248 | 0,357 | 0,041951508 | 4 |
| *CG1703* | 3,53E-06 | 0,488147452 | 0,283 | 0,174 | 0,042104402 | 4 |
| *Gbeta13F.1* | 3,56E-06 | -0,503264616 | 0,424 | 0,541 | 0,042448855 | 4 |
| *mop.1* | 3,60E-06 | 0,487464381 | 0,283 | 0,175 | 0,042969419 | 4 |
| *mei-P26.2* | 3,97E-06 | -0,593169113 | 0,324 | 0,448 | 0,047371883 | 4 |
| *CG6966.3* | 4,09E-06 | 0,435399278 | 0,562 | 0,437 | 0,048821876 | 4 |
| *milt.3* | 4,10E-06 | 0,439209962 | 0,541 | 0,406 | 0,048942491 | 4 |
| *RpS17.2* | 4,59E-06 | -0,547211445 | 0,441 | 0,55 | 0,054786281 | 4 |
| *RpL32.2* | 4,89E-06 | -0,594346387 | 0,545 | 0,639 | 0,058281092 | 4 |
| *GstE9.2* | 5,48E-06 | 0,437595917 | 0,362 | 0,243 | 0,065423295 | 4 |
| *bol.2* | 6,47E-06 | -0,71474567 | 0,159 | 0,275 | 0,077124123 | 4 |
| *RpL22* | 6,49E-06 | -0,55077067 | 0,507 | 0,603 | 0,077462099 | 4 |

| | | | | | | |
|---|---|---|---|---|---|---|
| *Bruce.2* | 6,79E-06 | 0,51052816 | 0,466 | 0,361 | 0,081014336 | 4 |
| *Blimp-1.3* | 7,04E-06 | -0,487338152 | 0,641 | 0,705 | 0,083950777 | 4 |
| *tou* | 7,48E-06 | 0,365853137 | 0,307 | 0,192 | 0,089191412 | 4 |
| *Ets98B.2* | 8,14E-06 | -0,533790887 | 0,424 | 0,523 | 0,097061649 | 4 |
| *Bacc.3* | 8,32E-06 | -0,472600569 | 0,728 | 0,758 | 0,099213256 | 4 |
| *Tret1-1.3* | 8,36E-06 | -0,634828345 | 0,469 | 0,571 | 0,099710303 | 4 |
| *HDAC4.1* | 9,48E-06 | 0,462947885 | 0,462 | 0,34 | 0,11308509 | 4 |
| *RpS10b.2* | 9,55E-06 | -0,650732342 | 0,603 | 0,645 | 0,113877392 | 4 |
| *CG1677.2* | 9,77E-06 | -0,637126753 | 0,183 | 0,299 | 0,116550478 | 4 |
| *AP-2alpha* | 1,00E-05 | 0,394429595 | 0,266 | 0,163 | 0,119242339 | 4 |
| *Fkbp14.1* | 1,01E-05 | 0,391297576 | 0,383 | 0,262 | 0,120698224 | 4 |
| *RpL17.3* | 1,08E-05 | -0,542188195 | 0,621 | 0,682 | 0,128644429 | 4 |
| *DIP-lambda.1* | 1,08E-05 | -1,344979752 | 0,183 | 0,285 | 0,129402454 | 4 |
| *RpL13A.3* | 1,15E-05 | -0,460496146 | 0,562 | 0,658 | 0,136670878 | 4 |
| *sxc.2* | 1,15E-05 | -0,605844768 | 0,241 | 0,351 | 0,137109384 | 4 |
| *Flo2.3* | 1,17E-05 | -0,785631275 | 0,207 | 0,311 | 0,139640966 | 4 |
| *Rm62.1* | 1,17E-05 | -0,360326412 | 0,634 | 0,718 | 0,13991484 | 4 |
| *Cdk12* | 1,23E-05 | 0,51076301 | 0,283 | 0,181 | 0,147258572 | 4 |
| *Gyf.1* | 1,31E-05 | -0,481243615 | 0,307 | 0,432 | 0,156563732 | 4 |
| *PlexA.1* | 1,33E-05 | 0,405904591 | 0,548 | 0,427 | 0,158350655 | 4 |
| *RpS16.1* | 1,36E-05 | -0,522747161 | 0,462 | 0,565 | 0,162069157 | 4 |
| *p120ctn.2* | 1,39E-05 | -0,570896543 | 0,252 | 0,365 | 0,166320286 | 4 |
| *RpS6.3* | 1,59E-05 | -0,653188397 | 0,424 | 0,499 | 0,189769606 | 4 |
| *ValRS.1* | 1,65E-05 | 0,335109137 | 0,255 | 0,154 | 0,197393934 | 4 |
| *Samuel* | 1,70E-05 | 0,385078564 | 0,734 | 0,627 | 0,202328799 | 4 |
| *PHGPx.2* | 1,78E-05 | 0,442705694 | 0,493 | 0,365 | 0,212468064 | 4 |
| *Gp93* | 1,79E-05 | 0,320716336 | 0,255 | 0,156 | 0,213941926 | 4 |
| *RpL34b.3* | 2,06E-05 | -0,620416687 | 0,462 | 0,544 | 0,245186399 | 4 |
| *Mapmodulin.2* | 2,16E-05 | -0,68523945 | 0,155 | 0,256 | 0,257123853 | 4 |
| *Pep.1* | 2,37E-05 | 0,370726318 | 0,452 | 0,326 | 0,283265006 | 4 |
| *mam.2* | 2,97E-05 | -0,560270129 | 0,29 | 0,398 | 0,354804654 | 4 |
| *larp.2* | 3,13E-05 | -0,522729457 | 0,493 | 0,595 | 0,373139326 | 4 |
| *E(bx).2* | 3,14E-05 | -0,590158957 | 0,172 | 0,278 | 0,375111364 | 4 |
| *Larp4B* | 3,47E-05 | 0,357896318 | 0,741 | 0,662 | 0,41427075 | 4 |
| *hfp.2* | 3,66E-05 | -0,597945327 | 0,21 | 0,312 | 0,436635435 | 4 |
| *CG32066.2* | 3,70E-05 | 0,383099471 | 0,528 | 0,428 | 0,441782508 | 4 |
| *CG9776* | 3,73E-05 | 0,454532885 | 0,272 | 0,176 | 0,444722623 | 4 |
| *RpL40.2* | 3,73E-05 | -0,657945643 | 0,39 | 0,476 | 0,445153487 | 4 |
| *cta.1* | 3,77E-05 | 0,399990765 | 0,569 | 0,438 | 0,449378699 | 4 |
| *COX6B* | 4,44E-05 | -0,541258988 | 0,159 | 0,26 | 0,529144737 | 4 |
| *cu.2* | 4,77E-05 | 0,298022454 | 0,597 | 0,478 | 0,569371429 | 4 |
| *HnRNP-K* | 5,31E-05 | 0,487314736 | 0,414 | 0,307 | 0,633680441 | 4 |
| *schlank* | 5,39E-05 | 0,433018718 | 0,29 | 0,196 | 0,642417543 | 4 |
| *RpLP0* | 5,60E-05 | -0,616215017 | 0,393 | 0,466 | 0,668235775 | 4 |
| *RpS25.2* | 5,62E-05 | -0,389051566 | 0,466 | 0,587 | 0,670689733 | 4 |
| *Snx6.1* | 5,67E-05 | 0,384728898 | 0,283 | 0,185 | 0,676643179 | 4 |
| *Gs1.2* | 5,86E-05 | -0,677955323 | 0,203 | 0,295 | 0,69938349 | 4 |
| *tay.2* | 6,34E-05 | -0,498640045 | 0,328 | 0,423 | 0,756022374 | 4 |
| *Su(dx)* | 6,39E-05 | 0,483809325 | 0,355 | 0,256 | 0,761720215 | 4 |
| *syd* | 6,40E-05 | 0,477346013 | 0,3 | 0,206 | 0,763603084 | 4 |
| *alpha-Spec.1* | 6,70E-05 | 0,395796022 | 0,334 | 0,232 | 0,799177973 | 4 |
| *CG11400.3* | 6,90E-05 | -0,578179222 | 0,3 | 0,394 | 0,822804184 | 4 |
| *Calr* | 6,95E-05 | 0,363620069 | 0,421 | 0,312 | 0,829056898 | 4 |
| *Rab2.1* | 7,17E-05 | 0,365705901 | 0,29 | 0,189 | 0,855208224 | 4 |
| *RpL21.2* | 7,34E-05 | -0,501182402 | 0,569 | 0,62 | 0,875779777 | 4 |
| *dom* | 7,44E-05 | -0,508760905 | 0,276 | 0,373 | 0,887282145 | 4 |
| *Sirt1.1* | 7,51E-05 | 0,41167241 | 0,31 | 0,213 | 0,896455604 | 4 |
| *CG1673.3* | 7,56E-05 | -0,622929559 | 0,566 | 0,588 | 0,902425199 | 4 |
| *RpS13.1* | 7,67E-05 | -0,475213724 | 0,524 | 0,609 | 0,915528954 | 4 |
| *ATPsynC.2* | 7,81E-05 | -0,486479734 | 0,379 | 0,467 | 0,931310768 | 4 |
| *CG43658.1* | 8,22E-05 | -0,604157031 | 0,652 | 0,677 | 0,980259 | 4 |

| | | | | | | |
|---|---|---|---|---|---|---|
| *tral* | 8,31E-05 | 0,475322141 | 0,476 | 0,386 | 0,991072876 | 4 |
| *magu* | 9,23E-05 | 0,391995112 | 0,269 | 0,174 | 1 | 4 |
| *CG9044.1* | 9,44E-05 | -0,615063013 | 0,193 | 0,284 | 1 | 4 |
| *Snp.1* | 9,92E-05 | -0,529259533 | 0,3 | 0,408 | 1 | 4 |
| *Sfxn1-3.2* | 9,96E-05 | -0,512150129 | 0,269 | 0,373 | 1 | 4 |
| *babo.1* | 0,000101429 | -0,515251686 | 0,21 | 0,306 | 1 | 4 |
| *Rab5.2* | 0,000103859 | 0,338649709 | 0,455 | 0,336 | 1 | 4 |
| *RpL15.2* | 0,000104299 | -0,514618177 | 0,586 | 0,654 | 1 | 4 |
| *anne.2* | 0,000104323 | -0,58368296 | 0,176 | 0,27 | 1 | 4 |
| *Pgm1.3* | 0,000107685 | -0,608708721 | 0,169 | 0,263 | 1 | 4 |
| *tweek.1* | 0,000107719 | 0,412014185 | 0,431 | 0,328 | 1 | 4 |
| *Sik3.2* | 0,000110797 | -0,460211351 | 0,424 | 0,51 | 1 | 4 |
| *CG42663.2* | 0,000111333 | -0,610970342 | 0,186 | 0,287 | 1 | 4 |
| *CG9331.2* | 0,000112525 | 0,369548169 | 0,4 | 0,291 | 1 | 4 |
| *Cka.2* | 0,000114473 | -0,606204449 | 0,234 | 0,327 | 1 | 4 |
| *Rac2.2* | 0,000117583 | 0,505260206 | 0,407 | 0,299 | 1 | 4 |
| *HmgZ* | 0,000121158 | -0,670653122 | 0,179 | 0,267 | 1 | 4 |
| *Pdcd4* | 0,000127111 | 0,497408501 | 0,266 | 0,182 | 1 | 4 |
| *Hex-A* | 0,000128616 | 0,352547912 | 0,252 | 0,162 | 1 | 4 |
| *CAH1.2* | 0,000128905 | -0,814466742 | 0,176 | 0,267 | 1 | 4 |
| *hzg.2* | 0,000132255 | -0,545741341 | 0,217 | 0,311 | 1 | 4 |
| *mnb.2* | 0,000135599 | -0,567017767 | 0,324 | 0,415 | 1 | 4 |
| *Pcyt1.3* | 0,000145126 | 0,262690127 | 0,586 | 0,47 | 1 | 4 |
| *Tctp.1* | 0,000148387 | -0,495589163 | 0,303 | 0,412 | 1 | 4 |
| *RpL23A.1* | 0,000153206 | -0,461391653 | 0,393 | 0,479 | 1 | 4 |
| *RpS23.2* | 0,000163764 | -0,433708414 | 0,61 | 0,684 | 1 | 4 |
| *CG44008.2* | 0,000173047 | 0,338921545 | 0,355 | 0,246 | 1 | 4 |
| *Letm1.2* | 0,000203124 | 0,294074719 | 0,362 | 0,252 | 1 | 4 |
| *Hrb87F* | 0,000223419 | 0,43713845 | 0,276 | 0,187 | 1 | 4 |
| *Mdh1.1* | 0,000232408 | -0,494817465 | 0,29 | 0,382 | 1 | 4 |
| *RpS9.2* | 0,000233956 | -0,406636039 | 0,51 | 0,6 | 1 | 4 |
| *aay.2* | 0,000242668 | -0,730642592 | 0,531 | 0,565 | 1 | 4 |
| *RpS30.3* | 0,000265105 | -0,520706796 | 0,448 | 0,533 | 1 | 4 |
| *kuz.1* | 0,000267653 | 0,595632132 | 0,386 | 0,298 | 1 | 4 |
| *Meltrin.1* | 0,000276607 | -0,489983092 | 0,179 | 0,273 | 1 | 4 |
| *Lsd-2.2* | 0,000288459 | -0,603288037 | 0,855 | 0,815 | 1 | 4 |
| *CG33144.1* | 0,000293519 | 0,425572013 | 0,321 | 0,229 | 1 | 4 |
| *nsl1* | 0,000300603 | 0,288029292 | 0,421 | 0,314 | 1 | 4 |
| *Pcf11.2* | 0,000301819 | -0,350250909 | 0,334 | 0,443 | 1 | 4 |
| *Akt1.2* | 0,000303371 | 0,369000495 | 0,469 | 0,367 | 1 | 4 |
| *dco.2* | 0,000313653 | 0,429947433 | 0,386 | 0,288 | 1 | 4 |
| *daw.2* | 0,000319972 | -0,539345256 | 0,307 | 0,398 | 1 | 4 |
| *CG7029.1* | 0,000366249 | 0,381325676 | 0,472 | 0,384 | 1 | 4 |
| *Jabba.3* | 0,000378193 | -0,701089579 | 0,21 | 0,291 | 1 | 4 |
| *Kdm4B.1* | 0,000382617 | -0,508121991 | 0,186 | 0,278 | 1 | 4 |
| *CkIIbeta.2* | 0,000387514 | -0,507168473 | 0,221 | 0,307 | 1 | 4 |
| *Vrp1* | 0,00041048 | 0,43002696 | 0,276 | 0,195 | 1 | 4 |
| *RpL9.3* | 0,000416559 | -0,529192142 | 0,483 | 0,557 | 1 | 4 |
| *dsd* | 0,000426149 | 0,416761459 | 0,276 | 0,193 | 1 | 4 |
| *mrva.2* | 0,000426908 | -0,43414411 | 0,21 | 0,308 | 1 | 4 |
| *Non1.1* | 0,000453254 | -0,444847701 | 0,269 | 0,358 | 1 | 4 |
| *Pka-R2.2* | 0,00045418 | 0,256101141 | 0,397 | 0,286 | 1 | 4 |
| *klu.2* | 0,000456598 | -0,659860328 | 0,431 | 0,495 | 1 | 4 |
| *fs(1)h.2* | 0,00047447 | -0,283061602 | 0,693 | 0,728 | 1 | 4 |
| *tim.2* | 0,000485806 | -0,351543102 | 0,569 | 0,634 | 1 | 4 |
| *Gs2.2* | 0,000493229 | -0,778810047 | 0,252 | 0,335 | 1 | 4 |
| *AP-1gamma.1* | 0,000504934 | 0,341145121 | 0,355 | 0,261 | 1 | 4 |
| *RpL36A.2* | 0,00053722 | -0,434091058 | 0,514 | 0,579 | 1 | 4 |
| *RpL13.1* | 0,000555516 | -0,428246145 | 0,486 | 0,576 | 1 | 4 |
| *ltl.2* | 0,000556527 | 0,464192092 | 0,297 | 0,216 | 1 | 4 |
| *galectin.2* | 0,000567299 | -0,484085102 | 0,193 | 0,274 | 1 | 4 |

| | | | | | | |
|---|---|---|---|---|---|---|
| *COX5A* | 0,000571376 | -0,531276032 | 0,172 | 0,253 | 1 | 4 |
| *Cyt-c-p* | 0,000663764 | -0,43040319 | 0,39 | 0,472 | 1 | 4 |
| *RhoGAP68F.2* | 0,000685852 | 0,274774163 | 0,303 | 0,213 | 1 | 4 |
| *RpS24.3* | 0,000705045 | -0,48055113 | 0,321 | 0,409 | 1 | 4 |
| *CG32695.1* | 0,000705258 | -0,587168124 | 0,297 | 0,374 | 1 | 4 |
| *sta* | 0,000738814 | -0,456338932 | 0,497 | 0,557 | 1 | 4 |
| *tud* | 0,000740494 | -0,429604109 | 0,172 | 0,256 | 1 | 4 |
| *Nipped-A.2* | 0,000824314 | -0,443112989 | 0,207 | 0,294 | 1 | 4 |
| *Ttd14.2* | 0,000849349 | -0,611855198 | 0,217 | 0,287 | 1 | 4 |
| *lolal.2* | 0,000912131 | -0,371278187 | 0,228 | 0,314 | 1 | 4 |
| *Galphao.2* | 0,000912947 | -0,364601686 | 0,49 | 0,566 | 1 | 4 |
| *RpL11.3* | 0,000960547 | -0,511436809 | 0,466 | 0,517 | 1 | 4 |
| *RpL31.2* | 0,001036098 | -0,428410707 | 0,521 | 0,59 | 1 | 4 |
| *lbk.2* | 0,001075523 | 0,322877572 | 0,352 | 0,259 | 1 | 4 |
| *Mob2.3* | 0,001111191 | -0,5357441 | 0,752 | 0,753 | 1 | 4 |
| *Nap1.2* | 0,001136824 | -0,438450985 | 0,183 | 0,263 | 1 | 4 |
| *Slik.1* | 0,001190782 | 0,285869551 | 0,507 | 0,413 | 1 | 4 |
| *Hrb98DE* | 0,001278626 | -0,4720511 | 0,255 | 0,324 | 1 | 4 |
| *CG7139* | 0,001284408 | 0,318957372 | 0,338 | 0,25 | 1 | 4 |
| *CG15771.2* | 0,001366423 | -0,40594165 | 0,214 | 0,298 | 1 | 4 |
| *lectin-28C* | 0,001655407 | -0,428378911 | 0,255 | 0,337 | 1 | 4 |
| *Lamp1.1* | 0,0016819 | -0,491314291 | 0,31 | 0,373 | 1 | 4 |
| *CG6428.2* | 0,001698388 | 0,334583296 | 0,352 | 0,264 | 1 | 4 |
| *tacc* | 0,001856574 | 0,299013733 | 0,348 | 0,259 | 1 | 4 |
| *ogre.2* | 0,001891795 | -0,347017897 | 0,193 | 0,269 | 1 | 4 |
| *vib* | 0,00193603 | 0,383623946 | 0,39 | 0,31 | 1 | 4 |
| *Ist1.2* | 0,002003576 | 0,269180146 | 0,372 | 0,288 | 1 | 4 |
| *olf186-M.2* | 0,002044869 | -0,481721908 | 0,334 | 0,416 | 1 | 4 |
| *Pzl.2* | 0,002050133 | -0,519297122 | 0,197 | 0,268 | 1 | 4 |
| *RpL35.2* | 0,002095274 | -0,308592962 | 0,479 | 0,549 | 1 | 4 |
| *RpS20.3* | 0,00211524 | -0,545552602 | 0,524 | 0,571 | 1 | 4 |
| *RpS21.3* | 0,0021834 | -0,376633458 | 0,507 | 0,577 | 1 | 4 |
| *RpS19a.1* | 0,002188899 | -0,474616552 | 0,438 | 0,487 | 1 | 4 |
| *how.1* | 0,002199232 | 0,345717508 | 0,607 | 0,536 | 1 | 4 |
| *CG4538.3* | 0,002208712 | -0,389243514 | 0,217 | 0,295 | 1 | 4 |
| *tws* | 0,002321259 | -0,507960305 | 0,266 | 0,336 | 1 | 4 |
| *Paip2.2* | 0,002376424 | -0,536166591 | 0,303 | 0,38 | 1 | 4 |
| *Sox14* | 0,002383769 | 0,346765219 | 0,428 | 0,347 | 1 | 4 |
| *eEF5.3* | 0,002578436 | -0,546275118 | 0,497 | 0,546 | 1 | 4 |
| *gpp.2* | 0,002621489 | -0,379884805 | 0,576 | 0,622 | 1 | 4 |
| *CG7611* | 0,002659266 | -0,467213339 | 0,207 | 0,276 | 1 | 4 |
| *arr.1* | 0,00268315 | -0,554029359 | 0,255 | 0,324 | 1 | 4 |
| *ATPsynF.1* | 0,002698677 | -0,474196418 | 0,19 | 0,258 | 1 | 4 |
| *Hrb27C.2* | 0,002826634 | -0,280407427 | 0,707 | 0,757 | 1 | 4 |
| *B4.2* | 0,002939257 | -0,380576421 | 0,424 | 0,49 | 1 | 4 |
| *osa* | 0,003018739 | 0,256925866 | 0,503 | 0,408 | 1 | 4 |
| *Mical.2* | 0,003282416 | -0,611375749 | 0,252 | 0,313 | 1 | 4 |
| *simj.1* | 0,003318667 | 0,372453328 | 0,348 | 0,269 | 1 | 4 |
| *blw.2* | 0,003403299 | -0,446900145 | 0,366 | 0,424 | 1 | 4 |
| *CG9003* | 0,00361136 | -0,436895112 | 0,197 | 0,266 | 1 | 4 |
| *Patronin* | 0,003634927 | 0,282563212 | 0,428 | 0,343 | 1 | 4 |
| *RpL4.2* | 0,004433396 | -0,422235179 | 0,514 | 0,551 | 1 | 4 |
| *Nrg.1* | 0,004485306 | 0,412513546 | 0,331 | 0,262 | 1 | 4 |
| *Cyt-b5.1* | 0,004505315 | -0,260221673 | 0,424 | 0,495 | 1 | 4 |
| *RpL26.2* | 0,004556408 | -0,449615645 | 0,572 | 0,611 | 1 | 4 |
| *sesB.2* | 0,004641488 | -0,379237706 | 0,452 | 0,496 | 1 | 4 |
| *PIP4K* | 0,004772637 | -0,392062287 | 0,197 | 0,265 | 1 | 4 |
| *cindr.1* | 0,005051858 | 0,264276736 | 0,6 | 0,521 | 1 | 4 |
| *Graf* | 0,005078095 | 0,295094299 | 0,307 | 0,238 | 1 | 4 |
| *14-3-3epsilon* | 0,005088389 | 0,30197703 | 0,659 | 0,618 | 1 | 4 |
| *Rab1.2* | 0,005393854 | 0,298430878 | 0,376 | 0,304 | 1 | 4 |

| | | | | | | |
|---|---|---|---|---|---|---|
| *CG13868.3* | 0,005412461 | -0,266995483 | 0,693 | 0,752 | 1 | 4 |
| *Dyrk3.1* | 0,005559443 | -0,328917962 | 0,448 | 0,514 | 1 | 4 |
| *CrebB.2* | 0,005619871 | -0,409116389 | 0,203 | 0,265 | 1 | 4 |
| *RpL38.3* | 0,005837683 | -0,46098037 | 0,338 | 0,394 | 1 | 4 |
| *RpL23.3* | 0,005976483 | -0,321301776 | 0,655 | 0,697 | 1 | 4 |
| *CG40191* | 0,006829848 | -0,274162647 | 0,255 | 0,328 | 1 | 4 |
| *shi* | 0,007137773 | 0,313399095 | 0,283 | 0,22 | 1 | 4 |
| *CG13631* | 0,008349245 | 0,267553093 | 0,286 | 0,216 | 1 | 4 |
| *Thd1.2* | 0,009385299 | -0,309545985 | 0,407 | 0,459 | 1 | 4 |
| *BomS2.1* | 0,009719763 | -0,71610596 | 0,348 | 0,4 | 1 | 4 |
| *hth* | 1,16E-127 | 2,821169365 | 0,728 | 0,093 | 1,39E-123 | 5 |
| *disco-r* | 2,36E-124 | 2,237506361 | 0,506 | 0,038 | 2,81E-120 | 5 |
| *MTA1-like.3* | 1,34E-72 | 2,491823271 | 0,943 | 0,406 | 1,60E-68 | 5 |
| *CG15143* | 7,49E-58 | 1,414466034 | 0,297 | 0,03 | 8,93E-54 | 5 |
| *ome* | 4,34E-57 | 1,87696616 | 0,532 | 0,107 | 5,17E-53 | 5 |
| *Svil.2* | 1,94E-53 | 1,941748738 | 0,924 | 0,528 | 2,31E-49 | 5 |
| *Helz.3* | 2,70E-51 | 1,409782434 | 0,987 | 0,905 | 3,22E-47 | 5 |
| *CG15145* | 3,88E-49 | 1,466985493 | 0,259 | 0,028 | 4,62E-45 | 5 |
| *Socs36E.4* | 3,76E-48 | 1,960945915 | 0,785 | 0,299 | 4,49E-44 | 5 |
| *LRR.3* | 3,83E-47 | 1,713010518 | 0,918 | 0,503 | 4,57E-43 | 5 |
| *hdc.2* | 7,57E-47 | 2,054824141 | 0,614 | 0,172 | 9,03E-43 | 5 |
| *Sytalpha* | 3,14E-42 | 1,496875045 | 0,31 | 0,048 | 3,75E-38 | 5 |
| *ImpL2.4* | 9,41E-42 | 1,835750961 | 0,671 | 0,225 | 1,12E-37 | 5 |
| *cv-c.3* | 4,45E-40 | 1,368797212 | 0,962 | 0,634 | 5,30E-36 | 5 |
| *Ubx.3* | 1,50E-39 | -1,792801874 | 0,31 | 0,885 | 1,79E-35 | 5 |
| *Pvf2.1* | 5,23E-38 | 1,646977902 | 0,297 | 0,047 | 6,24E-34 | 5 |
| *Ugt49C1* | 2,35E-35 | 1,39360294 | 0,329 | 0,063 | 2,80E-31 | 5 |
| *wb.1* | 9,90E-35 | 1,899934882 | 0,703 | 0,32 | 1,18E-30 | 5 |
| *LpR2.3* | 1,19E-32 | -1,40718907 | 0,386 | 0,871 | 1,42E-28 | 5 |
| *CG5346* | 4,48E-31 | 1,763567972 | 0,392 | 0,103 | 5,35E-27 | 5 |
| *Mmp2.2* | 7,77E-31 | 1,60023448 | 0,557 | 0,197 | 9,27E-27 | 5 |
| *CG34166.3* | 1,16E-30 | -3,074605218 | 0,487 | 0,819 | 1,39E-26 | 5 |
| *Fmo-2.2* | 3,52E-27 | 1,232105086 | 0,576 | 0,213 | 4,20E-23 | 5 |
| *Nep4.2* | 3,95E-26 | 1,554668798 | 0,538 | 0,203 | 4,71E-22 | 5 |
| *CG6770.3* | 2,18E-25 | -1,362964213 | 0,772 | 0,927 | 2,60E-21 | 5 |
| *NK7.1.2* | 1,13E-24 | 1,384539559 | 0,544 | 0,212 | 1,35E-20 | 5 |
| *FASN1.3* | 1,74E-24 | -2,315614217 | 0,563 | 0,803 | 2,07E-20 | 5 |
| *MESK2.1* | 3,52E-24 | 1,3600035 | 0,684 | 0,333 | 4,20E-20 | 5 |
| *Cyp28a5.2* | 3,00E-23 | 1,374175549 | 0,462 | 0,165 | 3,58E-19 | 5 |
| *bnl.3* | 8,40E-22 | 2,419892269 | 0,614 | 0,313 | 1,00E-17 | 5 |
| *CycG.3* | 8,44E-22 | 0,663948312 | 0,981 | 0,946 | 1,01E-17 | 5 |
| *zfh1.2* | 3,42E-21 | 1,166334595 | 0,709 | 0,368 | 4,08E-17 | 5 |
| *atk.2* | 9,43E-21 | 1,129595192 | 0,443 | 0,159 | 1,12E-16 | 5 |
| *Fas3.1* | 3,80E-20 | 1,659283343 | 0,411 | 0,14 | 4,54E-16 | 5 |
| *loco.3* | 1,16E-19 | 1,294265631 | 0,563 | 0,259 | 1,38E-15 | 5 |
| *CG9674.4* | 7,13E-19 | 0,946950837 | 0,905 | 0,619 | 8,51E-15 | 5 |
| *CG33969.2* | 1,05E-18 | 1,089758372 | 0,386 | 0,132 | 1,25E-14 | 5 |
| *CG34054.2* | 4,10E-18 | 0,903518816 | 0,456 | 0,176 | 4,90E-14 | 5 |
| *Gprk2.3* | 1,25E-17 | 1,097838039 | 0,892 | 0,686 | 1,49E-13 | 5 |
| *CG10082.4* | 1,34E-17 | -1,720605178 | 0,316 | 0,619 | 1,60E-13 | 5 |
| *cher.3* | 5,59E-17 | 1,377786033 | 0,563 | 0,277 | 6,67E-13 | 5 |
| *Mob2.4* | 8,74E-17 | 0,865386156 | 0,911 | 0,744 | 1,04E-12 | 5 |
| *DIP-lambda.2* | 1,14E-16 | 1,485213903 | 0,551 | 0,261 | 1,36E-12 | 5 |
| *Abl.4* | 4,20E-16 | 1,000431729 | 0,671 | 0,356 | 5,01E-12 | 5 |
| *jing.1* | 4,21E-16 | 1,03717468 | 0,494 | 0,218 | 5,02E-12 | 5 |
| *CG14823.4* | 5,53E-16 | -2,190708576 | 0,051 | 0,361 | 6,59E-12 | 5 |
| *Sdc.3* | 7,10E-16 | 0,611327333 | 1 | 0,946 | 8,47E-12 | 5 |
| *mld.2* | 1,34E-15 | 1,004535112 | 0,576 | 0,293 | 1,59E-11 | 5 |
| *Atf3.3* | 2,96E-15 | 1,010707719 | 0,551 | 0,281 | 3,53E-11 | 5 |
| *Gbs-70E.4* | 3,38E-15 | -2,757029397 | 0,285 | 0,542 | 4,03E-11 | 5 |
| *CG34347.4* | 4,52E-15 | 0,956838323 | 0,658 | 0,363 | 5,40E-11 | 5 |

| | | | | | | |
|---|---|---|---|---|---|---|
| *Src64B.3* | 7,47E-15 | 0,978545484 | 0,804 | 0,52 | 8,91E-11 | 5 |
| *CG1648.4* | 9,17E-15 | -1,673621271 | 0,38 | 0,623 | 1,09E-10 | 5 |
| *l(2)41Ab.2* | 2,33E-14 | 1,538604773 | 0,589 | 0,354 | 2,78E-10 | 5 |
| *InR.3* | 3,34E-14 | 0,583186647 | 0,994 | 0,908 | 3,98E-10 | 5 |
| *Bacc.4* | 8,25E-14 | -1,038366531 | 0,538 | 0,767 | 9,84E-10 | 5 |
| *CG42788.3* | 1,23E-13 | -2,249599476 | 0,222 | 0,496 | 1,47E-09 | 5 |
| *fry.2* | 1,43E-13 | 0,950865629 | 0,443 | 0,206 | 1,70E-09 | 5 |
| *GEFmeso.3* | 1,73E-13 | 0,906910393 | 0,703 | 0,424 | 2,06E-09 | 5 |
| *Swim.1* | 3,93E-13 | 1,226948915 | 0,272 | 0,094 | 4,69E-09 | 5 |
| *AcCoAS.3* | 4,25E-13 | -1,287666032 | 0,468 | 0,695 | 5,07E-09 | 5 |
| *Nplp2.4* | 4,88E-13 | -1,765414372 | 0,506 | 0,676 | 5,82E-09 | 5 |
| *CG31678.3* | 4,89E-13 | 0,909773929 | 0,449 | 0,208 | 5,84E-09 | 5 |
| *Gpdh1.4* | 7,21E-13 | -1,919085868 | 0,101 | 0,376 | 8,60E-09 | 5 |
| *Pka-C1.3* | 9,27E-13 | 0,752117054 | 0,861 | 0,657 | 1,11E-08 | 5 |
| *CG8745.4* | 1,07E-12 | -2,384582064 | 0,108 | 0,381 | 1,28E-08 | 5 |
| *Hers.4* | 1,45E-12 | 0,697421627 | 0,854 | 0,61 | 1,73E-08 | 5 |
| *RhoGAP71E.3* | 1,63E-12 | 1,129853951 | 0,671 | 0,414 | 1,94E-08 | 5 |
| *Ufd4.3* | 1,64E-12 | 0,746175015 | 0,589 | 0,333 | 1,95E-08 | 5 |
| *Lip4.2* | 1,75E-12 | 0,948184778 | 0,646 | 0,395 | 2,08E-08 | 5 |
| *CG9451.3* | 2,27E-12 | 1,128037339 | 0,443 | 0,22 | 2,71E-08 | 5 |
| *bun.4* | 2,60E-12 | -1,39718497 | 0,778 | 0,847 | 3,10E-08 | 5 |
| *GstT4.2* | 3,23E-12 | -1,874608679 | 0,095 | 0,354 | 3,85E-08 | 5 |
| *Kdm4B.2* | 3,27E-12 | 0,880083835 | 0,5 | 0,257 | 3,90E-08 | 5 |
| *CG1213.3* | 4,68E-12 | -1,878147864 | 0,095 | 0,354 | 5,59E-08 | 5 |
| *nmo.3* | 6,20E-12 | 0,865766252 | 0,778 | 0,567 | 7,40E-08 | 5 |
| *sfl.3* | 1,06E-11 | 0,898230834 | 0,715 | 0,503 | 1,27E-07 | 5 |
| *spir.3* | 1,68E-11 | 0,86366379 | 0,595 | 0,359 | 2,00E-07 | 5 |
| *Galk.3* | 1,81E-11 | -1,322460968 | 0,278 | 0,514 | 2,16E-07 | 5 |
| *Cyp4e3* | 1,95E-11 | 0,906567464 | 0,335 | 0,138 | 2,33E-07 | 5 |
| *Nc73EF.3* | 2,65E-11 | -1,154513587 | 0,487 | 0,662 | 3,16E-07 | 5 |
| *klu.3* | 2,86E-11 | 0,886641616 | 0,715 | 0,477 | 3,42E-07 | 5 |
| *Frl.4* | 5,08E-11 | 0,648086633 | 0,753 | 0,501 | 6,05E-07 | 5 |
| *Pde6.4* | 5,45E-11 | -1,257845138 | 0,285 | 0,524 | 6,50E-07 | 5 |
| *CG3036.4* | 6,48E-11 | 0,838491987 | 0,829 | 0,642 | 7,73E-07 | 5 |
| *CG14154.3* | 7,00E-11 | 0,814429603 | 0,665 | 0,452 | 8,36E-07 | 5 |
| *promL* | 8,56E-11 | 0,80236906 | 0,411 | 0,194 | 1,02E-06 | 5 |
| *spen.1* | 9,79E-11 | 0,433120436 | 0,981 | 0,959 | 1,17E-06 | 5 |
| *CG1703.1* | 1,28E-10 | 0,742572948 | 0,38 | 0,174 | 1,52E-06 | 5 |
| *for.3* | 1,71E-10 | -0,819044662 | 0,582 | 0,745 | 2,05E-06 | 5 |
| *c11.1.3* | 2,51E-10 | 0,697922895 | 0,462 | 0,235 | 3,00E-06 | 5 |
| *Obp99b* | 2,98E-10 | 2,154641132 | 0,285 | 0,118 | 3,55E-06 | 5 |
| *AdipoR.3* | 3,56E-10 | -1,274133622 | 0,361 | 0,547 | 4,24E-06 | 5 |
| *msn.2* | 3,80E-10 | 0,640925703 | 0,759 | 0,576 | 4,53E-06 | 5 |
| *fbp.4* | 4,51E-10 | -1,370448283 | 0,101 | 0,328 | 5,37E-06 | 5 |
| *Ldh.1* | 5,21E-10 | 0,934338983 | 0,31 | 0,138 | 6,22E-06 | 5 |
| *CG15117* | 7,15E-10 | 0,661759976 | 0,259 | 0,1 | 8,53E-06 | 5 |
| *chb.3* | 7,53E-10 | 0,788100431 | 0,5 | 0,283 | 8,99E-06 | 5 |
| *CG42668.4* | 8,11E-10 | 0,642798998 | 0,791 | 0,589 | 9,68E-06 | 5 |
| *CG8312.4* | 9,17E-10 | 0,88405821 | 0,551 | 0,315 | 1,09E-05 | 5 |
| *CG18135.2* | 1,19E-09 | -0,949002833 | 0,576 | 0,729 | 1,42E-05 | 5 |
| *DOR.4* | 1,26E-09 | -1,076320373 | 0,424 | 0,618 | 1,50E-05 | 5 |
| *Gug.4* | 1,29E-09 | -0,783940493 | 0,468 | 0,651 | 1,53E-05 | 5 |
| *Ack-like* | 1,44E-09 | 0,776786553 | 0,285 | 0,122 | 1,72E-05 | 5 |
| *CG6512.2* | 1,45E-09 | 0,70376251 | 0,354 | 0,164 | 1,73E-05 | 5 |
| *Tsp42Ea.1* | 1,53E-09 | 0,788679776 | 0,323 | 0,144 | 1,83E-05 | 5 |
| *CG3168.3* | 2,62E-09 | 0,525833114 | 0,525 | 0,282 | 3,12E-05 | 5 |
| *CCHa2.4* | 2,92E-09 | -1,578731665 | 0,063 | 0,273 | 3,48E-05 | 5 |
| *drpr.4* | 3,12E-09 | 0,692347491 | 0,684 | 0,445 | 3,72E-05 | 5 |
| *Ptp4E* | 3,52E-09 | 0,777146113 | 0,316 | 0,145 | 4,20E-05 | 5 |
| *PGRP-LC* | 3,56E-09 | 0,70920577 | 0,316 | 0,143 | 4,25E-05 | 5 |
| *Btk29A.1* | 4,60E-09 | 0,721330244 | 0,43 | 0,224 | 5,48E-05 | 5 |

| | | | | | | |
|---|---|---|---|---|---|---|
| *Men.3* | 5,50E-09 | -1,225519268 | 0,532 | 0,68 | 6,56E-05 | 5 |
| *CG8112.1* | 7,22E-09 | 0,720728827 | 0,335 | 0,158 | 8,61E-05 | 5 |
| *Tsp42Ed.3* | 7,39E-09 | 0,539671238 | 0,696 | 0,423 | 8,81E-05 | 5 |
| *trbl.3* | 7,82E-09 | 0,528364393 | 0,747 | 0,5 | 9,32E-05 | 5 |
| *Hipk.3* | 8,07E-09 | 0,613092712 | 0,854 | 0,678 | 9,62E-05 | 5 |
| *ACC.4* | 8,90E-09 | -0,985218777 | 0,57 | 0,711 | 0,000106225 | 5 |
| *Cam.2* | 8,93E-09 | 0,694808258 | 0,899 | 0,816 | 0,000106511 | 5 |
| *Invadolysin.4* | 9,27E-09 | -1,92552113 | 0,082 | 0,283 | 0,000110633 | 5 |
| *CG31523.3* | 9,82E-09 | 0,73057449 | 0,449 | 0,254 | 0,0001171 | 5 |
| *Meltrin.2* | 1,10E-08 | 0,824533523 | 0,449 | 0,254 | 0,000130838 | 5 |
| *Idh.4* | 1,18E-08 | -1,291190154 | 0,234 | 0,429 | 0,000140345 | 5 |
| *oys.4* | 1,25E-08 | 0,516996124 | 0,557 | 0,32 | 0,00014868 | 5 |
| *CG17841.3* | 1,30E-08 | -1,182004627 | 0,304 | 0,485 | 0,00015479 | 5 |
| *l(2)37Cc* | 1,40E-08 | 0,645060809 | 0,253 | 0,107 | 0,000167174 | 5 |
| *Pdp1.4* | 1,46E-08 | -0,459169271 | 0,994 | 0,993 | 0,000173875 | 5 |
| *Usp15-31* | 1,54E-08 | 0,778031229 | 0,272 | 0,12 | 0,000183311 | 5 |
| *CG6910.4* | 1,54E-08 | -0,961047935 | 0,633 | 0,765 | 0,000183437 | 5 |
| *E(spl)malpha-BFM.* | 1,64E-08 | 0,7232367 | 0,538 | 0,336 | 0,00019515 | 5 |
| *Eip74EF.1* | 1,86E-08 | 0,534765271 | 0,804 | 0,617 | 0,000221947 | 5 |
| *LPCAT.3* | 1,97E-08 | 0,816541326 | 0,373 | 0,196 | 0,000234947 | 5 |
| *nuf.4* | 2,04E-08 | 0,582551076 | 0,722 | 0,539 | 0,000243659 | 5 |
| *CD98hc.3* | 2,12E-08 | 0,617808312 | 0,405 | 0,213 | 0,000252989 | 5 |
| *Cyp6g1.3* | 2,22E-08 | -1,103224497 | 0,209 | 0,413 | 0,000264359 | 5 |
| *CG16926.3* | 2,22E-08 | -0,951807228 | 0,481 | 0,63 | 0,000264684 | 5 |
| *par-1.3* | 2,35E-08 | 0,415948151 | 0,987 | 0,939 | 0,000280872 | 5 |
| *sky.3* | 2,84E-08 | 0,801482881 | 0,424 | 0,233 | 0,000338729 | 5 |
| *CG7720.4* | 3,48E-08 | -2,016897792 | 0,171 | 0,365 | 0,0004152 | 5 |
| *CG18622* | 3,66E-08 | 0,754663941 | 0,285 | 0,133 | 0,000436726 | 5 |
| *alt.3* | 3,94E-08 | 0,865367796 | 0,532 | 0,349 | 0,000470229 | 5 |
| *CG43402.1* | 3,98E-08 | 0,984495626 | 0,373 | 0,203 | 0,000475332 | 5 |
| *Mical.3* | 4,20E-08 | 0,613820748 | 0,506 | 0,296 | 0,000501596 | 5 |
| *pHCl-2.1* | 4,42E-08 | 0,544171481 | 0,329 | 0,158 | 0,000527548 | 5 |
| *CG11089.3* | 4,51E-08 | 0,496881107 | 0,88 | 0,649 | 0,000537582 | 5 |
| *CG1572* | 4,64E-08 | 0,705402853 | 0,266 | 0,117 | 0,000553919 | 5 |
| *Cyt-b5-r.3* | 5,17E-08 | -0,999330029 | 0,399 | 0,573 | 0,000616682 | 5 |
| *UGP.3* | 5,89E-08 | -1,262581067 | 0,234 | 0,432 | 0,000702468 | 5 |
| *CG15096.4* | 6,09E-08 | -1,108763198 | 0,07 | 0,262 | 0,000726612 | 5 |
| *Drak.3* | 6,98E-08 | -1,13970823 | 0,462 | 0,63 | 0,000833132 | 5 |
| *h.2* | 7,02E-08 | 0,469911146 | 0,804 | 0,625 | 0,000837938 | 5 |
| *Tret1-1.4* | 7,77E-08 | -1,218048367 | 0,43 | 0,569 | 0,000926977 | 5 |
| *mtgo.1* | 7,88E-08 | 1,61088735 | 0,411 | 0,257 | 0,00093979 | 5 |
| *CG32767.4* | 8,09E-08 | -1,063213394 | 0,203 | 0,397 | 0,000964879 | 5 |
| *CG8034.4* | 9,30E-08 | -1,083741016 | 0,443 | 0,613 | 0,001109372 | 5 |
| *CG12991* | 1,08E-07 | 0,819325748 | 0,259 | 0,116 | 0,00128542 | 5 |
| *Acbp2.3* | 1,13E-07 | -1,190272378 | 0,133 | 0,323 | 0,001349604 | 5 |
| *ckn.2* | 1,23E-07 | 0,68608106 | 0,519 | 0,334 | 0,001468978 | 5 |
| *puc.4* | 1,27E-07 | 0,497641232 | 0,873 | 0,684 | 0,001517479 | 5 |
| *Nmdmc.3* | 1,35E-07 | 0,487730644 | 0,886 | 0,631 | 0,001612057 | 5 |
| *olf186-M.3* | 1,36E-07 | -1,283108745 | 0,234 | 0,418 | 0,00161867 | 5 |
| *Rala.2* | 1,37E-07 | 0,582416186 | 0,677 | 0,482 | 0,001630644 | 5 |
| *ValRS.2* | 1,54E-07 | 0,619853591 | 0,31 | 0,156 | 0,001838257 | 5 |
| *RhoGAP19D.4* | 1,76E-07 | 0,589862419 | 0,665 | 0,472 | 0,002098638 | 5 |
| *inaE.1* | 1,84E-07 | 0,727775391 | 0,291 | 0,14 | 0,002200429 | 5 |
| *Culd.4* | 1,99E-07 | -1,201249865 | 0,171 | 0,353 | 0,002372008 | 5 |
| *Tspo* | 2,08E-07 | 0,499759379 | 0,266 | 0,121 | 0,002476575 | 5 |
| *cic.4* | 2,20E-07 | -0,961327627 | 0,532 | 0,671 | 0,002625666 | 5 |
| *cac.3* | 2,34E-07 | 1,418230672 | 0,608 | 0,447 | 0,002796482 | 5 |
| *Rac2.3* | 2,74E-07 | 0,617543004 | 0,494 | 0,299 | 0,003268351 | 5 |
| *Eip93F.1* | 2,91E-07 | 0,316999194 | 1 | 0,987 | 0,003476567 | 5 |
| *goe.4* | 3,79E-07 | -1,223870713 | 0,101 | 0,28 | 0,004522264 | 5 |
| *elB.4* | 4,07E-07 | 0,384681657 | 0,519 | 0,3 | 0,004858356 | 5 |

| | | | | | | |
|---|---|---|---|---|---|---|
| *Chd64.3* | 5,48E-07 | 0,50901554 | 0,665 | 0,445 | 0,006536831 | 5 |
| *sug.4* | 5,64E-07 | -1,776600468 | 0,165 | 0,333 | 0,006728192 | 5 |
| *crc.3* | 7,06E-07 | 0,512869567 | 0,57 | 0,363 | 0,008426505 | 5 |
| *Jabba.4* | 7,19E-07 | -1,229980365 | 0,12 | 0,292 | 0,008575021 | 5 |
| *Apoltp.3* | 7,24E-07 | 0,313133028 | 0,924 | 0,787 | 0,008634175 | 5 |
| *CG2124* | 7,26E-07 | 0,49884792 | 0,266 | 0,126 | 0,008664939 | 5 |
| *Exn.2* | 7,59E-07 | 0,569344981 | 0,684 | 0,498 | 0,00905955 | 5 |
| *CG7920.4* | 8,78E-07 | -1,061226907 | 0,133 | 0,304 | 0,010475391 | 5 |
| *CG31635.4* | 9,16E-07 | -0,806893304 | 0,392 | 0,559 | 0,010929757 | 5 |
| *l(1)G0289.3* | 9,34E-07 | 0,629627984 | 0,658 | 0,463 | 0,011147255 | 5 |
| *CG14153.1* | 1,18E-06 | 0,633173417 | 0,297 | 0,152 | 0,014059231 | 5 |
| *kay.4* | 1,23E-06 | 0,399568214 | 0,709 | 0,507 | 0,014654286 | 5 |
| *ps.3* | 1,24E-06 | -0,461681078 | 0,994 | 0,983 | 0,014764867 | 5 |
| *PH4alphaEFB.1* | 1,27E-06 | 0,583268012 | 0,316 | 0,169 | 0,015161509 | 5 |
| *flw.2* | 1,40E-06 | 0,575204804 | 0,601 | 0,411 | 0,016739257 | 5 |
| *Pmp70.4* | 1,48E-06 | -1,052250468 | 0,108 | 0,279 | 0,017668358 | 5 |
| *baz.3* | 1,59E-06 | 0,588691925 | 0,405 | 0,227 | 0,019003295 | 5 |
| *Thor.3* | 1,86E-06 | -1,109654575 | 0,418 | 0,543 | 0,022161995 | 5 |
| *CG17646.3* | 1,89E-06 | -0,510201612 | 0,715 | 0,859 | 0,022559411 | 5 |
| *Mkp3.2* | 2,00E-06 | -0,744901027 | 0,234 | 0,409 | 0,023875106 | 5 |
| *Rel.3* | 2,05E-06 | 0,488651675 | 0,703 | 0,528 | 0,024464431 | 5 |
| *retm.4* | 2,12E-06 | -0,994051695 | 0,127 | 0,295 | 0,025347937 | 5 |
| *pst.3* | 2,13E-06 | 0,574660856 | 0,797 | 0,573 | 0,025411954 | 5 |
| *bbg.2* | 2,33E-06 | -1,095456424 | 0,177 | 0,346 | 0,027840223 | 5 |
| *Hex-A.1* | 2,45E-06 | 0,540914171 | 0,304 | 0,163 | 0,029255996 | 5 |
| *gce.4* | 2,48E-06 | -0,943362698 | 0,342 | 0,49 | 0,029565268 | 5 |
| *Bsg.1* | 2,59E-06 | 0,330006245 | 0,994 | 0,982 | 0,030943683 | 5 |
| *Letm1.3* | 2,76E-06 | 0,498508994 | 0,43 | 0,253 | 0,032979817 | 5 |
| *Gdap2.3* | 2,80E-06 | 0,517086879 | 0,582 | 0,398 | 0,033358113 | 5 |
| *HDAC4.2* | 2,99E-06 | 0,515251306 | 0,532 | 0,341 | 0,035713292 | 5 |
| *CG8485.4* | 3,46E-06 | -1,036987338 | 0,095 | 0,255 | 0,041254638 | 5 |
| *Usp10.1* | 3,50E-06 | 0,509234099 | 0,551 | 0,372 | 0,041788725 | 5 |
| *CG13868.4* | 3,64E-06 | -0,55194206 | 0,658 | 0,751 | 0,043390221 | 5 |
| *Tsf1.4* | 3,73E-06 | -1,273134243 | 0,146 | 0,302 | 0,044449484 | 5 |
| *Moe.2* | 4,01E-06 | 0,647550394 | 0,487 | 0,316 | 0,047786926 | 5 |
| *Eglp4.4* | 4,13E-06 | -1,267004586 | 0,234 | 0,377 | 0,049233878 | 5 |
| *Gpo1.3* | 4,28E-06 | -1,298760091 | 0,133 | 0,286 | 0,051062112 | 5 |
| *Cdc42* | 4,56E-06 | 0,528936665 | 0,316 | 0,173 | 0,054377191 | 5 |
| *Sodh-1.4* | 5,03E-06 | -1,218313389 | 0,152 | 0,303 | 0,059969369 | 5 |
| *spoon.4* | 5,06E-06 | -0,90752312 | 0,285 | 0,438 | 0,060359207 | 5 |
| *fok.2* | 5,09E-06 | 0,586072672 | 0,696 | 0,512 | 0,060765556 | 5 |
| *CG13887.2* | 5,15E-06 | 0,548483761 | 0,291 | 0,155 | 0,061405364 | 5 |
| *Cyp28d1.3* | 5,22E-06 | 0,424722749 | 0,373 | 0,21 | 0,062320818 | 5 |
| *REPTOR.3* | 5,41E-06 | -0,641082248 | 0,627 | 0,722 | 0,064487066 | 5 |
| *if.4* | 5,84E-06 | -1,324781837 | 0,203 | 0,356 | 0,069635637 | 5 |
| *caps.3* | 6,40E-06 | 0,533589331 | 0,677 | 0,492 | 0,076352399 | 5 |
| *CG5966.2* | 6,67E-06 | 0,566610883 | 0,614 | 0,443 | 0,079619437 | 5 |
| *Spn43Ab.3* | 7,28E-06 | -0,953735909 | 0,228 | 0,368 | 0,086833719 | 5 |
| *Cyp4p1.3* | 7,86E-06 | 0,495662144 | 0,475 | 0,301 | 0,093727519 | 5 |
| *Snx6.2* | 8,05E-06 | 0,441003559 | 0,335 | 0,186 | 0,096053117 | 5 |
| *CG6115.3* | 8,61E-06 | 0,502934942 | 0,595 | 0,42 | 0,102745047 | 5 |
| *CG44325.4* | 9,57E-06 | 0,752186146 | 0,475 | 0,309 | 0,11418731 | 5 |
| *beta-Spec* | 9,92E-06 | 0,531249525 | 0,278 | 0,145 | 0,118327518 | 5 |
| *tkv.3* | 1,01E-05 | -1,063791074 | 0,272 | 0,399 | 0,120130869 | 5 |
| *Pdk.2* | 1,02E-05 | -0,413451699 | 0,791 | 0,866 | 0,121211234 | 5 |
| *GstE9.3* | 1,11E-05 | 0,595333299 | 0,399 | 0,246 | 0,132503936 | 5 |
| *CG7766.3* | 1,24E-05 | -0,810250671 | 0,475 | 0,578 | 0,147915353 | 5 |
| *CG13631.1* | 1,24E-05 | 0,476430736 | 0,367 | 0,215 | 0,148245747 | 5 |
| *Ptpmeg2.1* | 1,33E-05 | 0,739631588 | 0,361 | 0,222 | 0,15806209 | 5 |
| *Nop17l.4* | 1,35E-05 | -1,081060599 | 0,297 | 0,42 | 0,160445069 | 5 |
| *CG10621.4* | 1,40E-05 | -1,322071895 | 0,127 | 0,276 | 0,166789414 | 5 |

| | | | | | | |
|---|---|---|---|---|---|---|
| *Ork1.3* | 1,43E-05 | -0,977979356 | 0,196 | 0,345 | 0,170385716 | 5 |
| *Snp.2* | 1,52E-05 | -0,823787659 | 0,247 | 0,406 | 0,181387271 | 5 |
| *kdn.3* | 1,53E-05 | -1,137757092 | 0,158 | 0,306 | 0,182374914 | 5 |
| *shn.2* | 1,54E-05 | 0,539703609 | 0,728 | 0,611 | 0,183200922 | 5 |
| *Hsp27.4* | 1,54E-05 | -2,360526014 | 0,215 | 0,348 | 0,18383092 | 5 |
| *Rtnl1.2* | 1,54E-05 | 0,318049896 | 0,532 | 0,34 | 0,184150395 | 5 |
| *Nipped-B.2* | 1,62E-05 | 0,52369905 | 0,854 | 0,745 | 0,193663485 | 5 |
| *hts.1* | 1,63E-05 | 0,45387306 | 0,658 | 0,479 | 0,193883628 | 5 |
| *Acsl.3* | 1,64E-05 | -1,075099283 | 0,481 | 0,56 | 0,19528074 | 5 |
| *Rpn13.3* | 1,65E-05 | 0,443681186 | 0,437 | 0,266 | 0,196392821 | 5 |
| *bmm.3* | 1,70E-05 | 0,380618414 | 0,88 | 0,756 | 0,202564409 | 5 |
| *Sam-S.3* | 1,77E-05 | -0,837210873 | 0,487 | 0,62 | 0,210650368 | 5 |
| *betaTub97EF.2* | 1,85E-05 | 0,492747584 | 0,43 | 0,273 | 0,22052471 | 5 |
| *GIIIspla2.2* | 2,05E-05 | 0,377028953 | 0,608 | 0,407 | 0,244806956 | 5 |
| *mew.3* | 2,26E-05 | -1,029584971 | 0,253 | 0,396 | 0,269891602 | 5 |
| *D2hgdh.3* | 2,34E-05 | -0,92020295 | 0,171 | 0,312 | 0,279509366 | 5 |
| *CG34423.1* | 2,49E-05 | 0,595414034 | 0,253 | 0,136 | 0,296963963 | 5 |
| *14-3-3epsilon.1* | 2,59E-05 | 0,363633839 | 0,772 | 0,614 | 0,30909159 | 5 |
| *cact.4* | 2,61E-05 | 0,473835297 | 0,766 | 0,635 | 0,311667705 | 5 |
| *CG1673.4* | 2,65E-05 | 0,35443212 | 0,766 | 0,576 | 0,316105337 | 5 |
| *Phb2* | 2,87E-05 | 0,460317006 | 0,259 | 0,139 | 0,341994329 | 5 |
| *qless* | 2,97E-05 | 0,50843045 | 0,259 | 0,139 | 0,353857144 | 5 |
| *IP3K2.3* | 3,20E-05 | 0,473920474 | 0,791 | 0,652 | 0,38141308 | 5 |
| *Cyp309a1.3* | 3,45E-05 | 0,541085012 | 0,373 | 0,232 | 0,411480367 | 5 |
| *Ubqn.3* | 3,72E-05 | 0,494880147 | 0,399 | 0,249 | 0,444195661 | 5 |
| *cdi.2* | 3,77E-05 | 0,864903771 | 0,532 | 0,395 | 0,450164045 | 5 |
| *Nop60B* | 4,02E-05 | 0,27736733 | 0,278 | 0,148 | 0,478972645 | 5 |
| *Vinc.2* | 4,16E-05 | 0,489830767 | 0,304 | 0,176 | 0,495757528 | 5 |
| *Nhe3* | 4,17E-05 | 0,476145635 | 0,253 | 0,134 | 0,496950926 | 5 |
| *Oda.3* | 4,21E-05 | -0,470707938 | 0,797 | 0,824 | 0,502018854 | 5 |
| *Flo2.4* | 4,25E-05 | 0,476520608 | 0,456 | 0,293 | 0,506746188 | 5 |
| *Ire1.3* | 4,27E-05 | 0,428470771 | 0,481 | 0,308 | 0,509358478 | 5 |
| *AnxB9.4* | 4,37E-05 | 0,260161545 | 0,703 | 0,485 | 0,521860674 | 5 |
| *SNF4Agamma.3* | 4,51E-05 | 0,515156436 | 0,949 | 0,889 | 0,538040718 | 5 |
| *CG5853.2* | 4,79E-05 | 0,802155407 | 0,43 | 0,29 | 0,571193034 | 5 |
| *Nrg.2* | 4,90E-05 | 0,479422393 | 0,418 | 0,26 | 0,58455389 | 5 |
| *CG5958.4* | 5,01E-05 | 0,665510495 | 0,551 | 0,405 | 0,598053275 | 5 |
| *CG3764.3* | 5,04E-05 | -0,917726242 | 0,165 | 0,305 | 0,601757552 | 5 |
| *cwo.3* | 5,05E-05 | -0,501647314 | 0,892 | 0,878 | 0,602715738 | 5 |
| *Ndae1.3* | 5,18E-05 | -0,968749071 | 0,196 | 0,346 | 0,618437042 | 5 |
| *KrT95D.4* | 5,19E-05 | 0,554524284 | 0,759 | 0,68 | 0,618621234 | 5 |
| *CG8671.1* | 5,38E-05 | 0,37876563 | 0,468 | 0,302 | 0,641710428 | 5 |
| *CG6067.3* | 5,47E-05 | -0,874521903 | 0,196 | 0,339 | 0,652174724 | 5 |
| *CG4199.2* | 5,52E-05 | 0,509095352 | 0,38 | 0,235 | 0,658614212 | 5 |
| *Aldh.2* | 5,56E-05 | -0,696770368 | 0,424 | 0,549 | 0,663441677 | 5 |
| *Msp300.3* | 5,63E-05 | -0,355794485 | 0,823 | 0,885 | 0,672101301 | 5 |
| *Zdhhc8* | 5,66E-05 | 0,710264489 | 0,614 | 0,458 | 0,67541624 | 5 |
| *B4.3* | 5,96E-05 | 0,432654515 | 0,646 | 0,476 | 0,711051807 | 5 |
| *CG17124.4* | 5,99E-05 | -0,718033949 | 0,696 | 0,751 | 0,714485131 | 5 |
| *kek5.3* | 6,10E-05 | 0,266064195 | 0,791 | 0,592 | 0,728034022 | 5 |
| *CG13784.2* | 6,46E-05 | 0,393976669 | 0,766 | 0,61 | 0,770330288 | 5 |
| *Stat92E.4* | 7,51E-05 | -0,972784221 | 0,525 | 0,593 | 0,896070408 | 5 |
| *Blimp-1.4* | 7,57E-05 | -0,669823729 | 0,639 | 0,702 | 0,902757695 | 5 |
| *Gs1.3* | 7,59E-05 | 0,538155756 | 0,424 | 0,279 | 0,905324957 | 5 |
| *CG7668.1* | 7,91E-05 | 0,369843083 | 0,272 | 0,148 | 0,943028935 | 5 |
| *Ten-m.1* | 8,32E-05 | 0,618091747 | 0,671 | 0,57 | 0,99240728 | 5 |
| *ninaE.3* | 8,45E-05 | 0,470836285 | 0,38 | 0,239 | 1 | 5 |
| *IscU.2* | 8,46E-05 | 0,391136603 | 0,595 | 0,413 | 1 | 5 |
| *Cyt-c-p.1* | 8,83E-05 | 0,449759872 | 0,608 | 0,457 | 1 | 5 |
| *Khc.1* | 9,23E-05 | 0,454900242 | 0,291 | 0,171 | 1 | 5 |
| *Act5C.4* | 9,82E-05 | 0,328711741 | 0,722 | 0,537 | 1 | 5 |

| | | | | | | |
|---|---|---|---|---|---|---|
| *crol* | 9,97E-05 | 0,366473895 | 0,722 | 0,575 | 1 | 5 |
| *cl.1* | 0,00010135 | 0,3584315 | 0,259 | 0,143 | 1 | 5 |
| *cbt.3* | 0,000101991 | -0,750741328 | 0,146 | 0,284 | 1 | 5 |
| *CG1578.3* | 0,000104732 | -0,902362479 | 0,253 | 0,388 | 1 | 5 |
| *CG14207.3* | 0,000105872 | 0,476350834 | 0,734 | 0,579 | 1 | 5 |
| *CG12012.2* | 0,00010963 | 0,485224869 | 0,411 | 0,269 | 1 | 5 |
| *Nedd4.3* | 0,000124601 | 0,474334877 | 0,468 | 0,325 | 1 | 5 |
| *vtd* | 0,000137642 | 0,317786295 | 0,354 | 0,212 | 1 | 5 |
| *mtd.3* | 0,000141019 | -0,786773613 | 0,639 | 0,694 | 1 | 5 |
| *AkhR.4* | 0,000149756 | -0,979857501 | 0,209 | 0,325 | 1 | 5 |
| *CG17549.3* | 0,000150843 | -0,76311113 | 0,253 | 0,379 | 1 | 5 |
| *CG12065.3* | 0,000166516 | 0,50945446 | 0,43 | 0,288 | 1 | 5 |
| *IM4.4* | 0,000167609 | -1,29966007 | 0,19 | 0,313 | 1 | 5 |
| *Dif.3* | 0,000172674 | -0,720957155 | 0,266 | 0,4 | 1 | 5 |
| *Gbs-76A.4* | 0,000173002 | -0,691962524 | 0,278 | 0,42 | 1 | 5 |
| *edl.1* | 0,000176842 | 0,357768359 | 0,266 | 0,148 | 1 | 5 |
| *lost.3* | 0,000178803 | 0,372068539 | 0,538 | 0,384 | 1 | 5 |
| *CG3902.3* | 0,000181402 | -0,940387278 | 0,19 | 0,314 | 1 | 5 |
| *whd.3* | 0,00020342 | -0,698796572 | 0,392 | 0,523 | 1 | 5 |
| *AdamTS-A.4* | 0,000210326 | -1,319935981 | 0,158 | 0,277 | 1 | 5 |
| *CG32486.3* | 0,000215212 | -0,76784345 | 0,373 | 0,482 | 1 | 5 |
| *Spn88Eb.2* | 0,000217083 | 0,410315315 | 0,323 | 0,195 | 1 | 5 |
| *CG10077.2* | 0,000220933 | -0,494196458 | 0,563 | 0,63 | 1 | 5 |
| *CG10799.2* | 0,000221283 | 0,448856781 | 0,639 | 0,496 | 1 | 5 |
| *Mvl.1* | 0,000223427 | 0,491937607 | 0,291 | 0,173 | 1 | 5 |
| *Cirl.2* | 0,00022673 | 0,399257816 | 0,506 | 0,361 | 1 | 5 |
| *CG40160* | 0,000237006 | 0,365717492 | 0,259 | 0,146 | 1 | 5 |
| *Pp1alpha-96A.2* | 0,000250017 | -0,772354936 | 0,519 | 0,592 | 1 | 5 |
| *dco.3* | 0,000259875 | 0,511644608 | 0,43 | 0,29 | 1 | 5 |
| *cv-2.2* | 0,000265733 | -0,715813221 | 0,373 | 0,481 | 1 | 5 |
| *CG42674.2* | 0,000268859 | 0,512297142 | 0,519 | 0,393 | 1 | 5 |
| *ALiX.3* | 0,000284364 | 0,425166191 | 0,348 | 0,219 | 1 | 5 |
| *14-3-3zeta.1* | 0,000289517 | 0,359842622 | 0,741 | 0,562 | 1 | 5 |
| *pes.2* | 0,000291718 | 0,430871177 | 0,475 | 0,334 | 1 | 5 |
| *hppy* | 0,000303482 | 0,419204969 | 0,652 | 0,513 | 1 | 5 |
| *CG14245* | 0,000306606 | 0,416130513 | 0,272 | 0,16 | 1 | 5 |
| *Eb1.3* | 0,000322393 | 0,442943004 | 0,373 | 0,242 | 1 | 5 |
| *GstE12.4* | 0,000323851 | -0,596499996 | 0,133 | 0,259 | 1 | 5 |
| *Gapdh2* | 0,000339942 | 0,435181223 | 0,354 | 0,236 | 1 | 5 |
| *CG7324.1* | 0,000340046 | 0,393482496 | 0,405 | 0,267 | 1 | 5 |
| *Lmpt.2* | 0,000347496 | 0,434240785 | 0,563 | 0,447 | 1 | 5 |
| *BomT3.4* | 0,000348797 | -1,21129109 | 0,329 | 0,432 | 1 | 5 |
| *larp.3* | 0,000358862 | -0,615157834 | 0,494 | 0,591 | 1 | 5 |
| *Nuak1.3* | 0,000360553 | 0,521314101 | 0,399 | 0,274 | 1 | 5 |
| *CG33144.2* | 0,000368831 | 0,26100501 | 0,367 | 0,231 | 1 | 5 |
| *ttv.1* | 0,000376882 | 0,337237331 | 0,456 | 0,304 | 1 | 5 |
| *PhKgamma.3* | 0,000378147 | -0,690634998 | 0,551 | 0,608 | 1 | 5 |
| *Ubc6.3* | 0,000394881 | 0,431650516 | 0,399 | 0,263 | 1 | 5 |
| *CG1943.3* | 0,000399628 | 0,451761186 | 0,329 | 0,206 | 1 | 5 |
| *CanA-14F.1* | 0,000408418 | 0,434947152 | 0,57 | 0,426 | 1 | 5 |
| *svp.3* | 0,000432243 | 0,38070953 | 0,639 | 0,522 | 1 | 5 |
| *mop.2* | 0,000442094 | 0,459699036 | 0,285 | 0,179 | 1 | 5 |
| *frma.3* | 0,000479228 | -0,777884087 | 0,177 | 0,289 | 1 | 5 |
| *scny* | 0,000498048 | 0,408656096 | 0,329 | 0,206 | 1 | 5 |
| *Gclm.4* | 0,000505584 | -0,947921453 | 0,259 | 0,367 | 1 | 5 |
| *CG5151.3* | 0,000530046 | -1,02287541 | 0,614 | 0,655 | 1 | 5 |
| *CG5955.3* | 0,000533309 | 0,415835715 | 0,316 | 0,197 | 1 | 5 |
| *CG3662* | 0,000538037 | 0,361783029 | 0,31 | 0,2 | 1 | 5 |
| *CG5910.4* | 0,00056074 | -0,625848519 | 0,272 | 0,405 | 1 | 5 |
| *Pits* | 0,000579828 | -0,57325772 | 0,335 | 0,454 | 1 | 5 |
| *Scamp.2* | 0,000588955 | 0,476475708 | 0,367 | 0,247 | 1 | 5 |

| | | | | | | |
|---|---|---|---|---|---|---|
| *CG3164.3* | 0,000637528 | 0,380398947 | 0,639 | 0,513 | 1 | 5 |
| *CG17574.1* | 0,000644429 | 0,656796418 | 0,354 | 0,243 | 1 | 5 |
| *CG31769* | 0,000701153 | 0,37820491 | 0,373 | 0,244 | 1 | 5 |
| *sra.3* | 0,000730418 | 0,364898017 | 0,399 | 0,263 | 1 | 5 |
| *RN-tre.2* | 0,000753983 | 0,378400979 | 0,323 | 0,209 | 1 | 5 |
| *zormin.3* | 0,000754764 | 0,391782016 | 0,551 | 0,403 | 1 | 5 |
| *Ntan1.4* | 0,000767208 | 0,344089987 | 0,766 | 0,597 | 1 | 5 |
| *His3.3B.1* | 0,000776149 | -0,498236138 | 0,278 | 0,402 | 1 | 5 |
| *CG11658* | 0,000780189 | 0,464934355 | 0,304 | 0,199 | 1 | 5 |
| *IM33.4* | 0,000790145 | -1,056387042 | 0,278 | 0,373 | 1 | 5 |
| *Syp.3* | 0,000791425 | -0,477379061 | 0,829 | 0,814 | 1 | 5 |
| *Ist1.3* | 0,000803037 | 0,383859422 | 0,424 | 0,289 | 1 | 5 |
| *MCU.3* | 0,000819936 | 0,500810012 | 0,405 | 0,279 | 1 | 5 |
| *mamo.4* | 0,000821228 | -0,724557029 | 0,494 | 0,559 | 1 | 5 |
| *CG43658.2* | 0,000822153 | -0,612502535 | 0,627 | 0,677 | 1 | 5 |
| *teq.4* | 0,000870537 | -0,745452981 | 0,146 | 0,255 | 1 | 5 |
| *Npl4.4* | 0,000910514 | 0,365792373 | 0,329 | 0,21 | 1 | 5 |
| *Hnf4.3* | 0,000971467 | -0,49742872 | 0,582 | 0,648 | 1 | 5 |
| *Fkbp12* | 0,000978257 | -0,546362567 | 0,152 | 0,261 | 1 | 5 |
| *ced-6.3* | 0,001003102 | -0,4092979 | 0,361 | 0,484 | 1 | 5 |
| *CG46385.3* | 0,001050375 | 0,310031025 | 0,975 | 0,956 | 1 | 5 |
| *MSBP.3* | 0,001071216 | 0,342944955 | 0,399 | 0,28 | 1 | 5 |
| *kst.4* | 0,001072007 | -0,839518649 | 0,184 | 0,293 | 1 | 5 |
| *CG33494.4* | 0,00108736 | 0,404552605 | 0,437 | 0,314 | 1 | 5 |
| *wdb.3* | 0,001102565 | -0,398733196 | 0,563 | 0,656 | 1 | 5 |
| *Pli.3* | 0,001118286 | -0,616667726 | 0,475 | 0,569 | 1 | 5 |
| *Rpt5.3* | 0,001132136 | 0,274314171 | 0,323 | 0,206 | 1 | 5 |
| *alpha-Spec.2* | 0,00115078 | 0,392869739 | 0,354 | 0,235 | 1 | 5 |
| *CG12116.4* | 0,001159732 | -0,674199757 | 0,31 | 0,418 | 1 | 5 |
| *CG30015.3* | 0,001276632 | 0,313341976 | 0,88 | 0,784 | 1 | 5 |
| *spin.3* | 0,00138954 | -0,601135011 | 0,557 | 0,633 | 1 | 5 |
| *CG42588.3* | 0,001390428 | 0,272215548 | 0,449 | 0,311 | 1 | 5 |
| *Pgm1.4* | 0,001396694 | -0,678117043 | 0,158 | 0,26 | 1 | 5 |
| *sgg.2* | 0,001416073 | -0,315962223 | 0,943 | 0,949 | 1 | 5 |
| *CG1640.3* | 0,001565377 | -0,551681687 | 0,165 | 0,274 | 1 | 5 |
| *mura.3* | 0,001588356 | -0,5506858 | 0,424 | 0,506 | 1 | 5 |
| *4E-T* | 0,001599203 | 0,332866905 | 0,285 | 0,181 | 1 | 5 |
| *MFS14.3* | 0,001660726 | -0,624949619 | 0,304 | 0,406 | 1 | 5 |
| *robo2.3* | 0,001691693 | 0,700537047 | 0,5 | 0,418 | 1 | 5 |
| *RpL7.3* | 0,001703384 | 0,349500091 | 0,709 | 0,601 | 1 | 5 |
| *sd.1* | 0,001770526 | 0,405785931 | 0,266 | 0,164 | 1 | 5 |
| *Rga* | 0,001791935 | 0,25547648 | 0,291 | 0,184 | 1 | 5 |
| *ogre.3* | 0,001844096 | -0,580041872 | 0,158 | 0,268 | 1 | 5 |
| *CG15098.2* | 0,00192948 | 0,373539726 | 0,5 | 0,382 | 1 | 5 |
| *Fer1HCH.3* | 0,001944784 | -0,489606887 | 0,81 | 0,802 | 1 | 5 |
| *Akap200.4* | 0,001973339 | -0,685174934 | 0,506 | 0,557 | 1 | 5 |
| *Tm1.1* | 0,001985395 | -0,437052115 | 0,551 | 0,632 | 1 | 5 |
| *CG42524.4* | 0,00199908 | -0,773638152 | 0,411 | 0,474 | 1 | 5 |
| *LRP1.3* | 0,001999476 | -0,785585778 | 0,335 | 0,43 | 1 | 5 |
| *CG17278.3* | 0,002036454 | 0,488367431 | 0,399 | 0,286 | 1 | 5 |
| *Sxl.1* | 0,002174318 | 0,334754049 | 0,867 | 0,798 | 1 | 5 |
| *Gel.3* | 0,002214016 | -0,608985691 | 0,38 | 0,466 | 1 | 5 |
| *cnn.3* | 0,002576788 | 0,268832206 | 0,399 | 0,27 | 1 | 5 |
| *Fs(2)Ket* | 0,002597774 | 0,289046926 | 0,329 | 0,22 | 1 | 5 |
| *Ac76E.2* | 0,002786749 | -0,625787726 | 0,278 | 0,377 | 1 | 5 |
| *Idgf1.2* | 0,002887366 | 0,437280846 | 0,386 | 0,279 | 1 | 5 |
| *raw.3* | 0,00293042 | 0,250691651 | 0,772 | 0,627 | 1 | 5 |
| *cpx.2* | 0,003022422 | 0,260412257 | 0,456 | 0,328 | 1 | 5 |
| *CG14478.2* | 0,003144383 | 0,268109418 | 0,671 | 0,504 | 1 | 5 |
| *lectin-28C.1* | 0,00316254 | -0,62214718 | 0,234 | 0,335 | 1 | 5 |
| *CG9044.2* | 0,003178311 | -0,602796915 | 0,184 | 0,281 | 1 | 5 |

| | | | | | | |
|---|---|---|---|---|---|---|
| *CG34325* | 0,003392627 | 0,364798573 | 0,348 | 0,242 | 1 | 5 |
| *CrebB.3* | 0,003404237 | 0,388108386 | 0,354 | 0,255 | 1 | 5 |
| *CG32066.3* | 0,003570682 | 0,403668819 | 0,57 | 0,43 | 1 | 5 |
| *Tango1.3* | 0,003599617 | 0,331677823 | 0,31 | 0,208 | 1 | 5 |
| *CG33229.1* | 0,003699332 | 0,283218287 | 0,791 | 0,676 | 1 | 5 |
| *lilli.3* | 0,003949999 | -0,329965194 | 0,646 | 0,736 | 1 | 5 |
| *sowah.3* | 0,004097902 | 0,448147734 | 0,424 | 0,324 | 1 | 5 |
| *jvl.3* | 0,004105085 | 0,334447544 | 0,823 | 0,746 | 1 | 5 |
| *Pfrx.4* | 0,00418758 | -0,558326219 | 0,203 | 0,301 | 1 | 5 |
| *CG6503.3* | 0,004271962 | -0,667584535 | 0,797 | 0,8 | 1 | 5 |
| *tim.3* | 0,004611145 | -0,450359074 | 0,608 | 0,629 | 1 | 5 |
| *PKD.4* | 0,004850586 | -0,653158735 | 0,209 | 0,295 | 1 | 5 |
| *CG9691.3* | 0,004908081 | -0,579358918 | 0,247 | 0,348 | 1 | 5 |
| *dnr1.2* | 0,005002525 | 0,572268082 | 0,335 | 0,241 | 1 | 5 |
| *Plc21C.2* | 0,005096931 | 0,258578053 | 0,671 | 0,568 | 1 | 5 |
| *Mef2.2* | 0,005102619 | -0,408686512 | 0,456 | 0,548 | 1 | 5 |
| *DIP-alpha.2* | 0,005775869 | 0,357061985 | 0,956 | 0,896 | 1 | 5 |
| *Pka-R1.1* | 0,005969906 | 0,350990699 | 0,367 | 0,274 | 1 | 5 |
| *CG44014.2* | 0,006034048 | 0,483902763 | 0,259 | 0,174 | 1 | 5 |
| *CG11400.4* | 0,006161826 | -0,527104239 | 0,297 | 0,39 | 1 | 5 |
| *CG6428.3* | 0,006354136 | -0,666861932 | 0,19 | 0,276 | 1 | 5 |
| *wnd.2* | 0,00637949 | 0,47968411 | 0,418 | 0,327 | 1 | 5 |
| *CG13315.3* | 0,006538174 | -0,636258228 | 0,835 | 0,821 | 1 | 5 |
| *atl* | 0,006638085 | 0,378637726 | 0,272 | 0,184 | 1 | 5 |
| *CG6870.2* | 0,006677619 | 0,298020226 | 0,291 | 0,2 | 1 | 5 |
| *Bruce.3* | 0,006794571 | 0,258569249 | 0,481 | 0,365 | 1 | 5 |
| *Hex-C.4* | 0,007064035 | -0,588308579 | 0,38 | 0,449 | 1 | 5 |
| *Pect.4* | 0,007345389 | -0,296818437 | 0,468 | 0,564 | 1 | 5 |
| *EcR.3* | 0,007412945 | -0,45786433 | 0,392 | 0,474 | 1 | 5 |
| *srp.3* | 0,007461178 | -0,387262082 | 0,576 | 0,634 | 1 | 5 |
| *eIF4EHP* | 0,007617869 | 0,26301863 | 0,62 | 0,496 | 1 | 5 |
| *UK114.3* | 0,007641499 | -0,618003492 | 0,348 | 0,424 | 1 | 5 |
| *cta.2* | 0,007642961 | 0,294056076 | 0,563 | 0,444 | 1 | 5 |
| *HDAC6.3* | 0,007821631 | -0,615510938 | 0,272 | 0,344 | 1 | 5 |
| *CG11873* | 0,007951365 | 0,316337517 | 0,449 | 0,353 | 1 | 5 |
| *DnaJ-1.4* | 0,008107104 | -0,545141069 | 0,481 | 0,537 | 1 | 5 |
| *BomS3.3* | 0,008370377 | -0,76596456 | 0,348 | 0,437 | 1 | 5 |
| *ATPCL.3* | 0,008380644 | -0,585740969 | 0,335 | 0,414 | 1 | 5 |
| *krz.2* | 0,008713632 | 0,399879592 | 0,304 | 0,221 | 1 | 5 |
| *cg* | 0,009238996 | 0,286231773 | 0,323 | 0,22 | 1 | 5 |
| *Sfxn1-3.3* | 0,009306328 | 0,308218364 | 0,475 | 0,357 | 1 | 5 |
| *dsb* | 0 | 5,417262648 | 0,867 | 0,016 | 0 | 6 |
| *disco* | 8,03E-197 | 2,603684116 | 0,571 | 0,017 | 9,57E-193 | 6 |
| *hth.1* | 1,85E-184 | 4,946343915 | 1 | 0,097 | 2,21E-180 | 6 |
| *slo* | 1,05E-178 | 4,974129109 | 0,765 | 0,048 | 1,25E-174 | 6 |
| *disco-r.1* | 5,64E-165 | 3,595510729 | 0,704 | 0,041 | 6,72E-161 | 6 |
| *Rbp6* | 3,03E-96 | 3,258797981 | 0,796 | 0,118 | 3,61E-92 | 6 |
| *CG3961* | 7,69E-81 | 2,342157342 | 0,531 | 0,052 | 9,18E-77 | 6 |
| *Nost* | 6,94E-80 | 3,22235515 | 0,765 | 0,131 | 8,28E-76 | 6 |
| *CG17108.4* | 1,08E-65 | 3,187514283 | 0,98 | 0,382 | 1,29E-61 | 6 |
| *apolpp.3* | 3,32E-57 | 2,886303147 | 1 | 0,798 | 3,96E-53 | 6 |
| *ome.1* | 1,19E-49 | 2,908296409 | 0,582 | 0,114 | 1,41E-45 | 6 |
| *Ubx.4* | 3,54E-49 | -4,865387867 | 0,051 | 0,882 | 4,22E-45 | 6 |
| *CG14669* | 8,44E-49 | 1,744756916 | 0,347 | 0,037 | 1,01E-44 | 6 |
| *Octbeta2R* | 1,23E-47 | 2,344827741 | 0,449 | 0,066 | 1,46E-43 | 6 |
| *fz2.2* | 1,67E-45 | 2,758425826 | 0,776 | 0,265 | 1,99E-41 | 6 |
| *CG42524.5* | 5,98E-44 | 2,047035369 | 0,959 | 0,455 | 7,13E-40 | 6 |
| *LpR2.4* | 6,07E-44 | -4,276228788 | 0,163 | 0,869 | 7,25E-40 | 6 |
| *l(2)41Ab.3* | 9,49E-38 | 2,456837417 | 0,847 | 0,35 | 1,13E-33 | 6 |
| *ap.3* | 1,85E-37 | 2,204668227 | 0,827 | 0,328 | 2,20E-33 | 6 |
| *kek5.4* | 3,75E-34 | 1,995269198 | 0,918 | 0,591 | 4,47E-30 | 6 |

| | | | | | | |
|---|---|---|---|---|---|---|
| *CG7470.3* | 3,95E-34 | 1,547913977 | 0,98 | 0,72 | 4,71E-30 | 6 |
| *CG34347.5* | 9,63E-31 | 1,9547168 | 0,847 | 0,363 | 1,15E-26 | 6 |
| *cpx.3* | 2,38E-30 | 1,909111633 | 0,765 | 0,32 | 2,83E-26 | 6 |
| *InR.4* | 8,66E-29 | -2,23642622 | 0,602 | 0,923 | 1,03E-24 | 6 |
| *CG17124.5* | 7,82E-28 | 1,522667502 | 0,949 | 0,742 | 9,33E-24 | 6 |
| *alpha-Est9.1* | 5,20E-27 | 1,646388178 | 0,582 | 0,17 | 6,20E-23 | 6 |
| *CG14762.3* | 8,74E-27 | 1,818770315 | 0,776 | 0,381 | 1,04E-22 | 6 |
| *CG30015.4* | 1,05E-25 | -2,234897281 | 0,306 | 0,804 | 1,25E-21 | 6 |
| *CARPB.1* | 3,65E-25 | 1,715112426 | 0,582 | 0,19 | 4,35E-21 | 6 |
| *BomT3.5* | 5,34E-25 | 1,598599895 | 0,837 | 0,413 | 6,37E-21 | 6 |
| *Nplp2.5* | 9,87E-25 | 1,548637145 | 0,99 | 0,658 | 1,18E-20 | 6 |
| *DIP-lambda.3* | 1,06E-24 | 2,226938693 | 0,653 | 0,263 | 1,27E-20 | 6 |
| *Ih* | 1,17E-24 | 1,56485837 | 0,459 | 0,123 | 1,40E-20 | 6 |
| *sbb.4* | 3,39E-24 | 1,398645599 | 0,898 | 0,579 | 4,05E-20 | 6 |
| *IM4.5* | 4,72E-23 | 1,584038829 | 0,704 | 0,294 | 5,64E-19 | 6 |
| *CG5953.3* | 6,36E-23 | -3,044195252 | 0,194 | 0,695 | 7,59E-19 | 6 |
| *pyd.3* | 7,13E-23 | 1,55946415 | 0,857 | 0,559 | 8,50E-19 | 6 |
| *CG6503.4* | 4,22E-22 | 1,108971776 | 0,98 | 0,794 | 5,03E-18 | 6 |
| *CG32521.3* | 8,79E-22 | 1,071768346 | 0,969 | 0,763 | 1,05E-17 | 6 |
| *IP3K1.4* | 1,08E-21 | -3,086737006 | 0,041 | 0,575 | 1,29E-17 | 6 |
| *Mmp2.3* | 1,49E-21 | 1,583544464 | 0,571 | 0,203 | 1,78E-17 | 6 |
| *osp.4* | 2,67E-21 | 1,697715593 | 0,673 | 0,307 | 3,18E-17 | 6 |
| *Nmdmc.4* | 2,86E-21 | -2,681018836 | 0,184 | 0,659 | 3,41E-17 | 6 |
| *CG32647.3* | 3,81E-21 | 1,645381674 | 0,827 | 0,556 | 4,55E-17 | 6 |
| *puc.5* | 4,66E-21 | -2,709325533 | 0,276 | 0,707 | 5,56E-17 | 6 |
| *comm3* | 9,11E-21 | 1,733945962 | 0,388 | 0,102 | 1,09E-16 | 6 |
| *Lst.4* | 1,01E-20 | -1,924765366 | 0,398 | 0,806 | 1,21E-16 | 6 |
| *bmm.4* | 1,80E-20 | -1,865245001 | 0,347 | 0,776 | 2,15E-16 | 6 |
| *CG4822* | 2,75E-20 | 1,550897758 | 0,439 | 0,132 | 3,27E-16 | 6 |
| *CG7720.5* | 4,10E-20 | 1,475012073 | 0,724 | 0,344 | 4,89E-16 | 6 |
| *GLaz* | 8,41E-20 | 1,35807561 | 0,296 | 0,063 | 1,00E-15 | 6 |
| *alpha-Est7* | 2,61E-19 | 1,434733112 | 0,429 | 0,133 | 3,11E-15 | 6 |
| *Ubi-p63E.4* | 2,85E-19 | -2,532701244 | 0,153 | 0,627 | 3,40E-15 | 6 |
| *CG31689.2* | 7,49E-19 | 1,218231469 | 0,898 | 0,67 | 8,93E-15 | 6 |
| *GstD1.3* | 1,19E-18 | -2,588470754 | 0,214 | 0,667 | 1,43E-14 | 6 |
| *bru1.3* | 1,34E-18 | 1,063699292 | 0,969 | 0,794 | 1,60E-14 | 6 |
| *CG14207.4* | 2,26E-18 | -2,659864843 | 0,143 | 0,601 | 2,70E-14 | 6 |
| *Ac13E.3* | 2,69E-18 | 1,692067692 | 0,571 | 0,25 | 3,21E-14 | 6 |
| *px.3* | 5,93E-18 | 1,250171104 | 0,908 | 0,726 | 7,07E-14 | 6 |
| *Obp99b.1* | 7,46E-18 | 1,642441429 | 0,398 | 0,118 | 8,90E-14 | 6 |
| *CG10960.4* | 1,05E-17 | -1,437512496 | 0,439 | 0,845 | 1,25E-13 | 6 |
| *mtd.4* | 1,59E-17 | 1,102543486 | 0,918 | 0,684 | 1,90E-13 | 6 |
| *Stat92E.5* | 2,09E-17 | 1,555569193 | 0,827 | 0,582 | 2,49E-13 | 6 |
| *Dmtn.4* | 2,17E-17 | 1,025580791 | 0,969 | 0,742 | 2,58E-13 | 6 |
| *CG10383.3* | 2,77E-17 | -3,16139122 | 0,031 | 0,486 | 3,30E-13 | 6 |
| *Syp.4* | 3,10E-17 | 0,930141904 | 0,959 | 0,81 | 3,70E-13 | 6 |
| *CG11089.4* | 3,47E-17 | -2,528188629 | 0,296 | 0,672 | 4,14E-13 | 6 |
| *Pdp1.5* | 4,32E-17 | -0,892615964 | 0,99 | 0,993 | 5,15E-13 | 6 |
| *Xrp1.3* | 5,32E-17 | -1,385811619 | 0,735 | 0,916 | 6,34E-13 | 6 |
| *Sarm.1* | 5,39E-17 | 1,632439228 | 0,806 | 0,565 | 6,43E-13 | 6 |
| *ssp7.3* | 6,49E-17 | 1,318815394 | 0,755 | 0,448 | 7,75E-13 | 6 |
| *Egfr.3* | 1,43E-16 | 0,904841834 | 0,959 | 0,768 | 1,71E-12 | 6 |
| *CG9928* | 2,18E-16 | 1,43913614 | 0,418 | 0,138 | 2,60E-12 | 6 |
| *CG10184* | 5,60E-16 | 1,333081482 | 0,265 | 0,062 | 6,68E-12 | 6 |
| *Smg5.3* | 1,05E-15 | -2,325995022 | 0,071 | 0,499 | 1,25E-11 | 6 |
| *Eglp4.5* | 1,28E-15 | 1,216858914 | 0,704 | 0,359 | 1,52E-11 | 6 |
| *Glt* | 1,33E-15 | 1,392703957 | 0,429 | 0,147 | 1,58E-11 | 6 |
| *AnxB9.5* | 1,58E-15 | -3,070322833 | 0,092 | 0,509 | 1,89E-11 | 6 |
| *CG2736* | 2,69E-15 | 1,329879007 | 0,357 | 0,108 | 3,20E-11 | 6 |
| *lola.3* | 3,20E-15 | 0,801064225 | 1 | 0,859 | 3,82E-11 | 6 |
| *CG1673.5* | 3,76E-15 | -2,039129641 | 0,204 | 0,598 | 4,48E-11 | 6 |

| | | | | | | |
|---|---|---|---|---|---|---|
| *eIF2beta.3* | 4,71E-15 | -2,392025396 | 0,143 | 0,54 | 5,61E-11 | 6 |
| *Helz.4* | 4,75E-15 | -1,438379206 | 0,694 | 0,916 | 5,67E-11 | 6 |
| *Pfas.3* | 6,08E-15 | -2,201131692 | 0,184 | 0,578 | 7,25E-11 | 6 |
| *Tsp42Ed.4* | 7,35E-15 | -2,431703903 | 0,041 | 0,449 | 8,77E-11 | 6 |
| *BomBc3* | 1,29E-14 | 1,243613611 | 0,306 | 0,084 | 1,54E-10 | 6 |
| *babos* | 1,65E-14 | 1,498918092 | 0,378 | 0,128 | 1,97E-10 | 6 |
| *LRP1.4* | 2,12E-14 | 1,471578872 | 0,704 | 0,416 | 2,53E-10 | 6 |
| *Chchd2.4* | 2,13E-14 | -1,800928468 | 0,255 | 0,629 | 2,54E-10 | 6 |
| *CG5773.4* | 6,76E-14 | 1,43145099 | 0,51 | 0,222 | 8,06E-10 | 6 |
| *CG6040.1* | 9,97E-14 | 1,526451999 | 0,408 | 0,149 | 1,19E-09 | 6 |
| *CG16898.3* | 1,31E-13 | -2,610611973 | 0,071 | 0,449 | 1,56E-09 | 6 |
| *trbl.4* | 1,53E-13 | -2,018905092 | 0,143 | 0,524 | 1,82E-09 | 6 |
| *Eaat1.3* | 1,58E-13 | 1,551145636 | 0,612 | 0,327 | 1,88E-09 | 6 |
| *MFS17.3* | 2,75E-13 | 1,120132987 | 0,867 | 0,69 | 3,28E-09 | 6 |
| *CG41378.3* | 4,90E-13 | 1,128835263 | 0,786 | 0,536 | 5,84E-09 | 6 |
| *pHCl-2.2* | 5,09E-13 | 1,247747602 | 0,418 | 0,159 | 6,07E-09 | 6 |
| *rgn.4* | 7,91E-13 | -2,213165289 | 0,082 | 0,472 | 9,43E-09 | 6 |
| *IM14.2* | 8,38E-13 | 0,97787661 | 0,48 | 0,2 | 9,99E-09 | 6 |
| *Tsf1.5* | 8,94E-13 | 1,291889631 | 0,582 | 0,285 | 1,07E-08 | 6 |
| *Oatp74D.3* | 1,09E-12 | 1,076786385 | 0,643 | 0,352 | 1,30E-08 | 6 |
| *AkhR.5* | 1,09E-12 | 1,08483188 | 0,602 | 0,31 | 1,31E-08 | 6 |
| *Atg8a.3* | 1,14E-12 | -1,367805791 | 0,388 | 0,704 | 1,36E-08 | 6 |
| *h.3* | 1,24E-12 | -1,575814077 | 0,265 | 0,646 | 1,48E-08 | 6 |
| *phu.1* | 1,92E-12 | 0,985959633 | 0,367 | 0,125 | 2,28E-08 | 6 |
| *GIIIspla2.3* | 2,06E-12 | -2,360921126 | 0,071 | 0,428 | 2,46E-08 | 6 |
| *vri.4* | 2,09E-12 | -1,793060782 | 0,143 | 0,512 | 2,50E-08 | 6 |
| *ref(2)P.3* | 2,24E-12 | -1,872520651 | 0,327 | 0,646 | 2,67E-08 | 6 |
| *CG12795.4* | 2,41E-12 | -2,734431207 | 0,01 | 0,36 | 2,88E-08 | 6 |
| *CG32369.4* | 2,94E-12 | -1,524800196 | 0,347 | 0,683 | 3,51E-08 | 6 |
| *raw.4* | 4,08E-12 | -1,609154072 | 0,306 | 0,645 | 4,86E-08 | 6 |
| *pum.3* | 5,60E-12 | 1,215065494 | 0,735 | 0,531 | 6,68E-08 | 6 |
| *GEFmeso.4* | 5,76E-12 | -1,988483155 | 0,092 | 0,45 | 6,87E-08 | 6 |
| *Treh.1* | 5,77E-12 | 1,561352309 | 0,449 | 0,199 | 6,89E-08 | 6 |
| *mamo.5* | 9,15E-12 | 0,910900704 | 0,816 | 0,547 | 1,09E-07 | 6 |
| *Pli.4* | 9,90E-12 | 1,249762031 | 0,796 | 0,557 | 1,18E-07 | 6 |
| *Gpdh1.5* | 1,07E-11 | 0,972630819 | 0,653 | 0,353 | 1,28E-07 | 6 |
| *PCB.4* | 1,56E-11 | 1,016151035 | 0,806 | 0,606 | 1,86E-07 | 6 |
| *Fer2LCH.4* | 1,85E-11 | -1,323456036 | 0,48 | 0,753 | 2,20E-07 | 6 |
| *scyl.3* | 2,00E-11 | -1,258443613 | 0,561 | 0,816 | 2,39E-07 | 6 |
| *Frl.5* | 2,03E-11 | -1,82298598 | 0,173 | 0,525 | 2,42E-07 | 6 |
| *bnl.4* | 2,96E-11 | -3,046721177 | 0,01 | 0,338 | 3,54E-07 | 6 |
| *CG43340.3* | 3,04E-11 | 0,952999893 | 0,684 | 0,406 | 3,63E-07 | 6 |
| *svp.4* | 3,81E-11 | 1,0566324 | 0,724 | 0,521 | 4,54E-07 | 6 |
| *AdSS.3* | 4,22E-11 | -1,939948468 | 0,092 | 0,426 | 5,04E-07 | 6 |
| *Atg18b.3* | 4,46E-11 | -1,771638566 | 0,112 | 0,454 | 5,32E-07 | 6 |
| *BomBc2.3* | 5,77E-11 | 1,136620927 | 0,765 | 0,551 | 6,89E-07 | 6 |
| *stv.3* | 6,15E-11 | -2,184943631 | 0,327 | 0,628 | 7,34E-07 | 6 |
| *CG42588.4* | 6,72E-11 | -2,151564523 | 0,01 | 0,328 | 8,02E-07 | 6 |
| *Lip4.3* | 6,86E-11 | -1,882114128 | 0,082 | 0,418 | 8,18E-07 | 6 |
| *Hsp26.4* | 6,90E-11 | -3,07546829 | 0,102 | 0,431 | 8,23E-07 | 6 |
| *MtnA.4* | 6,92E-11 | -1,856246091 | 0,255 | 0,578 | 8,25E-07 | 6 |
| *BomT2* | 7,43E-11 | 1,220724409 | 0,429 | 0,196 | 8,87E-07 | 6 |
| *CG5966.3* | 9,00E-11 | -1,700855655 | 0,112 | 0,463 | 1,07E-06 | 6 |
| *mino.3* | 9,12E-11 | -1,093072067 | 0,337 | 0,686 | 1,09E-06 | 6 |
| *Pp1alpha-96A.3* | 9,52E-11 | -1,492084106 | 0,276 | 0,598 | 1,14E-06 | 6 |
| *CG31145.3* | 1,06E-10 | 1,008666689 | 0,867 | 0,729 | 1,27E-06 | 6 |
| *Est-6* | 1,28E-10 | 1,044049385 | 0,265 | 0,083 | 1,53E-06 | 6 |
| *CG3376.2* | 1,37E-10 | -1,420961176 | 0,255 | 0,585 | 1,64E-06 | 6 |
| *sdt* | 1,41E-10 | 1,405448132 | 0,429 | 0,196 | 1,68E-06 | 6 |
| *cic.5* | 1,53E-10 | 0,877464704 | 0,847 | 0,659 | 1,82E-06 | 6 |
| *GstE1.3* | 1,63E-10 | -2,11974131 | 0,061 | 0,387 | 1,95E-06 | 6 |

| | | | | | | |
|---|---|---|---|---|---|---|
| *Gp150.2* | 1,77E-10 | -1,405655064 | 0,204 | 0,534 | 2,11E-06 | 6 |
| *cwo.4* | 2,19E-10 | -0,964711402 | 0,735 | 0,884 | 2,61E-06 | 6 |
| *GstT4.3* | 2,25E-10 | -1,9043166 | 0,031 | 0,351 | 2,68E-06 | 6 |
| *robo2.4* | 2,29E-10 | -1,811359351 | 0,102 | 0,433 | 2,73E-06 | 6 |
| *cher.4* | 2,48E-10 | -2,374178052 | 0 | 0,301 | 2,96E-06 | 6 |
| *CG16926.4* | 2,81E-10 | -1,216818504 | 0,306 | 0,633 | 3,35E-06 | 6 |
| *oys.5* | 2,86E-10 | -2,278399227 | 0,041 | 0,341 | 3,41E-06 | 6 |
| *Tab2.3* | 3,93E-10 | -1,555381991 | 0,092 | 0,415 | 4,69E-06 | 6 |
| *Chd64.4* | 4,44E-10 | -1,711411465 | 0,143 | 0,466 | 5,29E-06 | 6 |
| *Fur1.2* | 4,56E-10 | 0,784652737 | 0,867 | 0,713 | 5,44E-06 | 6 |
| *rhea.4* | 4,73E-10 | -2,018342923 | 0,184 | 0,504 | 5,64E-06 | 6 |
| *Xbp1.3* | 5,59E-10 | -1,293631146 | 0,255 | 0,584 | 6,67E-06 | 6 |
| *Rpn6.3* | 6,16E-10 | -1,559493947 | 0,082 | 0,399 | 7,35E-06 | 6 |
| *pyr.3* | 6,62E-10 | -2,082497788 | 0,01 | 0,306 | 7,90E-06 | 6 |
| *Act5C.5* | 7,57E-10 | -1,730095746 | 0,276 | 0,555 | 9,03E-06 | 6 |
| *CG45050.3* | 1,03E-09 | -1,09352464 | 0,806 | 0,905 | 1,23E-05 | 6 |
| *Pisd.1* | 1,17E-09 | 1,087298498 | 0,347 | 0,141 | 1,40E-05 | 6 |
| *BomS2.2* | 1,53E-09 | 1,07621898 | 0,612 | 0,388 | 1,83E-05 | 6 |
| *CCHa2.5* | 1,55E-09 | 1,163634862 | 0,5 | 0,255 | 1,85E-05 | 6 |
| *goe.5* | 1,62E-09 | -1,966408362 | 0 | 0,28 | 1,93E-05 | 6 |
| *CG8468.2* | 1,71E-09 | -1,183678921 | 0,367 | 0,684 | 2,03E-05 | 6 |
| *CG5793* | 1,75E-09 | 1,224835518 | 0,255 | 0,087 | 2,09E-05 | 6 |
| *sdk.1* | 1,78E-09 | 1,109206433 | 0,418 | 0,196 | 2,13E-05 | 6 |
| *Gdh.4* | 1,95E-09 | -1,819367346 | 0,092 | 0,394 | 2,33E-05 | 6 |
| *Iml1* | 2,26E-09 | 1,240899794 | 0,357 | 0,157 | 2,70E-05 | 6 |
| *CG13784.3* | 2,49E-09 | 0,987679917 | 0,755 | 0,614 | 2,97E-05 | 6 |
| *Tpr2.3* | 3,18E-09 | -1,369413277 | 0,316 | 0,606 | 3,80E-05 | 6 |
| *Culd.5* | 3,36E-09 | 0,823294726 | 0,602 | 0,336 | 4,01E-05 | 6 |
| *CG5151.4* | 3,47E-09 | 0,86880089 | 0,837 | 0,647 | 4,14E-05 | 6 |
| *MTA1-like.4* | 3,73E-09 | -2,265145339 | 0,153 | 0,442 | 4,44E-05 | 6 |
| *chic.4* | 4,21E-09 | -1,659943591 | 0,133 | 0,427 | 5,02E-05 | 6 |
| *CG31751* | 5,55E-09 | 1,121687904 | 0,286 | 0,11 | 6,62E-05 | 6 |
| *CG17841.4* | 5,61E-09 | 0,683803012 | 0,724 | 0,468 | 6,69E-05 | 6 |
| *Hsp27.5* | 5,66E-09 | -3,348776551 | 0,071 | 0,35 | 6,75E-05 | 6 |
| *IM33.5* | 5,66E-09 | 0,898220581 | 0,602 | 0,361 | 6,75E-05 | 6 |
| *Fer1HCH.4* | 5,85E-09 | -1,078647336 | 0,622 | 0,808 | 6,97E-05 | 6 |
| *Ufd4.4* | 6,02E-09 | -1,535612867 | 0,071 | 0,354 | 7,18E-05 | 6 |
| *LRR.4* | 6,18E-09 | -1,307491672 | 0,224 | 0,533 | 7,37E-05 | 6 |
| *SNF4Agamma.4* | 6,52E-09 | -0,858816527 | 0,735 | 0,897 | 7,78E-05 | 6 |
| *aralar1.3* | 7,43E-09 | -1,056864903 | 0,337 | 0,644 | 8,87E-05 | 6 |
| *cv-2.3* | 7,71E-09 | -1,293561875 | 0,173 | 0,485 | 9,20E-05 | 6 |
| *alpha-Est8* | 7,95E-09 | 1,114313549 | 0,296 | 0,115 | 9,48E-05 | 6 |
| *kay.5* | 8,15E-09 | -1,595632097 | 0,245 | 0,526 | 9,72E-05 | 6 |
| *cu.3* | 9,47E-09 | -1,130318347 | 0,184 | 0,498 | 0,000112923 | 6 |
| *Mob2.5* | 1,02E-08 | -1,255604744 | 0,531 | 0,76 | 0,000121267 | 6 |
| *Cyt-b5-r.4* | 1,05E-08 | 0,794250458 | 0,786 | 0,557 | 0,000125674 | 6 |
| *Hsc70-4.4* | 1,06E-08 | -1,081963286 | 0,48 | 0,748 | 0,000126764 | 6 |
| *Rel.4* | 1,17E-08 | -1,339199377 | 0,255 | 0,546 | 0,000139315 | 6 |
| *drpr.5* | 1,22E-08 | -1,566186039 | 0,184 | 0,465 | 0,000145597 | 6 |
| *Bsg.2* | 1,60E-08 | -0,589269859 | 0,959 | 0,983 | 0,000190332 | 6 |
| *Tps1.2* | 1,61E-08 | -0,706821576 | 0,827 | 0,946 | 0,000192138 | 6 |
| *lilli.4* | 1,62E-08 | 0,719396266 | 0,847 | 0,727 | 0,000192737 | 6 |
| *CG7130.4* | 1,62E-08 | -2,06224541 | 0,02 | 0,279 | 0,000192908 | 6 |
| *prage.1* | 1,64E-08 | -0,910104259 | 0,429 | 0,749 | 0,000195853 | 6 |
| *SCaMC.4* | 1,80E-08 | -0,943598727 | 0,541 | 0,772 | 0,000214462 | 6 |
| *Plod.4* | 1,85E-08 | -1,686470823 | 0,031 | 0,291 | 0,000220964 | 6 |
| *klu.4* | 2,00E-08 | -1,29987816 | 0,194 | 0,499 | 0,000239139 | 6 |
| *mbf1.4* | 2,66E-08 | -1,644739196 | 0,082 | 0,359 | 0,000317222 | 6 |
| *Sxl.2* | 3,03E-08 | -0,918634463 | 0,592 | 0,808 | 0,000361458 | 6 |
| *loco.4* | 3,32E-08 | -1,958902645 | 0,031 | 0,282 | 0,000395994 | 6 |
| *Vps13.3* | 3,36E-08 | -1,571889387 | 0,102 | 0,371 | 0,000401285 | 6 |

| | | | | | | |
|---|---|---|---|---|---|---|
| *Mal-B2.4* | 3,94E-08 | -1,229056815 | 0,245 | 0,534 | 0,000469541 | 6 |
| *CG11594.2* | 4,65E-08 | 1,05394238 | 0,398 | 0,198 | 0,00055509 | 6 |
| *CG6910.5* | 4,70E-08 | -1,211996437 | 0,561 | 0,765 | 0,000560512 | 6 |
| *Rpn2.3* | 5,13E-08 | -1,383769161 | 0,051 | 0,315 | 0,000612138 | 6 |
| *Msp300.4* | 6,95E-08 | 0,657633706 | 0,929 | 0,88 | 0,00082884 | 6 |
| *drongo.3* | 8,48E-08 | -1,21804042 | 0,316 | 0,589 | 0,001011452 | 6 |
| *TER94.3* | 8,51E-08 | -1,288143028 | 0,235 | 0,501 | 0,001015726 | 6 |
| *CG15099.4* | 8,59E-08 | -1,393375476 | 0,051 | 0,305 | 0,001025108 | 6 |
| *Snp.3* | 9,25E-08 | -1,204872584 | 0,122 | 0,407 | 0,001103989 | 6 |
| *Trxr-1.4* | 9,47E-08 | -1,411239835 | 0,133 | 0,399 | 0,001130254 | 6 |
| *Gdap2.4* | 9,50E-08 | -1,335471857 | 0,143 | 0,415 | 0,001133531 | 6 |
| *CG16756.1* | 1,02E-07 | 0,908710576 | 0,347 | 0,157 | 0,001216129 | 6 |
| *CG14823.5* | 1,03E-07 | 0,602979851 | 0,612 | 0,337 | 0,00123225 | 6 |
| *CG34136.3* | 1,04E-07 | -1,474013551 | 0,031 | 0,281 | 0,001234777 | 6 |
| *REPTOR.4* | 1,08E-07 | -0,972392424 | 0,469 | 0,725 | 0,001288026 | 6 |
| *CG7920.5* | 1,17E-07 | 0,847436209 | 0,51 | 0,289 | 0,001393689 | 6 |
| *c11.1.4* | 1,36E-07 | -1,459235085 | 0,02 | 0,254 | 0,001626938 | 6 |
| *eas.4* | 1,39E-07 | 1,154179665 | 0,633 | 0,465 | 0,001661418 | 6 |
| *Nadsyn.3* | 1,39E-07 | 0,936160682 | 0,49 | 0,28 | 0,001663081 | 6 |
| *sra.4* | 1,47E-07 | -1,631975715 | 0,041 | 0,278 | 0,001748056 | 6 |
| *CG31694.4* | 1,68E-07 | -1,494801122 | 0,122 | 0,384 | 0,002008682 | 6 |
| *Prosalpha3.3* | 1,73E-07 | -1,584425644 | 0,041 | 0,276 | 0,002065817 | 6 |
| *Jabba.5* | 1,78E-07 | 0,705143232 | 0,51 | 0,277 | 0,002118996 | 6 |
| *bbg.3* | 1,86E-07 | 1,557929263 | 0,51 | 0,332 | 0,002220865 | 6 |
| *AdenoK* | 2,06E-07 | 0,926695222 | 0,378 | 0,19 | 0,00245338 | 6 |
| *CG6428.4* | 2,19E-07 | -1,46237974 | 0,041 | 0,279 | 0,002617437 | 6 |
| *RhoGAP19D.5* | 2,21E-07 | 0,928376201 | 0,663 | 0,476 | 0,002632444 | 6 |
| *CG6426* | 2,37E-07 | 1,305431393 | 0,469 | 0,275 | 0,0028224 | 6 |
| *MCU.4* | 2,65E-07 | -1,504928279 | 0,051 | 0,293 | 0,003161742 | 6 |
| *l(1)G0289.4* | 2,80E-07 | -1,199550664 | 0,214 | 0,481 | 0,003340599 | 6 |
| *Pdfr.3* | 3,09E-07 | 0,976337296 | 0,643 | 0,472 | 0,003686997 | 6 |
| *teq.5* | 3,15E-07 | 1,018522325 | 0,429 | 0,244 | 0,003762894 | 6 |
| *Atg1.2* | 3,59E-07 | -1,290443769 | 0,245 | 0,512 | 0,004280634 | 6 |
| *Irc.4* | 3,64E-07 | -1,018182268 | 0,255 | 0,524 | 0,004341767 | 6 |
| *sgg.3* | 4,02E-07 | -0,648423083 | 0,939 | 0,949 | 0,004795095 | 6 |
| *NaCP60E* | 4,19E-07 | 1,013505822 | 0,255 | 0,105 | 0,005000898 | 6 |
| *l(3)80Fg.3* | 4,70E-07 | 0,844565651 | 0,735 | 0,607 | 0,005608794 | 6 |
| *Dbp80.3* | 4,79E-07 | 0,548611063 | 0,827 | 0,652 | 0,005715613 | 6 |
| *Idgf1.3* | 5,22E-07 | -1,399006983 | 0,061 | 0,292 | 0,006228247 | 6 |
| *Abl.5* | 5,88E-07 | -1,622291499 | 0,143 | 0,379 | 0,007017011 | 6 |
| *Pcyt1.4* | 6,22E-07 | -1,07041599 | 0,224 | 0,489 | 0,007415066 | 6 |
| *Npc2g.3* | 6,66E-07 | 0,992452876 | 0,541 | 0,34 | 0,007946014 | 6 |
| *Gart.3* | 6,99E-07 | -1,028748672 | 0,459 | 0,672 | 0,008344291 | 6 |
| *Btk29A.2* | 7,68E-07 | 1,074704712 | 0,408 | 0,229 | 0,009166323 | 6 |
| *Thor.4* | 8,04E-07 | -0,95146171 | 0,265 | 0,546 | 0,009587428 | 6 |
| *CG44325.5* | 8,22E-07 | -1,253666221 | 0,082 | 0,325 | 0,009802467 | 6 |
| *Diap1.4* | 8,48E-07 | -0,801421005 | 0,5 | 0,756 | 0,010117102 | 6 |
| *LamC.4* | 9,09E-07 | -1,101840325 | 0,041 | 0,273 | 0,010843168 | 6 |
| *pAbp.3* | 9,10E-07 | -0,817231446 | 0,622 | 0,808 | 0,010853876 | 6 |
| *sima.3* | 9,19E-07 | 0,504776459 | 0,98 | 0,925 | 0,010966036 | 6 |
| *nkd.3* | 9,92E-07 | -1,278810022 | 0,204 | 0,456 | 0,011832689 | 6 |
| *MRP.3* | 1,09E-06 | -1,74557929 | 0,194 | 0,435 | 0,012968066 | 6 |
| *gce.5* | 1,10E-06 | 0,672330193 | 0,673 | 0,477 | 0,013163125 | 6 |
| *CG32687.3* | 1,17E-06 | -0,859411915 | 0,214 | 0,499 | 0,013914595 | 6 |
| *Exn.3* | 1,24E-06 | -0,935728936 | 0,255 | 0,515 | 0,014794126 | 6 |
| *aqz.2* | 1,43E-06 | -0,555367752 | 0,816 | 0,952 | 0,017067301 | 6 |
| *Arf79F.3* | 1,70E-06 | -0,996481296 | 0,235 | 0,492 | 0,020293783 | 6 |
| *luna.4* | 1,76E-06 | -0,753460204 | 0,49 | 0,727 | 0,020992008 | 6 |
| *sug.5* | 1,85E-06 | 0,819198308 | 0,541 | 0,318 | 0,022075912 | 6 |
| *Rpn5.3* | 1,94E-06 | -1,205818907 | 0,051 | 0,272 | 0,023114434 | 6 |
| *Lpin.3* | 1,99E-06 | -1,023887845 | 0,235 | 0,49 | 0,023721413 | 6 |

| | | | | | | |
|---|---|---|---|---|---|---|
| *cact.5* | 2,06E-06 | -1,066242198 | 0,459 | 0,647 | 0,024601492 | 6 |
| *CG1648.5* | 2,19E-06 | -1,592641462 | 0,459 | 0,616 | 0,026138165 | 6 |
| *Parp.4* | 2,25E-06 | 0,740636071 | 0,776 | 0,676 | 0,02679344 | 6 |
| *RpL35.3* | 2,26E-06 | -0,926332802 | 0,306 | 0,55 | 0,02693857 | 6 |
| *CG8079.3* | 2,32E-06 | -1,182719025 | 0,071 | 0,298 | 0,027657379 | 6 |
| *Gbs-70E.5* | 2,39E-06 | 0,583918322 | 0,765 | 0,522 | 0,028456108 | 6 |
| *Prosbeta7.3* | 2,44E-06 | -1,302673723 | 0,051 | 0,256 | 0,029112009 | 6 |
| *Akt1.3* | 2,47E-06 | -0,7685834 | 0,122 | 0,385 | 0,029439614 | 6 |
| *Socs36E.5* | 2,47E-06 | -1,759411362 | 0,112 | 0,33 | 0,029495256 | 6 |
| *CG32695.2* | 2,60E-06 | -1,190632032 | 0,143 | 0,374 | 0,031000077 | 6 |
| *CG4629.4* | 2,75E-06 | 0,93393998 | 0,429 | 0,247 | 0,032787109 | 6 |
| *Rpn10.3* | 2,76E-06 | -1,320944961 | 0,051 | 0,255 | 0,032976836 | 6 |
| *Dif.4* | 2,81E-06 | 0,899930495 | 0,551 | 0,388 | 0,033564604 | 6 |
| *CG3638.4* | 2,99E-06 | 0,677164275 | 0,653 | 0,471 | 0,035651402 | 6 |
| *par-1.4* | 3,12E-06 | -0,545216818 | 0,898 | 0,943 | 0,037267916 | 6 |
| *Su(Tpl)* | 3,19E-06 | -0,859340249 | 0,429 | 0,637 | 0,038055703 | 6 |
| *Cyp4g1.2* | 3,25E-06 | -2,400412829 | 0,163 | 0,376 | 0,038776445 | 6 |
| *Sox102F.3* | 3,27E-06 | 0,63655741 | 0,878 | 0,743 | 0,038965933 | 6 |
| *Ork1.4* | 3,63E-06 | 0,651907411 | 0,531 | 0,331 | 0,04325766 | 6 |
| *Rab7.3* | 3,71E-06 | -1,118448506 | 0,092 | 0,306 | 0,044280624 | 6 |
| *CG12065.4* | 3,82E-06 | -0,975700013 | 0,071 | 0,302 | 0,045535526 | 6 |
| *Ttd14.3* | 3,91E-06 | 1,07899643 | 0,439 | 0,276 | 0,046645942 | 6 |
| *scrib.3* | 3,94E-06 | 1,028540228 | 0,531 | 0,371 | 0,047001577 | 6 |
| *Cyp28d1.4* | 4,36E-06 | 0,835023433 | 0,388 | 0,213 | 0,052010679 | 6 |
| *LManII* | 4,90E-06 | 1,051090981 | 0,316 | 0,166 | 0,058504209 | 6 |
| *pcs.4* | 4,92E-06 | -1,444944462 | 0,204 | 0,431 | 0,058653509 | 6 |
| *Hers.5* | 5,13E-06 | -0,898736935 | 0,398 | 0,63 | 0,061197472 | 6 |
| *Rm62.2* | 5,14E-06 | -0,625640026 | 0,459 | 0,719 | 0,061320593 | 6 |
| *NAT1.4* | 5,30E-06 | -0,65205441 | 0,439 | 0,703 | 0,063221034 | 6 |
| *Pdk1.3* | 5,44E-06 | -0,749099772 | 0,531 | 0,738 | 0,064872475 | 6 |
| *Gug.5* | 5,58E-06 | -0,74253925 | 0,418 | 0,649 | 0,066612435 | 6 |
| *CG14629* | 5,61E-06 | 0,792600177 | 0,306 | 0,149 | 0,066879098 | 6 |
| *CG12012.3* | 5,91E-06 | -1,113504803 | 0,071 | 0,283 | 0,070500121 | 6 |
| *RpS8.3* | 6,23E-06 | -0,890441784 | 0,52 | 0,719 | 0,074351038 | 6 |
| *vir-1.4* | 6,42E-06 | -1,001186148 | 0,551 | 0,744 | 0,076607617 | 6 |
| *CG14478.3* | 6,96E-06 | -0,763313748 | 0,255 | 0,52 | 0,083019051 | 6 |
| *Mur2B.4* | 7,24E-06 | 0,908203353 | 0,571 | 0,431 | 0,08640862 | 6 |
| *kst.5* | 7,95E-06 | -1,181113736 | 0,082 | 0,294 | 0,094785777 | 6 |
| *nmo.4* | 7,99E-06 | -0,891527954 | 0,367 | 0,584 | 0,095286581 | 6 |
| *CG30431* | 8,50E-06 | 0,889406865 | 0,255 | 0,117 | 0,101395219 | 6 |
| *CG12163.3* | 8,70E-06 | 0,834831499 | 0,643 | 0,498 | 0,103787819 | 6 |
| *Pmp70.5* | 9,26E-06 | 0,750820554 | 0,449 | 0,265 | 0,110509833 | 6 |
| *fray.2* | 1,01E-05 | -1,153152784 | 0,071 | 0,272 | 0,120270734 | 6 |
| *BomS3.4* | 1,05E-05 | 0,721825904 | 0,561 | 0,429 | 0,124682299 | 6 |
| *Tlk.3* | 1,06E-05 | 0,684618551 | 0,684 | 0,561 | 0,126499739 | 6 |
| *shep.3* | 1,09E-05 | 0,580974298 | 0,816 | 0,746 | 0,129730216 | 6 |
| *Syb.2* | 1,13E-05 | -1,041213337 | 0,112 | 0,325 | 0,135004267 | 6 |
| *tsr.4* | 1,14E-05 | -1,05817771 | 0,061 | 0,256 | 0,135747077 | 6 |
| *Yeti.4* | 1,23E-05 | -0,850011793 | 0,398 | 0,622 | 0,146316233 | 6 |
| *crc.4* | 1,25E-05 | -0,760669874 | 0,143 | 0,381 | 0,149335976 | 6 |
| *Nmda1.4* | 1,25E-05 | -0,968973308 | 0,133 | 0,359 | 0,149386622 | 6 |
| *Rad23.4* | 1,32E-05 | -0,974502573 | 0,122 | 0,339 | 0,157002409 | 6 |
| *RpL11.4* | 1,33E-05 | -0,906216127 | 0,306 | 0,519 | 0,158753777 | 6 |
| *CtsB1.2* | 1,34E-05 | -0,890446369 | 0,286 | 0,512 | 0,159539475 | 6 |
| *Rab11.4* | 1,35E-05 | -0,918945999 | 0,153 | 0,373 | 0,161632173 | 6 |
| *pigs* | 1,63E-05 | 0,928489538 | 0,347 | 0,194 | 0,194586172 | 6 |
| *Khc-73.4* | 1,81E-05 | -0,988119953 | 0,163 | 0,38 | 0,216038295 | 6 |
| *Cyp4p1.4* | 1,89E-05 | -1,122753255 | 0,112 | 0,317 | 0,225064988 | 6 |
| *CG31183.2* | 1,93E-05 | 0,666337007 | 0,684 | 0,558 | 0,230182825 | 6 |
| *Pvr.4* | 1,95E-05 | -1,656649682 | 0,235 | 0,443 | 0,232637513 | 6 |
| *ZnT41F* | 2,04E-05 | 0,883193311 | 0,306 | 0,163 | 0,243388602 | 6 |

| | | | | | | |
|---|---|---|---|---|---|---|
| *RpL18A.4* | 2,07E-05 | -0,767809592 | 0,418 | 0,627 | 0,247324945 | 6 |
| *tai.2* | 2,07E-05 | 0,535857667 | 0,918 | 0,845 | 0,247511515 | 6 |
| *Ist1.4* | 2,08E-05 | -1,042259005 | 0,102 | 0,302 | 0,248594779 | 6 |
| *CG34325.1* | 2,09E-05 | -1,119805656 | 0,061 | 0,253 | 0,249106112 | 6 |
| *Kank.1* | 2,09E-05 | 0,952009698 | 0,306 | 0,159 | 0,249457691 | 6 |
| *Rac2.4* | 2,21E-05 | -1,047726643 | 0,112 | 0,315 | 0,2635601 | 6 |
| *Pect.5* | 2,32E-05 | -1,007036688 | 0,357 | 0,566 | 0,276797923 | 6 |
| *ftz-f1.4* | 2,54E-05 | 0,683534415 | 0,704 | 0,612 | 0,302828999 | 6 |
| *step.3* | 2,57E-05 | -0,88246818 | 0,214 | 0,443 | 0,306239155 | 6 |
| *whd.4* | 2,63E-05 | 0,892182248 | 0,653 | 0,512 | 0,313957645 | 6 |
| *Csk.3* | 2,76E-05 | -0,845831459 | 0,245 | 0,488 | 0,32879704 | 6 |
| *Pak3.4* | 2,83E-05 | -0,995751977 | 0,092 | 0,295 | 0,338112088 | 6 |
| *Usp10.2* | 2,86E-05 | -0,819797644 | 0,173 | 0,388 | 0,34162433 | 6 |
| *kra.3* | 3,18E-05 | -0,744382778 | 0,327 | 0,554 | 0,379298686 | 6 |
| *18w.1* | 3,26E-05 | 0,797848173 | 0,357 | 0,2 | 0,389405498 | 6 |
| *Larp4B.1* | 3,29E-05 | -0,579886592 | 0,469 | 0,675 | 0,392782041 | 6 |
| *AdipoR.4* | 3,73E-05 | 0,435818733 | 0,735 | 0,532 | 0,444727634 | 6 |
| *pst.4* | 3,96E-05 | -0,751986259 | 0,388 | 0,59 | 0,472815158 | 6 |
| *Nha2.1* | 3,97E-05 | 0,81746164 | 0,296 | 0,157 | 0,47354557 | 6 |
| *RpS3A.2* | 3,99E-05 | -0,556064167 | 0,327 | 0,578 | 0,476241753 | 6 |
| *Myc.3* | 4,00E-05 | -0,8342451 | 0,459 | 0,665 | 0,476752866 | 6 |
| *BomS6.1* | 4,26E-05 | 0,909709094 | 0,347 | 0,207 | 0,508664245 | 6 |
| *CG33494.5* | 4,28E-05 | -0,911399209 | 0,122 | 0,326 | 0,510468895 | 6 |
| *spir.4* | 4,37E-05 | -0,913523996 | 0,173 | 0,377 | 0,521120514 | 6 |
| *CG31705.3* | 4,43E-05 | -0,77773669 | 0,367 | 0,577 | 0,5279463 | 6 |
| *mdy.1* | 4,47E-05 | 0,979969805 | 0,296 | 0,159 | 0,532898164 | 6 |
| *elB.5* | 4,50E-05 | -1,192180564 | 0,122 | 0,317 | 0,537083635 | 6 |
| *RpL5.2* | 4,51E-05 | -0,507394441 | 0,388 | 0,604 | 0,537816546 | 6 |
| *Hsp68.4* | 4,98E-05 | -1,929106877 | 0,122 | 0,305 | 0,593973979 | 6 |
| *Letm1.4* | 4,99E-05 | -0,990991799 | 0,082 | 0,268 | 0,595289279 | 6 |
| *Droj2.3* | 5,01E-05 | -0,861112176 | 0,214 | 0,421 | 0,597067792 | 6 |
| *EcR.4* | 5,10E-05 | 0,680801643 | 0,612 | 0,466 | 0,608252909 | 6 |
| *Prps.2* | 5,27E-05 | -0,558035617 | 0,918 | 0,925 | 0,629035981 | 6 |
| *Mdh2.2* | 5,58E-05 | -0,647880088 | 0,306 | 0,549 | 0,665164993 | 6 |
| *Non1.2* | 5,63E-05 | -0,830540177 | 0,153 | 0,356 | 0,671918923 | 6 |
| *CG3902.4* | 5,86E-05 | 0,939702162 | 0,449 | 0,303 | 0,699410379 | 6 |
| *Dad.3* | 5,99E-05 | -0,724980327 | 0,255 | 0,477 | 0,71500864 | 6 |
| *CG9932.2* | 6,01E-05 | 0,347269421 | 0,99 | 0,947 | 0,717348758 | 6 |
| *if.5* | 6,04E-05 | 0,659441984 | 0,51 | 0,343 | 0,720186794 | 6 |
| *PRL-1.4* | 6,16E-05 | -0,926485881 | 0,112 | 0,305 | 0,735023234 | 6 |
| *PHGPx.3* | 6,30E-05 | -0,956778276 | 0,184 | 0,383 | 0,751901681 | 6 |
| *Hsp83.4* | 6,52E-05 | -1,163383818 | 0,622 | 0,738 | 0,778107064 | 6 |
| *kirre.1* | 6,55E-05 | 1,143506365 | 0,316 | 0,187 | 0,780963284 | 6 |
| *poe.3* | 6,93E-05 | -0,899617945 | 0,143 | 0,346 | 0,826423978 | 6 |
| *AOX1.3* | 7,01E-05 | -0,983915264 | 0,153 | 0,34 | 0,836084143 | 6 |
| *RpL8.3* | 7,12E-05 | -0,752881819 | 0,48 | 0,673 | 0,849060447 | 6 |
| *mam.3* | 7,42E-05 | 0,785320509 | 0,52 | 0,384 | 0,885021619 | 6 |
| *CG3168.4* | 7,49E-05 | -1,125767799 | 0,112 | 0,3 | 0,893084919 | 6 |
| *Atpalpha.3* | 7,96E-05 | 0,397398104 | 0,857 | 0,733 | 0,949405119 | 6 |
| *CG12004.3* | 8,09E-05 | -0,828392679 | 0,092 | 0,275 | 0,965291401 | 6 |
| *eIF4A.3* | 9,43E-05 | -0,603429492 | 0,378 | 0,613 | 1 | 6 |
| *Pax.1* | 9,60E-05 | 0,664552681 | 0,367 | 0,216 | 1 | 6 |
| *Mkp3.3* | 9,67E-05 | -0,938705579 | 0,214 | 0,407 | 1 | 6 |
| *jim.3* | 0,00010167 | 0,812788316 | 0,612 | 0,513 | 1 | 6 |
| *CAH1.3* | 0,000102526 | 0,963446194 | 0,398 | 0,254 | 1 | 6 |
| *CG32486.4* | 0,000106113 | 0,564038622 | 0,622 | 0,471 | 1 | 6 |
| *ens.3* | 0,000106582 | -0,732456184 | 0,296 | 0,495 | 1 | 6 |
| *alpha-Man-Ia.3* | 0,000107755 | 0,809376433 | 0,5 | 0,356 | 1 | 6 |
| *Hipk.4* | 0,000112968 | -0,688325211 | 0,531 | 0,692 | 1 | 6 |
| *mys.4* | 0,000117655 | -0,698624271 | 0,194 | 0,399 | 1 | 6 |
| *NFAT.3* | 0,000117834 | 0,462289724 | 0,806 | 0,704 | 1 | 6 |

| | | | | | | |
|---|---|---|---|---|---|---|
| *yuri.3* | 0,000118302 | -0,780888534 | 0,122 | 0,314 | 1 | 6 |
| *pes.3* | 0,000120534 | -0,864420849 | 0,153 | 0,347 | 1 | 6 |
| *PAPLA1.3* | 0,000121972 | -0,915051904 | 0,286 | 0,5 | 1 | 6 |
| *Paics.3* | 0,000139238 | -0,596054815 | 0,622 | 0,789 | 1 | 6 |
| *retm.5* | 0,000142156 | 0,699793696 | 0,439 | 0,281 | 1 | 6 |
| *CG42674.3* | 0,000146501 | 0,632116478 | 0,541 | 0,395 | 1 | 6 |
| *Taldo.2* | 0,000147392 | 0,771347842 | 0,429 | 0,297 | 1 | 6 |
| *Plc21C.3* | 0,000147415 | 0,730116684 | 0,653 | 0,57 | 1 | 6 |
| *PGRP-LC.1* | 0,00014782 | 0,616697003 | 0,276 | 0,148 | 1 | 6 |
| *Gdi.2* | 0,00015464 | -0,874009759 | 0,143 | 0,321 | 1 | 6 |
| *CG44008.3* | 0,000180035 | -0,836336065 | 0,092 | 0,261 | 1 | 6 |
| *wb.2* | 0,000185171 | 0,920837506 | 0,48 | 0,335 | 1 | 6 |
| *Ahcy.3* | 0,000187166 | -0,846576529 | 0,122 | 0,299 | 1 | 6 |
| *Rpn13.4* | 0,000190812 | -0,758599603 | 0,102 | 0,28 | 1 | 6 |
| *Sam-S.4* | 0,000193612 | 0,812210441 | 0,714 | 0,61 | 1 | 6 |
| *Gclm.5* | 0,00019453 | -1,069179015 | 0,184 | 0,367 | 1 | 6 |
| *RpL21.3* | 0,000206778 | -0,713168328 | 0,429 | 0,621 | 1 | 6 |
| *cnn.4* | 0,000210625 | -0,779007088 | 0,102 | 0,282 | 1 | 6 |
| *RpL14.3* | 0,00021467 | -0,617683437 | 0,49 | 0,676 | 1 | 6 |
| *Map205.3* | 0,000214913 | -0,730011319 | 0,347 | 0,547 | 1 | 6 |
| *RpS11.3* | 0,000218417 | -0,585901751 | 0,439 | 0,633 | 1 | 6 |
| *mbc.3* | 0,000226249 | 0,664232077 | 0,388 | 0,257 | 1 | 6 |
| *CG6330.4* | 0,000229641 | -0,980220036 | 0,122 | 0,301 | 1 | 6 |
| *klar.1* | 0,000234764 | 0,839233516 | 0,571 | 0,471 | 1 | 6 |
| *pan.3* | 0,000243972 | 0,446700379 | 0,714 | 0,57 | 1 | 6 |
| *nuf.5* | 0,000251714 | -0,70430256 | 0,347 | 0,555 | 1 | 6 |
| *RpL26.3* | 0,000252381 | -0,583276772 | 0,418 | 0,614 | 1 | 6 |
| *uex.3* | 0,000264538 | 0,639739182 | 0,714 | 0,671 | 1 | 6 |
| *Stam.3* | 0,000266697 | -0,827600787 | 0,112 | 0,28 | 1 | 6 |
| *CycG.4* | 0,00028172 | -0,464382327 | 0,949 | 0,947 | 1 | 6 |
| *app.4* | 0,000283668 | 0,7561612 | 0,52 | 0,385 | 1 | 6 |
| *CG44774.3* | 0,000293637 | -0,670807764 | 0,092 | 0,262 | 1 | 6 |
| *srp.4* | 0,000297255 | 0,493001051 | 0,735 | 0,628 | 1 | 6 |
| *Gel.4* | 0,000304679 | 0,632397398 | 0,582 | 0,457 | 1 | 6 |
| *CG5958.5* | 0,000348249 | -1,012150888 | 0,245 | 0,418 | 1 | 6 |
| *jing.2* | 0,000353131 | 0,859851236 | 0,347 | 0,228 | 1 | 6 |
| *Scamp.3* | 0,000358884 | -0,71593358 | 0,092 | 0,258 | 1 | 6 |
| *RpL4.3* | 0,000370166 | -0,429118161 | 0,367 | 0,553 | 1 | 6 |
| *BomS1* | 0,000379397 | 0,272794348 | 0,255 | 0,133 | 1 | 6 |
| *CG11791.4* | 0,000380245 | -1,015761115 | 0,276 | 0,438 | 1 | 6 |
| *zfh1.3* | 0,000381086 | -0,762338516 | 0,204 | 0,391 | 1 | 6 |
| *Sap-r.3* | 0,000393969 | -0,625365267 | 0,378 | 0,559 | 1 | 6 |
| *Got2.4* | 0,000419807 | -0,590417011 | 0,347 | 0,556 | 1 | 6 |
| *CenG1A.3* | 0,000424508 | -1,521146402 | 0,418 | 0,552 | 1 | 6 |
| *Kdm4B.3* | 0,000462177 | -0,774962231 | 0,112 | 0,274 | 1 | 6 |
| *Indy.2* | 0,000465052 | -0,500123777 | 0,429 | 0,642 | 1 | 6 |
| *RpL31.3* | 0,000466366 | -0,426612659 | 0,367 | 0,59 | 1 | 6 |
| *Pkn.3* | 0,000484307 | 0,523853114 | 0,673 | 0,575 | 1 | 6 |
| *Gprk1.3* | 0,000486294 | 0,63423745 | 0,48 | 0,342 | 1 | 6 |
| *CG8036.2* | 0,000486759 | 0,788392281 | 0,48 | 0,35 | 1 | 6 |
| *Smr.4* | 0,000492901 | 0,455921939 | 0,816 | 0,746 | 1 | 6 |
| *zormin.4* | 0,000508996 | -0,635235015 | 0,235 | 0,416 | 1 | 6 |
| *Obp99c.3* | 0,000513803 | 0,610045181 | 0,837 | 0,75 | 1 | 6 |
| *unc-13.3* | 0,000521581 | 0,583927227 | 0,592 | 0,455 | 1 | 6 |
| *Syx1A.4* | 0,000528405 | -0,608562008 | 0,367 | 0,554 | 1 | 6 |
| *cac.4* | 0,000543705 | 0,519111025 | 0,602 | 0,451 | 1 | 6 |
| *Mst84Da.3* | 0,000554271 | -0,726779743 | 0,153 | 0,326 | 1 | 6 |
| *CG15096.5* | 0,000568933 | 0,752606747 | 0,388 | 0,248 | 1 | 6 |
| *dnr1.3* | 0,000571667 | -0,864981819 | 0,092 | 0,251 | 1 | 6 |
| *KrT95D.5* | 0,000574 | -0,638244882 | 0,49 | 0,691 | 1 | 6 |
| *CG9044.3* | 0,000575822 | -0,61874336 | 0,112 | 0,281 | 1 | 6 |

| | | | | | | |
|---|---|---|---|---|---|---|
| *Samuel.1* | 0,000585736 | 0,414238016 | 0,724 | 0,634 | 1 | 6 |
| *CG42668.5* | 0,000590651 | -0,509887922 | 0,439 | 0,604 | 1 | 6 |
| *Karybeta3* | 0,000596391 | -0,698438389 | 0,112 | 0,274 | 1 | 6 |
| *CCT2.4* | 0,000600464 | -0,812564052 | 0,102 | 0,261 | 1 | 6 |
| *Gclc.3* | 0,000645201 | -0,860947748 | 0,347 | 0,536 | 1 | 6 |
| *RpS27A.3* | 0,00064812 | -0,557107965 | 0,469 | 0,66 | 1 | 6 |
| *RpL15.3* | 0,000684754 | -0,551691772 | 0,469 | 0,654 | 1 | 6 |
| *Cyt-b5.2* | 0,000716805 | -0,754501776 | 0,337 | 0,493 | 1 | 6 |
| *csw.3* | 0,000748097 | -0,774795078 | 0,286 | 0,471 | 1 | 6 |
| *eEF2.3* | 0,000773862 | -0,385789022 | 0,582 | 0,762 | 1 | 6 |
| *IscU.3* | 0,00079263 | -0,726432962 | 0,265 | 0,427 | 1 | 6 |
| *akirin.2* | 0,000821651 | -0,730609802 | 0,163 | 0,326 | 1 | 6 |
| *dnc.4* | 0,000822355 | 0,498953328 | 0,847 | 0,749 | 1 | 6 |
| *CG42238.2* | 0,000851292 | 0,766983561 | 0,5 | 0,381 | 1 | 6 |
| *RpS28b.2* | 0,000915904 | -0,619268294 | 0,327 | 0,502 | 1 | 6 |
| *eEF5.4* | 0,000960245 | -0,558026951 | 0,378 | 0,547 | 1 | 6 |
| *Plp* | 0,000979793 | 0,404705029 | 0,796 | 0,727 | 1 | 6 |
| *lqf.3* | 0,001012794 | -0,651731793 | 0,388 | 0,565 | 1 | 6 |
| *Nep4.3* | 0,001016417 | 0,692010825 | 0,337 | 0,216 | 1 | 6 |
| *CG8665.1* | 0,001022553 | 0,818138726 | 0,316 | 0,202 | 1 | 6 |
| *Sec16* | 0,001085045 | 0,661930849 | 0,316 | 0,205 | 1 | 6 |
| *Rtnl1.3* | 0,00113905 | -0,88332376 | 0,194 | 0,355 | 1 | 6 |
| *Src42A* | 0,001161695 | 0,649949586 | 0,337 | 0,219 | 1 | 6 |
| *CG4080.3* | 0,001196955 | -0,77528737 | 0,122 | 0,277 | 1 | 6 |
| *Hsc70-5.3* | 0,001242146 | -0,671920908 | 0,133 | 0,287 | 1 | 6 |
| *Rab5.3* | 0,001260675 | -0,622590701 | 0,184 | 0,352 | 1 | 6 |
| *kibra.3* | 0,001271328 | -0,836175843 | 0,296 | 0,45 | 1 | 6 |
| *MCPH1.3* | 0,001297433 | -0,582641163 | 0,153 | 0,32 | 1 | 6 |
| *Ubc6.4* | 0,001327999 | -0,650807257 | 0,122 | 0,275 | 1 | 6 |
| *stas.2* | 0,001335463 | -0,713883123 | 0,143 | 0,298 | 1 | 6 |
| *l(1)G0196.3* | 0,001380111 | 0,621861587 | 0,541 | 0,43 | 1 | 6 |
| *Ubqn.4* | 0,001417955 | -0,608215256 | 0,112 | 0,261 | 1 | 6 |
| *CG42268.2* | 0,001473915 | -0,415259144 | 0,276 | 0,476 | 1 | 6 |
| *AGO3.3* | 0,001572526 | 0,867699509 | 0,347 | 0,233 | 1 | 6 |
| *shrb.3* | 0,001583762 | -0,657835595 | 0,143 | 0,294 | 1 | 6 |
| *Not1.3* | 0,001643272 | -0,455444607 | 0,592 | 0,749 | 1 | 6 |
| *larp.4* | 0,001649698 | 0,514961906 | 0,653 | 0,584 | 1 | 6 |
| *tsl.1* | 0,001665172 | 0,768720735 | 0,265 | 0,157 | 1 | 6 |
| *Oatp30B.4* | 0,001675011 | 0,585050594 | 0,684 | 0,584 | 1 | 6 |
| *Vha26.2* | 0,001707806 | -0,484693294 | 0,112 | 0,266 | 1 | 6 |
| *fus.3* | 0,001748712 | 0,463474437 | 0,51 | 0,396 | 1 | 6 |
| *dsx.4* | 0,001883465 | 0,479600225 | 0,633 | 0,529 | 1 | 6 |
| *lolal.3* | 0,001884168 | 0,585999947 | 0,429 | 0,302 | 1 | 6 |
| *NKAIN.2* | 0,001920629 | -0,589131682 | 0,153 | 0,316 | 1 | 6 |
| *RpS15.4* | 0,001942559 | -0,567949429 | 0,52 | 0,685 | 1 | 6 |
| *RhoGAP18B.2* | 0,001943814 | -0,562249135 | 0,337 | 0,514 | 1 | 6 |
| *Slik.2* | 0,002008131 | -0,532702524 | 0,255 | 0,427 | 1 | 6 |
| *Cyp6v1.1* | 0,002124337 | 0,714115291 | 0,276 | 0,17 | 1 | 6 |
| *Nacalpha.2* | 0,002132142 | -0,470368496 | 0,194 | 0,363 | 1 | 6 |
| *Aldh-III.1* | 0,002223652 | 0,67096896 | 0,265 | 0,161 | 1 | 6 |
| *CycT* | 0,002235627 | -0,597964817 | 0,194 | 0,351 | 1 | 6 |
| *CG32066.4* | 0,002238542 | 0,563854844 | 0,541 | 0,434 | 1 | 6 |
| *CG13917* | 0,002284571 | 0,739750401 | 0,296 | 0,188 | 1 | 6 |
| *Cirl.3* | 0,002343242 | -0,615242605 | 0,214 | 0,373 | 1 | 6 |
| *RpL39.2* | 0,002385626 | -0,556603519 | 0,347 | 0,491 | 1 | 6 |
| *Pde9.3* | 0,002389432 | 0,419478837 | 0,816 | 0,743 | 1 | 6 |
| *CG2201.2* | 0,00244663 | -0,531981742 | 0,204 | 0,364 | 1 | 6 |
| *Src64B.4* | 0,002466308 | -0,972573902 | 0,408 | 0,539 | 1 | 6 |
| *psq.3* | 0,002471846 | 0,576339649 | 0,582 | 0,507 | 1 | 6 |
| *wnd.3* | 0,002494105 | -0,638161463 | 0,184 | 0,337 | 1 | 6 |
| *l(2)gl* | 0,002495115 | -0,537526049 | 0,276 | 0,456 | 1 | 6 |

| | | | | | | |
|---|---|---|---|---|---|---|
| *Dic1.1* | 0,002644094 | 0,701971067 | 0,296 | 0,19 | 1 | 6 |
| *CrebB.4* | 0,00266039 | -0,638545054 | 0,122 | 0,264 | 1 | 6 |
| *CAHbeta* | 0,00268072 | 0,703764235 | 0,398 | 0,292 | 1 | 6 |
| *CG17646.4* | 0,00273769 | -0,360811504 | 0,714 | 0,856 | 1 | 6 |
| *SREBP.2* | 0,002802907 | -0,539121801 | 0,255 | 0,423 | 1 | 6 |
| *RpS19a.2* | 0,002869096 | -0,576118048 | 0,327 | 0,487 | 1 | 6 |
| *CAH3* | 0,002946909 | 0,849778759 | 0,286 | 0,191 | 1 | 6 |
| *rdx.3* | 0,002954868 | 0,395736302 | 0,684 | 0,594 | 1 | 6 |
| *RpL13A.4* | 0,003004622 | -0,453780023 | 0,49 | 0,654 | 1 | 6 |
| *Ref1.2* | 0,003035775 | -0,529519931 | 0,122 | 0,264 | 1 | 6 |
| *Eb1.4* | 0,003044074 | -0,748023418 | 0,122 | 0,253 | 1 | 6 |
| *CG32264.1* | 0,003111753 | -0,527679553 | 0,388 | 0,569 | 1 | 6 |
| *CG6701.2* | 0,003113013 | -0,697334147 | 0,133 | 0,271 | 1 | 6 |
| *RpS24.4* | 0,003126888 | -0,424437953 | 0,245 | 0,406 | 1 | 6 |
| *RpL7.4* | 0,003152567 | -0,504612817 | 0,469 | 0,611 | 1 | 6 |
| *fbl* | 0,00329862 | -0,533998591 | 0,337 | 0,51 | 1 | 6 |
| *CG6966.4* | 0,003330346 | -0,755336472 | 0,306 | 0,453 | 1 | 6 |
| *RpS16.2* | 0,003394989 | -0,397167576 | 0,388 | 0,561 | 1 | 6 |
| *Hex-C.5* | 0,003463817 | -0,642083042 | 0,306 | 0,45 | 1 | 6 |
| *Tm1.2* | 0,003572882 | -0,426985272 | 0,459 | 0,633 | 1 | 6 |
| *CG6115.4* | 0,003658979 | -0,611705097 | 0,286 | 0,433 | 1 | 6 |
| *SCAP* | 0,003745614 | -0,53156255 | 0,153 | 0,299 | 1 | 6 |
| *RpS29.4* | 0,003784254 | -0,487839193 | 0,429 | 0,579 | 1 | 6 |
| *Ire1.4* | 0,004004158 | -0,637329695 | 0,184 | 0,321 | 1 | 6 |
| *Oda.4* | 0,004037028 | -0,404651315 | 0,704 | 0,827 | 1 | 6 |
| *CG33144.3* | 0,004115743 | 0,592600552 | 0,337 | 0,234 | 1 | 6 |
| *Ubi-p5E.3* | 0,004147733 | -0,422360197 | 0,133 | 0,273 | 1 | 6 |
| *Acsl.4* | 0,004272205 | 0,273616186 | 0,684 | 0,552 | 1 | 6 |
| *Rab1.3* | 0,004292721 | -0,56352562 | 0,173 | 0,315 | 1 | 6 |
| *bic.2* | 0,004310763 | -0,52125741 | 0,245 | 0,398 | 1 | 6 |
| *hep.2* | 0,00443083 | -0,535217666 | 0,173 | 0,323 | 1 | 6 |
| *fok.3* | 0,004478103 | -0,559084023 | 0,378 | 0,526 | 1 | 6 |
| *AP-1gamma.2* | 0,00452765 | -0,541360033 | 0,133 | 0,274 | 1 | 6 |
| *Rack1.2* | 0,004581389 | -0,424750205 | 0,327 | 0,487 | 1 | 6 |
| *Idgf6.1* | 0,004606245 | 0,48638617 | 0,306 | 0,202 | 1 | 6 |
| *Ac76E.3* | 0,004712807 | -0,700700136 | 0,235 | 0,377 | 1 | 6 |
| *daw.3* | 0,004815009 | 0,773817521 | 0,469 | 0,387 | 1 | 6 |
| *RpL23A.2* | 0,004836804 | -0,387176858 | 0,327 | 0,476 | 1 | 6 |
| *Lis-1.4* | 0,004875207 | -0,42615897 | 0,214 | 0,37 | 1 | 6 |
| *IP3K2.4* | 0,004947457 | -0,497985367 | 0,5 | 0,664 | 1 | 6 |
| *Lamp1.2* | 0,004999965 | -0,386644911 | 0,214 | 0,372 | 1 | 6 |
| *wdp.2* | 0,00520851 | -0,804586676 | 0,184 | 0,317 | 1 | 6 |
| *CG3164.4* | 0,005278105 | 0,50184213 | 0,592 | 0,517 | 1 | 6 |
| *CG42663.3* | 0,005406239 | -0,769021174 | 0,153 | 0,281 | 1 | 6 |
| *COX6B.1* | 0,00544323 | 0,512129697 | 0,347 | 0,247 | 1 | 6 |
| *lost.4* | 0,005447114 | -0,545977607 | 0,255 | 0,396 | 1 | 6 |
| *Dys.4* | 0,005460062 | 0,419733484 | 0,663 | 0,563 | 1 | 6 |
| *Gs2.3* | 0,00551348 | 0,532700843 | 0,429 | 0,324 | 1 | 6 |
| *Cyp6d5.3* | 0,00561617 | -0,889932065 | 0,224 | 0,351 | 1 | 6 |
| *Kr-h1.3* | 0,00591724 | -0,504736907 | 0,316 | 0,475 | 1 | 6 |
| *tay.3* | 0,005929253 | 0,611461676 | 0,48 | 0,412 | 1 | 6 |
| *Ncoa6* | 0,005961019 | -0,412263477 | 0,235 | 0,398 | 1 | 6 |
| *CG1578.4* | 0,00597927 | 0,491701755 | 0,49 | 0,378 | 1 | 6 |
| *CAP.2* | 0,006012916 | -0,516526296 | 0,173 | 0,317 | 1 | 6 |
| *CG15098.3* | 0,006021068 | -0,584067783 | 0,255 | 0,392 | 1 | 6 |
| *CG18135.3* | 0,006049525 | -0,588494004 | 0,612 | 0,725 | 1 | 6 |
| *CG18067.1* | 0,00605997 | 0,71538773 | 0,531 | 0,481 | 1 | 6 |
| *eEF1gamma.2* | 0,006069319 | -0,381094011 | 0,173 | 0,324 | 1 | 6 |
| *Nedd4.4* | 0,006147598 | -0,573183185 | 0,194 | 0,336 | 1 | 6 |
| *E(Pc).2* | 0,006268448 | -0,513204985 | 0,133 | 0,271 | 1 | 6 |
| *Rbp1-like.3* | 0,006337923 | 0,491716776 | 0,378 | 0,262 | 1 | 6 |

| | | | | | | |
|---|---|---|---|---|---|---|
| *Galphai.1* | 0,00637106 | -0,373125838 | 0,173 | 0,317 | 1 | 6 |
| *Glut4EF.2* | 0,006507276 | -0,481843686 | 0,735 | 0,824 | 1 | 6 |
| *CG10082.5* | 0,006518615 | 0,417469956 | 0,673 | 0,602 | 1 | 6 |
| *Spat.3* | 0,006554108 | -0,703659529 | 0,316 | 0,468 | 1 | 6 |
| *Chc.2* | 0,006599532 | -0,397501788 | 0,163 | 0,308 | 1 | 6 |
| *Galk.4* | 0,006701622 | 0,34779104 | 0,602 | 0,499 | 1 | 6 |
| *RpL19.2* | 0,00708163 | -0,273803296 | 0,49 | 0,69 | 1 | 6 |
| *nsl1.1* | 0,007125922 | -0,466028769 | 0,194 | 0,328 | 1 | 6 |
| *stx.3* | 0,007178386 | 0,529610643 | 0,469 | 0,394 | 1 | 6 |
| *Ehbp1* | 0,007880814 | -0,484024644 | 0,143 | 0,273 | 1 | 6 |
| *glob1.2* | 0,007887073 | -0,458520439 | 0,592 | 0,746 | 1 | 6 |
| *mrva.3* | 0,00812526 | -0,62006392 | 0,184 | 0,303 | 1 | 6 |
| *dco.4* | 0,008316371 | -0,481073261 | 0,163 | 0,301 | 1 | 6 |
| *CG1213.4* | 0,00836144 | -0,820629944 | 0,224 | 0,344 | 1 | 6 |
| *GstE12.5* | 0,008931277 | 0,490651767 | 0,357 | 0,25 | 1 | 6 |
| *RpS25.3* | 0,00929888 | -0,495458213 | 0,429 | 0,58 | 1 | 6 |
| *RpS10b.3* | 0,009505851 | -0,402964738 | 0,49 | 0,646 | 1 | 6 |
| *Cyt-c-p.2* | 0,009519336 | -0,396975146 | 0,316 | 0,47 | 1 | 6 |
| *Mef2.3* | 0,009543964 | 0,386946003 | 0,633 | 0,541 | 1 | 6 |
| *Pka-C1.4* | 0,009545988 | -0,579138588 | 0,571 | 0,67 | 1 | 6 |
| *RpS12.3* | 0,009556633 | -0,524040057 | 0,48 | 0,603 | 1 | 6 |
| *swm.4* | 0,00966325 | -0,582843681 | 0,204 | 0,333 | 1 | 6 |
| *RpL23.4* | 0,009690596 | -0,566204921 | 0,592 | 0,696 | 1 | 6 |
| *chb.4* | 0,009777211 | -0,62147306 | 0,173 | 0,298 | 1 | 6 |
| *RpL24.3* | 0,00980782 | -0,324307812 | 0,469 | 0,61 | 1 | 6 |
| *Tep2.2* | 0,009997947 | 0,475765827 | 0,378 | 0,29 | 1 | 6 |
| *msi* | 6,93E-153 | 2,639348276 | 0,585 | 0,024 | 8,27E-149 | 7 |
| *Prosap* | 2,34E-138 | 4,578653779 | 0,537 | 0,023 | 2,79E-134 | 7 |
| *Ppn* | 7,44E-109 | 3,718176086 | 0,256 | 0,004 | 8,87E-105 | 7 |
| *ct* | 4,10E-103 | 2,371684842 | 0,524 | 0,032 | 4,89E-99 | 7 |
| *sr* | 1,06E-102 | 2,043272989 | 0,366 | 0,013 | 1,27E-98 | 7 |
| *Pgant9* | 2,15E-62 | 1,523200963 | 0,317 | 0,019 | 2,56E-58 | 7 |
| *Spn* | 2,21E-50 | 2,050149893 | 0,573 | 0,087 | 2,63E-46 | 7 |
| *CG40006* | 2,49E-32 | 1,967007273 | 0,415 | 0,069 | 2,97E-28 | 7 |
| *cpo* | 2,66E-32 | 1,650906507 | 0,439 | 0,076 | 3,17E-28 | 7 |
| *mtgo.2* | 6,95E-32 | 2,545298135 | 0,732 | 0,252 | 8,29E-28 | 7 |
| *ush* | 3,69E-31 | 1,679680799 | 0,427 | 0,075 | 4,41E-27 | 7 |
| *MYPT-75D* | 1,38E-30 | 1,563543998 | 0,341 | 0,048 | 1,64E-26 | 7 |
| *vkg* | 1,23E-29 | 2,725658048 | 0,463 | 0,097 | 1,47E-25 | 7 |
| *Col4a1.3* | 2,25E-29 | 2,473821613 | 0,671 | 0,217 | 2,69E-25 | 7 |
| *pum.4* | 3,40E-28 | 2,291091923 | 0,902 | 0,528 | 4,06E-24 | 7 |
| *pnt.2* | 4,38E-28 | 2,047008904 | 0,732 | 0,272 | 5,22E-24 | 7 |
| *Sh* | 2,16E-27 | 2,092257531 | 0,256 | 0,031 | 2,58E-23 | 7 |
| *Ten-m.2* | 8,03E-27 | 2,496986034 | 0,878 | 0,567 | 9,58E-23 | 7 |
| *Octbeta2R.1* | 1,24E-25 | 1,598468112 | 0,378 | 0,069 | 1,48E-21 | 7 |
| *wake* | 9,64E-24 | 1,409892515 | 0,305 | 0,048 | 1,15E-19 | 7 |
| *CG9328* | 3,52E-23 | 1,415201457 | 0,268 | 0,039 | 4,20E-19 | 7 |
| *hdc.3* | 4,07E-23 | 2,086615247 | 0,585 | 0,184 | 4,86E-19 | 7 |
| *Dyrk2* | 1,62E-22 | 1,495787672 | 0,427 | 0,098 | 1,93E-18 | 7 |
| *Jupiter.1* | 1,63E-22 | 2,271506275 | 0,524 | 0,158 | 1,94E-18 | 7 |
| *Snap25* | 9,80E-21 | 1,530290193 | 0,268 | 0,045 | 1,17E-16 | 7 |
| *fax* | 3,03E-20 | 1,059014833 | 0,439 | 0,107 | 3,61E-16 | 7 |
| *Btk29A.3* | 1,10E-19 | 1,472248774 | 0,622 | 0,224 | 1,31E-15 | 7 |
| *sty.1* | 2,52E-19 | 2,015551535 | 0,61 | 0,233 | 3,00E-15 | 7 |
| *CG10543.1* | 5,03E-18 | 1,369443315 | 0,707 | 0,31 | 6,00E-14 | 7 |
| *CG3328* | 1,43E-17 | 1,049257401 | 0,305 | 0,063 | 1,70E-13 | 7 |
| *cic.6* | 6,76E-16 | 1,731227956 | 0,866 | 0,659 | 8,07E-12 | 7 |
| *Zasp52.1* | 1,67E-15 | 1,499800409 | 0,488 | 0,167 | 1,99E-11 | 7 |
| *zfh1.4* | 3,57E-15 | 1,83994308 | 0,695 | 0,377 | 4,25E-11 | 7 |
| *CG3961.1* | 1,17E-13 | 1,258928236 | 0,268 | 0,061 | 1,39E-09 | 7 |
| *norpA* | 5,43E-13 | 0,876430745 | 0,476 | 0,162 | 6,47E-09 | 7 |

| | | | | | | |
|---|---|---|---|---|---|---|
| *shep.4* | 5,74E-13 | 1,253334095 | 0,915 | 0,744 | 6,85E-09 | 7 |
| *Sema1b* | 6,42E-13 | 1,146164133 | 0,463 | 0,16 | 7,66E-09 | 7 |
| *Rbp6.1* | 1,29E-12 | 1,364001195 | 0,402 | 0,132 | 1,54E-08 | 7 |
| *slo.1* | 1,39E-12 | 0,996204115 | 0,268 | 0,065 | 1,66E-08 | 7 |
| *Ptp99A* | 2,78E-12 | 0,953015441 | 0,402 | 0,134 | 3,32E-08 | 7 |
| *Hk* | 3,23E-12 | 1,024461677 | 0,256 | 0,063 | 3,85E-08 | 7 |
| *CG32767.5* | 1,03E-11 | 0,977995744 | 0,72 | 0,378 | 1,23E-07 | 7 |
| *NaCP60E.1* | 1,90E-11 | 0,787264296 | 0,341 | 0,103 | 2,27E-07 | 7 |
| *ena* | 2,54E-11 | 1,429655256 | 0,329 | 0,107 | 3,03E-07 | 7 |
| *fog.2* | 3,22E-11 | 1,29602231 | 0,512 | 0,222 | 3,85E-07 | 7 |
| *CG3726.1* | 4,26E-11 | 1,378201821 | 0,573 | 0,286 | 5,08E-07 | 7 |
| *mamo.6* | 4,43E-11 | 0,975134843 | 0,854 | 0,548 | 5,29E-07 | 7 |
| *kuz.2* | 4,67E-11 | 1,254494878 | 0,598 | 0,298 | 5,57E-07 | 7 |
| *CG12065.5* | 2,03E-10 | 0,978551756 | 0,585 | 0,287 | 2,42E-06 | 7 |
| *mew.4* | 2,55E-10 | 1,009071944 | 0,695 | 0,381 | 3,04E-06 | 7 |
| *dpp* | 2,65E-10 | 0,940541672 | 0,317 | 0,098 | 3,16E-06 | 7 |
| *srp.5* | 3,45E-10 | 1,163050176 | 0,817 | 0,626 | 4,11E-06 | 7 |
| *CG5080* | 1,02E-09 | 1,02374483 | 0,415 | 0,16 | 1,22E-05 | 7 |
| *pan.4* | 1,27E-09 | 0,89717595 | 0,829 | 0,567 | 1,51E-05 | 7 |
| *Mmp2.4* | 1,61E-09 | 1,192307029 | 0,463 | 0,208 | 1,92E-05 | 7 |
| *CG31637* | 2,66E-09 | 1,078968445 | 0,268 | 0,083 | 3,17E-05 | 7 |
| *l(3)05822.3* | 4,79E-09 | 1,154846822 | 0,463 | 0,204 | 5,72E-05 | 7 |
| *CG5888* | 8,79E-09 | 1,034315041 | 0,268 | 0,086 | 0,000104872 | 7 |
| *sbb.5* | 1,25E-08 | 0,806694381 | 0,841 | 0,582 | 0,000149098 | 7 |
| *unc-13.4* | 1,85E-08 | 0,888443793 | 0,72 | 0,453 | 0,000220925 | 7 |
| *sky.4* | 3,24E-08 | 0,741073205 | 0,512 | 0,236 | 0,00038651 | 7 |
| *fz2.3* | 3,54E-08 | 0,833887204 | 0,561 | 0,274 | 0,00042221 | 7 |
| *CG32982.1* | 4,44E-08 | 1,254487099 | 0,305 | 0,113 | 0,000529737 | 7 |
| *Ten-a* | 6,39E-08 | 1,266807165 | 0,354 | 0,147 | 0,00076243 | 7 |
| *RapGAP1.3* | 7,93E-08 | 0,913974909 | 0,512 | 0,246 | 0,000945602 | 7 |
| *psq.4* | 1,24E-07 | 0,824164234 | 0,72 | 0,504 | 0,001480072 | 7 |
| *Pvr.5* | 1,25E-07 | 0,634891297 | 0,707 | 0,429 | 0,00149334 | 7 |
| *Glut4EF.3* | 1,59E-07 | 0,680607478 | 0,902 | 0,819 | 0,001901409 | 7 |
| *CG17574.2* | 2,27E-07 | 0,913511358 | 0,476 | 0,243 | 0,002703285 | 7 |
| *CG34401* | 2,73E-07 | 0,716657706 | 0,341 | 0,139 | 0,003255356 | 7 |
| *Gprk1.4* | 2,97E-07 | 0,649479234 | 0,622 | 0,339 | 0,003543083 | 7 |
| *MESR3.1* | 3,20E-07 | 0,935300916 | 0,439 | 0,209 | 0,003811944 | 7 |
| *l(2)41Ab.4* | 4,34E-07 | 0,636270817 | 0,646 | 0,358 | 0,005177748 | 7 |
| *Nuak1.4* | 5,32E-07 | 0,812933865 | 0,524 | 0,273 | 0,006342288 | 7 |
| *Cip4.1* | 6,34E-07 | 1,023685829 | 0,5 | 0,262 | 0,007561481 | 7 |
| *eIF4EHP.1* | 7,50E-07 | 0,581743072 | 0,768 | 0,495 | 0,008948902 | 7 |
| *baz.4* | 8,88E-07 | 0,719937648 | 0,463 | 0,23 | 0,010587106 | 7 |
| *msn.3* | 9,92E-07 | 0,734113806 | 0,793 | 0,58 | 0,011835656 | 7 |
| *dikar* | 1,16E-06 | 0,853618254 | 0,476 | 0,243 | 0,01385846 | 7 |
| *pod1* | 1,91E-06 | 0,529237016 | 0,268 | 0,1 | 0,022772101 | 7 |
| *RpS15.5* | 4,40E-06 | -0,958570338 | 0,476 | 0,685 | 0,052523304 | 7 |
| *CG3638.5* | 5,42E-06 | 0,716694871 | 0,707 | 0,471 | 0,064631913 | 7 |
| *RhoGAPp190.1* | 5,74E-06 | 0,672982794 | 0,415 | 0,208 | 0,068438127 | 7 |
| *Zdhhc8.1* | 5,87E-06 | 0,688745629 | 0,683 | 0,46 | 0,070078416 | 7 |
| *Ptpmeg2.2* | 9,11E-06 | 0,634809949 | 0,439 | 0,224 | 0,108680807 | 7 |
| *Smr.5* | 1,05E-05 | 0,436253929 | 0,878 | 0,745 | 0,124781485 | 7 |
| *CG31324* | 1,12E-05 | 0,910384633 | 0,317 | 0,144 | 0,133562479 | 7 |
| *pigs.1* | 1,43E-05 | 0,515035758 | 0,402 | 0,194 | 0,170073328 | 7 |
| *cora.1* | 1,54E-05 | 0,764471474 | 0,366 | 0,176 | 0,183795818 | 7 |
| *lbk.3* | 1,64E-05 | 0,808995248 | 0,451 | 0,262 | 0,195929299 | 7 |
| *Flo2.5* | 1,81E-05 | 0,608066691 | 0,512 | 0,296 | 0,216342565 | 7 |
| *lectin-28C.2* | 1,87E-05 | 0,730709228 | 0,524 | 0,324 | 0,22270016 | 7 |
| *Trpm.2* | 1,87E-05 | 0,720012408 | 0,549 | 0,327 | 0,223515567 | 7 |
| *Ptp61F.1* | 1,96E-05 | 0,766588608 | 0,293 | 0,128 | 0,233634964 | 7 |
| *rl.3* | 3,68E-05 | 0,440593682 | 0,927 | 0,897 | 0,439472344 | 7 |
| *MESK2.2* | 3,91E-05 | 0,789597657 | 0,524 | 0,346 | 0,466819283 | 7 |

| | | | | | | |
|---|---|---|---|---|---|---|
| *mei-P26.3* | 4,01E-05 | 0,555114838 | 0,671 | 0,431 | 0,478928555 | 7 |
| *Fim.1* | 4,40E-05 | 0,646950171 | 0,329 | 0,154 | 0,52455629 | 7 |
| *cno* | 4,51E-05 | 0,622872887 | 0,256 | 0,111 | 0,537582433 | 7 |
| *CG33298* | 5,71E-05 | 0,720534637 | 0,268 | 0,122 | 0,681158153 | 7 |
| *eEF1alpha1.3* | 5,98E-05 | -0,476126136 | 0,878 | 0,91 | 0,712853605 | 7 |
| *vsg.2* | 6,05E-05 | 0,793200219 | 0,488 | 0,292 | 0,721187402 | 7 |
| *Hex-A.2* | 6,74E-05 | 0,674238713 | 0,329 | 0,166 | 0,803553917 | 7 |
| *lola.4* | 8,13E-05 | 0,395748037 | 0,927 | 0,862 | 0,969960148 | 7 |
| *mub.2* | 8,14E-05 | 0,441655259 | 0,415 | 0,219 | 0,971525685 | 7 |
| *heph* | 9,58E-05 | 0,608324936 | 0,341 | 0,17 | 1 | 7 |
| *cv-c.4* | 0,000106824 | 0,441224397 | 0,841 | 0,645 | 1 | 7 |
| *Syx1A.5* | 0,000108594 | 0,520811918 | 0,707 | 0,544 | 1 | 7 |
| *Sap47.2* | 0,000120743 | 0,495962135 | 0,573 | 0,358 | 1 | 7 |
| *sbr* | 0,000127256 | 0,441487155 | 0,524 | 0,291 | 1 | 7 |
| *CG11873.1* | 0,000153632 | 0,583545544 | 0,549 | 0,352 | 1 | 7 |
| *Tlk.4* | 0,000165362 | 0,482121002 | 0,72 | 0,561 | 1 | 7 |
| *Tomosyn.1* | 0,000165712 | 0,451212817 | 0,549 | 0,357 | 1 | 7 |
| *upSET.2* | 0,000177467 | 0,537766219 | 0,634 | 0,413 | 1 | 7 |
| *Abl.6* | 0,000178215 | 0,548723065 | 0,573 | 0,367 | 1 | 7 |
| *Rbfox1.3* | 0,000183735 | 0,630410513 | 0,683 | 0,511 | 1 | 7 |
| *Mob2.6* | 0,000196508 | 0,525595887 | 0,89 | 0,749 | 1 | 7 |
| *egh* | 0,000211648 | 0,376353545 | 0,28 | 0,132 | 1 | 7 |
| *nocte* | 0,000233021 | 0,763706645 | 0,463 | 0,281 | 1 | 7 |
| *Meltrin.3* | 0,000245034 | 0,54221796 | 0,451 | 0,259 | 1 | 7 |
| *RpS26.2* | 0,000249052 | -1,05834127 | 0,415 | 0,549 | 1 | 7 |
| *mtd.5* | 0,000295295 | 0,328589866 | 0,817 | 0,688 | 1 | 7 |
| *CG31211* | 0,000303191 | 0,606177672 | 0,293 | 0,149 | 1 | 7 |
| *PDZ-GEF* | 0,000330285 | 0,279926956 | 0,317 | 0,15 | 1 | 7 |
| *CG18171.1* | 0,000431533 | 0,479604091 | 0,28 | 0,135 | 1 | 7 |
| *CG41099* | 0,000439989 | 0,471371628 | 0,354 | 0,187 | 1 | 7 |
| *l(3)L1231.3* | 0,000490631 | 0,466444944 | 0,756 | 0,586 | 1 | 7 |
| *gce.6* | 0,000495314 | 0,481968114 | 0,659 | 0,478 | 1 | 7 |
| *Sirup* | 0,000579913 | 0,768383348 | 0,305 | 0,159 | 1 | 7 |
| *RpL27.4* | 0,000633831 | -0,695207602 | 0,488 | 0,612 | 1 | 7 |
| *tai.3* | 0,000637959 | 0,387090505 | 0,927 | 0,846 | 1 | 7 |
| *Fmr1* | 0,000683119 | 0,475883141 | 0,28 | 0,14 | 1 | 7 |
| *CanA-14F.2* | 0,000715054 | 0,405831806 | 0,622 | 0,429 | 1 | 7 |
| *CG33144.4* | 0,000722063 | 0,800363701 | 0,378 | 0,234 | 1 | 7 |
| *brat* | 0,000733986 | 0,370250113 | 0,256 | 0,123 | 1 | 7 |
| *zip.1* | 0,000751036 | 0,384979289 | 0,354 | 0,189 | 1 | 7 |
| *Prosalpha3.4* | 0,000872248 | -1,176570122 | 0,122 | 0,272 | 1 | 7 |
| *Pur-alpha.3* | 0,000997814 | 0,505988751 | 0,61 | 0,445 | 1 | 7 |
| *CG15611.1* | 0,001109013 | 0,562733507 | 0,268 | 0,137 | 1 | 7 |
| *cpx.4* | 0,001271431 | 0,532786864 | 0,488 | 0,33 | 1 | 7 |
| *Atx2.1* | 0,001416683 | 0,539095266 | 0,39 | 0,243 | 1 | 7 |
| *BicD.1* | 0,001489872 | 0,581698513 | 0,341 | 0,202 | 1 | 7 |
| *mbc.4* | 0,001492005 | 0,515965372 | 0,415 | 0,257 | 1 | 7 |
| *lap.3* | 0,001586727 | 0,458813394 | 0,476 | 0,301 | 1 | 7 |
| *CG32486.5* | 0,001696357 | 0,400961655 | 0,646 | 0,472 | 1 | 7 |
| *CG5151.5* | 0,001698737 | 0,420597571 | 0,793 | 0,649 | 1 | 7 |
| *CG10433.3* | 0,001786376 | -0,765081157 | 0,622 | 0,678 | 1 | 7 |
| *CG17698* | 0,00185044 | 0,293523436 | 0,28 | 0,144 | 1 | 7 |
| *Crtc.3* | 0,002001804 | 0,374645737 | 0,671 | 0,501 | 1 | 7 |
| *Srrm234* | 0,002034021 | 0,597826607 | 0,329 | 0,193 | 1 | 7 |
| *hang.1* | 0,002047347 | 0,495161486 | 0,463 | 0,296 | 1 | 7 |
| *SPARC.4* | 0,002054073 | 0,632052018 | 0,488 | 0,301 | 1 | 7 |
| *Atet.1* | 0,002167882 | 0,292448949 | 0,354 | 0,199 | 1 | 7 |
| *RpS3.2* | 0,00218473 | -0,738671369 | 0,366 | 0,495 | 1 | 7 |
| *CG31145.4* | 0,002240325 | 0,450764202 | 0,817 | 0,731 | 1 | 7 |
| *ewg* | 0,002364611 | 0,42002601 | 0,268 | 0,14 | 1 | 7 |
| *CaMKII.3* | 0,002418547 | 0,45973272 | 0,476 | 0,312 | 1 | 7 |

| | | | | | | |
|---|---|---|---|---|---|---|
| *CG1637* | 0,002450516 | 0,318254024 | 0,293 | 0,162 | 1 | 7 |
| *Chd64.5* | 0,002635431 | 0,366533727 | 0,634 | 0,451 | 1 | 7 |
| *mnb.3* | 0,002717585 | 0,275787886 | 0,585 | 0,402 | 1 | 7 |
| *nej.3* | 0,002773974 | 0,444796273 | 0,695 | 0,509 | 1 | 7 |
| *blot.3* | 0,002957132 | 0,469897485 | 0,537 | 0,394 | 1 | 7 |
| *RN-tre.3* | 0,002989825 | 0,378669234 | 0,354 | 0,211 | 1 | 7 |
| *jim.4* | 0,002993742 | 0,55476512 | 0,671 | 0,512 | 1 | 7 |
| *RpL13A.5* | 0,003062018 | -0,544152446 | 0,537 | 0,652 | 1 | 7 |
| *Pitslre.1* | 0,003198191 | 0,426360656 | 0,573 | 0,41 | 1 | 7 |
| *RpS29.5* | 0,003270824 | -0,654900784 | 0,451 | 0,578 | 1 | 7 |
| *be.1* | 0,00339516 | 0,402366963 | 0,268 | 0,146 | 1 | 7 |
| *ftz-f1.5* | 0,003701905 | 0,459838723 | 0,72 | 0,612 | 1 | 7 |
| *mam.4* | 0,003727305 | 0,382440091 | 0,549 | 0,384 | 1 | 7 |
| *eIF4A.4* | 0,003752616 | -0,594031374 | 0,5 | 0,609 | 1 | 7 |
| *RpS11.4* | 0,003949206 | -0,510132634 | 0,549 | 0,629 | 1 | 7 |
| *sdk.2* | 0,004030934 | 0,446699297 | 0,329 | 0,2 | 1 | 7 |
| *RpS7.3* | 0,004541013 | -0,505575731 | 0,598 | 0,682 | 1 | 7 |
| *CG31998* | 0,004895248 | 0,386731743 | 0,329 | 0,199 | 1 | 7 |
| *Mal-B2.5* | 0,004921173 | -0,679646058 | 0,415 | 0,528 | 1 | 7 |
| *CG6428.5* | 0,005149871 | -0,860113023 | 0,146 | 0,275 | 1 | 7 |
| *CG9003.1* | 0,005343615 | 0,43356939 | 0,39 | 0,257 | 1 | 7 |
| *Pfas.4* | 0,006008226 | -0,649408439 | 0,463 | 0,568 | 1 | 7 |
| *CG10960.5* | 0,006043737 | -0,385108581 | 0,768 | 0,834 | 1 | 7 |
| *wts* | 0,006349429 | 0,55249081 | 0,451 | 0,311 | 1 | 7 |
| *Usp47* | 0,006465202 | 0,278338999 | 0,268 | 0,15 | 1 | 7 |
| *mod(mdg4)* | 0,007036443 | 0,475042943 | 0,28 | 0,17 | 1 | 7 |
| *cnc.1* | 0,007271229 | -0,366673524 | 0,951 | 0,963 | 1 | 7 |
| *hppy.1* | 0,007430036 | 0,408783466 | 0,622 | 0,518 | 1 | 7 |
| *GramD1B.2* | 0,007921779 | -0,442292426 | 0,415 | 0,556 | 1 | 7 |
| *Ars2* | 0,008028229 | 0,615580944 | 0,256 | 0,154 | 1 | 7 |
| *Irc.5* | 0,008193761 | -0,666694511 | 0,415 | 0,518 | 1 | 7 |
| *CG10799.3* | 0,008595181 | -0,465992495 | 0,39 | 0,506 | 1 | 7 |
| *Trf2* | 0,008866953 | 0,292290721 | 0,622 | 0,474 | 1 | 7 |
| *sm.4* | 0,009037355 | 0,349704241 | 0,439 | 0,3 | 1 | 7 |
| *alphaTub84B.2* | 0,009154692 | 0,572309173 | 0,476 | 0,351 | 1 | 7 |
| *DIP-lambda.4* | 0,009255379 | 0,403817861 | 0,39 | 0,272 | 1 | 7 |
| *AGO3.4* | 0,00962078 | 0,319033447 | 0,366 | 0,233 | 1 | 7 |
| *CG6701.3* | 0,009720623 | 0,562300408 | 0,39 | 0,263 | 1 | 7 |
| *rin.1* | 0,009817373 | 0,369363424 | 0,524 | 0,387 | 1 | 7 |
| *BomBc1* | 4,01E-204 | 4,539199608 | 0,527 | 0,009 | 4,79E-200 | 8 |
| *BomT1* | 8,06E-195 | 3,588040984 | 0,541 | 0,011 | 9,62E-191 | 8 |
| *DptB* | 6,10E-150 | 4,08502087 | 0,541 | 0,018 | 7,28E-146 | 8 |
| *AttC* | 1,05E-104 | 4,036727181 | 0,541 | 0,031 | 1,25E-100 | 8 |
| *Mtk* | 1,27E-102 | 2,917377013 | 0,378 | 0,013 | 1,52E-98 | 8 |
| *BomBc3.1* | 4,26E-99 | 3,122658489 | 0,757 | 0,075 | 5,08E-95 | 8 |
| *GNBP-like3* | 1,78E-87 | 3,903287791 | 0,851 | 0,122 | 2,12E-83 | 8 |
| *DptA* | 9,68E-85 | 3,540120924 | 0,392 | 0,019 | 1,15E-80 | 8 |
| *BomS1.1* | 1,17E-76 | 4,606597064 | 0,797 | 0,121 | 1,40E-72 | 8 |
| *PGRP-SB1* | 1,22E-72 | 2,270396239 | 0,311 | 0,013 | 1,46E-68 | 8 |
| *Dro* | 4,62E-72 | 3,700463118 | 0,514 | 0,045 | 5,51E-68 | 8 |
| *IM14.3* | 1,02E-66 | 3,616280047 | 0,892 | 0,192 | 1,21E-62 | 8 |
| *Drs* | 2,40E-56 | 4,603389616 | 0,554 | 0,071 | 2,86E-52 | 8 |
| *IM4.6* | 6,02E-56 | 3,434413678 | 0,959 | 0,291 | 7,18E-52 | 8 |
| *BomS5* | 6,92E-54 | 1,385329101 | 0,257 | 0,013 | 8,26E-50 | 8 |
| *BomS2.3* | 1,36E-48 | 3,52267233 | 0,946 | 0,382 | 1,62E-44 | 8 |
| *BomS3.5* | 1,03E-47 | 3,383571251 | 0,973 | 0,42 | 1,22E-43 | 8 |
| *CG16772* | 4,43E-45 | 3,621533624 | 0,486 | 0,066 | 5,28E-41 | 8 |
| *Tsf1.6* | 2,42E-44 | 2,3167839 | 0,919 | 0,279 | 2,88E-40 | 8 |
| *CG16713* | 1,30E-39 | 2,082209843 | 0,676 | 0,142 | 1,55E-35 | 8 |
| *CecA2* | 1,56E-39 | 2,957255624 | 0,257 | 0,019 | 1,87E-35 | 8 |
| *BomBc2.4* | 1,28E-38 | 2,880998986 | 0,973 | 0,547 | 1,53E-34 | 8 |

| | | | | | | |
|---|---|---|---|---|---|---|
| *BomT3.6* | 1,30E-38 | 3,014113554 | 0,932 | 0,414 | 1,55E-34 | 8 |
| *cue.1* | 1,86E-38 | 1,970687046 | 0,689 | 0,15 | 2,22E-34 | 8 |
| *CG18067.2* | 2,69E-38 | 4,06907139 | 0,932 | 0,472 | 3,21E-34 | 8 |
| *CG1358.2* | 7,19E-37 | 2,269699485 | 0,743 | 0,192 | 8,58E-33 | 8 |
| *NimB1* | 1,95E-34 | 1,660563237 | 0,405 | 0,056 | 2,32E-30 | 8 |
| *CG30002* | 2,29E-34 | 1,543494555 | 0,378 | 0,049 | 2,73E-30 | 8 |
| *IM33.6* | 1,44E-27 | 2,40697442 | 0,838 | 0,357 | 1,72E-23 | 8 |
| *fra* | 1,32E-26 | 1,696847012 | 0,405 | 0,072 | 1,57E-22 | 8 |
| *Stacl* | 5,39E-26 | 1,94643593 | 0,284 | 0,037 | 6,43E-22 | 8 |
| *CrebA.3* | 1,29E-24 | 3,384596109 | 0,689 | 0,242 | 1,54E-20 | 8 |
| *Ipk1* | 9,60E-23 | 1,332846653 | 0,257 | 0,034 | 1,15E-18 | 8 |
| *Tep2.3* | 6,02E-22 | 1,900507222 | 0,757 | 0,282 | 7,18E-18 | 8 |
| *Dif.5* | 2,71E-20 | 2,278125445 | 0,77 | 0,384 | 3,23E-16 | 8 |
| *SPE* | 3,25E-19 | 1,254909185 | 0,392 | 0,085 | 3,87E-15 | 8 |
| *Sid* | 1,87E-18 | 1,413628469 | 0,284 | 0,051 | 2,23E-14 | 8 |
| *CG32521.4* | 7,33E-18 | 1,393591706 | 0,959 | 0,765 | 8,75E-14 | 8 |
| *LpR2.5* | 1,12E-17 | 1,550308302 | 0,932 | 0,845 | 1,34E-13 | 8 |
| *fon.4* | 3,27E-17 | 1,459197929 | 0,703 | 0,303 | 3,90E-13 | 8 |
| *CG9928.1* | 6,75E-17 | 1,242432116 | 0,486 | 0,139 | 8,06E-13 | 8 |
| *BomT2.1* | 1,26E-16 | 1,584159522 | 0,554 | 0,195 | 1,51E-12 | 8 |
| *Gart.4* | 1,64E-16 | -2,300847263 | 0,189 | 0,677 | 1,95E-12 | 8 |
| *shf.1* | 5,04E-16 | 1,489550186 | 0,486 | 0,151 | 6,01E-12 | 8 |
| *Gbs-70E.6* | 2,03E-15 | 1,195468333 | 0,919 | 0,52 | 2,42E-11 | 8 |
| *CG9674.5* | 3,06E-15 | -2,542577943 | 0,189 | 0,644 | 3,65E-11 | 8 |
| *Nmdmc.5* | 6,50E-15 | -2,456263904 | 0,216 | 0,654 | 7,75E-11 | 8 |
| *CG14762.4* | 1,32E-14 | 1,488638724 | 0,73 | 0,386 | 1,58E-10 | 8 |
| *sug.6* | 2,69E-14 | 1,124024058 | 0,73 | 0,315 | 3,21E-10 | 8 |
| *pnt.3* | 2,85E-13 | 1,075713607 | 0,649 | 0,276 | 3,40E-09 | 8 |
| *dl.1* | 3,19E-13 | 1,477166242 | 0,365 | 0,105 | 3,81E-09 | 8 |
| *CG30026.1* | 4,81E-13 | 1,147426414 | 0,473 | 0,164 | 5,73E-09 | 8 |
| *osp.5* | 1,78E-12 | 1,47941153 | 0,635 | 0,31 | 2,13E-08 | 8 |
| *ref(2)P.4* | 1,97E-12 | -2,345441202 | 0,284 | 0,645 | 2,35E-08 | 8 |
| *Nha2.2* | 3,92E-12 | 1,132125558 | 0,459 | 0,154 | 4,67E-08 | 8 |
| *Xrp1.4* | 4,54E-12 | -1,299842922 | 0,824 | 0,913 | 5,41E-08 | 8 |
| *CG11841* | 8,16E-12 | 1,073900851 | 0,257 | 0,06 | 9,74E-08 | 8 |
| *Vago* | 2,61E-11 | 1,685204995 | 0,284 | 0,076 | 3,11E-07 | 8 |
| *Tapdelta* | 4,06E-11 | 0,807669655 | 0,284 | 0,074 | 4,85E-07 | 8 |
| *CG11089.5* | 4,12E-11 | -2,305273521 | 0,392 | 0,667 | 4,92E-07 | 8 |
| *prage.2* | 5,77E-11 | -1,527512917 | 0,486 | 0,745 | 6,88E-07 | 8 |
| *CG16898.4* | 6,38E-11 | -2,379691713 | 0,054 | 0,446 | 7,61E-07 | 8 |
| *ergic53* | 8,02E-11 | 0,843482389 | 0,27 | 0,07 | 9,56E-07 | 8 |
| *Pur-alpha.4* | 9,95E-11 | 0,980358058 | 0,784 | 0,441 | 1,19E-06 | 8 |
| *CG10383.4* | 1,01E-10 | -2,606489336 | 0,095 | 0,48 | 1,21E-06 | 8 |
| *l(1)G0320* | 1,06E-10 | 0,947499836 | 0,257 | 0,066 | 1,26E-06 | 8 |
| *Sdc.4* | 1,23E-10 | 0,765113579 | 0,986 | 0,948 | 1,47E-06 | 8 |
| *Hsc70-3.2* | 1,82E-10 | 2,308056741 | 0,662 | 0,405 | 2,17E-06 | 8 |
| *Mec2* | 2,07E-10 | 0,954299949 | 0,257 | 0,067 | 2,47E-06 | 8 |
| *Calr.1* | 4,96E-10 | 1,626760895 | 0,595 | 0,315 | 5,92E-06 | 8 |
| *CG6426.1* | 5,35E-10 | 1,271892342 | 0,568 | 0,274 | 6,38E-06 | 8 |
| *Caper.3* | 6,29E-10 | 0,894571021 | 0,851 | 0,667 | 7,51E-06 | 8 |
| *CG3036.5* | 7,37E-10 | -2,011861913 | 0,365 | 0,658 | 8,79E-06 | 8 |
| *GstE1.4* | 7,91E-10 | -2,554051229 | 0,027 | 0,385 | 9,44E-06 | 8 |
| *shep.5* | 8,22E-10 | 0,841920223 | 0,973 | 0,743 | 9,80E-06 | 8 |
| *Desat1.4* | 1,08E-09 | 0,708039171 | 1 | 0,89 | 1,29E-05 | 8 |
| *gukh* | 1,57E-09 | 1,006738261 | 0,311 | 0,098 | 1,87E-05 | 8 |
| *CG12795.5* | 1,90E-09 | -2,676028694 | 0,014 | 0,358 | 2,26E-05 | 8 |
| *CG13360.1* | 1,92E-09 | 0,817950975 | 0,405 | 0,146 | 2,29E-05 | 8 |
| *CG5151.6* | 2,01E-09 | 0,732075367 | 0,905 | 0,647 | 2,40E-05 | 8 |
| *trbl.5* | 2,15E-09 | -1,859714688 | 0,189 | 0,52 | 2,57E-05 | 8 |
| *CG16758.3* | 2,53E-09 | -1,100113185 | 0,784 | 0,925 | 3,02E-05 | 8 |
| *Tsp42Ed.5* | 2,63E-09 | -2,132702294 | 0,108 | 0,444 | 3,14E-05 | 8 |

| | | | | | | |
|---|---|---|---|---|---|---|
| *Smg5.4* | 2,83E-09 | -1,864348425 | 0,149 | 0,493 | 3,37E-05 | 8 |
| *BomS6.2* | 3,19E-09 | 1,248093708 | 0,459 | 0,205 | 3,81E-05 | 8 |
| *Hayan* | 4,89E-09 | 1,252303907 | 0,392 | 0,151 | 5,83E-05 | 8 |
| *Pdi.1* | 6,40E-09 | 1,596582201 | 0,5 | 0,247 | 7,63E-05 | 8 |
| *stv.4* | 6,67E-09 | -2,395640047 | 0,338 | 0,625 | 7,96E-05 | 8 |
| *Sec16.1* | 8,04E-09 | 0,969076501 | 0,459 | 0,202 | 9,59E-05 | 8 |
| *CG2918* | 8,82E-09 | 0,998658915 | 0,284 | 0,089 | 0,000105175 | 8 |
| *CG14207.5* | 1,83E-08 | -1,928069901 | 0,324 | 0,593 | 0,000218304 | 8 |
| *Chchd2.5* | 2,07E-08 | -1,585457772 | 0,365 | 0,624 | 0,000247419 | 8 |
| *Lpin.4* | 2,09E-08 | -1,672608644 | 0,162 | 0,489 | 0,000248749 | 8 |
| *dnc.5* | 2,19E-08 | 0,634564888 | 0,959 | 0,747 | 0,000261256 | 8 |
| *CG42588.5* | 2,36E-08 | -2,102460089 | 0,014 | 0,325 | 0,000282084 | 8 |
| *CG10680.4* | 4,80E-08 | 0,914716706 | 0,703 | 0,411 | 0,000572414 | 8 |
| *Pka-C1.5* | 5,33E-08 | -1,402912208 | 0,459 | 0,672 | 0,000636125 | 8 |
| *TBC1D5* | 5,97E-08 | 0,970006717 | 0,297 | 0,101 | 0,000712381 | 8 |
| *Pfas.5* | 6,00E-08 | -1,575840436 | 0,297 | 0,572 | 0,000715506 | 8 |
| *scyl.4* | 6,02E-08 | -1,184336203 | 0,649 | 0,812 | 0,000717952 | 8 |
| *trol* | 6,02E-08 | 1,118345965 | 0,257 | 0,082 | 0,000718074 | 8 |
| *AdSS.4* | 6,62E-08 | -1,862682976 | 0,122 | 0,423 | 0,000789713 | 8 |
| *CG5850* | 6,82E-08 | 0,678084303 | 0,311 | 0,107 | 0,000813939 | 8 |
| *Hsp27.6* | 7,12E-08 | -3,73304312 | 0,054 | 0,348 | 0,000849573 | 8 |
| *Lk6.4* | 7,37E-08 | -0,915081204 | 0,865 | 0,897 | 0,000879435 | 8 |
| *CG45050.4* | 8,40E-08 | -1,151034841 | 0,797 | 0,905 | 0,001001645 | 8 |
| *nec* | 9,36E-08 | 1,043537393 | 0,405 | 0,184 | 0,001116659 | 8 |
| *Bsg.3* | 9,75E-08 | -0,617710369 | 0,986 | 0,982 | 0,001163495 | 8 |
| *oys.6* | 1,04E-07 | -1,992925699 | 0,041 | 0,339 | 0,001240761 | 8 |
| *cac.5* | 1,05E-07 | -2,245657614 | 0,162 | 0,462 | 0,001256201 | 8 |
| *Ald1.2* | 1,28E-07 | -0,936598193 | 0,838 | 0,883 | 0,001528241 | 8 |
| *Atg1.3* | 1,30E-07 | -1,812497516 | 0,243 | 0,51 | 0,001544924 | 8 |
| *Egfr.4* | 1,33E-07 | 0,597303933 | 0,973 | 0,769 | 0,001582545 | 8 |
| *emc.1* | 1,52E-07 | 0,805460092 | 0,432 | 0,192 | 0,001814153 | 8 |
| *Spn43Ab.4* | 1,53E-07 | 0,882436727 | 0,635 | 0,354 | 0,001820241 | 8 |
| *TER94.4* | 1,54E-07 | -1,545227238 | 0,216 | 0,499 | 0,001836921 | 8 |
| *MP1* | 1,63E-07 | 0,859306883 | 0,284 | 0,099 | 0,001950097 | 8 |
| *AOX1.4* | 1,68E-07 | -2,048398196 | 0,054 | 0,341 | 0,002009613 | 8 |
| *l(3)80Fg.4* | 1,88E-07 | 0,748734322 | 0,851 | 0,605 | 0,002240132 | 8 |
| *puc.6* | 2,04E-07 | -1,50630439 | 0,527 | 0,697 | 0,00243486 | 8 |
| *teq.6* | 2,25E-07 | 0,840797106 | 0,5 | 0,244 | 0,002688623 | 8 |
| *eIF5B* | 2,48E-07 | 0,832274209 | 0,419 | 0,184 | 0,002956088 | 8 |
| *Gbp2.1* | 3,50E-07 | 0,87599572 | 0,581 | 0,305 | 0,004174284 | 8 |
| *mtd.6* | 3,99E-07 | 0,429231804 | 0,932 | 0,686 | 0,004760894 | 8 |
| *MtnA.5* | 4,11E-07 | -1,314581206 | 0,27 | 0,575 | 0,004898167 | 8 |
| *Amph* | 4,15E-07 | 0,765909863 | 0,297 | 0,107 | 0,004955207 | 8 |
| *Idgf6.2* | 4,72E-07 | 0,997774541 | 0,419 | 0,201 | 0,005627175 | 8 |
| *CG15293.4* | 5,59E-07 | 0,810294074 | 0,703 | 0,441 | 0,006668744 | 8 |
| *CG7130.5* | 6,57E-07 | -2,051726294 | 0,014 | 0,277 | 0,007836095 | 8 |
| *Fas3.2* | 7,38E-07 | 1,185619346 | 0,351 | 0,149 | 0,008806746 | 8 |
| *CG5773.5* | 7,55E-07 | 0,810230096 | 0,486 | 0,225 | 0,009003206 | 8 |
| *mamo.7* | 8,19E-07 | 0,474351513 | 0,865 | 0,548 | 0,009764734 | 8 |
| *Hsp26.5* | 8,22E-07 | -3,211905926 | 0,176 | 0,427 | 0,009808851 | 8 |
| *CenG1A.4* | 9,07E-07 | -2,212952953 | 0,324 | 0,553 | 0,010815755 | 8 |
| *Pdp1.6* | 9,37E-07 | -0,575816613 | 0,973 | 0,993 | 0,011171812 | 8 |
| *CG10031* | 9,56E-07 | 0,808762054 | 0,257 | 0,09 | 0,011403718 | 8 |
| *MFS17.4* | 1,13E-06 | 0,575434695 | 0,892 | 0,691 | 0,013473139 | 8 |
| *sbb.6* | 1,16E-06 | 0,707491539 | 0,838 | 0,583 | 0,013859255 | 8 |
| *Ubi-p63E.5* | 1,21E-06 | -1,553823741 | 0,432 | 0,617 | 0,014481985 | 8 |
| *Iml1.1* | 1,24E-06 | 1,021013244 | 0,351 | 0,159 | 0,014751611 | 8 |
| *l(1)G0007* | 1,52E-06 | 0,726652897 | 0,311 | 0,121 | 0,018139179 | 8 |
| *Prps.3* | 1,55E-06 | -0,754246954 | 0,824 | 0,927 | 0,018472675 | 8 |
| *spoon.5* | 1,73E-06 | 0,686581381 | 0,703 | 0,424 | 0,020622341 | 8 |
| *Pdk.3* | 1,89E-06 | -0,748237149 | 0,797 | 0,864 | 0,022594977 | 8 |

| | | | | | | |
|---|---|---|---|---|---|---|
| *CG5059.2* | 2,15E-06 | 0,682600439 | 0,865 | 0,69 | 0,025677532 | 8 |
| *CG8086.3* | 2,44E-06 | -1,570449573 | 0,068 | 0,331 | 0,029068889 | 8 |
| *bru1.4* | 2,45E-06 | 0,535893906 | 0,946 | 0,796 | 0,029221393 | 8 |
| *Spn77Ba* | 2,50E-06 | 0,769362101 | 0,284 | 0,111 | 0,029879985 | 8 |
| *cher.5* | 2,58E-06 | -2,100549191 | 0,054 | 0,297 | 0,030829395 | 8 |
| *mbf1.5* | 2,82E-06 | -1,755246994 | 0,108 | 0,356 | 0,033650829 | 8 |
| *Shmt.4* | 2,98E-06 | -1,202215435 | 0,432 | 0,618 | 0,035558762 | 8 |
| *Helz.5* | 3,04E-06 | -1,031541663 | 0,865 | 0,91 | 0,036252652 | 8 |
| *Gnmt.2* | 3,06E-06 | 0,822852209 | 0,824 | 0,641 | 0,036547285 | 8 |
| *cv-d.1* | 3,30E-06 | 0,672080093 | 0,392 | 0,182 | 0,039364177 | 8 |
| *CG6040.2* | 3,31E-06 | 0,774214862 | 0,351 | 0,152 | 0,039429893 | 8 |
| *Pvr.6* | 3,40E-06 | -2,188421437 | 0,189 | 0,442 | 0,040588115 | 8 |
| *Sox102F.4* | 3,52E-06 | 0,482409897 | 0,919 | 0,743 | 0,042020673 | 8 |
| *Cyp4p1.5* | 3,85E-06 | -1,532481108 | 0,068 | 0,316 | 0,04592366 | 8 |
| *Rpn6.4* | 3,94E-06 | -1,347083162 | 0,135 | 0,395 | 0,046996337 | 8 |
| *raw.5* | 4,10E-06 | -1,17131412 | 0,459 | 0,638 | 0,048881349 | 8 |
| *sra.5* | 4,99E-06 | -1,643523138 | 0,041 | 0,276 | 0,059528937 | 8 |
| *Hsc70-4.5* | 5,56E-06 | -1,045846445 | 0,595 | 0,743 | 0,066314326 | 8 |
| *Hers.6* | 5,59E-06 | -1,182561782 | 0,459 | 0,626 | 0,066681765 | 8 |
| *Nep4.4* | 6,58E-06 | 0,957024051 | 0,419 | 0,215 | 0,078525192 | 8 |
| *Cyp6d5.4* | 6,62E-06 | -1,389373373 | 0,108 | 0,352 | 0,07893747 | 8 |
| *bgm.3* | 7,82E-06 | 0,876887161 | 0,811 | 0,631 | 0,093241519 | 8 |
| *ap.4* | 8,29E-06 | 0,803853541 | 0,568 | 0,338 | 0,098886809 | 8 |
| *loco.5* | 8,65E-06 | -1,447469097 | 0,041 | 0,28 | 0,10315263 | 8 |
| *Syp.5* | 9,01E-06 | 0,487676524 | 0,946 | 0,811 | 0,107450644 | 8 |
| *CG7115* | 9,12E-06 | 1,136964846 | 0,257 | 0,1 | 0,108818407 | 8 |
| *Invadolysin.5* | 9,16E-06 | 0,613197617 | 0,527 | 0,267 | 0,109254094 | 8 |
| *CG17549.4* | 9,33E-06 | 0,729168665 | 0,595 | 0,367 | 0,111309588 | 8 |
| *NFAT.4* | 9,57E-06 | 0,559031346 | 0,905 | 0,702 | 0,114187028 | 8 |
| *CG16704* | 9,89E-06 | 0,715909791 | 0,297 | 0,123 | 0,117941929 | 8 |
| *rhea.5* | 1,02E-05 | -1,846164949 | 0,297 | 0,499 | 0,121991649 | 8 |
| *Trxr-1.5* | 1,08E-05 | -1,44782864 | 0,162 | 0,397 | 0,129276702 | 8 |
| *Ndae1.4* | 1,08E-05 | 0,657098136 | 0,581 | 0,333 | 0,129293125 | 8 |
| *cact.6* | 1,08E-05 | 1,517609346 | 0,77 | 0,638 | 0,12940755 | 8 |
| *Frl.6* | 1,13E-05 | -1,596289276 | 0,338 | 0,518 | 0,134518123 | 8 |
| *MTA1-like.5* | 1,17E-05 | -2,258508216 | 0,243 | 0,438 | 0,140039276 | 8 |
| *Abl.7* | 1,21E-05 | -1,868464143 | 0,162 | 0,377 | 0,144019585 | 8 |
| *Fkbp14.2* | 1,23E-05 | 0,723105014 | 0,5 | 0,268 | 0,147071591 | 8 |
| *CG1607* | 1,27E-05 | 0,710960021 | 0,27 | 0,11 | 0,151444382 | 8 |
| *eIF2beta.4* | 1,45E-05 | -1,422118547 | 0,324 | 0,533 | 0,172923129 | 8 |
| *Hsp83.5* | 1,46E-05 | -1,2609763 | 0,622 | 0,737 | 0,173573845 | 8 |
| *CG17124.6* | 1,47E-05 | 0,589739129 | 0,932 | 0,744 | 0,174807979 | 8 |
| *CG12054.3* | 1,48E-05 | 0,734522505 | 0,608 | 0,383 | 0,176326433 | 8 |
| *GIIIspla2.4* | 1,59E-05 | -1,580957267 | 0,189 | 0,422 | 0,189331671 | 8 |
| *Hsp68.5* | 1,63E-05 | -2,401791395 | 0,081 | 0,304 | 0,193946053 | 8 |
| *Rbp1-like.4* | 1,79E-05 | 0,695149408 | 0,486 | 0,26 | 0,213397059 | 8 |
| *Lip4.4* | 1,95E-05 | -1,411315408 | 0,176 | 0,413 | 0,233034444 | 8 |
| *Adk1* | 2,21E-05 | 0,910819613 | 0,297 | 0,132 | 0,264005227 | 8 |
| *CCHa2.6* | 2,25E-05 | 0,474828916 | 0,5 | 0,257 | 0,268771287 | 8 |
| *Src64B.5* | 2,37E-05 | -1,613162614 | 0,365 | 0,539 | 0,283306497 | 8 |
| *cv-c.5* | 2,43E-05 | -1,060468551 | 0,459 | 0,655 | 0,290351329 | 8 |
| *Ttd14.4* | 2,55E-05 | 0,894014319 | 0,473 | 0,276 | 0,304281157 | 8 |
| *CG43103* | 2,64E-05 | 0,582324874 | 0,297 | 0,126 | 0,315203054 | 8 |
| *Spn88Eb.3* | 2,68E-05 | 0,715802543 | 0,392 | 0,197 | 0,320002627 | 8 |
| *alpha-Est9.2* | 2,74E-05 | 0,786323053 | 0,365 | 0,178 | 0,326555769 | 8 |
| *InR.5* | 2,79E-05 | -0,913408574 | 0,878 | 0,914 | 0,332409811 | 8 |
| *CG1673.6* | 2,91E-05 | -1,132998213 | 0,392 | 0,59 | 0,347022691 | 8 |
| *Pino.2* | 2,98E-05 | 0,635165869 | 0,878 | 0,764 | 0,355184055 | 8 |
| *Kr-h1.4* | 3,09E-05 | -0,511558548 | 0,203 | 0,476 | 0,368529219 | 8 |
| *Tet.3* | 3,11E-05 | -1,045340845 | 0,365 | 0,571 | 0,371577481 | 8 |
| *pug.2* | 3,17E-05 | -1,066698461 | 0,351 | 0,572 | 0,377559531 | 8 |

| | | | | | | |
|---|---|---|---|---|---|---|
| *CG8468.3* | 3,31E-05 | -0,706079329 | 0,405 | 0,681 | 0,394665364 | 8 |
| *Pgant5.2* | 3,62E-05 | 0,979558899 | 0,446 | 0,263 | 0,431253398 | 8 |
| *CG5953.4* | 3,90E-05 | -1,519768278 | 0,608 | 0,681 | 0,465524369 | 8 |
| *Rad23.5* | 4,09E-05 | -1,331040897 | 0,135 | 0,337 | 0,488477807 | 8 |
| *Ets98B.3* | 4,11E-05 | 0,507509846 | 0,757 | 0,508 | 0,489982326 | 8 |
| *tth* | 4,29E-05 | 0,563679993 | 0,351 | 0,166 | 0,51185102 | 8 |
| *AnxB9.6* | 4,43E-05 | -1,821013699 | 0,338 | 0,5 | 0,528237411 | 8 |
| *CG9932.3* | 4,45E-05 | 0,473736586 | 0,973 | 0,948 | 0,531087383 | 8 |
| *lola.5* | 5,12E-05 | 0,460257735 | 0,946 | 0,862 | 0,610873998 | 8 |
| *spin.4* | 5,18E-05 | -0,844313823 | 0,419 | 0,634 | 0,617769647 | 8 |
| *CG12290.3* | 5,28E-05 | -1,402559713 | 0,068 | 0,282 | 0,629804258 | 8 |
| *Sec63* | 5,42E-05 | 0,660191953 | 0,351 | 0,165 | 0,64703762 | 8 |
| *Nplp2.6* | 5,65E-05 | 0,638951708 | 0,919 | 0,662 | 0,674072162 | 8 |
| *MRP.4* | 5,69E-05 | -1,754550325 | 0,23 | 0,432 | 0,678990331 | 8 |
| *SNF4Agamma.5* | 5,71E-05 | -0,616659143 | 0,892 | 0,892 | 0,680978263 | 8 |
| *Oda.5* | 6,47E-05 | -0,769330389 | 0,784 | 0,824 | 0,772319508 | 8 |
| *CG16799.1* | 6,54E-05 | 0,772485204 | 0,365 | 0,186 | 0,779612879 | 8 |
| *CG33493.2* | 6,59E-05 | 0,702642789 | 0,405 | 0,219 | 0,785650017 | 8 |
| *Fur1.3* | 6,97E-05 | 0,371569127 | 0,919 | 0,713 | 0,831904105 | 8 |
| *Lmpt.3* | 6,98E-05 | -1,03487397 | 0,243 | 0,457 | 0,833201113 | 8 |
| *lilli.5* | 7,01E-05 | 0,526267132 | 0,878 | 0,728 | 0,836576418 | 8 |
| *Hs6st* | 7,11E-05 | 0,644177251 | 0,351 | 0,176 | 0,847950811 | 8 |
| *fus.4* | 7,17E-05 | 0,431167095 | 0,635 | 0,394 | 0,855743044 | 8 |
| *Pde6.5* | 7,33E-05 | 0,469410133 | 0,743 | 0,506 | 0,87389462 | 8 |
| *step.4* | 7,35E-05 | -1,156837953 | 0,257 | 0,44 | 0,877003925 | 8 |
| *Irp-1B.3* | 7,44E-05 | -1,186811124 | 0,081 | 0,285 | 0,887951677 | 8 |
| *IP3K1.5* | 8,00E-05 | -1,206860087 | 0,378 | 0,563 | 0,954653121 | 8 |
| *Culd.6* | 8,39E-05 | 0,438580239 | 0,581 | 0,338 | 1 | 8 |
| *cnn.5* | 8,59E-05 | -1,225644853 | 0,081 | 0,282 | 1 | 8 |
| *CG15099.5* | 8,60E-05 | -1,300809619 | 0,108 | 0,302 | 1 | 8 |
| *Tl.3* | 8,64E-05 | 0,825284561 | 0,757 | 0,587 | 1 | 8 |
| *CG8665.2* | 9,57E-05 | 0,770490607 | 0,392 | 0,201 | 1 | 8 |
| *CG34054.3* | 9,93E-05 | 0,701888756 | 0,365 | 0,186 | 1 | 8 |
| *Cp110* | 0,000104561 | 0,650352266 | 0,27 | 0,119 | 1 | 8 |
| *Droj2.4* | 0,000111284 | -1,073950712 | 0,203 | 0,42 | 1 | 8 |
| *CG10433.4* | 0,00011624 | 0,46027998 | 0,851 | 0,672 | 1 | 8 |
| *bnl.5* | 0,000118665 | -2,156177637 | 0,135 | 0,333 | 1 | 8 |
| *Gdap2.5* | 0,000120019 | -1,327514858 | 0,23 | 0,411 | 1 | 8 |
| *NimB2* | 0,000125611 | 0,566740004 | 0,527 | 0,315 | 1 | 8 |
| *chic.5* | 0,000128052 | -1,338900493 | 0,23 | 0,422 | 1 | 8 |
| *MCPH1.4* | 0,0001289 | -1,19297694 | 0,122 | 0,32 | 1 | 8 |
| *cpx.5* | 0,000128912 | 0,804706417 | 0,514 | 0,33 | 1 | 8 |
| *Dp1.2* | 0,000130318 | 0,673597448 | 0,419 | 0,24 | 1 | 8 |
| *CG13315.4* | 0,000130388 | 0,689075062 | 0,946 | 0,819 | 1 | 8 |
| *aqz.3* | 0,000135767 | -0,581429536 | 0,905 | 0,949 | 1 | 8 |
| *Prosalpha3.5* | 0,000141486 | -1,258197077 | 0,081 | 0,273 | 1 | 8 |
| *l(3)80Fj.3* | 0,00015113 | 0,436722116 | 0,905 | 0,75 | 1 | 8 |
| *Nop17l.5* | 0,000157998 | -1,124708252 | 0,203 | 0,419 | 1 | 8 |
| *CG16756.2* | 0,000158264 | 0,618393768 | 0,324 | 0,159 | 1 | 8 |
| *norpA.1* | 0,00016732 | 0,640039563 | 0,324 | 0,167 | 1 | 8 |
| *CG5958.6* | 0,000188491 | -1,186031646 | 0,216 | 0,417 | 1 | 8 |
| *Nedd4.5* | 0,000192963 | -0,961723311 | 0,135 | 0,336 | 1 | 8 |
| *GstD1.4* | 0,000195076 | -1,338967606 | 0,541 | 0,656 | 1 | 8 |
| *pcs.5* | 0,000196371 | -1,492280813 | 0,243 | 0,428 | 1 | 8 |
| *Dmtn.5* | 0,000210197 | 0,522798784 | 0,878 | 0,746 | 1 | 8 |
| *Hsc70-5.4* | 0,000219982 | -1,149700057 | 0,095 | 0,286 | 1 | 8 |
| *Rpn10.4* | 0,000228919 | -1,130302378 | 0,068 | 0,253 | 1 | 8 |
| *elB.6* | 0,000232584 | -1,364616906 | 0,122 | 0,316 | 1 | 8 |
| *bun.5* | 0,000234714 | 0,255822289 | 0,932 | 0,841 | 1 | 8 |
| *dnr1.4* | 0,000252872 | -1,563485253 | 0,068 | 0,25 | 1 | 8 |
| *Atg8a.4* | 0,00025435 | -0,792053564 | 0,554 | 0,697 | 1 | 8 |

| | | | | | | |
|---|---|---|---|---|---|---|
| *Jafrac1.3* | 0,000272923 | -1,043757999 | 0,216 | 0,396 | 1 | 8 |
| *REPTOR.5* | 0,000274601 | -0,667168783 | 0,554 | 0,721 | 1 | 8 |
| *RapGAP1.4* | 0,000286629 | 0,358596516 | 0,446 | 0,248 | 1 | 8 |
| *lqf.4* | 0,000303383 | -0,962612475 | 0,432 | 0,562 | 1 | 8 |
| *CLIP-190.2* | 0,000306951 | 0,324667937 | 0,838 | 0,671 | 1 | 8 |
| *su(w[a]).1* | 0,000329425 | 0,702925414 | 0,311 | 0,159 | 1 | 8 |
| *h.4* | 0,000330026 | -1,105423398 | 0,473 | 0,638 | 1 | 8 |
| *Indy.3* | 0,000337019 | -0,769390147 | 0,446 | 0,64 | 1 | 8 |
| *CG18135.4* | 0,000350469 | -0,53337364 | 0,595 | 0,724 | 1 | 8 |
| *CG42524.6* | 0,000360114 | 0,417171575 | 0,689 | 0,466 | 1 | 8 |
| *eEF2.4* | 0,000364632 | -0,660553506 | 0,676 | 0,758 | 1 | 8 |
| *Naprt.4* | 0,000407817 | -1,08492667 | 0,189 | 0,368 | 1 | 8 |
| *CG6910.6* | 0,000415466 | -0,783929109 | 0,635 | 0,762 | 1 | 8 |
| *pan.5* | 0,000420974 | 0,509584508 | 0,77 | 0,57 | 1 | 8 |
| *Gp93.1* | 0,000425878 | 1,087676094 | 0,311 | 0,162 | 1 | 8 |
| *JIL-1.1* | 0,000448809 | 0,588969386 | 0,405 | 0,225 | 1 | 8 |
| *CG6428.6* | 0,000475467 | -1,236930448 | 0,108 | 0,276 | 1 | 8 |
| *Atg18b.4* | 0,000481759 | -1,067183479 | 0,27 | 0,448 | 1 | 8 |
| *path.4* | 0,000501586 | -0,372634088 | 0,757 | 0,891 | 1 | 8 |
| *Mi-2* | 0,000505256 | 0,581555797 | 0,405 | 0,233 | 1 | 8 |
| *Nipped-B.3* | 0,000514301 | -0,677803642 | 0,662 | 0,752 | 1 | 8 |
| *AkhR.6* | 0,000535098 | 0,320795111 | 0,541 | 0,314 | 1 | 8 |
| *tsl.2* | 0,000551531 | 0,535483529 | 0,311 | 0,157 | 1 | 8 |
| *CG2852.2* | 0,000556624 | 0,505103109 | 0,446 | 0,261 | 1 | 8 |
| *mei-P26.4* | 0,000574734 | 0,792152018 | 0,608 | 0,433 | 1 | 8 |
| *SCaMC.5* | 0,00057534 | -0,64357589 | 0,662 | 0,768 | 1 | 8 |
| *SPARC.5* | 0,000664954 | 0,339381125 | 0,5 | 0,302 | 1 | 8 |
| *shrb.4* | 0,000683928 | -1,039005537 | 0,122 | 0,294 | 1 | 8 |
| *PMCA.4* | 0,000709406 | 0,529009931 | 0,73 | 0,579 | 1 | 8 |
| *Dad.4* | 0,000755714 | -0,909619064 | 0,297 | 0,475 | 1 | 8 |
| *MSBP.4* | 0,00076095 | -0,98620493 | 0,122 | 0,29 | 1 | 8 |
| *Tret1-1.5* | 0,000770705 | 0,329525096 | 0,757 | 0,557 | 1 | 8 |
| *Cam.3* | 0,000805105 | -0,596549959 | 0,716 | 0,823 | 1 | 8 |
| *CG8312.5* | 0,000825763 | -1,740462053 | 0,176 | 0,33 | 1 | 8 |
| *pyd.4* | 0,000846042 | 0,378561177 | 0,77 | 0,563 | 1 | 8 |
| *Prat2.2* | 0,000856924 | -0,800232309 | 0,595 | 0,731 | 1 | 8 |
| *vir-1.5* | 0,000870778 | -0,987314719 | 0,649 | 0,741 | 1 | 8 |
| *Thd1.3* | 0,000891427 | 0,40165923 | 0,676 | 0,449 | 1 | 8 |
| *CG31694.5* | 0,000901741 | -0,954752847 | 0,216 | 0,38 | 1 | 8 |
| *Khc-73.5* | 0,000918989 | -0,920380554 | 0,189 | 0,378 | 1 | 8 |
| *Pmp70.6* | 0,000936701 | 0,431529074 | 0,446 | 0,267 | 1 | 8 |
| *CG5080.1* | 0,000982501 | 0,427148632 | 0,311 | 0,163 | 1 | 8 |
| *CG6115.5* | 0,000983594 | -0,664256961 | 0,23 | 0,434 | 1 | 8 |
| *CG46385.4* | 0,000986259 | 0,297073115 | 1 | 0,955 | 1 | 8 |
| *CG32687.4* | 0,001019609 | -0,898953109 | 0,351 | 0,493 | 1 | 8 |
| *alpha-Cat.1* | 0,001027321 | 0,517057355 | 0,405 | 0,237 | 1 | 8 |
| *IscU.4* | 0,001027892 | -0,899869269 | 0,257 | 0,426 | 1 | 8 |
| *drpr.6* | 0,001053034 | -1,008743145 | 0,284 | 0,461 | 1 | 8 |
| *pyr.4* | 0,001053825 | -1,266564758 | 0,135 | 0,301 | 1 | 8 |
| *CG3262* | 0,001055111 | 0,426135083 | 0,257 | 0,125 | 1 | 8 |
| *Exn.4* | 0,001058406 | -0,995591696 | 0,392 | 0,51 | 1 | 8 |
| *CG32276.2* | 0,001109377 | 0,713716886 | 0,446 | 0,287 | 1 | 8 |
| *pAbp.4* | 0,001117827 | -0,699660933 | 0,743 | 0,804 | 1 | 8 |
| *apolpp.4* | 0,001140478 | 0,64362968 | 0,959 | 0,8 | 1 | 8 |
| *Ire1.5* | 0,001142175 | -1,019076326 | 0,162 | 0,321 | 1 | 8 |
| *CG12065.6* | 0,00118911 | -1,034233695 | 0,135 | 0,299 | 1 | 8 |
| *CG7920.6* | 0,00119695 | 0,561738182 | 0,459 | 0,292 | 1 | 8 |
| *spir.5* | 0,001232076 | -0,928827366 | 0,203 | 0,375 | 1 | 8 |
| *zormin.5* | 0,001372364 | -0,915674889 | 0,243 | 0,415 | 1 | 8 |
| *pst.5* | 0,001401199 | 0,406633579 | 0,784 | 0,579 | 1 | 8 |
| *Gbp3.3* | 0,001431832 | 0,42546661 | 0,446 | 0,264 | 1 | 8 |

| | | | | | | |
|---|---|---|---|---|---|---|
| *Tpr2.4* | 0,001457989 | -0,859286713 | 0,514 | 0,599 | 1 | 8 |
| *AttB.3* | 0,001499816 | 1,645076597 | 0,324 | 0,19 | 1 | 8 |
| *klu.5* | 0,001560276 | -0,940720097 | 0,351 | 0,493 | 1 | 8 |
| *alpha-Man-Ia.4* | 0,001633489 | 0,478099809 | 0,541 | 0,356 | 1 | 8 |
| *Pgd* | 0,001636922 | 0,445696776 | 0,297 | 0,156 | 1 | 8 |
| *dlg1.2* | 0,001653965 | 0,53110087 | 0,568 | 0,396 | 1 | 8 |
| *CG3376.3* | 0,001675659 | -0,668659371 | 0,405 | 0,579 | 1 | 8 |
| *Rpn5.4* | 0,001703302 | -0,884019686 | 0,108 | 0,269 | 1 | 8 |
| *c11.1.5* | 0,001723165 | -0,947621746 | 0,095 | 0,25 | 1 | 8 |
| *PNUTS.2* | 0,001751501 | 0,433827571 | 0,757 | 0,606 | 1 | 8 |
| *CG4538.4* | 0,001795039 | -0,880747667 | 0,135 | 0,291 | 1 | 8 |
| *cnc.2* | 0,001829645 | -0,427028159 | 0,959 | 0,963 | 1 | 8 |
| *Npc2g.4* | 0,001862742 | 0,391041696 | 0,514 | 0,342 | 1 | 8 |
| *Got2.5* | 0,001906405 | -0,704634892 | 0,419 | 0,553 | 1 | 8 |
| *Hsc70Cb.4* | 0,001960703 | -0,923304521 | 0,378 | 0,532 | 1 | 8 |
| *l(1)G0196.4* | 0,002029847 | 0,462569605 | 0,595 | 0,429 | 1 | 8 |
| *E(bx).3* | 0,002030549 | 0,475478686 | 0,419 | 0,265 | 1 | 8 |
| *Chd64.6* | 0,002150021 | -0,758396632 | 0,297 | 0,46 | 1 | 8 |
| *Zasp52.2* | 0,002177662 | 0,384978731 | 0,311 | 0,172 | 1 | 8 |
| *Spat.4* | 0,002195729 | -0,951542058 | 0,297 | 0,467 | 1 | 8 |
| *spri.3* | 0,002304913 | 0,37333043 | 0,757 | 0,635 | 1 | 8 |
| *eff.3* | 0,00239614 | -0,87647226 | 0,486 | 0,593 | 1 | 8 |
| *egh.1* | 0,002426585 | 0,412609768 | 0,257 | 0,133 | 1 | 8 |
| *Su(Tpl).1* | 0,002475435 | 0,497130451 | 0,716 | 0,628 | 1 | 8 |
| *LamC.5* | 0,002478647 | -1,071725352 | 0,122 | 0,269 | 1 | 8 |
| *Ptpmeg.2* | 0,00252557 | -0,730743755 | 0,392 | 0,544 | 1 | 8 |
| *CG14073.3* | 0,002594422 | -0,859629005 | 0,257 | 0,401 | 1 | 8 |
| *wdp.3* | 0,002697202 | -1,131307184 | 0,176 | 0,317 | 1 | 8 |
| *Prosbeta7.4* | 0,002741387 | -0,946675224 | 0,108 | 0,253 | 1 | 8 |
| *NAT1.5* | 0,002763531 | -0,586382909 | 0,581 | 0,698 | 1 | 8 |
| *fz2.4* | 0,002808761 | 0,500521359 | 0,432 | 0,278 | 1 | 8 |
| *poe.4* | 0,002845676 | -0,654343868 | 0,176 | 0,344 | 1 | 8 |
| *DOR.5* | 0,002902271 | 0,429802665 | 0,77 | 0,605 | 1 | 8 |
| *lectin-28C.3* | 0,002917994 | -0,85648679 | 0,176 | 0,333 | 1 | 8 |
| *CG32369.5* | 0,003070947 | -0,847383541 | 0,581 | 0,675 | 1 | 8 |
| *Idgf4.3* | 0,003096986 | 0,60484587 | 0,432 | 0,285 | 1 | 8 |
| *nuf.6* | 0,003159109 | -0,680267713 | 0,405 | 0,552 | 1 | 8 |
| *aralar1.4* | 0,003225855 | -0,622573518 | 0,527 | 0,637 | 1 | 8 |
| *sqd.3* | 0,003248667 | 0,28811081 | 0,959 | 0,83 | 1 | 8 |
| *ImpL2.5* | 0,003256753 | -1,373982923 | 0,122 | 0,251 | 1 | 8 |
| *Gp150.3* | 0,003289811 | -0,763918198 | 0,419 | 0,526 | 1 | 8 |
| *CG10799.4* | 0,003339444 | 0,590349802 | 0,649 | 0,499 | 1 | 8 |
| *Non1.3* | 0,003423671 | -0,708345708 | 0,189 | 0,354 | 1 | 8 |
| *Pdfr.4* | 0,003474954 | 0,350837625 | 0,635 | 0,473 | 1 | 8 |
| *spen.2* | 0,00355247 | -0,355389057 | 0,973 | 0,96 | 1 | 8 |
| *Ac13E.4* | 0,003675069 | 0,530648636 | 0,392 | 0,257 | 1 | 8 |
| *tim.4* | 0,003741539 | -0,624788548 | 0,486 | 0,632 | 1 | 8 |
| *Oatp30B.5* | 0,003818046 | -0,678141662 | 0,473 | 0,59 | 1 | 8 |
| *CG43736.2* | 0,003863745 | 0,55773141 | 0,473 | 0,324 | 1 | 8 |
| *Ufd4.5* | 0,00398531 | -0,778629789 | 0,203 | 0,349 | 1 | 8 |
| *fok.4* | 0,00400908 | -0,621565622 | 0,378 | 0,524 | 1 | 8 |
| *Ars2.1* | 0,004308917 | 0,527092237 | 0,27 | 0,154 | 1 | 8 |
| *CkIIalpha* | 0,004408205 | 0,50051838 | 0,284 | 0,162 | 1 | 8 |
| *CG14629.1* | 0,004467692 | 0,436150182 | 0,27 | 0,151 | 1 | 8 |
| *CG31769.1* | 0,004495823 | 0,556780634 | 0,378 | 0,247 | 1 | 8 |
| *Mst84Da.4* | 0,004502183 | -1,062037434 | 0,189 | 0,324 | 1 | 8 |
| *jvl.4* | 0,004516318 | -0,521400228 | 0,649 | 0,752 | 1 | 8 |
| *Mvl.2* | 0,004593711 | 0,756385682 | 0,297 | 0,177 | 1 | 8 |
| *Mical.4* | 0,004600631 | -0,79826891 | 0,162 | 0,31 | 1 | 8 |
| *Smr.6* | 0,004638712 | 0,350238652 | 0,932 | 0,744 | 1 | 8 |
| *CG34120* | 0,004679761 | 0,463592563 | 0,311 | 0,18 | 1 | 8 |

| | | | | | | |
|---|---|---|---|---|---|---|
| *Gdh.5* | 0,004702045 | -0,938293511 | 0,243 | 0,388 | 1 | 8 |
| *Myd88.2* | 0,004811581 | -0,735718362 | 0,216 | 0,367 | 1 | 8 |
| *Gprk1.5* | 0,004874435 | 0,363138729 | 0,514 | 0,343 | 1 | 8 |
| *Gmap.2* | 0,004877106 | 0,423453686 | 0,635 | 0,473 | 1 | 8 |
| *luna.5* | 0,004968347 | 0,388663166 | 0,824 | 0,717 | 1 | 8 |
| *sgg.4* | 0,005070821 | -0,345781669 | 0,959 | 0,948 | 1 | 8 |
| *bmm.5* | 0,005211443 | -0,585727891 | 0,703 | 0,764 | 1 | 8 |
| *Rpn2.4* | 0,005296051 | -0,900110523 | 0,176 | 0,31 | 1 | 8 |
| *Rpn13.5* | 0,005398709 | -0,783999138 | 0,135 | 0,278 | 1 | 8 |
| *ZnT41F.1* | 0,005558126 | 0,659549886 | 0,284 | 0,165 | 1 | 8 |
| *ogre.4* | 0,005564721 | 0,428478509 | 0,405 | 0,259 | 1 | 8 |
| *CG12116.5* | 0,005590714 | 0,452859504 | 0,568 | 0,409 | 1 | 8 |
| *cwo.5* | 0,005672285 | -0,415967693 | 0,865 | 0,879 | 1 | 8 |
| *Parp.5* | 0,005791096 | 0,322817727 | 0,838 | 0,675 | 1 | 8 |
| *CG9005.4* | 0,005862341 | 0,426163402 | 0,527 | 0,366 | 1 | 8 |
| *CG3168.5* | 0,005867362 | -1,049485476 | 0,162 | 0,298 | 1 | 8 |
| *CG31211.1* | 0,00588749 | 0,371537142 | 0,27 | 0,15 | 1 | 8 |
| *Svil.3* | 0,006302386 | -0,919282392 | 0,459 | 0,55 | 1 | 8 |
| *Cyp6w1.2* | 0,006461194 | 0,732294743 | 0,338 | 0,213 | 1 | 8 |
| *kek5.5* | 0,006724343 | 0,38142627 | 0,757 | 0,598 | 1 | 8 |
| *CG34136.4* | 0,006840932 | -0,646849925 | 0,135 | 0,276 | 1 | 8 |
| *Pdk1.4* | 0,006850232 | -0,604731454 | 0,676 | 0,733 | 1 | 8 |
| *cathD.2* | 0,006867714 | 0,419144995 | 0,419 | 0,276 | 1 | 8 |
| *Gclm.6* | 0,006910839 | -1,01028877 | 0,23 | 0,365 | 1 | 8 |
| *CG43367* | 0,00694566 | 0,330731001 | 0,257 | 0,139 | 1 | 8 |
| *Plod.5* | 0,007102514 | -0,665488207 | 0,149 | 0,286 | 1 | 8 |
| *Pisd.2* | 0,007199699 | 0,413763672 | 0,257 | 0,145 | 1 | 8 |
| *CG10621.5* | 0,007658473 | -1,280822341 | 0,149 | 0,271 | 1 | 8 |
| *PRL-1.5* | 0,00781238 | -0,872617925 | 0,176 | 0,302 | 1 | 8 |
| *Cyp6g1.4* | 0,008026213 | -0,847128084 | 0,284 | 0,406 | 1 | 8 |
| *comm2.4* | 0,008132259 | -0,842902084 | 0,135 | 0,266 | 1 | 8 |
| *alt.4* | 0,008138025 | 0,520862183 | 0,527 | 0,354 | 1 | 8 |
| *CG41099.1* | 0,00822778 | 0,531278044 | 0,311 | 0,189 | 1 | 8 |
| *Pli.5* | 0,008417853 | 0,342095269 | 0,676 | 0,562 | 1 | 8 |
| *CG11400.5* | 0,008451698 | 0,352666959 | 0,541 | 0,382 | 1 | 8 |
| *smash.2* | 0,008469817 | -0,629788211 | 0,554 | 0,641 | 1 | 8 |
| *kibra.4* | 0,008788902 | -0,893380581 | 0,338 | 0,448 | 1 | 8 |
| *Trp1.3* | 0,009174871 | 0,606737995 | 0,351 | 0,232 | 1 | 8 |
| *Tis11.2* | 0,009341502 | -0,412441156 | 0,865 | 0,913 | 1 | 8 |
| *RasGAP1.4* | 0,009447114 | -0,804936922 | 0,149 | 0,281 | 1 | 8 |
| *Cat.3* | 0,009478762 | -0,607983499 | 0,216 | 0,352 | 1 | 8 |
| *bin3* | 0,009501555 | 0,57334967 | 0,459 | 0,337 | 1 | 8 |
| *GEFmeso.5* | 0,009508655 | -0,606773667 | 0,297 | 0,442 | 1 | 8 |
| *rgn.5* | 0,009535514 | -1,241347951 | 0,351 | 0,463 | 1 | 8 |
| *Fer1HCH.5* | 0,009730565 | -0,534667686 | 0,743 | 0,804 | 1 | 8 |

ntity
