## Supplementary material for "DNA damage signaling in *Drosophila* macrophages modulates systemic cytokine levels in response to oxidative stress": Table S3

| List of *Drosophila* lines used in this study | Source | ID |
|---|---|---|
| *$w^{1118}$;HmlΔ -Gal4,UAS-2xeGFP* | Gift from Marc S. Dionne | N/A |
| *$w^{1118}$;UAS-tefu-IR* | Vienna Drosophila Research Center | VDRC ID: 108074 |
| *$w^{1118}$;UAS-mei41-IR* | Vienna Drosophila Research Center | VDRC ID: 11251 |
| *$w^{1118}$;UAS-nbs-IR* | Vienna Drosophila Research Center | VDRC ID:110366 |
| *$w^{1118}$;UAS-mei41-IR;UAS-tefu-IR* | made for this study | N/A |
| *$w^{1118}$;UAS-bsk-DN* | Vienna Drosophila Research Center | VDRC ID: 104569 |
| *$w^{1118}$;;HmlΔ-DsRed.nuc* | Gift from Katie J. Woodcock | N/A |
| *$w^{1118}$;UAS-upd3-IR* | Vienna Drosophila Research Center | VDRC ID: 106869 |
| *$w^{1118}$;HmlΔ -Gal4,UAS-2xeGFP;tubGal80/tm6csb´* | Gift from Katie J. Woodcock | N/A |
| *$w^{1118}$;UAS-hep[act]/CyO; Dr/tm6csb'* | Gift from Anne-Kathrin Classen | N/A |
| *$w^{1118}$;UAS-upd3/Sm6a* | Gift from Katie J. Woodcock | N/A |
| *w[*] upd3[Δ]* | Bloomington Drosophila Stock Center | BDSC ID: 55728 |
| *w´upd3Δ; UAS-upd3/Sm6a* | made for this study | N/A |
| *$w^{1118}$;tub-Gal80$^{ts}$/Sm6a;crq-Gal4/TM6 c, Sb'* | Gift from Marc S. Dionne | N/A |
| *w1118;;UAS-rpr/TM6 c, Sb1* | Bloomington Drosophila Stock Center | BDSC ID: 5824 |
| *w1118;UAS-CD8-mCherry* | Bloomington Drosophila Stock Center | BDSC ID: 27391 |

| List of RTqPCR primers used in this study | Source |
|---|---|
| Primer: Rpl1_for 5'-TCCACCTTGAAGAAGGGCTA-3' | Designed for this study |
| Primer: Rpl1_rev 5'-TTGCGGATCTCCTCAGACTT-3' | Designed for this study |
| Primer: Upd1_for 5'-GCACACTGATTTCGATACGG-3' | Designed for this study |
| Primer: Upd1_rev 5'-CTGCCGTGGTGCTGTTTT-3' | Designed for this study |
| Primer: Upd2_for 5'-CGGAACATCACGATGAGCGAAT-3' | Designed for this study |
| Primer: Upd2_rev 5'-TCGGCAGGAACTTGTACTCG-3' | Designed for this study |
| Primer: Upd3_for 5'-ACTGGGAGAACACCTGCAAT-3' | Designed for this study |
| Primer: Upd3_rev 5'-GCCCGTTTGGTTCTGTAGAT-3' | Designed for this study |
| Primer: Socs36E_for 5'-AAAAAGCCAGCAAACCAAAA-3' | Designed for this study |
| Primer: Socs36E_rev 5'-AGGTGATGACCCATTGGAAG-3' | Designed for this study |
| Primer: TotA_for 5'-CCAAAATGAATTCTTCAACTGC-3' | Designed for this study |
| Primer: TotA_rev 5'-GAATAGCCCATGCATAGAGGAC-3' | Designed for this study |
| Primer: Puc_for 5'-CGTCATCATCAACGGCAAT-3' | Designed for this study |
| Primer: Puc_rev 5'-AGGCGGGGTGTGTTTCTAT-3' | Designed for this study |
| Primer: Ilp-2_for 5'-ATCCCGTGATTCCACCACAAG-3' | Designed for this study |
| Primer: Ilp-2_rev 5'-GCGGTTCCGATATCGAGTTA-3' | Designed for this study |
| Primer: Ilp-3_for 5'- CAACGCAATGACCAAGAGAA-3' | Designed for this study |
| Primer: Ilp-3_rev 5'-TGAGCATCTGAACCGAACT-3' | Designed for this study |
| Primer: Ilp-5_for 5'-GCCTTGATGGACATGCTGA-3' | Designed for this study |
| Primer: Ilp-5_rev 5'-AGCTATCCAAATCCGCCA-3' | Designed for this study |
| Primer: Thor_for 5'- CAGGAAGGTTGTCATCTCGGA-3' | Designed for this study |
| Primer: Thor_rev 5'-GGAGTGGTGGAGTAGAGGGTT-3' | Designed for this study |
| Primer: Pepck_for 5'-GGATAAGGTGGACGTGAAG-3' | Designed for this study |
| Primer: Pepck_rev 5'-ACCTCCTGCGACCAGAACT-3' | Designed for this study |
| Primer: InR_for 5'-GCACCATTATAACCGGAACC-3' | Designed for this study |

| | |
|---|---|
| Primer: InR_rev 5’-TTAATTCATCCATGACGTGAGC-3’ | Designed for this study |
| Primer: Def_for 5’-TTCTCGTGGCTATCGCTTTT-3’ | Designed for this study |
| Primer: Def_rev 5’-GGAGAGTAGGTCGCATGTGG-3’ | Designed for this study |
| Primer: AttA_for 5’-CACAATGTGGTGGGTCAGG | Designed for this study |
| Primer: AttA_rev 5’-GGCACCATGACCACCATT | Designed for this study |
| Primer: DiptA_for 5’-ACCGCAGTACCCACTCAATC | Designed for this study |
| Primer: DiptA_rev 5’-CCCAAGTGCTGTCCATATCC | Designed for this study |
| Primer: Mtk_for 5’-TCTTGGAGCGATTTTTCTGG | Designed for this study |
| Primer: Mtk_rev 5’-TCTGCCAGCACTGATGTAGC | Designed for this study |
| Primer: Dro_for 5’-CCATCGAGGATCACCTGACT | Designed for this study |
| Primer: Dro_rev 5’-CTTTAGGCGGGCAGAATG | Designed for this study |
| Primer: Drs_for 5’-GTACTTGTTCGCCCTCTTCG | Designed for this study |
| Primer: Drs_rev 5’-CTTGCACACACGACGACAG | Designed for this study |
| Primer: daw_for 5’-CAGTCGGTGAATGTGCAAGA | Designed for this study |
| Primer: daw_rev 5’-GAGGTGGCTCAGCTCGTG | Designed for this study |
| Primer: dpp_for 5’-GCTGATTGGATACATAATTCTCAGG | Designed for this study |
| Primer: dpp_rev 5’-TTTGAAAAGTCGCCAGCAC | Designed for this study |
| Primer: eiger_for 5’-CGACGAGTTCCAAAAGGAGT | Designed for this study |
| Primer: eiger_rev 5’-GTCGTCGTCCTCCTCATC | Designed for this study |
